# The methyltransferase NmbA methylates the low-molecular weight thiol bacillithiol, and displays a specific structural architecture

**DOI:** 10.1101/2025.07.06.663066

**Authors:** Marta Hammerstad, Erlend Steinvik, Hans-Petter Hersleth

## Abstract

Low-molecular weight (LMW) thiols maintain the cellular redox balance, and play a protective role against reactive species, heavy metals, toxins, and antibiotics. Despite serving similar metabolic functions, structurally distinct LMW thiols are widespread in nature, with bacillithiol (BSH) being the predominant LMW thiol in bacteria. The LMW thiol *N*-methyl-BSH (*N*-Me-BSH) has been characterized in the green sulfur bacterium *Chlorobaculum tepidum*, unveiling the presence and role of a putative *S*-adenosyl-L-methionine (SAM)-dependent methyltransferase (MT), NmbA, catalyzing the final biosynthetic step of *N*-Me-BSH. In this work, we report biochemical evidence for NmbAs function as an MT of the N-atom of the BSH cysteine moiety, showing substrate specificity, which together with our bioinformatics analyses classifies NmbA and its outspread homologs as *N*-directed natural product MTs (NPMTs) of the LMW thiol BSH. We also present the crystal structure of NmbA, which shows that NmbA is a Class I SAM-dependent MT, however, displaying a unique three-dimensional architecture unlike that observed for other NPMTs. The NmbA active site adopts a narrow molecular basket structure due to an unusual organization of the variable Cap domain, which together with our docking calculations suggests that it may specifically accommodate the BSH substrate. Our studies provide a valuable overview of the phylogenetic distribution of *N*-Me-BSH in bacteria, accompanied by important functional and structural insight into a new class of NPMTs. Our findings contribute to the field of SAM-dependent MTs, as well as possible applications for targeting distinct bacterial defense mechanisms involving LMW thiols with potential environmental, biotechnological, and medical implications.

## INTRODUCTION

Low-molecular-weight (LMW) thiols, a group of reactive sulfhydryl-containing metabolites, play a critical role in mediating redox homeostasis in nearly all organisms. These antioxidants maintain a reducing intracellular environment, function as thiol cofactors of many enzymes in scavenging of reactive species, in detoxification of toxins and antibiotics, in protection against heavy metals and metal storage, and regulate and protect exposed protein thiols through *S*-thiolation ^1-4^. Despite playing analogous roles as redox buffers in cells, LMW thiols display large structural diversity across species. Eukaryotes and most Gram-negative bacteria use the extensively studied glutathione (GSH) as their major LMW thiol ^5-6^. Studies have demonstrated that most bacteria (and some eukaryotes) do not produce GSH, and rely on alternative and unique thiol-redox buffers ^7-8^. These include the structurally distinct LMW thiols trypanothione, γ-glutamylcysteine, ergothioneine ^1^, ovothiol ^1^, and glutathione amide ^9^. Furthermore, Gram-positive bacteria mainly produce two major structurally distinct LMW thiols, involved in similar metabolic processes. In Actinobacteria (high-G+C Gram-positive bacteria), such as mycobacteria, *Streptomycetes* and corynebacteria, mycothiol (MSH) serves as the predominant LMW thiol ^10-13^. Firmicutes (Bacillota) (low-G+C Gram-positive bacteria), including *Staphylococcus aureus* and many bacilli, utilize bacillithiol (BSH, Cys-GlcN-Mal) ^14-16^. To limit the extent of oxidative damage (e.g caused by reactive oxygen species (ROS) generated by human neutrophils and macrophages during infection, as well as from external sources such as antibiotics), pathogenic Firmicutes rely on mechanisms involving BSH. Another important role of BSH is the protection of redox-sensitive protein thiols, through the reversible formation of mixed disulfides with BSH, termed *S*-bacillithiolation. This protein *S*-thiolation under oxidative stress conditions is analogous to *S*-glutathionylation in eukaryotes, and the de-thiolation is catalyzed by glutaredoxins or bacilliredoxins in cells relying on GSH and BSH, respectively. In the latter case, this ultimately leads to the formation of the bacillithiol disulfide (BSSB) ^17-19^. Alternatively, BSH can react directly with ROS, again leading to the oxidation of BSH to BSSB ^20^. In order to maintain the high intracellular BSH:BSSH redox ratio, BSH is recycled by the FAD-containing NADPH-dependent oxidoreductase Bdr, present only in BSH-containing bacteria ^21-23^.

The vast and increasing number of reported LMW thiols, particularly across bacteria, highlights the importance to further elucidate the role of, and discover new functionally similar, yet structurally distinct redox modulators. Through evolution, LMW thiols have evolved to participate in crucial and similar cellular processes, yet diverged into specialized molecules with unique structural and biophysical properties across organisms. Hence, a more detailed understanding of the enzymatic and cellular processes involving these key metabolites may contribute to future antimicrobial drug development. Alternatively, a broader insight into the microbiology of bacteria evolved to utilize specialized forms of LMW thiols, often found in extreme environments, could unveil and establish new biotechnological applications. In 2018, Hiras *et al*. reported the existence of yet another novel thiol from the anaerobic green sulfur bacterium *Chlorobium tepidum, N*-methyl-bacillithiol (*N*-Me-BSH), a BSH derivative modified by *N*-methylation of the amino group of the cysteine moiety of BSH ^24^ (Scheme 1).

**Scheme 1.**
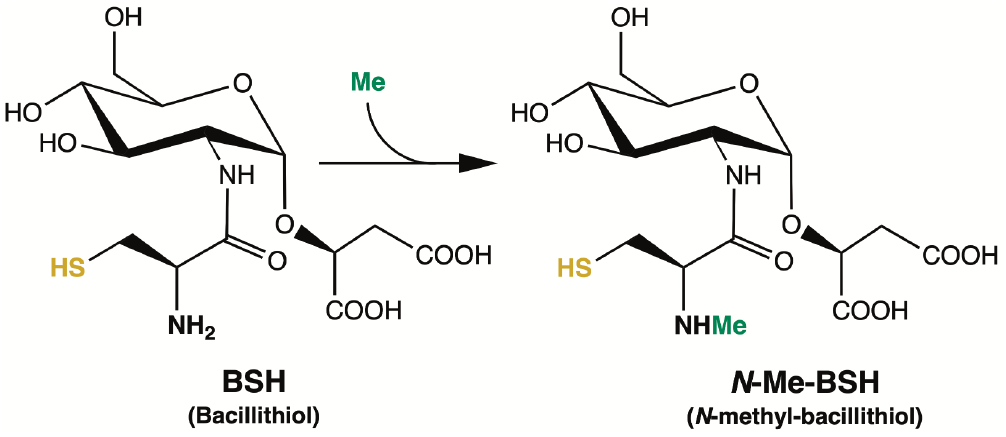
Structures of BSH and *N*-Me-BSH, with the redox-active sulfhydryl group depicted in yellow and the methyl group of *N*-Me-BSH highlighted in green.

The *N*-Me-BSH pool size was found to be highest in the stationary growth phase, and strongly correlated to the cellular biomass. *C. tepidum* cultures grown in low light contained a fivefold more *N*-Me-BSH. Moreover, *N*-Me-BSH was found to exist predominantly in its reduced state, which together suggests that *N*-Me-BSH is an important redox buffer in *C. tepidum*.

In addition to the BSH biosynthesis genes bshA, bshB, and bshC ^15, 25^, *C. tepidum* was found to contain two orthologs of putative *S*-adenosyl-l-methionine (SAM)-dependent methyltransferases (MTs), CT1213 and CT1040, where the latter was identified as the *in vivo* BSH MT, designated NmbA (*<u>N</u>*-<u>M</u>e-<u>B</u>SH synthase <u>A</u>) ^24^. The mutant knock-out strains showed no BSH methylation and instead BSH levels similar to those of *N*-Me-BSH in the wild-type, and strains lacking CT1040 grew 20 % slower than the parental strain in media with sulfide and thiosulfate as electron donors. Although bacteria producing *N*-Me-BSH often contain BSH as well, the methylated form appears to be the physiologically preferred form due to its higher levels. However, detailed information about the biosynthesis of *N*-Me-BSH, as well as the functional role of *N*-Me-BSH in Chlorobiota and other bacteria encoding an NmbA homolog is currently unknown, emphasizing the need for further investigations.

MTs catalyze the transfer of a methyl group from SAM to a given substrate, leaving the product *S*-adenosyl-L-homocysteine (SAH). In the biosynthesis of natural products, methylation of primary and secondary metabolites is a common chemical modification, catalyzed by small molecule, or natural product, SAM-dependent MTs (NPMTs) ^26^. MTs are classified based on the methyl-accepting atom, with the most abundant ones being *O*-directed MTs, but NPMTs also commonly methylate *N, C*, or less frequently *S*, in addition to other less typical types of acceptors ^27-29^. As a proposed MT of BSH ^24^, NmbA can be typified as an *N*-directed NPMT. *N*-methylation of cysteine, however, is rare, and *N*-Me-BSH is regarded as the first reported case of cysteine *N*-methylation catalyzed by a putative standalone methyl transferase ^30-32^, which highlights the importance to further characterize NmbA. In this work, we aimed to conduct a broader investigation of the NmbA enzyme in terms of phylogenetic distribution and provide biochemical evidence for the role of NmbA as an MT of BSH. Moreover, we report the first crystal structure of NmbA, revealing a unique structural architecture providing insight into its mode of action.

## RESULTS AND DISCUSSION

### The phylogenetic distribution of NmbA displays a new class of MTs

To generally characterize NmbA with respect to other homologous MTs, and to identify its distribution among organisms containing BSH, we performed a bioinformatics analysis. We searched through the RefSeq (NCBI Reference Sequence) Database for organisms encoding the complete *N*-Me-BSH biosynthetic pathway with orthologs of NmbA, BshA, BshB, and BshC, using *C. tepidum* as a reference. No eukaryotes or archaea were identified to contain the complete *N*-Me-BSH nor BSH biosynthetic pathway. For the microbials, all phyla that have been validly published according to the Bacteriological Code ^33-34^ were searched. A total of 3842 unique species were identified to contain the complete BSH biosynthetic pathway and among these, 1765 contained NmbA orthologs (Figure 1, Table S1, and Supplementary Dataset 1). All Chlorobiota species containing the BSH biosynthetic enzymes also encode NmbA. The phyla containing the largest number of species encoding the full BSH biosynthetic pathway are Bacillota (1466) and Bacteroidota (2111), and among these, 25% and 63% also contain NmbA, respectively.

**Figure 1.**
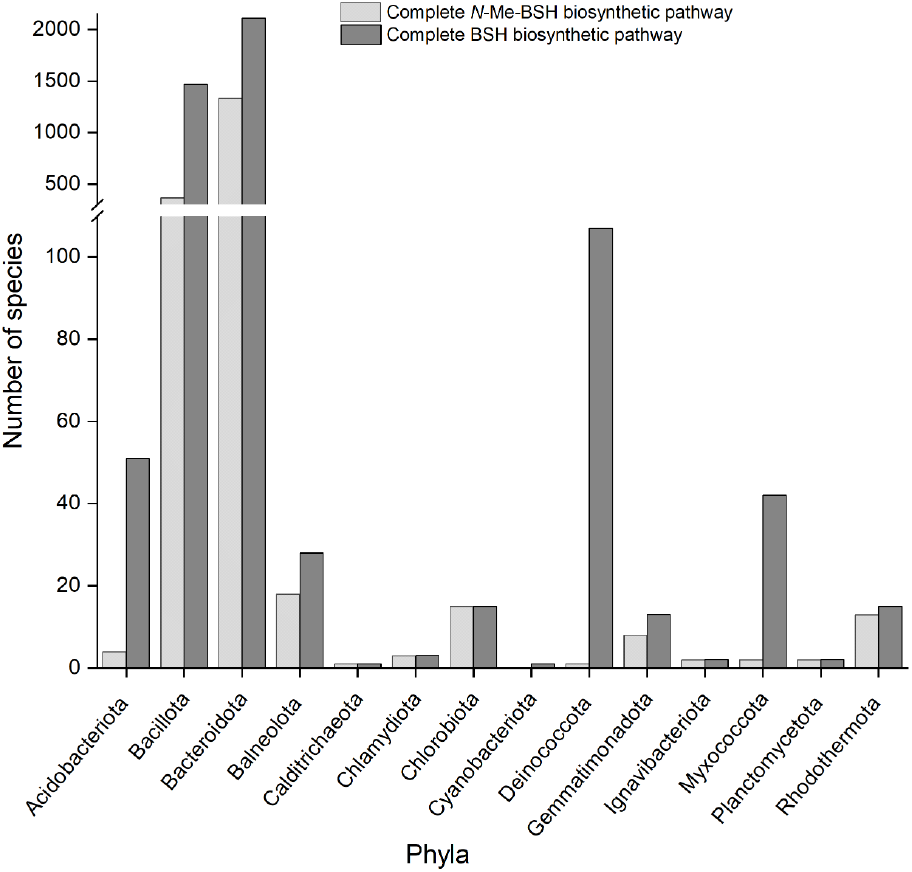
Phylogenetic distribution of bacteria encoding *N*-Me-BSH or BSH. The bar graph shows the amount of unique bacterial species encoding the full *N*-Me-BSH biosynthetic pathway (BshA, BshB, BshC, and NmbA) (light gray) and the BSH biosynthetic pathway (BshA, BshB, and BshC) (dark gray).

The presence of a complete pathway for *N*-Me-BSH biosynthesis in bacterial members of these phyla is in agreement with what was previously reported ^24^, however, our findings show that *N*-Me-BSH is likely utilized by a considerably larger number of species within all three phyla. Other phyla with a high propensity for having NmbA (in addition to BshA, BshB, and BshC) include Balneolota, Gemmatimonadota, and Rhodothermota, while single or sparse NmbA-encoding members are identified among several phyla (Figure 1 and Table S1). Notably, most members of the Deinococcota, Myxococcota, and Acidobacteriota containing the BSH pathway do not encode NmbA. Among the Bacteriodota, the genes for *N*-Me-BSH biosynthesis are most heavily represented in the Families Flavobacteriaceae, Cytophagaceae, Sphingobacteriaceae, Chitinophagaceae, and Weeksellaceae, whereas Families within the Paenibacillaceae and Thermoactinomycetaceae comprise the majority of hits in the Bacillota cluster. Common nominators for the seeming *N*-Me-BSH-synthesizing bacteria are that many of these live in aquatic and marine, as well as extreme environments, including several thermophiles and halophiles. Moreover, some pathogenic bacteria are found to encode the complete *N*-Me-BSH biosynthetic pathway, of which many are aquatic pathogens. These include e.g. *Flavobacterium columnare*, the causative agent of columnaris disease in freshwater fish ^35^, displaying a serious threat to fish farming and fish populations worldwide. Among bacteria containing only BshA, BshB, and BshC, a broader ecological trait is observed, e.g., Bacillota, where families Staphylococcaceae and Bacillaceae include several human pathogens. In summary, we report the presence of BshA, BshB, BshC, and NmbA orthologs in multiple members among several bacterial phyla of environmental and medical relevance (Supplementary Dataset 1), predicting that they should synthesize *N*-Me-BSH.

To further examine the distribution and properties of NmbA, we generated a sequence-similarity network (SSN) with the EFI - Enzyme Similarity Tool using 35-95% sequence identity criteria of homologs in the UniProt90 database. The SSN displays a clustering of NmbA homologs, and the largest clusters were selected and assembled into nine phyla groups (Figure S1), with Bacteriodota and Bacillota comprising the largest phyla, resulting in the same phyla as from the bioinformatics analysis above, when searching for the full *N*-Me-BSH biosynthetic pathway. None of the NmbA homologs from these top clusters have other annotated functions, making them likely candidates as BSH MTs. The SSN also identified some additional sequences appearing as single entities with the alignment score chosen, not clustering with the major groups. Thus, these MTs might methylate other acceptor molecules. These findings suggest that NmbA and its homologs constitute a new and distinct class of NPMTs.

### NmbA is a specific methylase of BSH

#### MT activity

Whereas NmbA presumably employs SAM for activity, biochemical evidence confirming that NmbA acts as a specific BSH methylase *in vitro* has been missing. We were curious to elucidate this, not only for NmbA from the green sulfur bacterium *C. tepidum* (Chlorobiota), but also from a representative of a putative *N*-Me-BSH-utilizing Bacilli, *P. polymyxa* NmbA (*Pp*NmbA) (Bacillota) (Figure S1). Conversion of BSH to *N*-Me-BSH was investigated through enzymatic assays and relative product formation was analyzed and quantified by mass spectrometry (MS). In the presence of BSH and SAM, our results show that *Ct*NmbA and *Pp*NmbA catalyze the methylation of BSH. Omitting either SAM, BSH or enzyme resulted in no product formation. Analysis of reaction products run under identical, as well as varying conditions showed that *Ct*NmbA and *Pp*NmbA catalyze the methylation of BSH to approximately equal product yields (Figure 2), confirming their role as BSH MTs, and supporting the use of *N*-Me-BSH among different bacterial phyla.

**Figure 2.**
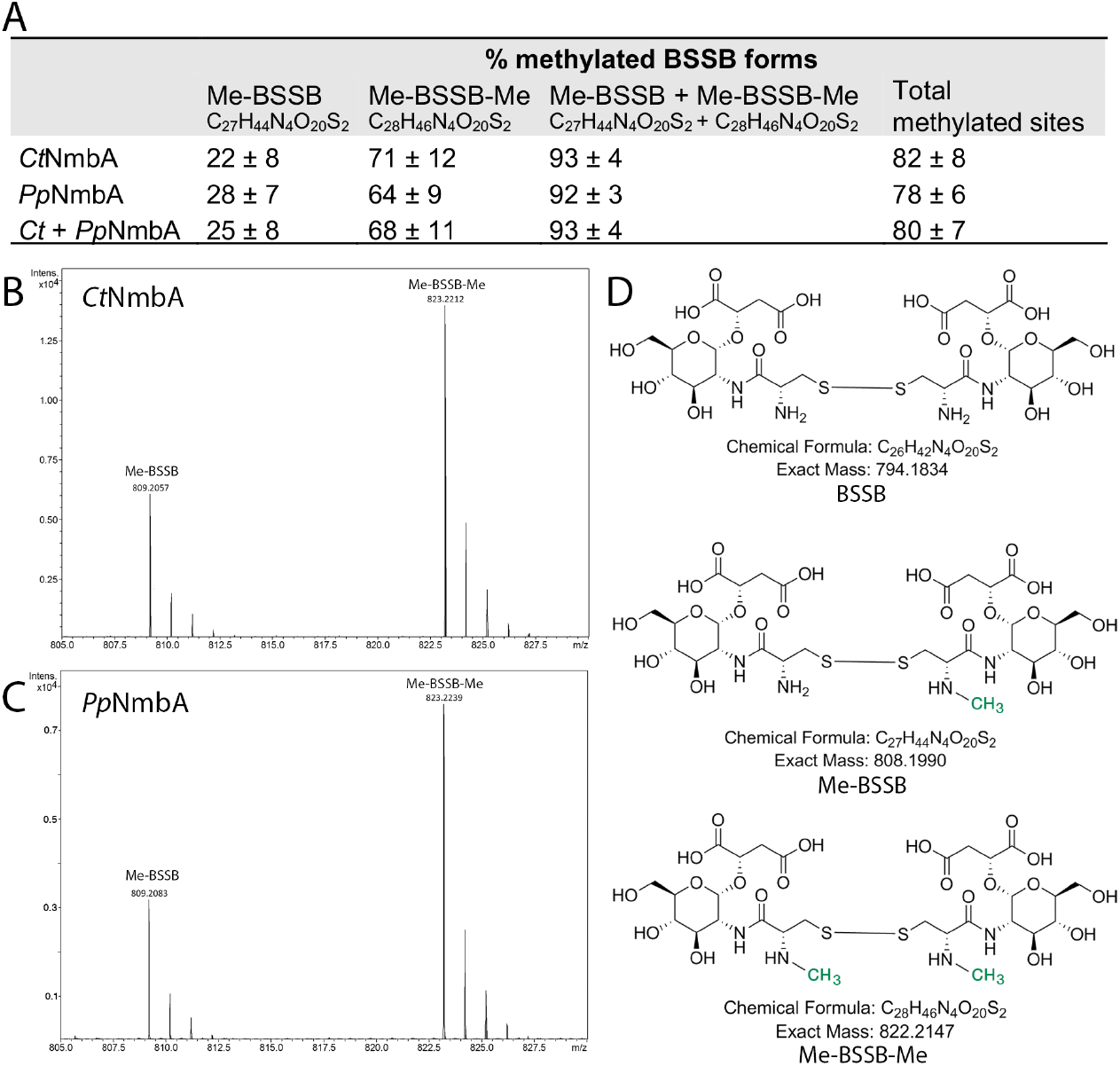
Mass spectrometry analysis of methylated BSSB products. (**A**) The calculated percentage of methylated BSSB forms from the enzymatic reactions using BSH as a substrate and SAM as co-substrate, catalyzed by *Ct*NmbA or *Pp*NmbA. The numbers are calculated from the total BSSB pool (unmethylated BSSB, Me-BSSB, and Me-BSSB-Me) from the enzymatic reactions. The total amount of methylated sites is given based on the total amount of possible methylation sites (2/BSSB dimer). (**B**) and (**C**) Representative mass spectra of reactions catalyzed by *Ct*NmbA and *Pp*NmbA, respectively, showing the masses of Me-BSSB [M + H]^+^809.2057/809.2083 and Me-BSSB-Me [M + H]^+^823.2212/823.2239 (*Ct*NmbA/*Pp*NmbA). Structures of the three BSSB forms; unmethylated, mono-methylated, and di-methylated, are shown in (**D**), with the methyl groups (-CH_3_) highlighted in green.

#### Methylation products

As the aqueous solution of the synthetic BSH used in the reactions is readily oxidized in the MS analysis, all resulting methylated product species include the methylated disulfide form of BSSB, except for trace amounts of methylated BSH. Relative product formation was quantified by MS, with the ion count for all products (and by-products) set to 100%, adjusted according to the BSH substrate count, and with all corresponding values scaled accordingly. *Ct*NmbA and *Pp*NmbA methylate the N-atom of the BSH cysteine with similar ability. Generally, all reaction conditions result in the detection of two physiologically significant product species, the mono– and di-methylated forms of BSSB; Me-BSSB (C_27_H_44_N_4_O_20_S_2_, m/z 808.1990) and Me-BSSB-Me (C_28_H_46_N_4_O_20_S_2_, m/z 822.2147) (Figure 2). For all reactions catalyzed by both *Ct*NmbA and *Pp*NmbA, an excess of the di-methylated species is detected. The percentage of Me-BSSB constitutes 25 ± 8 % of the total pool of BSSB species, whereas Me-BSSB-Me comprises 68 ± 11 % of the total BSSB content. Ultimately, this results in in a total of 93 ± 4 % product formation, equivalent to 80 ± 7 % available BSSB-sites for methylation (100 % for Me-BSSB-Me and 50 % for Me-BSSB), leaving <10 % unmethylated BSSB substrate (C_26_H_42_N_4_O_20_S_2_, m/z 794.1834) (Figure 2A).

#### By-products

Due to the hydrolytic instability of BSH, as well as possible side reactions, we observed the presence of three by-products in all *in vitro* reactions (Figure S2). C_17_H_30_N_2_O_10_S (m/z 454.1621), a likely butylated form of BSH also present in the control samples only containing the synthetic substrate, is a possible by-product or side product from the BSH synthesis, found in low amounts (10.2 ± 4.1 %). The second species, C_14_H_22_N_2_O_10_S (m/z 410.0995) (<1 %) (Figure S2B), predicted to be a BSH species with a methyl-bridge formed between the cysteine amine and sulfide group of BSH from a potential radical reaction between BSH and SAM in a non-enzymatic reaction is detected in trace amounts in all reactions, including control reactions where enzyme was omitted. Interestingly, the last species, C_15_H_24_N_2_O_10_S (m/z 424.1152) (Figure S2B), a likely methylated form of C_14_H_22_N_2_O_10_S, also on the cysteine N-atom, is present in noticeable amounts (11.7 ± 9.2 %) only in the enzymatically catalyzed reactions containing NmbA, indicating that NmbA is able to methylate a secondary amine in addition to the physiologically relevant primary N-atom of the BSH cysteine. For NmbA-catalyzed reactions, this species constitutes a comparable average product yield as compared to the physiologically relevant mono-methylated BSSB species, however, together with the two latter by-products, was not included in the final calculations of physiologically significant product yields presented in Figure 2A (Me-BSSB and Me-BSSB-Me).

#### Substrate oxidation state

To explore substrate permissiveness further, we performed the enzymatic assays on oxidized BSSB as a substrate for methylation. Under no conditions was methylation of BSSB observed, leaving a 100 % content of unmethylated BSSB (C_26_H_42_N_4_O_20_S_2_) (or > 90 % including the by-products), suggesting that NmbA is highly selective for BSH over BSSB. Thus, BSH is methylated to Me-BSSB before being oxidized to Me-BSSB-Me, and due to the redox instability of BSH and Me-BSH *in vitro*, Me-BSH can further form a disulfide bridge with the unmethylated BSH, ultimately leading to the mono-methylated Me-BSSB form. As only the reduced form of BSH is methylated by NmbA, these results suggest that *N*-Me-BSH is the redox-responsive LMW thiol in bacteria encoding the *N*-Me-BSH biosynthesis genes, and that the methylated form of BSH is utilized in LMW thiol-dependent biochemical processes prior to its oxidation to the disulfide form, and before being recycled back to the reduced state. To our knowledge, there are no reports on a specific *N*-Me-BSH de-methylase in the literature. It would be of interest to elucidate whether the methylation step of BSH occurs as a final step of the biosynthesis of *N*-Me-BSH to a given proportion of the BSH pool, acting as a permanent and stable modification throughout the lifespan of *N*-Me-BSH as a redox buffer, or if a de-methylation (and successive re-methylation) event occurs under certain conditions or with respect to specific cellular processes.

#### Substrate specificity

Moreover, we wanted to gain more insight into substrate specificity and assess whether NmbA can act upon diverse substrates *in vitro*. Based on the high structural similarity to glycine N-MTs, the small size of the substrate, and to investigate if NmbA is limited to the BSH cysteine N-atom in substrate scope, the amino acid glycine was tested as a substrate. As an additional control with respect to an alternative LMW substrate, enzymatic activity of NmbA on GSH was also tested. No reaction products of the two latter substrates were detected by MS. In summary, *Ct*NmbA and *Pp*NmbA have the ability to methylate 93 and 92 %, respectively, of the total available substrate content, resulting in either Me-BSSB or Me-BSSB-Me, which *in vitro* accounts for an average 80 % of the total methyl-accepting sites in BSH. Our findings suggest that NmbA acts as a selective *N*-directed MTase of the cysteine moiety of BSH, catalyzing the last step of *N*-Me-BSH biosynthesis (Figure S3).

### NmbA – a seven β-strand NPMT with a unique architecture

#### Overall fold of *Ct*NmbA

As a representative of a new proposed class of NPMTs, we were interested in getting insight into the structural aspects of NmbA as a specific methylase of a novel LMW thiol. Despite the broad chemical and structural diversity of NPMT substrates across all domains of life, all currently known NPMT structures can be classified as members of either Class I or Class III MT-superfamily fold enzymes ^36-37^. The majority belong to the large Class I MTs, dominated by a Rossmann-like fold, which was also seen for the first reported crystal structure of an NPMT, catechol O-MT (COMT) ^38^. The fold is made up by alternating α-helices and β-strands, with a core seven β-strand (7BS) sheet sandwiched by the α-helices, serving as a SAM-binding domain ^36, 39^. A series of regions across the core Rossmann-like fold, designated Motifs I-VI, contain conserved sequence blocks, where several nearly invariant residues along Motifs I-III are directly associated with SAM co-substrate binding. In addition, most MTs contain supplementary domains and structural elements, contributing to the variations in the substrate-binding regions, and hence the selectivity of individual MTs ^36^. Here, we present the first crystal structure of *Ct*NmbA (Table 1), contributing to a deeper understanding of the methylation of BSH, as well as the NPMTs selectivity towards a diverse range of methyl acceptors. Overall, the structure of NmbA is assembled into two domains, dominated by the canonical Class I MT Rossmann-like core domain (Figure 3). This domain consists of a twisted 7BS sheet with five alternating α-helices, where the sheet is sandwiched between two helical regions of two and three α-helices, respectively. Arranged in a sequential order across the core domain, six structural elements equivalent to the highly conserved Motifs I-VI are identified in the structure, confirming that NmbA is a SAM-binding enzyme. Motif I, which is found in the majority of MTs, spanning the first β-strand (β1) of the Rossmann-like fold and the preceding loop connecting it to the following α-helix, typically includes a conserved amino acid block with the consensus sequence (V/I/L)(L/V)(D/E)(V/I)-GxGxG. In NmbA, this sequence corresponds to VLDI and the C-terminal glycine-rich signature sequence AxGxG, where the G to A substitution can be seen in some MTs ^37, 40^. Furthermore, NmbA harbors the typical and partially conserved acidic residue D located at the C-terminal part of β2, and in the loop succeeding β3, encompassing Motifs II and III, respectively. Motif IV contains a partially conserved D at the N-terminus of β4, which is on the side of the strand opposite to the SAM-binding site. Aromatic residues are found in the loop adjoining Motif IV and the following α-helix (α4) (Motif V), where in particular Y134 and F135 are likely to be involved in the stabilization of the adenine moiety of SAM. Motif VI contains a conserved G residue located on the following tight turn preceding β5 of the Rossmann-like fold of NmbA.

**Table 1.**
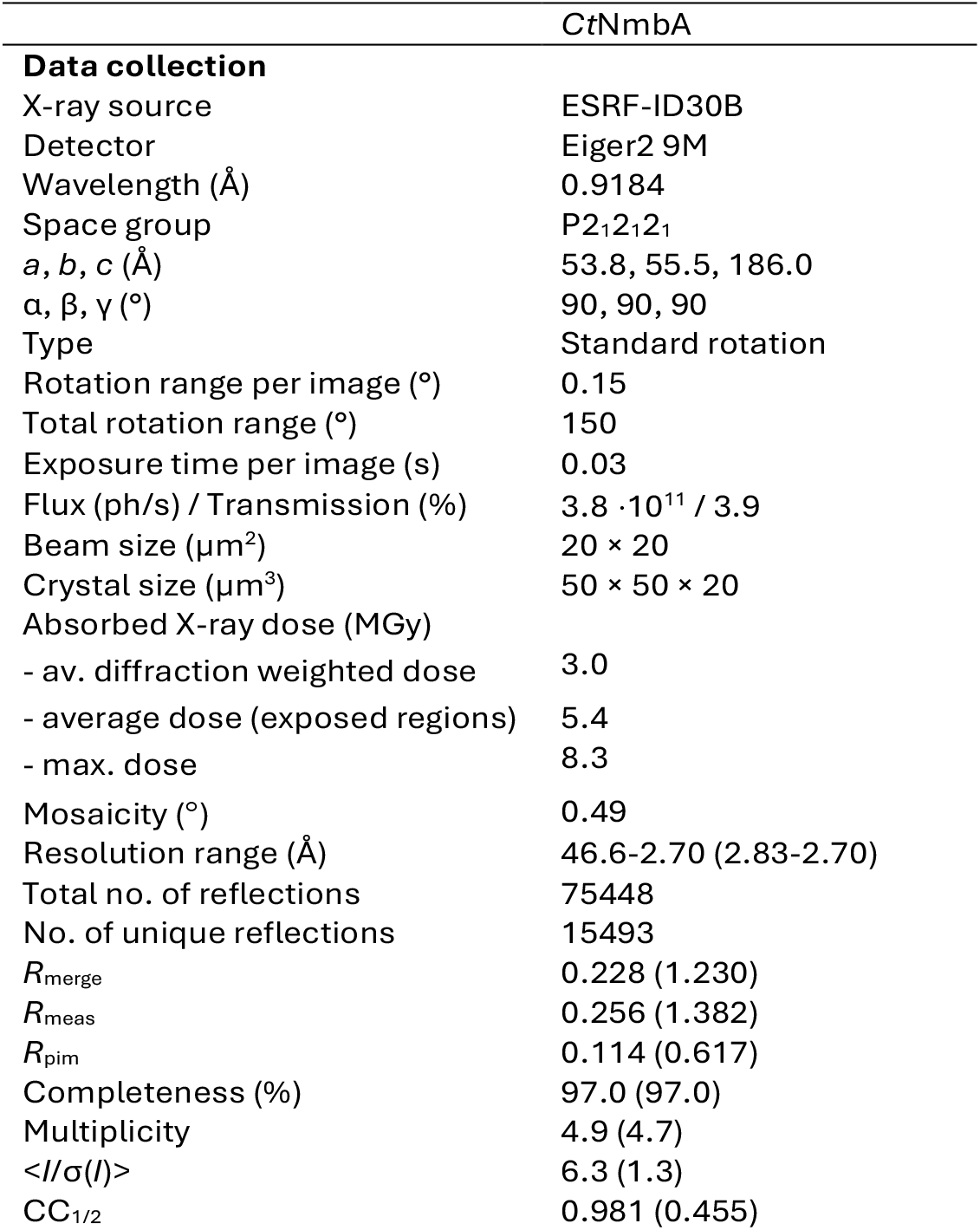
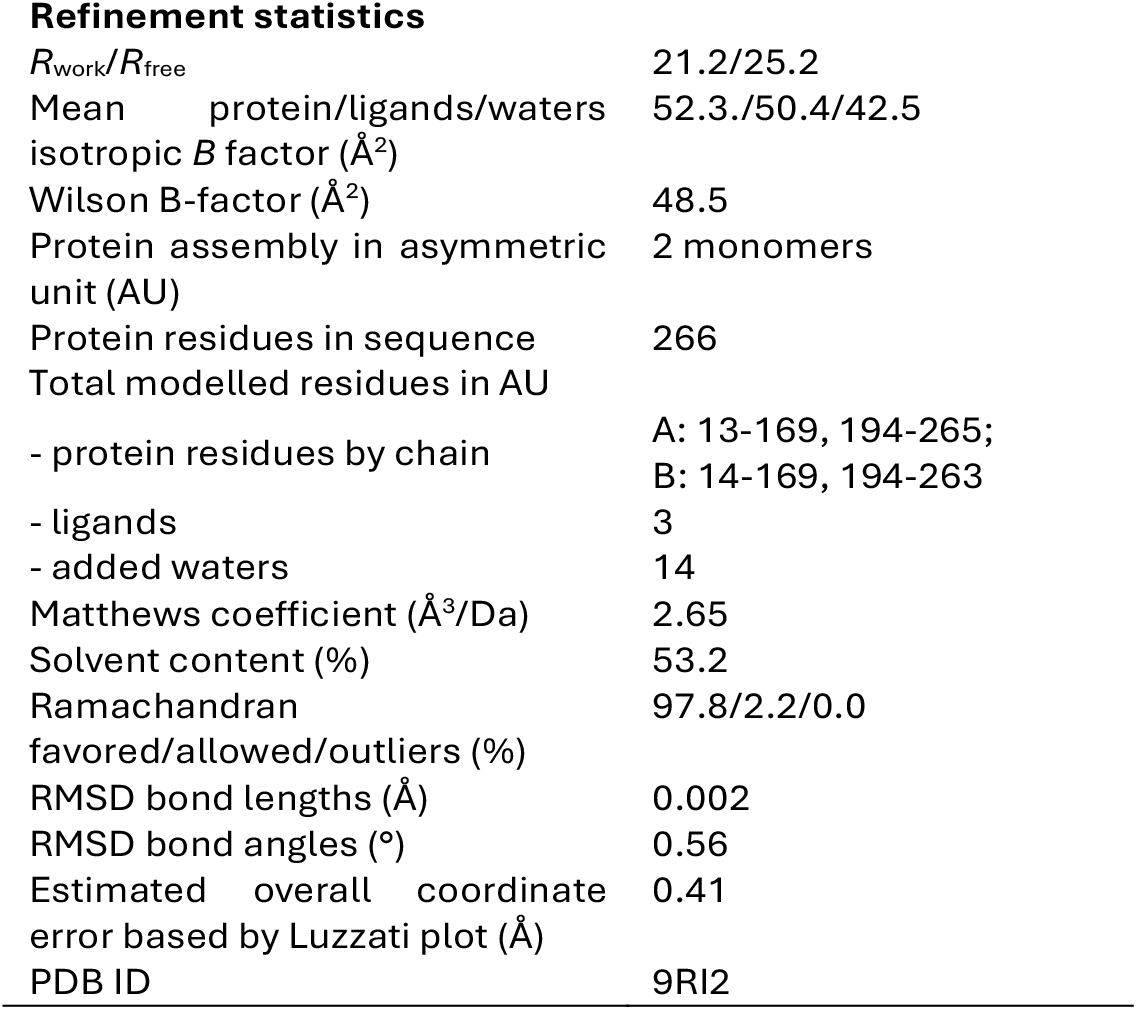
Crystallographic data-collection and refinement statistics. Values given in parentheses are for the outer shell.

**Figure 3.**
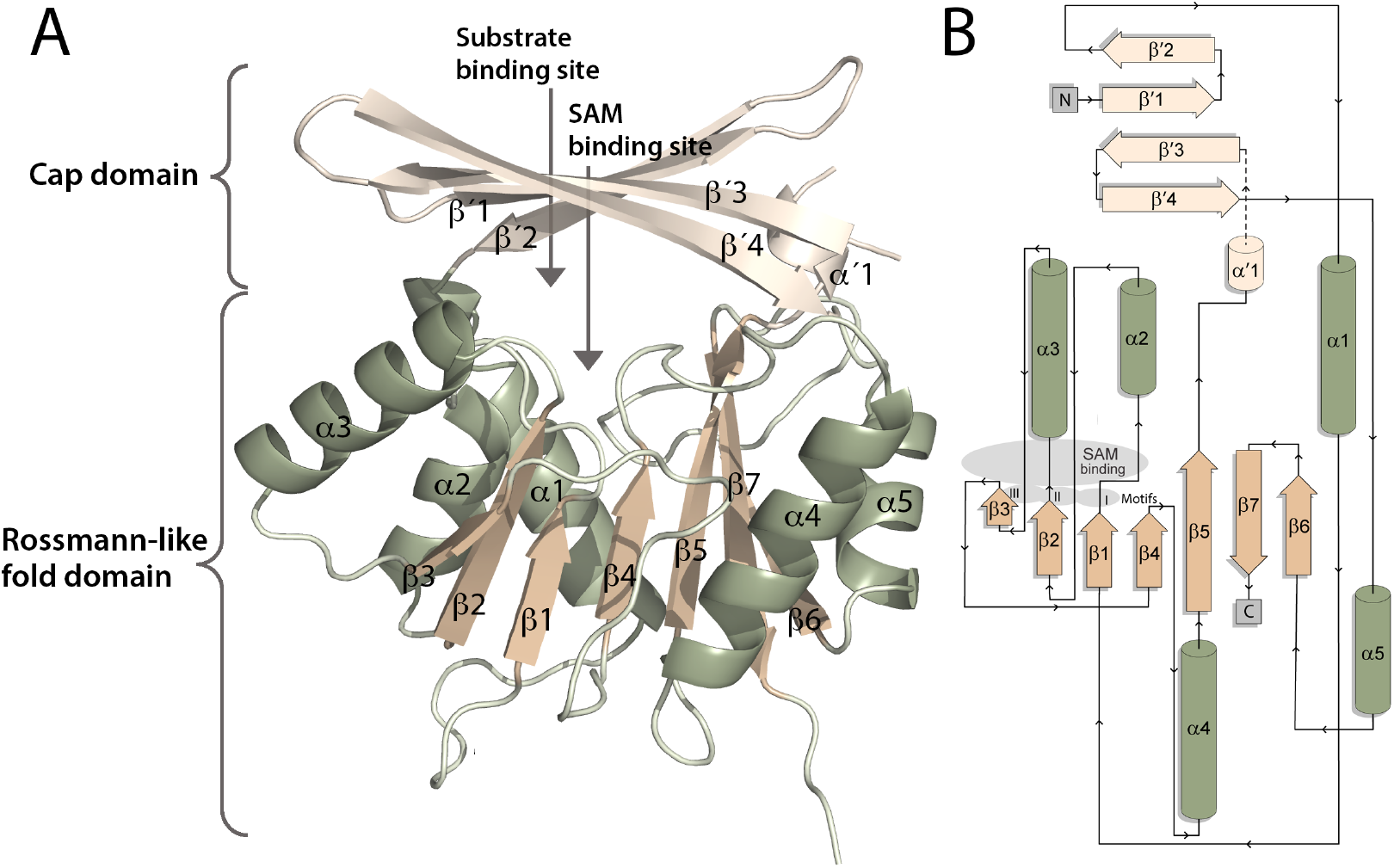
Crystal structure of *Ct*NmbA. (**A**) Overall structure of NmbA displaying the 7BS sheet (β1-7, dark beige) flanked by the α-helices (α1-5, green) of the Rossmann-like fold domain, and the Cap domain (α’1 and β’1-4, light beige). The SAM and substrate binding sites are wedged between the two domains. (**B**) Topology of *Ct*NmbA, colored according to the structure in (**A**).

#### Cap domain

As is often the case for 7BS-MTs, NmbA contains an additional domain at the top of the canonical Rossmann-like fold, typically involved in recognition of the substrate. The active site is wedged between the Rossmann-like fold domain and the substrate-binding domain, with the SAM cofactor binding site located on the lower side of the cleft, allowing it to interact with the conserved Motifs of the Rossmann-like fold. The variable substrate-binding domain consists of a twisted β-sheet of four antiparallel strands, acting as a cap enclosing the active site (Figure 3). Strikingly, due to an unusual architecture of the β-strands of this “Cap-domain”, NmbA adopts a unique tertiary structure. In *Ct*NmbA, the Cap domain encloses the active site to form a narrow molecular basket, consequently forming a different overall structure and a more enclosed active site than predicted by the *ab initio* AlphaFold (AF) 3 model of *Ct*NmbA (Figure 4). More importantly, the confined active site in *Ct*NmbA is caused by a unique organization of the β-strands making up the Cap domain; β’1-β’4. The N-terminal β’1 is directly followed by β’2, which proceeds on to α1 of the Rossmann-like fold domain. β’3-β’4 make up the second segment of the Cap domain (together with α’1 and an unmodelled region of the structure comprising 24 amino acids), succeeding β5 of the 7BS sheet, and continuing to α5 of the C-terminal part of the protein. This Cap-domain assembly results in a contorted and non-continuous four-stranded β-sheet of two subdivided segments, β’1-β’2 and β’3-β’4, where only the middle section of the β’1 and β’3 strands is connected through hydrogen bonds. This topology of the β’1-β’4 differs from the topology predicted by the AF model (Figure 4C), however, the organization in Figure 3 is confirmed by the electron density, as shown in Figure 5A. Here, a continues electron density map is consistent with the loops connecting these β-strands in the final modelled structure. Modelling the topology predicted by AF and re-refining the structure (Figure 5C) results in negative electron density for the alternative loop connections and positive electron density for the removed loop with respect to the refined crystal structure. Consequently, the organization of the β-strands in the experimental structure results in a narrower and more rigid Cap domain, which is unlikely to undergo a conformational rearrangement upon catalysis, unable to create a wider entry site for the substrate. This structural feature of the *Ct*NmbA structure might, however, promote the BSH substrate to adopt an appropriate conformation for the methylation reaction.

**Figure 4.**
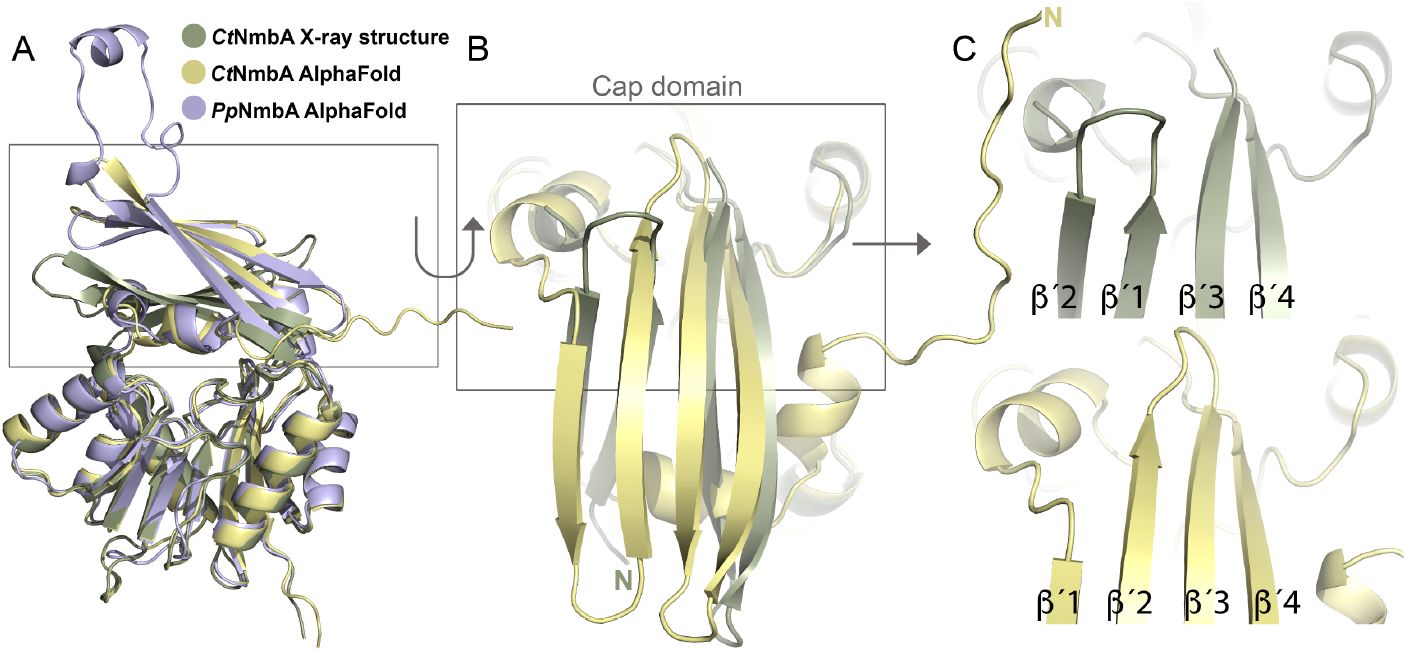
Structural alignment of *Ct*NmbA with AF models. (**A**) Alignment of the *Ct*NmbA crystal structure with the corresponding AF model, as well as the AF model of *Pp*NmbA, depicting the discrepancy in Cap domain architecture. (**B**) Close-up view of the *Ct*NmbA Cap domain showing the differences in the assembly of b-strands b′1-4 in the crystal structure (green) versus the AF model (yellow).

**Figure 5.**
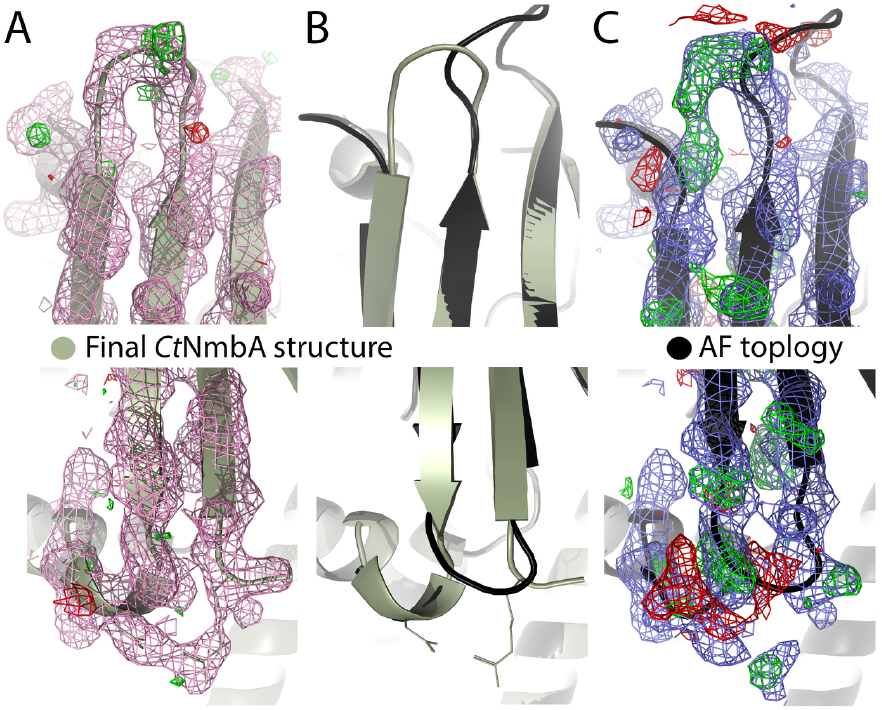
Electron density for the *Ct*NmbA Cap domain. The topology of the Cap domain (**A** and **B**), supported by the electron density, differs from the topology proposed by AF (**B** and **C**). The Cap domain architectures in the crystal structure (**A**) and AF model (**C**) are overlayed in (**B**), showing the discrepancy in the organization of the β-strands. The 2F_o_-F_c_ map is contoured at 1σ (colored in pink/blue) and the F_o_-F_c_ maps contoured at ±3σ (colored in green/red).

#### *Pp*NmbA

In order to gain more insight into the unique structural features of NmbA, we aimed to compare *Ct*NmbA to the structure of the homologous NmbA from *P. polymyxa,* a representative member of the Bacillota found to encode the complete biosynthetic pathway for *N*-Me-BSH (Supplementary Dataset 1, Figure S1) and shown to methylate BSH (Figure 2). Due to the lack of obtaining diffracting crystals, *Ct*NmbA was compared to the *Pp*NmbA AF model, which, as expected, displays the same overall structural architecture as the *Ct*NmbA AF model (in addition to an extended loop protruding from the Cap-domain between β’3 and β’4) (Figure 4A). Although a more disclosed overall organization of the Cap domain is seen in the crystal structure as compared to the AF models, there are examples of NPMTs with similarly confined substrate binding sites. Human phenylethanolamine *N*-methyltransferase (PNMT) (PDBid:3HCD), which catalyzes the conversion of noradrenaline to adrenaline, has adopted a tight substrate specificity pocket, exemplifying the ability of NPMTs to evolve and adapt to form specialized enzyme-ligand interactions.

#### Structural comparison

Structural comparison of *Ct*NmbA with deposited PDB structures using the *DALI* protein structure comparison server shows that the NmbA Rossmann-like core fold is highly similar to other homologous MTs (Table 2). Although high structural similarity is seen with the top selected hits presented in Table 2, the structural organization of the Cap domains differ, varying between wider substrate binding pockets due to a more open architecture, and Cap domains with a higher or exclusive content of α-helices (Figure 6). It is striking that the AF models of NmbA resemble the DALI hits in Figure 6B with a more open Cap domain and an α-helix positioned between the β’-sheet and the 7BS domain. Interestingly, the most similar MT crystal structures all show lower sequence identity with *Ct*NmbA as compared to the NmbA homologs in bacteria encoding the complete *N*-Me-BSH biosynthetic pathway in our bioinformatics investigations. The structures from the DALI hits (Table 2) do not cluster together with the main *N*-Me-BSH-containing phyla of the SSN network (Figure S1) when included in the search, but stand out as single entities in the SSN. Together, our findings suggest that *Ct*NmbA belongs to a new class of NPMTs, specifically targeting the cysteine N-atom of BSH.

**Table 2.**
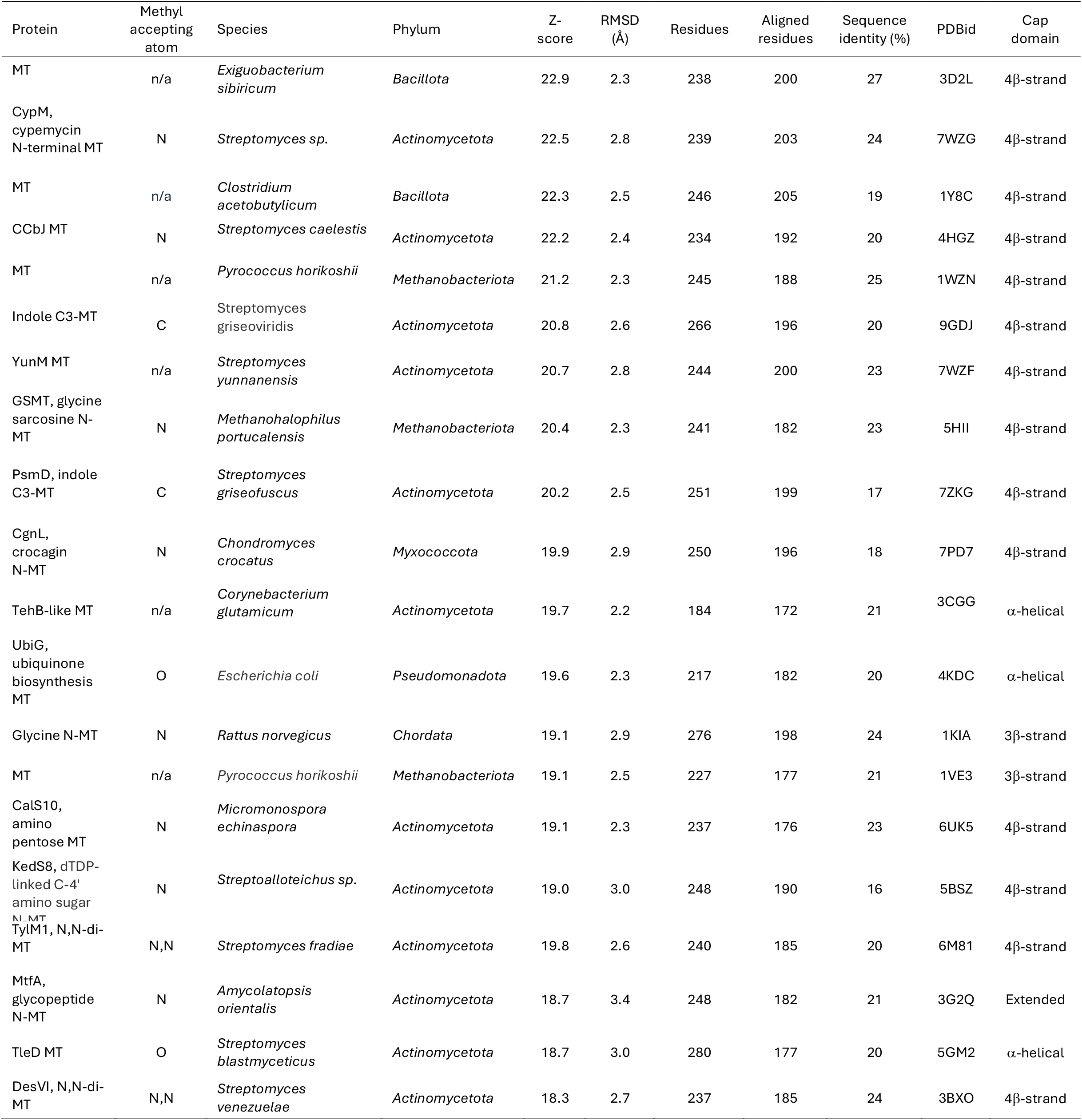
Structural comparison of *Ct*NmbA with selected similar MTs, based on Z-score, from the DALI search. All MTs are SAM-dependent.

**Figure 6.**
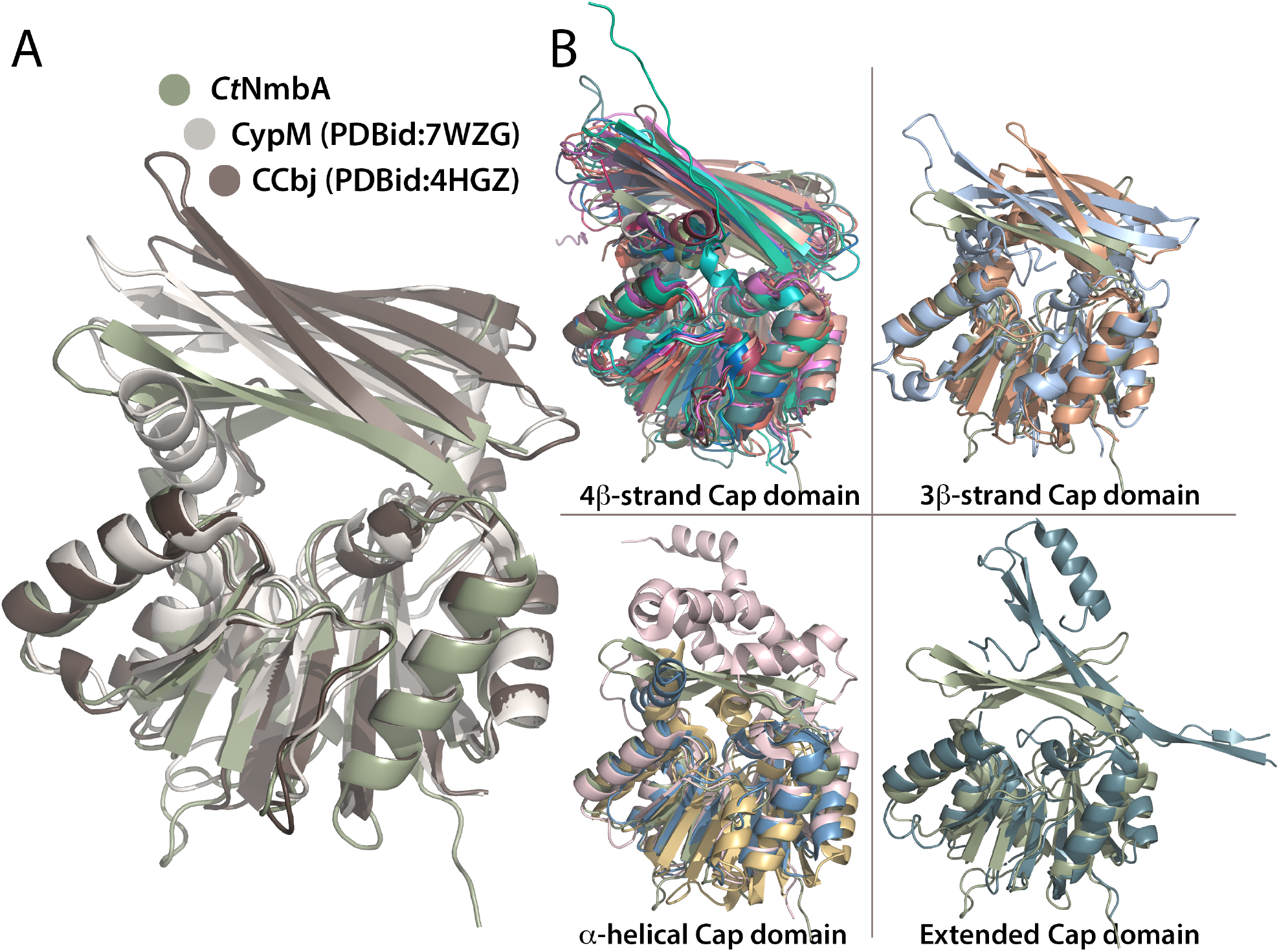
Different Cap domain architectures in MTs listed in Table 2. Structural alignment of MTs from the DALI-search (PDB codes listed in Table 2), similar to *Ct*NmbA, showing variations in the Cap-domain. (**A**) Alignment of *Ct*NmbA with MTs CypM and CCbj ^41^, illustrating the different sizes of the active sites. (**B**) Variations in the Cap domain architecture and substrate binding site emphasized through the structural alignments of *Ct*NmbA with homologous structures with Cap domains dominated by four β-strands (top left), three β-strands (top right), α-helices (bottom left), and an extended architecture (bottom right).

### The BSH binding site supports the methylation reaction

To further elucidate whether the *Ct*NmbA structure can accommodate and position the substrate for catalysis, the binding of BSH was examined through docking studies. Structure-based docking calculations result in binding of BSH in the expected substrate binding pocket of NmbA (Figure 7A), placing the N-atom of the BSH cysteine in proximity to the sulfonium group of SAM (Figure 7B). The binding pocket for SAM and BSH is lined with conserved residues. As described above, the Cap domain in NmbA generates a relatively tight basket that nicely accommodates BSH in a position supported by several putative polar contacts, which could reasonably facilitate catalysis (Figure 7C). SAM-dependent methylation occurs via a nucleophilic S_N_2-like substitution mechanism with a typical methyl donor-to-acceptor distance of 3-4 Å ^42^. Our structural data could comply with a classical, so-called “proximity and desolvation” mechanism, where the methyl acceptor is optimally positioned in close proximity to the methyl donor, facilitating the nucleophilic substitution. Such mechanism fits with the NmbA-BSH docked structure. An alternative MT mechanism involves acid/base catalysis, where a base is required for the deprotonation of the substrate prior to the nucleophilic attack. In the NmbA structure, conserved arginine residues are located in the substrate binding cleft, however, not in close proximity to the acceptor cysteine N-atom of the docked BSH substrate. Unraveling the exact mechanism employed by NmbA requires further studies.

**Figure 7.**
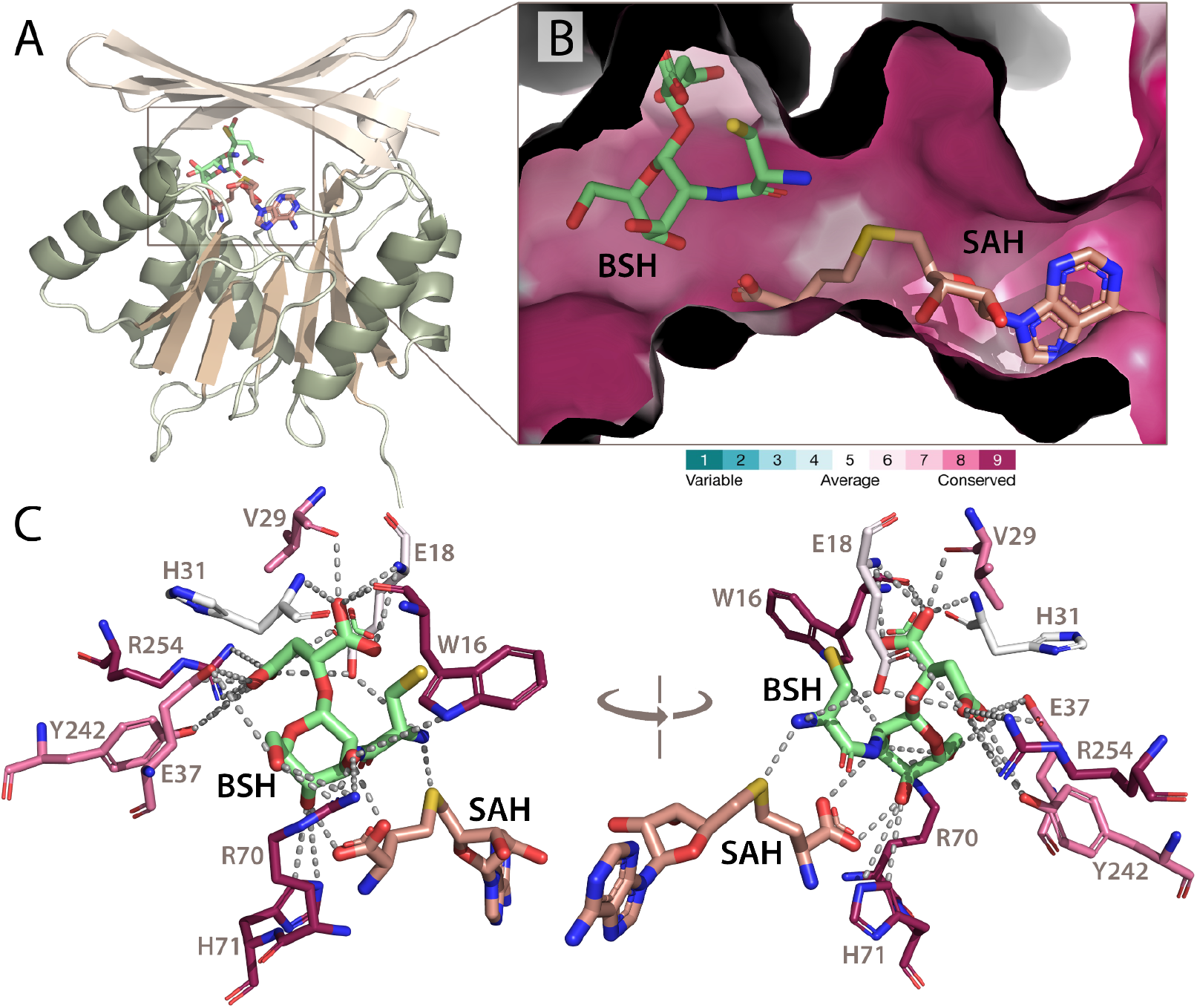
The putative SAM and BSH binding site in *Ct*NmbA. (**A**) Overall structure of *Ct*NmbA with the proposed binding site for the SAM co-substrate and the docked BSH substrate, wedged between the Rossmann-like fold domain and the Cap domain. (**B**) Surface representation of the binding pocket of *Ct*NmbA showing the cysteine N-atom of BSH bound in proximity to the sulfonium atom of SAH, which is formed by the demethylation of SAM. Amino acid residues lining the putative active site, proposed to be involved in electrostatic interactions and hydrogen bonds with the substrate, are shown in (**C**). In (**B**) and (**C**), the degree of conservation of residues is evaluated with *ConSurf*. Variable residues are colored turquoise and highly conserved residues are colored maroon. Cofactors, catalytic residues and substrates are represented as sticks and colored by atom type. Coordinates for SAH were taken from PDBid:7WZG.

## CONCLUSION

Our biochemical results established that NmbA is a novel SAM-dependent methylase of the LMW thiol BSH, which is the first reported case of a functionally important LMW thiol modified through methylation, and a rare instance of *N*-methylation of a cysteine in a secondary metabolite. NmbA shows substrate selectivity for BSH over its oxidized disulfide form, BSSB, suggesting that the methylated form of BSH is utilized in LMW thiol-dependent biochemical processes. NmbA does not show any methylase activity towards amino acid glycine nor GSH, proving its role as a specific *N*-directed MT of the cysteine moiety of BSH. The hereby presented three-dimensional structure of NmbA provides a detailed characterization of the enzyme, which together with our docking calculations and the proposed binding site for the BSH substrate may provide a structural platform for elucidating the mechanism of enzymatic catalysis. The unique architectural organization of the substrate binding domain of NmbA, together with the phylogenetic distribution of NmbA homologs, assign these enzymes to a new class of NPMTs, evolved to specifically methylate the biothiol BSH. Our bioinformatics analyses show that the biosynthetic machinery for *N*-Me-BSH is present in more than 1700 unique bacterial species distributed across Chlorobiota, Bacteriodota, Bacillota and several other phyla, suggesting that *N*-Me-BSH is a physiologically important LMW across many, in particular, aquatic and marine bacteria. From the latter findings, it is evident that through evolution, numerous bacterial species have adapted to utilize the methylated form of BSH. To which degree the biophysical properties of *N*-Me-BSH are altered relative to BSH remains to be determined. Methylation of the amino group of the BSH cysteine moiety could, through its electron-donating properties, stabilize the protonated state of the amine, which could consequently increase the nucleophilicity and reactivity of the cysteine thiol ^24^. Increased hydrophobicity and steric effects caused by the *N*-methylated cysteinyl amine might also influence its affinity and binding properties to target proteins. Besides their roles in maintaining the cellular redox balance, LMW thiols have shown emerging roles in various cellular processes including virulence regulation, host-microbe interactions, and generation of bioactive by-products following their degradation ^43^. Although further physiological studies are needed to elucidate how *N*-Me-BSH influences host physiology, our work provides important structural and biochemical knowledge to the growing record of alternative LMW thiols and their metabolism, and lays the basis for potential applications involving specialized bacteria utilizing *N*-Me-BSH.

## EXPERIMENTAL SECTION

### Bioinformatic analyses – sequence searches, analysis of conserved residues, and sequence similarity networks (SSNs)

The RefSeq: NCBI (National Center for Biotechnology Information) Reference Sequence Database, which is a comprehensive, integrated, non-redundant, well-annotated set of sequences was searched to identify organisms containing homologs of the bacillithiol synthesis enzymes BshA, BshB and BshC, and the bacillithiol MT NmbA. To identify the organisms, a Protein BLAST (Basic Local Alignment Search Tool) search was performed through the NCBI webpage. Individual searches were performed using the protein sequences of the enzymes from *Chlorobaculum tepidum*: *Ct*BshA (locus tag: CT0548), *Ct*BshB (locus tag: CT1419), *Ct*BshC (locus tag: CT1558), and *Ct*NmbA (locus tag: CT1040). No members of the Eukaroyta and Archaea showed to contain homologs of BshA, BshB, and BshC. Each of the 43 bacterial phyla that have been validly published according to the Bacteriological Code ^33-34^ were individually searched for homologs of CT0548, CT1419, CT1158, and CT1040, and for each phylum, the search was supplemented with one identified homolog within that phylum. A requirement of a minimum BLAST E-value of 1^-20^, a similar length of query and subject, coverage above 80%, and sequence identity above 25% was set. Unique organisms within each phyla containing the complete BSH biosynthetic pathway or the complete *N*-Me-BSH biosynthetic pathway were then identified requiring a unique bacterial name, and for bacteria with several strains only the member with the core bacterial name was included, otherwise the strains with the highest E-values. The degree of conservation of residues in the *Ct*NmbA structure was evaluated with ConSurf ^44-46^. The analysis was based on identification of homologous sequences from the UniRef90 database using the HMMER algorithm ^47^, and multiple sequence alignment with MAFFT ^48^. Of the 1511 homologous sequences passing the threshold of 35-95% sequence identity, ConSurf selected a sample of 150 representative sequences. A 9-bin colored scale was used to show the conservation of each residue, from most variable (turquoise) to the most conserved (maroon), when generating three-dimensional figures with PyMOL. A sequence similarity network (SSN) was generated with the Web-based Enzyme Function Initiative tool – Enzyme similarity tool EFI-EST (https://efi.igb.illinois.edu/efi-est/) ^49^ using *Ct*NmbA as a search sequence. The SNN was based on the 1511 homologous sequences generated by ConSurf (described above). Sequences were grouped with an alignment score of 50 and nodes representing sequences sharing >90% identity. The largest clusters of sequences were selected and assembled into nine phyla groups. Figures illustrating SSN analyses were created in Cytoscape (version 3.9) with the organic layout ^50^.

### Expression and purification of CtNmbA and PpNmbA

The genes for *Chlorobaculum tepidum* TLS NmbA (*Ct*NmbA, locus tag: CT1040) or *Paenibacillus polymyxa SC2* NmbA (*Pp*NmbA, locus_tag: PPSC2_19805) were codon optimized and cloned into a pET-22b(+) plasmid (constructed using NdeI and HindIII sites) (GenScript) and transformed into competent *Escherichia coli* One Shot^TM^ BL21 (DE3) cells (Invitrogen, Thermo Fischer Scientific). Cells were grown in Terrific Broth medium containing 100 μg/mL ampicillin, protein expression was induced by adding isopropyl β-D-thiogalactoside (IPTG) to a final concentration of 0.5 mM at OD_600nm_ = 0.7-0.8, and the cultures were incubated for 16 hours at 18°C with vigorous shaking before cells were harvested and frozen at -20°C. Cells were thawed and dissolved in 50 mM Tris-HCl pH 7.5, 2 mM DTT, 5 μg/mL DNase, cOmplete Protease Inhibitor Cocktail tablet (Roche) in a 1:4 cell we weight to buffer ratio and lysed by sonication. *Pp*NmbA was precipitated with 0.4 g/mL (NH_4_)_2_SO_4_, whereas for *Ct*NmbA, contaminant proteins were first precipitated with 0.2 g/mL (NH_4_)_2_SO_4_) before *Ct*NmbA was precipitated by adding (NH_4_)_2_SO_4_ to the remaining supernatant to a final concentration of 0.4 g/mL. Proteins were dissolved in 50 mM Tris-HCl pH 7.5, 2 mM DTT, and desalted through dialysis (SnakeSkin™ dialysis tubing, 10 kDa MWCO, ThermoFisher Scientific). Desalted proteins were filtered through a 0.45 μm filter (Sarstedt), applied to a HiTrap HP Q column (Cytiva) and eluted with linear 0-0.2 M KCl or 0-0.25 M KCl gradients for *Ct*NmbA and *Pp*NmbA, respectively, using 50 mM Tris-HCl pH 7.5, 1 mM DTT, 1 M KCl. Proteins were further purified on a Superdex 75 or 200 Increase 10/300 GL column (Cytiva) in 50 mM HEPES pH 7.5, 100 mM NaCl, indicating monomeric assemblies of *Ct*NmbA (30.5 kDa) and *Pp*NmbA (29.7 kDa). As a final polishing step prior to crystallization, proteins were purified on a MonoQ 5/50 GL column with linear 0-0.3 M KCl gradients using 50 mM Tris-HCl pH 7.5, 1 M KCl. Protein fractions were pooled, concentrated in Amicon Ultra-15 filter units (10 kDA MWCO, Merck-Millipore), flash-frozen in liq N_2_, and stored at – 80 °C. All chromatographic steps were performed on an Äkta purifier FPLC system (GE Healthcare) and all expression and purification steps were monitored by sodium dodecyl sulfate-polyacrylamide gel electrophoresis (SDS-PAGE). Protein concentrations were estimated using an Agilent Cary 60 spectrophotometer and the extinction coefficient for *Ct*NmbA, ε_280 nm_ = 32.43 mM^-1^cm^-1^ and for *Pp*NmbA, ε_280 nm_ = 44.35 mM^-1^cm^-1 51^.

### Preparation of S-adenosyl-L-methionine (SAM) and S-adenosyl-L-homocysteine (SAH) stock solutions

SAH (Sigma-Aldrich) was dissolved in dimethyl sulfoxide (DMSO) to a final concentration of 182 mM and incubated in an ultrasonic water bath until solution turned clear, whereas SAM (Sigma-Aldrich) was dissolved in mqH_2_O to a final concentration of 10 mM.

#### Protein crystallization

All initial crystallization screening of both NmbA proteins was performed using the sitting drop vapor diffusion crystallization method with a Mosquito robot (SPT Labtech). *Ct*NmbA and *Pp*NmbA were initially screened at various protein concentrations in their native, apo states as well as attempted co-crystallized with SAH in a 1:5 protein:SAH ratio. Native *Ct*NmbA crystals (13 mg/mL) were obtained with condition H4 from the PEG/Ion HT crystallization screen (Hampton Research) (0.03 M citric acid, 0.07 M BIS-TRIS propane, pH = 7.6, 20 % w/v polyethylene glycol 3,350) and native *Pp*NmbA crystals (11.5 mg/mL) were obtained with condition D11 from the Wizard Cryo crystallization screen (Molecular Dimensions) (0.1 M sodium acetate/acetic acid, pH = 4.5, 0.2 M lithium sulfate, 50 % w/v polyethylene glycol 400). Conditions that identified initial hits were further attempted optimized by systematic optimization using the sitting drop vapor diffusion method. Crystals were grown at 20 ºC, cryoprotected in 50% (w/v) polyethylene glycol 400 (*Pp*NmbA) or 25% (w/v) polyethylene glycol 400 (*Ct*NmbA) and flash-frozen in liquid nitrogen prior to data collection.

#### Crystal Data Collection, Processing, and Refinement

For the apparent native *Ct*NmbA and *Pp*NmbA crystals, diffraction data were collected at beamline ID30B at the European Synchrotron Radiation Facility (ESRF), Grenoble, France, and beamline BioMAX at MAX IV, Lund, Sweden, respectively. The *Ct*NmbA crystals diffracted beyond 3 Å, however, the *Pp*NmbA crystals did not diffract. The diffraction data of *Ct*NmbA crystals were indexed and integrated through auto-processing with autoPROC ^52^ and XDS ^53^, and scaled and merged with Aimless in the CCP4 package ^54^. The best dataset was scaled to 2.7 Å. The *Ct*NmbA structure was solved with molecular replacement (MR) with PHASER ^55^ using a search model of *Ct*NmbA from the AlphaFold Protein Structure Database ^56-57^ (Uniprot: Q8KDK7) with a TFZ score of 8.3 and 17.8 for the two molecules in the asymmetric unit. The 7BS Rossmann-like fold domain gave a good fit with the electron density, however, the Cap domains from the two monomers clashed into each other, and were therefore deleted in a step-wise manner and manually rebuilt. The Cap domains could be rebuilt to avoid clashing between the two monomers and with a good fit to the electron density, except for residues 170-193, which did not show any electron density. The disorder of these residues was also indicated by the D_2_P_2_: Database of Disorder Protein Predictions ^58^. The rebuilt model resulted in a different topology and orientation of the Cap domain as compared to the AlphaFold model, where the re-modelled topology for the β’1-β’4 strands is supported by the electron density. The first rounds of refinements were performed using restrained refinement in REFMAC5 ^59^ followed by several cycles of refinement with phenix.refine ^60^ in the Phenix suite ^61^ performed in iterative cycles with model building performed in Coot ^62^. The final refinement included refinement of XYZ in reciprocal space, translation-liberation-screw (TLS) rotation factors, individual *B*-factors and applying non-crystallographic symmetry (NCS) restraints and optimizing X-ray/stereochemistry/atomic displacement parameter weight. Model validation was performed using MolProbity ^63^. The absorbed X-ray dose was calculated using the program RADDOSE-3D ^64^. PDBePISA ^65^ was used to confirm that NmbA is a biological monomer, despite and having two monomers in the asymmetric unit, since analysis of the protein interfaces did not reveal any specific interactions that could result in the formation of a stable quaternary structure. All structure figures were prepared using PyMOL version 3.1.4.1 (Schrödinger, LLC). The secondary structure toplogy diagram for *Ct*NmbA was generated with PDBsum ^66^ and manually adjusted and colored.

### Structure comparison - structural alignment search with Distance-matrix ALIgnment

A search for similar structures to NmbA in the Protein Data Bank (PDB) was performed with the DALI (Distance-matrix ALIgnment) protein structure-comparison server ^67^ using the *Ct*NmbA structure as a search template.

#### Preparation of bacillitiol (BSH) and bacillithiol disulfide (BSSB)

A solution of reduced BSH was prepared by dissolving its trifluoroacetic salt (Jema Biosciences) in mqH_2_O. As oxidized BSSB is not commercially available, an oxidation of reduced BSH was performed, as described by Hamilton and coworkers ^16^. In short, a solution of NH_4_HCO_3_ was added to reduced BSH (Jema Biosciences) dissolved in water at room temperature and stirred with exposure to air for 90 min, flash-frozen in liq N_2_, and stored at - 80°C. Oxidation of BSH to BSSB was verified with 5,5’-dithiobis-(2-nitrobenzoic acid) (DTNB).

#### Enzymatic assays of NmbA methyltransferase activity and MS analysis

The *in vitro* methylation of BSH and BSSB, as well as glutathione (GSH, Sigma-Aldrich) and glycine (Sigma-Aldrich) was investigated through enzymatic assays, and analyzed and quantified using liquid chromatography-mass spectrometry (LC-MS). 100 μL reactions were set up in 20 mM Tris-HCl, pH 7.5, with varying concentrations of BSH (10 – 20 μM), BSSB (5-10 μM), GSH (10 μM) or glycin (10 μM), and SAM (15 – 50 μM). The reactions were initiated by the addition of 1-3 μM purified *Ct*NmbA or *Pp*NmbA and incubated at 25 °C for 60 or 120 min. Reactions were quenched and protein was precipitated through incubation at 60 °C for 15 min and precipitated protein was removed by centrifugation at 12,000 x g for 5 min. Control reactions where either of the reactants were omitted were also run and treated in the same manner. All reactions were run in multiple parallels. All LC-MS data were collected using a Bruker Daltonics maXis II QTOF high-resolution mass spectrometer and a Dionex UltiMate 3000 UHPLC system. Separation was achieved by using an Adsorbosphere XL CN column (100 x 4.6 mm, 3 μm; Alltech) held at 25 °C, mobile phases A: water +0.1 % formic acid, mobile phase B: methanol + 0.1% formic acid. The flow rate was held at 1 mL/min, at 90 % B for 1 min, followed by a linear gradient to 10 % B over 0.5 min, held at 10 % B for 7 min, then brought up to 90 % over 1 min and a hold at 90 % for 2 min to equilibrate the column (total run time 11.5 min). The injection volume was 5 μL. The source conditions for electrospray ionization MS (ESI-MS) were as follows: capillary voltage: 4kV, nebulizer gas flow: 5.0 bar, dry gas flow: 12.0 L/min at 250 °C. The sample composition was determined by relative quantification from the peak area of extracted ion chromatograms of each protonated target peak. (Bruker Compass DataAnalysis 4.3).

#### Docking analysis and protein-ligand interactions

Structure-based blind-docking calculations between *Ct*NmbA and BSH were performed with *CB-Dock*2 (*Cavity-detection guided Blind Docking*) ^68^. The rotamer geometry of selected amino acid side chains at the putative substrate/co-substrate binding and entry sites (W16, W20, H31, R70, and K198) were slightly adjusted and regularized with respect to stereochemical constraints in Coot and PyMOL. The SAH coordinates were taken from PDBid:7WZG, and included in the docking calculations.

## SUPPORTING INFORMATION

### Supporting Information is available

Bioinformatics analyses and SSN, LC-MS analysis, scheme of *N*-Me-BSH biosynthetic pathway Dataset 1; complete list of species containing the BSH/*N*-Me-BSH biosynthetic pathway.

## AUTHOR CONTRIBUTION

M.H. and H.P.H performed the bioinformatics, biochemical, and structural studies and analyzed the data. E.S performed the MS and analyzed the data. M.H. and H.P.H wrote the manuscript with input from E.S. All authors have given approval to the final version of the manuscript.

The authors declare no competing financial interest.

## ACKNOWLEDGMENT

This project has been funded by a grant from the Research Council of Norway (grant No. 301584) through support from the University of Oslo and the UiO Structural Biology Core Facilities (PX-Oslo). The authors thank the Regional Core Facility for Structural Biology at the Oslo University Hospital for access to crystallization screening. We acknowledge the European Synchrotron Radiation Facility (ESRF) and MAX IV for the provision of synchrotron-radiation facilities, and the staff of beamlines ID30B (ESRF) and BioMAX (MAX IV) for their assistance. The diffraction data for *Ct*NmbA were collected at the ESRF and have been archived in the ESRF depository (https://doi.org/10.15151/ESRF-DC-XXXX).

## ABBREVIATIONS

7BS: seven β-strand
AF: AlphaFold
BSH: bacillithiol
BSSB: bacillithiol disulfide
*Ct*: *Chlorobium tepidum*
DALI: Distance-matrix ALIgnment
GSH: glutathione
LMW: low-molecular weight
Me: methyl
MS: mass spectrometry
MSH: mycothiol
MT: methyltransferase
*N*-Me-BSH: *N*-methyl-bacillithiol
NmbA,: *N*-Me-BSH-synthase
A;NPMT: natural product methyltransferase
PDB: Protein Data Bank
*Pp*: *Paenibacillus polymyxa*
SAM: *S*-adenosyl-l-methionine
SAH: *S*-adenosyl-s-homocysteine
SSN: Sequence Similarity Network.

## Notes

### Competing Interest Statement

The authors have declared no competing interest.

