## Supporting Information for "The methyltransferase NmbA methylates the low-molecular weight thiol bacillithiol, and displays a specific structural architecture"

**Table S1. Distribution of unique species among bacterial phyla encoding the *N*-Me-BSH or BSH biosynthetic pathways*†.**

| Bacterial phylum | # species with homologs of NmbA, BshA, BshB, and BshC | # species with homologs of BshA, BshB, and BshC |
|---|---|---|
| Acidobacteriota | 4 | 51 |
| Bacillota | 366 | 1466 |
| Bacteroidota | 1329 | 2111 |
| Balneolota | 18 | 28 |
| Calditrichaeota | 1 | 1 |
| Chlamydiota | 3 | 3 |
| Chlorobiota | 15 | 15 |
| Cyanobacteriota | 0 | 1 |
| Deinococcota | 1 | 107 |
| Gemmatimonadota | 8 | 13 |
| Ignavibacteriota | 2 | 2 |
| Myxococcota | 2 | 42 |
| Planctomycetota | 2 | 2 |
| Rhodothermota | 13 | 15 |

* Searching remaining bacterial phyla validly published according to the Bacteriological Code gave no hits of members containing the complete BSH nor *N*-Me-BSH biosynthetic pathways. These included: Actinomycetota, Aquificota, Armatimonadota, Atribacterota, Bdellovibrionota, Caldisericota, Campylobacterota, Chloroflexota, Chrysiogenota, Coprothermobacterota, Deferribacterota, Dictyoglomota, Elusimicrobiota, Fibrobacterota, Fusobacteriota, Kiritimatiellota, Lentisphaerota, Mycoplasmatota, Nitrospinota, Nitrospirota, Pseudomonadota, Spirochaetota, Synergistota, Thermodesulfobacteriota, Thermomicrobiota, Thermoproteota, Thermotogota, and Verrucomicrobiota.

† A complete overview of all species in this table, including organism names and accession codes, is listed in Supplementary Dataset 1.

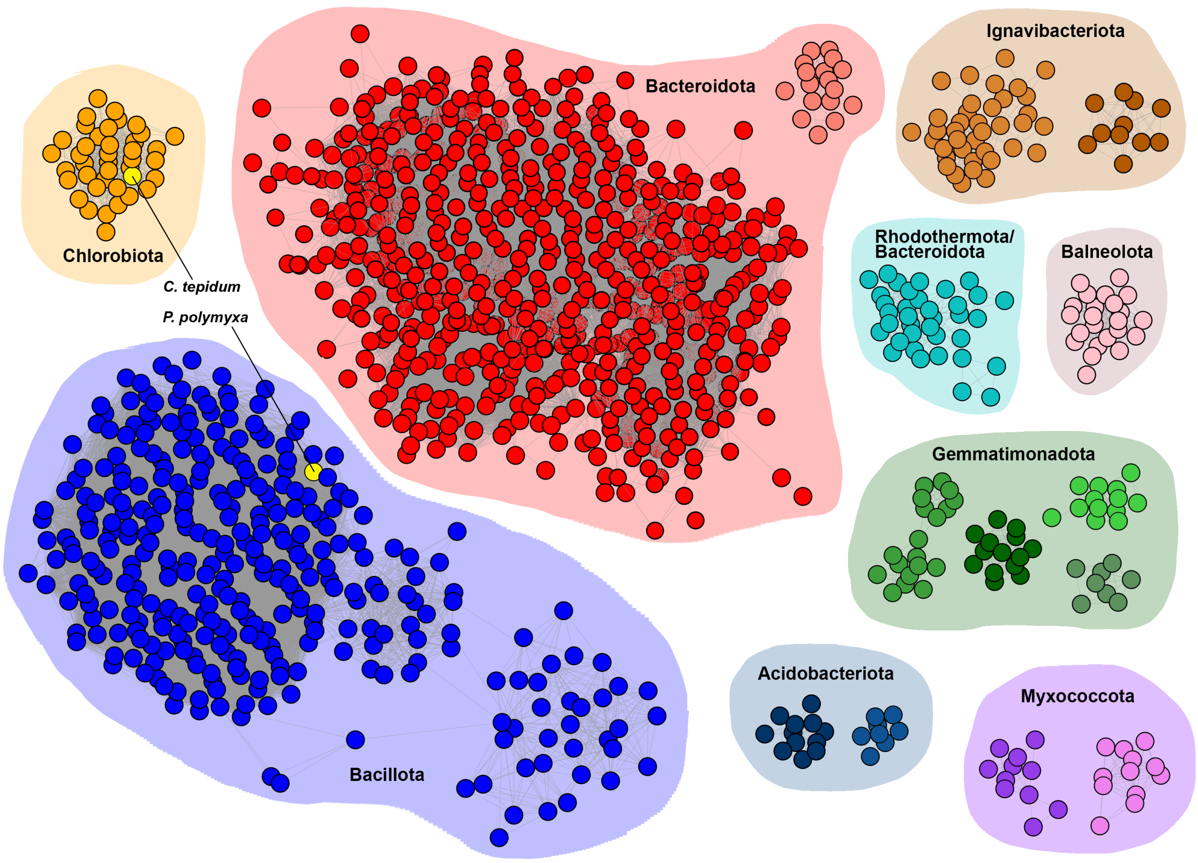


**Figure S1. Comparison of *Ct*NmbA with homologous MTs through sequence similarity networks (SSNs).** The SSN displays the nine largest clusters. The clusters are individually colored and assembled into groups with respect to bacterial phyla. Selected representative species from Chlorobiota and Bacillota encoding the *N*-Me-BSH biosynthetic genes are indicated.

A

| | Total % products, including by-products | | | | | |
|---|---|---|---|---|---|---|
| | Me-BSSB $C_{27}H_{44}N_4O_{20}S_2$ | Me-BSSB-Me $C_{28}H_{46}N_4O_{20}S_2$ | Me-BSSB + Me-BSSB-Me $C_{27}H_{44}N_4O_{20}S_2$ + $C_{28}H_{46}N_4O_{20}S_2$ | $C_{14}H_{22}N_2O_{10}S$ | $C_{15}H_{24}N_2O_{10}S$ | $C_{17}H_{30}N_2O_{10}S$ |
| *Ct*NmbA | 17 ± 7 | 57 ± 10 | 74 ± 7 | 0.5 ± 0.5 | 9.8 ± 9.7 | 10 ± 4.4 |
| *Pp*NmbA | 21 ± 5 | 48 ± 10 | 69 ± 9 | 0.7 ± 0.4 | 13.6 ± 9 | 10.4 ± 4.1 |
| *Ct* + *Pp*NmbA | 19 ± 6 | 53 ± 11 | 72 ± 8 | 0.6 ± 0.5 | 11.7 ± 9.2 | 10.2 ± 4.1 |

B

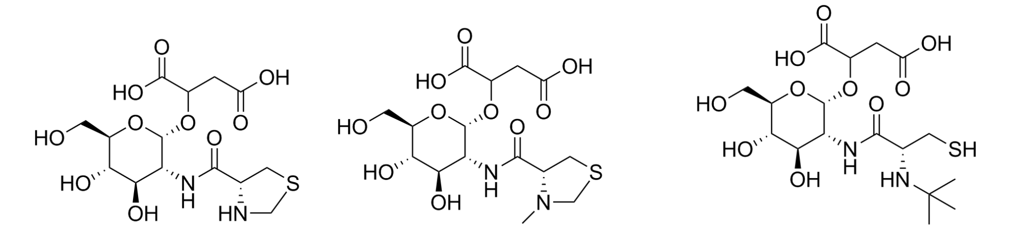


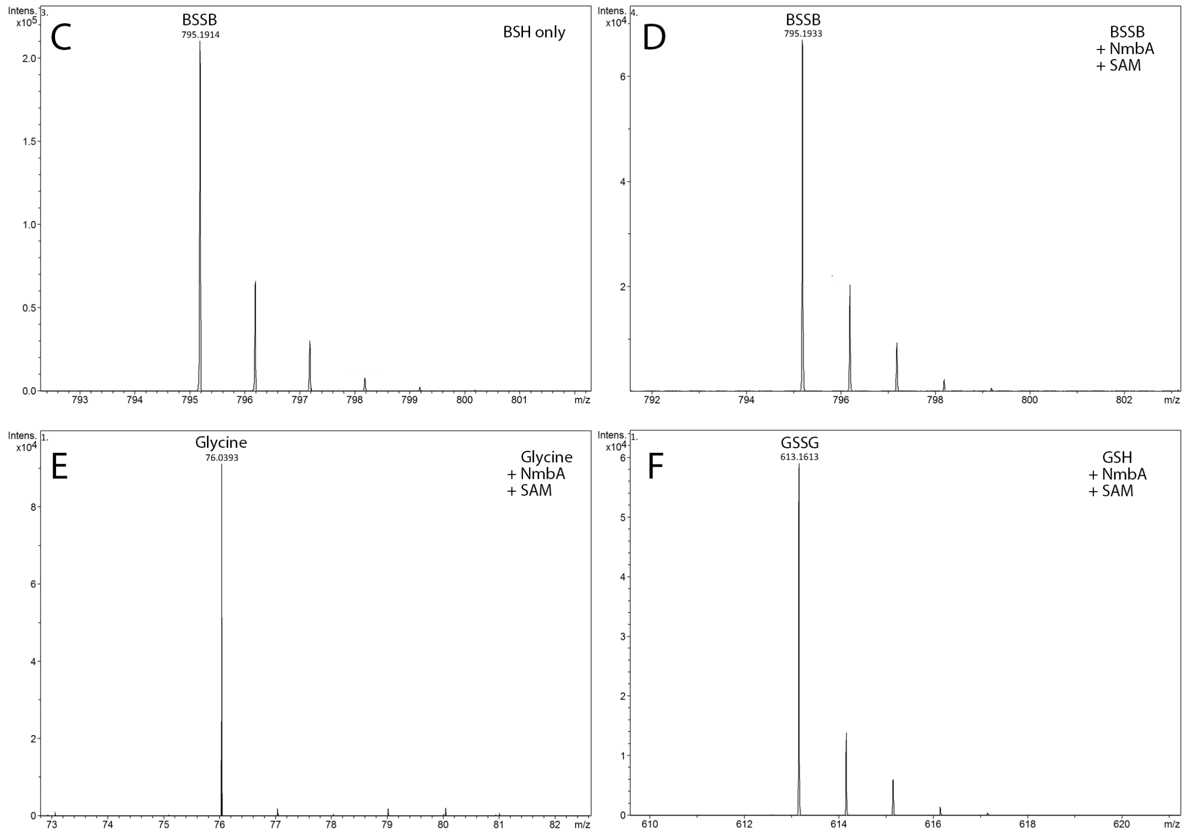


**Figure S2. Mass spectrometry analysis of methylated BSSB products, by-products, and alternative substrates. (A)** The calculated percentage of methylated BSSB forms from the enzymatic reactions using BSH as a substrate and SAM as co-substrate, catalyzed by *Ct*NmbA or *Pp*NmbA, including three by-products. The numbers are calculated from the total BSSB pool (unmethylated BSSB, Me-BSSB, and Me-BSSB-Me) as well as the by-products from the enzymatic reactions. **(B)** Proposed structures of the three detected by-products. **(C)** and **(D)** show representative mass spectra of unmethylated BSSB ($[M + H]^+$795.1914 or 795.1933), from a sample of BSH only **(C)** and from a reaction catalyzed by *Ct*NmbA using oxidized BSSB as a substrate **(D)**. Characteristic mass spectra from reactions using alternative substrates glycine ($[M + H]^+$76.0393) in **(E)** and using GSH (oxidized GSSG, $[M + H]^+$613.1613) in **(F)** catalyzed by *Pp*NmbA show no methylated products.

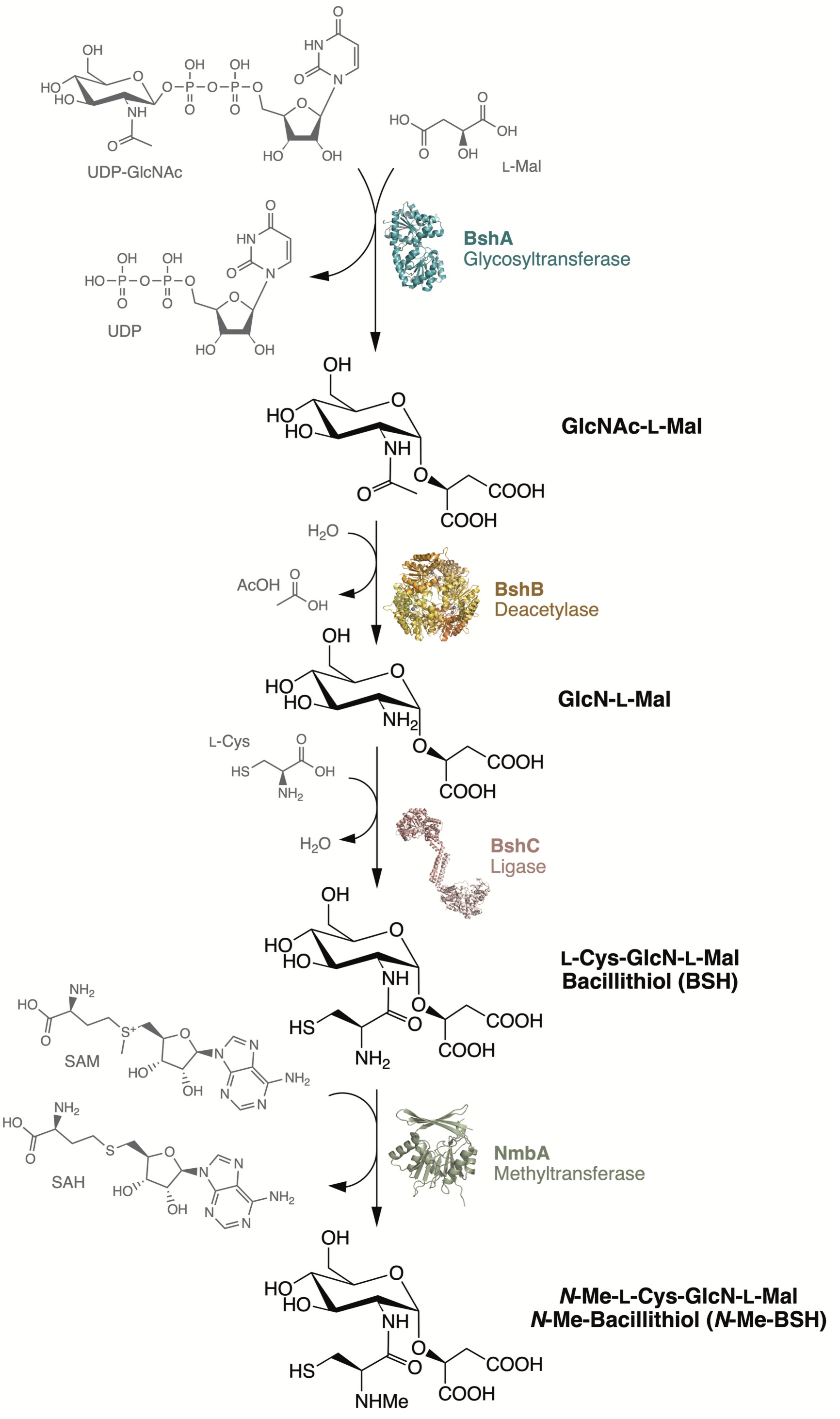


**Figure S3. *N*-Me-BSH biosynthesis pathway.** The glycosyltransferase BshA (PDBid:6D9T [1]) catalyzes the first step in (*N*-Me)-BSH biosynthesis, utilizing UDP-GlcNAc (uridine 5'-diphosphate-*N*-acetylglucosamine) and L-Mal (L-malate) as substrates, resulting in formation of GlcNAc-L-Mal (malyl-*N*-acetyl-D-glucosamine). GlcNAc-L-Mal is further deacetylated by deacetylase BshB (PDBid:6ULL [2]) to GlcN-L-Mal (malyl-D-glucosamine), which together with L-Cys (L-cysteine) serve as substrates for ligase BshC (PDBid:4WBD

[3]) in the formation of BSH. Lastly, in *N*-Me-BSH-producing bacteria, NPMT NmbA catalyzes methylation of the cysteine N-atom of BSH, resulting in *N*-Me-BSH.
