## Supplementary Dataset 1 for "The methyltransferase NmbA methylates the low-molecular weight thiol bacillithiol, and displays a specific structural architecture"

Supplementary Dataset 1A – Unique species containing the complete N-Me-BSH biosynthetic pathway. Phylum, species, and NCBI accession codes for NmbA, BshA, BshB, and BshC are listed for each organism.

| **Phylum** | **Species** | **Accession code NmbA** | **Accession code BshA** | **Accession code BshB** | **Accession code BshC** |
|---|---|---|---|---|---|
| Acidobacteriota | *Chloracidobacterium aggregatum* | WP_211427380.1 | WP_211422436.1 | WP_211423485.1 | WP_211426094.1 |
| Acidobacteriota | *Chloracidobacterium sp. D* | WP_211433143.1 | WP_211432135.1 | WP_211433022.1 | WP_211432012.1 |
| Acidobacteriota | *Chloracidobacterium thermophilum* | WP_014101269.1 | WP_014098778.1 | WP_058865916.1 | WP_014098599.1 |
| Acidobacteriota | *Chloracidobacterium validum* | WP_211430584.1 | WP_211428433.1 | WP_211430360.1 | WP_211428837.1 |
| Bacillota | *Aliibacillus thermotolerans* | WP_270895466.1 | WP_270897557.1 | WP_270897558.1 | WP_270898399.1 |
| Bacillota | *Alkalihalobacillus deserti* | WP_227937551.1 | WP_227936340.1 | WP_227936216.1 | WP_227935718.1 |
| Bacillota | *Alkalihalobacillus sp. LMS39* | WP_243527333.1 | WP_243527612.1 | WP_243527614.1 | WP_243528494.1 |
| Bacillota | *Alkalihalobacterium bogoriense* | WP_026671931.1 | WP_035178327.1 | WP_035178325.1 | WP_026674384.1 |
| Bacillota | *Alkalihalophilus lindianensis* | WP_317123171.1 | WP_317120797.1 | WP_317120796.1 | WP_317122523.1 |
| Bacillota | *Alkalihalophilus marmarensis* | WP_326239376.1 | WP_022627180.1 | WP_326238214.1 | WP_326240074.1 |
| Bacillota | *Alkalihalophilus pseudofirmus* | WP_289235703.1 | WP_075682216.1 | WP_012958526.1 | WP_075682601.1 |
| Bacillota | *Alkalihalophilus sp. As8PL* | WP_368505392.1 | WP_368504480.1 | WP_368504481.1 | WP_368503982.1 |
| Bacillota | *Alteribacillus bidgolensis* | WP_091579508.1 | WP_091579714.1 | WP_245917746.1 | WP_091582817.1 |
| Bacillota | *Alteribacillus iranensis* | WP_091656492.1 | WP_091656672.1 | WP_091656669.1 | WP_091657528.1 |
| Bacillota | *Alteribacillus persepolensis* | WP_091271002.1 | WP_091270857.1 | WP_245705149.1 | WP_091270224.1 |
| Bacillota | *Alteribacillus sp. JSM 102045* | WP_375006769.1 | WP_375006684.1 | WP_375006685.1 | WP_375005851.1 |
| Bacillota | *Ammoniphilus resinae* | WP_209809244.1 | WP_209811430.1 | WP_209811431.1 | WP_209808725.1 |
| Bacillota | *Ammoniphilus sp. YIM 78166* | WP_167577707.1 | WP_134699003.1 | WP_134699002.1 | WP_167577581.1 |
| Bacillota | *Anaerobacillus sp.* | WP_409848335.1 | WP_409852530.1 | WP_409852635.1 | WP_409849617.1 |
| Bacillota | *Aneurinibacillus aneurinilyticus* | WP_276915438.1 | WP_370951626.1 | WP_021622598.1 | WP_021624582.1 |
| Bacillota | *Aneurinibacillus migulanus* | WP_043066703.1 | WP_043067898.1 | WP_235355982.1 | WP_043066087.1 |
| Bacillota | *Aneurinibacillus sp. REN35* | WP_371148367.1 | WP_371146670.1 | WP_371146669.1 | WP_371147049.1 |
| Bacillota | *Aneurinibacillus terranovensis* | WP_027417799.1 | WP_027417519.1 | WP_035102034.1 | WP_027414484.1 |
| Bacillota | *Aneurinibacillus tyrosinisolvens* | WP_047152122.1 | WP_047150554.1 | WP_047152599.1 | WP_052947533.1 |

| | | | | | |
|---|---|---|---|---|---|
| Bacillota | *Aneurinibacillus uraniidurans* | WP_272562244.1 | WP_272560005.1 | WP_272560006.1 | WP_272562398.1 |
| Bacillota | *Anoxybacillus calidus* | WP_181537656.1 | WP_181535810.1 | WP_181535808.1 | WP_181536607.1 |
| Bacillota | *Bacillus alveayuensis* | WP_044894876.1 | WP_044748079.1 | WP_044748078.1 | WP_419151291.1 |
| Bacillota | *Bacillus cereus group* | WP_065212475.1 | WP_097817830.1 | WP_070172932.1 | WP_002165144.1 |
| Bacillota | *Bacillus piscicola* | WP_240377695.1 | WP_240375564.1 | WP_338021128.1 | WP_240375160.1 |
| Bacillota | *Bacillus solitudinis* | WP_100407558.1 | WP_100407236.1 | WP_100408361.1 | WP_100404574.1 |
| Bacillota | *Bacillus sp. FJAT-45037* | WP_100372031.1 | WP_100372728.1 | WP_100372727.1 | WP_157796518.1 |
| Bacillota | *Baia soyae* | WP_243649462.1 | WP_131848545.1 | WP_131848544.1 | WP_131847547.1 |
| Bacillota | *Brevibacillus agri* | WP_290426100.1 | WP_327944617.1 | WP_005831237.1 | WP_005833010.1 |
| Bacillota | *Brevibacillus antibioticus* | WP_137029627.1 | WP_137029308.1 | WP_137029309.1 | WP_137028103.1 |
| Bacillota | *Brevibacillus borstelensis* | WP_340257345.1 | WP_171506301.1 | WP_340250559.1 | WP_024983988.1 |
| Bacillota | *Brevibacillus brevis* | WP_310772139.1 | WP_144616023.1 | WP_310772836.1 | WP_106653219.1 |
| Bacillota | *Brevibacillus centrosporus* | WP_092267967.1 | WP_411504249.1 | WP_327930207.1 | WP_328211152.1 |
| Bacillota | *Brevibacillus choshinensis* | WP_327995948.1 | WP_203356916.1 | WP_327976965.1 | WP_055746698.1 |
| Bacillota | *Brevibacillus composti* | WP_198826645.1 | WP_198826360.1 | WP_198826361.1 | WP_198829395.1 |
| Bacillota | *Brevibacillus daliensis* | WP_232699265.1 | WP_232696377.1 | WP_232696376.1 | WP_232698373.1 |
| Bacillota | *Brevibacillus dissolubilis* | WP_139489532.1 | WP_139489136.1 | WP_139489137.1 | WP_139489967.1 |
| Bacillota | *Brevibacillus fluminis* | WP_122919557.1 | WP_122920747.1 | WP_122920746.1 | WP_122916979.1 |
| Bacillota | *Brevibacillus formosus* | WP_047068682.1 | WP_047071010.1 | WP_047069015.1 | WP_047068378.1 |
| Bacillota | *Brevibacillus fortis* | WP_409178351.1 | WP_106841588.1 | WP_327950005.1 | WP_106840056.1 |
| Bacillota | *Brevibacillus gelatini* | WP_122903973.1 | WP_122904259.1 | WP_122904258.1 | WP_122906096.1 |
| Bacillota | *Brevibacillus humidisoli* | WP_230046769.1 | WP_230046195.1 | WP_230046959.1 | WP_230045536.1 |
| Bacillota | *Brevibacillus invocatus* | WP_122911184.1 | WP_122908283.1 | WP_122908779.1 | WP_251243155.1 |
| Bacillota | *Brevibacillus laterosporus* | WP_121473740.1 | WP_031414565.1 | WP_003341662.1 | WP_104031805.1 |
| Bacillota | *Brevibacillus marinus* | WP_126427700.1 | WP_126427242.1 | WP_126427244.1 | WP_126425988.1 |
| Bacillota | *Brevibacillus massiliensis* | WP_019119801.1 | WP_019122419.1 | WP_019122420.1 | WP_019120713.1 |
| Bacillota | *Brevibacillus migulae* | WP_134682950.1 | WP_134684718.1 | WP_134686655.1 | WP_134687458.1 |
| Bacillota | *Brevibacillus nitrificans* | WP_310234934.1 | WP_310227961.1 | WP_310227962.1 | WP_328207014.1 |

| | | | | | |
|---|---|---|---|---|---|
| Bacillota | *Brevibacillus panacihumi* | WP_023554712.1 | WP_122912326.1 | WP_427329339.1 | WP_427329734.1 |
| Bacillota | *Brevibacillus parabrevis* | WP_063227760.1 | WP_063226980.1 | WP_063227869.1 | WP_063230376.1 |
| Bacillota | *Brevibacillus porteri* | WP_376897442.1 | WP_106833023.1 | WP_106835974.1 | WP_106835688.1 |
| Bacillota | *Brevibacillus reuszeri* | WP_103108679.1 | WP_049740083.1 | WP_049740084.1 | WP_213022732.1 |
| Bacillota | *Brevibacillus ruminantium* | WP_251870872.1 | WP_251870562.1 | WP_251870563.1 | WP_251874646.1 |
| Bacillota | *Brevibacillus sp. SKDU10* | WP_064018533.1 | WP_064018445.1 | WP_064018446.1 | WP_064018249.1 |
| Bacillota | *Brevibacillus thermoruber* | WP_029099275.1 | WP_029097619.1 | WP_029099570.1 | WP_271140081.1 |
| Bacillota | *Caldalkalibacillus mannanilyticus* | WP_025026615.1 | WP_025026795.1 | WP_025026794.1 | WP_025028057.1 |
| Bacillota | *Caldalkalibacillus salinus* | WP_202078626.1 | WP_272899181.1 | WP_202079975.1 | WP_202076947.1 |
| Bacillota | *Caldalkalibacillus thermarum* | WP_188622173.1 | WP_188623426.1 | WP_188623462.1 | WP_222822622.1 |
| Bacillota | *Caldalkalibacillus uzonensis* | WP_307340273.1 | WP_307334880.1 | WP_307334882.1 | WP_307338827.1 |
| Bacillota | *Calditerricola satsumensis* | WP_188816489.1 | WP_054672757.1 | WP_054672754.1 | WP_188816587.1 |
| Bacillota | *Chengkuizengella axinellae* | WP_305989955.1 | WP_305991132.1 | WP_305991131.1 | WP_305990434.1 |
| Bacillota | *Chengkuizengella marina* | WP_160645612.1 | WP_160647308.1 | WP_160647307.1 | WP_160646215.1 |
| Bacillota | *Chengkuizengella sediminis* | WP_162038533.1 | WP_162035760.1 | WP_162035759.1 | WP_162035053.1 |
| Bacillota | *Chengkuizengella sp. SCS-71B* | WP_349250319.1 | WP_349248815.1 | WP_349248814.1 | WP_349249855.1 |
| Bacillota | *Cohnella abietis* | WP_130606950.1 | WP_130608606.1 | WP_157994104.1 | WP_157994083.1 |
| Bacillota | *Cohnella boryungensis* | WP_204601817.1 | WP_204603271.1 | WP_204603272.1 | WP_204601225.1 |
| Bacillota | *Cohnella faecalis* | WP_119148594.1 | WP_119147875.1 | WP_119147874.1 | WP_158593973.1 |
| Bacillota | *Cohnella herbarum* | WP_169278050.1 | WP_169278750.1 | WP_169278749.1 | WP_169279893.1 |
| Bacillota | *Cohnella hongkongensis* | WP_378102549.1 | WP_378096723.1 | WP_378096104.1 | WP_378097115.1 |
| Bacillota | *Cohnella luojiensis* | WP_135152141.1 | WP_135150944.1 | WP_135150943.1 | WP_167746931.1 |
| Bacillota | *Cohnella lupini* | WP_115992159.1 | WP_115993120.1 | WP_115993121.1 | WP_115990748.1 |
| Bacillota | *Cohnella mopanensis* | WP_239615160.1 | WP_239614212.1 | WP_239614211.1 | WP_239617380.1 |
| Bacillota | *Cohnella silvisoli* | WP_232187555.1 | WP_232183992.1 | WP_232186837.1 | WP_232187431.1 |
| Bacillota | *Cohnella sp.* | WP_287021570.1 | WP_372634402.1 | WP_287022233.1 | WP_287021775.1 |
| Bacillota | *Croceifilum oryzae* | WP_307251400.1 | WP_307254358.1 | WP_307254361.1 | WP_307250766.1 |
| Bacillota | *Desertibacillus haloalkaliphilus* | WP_217221480.1 | WP_217224796.1 | WP_217224795.1 | WP_217224580.1 |

| | | | | | |
|---|---|---|---|---|---|
| Bacillota | *Desmospora activa* | WP_107724566.1 | WP_107725504.1 | WP_245891082.1 | WP_107726050.1 |
| Bacillota | *Desmospora profundinema* | WP_309865860.1 | WP_309861822.1 | WP_309861824.1 | WP_309862629.1 |
| Bacillota | *Fontibacillus panacisegetis* | WP_091228678.1 | WP_091229626.1 | WP_091229628.1 | WP_091226566.1 |
| Bacillota | *Fontibacillus phaseoli* | WP_114497056.1 | WP_114495195.1 | WP_114495194.1 | WP_114494769.1 |
| Bacillota | *Fontibacillus solani* | WP_182534579.1 | WP_182535212.1 | WP_182535213.1 | WP_182535122.1 |
| Bacillota | *Fontibacillus sp. BL9* | WP_410771892.1 | WP_410768391.1 | WP_410768390.1 | WP_410768037.1 |
| Bacillota | *Gordoniibacillus kamchatkensis* | WP_052487363.1 | WP_156157817.1 | WP_041046861.1 | WP_041046255.1 |
| Bacillota | *Gorillibacterium sp. CAU 1737* | WP_347771175.1 | WP_347768118.1 | WP_347768119.1 | WP_347772139.1 |
| Bacillota | *Gorillibacterium timonense* | WP_058302272.1 | WP_058303859.1 | WP_058303858.1 | WP_082651929.1 |
| Bacillota | *Halalkalibacter alkalisediminis* | WP_273839419.1 | WP_273840757.1 | WP_273840758.1 | WP_273839757.1 |
| Bacillota | *Halalkalibacter flavus* | WP_332633351.1 | WP_332630148.1 | WP_332630147.1 | WP_332632203.1 |
| Bacillota | *Halalkalibacter kiskunsagensis* | WP_335963127.1 | WP_335958984.1 | WP_335958985.1 | WP_335960984.1 |
| Bacillota | *Halalkalibacter lacteus* | WP_332696835.1 | WP_332693267.1 | WP_332693270.1 | WP_332690015.1 |
| Bacillota | *Hazenella coriacea* | WP_131923295.1 | WP_131925631.1 | WP_131925633.1 | WP_131924031.1 |
| Bacillota | *Kroppenstedtia eburnea* | WP_380132810.1 | WP_076523778.1 | WP_040387154.1 | WP_380150828.1 |
| Bacillota | *Kroppenstedtia guangzhouensis* | WP_188431726.1 | WP_188430415.1 | WP_188430417.1 | WP_188428771.1 |
| Bacillota | *Kroppenstedtia pulmonis* | WP_173220744.1 | WP_173224572.1 | WP_173221991.1 | WP_173222804.1 |
| Bacillota | *Kroppenstedtia sanguinis* | WP_380162730.1 | WP_380163115.1 | WP_380163114.1 | WP_380164071.1 |
| Bacillota | *Laceyella putida* | WP_379862863.1 | WP_379863246.1 | WP_379865748.1 | WP_379866126.1 |
| Bacillota | *Laceyella sacchari* | WP_132219594.1 | WP_259436017.1 | WP_259436617.1 | WP_022736468.1 |
| Bacillota | *Laceyella tengchongensis* | WP_284723808.1 | WP_284724464.1 | WP_333639873.1 | WP_102992870.1 |
| Bacillota | *Lihuaxuella thermophila* | WP_089971102.1 | WP_244527535.1 | WP_089969006.1 | WP_089966921.1 |
| Bacillota | *Longirhabdus pacifica* | WP_128893698.1 | WP_128894412.1 | WP_128894413.1 | WP_128896012.1 |
| Bacillota | *Marinicrinis sediminis* | WP_379927926.1 | WP_379930489.1 | WP_379930488.1 | WP_379929885.1 |
| Bacillota | *Marininema halotolerans* | WP_091836498.1 | WP_245838860.1 | WP_091838700.1 | WP_176391884.1 |
| Bacillota | *Marininema mesophilum* | WP_091742882.1 | WP_091737720.1 | WP_091737717.1 | WP_091734794.1 |
| Bacillota | *Marinithermofilum abyssi* | WP_188646376.1 | WP_188646919.1 | WP_188646918.1 | WP_188648308.1 |
| Bacillota | *Marinococcus halophilus* | WP_079476624.1 | WP_079476570.1 | WP_079476571.1 | WP_079476090.1 |

| | | | | | |
|---|---|---|---|---|---|
| Bacillota | *Marinococcus halotolerans* | WP_022792304.1 | WP_022792248.1 | WP_022792249.1 | WP_022792687.1 |
| Bacillota | *Marinococcus luteus* | WP_322525176.1 | WP_322525124.1 | WP_322525125.1 | WP_091610196.1 |
| Bacillota | *Mechercharimyces sp. CAU 1602* | WP_258839993.1 | WP_258839241.1 | WP_258839240.1 | WP_258838868.1 |
| Bacillota | *Melghirimyces algeriensis* | WP_142506200.1 | WP_142504252.1 | WP_246064816.1 | WP_142504508.1 |
| Bacillota | *Melghirimyces profundicolus* | WP_108023950.1 | WP_108023223.1 | WP_108023221.1 | WP_108026146.1 |
| Bacillota | *Melghirimyces thermohalophilus* | WP_091568263.1 | WP_091569222.1 | WP_091569219.1 | WP_091565521.1 |
| Bacillota | *Metabacillus herbersteinensis* | WP_378939256.1 | WP_378932858.1 | WP_378933101.1 | WP_378931977.1 |
| Bacillota | *Microaerobacter geothermalis* | WP_236404968.1 | WP_236405557.1 | WP_407942200.1 | WP_236405793.1 |
| Bacillota | *Novibacillus thermophilus* | WP_077719752.1 | WP_077720594.1 | WP_077718877.1 | WP_077719370.1 |
| Bacillota | *Paenactinomyces guangxiensis* | WP_181750949.1 | WP_181751724.1 | WP_309506121.1 | WP_181751500.1 |
| Bacillota | *Paenibacillus abyssi* | WP_188533017.1 | WP_188528309.1 | WP_188528307.1 | WP_188531779.1 |
| Bacillota | *Paenibacillus aceris* | WP_167069014.1 | WP_167053849.1 | WP_167053847.1 | WP_167063709.1 |
| Bacillota | *Paenibacillus aceti* | WP_120465243.1 | WP_120462256.1 | WP_120462257.1 | WP_120462650.1 |
| Bacillota | *Paenibacillus agilis* | WP_186786190.1 | WP_144988852.1 | WP_144988849.1 | WP_144991373.1 |
| Bacillota | *Paenibacillus agri* | WP_175370962.1 | WP_175373667.1 | WP_175373669.1 | WP_175372874.1 |
| Bacillota | *Paenibacillus agricola* | WP_166148862.1 | WP_166145632.1 | WP_166145635.1 | WP_311764869.1 |
| Bacillota | *Paenibacillus alba* | WP_173223509.1 | WP_326071302.1 | WP_326071303.1 | WP_326073808.1 |
| Bacillota | *Paenibacillus albicereus* | WP_168908529.1 | WP_206110059.1 | WP_168907475.1 | WP_168908071.1 |
| Bacillota | *Paenibacillus albidus* | WP_189021651.1 | WP_215176829.1 | WP_189024419.1 | WP_215175915.1 |
| Bacillota | *Paenibacillus albus* | WP_126015480.1 | WP_126016142.1 | WP_126016140.1 | WP_126015742.1 |
| Bacillota | *Paenibacillus alginolyticus* | WP_029194617.1 | WP_268615698.1 | WP_029193045.1 | WP_268615491.1 |
| Bacillota | *Paenibacillus algorifonticola* | WP_046229419.1 | WP_046232076.1 | WP_046232075.1 | WP_046230403.1 |
| Bacillota | *Paenibacillus alkaliterrae* | WP_235234645.1 | WP_235234721.1 | WP_235235115.1 | WP_235238666.1 |
| Bacillota | *Paenibacillus alkalitolerans* | WP_199618403.1 | WP_199614925.1 | WP_199614927.1 | WP_199618942.1 |
| Bacillota | *Paenibacillus allorhizoplanae* | WP_236290221.1 | WP_236284611.1 | WP_236285355.1 | WP_236287531.1 |
| Bacillota | *Paenibacillus allorhizosphaerae* | WP_218100378.1 | WP_218100601.1 | WP_218100602.1 | WP_218097063.1 |
| Bacillota | *Paenibacillus alvei* | WP_171419366.1 | WP_138186444.1 | WP_410453185.1 | WP_268632461.1 |
| Bacillota | *Paenibacillus amylolyticus* | WP_353883419.1 | WP_123066039.1 | WP_133385331.1 | WP_347386579.1 |

| | | | | | |
|---|---|---|---|---|---|
| Bacillota | *Paenibacillus andongensis* | WP_261304790.1 | WP_261306455.1 | WP_261301090.1 | WP_261305761.1 |
| Bacillota | *Paenibacillus antibioticophila* | WP_101781156.1 | WP_212939118.1 | WP_212939119.1 | WP_044479289.1 |
| Bacillota | *Paenibacillus apiarius* | WP_206097645.1 | WP_206098075.1 | WP_206098076.1 | WP_206096386.1 |
| Bacillota | *Paenibacillus apii* | WP_165099327.1 | WP_165094357.1 | WP_165094360.1 | WP_165094978.1 |
| Bacillota | *Paenibacillus apis* | WP_301628741.1 | WP_301624040.1 | WP_301627953.1 | WP_301624390.1 |
| Bacillota | *Paenibacillus arenilitoris* | WP_190863467.1 | WP_190861985.1 | WP_190861983.1 | WP_190862815.1 |
| Bacillota | *Paenibacillus arenosi* | WP_192026591.1 | WP_192024005.1 | WP_192024006.1 | WP_192023195.1 |
| Bacillota | *Paenibacillus assamensis* | WP_028594151.1 | WP_028593560.1 | WP_028593559.1 | WP_028595832.1 |
| Bacillota | *Paenibacillus athensensis* | WP_134756811.1 | WP_134750921.1 | WP_134750231.1 | WP_134757304.1 |
| Bacillota | *Paenibacillus auburnensis* | WP_236335297.1 | WP_236331603.1 | WP_236331605.1 | WP_236332662.1 |
| Bacillota | *Paenibacillus aurantiacus* | WP_377492888.1 | WP_377500029.1 | WP_377500027.1 | WP_377491245.1 |
| Bacillota | *Paenibacillus aurantius* | WP_315606699.1 | WP_315603029.1 | WP_315607009.1 | WP_315607290.1 |
| Bacillota | *Paenibacillus baekrokdamisoli* | WP_232016416.1 | WP_125656665.1 | WP_164522785.1 | WP_125657417.1 |
| Bacillota | *Paenibacillus barcinonensis* | WP_216519699.1 | WP_110894264.1 | WP_110894265.1 | WP_110898516.1 |
| Bacillota | *Paenibacillus barengoltzii* | WP_085279631.1 | WP_016313974.1 | WP_016311834.1 | WP_085169625.1 |
| Bacillota | *Paenibacillus beijingensis* | WP_045671434.1 | WP_045672563.1 | WP_045672564.1 | WP_045671783.1 |
| Bacillota | *Paenibacillus borealis* | WP_042217159.1 | WP_042215472.1 | WP_042215475.1 | WP_042216727.1 |
| Bacillota | *Paenibacillus bouchesdurhonensis* | WP_110933759.1 | WP_110932522.1 | WP_110932521.1 | WP_110932164.1 |
| Bacillota | *Paenibacillus brevis* | WP_216479681.1 | WP_216481004.1 | WP_216481005.1 | WP_216481043.1 |
| Bacillota | *Paenibacillus camerounensis* | WP_042203043.1 | WP_042196114.1 | WP_042204296.1 | WP_042197491.1 |
| Bacillota | *Paenibacillus castaneae* | WP_208414390.1 | WP_102713920.1 | WP_102710846.1 | WP_102712600.1 |
| Bacillota | *Paenibacillus caui* | WP_223066141.1 | WP_223066654.1 | WP_223066691.1 | WP_223066358.1 |
| Bacillota | *Paenibacillus cellulosilyticus* | WP_174812767.1 | WP_110043359.1 | WP_110043360.1 | WP_110046471.1 |
| Bacillota | *Paenibacillus cellulositrophicus* | WP_152398643.1 | WP_251676995.1 | WP_251676997.1 | WP_152398424.1 |
| Bacillota | *Paenibacillus chibensis* | WP_127606199.1 | WP_328277584.1 | WP_127607166.1 | WP_127601684.1 |
| Bacillota | *Paenibacillus chitinolyticus* | WP_387540779.1 | WP_218229832.1 | WP_377648432.1 | WP_042232440.1 |
| Bacillota | *Paenibacillus chungangensis* | WP_377568795.1 | WP_377562651.1 | WP_377562179.1 | WP_377568441.1 |
| Bacillota | *Paenibacillus cineris* | WP_212971249.1 | WP_212984345.1 | WP_212967858.1 | WP_212984444.1 |

| | | | | | |
|---|---|---|---|---|---|
| Bacillota | *Paenibacillus contaminans* | WP_113030359.1 | WP_113033634.1 | WP_113032777.1 | WP_113032225.1 |
| Bacillota | *Paenibacillus cremeus* | WP_144842443.1 | WP_144842940.1 | WP_261381216.1 | WP_144847196.1 |
| Bacillota | *Paenibacillus curdlanolyticus* | WP_006039766.1 | WP_006039229.1 | WP_006039230.1 | WP_006036336.1 |
| Bacillota | *Paenibacillus cymbidii* | WP_135553327.1 | WP_135551466.1 | WP_135547907.1 | WP_135546844.1 |
| Bacillota | *Paenibacillus daejeonensis* | WP_020615915.1 | WP_020616645.1 | WP_020616646.1 | WP_020615593.1 |
| Bacillota | *Paenibacillus dakarensis* | WP_054956193.1 | WP_054957615.1 | WP_054957616.1 | WP_054956622.1 |
| Bacillota | *Paenibacillus dendritiformis* | WP_133382336.1 | WP_168181508.1 | WP_133381197.1 | WP_213469088.1 |
| Bacillota | *Paenibacillus dendrobii* | WP_160500621.1 | WP_202128686.1 | WP_160498316.1 | WP_160498813.1 |
| Bacillota | *Paenibacillus dokdonensis* | WP_326089625.1 | WP_136606159.1 | WP_136606160.1 | WP_326090758.1 |
| Bacillota | *Paenibacillus donghaensis* | WP_087917836.1 | WP_087914496.1 | WP_087916894.1 | WP_087917553.1 |
| Bacillota | *Paenibacillus durus* | WP_042207934.1 | WP_042205360.1 | WP_042207283.1 | WP_042207704.1 |
| Bacillota | *Paenibacillus ehimensis* | WP_326059397.1 | WP_127485603.1 | WP_025846315.1 | WP_326057552.1 |
| Bacillota | *Paenibacillus elgii* | WP_339371850.1 | WP_108531728.1 | WP_339370912.1 | WP_127459765.1 |
| Bacillota | *Paenibacillus endophyticus* | WP_183557533.1 | WP_183561091.1 | WP_183561093.1 | WP_183563669.1 |
| Bacillota | *Paenibacillus etheri* | WP_060621774.1 | WP_060622731.1 | WP_060622730.1 | WP_060622030.1 |
| Bacillota | *Paenibacillus eucommiae* | WP_209975024.1 | WP_209972764.1 | WP_209973263.1 | WP_209974125.1 |
| Bacillota | *Paenibacillus faecalis* | WP_106768550.1 | WP_106767939.1 | WP_106767940.1 | WP_106768318.1 |
| Bacillota | *Paenibacillus faecis* | WP_213508219.1 | WP_213504414.1 | WP_213504415.1 | WP_148451589.1 |
| Bacillota | *Paenibacillus farraposensis* | WP_229523210.1 | WP_229525501.1 | WP_229525500.1 | WP_229524566.1 |
| Bacillota | *Paenibacillus favisporus* | WP_163881487.1 | WP_163879295.1 | WP_354494891.1 | WP_354494692.1 |
| Bacillota | *Paenibacillus ferrarius* | WP_079419186.1 | WP_079420736.1 | WP_079420735.1 | WP_079420014.1 |
| Bacillota | *Paenibacillus filicis* | WP_341418433.1 | WP_341414414.1 | WP_341414415.1 | WP_341416827.1 |
| Bacillota | *Paenibacillus flagellatus* | WP_110842254.1 | WP_110839882.1 | WP_110839883.1 | WP_110840646.1 |
| Bacillota | *Paenibacillus foliorum* | WP_171654195.1 | WP_171651155.1 | WP_171655783.1 | WP_312886573.1 |
| Bacillota | *Paenibacillus fonticola* | WP_019638513.1 | WP_019640396.1 | WP_019640395.1 | WP_019636244.1 |
| Bacillota | *Paenibacillus forsythiae* | WP_025697225.1 | WP_025699187.1 | WP_025699479.1 | WP_025697264.1 |
| Bacillota | *Paenibacillus frigoriresistens* | WP_173186128.1 | WP_173184821.1 | WP_173184605.1 | WP_173186448.1 |
| Bacillota | *Paenibacillus ginsengihumi* | WP_019532785.1 | WP_019534029.1 | WP_019534030.1 | WP_019533053.1 |

| | | | | | |
|---|---|---|---|---|---|
| Bacillota | *Paenibacillus graminis* | WP_326082249.1 | WP_025707470.1 | WP_025707469.1 | WP_025708368.1 |
| Bacillota | *Paenibacillus guangzhouensis* | WP_152391647.1 | WP_152393925.1 | WP_152393926.1 | WP_265333443.1 |
| Bacillota | *Paenibacillus gyeongsangnamensis* | WP_269881790.1 | WP_269881438.1 | WP_269880074.1 | WP_269879583.1 |
| Bacillota | *Paenibacillus haidiansis* | WP_331848184.1 | WP_331844562.1 | WP_331844563.1 | WP_331844893.1 |
| Bacillota | *Paenibacillus hamazuiensis* | WP_248927459.1 | WP_248925463.1 | WP_248925462.1 | WP_248926615.1 |
| Bacillota | *Paenibacillus harenae* | WP_029192613.1 | WP_307201541.1 | WP_028610564.1 | WP_028611235.1 |
| Bacillota | *Paenibacillus hemerocallicola* | WP_139601970.1 | WP_139604983.1 | WP_139604984.1 | WP_139602082.1 |
| Bacillota | *Paenibacillus herberti* | WP_342745633.1 | WP_176444973.1 | WP_089523697.1 | WP_176444650.1 |
| Bacillota | *Paenibacillus hexagrammi* | WP_235118391.1 | WP_235118659.1 | WP_235122911.1 | WP_235121698.1 |
| Bacillota | *Paenibacillus ihbetae* | WP_099478905.1 | WP_077565455.1 | WP_099476792.1 | WP_099476993.1 |
| Bacillota | *Paenibacillus ihuae* | WP_054942784.1 | WP_054941331.1 | WP_054941332.1 | WP_054941758.1 |
| Bacillota | *Paenibacillus ihumii* | WP_055109149.1 | WP_055107795.1 | WP_055110100.1 | WP_055107462.1 |
| Bacillota | *Paenibacillus illinoisensis* | WP_221807689.1 | WP_336780754.1 | WP_402875603.1 | WP_337032811.1 |
| Bacillota | *Paenibacillus jilunlii* | WP_062523374.1 | WP_062519683.1 | WP_062519685.1 | WP_062527005.1 |
| Bacillota | *Paenibacillus kobensis* | WP_127529797.1 | WP_127530721.1 | WP_127530722.1 | WP_127533050.1 |
| Bacillota | *Paenibacillus koleovorans* | WP_127578709.1 | WP_127588578.1 | WP_127588579.1 | WP_127582757.1 |
| Bacillota | *Paenibacillus kribbensis* | WP_094154033.1 | WP_068499549.1 | WP_326109516.1 | WP_094154485.1 |
| Bacillota | *Paenibacillus lacisoli* | WP_305024135.1 | WP_305022224.1 | WP_305022225.1 | WP_305022581.1 |
| Bacillota | *Paenibacillus larvae* | WP_077995778.1 | WP_268573972.1 | WP_023482678.1 | WP_079940202.1 |
| Bacillota | *Paenibacillus lautus* | WP_220685281.1 | WP_127594050.1 | WP_127594126.1 | WP_127592034.1 |
| Bacillota | *Paenibacillus lemnae* | WP_169504005.1 | WP_169506788.1 | WP_169506789.1 | WP_169503480.1 |
| Bacillota | *Paenibacillus lentus* | WP_125084497.1 | WP_379392406.1 | WP_379392405.1 | WP_125082923.1 |
| Bacillota | *Paenibacillus lignilyticus* | WP_210656124.1 | WP_210659859.1 | WP_210659857.1 | WP_210659384.1 |
| Bacillota | *Paenibacillus luteus* | WP_141504106.1 | WP_141504020.1 | WP_141504734.1 | WP_141505818.1 |
| Bacillota | *Paenibacillus lutimineralis* | WP_127002345.1 | WP_126997891.1 | WP_126997889.1 | WP_126997225.1 |
| Bacillota | *Paenibacillus lutrae* | WP_235918181.1 | WP_157335440.1 | WP_157335441.1 | WP_157334343.1 |
| Bacillota | *Paenibacillus lycopersici* | WP_162357517.1 | WP_162356891.1 | WP_162356889.1 | WP_162356475.1 |
| Bacillota | *Paenibacillus macerans* | WP_278738955.1 | WP_251586340.1 | WP_326045291.1 | WP_328270886.1 |

| | | | | | |
|---|---|---|---|---|---|
| Bacillota | *Paenibacillus marchantiophytorum* | WP_189020885.1 | WP_189009728.1 | WP_189014345.1 | WP_189006243.1 |
| Bacillota | *Paenibacillus massiliensis* | WP_025676521.1 | WP_028589105.1 | WP_025677800.1 | WP_028588777.1 |
| Bacillota | *Paenibacillus maysiensis* | WP_025685211.1 | WP_025685875.1 | WP_025685876.1 | WP_025685742.1 |
| Bacillota | *Paenibacillus mellifer* | WP_248552883.1 | WP_248550581.1 | WP_248550580.1 | WP_248551931.1 |
| Bacillota | *Paenibacillus mendelii* | WP_204817996.1 | WP_204817258.1 | WP_204818322.1 | WP_204816513.1 |
| Bacillota | *Paenibacillus mesophilus* | WP_138877610.1 | WP_138878350.1 | WP_138878085.1 | WP_138882220.1 |
| Bacillota | *Paenibacillus mesotrionivorans* | WP_416215470.1 | WP_416214544.1 | WP_416214545.1 | WP_416219599.1 |
| Bacillota | *Paenibacillus methanolicus* | WP_148931812.1 | WP_148929581.1 | WP_148928225.1 | WP_148928442.1 |
| Bacillota | *Paenibacillus montanisoli* | WP_206098550.1 | WP_112882578.1 | WP_112882579.1 | WP_112882883.1 |
| Bacillota | *Paenibacillus montaniterrae* | WP_213516663.1 | WP_213513802.1 | WP_213513264.1 | WP_246563468.1 |
| Bacillota | *Paenibacillus monticola* | WP_154121649.1 | WP_154118393.1 | WP_154118394.1 | WP_154121344.1 |
| Bacillota | *Paenibacillus motobuensis* | WP_343862506.1 | WP_343863633.1 | WP_343863631.1 | WP_343865702.1 |
| Bacillota | *Paenibacillus mucilaginosus* | WP_013920168.1 | WP_013916540.1 | WP_013916539.1 | WP_014652281.1 |
| Bacillota | *Paenibacillus naphthalenovorans* | WP_074727122.1 | WP_175472024.1 | WP_074729390.1 | WP_062410860.1 |
| Bacillota | *Paenibacillus nasutitermitis* | WP_188993163.1 | WP_188992611.1 | WP_188992612.1 | WP_188992857.1 |
| Bacillota | *Paenibacillus oceani* | WP_190924490.1 | WP_190925107.1 | WP_190925109.1 | WP_190926136.1 |
| Bacillota | *Paenibacillus odorifer* | WP_094902091.1 | WP_312151840.1 | WP_248548496.1 | WP_076304292.1 |
| Bacillota | *Paenibacillus oenotherae* | WP_219874260.1 | WP_219870643.1 | WP_219870644.1 | WP_219870874.1 |
| Bacillota | *Paenibacillus oleatilyticus* | WP_216795578.1 | WP_216792181.1 | WP_216792182.1 | WP_216797443.1 |
| Bacillota | *Paenibacillus oralis* | WP_128634953.1 | WP_128634089.1 | WP_128634090.1 | WP_128634460.1 |
| Bacillota | *Paenibacillus oryzae* | WP_068679112.1 | WP_068683840.1 | WP_068683842.1 | WP_068684506.1 |
| Bacillota | *Paenibacillus oryzisoli* | WP_068665391.1 | WP_068670283.1 | WP_068670284.1 | WP_068670715.1 |
| Bacillota | *Paenibacillus ottowii* | WP_142613378.1 | WP_064796416.1 | WP_326400406.1 | WP_305164649.1 |
| Bacillota | *Paenibacillus pabuli* | WP_062327955.1 | WP_426251135.1 | WP_347384342.1 | WP_247898527.1 |
| Bacillota | *Paenibacillus paeoniae* | WP_116042437.1 | WP_116044696.1 | WP_116044698.1 | WP_240644312.1 |
| Bacillota | *Paenibacillus paridis* | WP_139991027.1 | WP_139999520.1 | WP_238404104.1 | WP_139997909.1 |
| Bacillota | *Paenibacillus pasadenensis* | WP_251559661.1 | WP_251560931.1 | WP_251560930.1 | WP_101809358.1 |
| Bacillota | *Paenibacillus pectinilyticus* | WP_065851700.1 | WP_065857228.1 | WP_065857226.1 | WP_244163060.1 |

| | | | | | |
|---|---|---|---|---|---|
| Bacillota | *Paenibacillus pedocola* | WP_310829968.1 | WP_310829302.1 | WP_310829303.1 | WP_310829725.1 |
| Bacillota | *Paenibacillus peoriae* | WP_338532767.1 | WP_010345798.1 | WP_076293050.1 | WP_104497052.1 |
| Bacillota | *Paenibacillus periandrae* | WP_240414419.1 | WP_240413846.1 | WP_240413847.1 | WP_240417094.1 |
| Bacillota | *Paenibacillus phocaensis* | WP_068783155.1 | WP_068785288.1 | WP_068785287.1 | WP_068784959.1 |
| Bacillota | *Paenibacillus phyllosphaerae* | WP_183596122.1 | WP_183596763.1 | WP_183598251.1 | WP_246427471.1 |
| Bacillota | *Paenibacillus physcomitrellae* | WP_094096216.1 | WP_094093303.1 | WP_094093304.1 | WP_094093583.1 |
| Bacillota | *Paenibacillus phytohabitans* | WP_336102449.1 | WP_171716664.1 | WP_171716665.1 | WP_171721012.1 |
| Bacillota | *Paenibacillus pinihumi* | WP_028561893.1 | WP_028561258.1 | WP_028561259.1 | WP_028561628.1 |
| Bacillota | *Paenibacillus pinisoli* | WP_120107911.1 | WP_120113277.1 | WP_424452460.1 | WP_243643891.1 |
| Bacillota | *Paenibacillus piri* | WP_133228799.1 | WP_133225913.1 | WP_133225912.1 | WP_342774573.1 |
| Bacillota | *Paenibacillus piscarius* | WP_238652198.1 | WP_238652982.1 | WP_238652983.1 | WP_238654633.1 |
| Bacillota | *Paenibacillus plantarum* | WP_171630428.1 | WP_171628794.1 | WP_171629110.1 | WP_171635300.1 |
| Bacillota | *Paenibacillus planticolens* | WP_171686840.1 | WP_171682055.1 | WP_171682056.1 | WP_246294449.1 |
| Bacillota | *Paenibacillus plantiphilus* | WP_236346401.1 | WP_236344839.1 | WP_371877750.1 | WP_236344089.1 |
| Bacillota | *Paenibacillus polymyxa* | WP_330706336.1 | WP_290402692.1 | WP_194843217.1 | WP_013311101.1 |
| Bacillota | *Paenibacillus polysaccharolyticus* | WP_251508739.1 | WP_251505656.1 | WP_251505659.1 | WP_253492605.1 |
| Bacillota | *Paenibacillus popilliae* | WP_142545952.1 | WP_006285465.1 | WP_142543324.1 | WP_006286443.1 |
| Bacillota | *Paenibacillus prosopidis* | WP_114382400.1 | WP_114382545.1 | WP_114378576.1 | WP_114381166.1 |
| Bacillota | *Paenibacillus pseudetheri* | WP_234537788.1 | WP_234531948.1 | WP_234531942.1 | WP_234533213.1 |
| Bacillota | *Paenibacillus psychroresistens* | WP_155702999.1 | WP_155701927.1 | WP_155704866.1 | WP_155700801.1 |
| Bacillota | *Paenibacillus puerhi* | WP_159881110.1 | WP_159883059.1 | WP_159883057.1 | WP_235941410.1 |
| Bacillota | *Paenibacillus puldeungensis* | WP_379321167.1 | WP_379321196.1 | WP_379321195.1 | WP_379318034.1 |
| Bacillota | *Paenibacillus qinlingensis* | WP_173115243.1 | WP_173110236.1 | WP_173110235.1 | WP_310226997.1 |
| Bacillota | *Paenibacillus radicibacter* | WP_258208379.1 | WP_258206429.1 | WP_258206430.1 | WP_258206078.1 |
| Bacillota | *Paenibacillus radicis (ex Xue et al. 2023)* | WP_258211272.1 | WP_258216056.1 | WP_258216055.1 | WP_258216523.1 |
| Bacillota | *Paenibacillus rhizophilus* | WP_124695119.1 | WP_124693619.1 | WP_124693618.1 | WP_124695680.1 |
| Bacillota | *Paenibacillus rhizoplanae* | WP_209993516.1 | WP_379256890.1 | WP_209993887.1 | WP_209987956.1 |
| Bacillota | *Paenibacillus rhizosphaerae* | WP_183587077.1 | WP_183581975.1 | WP_183581974.1 | WP_183581722.1 |

| | | | | | |
|---|---|---|---|---|---|
| Bacillota | *Paenibacillus rigui* | WP_094014449.1 | WP_094013285.1 | WP_094013286.1 | WP_094015209.1 |
| Bacillota | *Paenibacillus riograndensis* | WP_020430154.1 | WP_060861783.1 | WP_060861782.1 | WP_020427852.1 |
| Bacillota | *Paenibacillus sabinae* | WP_025336118.1 | WP_025335550.1 | WP_025335551.1 | WP_025335875.1 |
| Bacillota | *Paenibacillus sabuli* | WP_223869387.1 | WP_190916452.1 | WP_190916454.1 | WP_190915075.1 |
| Bacillota | *Paenibacillus sambharensis* | WP_181438790.1 | WP_111149231.1 | WP_111149230.1 | WP_111148711.1 |
| Bacillota | *Paenibacillus sanguinis* | WP_018752292.1 | WP_018751371.1 | WP_018751372.1 | WP_018753264.1 |
| Bacillota | *Paenibacillus sedimenti* | WP_188175441.1 | WP_188177267.1 | WP_188174343.1 | WP_188172947.1 |
| Bacillota | *Paenibacillus selenitireducens* | WP_078497635.1 | WP_078497211.1 | WP_078497210.1 | WP_078498745.1 |
| Bacillota | *Paenibacillus senegalensis* | WP_010271006.1 | WP_026021375.1 | WP_010273795.1 | WP_010270149.1 |
| Bacillota | *Paenibacillus senegalimassiliensis* | WP_059049591.1 | WP_059052566.1 | WP_059052564.1 | WP_059052170.1 |
| Bacillota | *Paenibacillus septentrionalis* | WP_379231073.1 | WP_379233282.1 | WP_379232422.1 | WP_379230547.1 |
| Bacillota | *Paenibacillus sepulcri* | WP_210046909.1 | WP_210041358.1 | WP_210045727.1 | WP_210037664.1 |
| Bacillota | *Paenibacillus silagei* | WP_209876089.1 | WP_209874571.1 | WP_209869954.1 | WP_209871997.1 |
| Bacillota | *Paenibacillus silvae* | WP_248063624.1 | WP_111271384.1 | WP_111271385.1 | WP_111271909.1 |
| Bacillota | *Paenibacillus silvestris* | WP_235960584.1 | WP_161408218.1 | WP_161408564.1 | WP_235959055.1 |
| Bacillota | *Paenibacillus silvisoli* | WP_308638710.1 | WP_308639168.1 | WP_308639167.1 | WP_308638903.1 |
| Bacillota | *Paenibacillus silviterrae* | WP_284644809.1 | WP_284643983.1 | WP_284640174.1 | WP_284638745.1 |
| Bacillota | *Paenibacillus sinopodophylli* | WP_342778467.1 | WP_138750981.1 | WP_138750980.1 | WP_138751928.1 |
| Bacillota | *Paenibacillus solanacearum* | WP_218092977.1 | WP_218091182.1 | WP_218091183.1 | WP_218090647.1 |
| Bacillota | *Paenibacillus solisilvae* | WP_379190145.1 | WP_379189360.1 | WP_379191972.1 | WP_379190327.1 |
| Bacillota | *Paenibacillus sonchi* | WP_039838104.1 | WP_039835063.1 | WP_039835064.1 | WP_233184314.1 |
| Bacillota | *Paenibacillus sophorae* | WP_036598261.1 | WP_036594357.1 | WP_036594356.1 | WP_036596753.1 |
| Bacillota | *Paenibacillus sp.* | WP_309122791.1 | WP_309118627.1 | WP_309118628.1 | WP_421885775.1 |
| Bacillota | *Paenibacillus spongiae* | WP_258387971.1 | WP_258387583.1 | WP_258388477.1 | WP_258388242.1 |
| Bacillota | *Paenibacillus stellifer* | WP_038698104.1 | WP_038697001.1 | WP_038697002.1 | WP_038697658.1 |
| Bacillota | *Paenibacillus swuensis* | WP_068607342.1 | WP_068605796.1 | WP_068604930.1 | WP_231891263.1 |
| Bacillota | *Paenibacillus taichungensis* | WP_113055901.1 | WP_175382996.1 | WP_413372315.1 | WP_113053843.1 |
| Bacillota | *Paenibacillus taihuensis* | WP_116189947.1 | WP_181909822.1 | WP_116187052.1 | WP_116187460.1 |

| | | | | | |
|---|---|---|---|---|---|
| Bacillota | *Paenibacillus taiwanensis* | WP_028547052.1 | WP_028544285.1 | WP_028544284.1 | WP_245596021.1 |
| Bacillota | *Paenibacillus tarimensis* | WP_235287205.1 | WP_235285858.1 | WP_235285857.1 | WP_235285404.1 |
| Bacillota | *Paenibacillus tengchongensis* | WP_151736597.1 | WP_151734500.1 | WP_151734499.1 | WP_151735851.1 |
| Bacillota | *Paenibacillus tepidiphilus* | WP_150269544.1 | WP_150273581.1 | WP_150273580.1 | WP_150266292.1 |
| Bacillota | *Paenibacillus terrae* | WP_145163154.1 | WP_014282065.1 | WP_014282066.1 | WP_149093315.1 |
| Bacillota | *Paenibacillus terricola* | WP_191201982.1 | WP_191203123.1 | WP_191203122.1 | WP_191202242.1 |
| Bacillota | *Paenibacillus terrigena* | WP_314587033.1 | WP_018755349.1 | WP_314585441.1 | WP_018756691.1 |
| Bacillota | *Paenibacillus thalictri* | WP_170318199.1 | WP_131012956.1 | WP_131012957.1 | WP_131014125.1 |
| Bacillota | *Paenibacillus thermoaerophilus* | WP_138788696.1 | WP_138787912.1 | WP_138787913.1 | WP_170209381.1 |
| Bacillota | *Paenibacillus thiaminolyticus* | WP_119796028.1 | WP_272045015.1 | WP_119792949.1 | WP_087442146.1 |
| Bacillota | *Paenibacillus tianjinensis* | WP_206101756.1 | WP_206100817.1 | WP_206100818.1 | WP_206101483.1 |
| Bacillota | *Paenibacillus tianmuensis* | WP_090669484.1 | WP_090665888.1 | WP_090665886.1 | WP_090670061.1 |
| Bacillota | *Paenibacillus timonensis* | WP_240271378.1 | WP_240268406.1 | WP_240268405.1 | WP_240269054.1 |
| Bacillota | *Paenibacillus tritici* | WP_173132030.1 | WP_173129013.1 | WP_211720201.1 | WP_173128240.1 |
| Bacillota | *Paenibacillus tundrae* | WP_307218044.1 | WP_307212880.1 | WP_307212877.1 | WP_338540183.1 |
| Bacillota | *Paenibacillus turicensis* | WP_210088697.1 | WP_210091285.1 | WP_210091286.1 | WP_210088043.1 |
| Bacillota | *Paenibacillus turpanensis* | WP_166240415.1 | WP_166239436.1 | WP_166239434.1 | WP_166243811.1 |
| Bacillota | *Paenibacillus typhae* | WP_090714882.1 | WP_090714162.1 | WP_221798010.1 | WP_090712285.1 |
| Bacillota | *Paenibacillus tyrfis* | WP_281937182.1 | WP_036680591.1 | WP_253409447.1 | WP_253412858.1 |
| Bacillota | *Paenibacillus uliginis* | WP_208915349.1 | WP_208914519.1 | WP_208914520.1 | WP_208915055.1 |
| Bacillota | *Paenibacillus vandeheii* | WP_301248911.1 | WP_301246512.1 | WP_301246510.1 | WP_301243803.1 |
| Bacillota | *Paenibacillus vietnamensis* | WP_224723386.1 | WP_224723467.1 | WP_224722472.1 | WP_224721890.1 |
| Bacillota | *Paenibacillus whitsoniae* | WP_126145122.1 | WP_126140510.1 | WP_126140511.1 | WP_126143135.1 |
| Bacillota | *Paenibacillus woosongensis* | WP_283927468.1 | WP_155610100.1 | WP_330163346.1 | WP_283928465.1 |
| Bacillota | *Paenibacillus wynnii* | WP_307589484.1 | WP_036657105.1 | WP_307586656.1 | WP_036657888.1 |
| Bacillota | *Paenibacillus xanthanilyticus* | WP_377721381.1 | WP_377722267.1 | WP_377722268.1 | WP_377722477.1 |
| Bacillota | *Paenibacillus xerothermodurans* | WP_089200934.1 | WP_089199061.1 | WP_089199062.1 | WP_243633136.1 |
| Bacillota | *Paenibacillus xylanexedens* | WP_315370738.1 | WP_145047451.1 | WP_145408863.1 | WP_124116014.1 |

| | | | | | |
|---|---|---|---|---|---|
| Bacillota | *Paenibacillus xylaniclasticus* | WP_127572600.1 | WP_127566875.1 | WP_127566877.1 | WP_127567498.1 |
| Bacillota | *Paenibacillus xylanilyticus* | WP_175394430.1 | WP_413035370.1 | WP_175397877.1 | WP_175397529.1 |
| Bacillota | *Paenibacillus xylanivorans* | WP_053783136.1 | WP_053780711.1 | WP_053780710.1 | WP_053781229.1 |
| Bacillota | *Paenibacillus yonginensis* | WP_068697326.1 | WP_068695883.1 | WP_068695885.1 | WP_068696453.1 |
| Bacillota | *Paenibacillus zanthoxyli* | WP_025693164.1 | WP_025692677.1 | WP_025692678.1 | WP_025691633.1 |
| Bacillota | *Paludifilum halophilum* | WP_169713717.1 | WP_094264071.1 | WP_094262790.1 | WP_240511973.1 |
| Bacillota | *Peribacillus butanolivorans* | WP_340455904.1 | WP_379155062.1 | WP_379155065.1 | WP_387587241.1 |
| Bacillota | *Peribacillus castrilensis* | WP_318291301.1 | WP_367409646.1 | WP_367407233.1 | WP_367407051.1 |
| Bacillota | *Peribacillus frigoritolerans* | WP_367235604.1 | WP_349728053.1 | WP_286166049.1 | WP_342615343.1 |
| Bacillota | *Peribacillus simplex* | WP_137020667.1 | WP_061144265.1 | WP_061144266.1 | WP_061144059.1 |
| Bacillota | *Planifilum fimeticola* | WP_106345446.1 | WP_106345631.1 | WP_106345581.1 | WP_106344368.1 |
| Bacillota | *Planifilum fulgidum* | WP_177199077.1 | WP_092035968.1 | WP_092035967.1 | WP_092041673.1 |
| Bacillota | *Planococcus antarcticus* | WP_006830683.1 | WP_040852325.1 | WP_006829302.1 | WP_006828840.1 |
| Bacillota | *Polycladomyces abyssicola* | WP_212774188.1 | WP_212772256.1 | WP_212772255.1 | WP_212772617.1 |
| Bacillota | *Polycladomyces subterraneus* | WP_301240606.1 | WP_301237983.1 | WP_301237984.1 | WP_301239880.1 |
| Bacillota | *Polycladospora coralii* | WP_191139217.1 | WP_347239741.1 | WP_191138248.1 | WP_191140021.1 |
| Bacillota | *Saccharibacillus alkalitolerans* | WP_166276593.1 | WP_166272347.1 | WP_166272345.1 | WP_166271645.1 |
| Bacillota | *Saccharibacillus deserti* | WP_172254369.1 | WP_172250470.1 | WP_172250468.1 | WP_172249784.1 |
| Bacillota | *Saccharibacillus endophyticus* | WP_172244410.1 | WP_172239876.1 | WP_172239879.1 | WP_172247082.1 |
| Bacillota | *Saccharibacillus kuerlensis* | WP_018978303.1 | WP_018976427.1 | WP_018976426.1 | WP_018975684.1 |
| Bacillota | *Saccharibacillus qingshengii* | WP_172196605.1 | WP_172196256.1 | WP_172196255.1 | WP_172194137.1 |
| Bacillota | *Saccharibacillus sp. O23* | WP_088488869.1 | WP_088489527.1 | WP_088489528.1 | WP_088489347.1 |
| Bacillota | *Salibacterium aidingense* | WP_026700176.1 | WP_026700929.1 | WP_422787443.1 | WP_422785923.1 |
| Bacillota | *Salibacterium halotolerans* | WP_093335510.1 | WP_093335998.1 | WP_093335997.1 | WP_212634930.1 |
| Bacillota | *Salibacterium lacus* | WP_380711669.1 | WP_380711740.1 | WP_380711739.1 | WP_380712172.1 |
| Bacillota | *Salibacterium qingdaonense* | WP_177195506.1 | WP_090925891.1 | WP_090925890.1 | WP_177195403.1 |
| Bacillota | *Salibacterium salarium* | WP_125557516.1 | WP_125555844.1 | WP_348639443.1 | WP_306897393.1 |
| Bacillota | *Salinithrix halophila* | WP_380702635.1 | WP_380705887.1 | WP_380705888.1 | WP_380703580.1 |

| | | | | | |
|---|---|---|---|---|---|
| Bacillota | *Salipaludibacillus sp. CUR1* | WP_230896231.1 | WP_230896188.1 | WP_230896189.1 | WP_230898551.1 |
| Bacillota | *Salsuginibacillus halophilus* | WP_106587779.1 | WP_106587852.1 | WP_106587851.1 | WP_106587505.1 |
| Bacillota | *Salsuginibacillus kocurii* | WP_018922183.1 | WP_018922278.1 | WP_018922277.1 | WP_018921352.1 |
| Bacillota | *Sinobaca qinghaiensis* | WP_120193293.1 | WP_120193227.1 | WP_120193228.1 | WP_120192335.1 |
| Bacillota | *Sutcliffiella horikoshii* | WP_148990256.1 | WP_226683573.1 | WP_404428349.1 | WP_187441928.1 |
| Bacillota | *Thermoactinomyces sp. DSM 45891* | WP_072336567.1 | WP_072329871.1 | WP_072329873.1 | WP_072331764.1 |
| Bacillota | *Thermoflavimicrobium daqui* | WP_113657606.1 | WP_113660335.1 | WP_113660334.1 | WP_113658619.1 |
| Bacillota | *Thermoflavimicrobium dichotomicum* | WP_093228129.1 | WP_093230852.1 | WP_093230851.1 | WP_175482318.1 |
| Bacillota | *Virgibacillus tibetensis* | WP_327609447.1 | WP_327606045.1 | WP_327608938.1 | WP_327606340.1 |
| Bacillota | *Xylanibacillus composti* | WP_244865042.1 | WP_213410179.1 | WP_213410178.1 | WP_213410594.1 |
| Bacteroidota | *Acidiluteibacter ferrifornacis* | WP_160633441.1 | WP_160634009.1 | WP_160633238.1 | WP_160632436.1 |
| Bacteroidota | *Adhaeribacter arboris* | WP_106931386.1 | WP_106925362.1 | WP_106929784.1 | WP_106933576.1 |
| Bacteroidota | *Adhaeribacter pallidiroseus* | WP_115374736.1 | WP_115373291.1 | WP_115373435.1 | WP_115375762.1 |
| Bacteroidota | *Adhaeribacter radiodurans* | WP_246343606.1 | WP_182414787.1 | WP_182412160.1 | WP_182414468.1 |
| Bacteroidota | *Adhaeribacter soli* | WP_150903137.1 | WP_150904862.1 | WP_150905911.1 | WP_150903363.1 |
| Bacteroidota | *Adhaeribacter swui* | WP_185273551.1 | WP_185270322.1 | WP_185274196.1 | WP_185272839.1 |
| Bacteroidota | *Adhaeribacter terreus* | WP_378017384.1 | WP_378015753.1 | WP_378016465.1 | WP_378017653.1 |
| Bacteroidota | *Adhaeribacter terrigena* | WP_200506402.1 | WP_200504364.1 | WP_200505621.1 | WP_200506036.1 |
| Bacteroidota | *Aequorivita antarctica* | WP_111843961.1 | WP_111843834.1 | WP_111845043.1 | WP_111844811.1 |
| Bacteroidota | *Aequorivita aquimaris* | WP_062623013.1 | WP_062621430.1 | WP_062619548.1 | WP_062619660.1 |
| Bacteroidota | *Aequorivita aurantiaca* | WP_290253495.1 | WP_290253605.1 | WP_290255165.1 | WP_290255086.1 |
| Bacteroidota | *Aequorivita capsosiphonis* | WP_026451993.1 | WP_026449899.1 | WP_026451927.1 | WP_026450721.1 |
| Bacteroidota | *Aequorivita ciconiae* | WP_128249725.1 | WP_128249060.1 | WP_128249027.1 | WP_128251637.1 |
| Bacteroidota | *Aequorivita lipolytica* | WP_111816085.1 | WP_111815969.1 | WP_111816248.1 | WP_111816891.1 |
| Bacteroidota | *Aequorivita marina* | WP_310993630.1 | WP_310991922.1 | WP_310992157.1 | WP_310993816.1 |
| Bacteroidota | *Aequorivita nionensis* | WP_410005368.1 | WP_410005228.1 | WP_410004785.1 | WP_410007104.1 |
| Bacteroidota | *Aequorivita soesokkakensis* | WP_068761931.1 | WP_068760878.1 | WP_068761363.1 | WP_068760990.1 |
| Bacteroidota | *Aequorivita sp. KMM 9714* | WP_165150480.1 | WP_165147784.1 | WP_165149664.1 | WP_165150353.1 |

| | | | | | |
|---|---|---|---|---|---|
| Bacteroidota | *Aequorivita sublithincola* | WP_042492077.1 | WP_014783441.1 | WP_014781031.1 | WP_014780829.1 |
| Bacteroidota | *Aequorivita todarodis* | WP_272854795.1 | WP_272854907.1 | WP_272856410.1 | WP_272856446.1 |
| Bacteroidota | *Aequorivita viscosa* | WP_073214506.1 | WP_073219450.1 | WP_073221462.1 | WP_073214181.1 |
| Bacteroidota | *Aequorivita vitellina* | WP_237602696.1 | WP_237602884.1 | WP_237604082.1 | WP_237604065.1 |
| Bacteroidota | *Aequorivita vladivostokensis* | WP_045081596.1 | WP_045080904.1 | WP_045080108.1 | WP_045081657.1 |
| Bacteroidota | *Aequorivita xiaoshiensis* | WP_237609006.1 | WP_237607055.1 | WP_237608260.1 | WP_237608850.1 |
| Bacteroidota | *Aestuariibaculum lutulentum* | WP_240573145.1 | WP_240572916.1 | WP_240574573.1 | WP_240573340.1 |
| Bacteroidota | *Aestuariibaculum marinum* | WP_188222212.1 | WP_188223804.1 | WP_188221776.1 | WP_188222367.1 |
| Bacteroidota | *Aestuariibaculum sediminum* | WP_188229559.1 | WP_188229943.1 | WP_188229495.1 | WP_188230458.1 |
| Bacteroidota | *Aestuariibaculum sp. M13* | WP_258228506.1 | WP_258227523.1 | WP_258227337.1 | WP_258228345.1 |
| Bacteroidota | *Aestuariibaculum suncheonense* | WP_188216222.1 | WP_188215830.1 | WP_188215657.1 | WP_188216389.1 |
| Bacteroidota | *Aestuariivivens insulae* | WP_242205529.1 | WP_242204282.1 | WP_242206092.1 | WP_242205612.1 |
| Bacteroidota | *Aestuariivivens marinum* | WP_242132968.1 | WP_242132290.1 | WP_242135394.1 | WP_242135503.1 |
| Bacteroidota | *Aestuariivivens sediminicola* | WP_242093504.1 | WP_242120336.1 | WP_242121371.1 | WP_242093034.1 |
| Bacteroidota | *Aestuariivivens sediminis* | WP_242084579.1 | WP_242086932.1 | WP_242158599.1 | WP_242157877.1 |
| Bacteroidota | *Aestuariivivens sp. NBU2969* | WP_223548038.1 | WP_223548296.1 | WP_223551508.1 | WP_223552062.1 |
| Bacteroidota | *Agaribacillus aureus* | WP_346761610.1 | WP_346757862.1 | WP_346755833.1 | WP_346761836.1 |
| Bacteroidota | *Albibacterium bauzanense* | WP_132221933.1 | WP_132222972.1 | WP_132221473.1 | WP_132222344.1 |
| Bacteroidota | *Albibacterium sp.* | WP_325360523.1 | WP_325357452.1 | WP_325358942.1 | WP_325359824.1 |
| Bacteroidota | *Algibacter agarivorans* | WP_345193075.1 | WP_345189876.1 | WP_345193043.1 | WP_345193815.1 |
| Bacteroidota | *Algibacter amylolyticus* | WP_144117106.1 | WP_144118081.1 | WP_144117049.1 | WP_144116980.1 |
| Bacteroidota | *Algibacter aquimarinus* | WP_345169948.1 | WP_345169810.1 | WP_345170338.1 | WP_345169783.1 |
| Bacteroidota | *Algibacter lectus* | WP_074940410.1 | WP_042494898.1 | WP_042497935.1 | WP_074936115.1 |
| Bacteroidota | *Algibacter luteus* | WP_019387202.1 | WP_289194008.1 | WP_019387259.1 | WP_289196303.1 |
| Bacteroidota | *Algibacter marinivivus* | WP_109353024.1 | WP_109353276.1 | WP_109353074.1 | WP_109353815.1 |
| Bacteroidota | *Algibacter mikhailovii* | WP_189361399.1 | WP_189358466.1 | WP_189362427.1 | WP_282124090.1 |
| Bacteroidota | *Algibacter miyuki* | WP_290273055.1 | WP_290273454.1 | WP_290271960.1 | WP_290273464.1 |
| Bacteroidota | *Algibacter pacificus* | WP_147677447.1 | WP_147678726.1 | WP_147678533.1 | WP_147678795.1 |

| | | | | | |
|---|---|---|---|---|---|
| Bacteroidota | *Algibacter pectinivorans* | WP_092852296.1 | WP_092853154.1 | WP_092852392.1 | WP_092851984.1 |
| Bacteroidota | *Algibacter sp.* | WP_300249043.1 | WP_300252120.1 | WP_348451855.1 | WP_348451771.1 |
| Bacteroidota | *Algivirga pacifica* | WP_345370542.1 | WP_345373764.1 | WP_345373213.1 | WP_345371319.1 |
| Bacteroidota | *Algoriella sp.* | WP_332546600.1 | WP_332544110.1 | WP_332546255.1 | WP_287443154.1 |
| Bacteroidota | *Allotamlana fucoidanivorans* | WP_139698527.1 | WP_139694982.1 | WP_139698433.1 | WP_139696462.1 |
| Bacteroidota | *Altibacter sp.* | WP_290625980.1 | WP_288003297.1 | WP_290628138.1 | WP_290627392.1 |
| Bacteroidota | *Anditalea andensis* | WP_035075125.1 | WP_035075984.1 | WP_035072741.1 | WP_035075153.1 |
| Bacteroidota | *Apibacter adventoris* | WP_105192749.1 | WP_105193923.1 | WP_181042806.1 | WP_105193663.1 |
| Bacteroidota | *Apibacter mensalis* | WP_303812563.1 | WP_303821584.1 | WP_055425757.1 | WP_303812704.1 |
| Bacteroidota | *Apibacter muscae* | WP_146310081.1 | WP_146261981.1 | WP_146261740.1 | WP_146311829.1 |
| Bacteroidota | *Apibacter sp.* | WP_293918652.1 | WP_293917054.1 | WP_293918312.1 | WP_293918330.1 |
| Bacteroidota | *Aquaticitalea lipolytica* | WP_283635384.1 | WP_188605702.1 | WP_283635546.1 | WP_188606921.1 |
| Bacteroidota | *Aquimarina acroporae* | WP_248333197.1 | WP_248325971.1 | WP_248327184.1 | WP_248329473.1 |
| Bacteroidota | *Aquimarina addita* | WP_344929695.1 | WP_344928121.1 | WP_344926945.1 | WP_344927438.1 |
| Bacteroidota | *Aquimarina agarilytica* | WP_010182218.1 | WP_010182235.1 | WP_010178350.1 | WP_010182594.1 |
| Bacteroidota | *Aquimarina agarivorans* | WP_010520948.1 | WP_010521832.1 | WP_010522891.1 | WP_010520711.1 |
| Bacteroidota | *Aquimarina algicola* | WP_140588301.1 | WP_140589169.1 | WP_140595569.1 | WP_140595083.1 |
| Bacteroidota | *Aquimarina amphilecti* | WP_091403738.1 | WP_091404647.1 | WP_091403964.1 | WP_091410641.1 |
| Bacteroidota | *Aquimarina aquimarini* | WP_234859132.1 | WP_108869272.1 | WP_108868242.1 | WP_108868066.1 |
| Bacteroidota | *Aquimarina atlantica* | WP_051575609.1 | WP_034238049.1 | WP_034241498.1 | WP_034246824.1 |
| Bacteroidota | *Aquimarina brevivitae* | WP_130287912.1 | WP_130286860.1 | WP_130287960.1 | WP_130286092.1 |
| Bacteroidota | *Aquimarina celericrescens* | WP_378321902.1 | WP_378318285.1 | WP_378318615.1 | WP_378321627.1 |
| Bacteroidota | *Aquimarina gracilis* | WP_324181944.1 | WP_324180621.1 | WP_324180101.1 | WP_324178525.1 |
| Bacteroidota | *Aquimarina intermedia* | WP_148783869.1 | WP_148782085.1 | WP_148782431.1 | WP_148783651.1 |
| Bacteroidota | *Aquimarina litoralis* | WP_219009173.1 | WP_343912639.1 | WP_299600296.1 | WP_343913483.1 |
| Bacteroidota | *Aquimarina longa* | WP_062055130.1 | WP_062057664.1 | WP_062058925.1 | WP_062059664.1 |
| Bacteroidota | *Aquimarina macrocephali* | WP_024772579.1 | WP_024770031.1 | WP_024771131.1 | WP_024771276.1 |
| Bacteroidota | *Aquimarina muelleri* | WP_027413955.1 | WP_027411125.1 | WP_027411711.1 | WP_027411300.1 |

| | | | | | |
|---|---|---|---|---|---|
| Bacteroidota | *Aquimarina mytili* | WP_201924405.1 | WP_201917632.1 | WP_201924142.1 | WP_201922323.1 |
| Bacteroidota | *Aquimarina pacifica* | WP_025740165.1 | WP_025742060.1 | WP_025743084.1 | WP_025742517.1 |
| Bacteroidota | *Aquimarina rubra* | WP_378288611.1 | WP_378290376.1 | WP_378288455.1 | WP_378295432.1 |
| Bacteroidota | *Aquimarina sediminis* | WP_103067938.1 | WP_103068351.1 | WP_103070573.1 | WP_103070728.1 |
| Bacteroidota | *Aquimarina sp. 2-A2* | WP_407140247.1 | WP_407139263.1 | WP_407140005.1 | WP_407139823.1 |
| Bacteroidota | *Aquimarina spongiae* | WP_073314334.1 | WP_073314409.1 | WP_073314355.1 | WP_073320496.1 |
| Bacteroidota | *Aquirufa antheringensis* | WP_130895386.1 | WP_130922775.1 | WP_269001602.1 | WP_269001114.1 |
| Bacteroidota | *Aquirufa aurantiipilula* | WP_223144145.1 | WP_223142915.1 | WP_276343803.1 | WP_223143335.1 |
| Bacteroidota | *Aquirufa avitistagni* | WP_377983087.1 | WP_377983723.1 | WP_377982913.1 | WP_377983456.1 |
| Bacteroidota | *Aquirufa beregesia* | WP_166229642.1 | WP_166230096.1 | WP_166228982.1 | WP_166229514.1 |
| Bacteroidota | *Aquirufa ecclesiirivi* | WP_166373820.1 | WP_166374907.1 | WP_166375689.1 | WP_422184414.1 |
| Bacteroidota | *Aquirufa echingensis* | WP_377974724.1 | WP_377976744.1 | WP_377976516.1 | WP_377974603.1 |
| Bacteroidota | *Aquirufa lenticrescens* | WP_223131557.1 | WP_223131088.1 | WP_223130902.1 | WP_223129962.1 |
| Bacteroidota | *Aquirufa nivalisilvae* | WP_109324317.1 | WP_109322091.1 | WP_269024951.1 | WP_130924003.1 |
| Bacteroidota | *Aquirufa novilacunae* | WP_406778192.1 | WP_406800710.1 | WP_406800059.1 | WP_406800572.1 |
| Bacteroidota | *Aquirufa originis* | WP_377978463.1 | WP_377979066.1 | WP_377978324.1 | WP_377978916.1 |
| Bacteroidota | *Aquirufa regiilacus* | WP_315574943.1 | WP_315576891.1 | WP_315575113.1 | WP_316070539.1 |
| Bacteroidota | *Aquirufa rosea* | WP_129027395.1 | WP_129026517.1 | WP_129027631.1 | WP_129027468.1 |
| Bacteroidota | *Aquirufa salirivi* | WP_406750953.1 | WP_406749496.1 | WP_406751867.1 | WP_406750838.1 |
| Bacteroidota | *Aquirufa sp.* | WP_395767647.1 | WP_395617283.1 | WP_395767809.1 | WP_395616701.1 |
| Bacteroidota | *Arcticibacter eurypsychrophilus* | WP_069659817.1 | WP_069660197.1 | WP_069658220.1 | WP_069658143.1 |
| Bacteroidota | *Arcticibacter pallidicorallinus* | WP_106290857.1 | WP_106295235.1 | WP_106293321.1 | WP_106293700.1 |
| Bacteroidota | *Arcticibacter sp.* | WP_407430282.1 | WP_407426349.1 | WP_407428255.1 | WP_407430250.1 |
| Bacteroidota | *Arcticibacter svalbardensis* | WP_016196807.1 | WP_016197349.1 | WP_016193496.1 | WP_016195818.1 |
| Bacteroidota | *Arcticibacter tournemirensis* | WP_128768377.1 | WP_128770802.1 | WP_128769017.1 | WP_128771266.1 |
| Bacteroidota | *Aridibaculum aurantiacum* | WP_207493959.1 | WP_207496171.1 | WP_207493540.1 | WP_207493862.1 |
| Bacteroidota | *Ascidiimonas aurantiaca* | WP_340067019.1 | WP_340063238.1 | WP_340065951.1 | WP_340063664.1 |
| Bacteroidota | *Ascidiimonas meishanensis* | WP_340202765.1 | WP_340202346.1 | WP_340201815.1 | WP_340203336.1 |

| | | | | | |
|---|---|---|---|---|---|
| Bacteroidota | *Asinibacterium sp. OR53* | WP_026770568.1 | WP_026768116.1 | WP_026770043.1 | WP_026770422.1 |
| Bacteroidota | *Asprobacillus argus* | WP_349241159.1 | WP_349240976.1 | WP_349241222.1 | WP_349242390.1 |
| Bacteroidota | *Aurantibacillus circumpalustris* | WP_317898783.1 | WP_317897266.1 | WP_317899147.1 | WP_317898617.1 |
| Bacteroidota | *Aurantibacter aestuarii* | WP_181256232.1 | WP_106463421.1 | WP_106463824.1 | WP_106463759.1 |
| Bacteroidota | *Aurantibacter sp.* | WP_375238534.1 | WP_375239262.1 | WP_348149237.1 | WP_375239007.1 |
| Bacteroidota | *Aureibacter tunicatorum* | WP_309938190.1 | WP_309937555.1 | WP_309941955.1 | WP_309937614.1 |
| Bacteroidota | *Aureibaculum algae* | WP_138949641.1 | WP_138950357.1 | WP_138950222.1 | WP_138948862.1 |
| Bacteroidota | *Aureibaculum flavum* | WP_198841374.1 | WP_198839813.1 | WP_198839693.1 | WP_198842913.1 |
| Bacteroidota | *Aureibaculum luteum* | WP_117880577.1 | WP_117883329.1 | WP_117883048.1 | WP_117885144.1 |
| Bacteroidota | *Aureibaculum marinum* | WP_123899280.1 | WP_123896962.1 | WP_123897081.1 | WP_123896527.1 |
| Bacteroidota | *Aureibaculum sp. 2308TA14-22* | WP_354204459.1 | WP_354205576.1 | WP_354205365.1 | WP_354206002.1 |
| Bacteroidota | *Aureisphaera galaxeae* | WP_272862643.1 | WP_272862417.1 | WP_272863078.1 | WP_272861625.1 |
| Bacteroidota | *Aureispira anguillae* | WP_264788052.1 | WP_264788426.1 | WP_264791552.1 | WP_264791511.1 |
| Bacteroidota | *Aureitalea sp. L0-47* | WP_265200652.1 | WP_265198881.1 | WP_265199191.1 | WP_265200177.1 |
| Bacteroidota | *Aureivirga marina* | WP_196892920.1 | WP_196892692.1 | WP_196894274.1 | WP_196894243.1 |
| Bacteroidota | *Aureivirga sp. CE67* | WP_196886436.1 | WP_196888611.1 | WP_196888714.1 | WP_196887798.1 |
| Bacteroidota | *Bergeyella cardium* | WP_120489054.1 | WP_120489413.1 | WP_120488367.1 | WP_120488308.1 |
| Bacteroidota | *Bergeyella porcorum* | WP_376834211.1 | WP_327985048.1 | WP_327984028.1 | WP_412677604.1 |
| Bacteroidota | *Bergeyella sp. RCAD1439* | WP_327609825.1 | WP_327609899.1 | WP_327610773.1 | WP_327610132.1 |
| Bacteroidota | *Bergeyella zoohelcum* | WP_002664476.1 | WP_394265388.1 | WP_002661560.1 | WP_002662668.1 |
| Bacteroidota | *Bernardetia litoralis* | WP_014798351.1 | WP_014799449.1 | WP_014796014.1 | WP_014797464.1 |
| Bacteroidota | *Bernardetia sp.* | WP_291723737.1 | WP_291727868.1 | WP_291727328.1 | WP_291721233.1 |
| Bacteroidota | *Bizionia arctica* | WP_188464709.1 | WP_188463855.1 | WP_188463365.1 | WP_188465515.1 |
| Bacteroidota | *Bizionia argentinensis* | WP_008636277.1 | WP_008634538.1 | WP_008636439.1 | WP_008637444.1 |
| Bacteroidota | *Bizionia echini* | WP_092208682.1 | WP_092207375.1 | WP_092206879.1 | WP_092209708.1 |
| Bacteroidota | *Bizionia hallyeonensis* | WP_376860896.1 | WP_376859852.1 | WP_376859305.1 | WP_376860208.1 |
| Bacteroidota | *Bizionia myxarmorum* | WP_148403551.1 | WP_148404171.1 | WP_148405154.1 | WP_148403799.1 |
| Bacteroidota | *Bizionia paragorgiae* | WP_092132826.1 | WP_318908187.1 | WP_092135828.1 | WP_318907152.1 |

| | | | | | |
|---|---|---|---|---|---|
| Bacteroidota | *Bizionia sediminis* | WP_376894251.1 | WP_376891938.1 | WP_376891554.1 | WP_376891096.1 |
| Bacteroidota | *Bizionia sp.* | WP_417196395.1 | WP_417195014.1 | WP_417199749.1 | WP_417238594.1 |
| Bacteroidota | *Botryobacter ruber* | WP_114782892.1 | WP_114781647.1 | WP_114781543.1 | WP_240676079.1 |
| Bacteroidota | *Brumimicrobium aurantiacum* | WP_116880677.1 | WP_116880684.1 | WP_116880515.1 | WP_116881141.1 |
| Bacteroidota | *Brumimicrobium glaciale* | WP_130094390.1 | WP_130094022.1 | WP_242494536.1 | WP_165366249.1 |
| Bacteroidota | *Brumimicrobium mesophilum* | WP_107039253.1 | WP_107040258.1 | WP_107039225.1 | WP_107039918.1 |
| Bacteroidota | *Brumimicrobium oceani* | WP_109360571.1 | WP_109358790.1 | WP_233244208.1 | WP_109360517.1 |
| Bacteroidota | *Brumimicrobium salinarum* | WP_101334249.1 | WP_101335045.1 | WP_101334281.1 | WP_101335294.1 |
| Bacteroidota | *Brumimicrobium sp.* | WP_417266258.1 | WP_417265319.1 | WP_417266280.1 | WP_417265785.1 |
| Bacteroidota | *Candidatus Kaistella beijingensis* | WP_224136972.1 | WP_224135010.1 | WP_224134386.1 | WP_224134929.1 |
| Bacteroidota | *Candidatus Ulvibacter alkanivorans* | WP_114490672.1 | WP_114491340.1 | WP_114491714.1 | WP_114491961.1 |
| Bacteroidota | *Cesiribacter sp. SM1* | WP_370634655.1 | WP_224999480.1 | WP_224998922.1 | WP_224999441.1 |
| Bacteroidota | *Changchengzhania lutea* | WP_142785652.1 | WP_142785956.1 | WP_142783246.1 | WP_142783659.1 |
| Bacteroidota | *Chishuiella changwenlii* | WP_072930933.1 | WP_072931684.1 | WP_072930379.1 | WP_143147299.1 |
| Bacteroidota | *Chishuiella sp.* | WP_313376097.1 | WP_313385780.1 | WP_313386052.1 | WP_313374642.1 |
| Bacteroidota | *Chitinophaga agri* | WP_238429994.1 | WP_162333742.1 | WP_162331142.1 | WP_238430125.1 |
| Bacteroidota | *Chitinophaga agrisoli* | WP_149839302.1 | WP_149836969.1 | WP_149837932.1 | WP_149840340.1 |
| Bacteroidota | *Chitinophaga alhagiae* | WP_119078929.1 | WP_119077216.1 | WP_119079661.1 | WP_157986662.1 |
| Bacteroidota | *Chitinophaga arvensicola* | WP_089902623.1 | WP_089889289.1 | WP_089896020.1 | WP_177192308.1 |
| Bacteroidota | *Chitinophaga barathri* | WP_120516762.1 | WP_120518200.1 | WP_120514463.1 | WP_162946241.1 |
| Bacteroidota | *Chitinophaga caeni* | WP_098193960.1 | WP_098192515.1 | WP_098194845.1 | WP_157760992.1 |
| Bacteroidota | *Chitinophaga caseinilytica* | WP_423737646.1 | WP_341842633.1 | WP_341839141.1 | WP_423734991.1 |
| Bacteroidota | *Chitinophaga costaii* | WP_089708494.1 | WP_089715512.1 | WP_089710726.1 | WP_089715615.1 |
| Bacteroidota | *Chitinophaga cymbidii* | WP_146862438.1 | WP_146858207.1 | WP_146865249.1 | WP_186831128.1 |
| Bacteroidota | *Chitinophaga defluvii* | WP_354663379.1 | WP_354661261.1 | WP_354660547.1 | WP_354659270.1 |
| Bacteroidota | *Chitinophaga deserti* | WP_109698414.1 | WP_109694755.1 | WP_109699103.1 | WP_157962811.1 |
| Bacteroidota | *Chitinophaga dinghuensis* | WP_245950740.1 | WP_111591532.1 | WP_111592684.1 | WP_111595453.1 |
| Bacteroidota | *Chitinophaga eiseniae* | WP_168741694.1 | WP_078670110.1 | WP_078670884.1 | WP_168741852.1 |

| | | | | | |
|---|---|---|---|---|---|
| Bacteroidota | *Chitinophaga filiformis* | WP_247813539.1 | WP_089834396.1 | WP_089836010.1 | WP_089830986.1 |
| Bacteroidota | *Chitinophaga flava* | WP_113617360.1 | WP_113616485.1 | WP_113614148.1 | WP_113618438.1 |
| Bacteroidota | *Chitinophaga fulva* | WP_169228654.1 | WP_169222926.1 | WP_169225324.1 | WP_169227571.1 |
| Bacteroidota | *Chitinophaga ginsengisoli* | WP_245901671.1 | WP_106600515.1 | WP_106605529.1 | WP_106600880.1 |
| Bacteroidota | *Chitinophaga horti* | WP_264281860.1 | WP_244842465.1 | WP_264282605.1 | WP_264282541.1 |
| Bacteroidota | *Chitinophaga hostae* | WP_211971874.1 | WP_211973759.1 | WP_211974175.1 | WP_211973914.1 |
| Bacteroidota | *Chitinophaga japonensis* | WP_145713553.1 | WP_145711536.1 | WP_145715336.1 | WP_145717070.1 |
| Bacteroidota | *Chitinophaga jiangningensis* | WP_073083571.1 | WP_073084114.1 | WP_073086198.1 | WP_073087053.1 |
| Bacteroidota | *Chitinophaga lutea* | WP_123848482.1 | WP_123845536.1 | WP_123849538.1 | WP_158618180.1 |
| Bacteroidota | *Chitinophaga niabensis* | WP_074239863.1 | WP_343304213.1 | WP_074238956.1 | WP_143197336.1 |
| Bacteroidota | *Chitinophaga niastensis* | WP_106528029.1 | WP_106526122.1 | WP_106528908.1 | WP_106530351.1 |
| Bacteroidota | *Chitinophaga nivalis* | WP_264727449.1 | WP_264735150.1 | WP_264728856.1 | WP_264732223.1 |
| Bacteroidota | *Chitinophaga oryzae* | WP_168806582.1 | WP_168807646.1 | WP_168859804.1 | WP_168804602.1 |
| Bacteroidota | *Chitinophaga oryziterrae* | WP_157299517.1 | WP_157302517.1 | WP_157302627.1 | WP_157300745.1 |
| Bacteroidota | *Chitinophaga parva* | WP_108686952.1 | WP_108684680.1 | WP_108687857.1 | WP_108687548.1 |
| Bacteroidota | *Chitinophaga pinensis* | WP_261387389.1 | WP_146305122.1 | WP_012790728.1 | WP_012792326.1 |
| Bacteroidota | *Chitinophaga pollutisoli* | WP_341835061.1 | WP_341838311.1 | WP_341836676.1 | WP_341835840.1 |
| Bacteroidota | *Chitinophaga polysaccharea* | WP_142686179.1 | WP_145660751.1 | WP_168763632.1 | WP_145670857.1 |
| Bacteroidota | *Chitinophaga qingshengii* | WP_188089689.1 | WP_188091016.1 | WP_188086724.1 | WP_188091470.1 |
| Bacteroidota | *Chitinophaga rhizophila* | WP_317197475.1 | WP_220251837.1 | WP_220253029.1 | WP_220252641.1 |
| Bacteroidota | *Chitinophaga rhizosphaerae* | WP_126246916.1 | WP_126245588.1 | WP_126249185.1 | WP_164714245.1 |
| Bacteroidota | *Chitinophaga rupis* | WP_089913986.1 | WP_089920443.1 | WP_089919493.1 | WP_089909602.1 |
| Bacteroidota | *Chitinophaga sancti* | WP_320580850.1 | WP_320580275.1 | WP_072357214.1 | WP_320573126.1 |
| Bacteroidota | *Chitinophaga silvatica* | WP_116977058.1 | WP_116975180.1 | WP_116973669.1 | WP_116976693.1 |
| Bacteroidota | *Chitinophaga silvisoli* | WP_233525504.1 | WP_116853605.1 | WP_116854710.1 | WP_116854545.1 |
| Bacteroidota | *Chitinophaga skermanii* | WP_111599050.1 | WP_111595753.1 | WP_111597493.1 | WP_111596564.1 |
| Bacteroidota | *Chitinophaga solisilvae* | WP_235938309.1 | WP_127037710.1 | WP_160712735.1 | WP_160716070.1 |
| Bacteroidota | *Chitinophaga sp.* | WP_325645883.1 | WP_343669784.1 | WP_326464793.1 | WP_298707121.1 |

| | | | | | |
|---|---|---|---|---|---|
| Bacteroidota | *Chitinophaga terrae (ex Kim and Jung 2007)* | WP_089759984.1 | WP_089758629.1 | WP_089759461.1 | WP_089760460.1 |
| Bacteroidota | *Chitinophaga tropicalis* | WP_232539105.1 | WP_157304928.1 | WP_157309866.1 | WP_157306277.1 |
| Bacteroidota | *Chitinophaga varians* | WP_188099639.1 | WP_188096056.1 | WP_188101673.1 | WP_168873033.1 |
| Bacteroidota | *Chitinophaga vietnamensis* | WP_143307862.1 | WP_143310755.1 | WP_143310472.1 | WP_143306111.1 |
| Bacteroidota | *Chryseobacterium angstadtii* | WP_048507973.1 | WP_048504691.1 | WP_048508403.1 | WP_048504839.1 |
| Bacteroidota | *Chryseobacterium aquaeductus* | WP_162088794.1 | WP_162087455.1 | WP_162088291.1 | WP_162087563.1 |
| Bacteroidota | *Chryseobacterium aquaticum* | WP_059136490.1 | WP_056017623.1 | WP_056013198.1 | WP_059135831.1 |
| Bacteroidota | *Chryseobacterium arachidis* | WP_072952915.1 | WP_072953422.1 | WP_072962042.1 | WP_072953694.1 |
| Bacteroidota | *Chryseobacterium arthrosphaerae* | WP_229047544.1 | WP_065401126.1 | WP_229045977.1 | WP_229046828.1 |
| Bacteroidota | *Chryseobacterium artocarpi* | WP_065393734.1 | WP_065393438.1 | WP_065395308.1 | WP_065395850.1 |
| Bacteroidota | *Chryseobacterium aureum* | WP_126654068.1 | WP_126653405.1 | WP_126652494.1 | WP_126653278.1 |
| Bacteroidota | *Chryseobacterium binzhouense* | WP_143884491.1 | WP_160812457.1 | WP_143885159.1 | WP_143885655.1 |
| Bacteroidota | *Chryseobacterium camelliae* | WP_309869801.1 | WP_271148067.1 | WP_100075739.1 | WP_100076441.1 |
| Bacteroidota | *Chryseobacterium carnipullorum* | WP_276729760.1 | WP_073335484.1 | WP_073329474.1 | WP_123877096.1 |
| Bacteroidota | *Chryseobacterium cheonjiense* | WP_169229803.1 | WP_169231258.1 | WP_169232082.1 | WP_169231368.1 |
| Bacteroidota | *Chryseobacterium contaminans* | WP_336959609.1 | WP_066699585.1 | WP_336963978.1 | WP_066696416.1 |
| Bacteroidota | *Chryseobacterium culicis* | WP_193545624.1 | WP_193546784.1 | WP_105681681.1 | WP_105683270.1 |
| Bacteroidota | *Chryseobacterium daecheongense* | WP_123263217.1 | WP_245142642.1 | WP_123263453.1 | WP_245142849.1 |
| Bacteroidota | *Chryseobacterium daeguense* | WP_027378272.1 | WP_027380310.1 | WP_027381351.1 | WP_027380381.1 |
| Bacteroidota | *Chryseobacterium defluvii* | WP_121461429.1 | WP_184192087.1 | WP_184190659.1 | WP_184188145.1 |
| Bacteroidota | *Chryseobacterium echinoideorum* | WP_144282438.1 | WP_144283498.1 | WP_144281767.1 | WP_144281870.1 |
| Bacteroidota | *Chryseobacterium edaphi* | WP_263001273.1 | WP_263004635.1 | WP_263003815.1 | WP_263003315.1 |
| Bacteroidota | *Chryseobacterium endalhagicum* | WP_202093532.1 | WP_202092377.1 | WP_202088381.1 | WP_202092137.1 |
| Bacteroidota | *Chryseobacterium flavum* | WP_115964410.1 | WP_115962526.1 | WP_115959980.1 | WP_333596648.1 |
| Bacteroidota | *Chryseobacterium fluminis* | WP_266174365.1 | WP_266173542.1 | WP_266172508.1 | WP_266173392.1 |
| Bacteroidota | *Chryseobacterium foetidum* | WP_262152037.1 | WP_262150941.1 | WP_262149356.1 | WP_262150698.1 |
| Bacteroidota | *Chryseobacterium formosense* | WP_034678774.1 | WP_034678117.1 | WP_034679053.1 | WP_034677951.1 |

| | | | | | |
|---|---|---|---|---|---|
| Bacteroidota | *Chryseobacterium formosus* | WP_267264994.1 | WP_267266397.1 | WP_267266997.1 | WP_267266551.1 |
| Bacteroidota | *Chryseobacterium gambrini* | WP_333781483.1 | WP_076395077.1 | WP_421745152.1 | WP_338613118.1 |
| Bacteroidota | *Chryseobacterium gilvum* | WP_262989096.1 | WP_262989998.1 | WP_262991445.1 | WP_262990122.1 |
| Bacteroidota | *Chryseobacterium ginsengisoli* | WP_345208418.1 | WP_345204417.1 | WP_345207700.1 | WP_345204007.1 |
| Bacteroidota | *Chryseobacterium ginsenosidimutans* | WP_344832014.1 | WP_344828526.1 | WP_344823491.1 | WP_344828136.1 |
| Bacteroidota | *Chryseobacterium gleum* | WP_002980773.1 | WP_343687245.1 | WP_131436182.1 | WP_002982977.1 |
| Bacteroidota | *Chryseobacterium gotjawalense* | WP_282905641.1 | WP_282903957.1 | WP_282903963.1 | WP_282903911.1 |
| Bacteroidota | *Chryseobacterium gregarium* | WP_027386164.1 | WP_027387127.1 | WP_027387487.1 | WP_027387596.1 |
| Bacteroidota | *Chryseobacterium group* | WP_063968116.1 | WP_063971113.1 | WP_063971246.1 | WP_063971373.1 |
| Bacteroidota | *Chryseobacterium gwangjuense* | WP_233111112.1 | WP_233112162.1 | WP_233112332.1 | WP_233112287.1 |
| Bacteroidota | *Chryseobacterium hagamense* | WP_146941259.1 | WP_146939999.1 | WP_146942032.1 | WP_146939862.1 |
| Bacteroidota | *Chryseobacterium herbae* | WP_259836630.1 | WP_259839341.1 | WP_259838793.1 | WP_259839482.1 |
| Bacteroidota | *Chryseobacterium indologenes* | WP_062696905.1 | WP_034737273.1 | WP_276283141.1 | WP_034734713.1 |
| Bacteroidota | *Chryseobacterium jejuense* | WP_089732913.1 | WP_209919674.1 | WP_209916422.1 | WP_089734761.1 |
| Bacteroidota | *Chryseobacterium koreense* | WP_304343934.1 | WP_048499434.1 | WP_048499442.1 | WP_048498815.1 |
| Bacteroidota | *Chryseobacterium kwangjuense* | WP_062652674.1 | WP_062650765.1 | WP_062650258.1 | WP_062651103.1 |
| Bacteroidota | *Chryseobacterium lactis* | WP_047097582.1 | WP_047098204.1 | WP_103291818.1 | WP_185290056.1 |
| Bacteroidota | *Chryseobacterium lacus* | WP_114303517.1 | WP_126730282.1 | WP_126728548.1 | WP_126731582.1 |
| Bacteroidota | *Chryseobacterium lathyri* | WP_307210283.1 | WP_306840365.1 | WP_306845405.1 | WP_111956929.1 |
| Bacteroidota | *Chryseobacterium limigenitum* | WP_072411935.1 | WP_072407513.1 | WP_072411150.1 | WP_072407276.1 |
| Bacteroidota | *Chryseobacterium luquanense* | WP_267280016.1 | WP_267280638.1 | WP_267282236.1 | WP_267280772.1 |
| Bacteroidota | *Chryseobacterium luteum* | WP_034704854.1 | WP_034702980.1 | WP_034701920.1 | WP_034702712.1 |
| Bacteroidota | *Chryseobacterium manosquense* | WP_188321635.1 | WP_123248881.1 | WP_188322246.1 | WP_188320621.1 |
| Bacteroidota | *Chryseobacterium mucoviscidosis* | WP_087708020.1 | WP_288460300.1 | WP_336717035.1 | WP_343663715.1 |
| Bacteroidota | *Chryseobacterium muglaense* | WP_191179782.1 | WP_191178277.1 | WP_191179160.1 | WP_191178394.1 |
| Bacteroidota | *Chryseobacterium nakagawai* | WP_123859852.1 | WP_123858886.1 | WP_123857931.1 | WP_123858751.1 |
| Bacteroidota | *Chryseobacterium nepalense* | WP_248392104.1 | WP_248390388.1 | WP_326988922.1 | WP_326988498.1 |
| Bacteroidota | *Chryseobacterium oleae* | WP_090024241.1 | WP_090025259.1 | WP_090025667.1 | WP_090025098.1 |

| | | | | | |
|---|---|---|---|---|---|
| Bacteroidota | *Chryseobacterium oranimense* | WP_343608012.1 | WP_040993637.1 | WP_040997539.1 | WP_343611841.1 |
| Bacteroidota | *Chryseobacterium oryctis* | WP_264743146.1 | WP_264743935.1 | WP_264742216.1 | WP_264744072.1 |
| Bacteroidota | *Chryseobacterium panacisoli* | WP_149388131.1 | WP_047382874.1 | WP_149387241.1 | WP_149387698.1 |
| Bacteroidota | *Chryseobacterium paridis* | WP_200244889.1 | WP_200241446.1 | WP_200248911.1 | WP_200241814.1 |
| Bacteroidota | *Chryseobacterium pennipullorum* | WP_115928125.1 | WP_115929148.1 | WP_115928623.1 | WP_115927319.1 |
| Bacteroidota | *Chryseobacterium phocaeense* | WP_080778555.1 | WP_080777849.1 | WP_080776683.1 | WP_080777716.1 |
| Bacteroidota | *Chryseobacterium phosphatilyticum* | WP_109712741.1 | WP_103248064.1 | WP_103247118.1 | WP_109713458.1 |
| Bacteroidota | *Chryseobacterium piperi* | WP_034685471.1 | WP_034687838.1 | WP_034681116.1 | WP_034680385.1 |
| Bacteroidota | *Chryseobacterium piscicola* | WP_076452458.1 | WP_076449729.1 | WP_076452144.1 | WP_076449478.1 |
| Bacteroidota | *Chryseobacterium polytrichastri* | WP_073293819.1 | WP_073295856.1 | WP_073293519.1 | WP_073296113.1 |
| Bacteroidota | *Chryseobacterium populi* | WP_007842630.1 | WP_007846994.1 | WP_034669435.1 | WP_007847155.1 |
| Bacteroidota | *Chryseobacterium profundimaris* | WP_283422386.1 | WP_283421978.1 | WP_283422706.1 | WP_283421026.1 |
| Bacteroidota | *Chryseobacterium salviniae* | WP_326322517.1 | WP_326320873.1 | WP_326319633.1 | WP_326321868.1 |
| Bacteroidota | *Chryseobacterium scophthalmum* | WP_210150663.1 | WP_411811350.1 | WP_074232060.1 | WP_210148717.1 |
| Bacteroidota | *Chryseobacterium sediminis* | WP_309875175.1 | WP_309876686.1 | WP_184557784.1 | WP_149833964.1 |
| Bacteroidota | *Chryseobacterium shandongense* | WP_123852139.1 | WP_123852675.1 | WP_123854981.1 | WP_123854495.1 |
| Bacteroidota | *Chryseobacterium shigense* | WP_184161100.1 | WP_076506939.1 | WP_184167674.1 | WP_076507142.1 |
| Bacteroidota | *Chryseobacterium soldanellicola* | WP_089753028.1 | WP_089754460.1 | WP_089756315.1 | WP_089754742.1 |
| Bacteroidota | *Chryseobacterium sp.* | WP_312075764.1 | WP_147756884.1 | WP_353148905.1 | WP_312389382.1 |
| Bacteroidota | *Chryseobacterium suipulveris* | WP_243550869.1 | WP_243548920.1 | WP_243548777.1 | WP_243550068.1 |
| Bacteroidota | *Chryseobacterium taeanense* | WP_374460200.1 | WP_089856114.1 | WP_374460750.1 | WP_374458606.1 |
| Bacteroidota | *Chryseobacterium tagetis* | WP_225685454.1 | WP_225686926.1 | WP_225690509.1 | WP_225687090.1 |
| Bacteroidota | *Chryseobacterium taichungense* | WP_312902571.1 | WP_312901422.1 | WP_089998393.1 | WP_312902982.1 |
| Bacteroidota | *Chryseobacterium taihuense* | WP_089742875.1 | WP_089744917.1 | WP_089744036.1 | WP_089741825.1 |
| Bacteroidota | *Chryseobacterium taiwanense* | WP_039366252.1 | WP_039364300.1 | WP_039365139.1 | WP_039364551.1 |
| Bacteroidota | *Chryseobacterium takakiae* | WP_072884494.1 | WP_072883998.1 | WP_072883461.1 | WP_072883882.1 |
| Bacteroidota | *Chryseobacterium taklimakanense* | WP_095070596.1 | WP_124806977.1 | WP_124807081.1 | WP_124785301.1 |
| Bacteroidota | *Chryseobacterium terrae* | WP_408087995.1 | WP_408090487.1 | WP_408091166.1 | WP_408090916.1 |

| | | | | | |
|---|---|---|---|---|---|
| Bacteroidota | *Chryseobacterium tructae* | WP_290300233.1 | WP_290301454.1 | WP_290295476.1 | WP_290301759.1 |
| Bacteroidota | *Chryseobacterium turcicum* | WP_230670334.1 | WP_230672020.1 | WP_230667182.1 | WP_230672276.1 |
| Bacteroidota | *Chryseobacterium ureilyticum* | WP_076552841.1 | WP_076553735.1 | WP_076551667.1 | WP_076552180.1 |
| Bacteroidota | *Chryseobacterium vietnamense* | WP_047432483.1 | WP_047429091.1 | WP_310405478.1 | WP_047427422.1 |
| Bacteroidota | *Chryseobacterium vrystaatense* | WP_034742985.1 | WP_034738641.1 | WP_073175554.1 | WP_034738968.1 |
| Bacteroidota | *Chryseobacterium wanjuense* | WP_089792497.1 | WP_089793124.1 | WP_089795693.1 | WP_089793430.1 |
| Bacteroidota | *Chryseobacterium zhengzhouense* | WP_378173883.1 | WP_378177090.1 | WP_378181010.1 | WP_378176574.1 |
| Bacteroidota | *Chryseolinea serpens* | WP_221408807.1 | WP_073133656.1 | WP_073130757.1 | WP_073134550.1 |
| Bacteroidota | *Chryseolinea soli* | WP_119756236.1 | WP_119754613.1 | WP_119757032.1 | WP_119754129.1 |
| Bacteroidota | *Cloacibacterium caeni* | WP_213195642.1 | WP_213190510.1 | WP_213196799.1 | WP_213196929.1 |
| Bacteroidota | *Cloacibacterium normanense* | WP_104792413.1 | WP_104794166.1 | WP_374451362.1 | WP_069800177.1 |
| Bacteroidota | *Cloacibacterium rupense* | WP_188617936.1 | WP_188618261.1 | WP_188616825.1 | WP_229663381.1 |
| Bacteroidota | *Cloacibacterium sp.* | WP_414984650.1 | WP_414983089.1 | WP_414982360.1 | WP_374362169.1 |
| Bacteroidota | *Cnuella takakiae* | WP_073042383.1 | WP_073048635.1 | WP_073047240.1 | WP_073046215.1 |
| Bacteroidota | *Cochleicola gelatinilyticus* | WP_068593090.1 | WP_068592651.1 | WP_068591820.1 | WP_068588824.1 |
| Bacteroidota | *Cognatitamlana onchidii* | WP_136480838.1 | WP_136481225.1 | WP_136481392.1 | WP_136480656.1 |
| Bacteroidota | *Confluentibacter citreus* | WP_100615263.1 | WP_100614548.1 | WP_100614611.1 | WP_100615531.1 |
| Bacteroidota | *Confluentibacter flavum* | WP_106658209.1 | WP_106660758.1 | WP_106658464.1 | WP_106659928.1 |
| Bacteroidota | *Confluentibacter lentus* | WP_100613067.1 | WP_100611433.1 | WP_100611373.1 | WP_100612254.1 |
| Bacteroidota | *Constantimarinum furrinae* | WP_186988948.1 | WP_186988759.1 | WP_186989389.1 | WP_186987917.1 |
| Bacteroidota | *Corallibacter sp.* | WP_418512035.1 | WP_418513145.1 | WP_418509762.1 | WP_418513932.1 |
| Bacteroidota | *Croceibacter atlanticus* | WP_341199413.1 | WP_327018545.1 | WP_341200636.1 | WP_013185972.1 |
| Bacteroidota | *Croceimicrobium hydrocarbonivorans* | WP_210759163.1 | WP_210758119.1 | WP_210758057.1 | WP_210758765.1 |
| Bacteroidota | *Croceimicrobium sp.* | WP_421754238.1 | WP_421753003.1 | WP_421752888.1 | WP_421753676.1 |
| Bacteroidota | *Crocinitomix algicola* | WP_066759458.1 | WP_070138319.1 | WP_066755789.1 | WP_066757945.1 |
| Bacteroidota | *Crocinitomix catalasitica* | WP_051568430.1 | WP_027419381.1 | WP_027421155.1 | WP_051568672.1 |
| Bacteroidota | *Cryomorpha ignava* | WP_163285727.1 | WP_163283374.1 | WP_163285353.1 | WP_163285151.1 |
| Bacteroidota | *Cytophaga aurantiaca* | WP_018341921.1 | WP_018341432.1 | WP_018344950.1 | WP_018344260.1 |

| | | | | | |
|---|---|---|---|---|---|
| Bacteroidota | *Cytophaga hutchinsonii* | WP_011583780.1 | WP_011583422.1 | WP_011585160.1 | WP_011584089.1 |
| Bacteroidota | *Daejeonella lutea* | WP_079701599.1 | WP_079702447.1 | WP_079700741.1 | WP_079703645.1 |
| Bacteroidota | *Daejeonella oryzae* | WP_026898753.1 | WP_026896934.1 | WP_026898669.1 | WP_026896820.1 |
| Bacteroidota | *Daejeonella rubra* | WP_090701791.1 | WP_090706014.1 | WP_090700380.1 | WP_090706315.1 |
| Bacteroidota | *Daejeonella sp.* | WP_305363755.1 | WP_340382233.1 | WP_395625608.1 | WP_424288394.1 |
| Bacteroidota | *Danxiaibacter flavus* | WP_369331907.1 | WP_369329328.1 | WP_369329440.1 | WP_369330972.1 |
| Bacteroidota | *Deminuibacter soli* | WP_116847103.1 | WP_116845410.1 | WP_116847565.1 | WP_116846911.1 |
| Bacteroidota | *Desertivirga arenae* | WP_207532248.1 | WP_207535454.1 | WP_207532598.1 | WP_207534650.1 |
| Bacteroidota | *Desertivirga brevis* | WP_207423621.1 | WP_207424218.1 | WP_207422904.1 | WP_207424873.1 |
| Bacteroidota | *Desertivirga xinjiangensis* | WP_256012415.1 | WP_256013323.1 | WP_256009515.1 | WP_256009196.1 |
| Bacteroidota | *Dinghuibacter silviterrae* | WP_133993125.1 | WP_133989948.1 | WP_133993427.1 | WP_133993337.1 |
| Bacteroidota | *Edaphocola aurantiacus* | WP_222165161.1 | WP_222166449.1 | WP_222167030.1 | WP_222166341.1 |
| Bacteroidota | *Edaphocola flava* | WP_129022374.1 | WP_129021451.1 | WP_129019585.1 | WP_161971320.1 |
| Bacteroidota | *Eisenibacter elegans* | WP_027000827.1 | WP_027000537.1 | WP_027000503.1 | WP_027000333.1 |
| Bacteroidota | *Elizabethkingia anophelis* | WP_035588455.1 | WP_407473270.1 | WP_021348450.1 | WP_289217335.1 |
| Bacteroidota | *Elizabethkingia argenteiflava* | WP_166520106.1 | WP_166519254.1 | WP_166519561.1 | WP_166520285.1 |
| Bacteroidota | *Elizabethkingia meningoseptica* | WP_016199328.1 | WP_087138152.1 | WP_249063179.1 | WP_407482187.1 |
| Bacteroidota | *Elizabethkingia sp. JS20170427COW* | WP_138983013.1 | WP_138982481.1 | WP_138982284.1 | WP_138981844.1 |
| Bacteroidota | *Elizabethkingia ursingii* | WP_249081401.1 | WP_059325009.1 | WP_249079041.1 | WP_249078125.1 |
| Bacteroidota | *Empedobacter brevis* | WP_353164418.1 | WP_312557397.1 | WP_312557845.1 | WP_314061045.1 |
| Bacteroidota | *Empedobacter falsenii* | WP_115002071.1 | WP_286478688.1 | WP_038336617.1 | WP_416403977.1 |
| Bacteroidota | *Empedobacter sedimenti* | WP_282628450.1 | WP_282628566.1 | WP_282630778.1 | WP_282629868.1 |
| Bacteroidota | *Empedobacter tilapiae* | WP_135835889.1 | WP_135834963.1 | WP_135834524.1 | WP_135835429.1 |
| Bacteroidota | *Emticicia aquatilis* | WP_188767157.1 | WP_188768899.1 | WP_188770815.1 | WP_188765468.1 |
| Bacteroidota | *Emticicia oligotrophica* | WP_015028859.1 | WP_015029864.1 | WP_015027285.1 | WP_015028296.1 |
| Bacteroidota | *Emticicia sp.* | WP_421768965.1 | WP_421769690.1 | WP_394993460.1 | WP_394994852.1 |
| Bacteroidota | *Epilithonimonas arachidiradicis* | WP_120213231.1 | WP_120213936.1 | WP_120213882.1 | WP_120212215.1 |
| Bacteroidota | *Epilithonimonas bovis* | WP_076784065.1 | WP_076781783.1 | WP_076783416.1 | WP_076783565.1 |

| | | | | | |
|---|---|---|---|---|---|
| Bacteroidota | *Epilithonimonas caeni* | WP_027381826.1 | WP_027383614.1 | WP_027383664.1 | WP_034671664.1 |
| Bacteroidota | *Epilithonimonas hispanica* | WP_116036308.1 | WP_116033190.1 | WP_116033058.1 | WP_116033454.1 |
| Bacteroidota | *Epilithonimonas hominis* | WP_312083011.1 | WP_313306334.1 | WP_312083915.1 | WP_312082323.1 |
| Bacteroidota | *Epilithonimonas hungarica* | WP_089874579.1 | WP_089874829.1 | WP_307311947.1 | WP_089870570.1 |
| Bacteroidota | *Epilithonimonas lactis* | WP_051879890.1 | WP_034976505.1 | WP_034976395.1 | WP_311316404.1 |
| Bacteroidota | *Epilithonimonas mollis* | WP_072995787.1 | WP_072997678.1 | WP_072996781.1 | WP_072997439.1 |
| Bacteroidota | *Epilithonimonas pallida* | WP_283415408.1 | WP_283416258.1 | WP_283416210.1 | WP_283415997.1 |
| Bacteroidota | *Epilithonimonas sp.* | WP_312418479.1 | WP_312823661.1 | WP_333852588.1 | WP_312823036.1 |
| Bacteroidota | *Epilithonimonas tenax* | WP_028123098.1 | WP_028121144.1 | WP_028121200.1 | WP_316928783.1 |
| Bacteroidota | *Epilithonimonas vandammei* | WP_124802385.1 | WP_312195794.1 | WP_124801530.1 | WP_317125027.1 |
| Bacteroidota | *Epilithonimonas xixisoli* | WP_133943661.1 | WP_133945463.1 | WP_133945376.1 | WP_133944863.1 |
| Bacteroidota | *Epilithonimonas zeae* | WP_248876890.1 | WP_248875836.1 | WP_248875783.1 | WP_317041454.1 |
| Bacteroidota | *Faecalibacter bovis* | WP_230475886.1 | WP_230477862.1 | WP_230476304.1 | WP_230477768.1 |
| Bacteroidota | *Faecalibacter macacae* | WP_121934179.1 | WP_121935498.1 | WP_121934504.1 | WP_121933187.1 |
| Bacteroidota | *Faecalibacter rhinopitheci* | WP_194182785.1 | WP_194183591.1 | WP_194181660.1 | WP_194183128.1 |
| Bacteroidota | *Faecalibacter sp. LW9* | WP_322970199.1 | WP_323674094.1 | WP_322969619.1 | WP_322971732.1 |
| Bacteroidota | *Ferruginibacter lapsinanis* | WP_229759503.1 | WP_229761199.1 | WP_229759758.1 | WP_229760830.1 |
| Bacteroidota | *Ferruginibacter sp.* | WP_301931299.1 | WP_301933928.1 | WP_301929043.1 | WP_301932499.1 |
| Bacteroidota | *Ferruginibacter yonginensis* | WP_379707491.1 | WP_379710127.1 | WP_379706315.1 | WP_379706877.1 |
| Bacteroidota | *Filimonas effusa* | WP_129002188.1 | WP_129005128.1 | WP_246022425.1 | WP_129001150.1 |
| Bacteroidota | *Filimonas lacunae* | WP_076381105.1 | WP_076379672.1 | WP_076378918.1 | WP_231940298.1 |
| Bacteroidota | *Filimonas zeae* | WP_188951172.1 | WP_188957660.1 | WP_188951939.1 | WP_188952473.1 |
| Bacteroidota | *Flammeovirga kamogawensis* | WP_144073603.1 | WP_144073762.1 | WP_144073022.1 | WP_144074686.1 |
| Bacteroidota | *Flammeovirga pacifica* | WP_044226230.1 | WP_044227781.1 | WP_044219798.1 | WP_044228724.1 |
| Bacteroidota | *Flammeovirga pectinis* | WP_126617619.1 | WP_126616529.1 | WP_126610898.1 | WP_126614491.1 |
| Bacteroidota | *Flammeovirga sp. EKP202* | WP_188170347.1 | WP_188169643.1 | WP_188168981.1 | WP_188168142.1 |
| Bacteroidota | *Flammeovirga yaeyamensis* | WP_169665818.1 | WP_169664813.1 | WP_169662846.1 | WP_169665136.1 |
| Bacteroidota | *Flaviaesturariibacter amylovorans* | WP_345256172.1 | WP_345252897.1 | WP_345255691.1 | WP_345253761.1 |

| | | | | | |
|---|---|---|---|---|---|
| Bacteroidota | *Flaviaesturariibacter aridisoli* | WP_131852011.1 | WP_131850298.1 | WP_131850986.1 | WP_131853005.1 |
| Bacteroidota | *Flaviaesturariibacter flavus* | WP_131450506.1 | WP_131449966.1 | WP_243647739.1 | WP_131449650.1 |
| Bacteroidota | *Flavicella marina* | WP_152287659.1 | WP_152286514.1 | WP_152286678.1 | WP_226913082.1 |
| Bacteroidota | *Flavicella sediminum* | WP_139959642.1 | WP_139959103.1 | WP_139959517.1 | WP_139959645.1 |
| Bacteroidota | *Flavicella sp.* | WP_336892167.1 | WP_300324116.1 | WP_336893444.1 | WP_300321354.1 |
| Bacteroidota | *Flavihumibacter cheonanensis* | WP_239596693.1 | WP_239602470.1 | WP_239595074.1 | WP_239595392.1 |
| Bacteroidota | *Flavihumibacter fluminis* | WP_234863950.1 | WP_234867994.1 | WP_234865708.1 | WP_234865854.1 |
| Bacteroidota | *Flavihumibacter fluvii* | WP_214457096.1 | WP_214458631.1 | WP_214457879.1 | WP_214457758.1 |
| Bacteroidota | *Flavihumibacter profundi* | WP_224115923.1 | WP_224117810.1 | WP_224115123.1 | WP_224115252.1 |
| Bacteroidota | *Flavihumibacter rivuli* | WP_214449044.1 | WP_214447400.1 | WP_214449400.1 | WP_239804314.1 |
| Bacteroidota | *Flavihumibacter solisilvae* | WP_039142204.1 | WP_039138289.1 | WP_039140031.1 | WP_039141623.1 |
| Bacteroidota | *Flavihumibacter sp.* | WP_332737307.1 | WP_332732135.1 | WP_332734476.1 | WP_332734753.1 |
| Bacteroidota | *Flavihumibacter stibioxidans* | WP_187257671.1 | WP_187255864.1 | WP_187255311.1 | WP_187255180.1 |
| Bacteroidota | *Flavipsychrobacter stenotrophus* | WP_105040525.1 | WP_105041010.1 | WP_105040190.1 | WP_105037087.1 |
| Bacteroidota | *Flaviramulus aquimarinus* | WP_345272025.1 | WP_345272414.1 | WP_345274344.1 | WP_345272394.1 |
| Bacteroidota | *Flaviramulus basaltis* | WP_072401373.1 | WP_072401320.1 | WP_171946619.1 | WP_072402259.1 |
| Bacteroidota | *Flaviramulus multivorans* | WP_237231591.1 | WP_237230794.1 | WP_237232560.1 | WP_237230925.1 |
| Bacteroidota | *Flaviramulus sp. BrNp1-15* | WP_238676220.1 | WP_238673743.1 | WP_238676151.1 | WP_238673896.1 |
| Bacteroidota | *Flavisolibacter ginsengisoli* | WP_072833833.1 | WP_072835562.1 | WP_072835367.1 | WP_072835393.1 |
| Bacteroidota | *Flavisolibacter ginsenosidimutans* | WP_146783909.1 | WP_146785371.1 | WP_146781711.1 | WP_146781592.1 |
| Bacteroidota | *Flavisolibacter nicotianae* | WP_121355392.1 | WP_121354209.1 | WP_121356021.1 | WP_121356571.1 |
| Bacteroidota | *Flavisolibacter tropicus* | WP_066409453.1 | WP_066404609.1 | WP_066401941.1 | WP_066402027.1 |
| Bacteroidota | *Flavivirga abyssicola* | WP_303315535.1 | WP_303317827.1 | WP_303315654.1 | WP_303319200.1 |
| Bacteroidota | *Flavivirga algicola* | WP_169669217.1 | WP_169673212.1 | WP_169669341.1 | WP_169676338.1 |
| Bacteroidota | *Flavivirga amylovorans* | WP_303283397.1 | WP_303282627.1 | WP_303283341.1 | WP_303283293.1 |
| Bacteroidota | *Flavivirga aquatica* | WP_069831156.1 | WP_069830633.1 | WP_069831205.1 | WP_069830554.1 |
| Bacteroidota | *Flavivirga aquimarina* | WP_303279537.1 | WP_303279803.1 | WP_303277252.1 | WP_303276492.1 |
| Bacteroidota | *Flavivirga eckloniae* | WP_102757772.1 | WP_102758028.1 | WP_102754615.1 | WP_102754426.1 |

| | | | | | |
|---|---|---|---|---|---|
| Bacteroidota | *Flavivirga jejuensis* | WP_303301060.1 | WP_303303193.1 | WP_303301116.1 | WP_303302142.1 |
| Bacteroidota | *Flavivirga rizhaonensis* | WP_135879042.1 | WP_135878189.1 | WP_135874426.1 | WP_135874598.1 |
| Bacteroidota | *Flavivirga sp. 57AJ16* | WP_274185376.1 | WP_274184694.1 | WP_274186157.1 | WP_274185721.1 |
| Bacteroidota | *Flavivirga spongiicola* | WP_303308613.1 | WP_303307549.1 | WP_303308519.1 | WP_303308337.1 |
| Bacteroidota | *Flavobacterium acetivorans* | WP_230062601.1 | WP_230062547.1 | WP_230061645.1 | WP_230061457.1 |
| Bacteroidota | *Flavobacterium aciduliphilum* | WP_112111896.1 | WP_112111771.1 | WP_112113984.1 | WP_112111999.1 |
| Bacteroidota | *Flavobacterium adhaerens* | WP_348799578.1 | WP_348800068.1 | WP_348799652.1 | WP_348800379.1 |
| Bacteroidota | *Flavobacterium aerium* | WP_306352775.1 | WP_306350197.1 | WP_306352814.1 | WP_306352934.1 |
| Bacteroidota | *Flavobacterium aestivum* | WP_268846035.1 | WP_268846490.1 | WP_268846139.1 | WP_281322463.1 |
| Bacteroidota | *Flavobacterium agri* | WP_176004311.1 | WP_176004808.1 | WP_176004297.1 | WP_176006078.1 |
| Bacteroidota | *Flavobacterium agrisoli* | WP_200104785.1 | WP_200104139.1 | WP_200104943.1 | WP_200105035.1 |
| Bacteroidota | *Flavobacterium ajazii* | WP_163411136.1 | WP_163410375.1 | WP_163408156.1 | WP_163408377.1 |
| Bacteroidota | *Flavobacterium akiainvivens* | WP_054406163.1 | WP_054406461.1 | WP_054409670.1 | WP_054409766.1 |
| Bacteroidota | *Flavobacterium album* | WP_108778736.1 | WP_108778230.1 | WP_108778512.1 | WP_108779563.1 |
| Bacteroidota | *Flavobacterium algicola* | WP_239861776.1 | WP_239865649.1 | WP_239867780.1 | WP_239866461.1 |
| Bacteroidota | *Flavobacterium alkalisoli* | WP_417352198.1 | WP_147583880.1 | WP_147583557.1 | WP_147584873.1 |
| Bacteroidota | *Flavobacterium alvei* | WP_103804954.1 | WP_304198350.1 | WP_103806427.1 | WP_103806165.1 |
| Bacteroidota | *Flavobacterium ammonificans* | WP_229323660.1 | WP_229329560.1 | WP_229329027.1 | WP_229329847.1 |
| Bacteroidota | *Flavobacterium ammoniigenes* | WP_229316983.1 | WP_229316756.1 | WP_229316411.1 | WP_229316318.1 |
| Bacteroidota | *Flavobacterium amnicola* | WP_129436564.1 | WP_129435686.1 | WP_129436520.1 | WP_129436066.1 |
| Bacteroidota | *Flavobacterium amniphilum* | WP_250595224.1 | WP_250595108.1 | WP_250597894.1 | WP_250598363.1 |
| Bacteroidota | *Flavobacterium anhuiense* | WP_129748659.1 | WP_091133020.1 | WP_129745604.1 | WP_091133591.1 |
| Bacteroidota | *Flavobacterium antarcticum* | WP_022828777.1 | WP_022827233.1 | WP_022828706.1 | WP_022828531.1 |
| Bacteroidota | *Flavobacterium aquariorum* | WP_111409190.1 | WP_111410466.1 | WP_111409112.1 | WP_111409004.1 |
| Bacteroidota | *Flavobacterium aquaticum* | WP_111565510.1 | WP_111567236.1 | WP_111565803.1 | WP_111565483.1 |
| Bacteroidota | *Flavobacterium aquatile* | WP_035126130.1 | WP_035127441.1 | WP_035126550.1 | WP_035126315.1 |
| Bacteroidota | *Flavobacterium aquicola* | WP_115809568.1 | WP_115812379.1 | WP_115809349.1 | WP_115810936.1 |
| Bacteroidota | *Flavobacterium aquidurense* | WP_310296061.1 | WP_310299635.1 | WP_310297587.1 | WP_395432764.1 |

| | | | | | |
|---|---|---|---|---|---|
| Bacteroidota | *Flavobacterium aquiphilum* | WP_281226196.1 | WP_281228934.1 | WP_281226080.1 | WP_281228415.1 |
| Bacteroidota | *Flavobacterium araucananum* | WP_089479424.1 | WP_089480192.1 | WP_089481795.1 | WP_089478665.1 |
| Bacteroidota | *Flavobacterium arcticum* | WP_114678501.1 | WP_114678538.1 | WP_114678788.1 | WP_114676491.1 |
| Bacteroidota | *Flavobacterium ardleyense* | WP_379803746.1 | WP_379807201.1 | WP_318640057.1 | WP_318639709.1 |
| Bacteroidota | *Flavobacterium arsenatis* | WP_310026947.1 | WP_310025109.1 | WP_310028268.1 | WP_310023609.1 |
| Bacteroidota | *Flavobacterium artemisiae* | WP_379813796.1 | WP_379816507.1 | WP_379817049.1 | WP_379816923.1 |
| Bacteroidota | *Flavobacterium arundinis* | WP_341696644.1 | WP_341695324.1 | WP_341696461.1 | WP_341696278.1 |
| Bacteroidota | *Flavobacterium aurantiibacter* | WP_094484980.1 | WP_094485081.1 | WP_094486620.1 | WP_094487510.1 |
| Bacteroidota | *Flavobacterium aureirubrum* | WP_342695876.1 | WP_342695397.1 | WP_342696707.1 | WP_342696298.1 |
| Bacteroidota | *Flavobacterium azizsancarii* | WP_271335569.1 | WP_271336029.1 | WP_271338417.1 | WP_271334397.1 |
| Bacteroidota | *Flavobacterium azooxidireducens* | WP_248433839.1 | WP_248434158.1 | WP_248433393.1 | WP_248433145.1 |
| Bacteroidota | *Flavobacterium beibuense* | WP_129749981.1 | WP_129751414.1 | WP_035132715.1 | WP_129751784.1 |
| Bacteroidota | *Flavobacterium bernardetii* | WP_166130803.1 | WP_166125315.1 | WP_166126073.1 | WP_166126115.1 |
| Bacteroidota | *Flavobacterium bizetiae* | WP_173972779.1 | WP_263759732.1 | WP_173971207.1 | WP_173971103.1 |
| Bacteroidota | *Flavobacterium branchiarum* | WP_290265837.1 | WP_290262884.1 | WP_290262510.1 | WP_290266136.1 |
| Bacteroidota | *Flavobacterium branchiicola* | WP_213255567.1 | WP_213258301.1 | WP_213258164.1 | WP_213257900.1 |
| Bacteroidota | *Flavobacterium branchiophilum* | WP_097553948.1 | WP_089081239.1 | WP_014083730.1 | WP_089081437.1 |
| Bacteroidota | *Flavobacterium buctense* | WP_187660344.1 | WP_187660824.1 | WP_187660755.1 | WP_187660569.1 |
| Bacteroidota | *Flavobacterium caeni* | WP_091144087.1 | WP_091141017.1 | WP_091140782.1 | WP_091139974.1 |
| Bacteroidota | *Flavobacterium calami* | WP_341694229.1 | WP_341690908.1 | WP_341692856.1 | WP_341692452.1 |
| Bacteroidota | *Flavobacterium capsici* | WP_313321799.1 | WP_313321563.1 | WP_313323035.1 | WP_313325783.1 |
| Bacteroidota | *Flavobacterium caseinilyticum* | WP_131909939.1 | WP_131908690.1 | WP_131910707.1 | WP_131909271.1 |
| Bacteroidota | *Flavobacterium cauense* | WP_023571930.1 | WP_023571266.1 | WP_023571697.1 | WP_023570087.1 |
| Bacteroidota | *Flavobacterium celericrescens* | WP_166236013.1 | WP_166235729.1 | WP_166236799.1 | WP_166235918.1 |
| Bacteroidota | *Flavobacterium cellulosilyticum* | WP_132009313.1 | WP_132006492.1 | WP_132006986.1 | WP_132004434.1 |
| Bacteroidota | *Flavobacterium cerinum* | WP_256551113.1 | WP_128388995.1 | WP_256549717.1 | WP_256552875.1 |
| Bacteroidota | *Flavobacterium channae* | WP_231836571.1 | WP_231834529.1 | WP_231836887.1 | WP_231836528.1 |
| Bacteroidota | *Flavobacterium cheongpyeongense* | WP_110306664.1 | WP_110307633.1 | WP_110304959.1 | WP_110304752.1 |

| | | | | | |
|---|---|---|---|---|---|
| Bacteroidota | *Flavobacterium chilense* | WP_068843172.1 | WP_068842693.1 | WP_068842835.1 | WP_068844503.1 |
| Bacteroidota | *Flavobacterium chuncheonense* | WP_379811074.1 | WP_379810082.1 | WP_379811473.1 | WP_379811128.1 |
| Bacteroidota | *Flavobacterium chungnamense* | WP_345089596.1 | WP_345094576.1 | WP_345095656.1 | WP_345093744.1 |
| Bacteroidota | *Flavobacterium circumlabens* | WP_132037869.1 | WP_132035216.1 | WP_348079458.1 | WP_132037989.1 |
| Bacteroidota | *Flavobacterium collinsii* | WP_173966751.1 | WP_281924965.1 | WP_263361526.1 | WP_395340080.1 |
| Bacteroidota | *Flavobacterium columnare* | WP_192188487.1 | WP_014165519.1 | WP_138424961.1 | WP_163444573.1 |
| Bacteroidota | *Flavobacterium coralii* | WP_348676667.1 | WP_223052048.1 | WP_223053230.1 | WP_348678002.1 |
| Bacteroidota | *Flavobacterium crassostreae* | WP_066334673.1 | WP_066335673.1 | WP_066334792.1 | WP_066334607.1 |
| Bacteroidota | *Flavobacterium croceum* | WP_245874595.1 | WP_282787780.1 | WP_103726256.1 | WP_103726419.1 |
| Bacteroidota | *Flavobacterium crocinum* | WP_109192690.1 | WP_109194175.1 | WP_109194355.1 | WP_109194604.1 |
| Bacteroidota | *Flavobacterium cucumis* | WP_073583744.1 | WP_073584520.1 | WP_073582549.1 | WP_073580360.1 |
| Bacteroidota | *Flavobacterium cupreum* | WP_127337142.1 | WP_127338736.1 | WP_127336437.1 | WP_127339709.1 |
| Bacteroidota | *Flavobacterium cutihirudinis* | WP_115886603.1 | WP_115888499.1 | WP_115888668.1 | WP_115888903.1 |
| Bacteroidota | *Flavobacterium cyanobacteriorum* | WP_094412888.1 | WP_094416373.1 | WP_094412463.1 | WP_094417039.1 |
| Bacteroidota | *Flavobacterium cyclinae* | WP_231841939.1 | WP_231843804.1 | WP_231841638.1 | WP_231841964.1 |
| Bacteroidota | *Flavobacterium daejeonense* | WP_026715165.1 | WP_026714153.1 | WP_026715500.1 | WP_026714113.1 |
| Bacteroidota | *Flavobacterium daemonense* | WP_144337204.1 | WP_144339228.1 | WP_144337841.1 | WP_144338222.1 |
| Bacteroidota | *Flavobacterium dankookense* | WP_133533032.1 | WP_133531808.1 | WP_133531489.1 | WP_133533978.1 |
| Bacteroidota | *Flavobacterium dauae* | WP_129758356.1 | WP_129758015.1 | WP_129757852.1 | WP_129757042.1 |
| Bacteroidota | *Flavobacterium defluvii* | WP_073413815.1 | WP_073416481.1 | WP_073416641.1 | WP_073417082.1 |
| Bacteroidota | *Flavobacterium degerlachei* | WP_091429533.1 | WP_091434817.1 | WP_091429243.1 | WP_091434165.1 |
| Bacteroidota | *Flavobacterium denitrificans* | WP_026729249.1 | WP_026730144.1 | WP_026730335.1 | WP_026728034.1 |
| Bacteroidota | *Flavobacterium difficile* | WP_166078192.1 | WP_166077145.1 | WP_166078006.1 | WP_166077778.1 |
| Bacteroidota | *Flavobacterium eburneipallidum* | WP_269225030.1 | WP_281336077.1 | WP_269225176.1 | WP_281337225.1 |
| Bacteroidota | *Flavobacterium enshiense* | WP_023574756.1 | WP_023572418.1 | WP_367230350.1 | WP_023574299.1 |
| Bacteroidota | *Flavobacterium erciyesense* | WP_210789057.1 | WP_210788498.1 | WP_210788461.1 | WP_210788629.1 |
| Bacteroidota | *Flavobacterium facile* | WP_329804406.1 | WP_329804786.1 | WP_329806052.1 | WP_329806076.1 |
| Bacteroidota | *Flavobacterium faecale* | WP_108741003.1 | WP_108741200.1 | WP_108740786.1 | WP_418263472.1 |

| | | | | | |
|---|---|---|---|---|---|
| Bacteroidota | *Flavobacterium filum* | WP_051220712.1 | WP_276979020.1 | WP_026709932.1 | WP_026711516.1 |
| Bacteroidota | *Flavobacterium flavigenum* | WP_269235311.1 | WP_269236635.1 | WP_269236927.1 | WP_269237110.1 |
| Bacteroidota | *Flavobacterium flavipallidum* | WP_341700562.1 | WP_341700249.1 | WP_341700978.1 | WP_341701690.1 |
| Bacteroidota | *Flavobacterium flevense* | WP_073245238.1 | WP_073245585.1 | WP_073245546.1 | WP_073244749.1 |
| Bacteroidota | *Flavobacterium fluviale* | WP_113678179.1 | WP_113679386.1 | WP_113680010.1 | WP_113679743.1 |
| Bacteroidota | *Flavobacterium fluviatile* | WP_163399140.1 | WP_163399805.1 | WP_163401290.1 | WP_163400519.1 |
| Bacteroidota | *Flavobacterium fluvii* | WP_073372003.1 | WP_073370950.1 | WP_073371333.1 | WP_073370273.1 |
| Bacteroidota | *Flavobacterium foetidum* | WP_135220181.1 | WP_135223692.1 | WP_135220578.1 | WP_135220849.1 |
| Bacteroidota | *Flavobacterium fontis* | WP_073363541.1 | WP_073362329.1 | WP_073365076.1 | WP_073362776.1 |
| Bacteroidota | *Flavobacterium fragile* | WP_250582634.1 | WP_250581805.1 | WP_250582693.1 | WP_250582588.1 |
| Bacteroidota | *Flavobacterium franklandianum* | WP_143389910.1 | WP_143391881.1 | WP_143391434.1 | WP_143391604.1 |
| Bacteroidota | *Flavobacterium frigidarium* | WP_339654646.1 | WP_026707310.1 | WP_371570404.1 | WP_371571863.1 |
| Bacteroidota | *Flavobacterium frigidimaris* | WP_074657062.1 | WP_074658449.1 | WP_074658704.1 | WP_074659117.1 |
| Bacteroidota | *Flavobacterium frigoris* | WP_007138811.1 | WP_007139028.1 | WP_074723038.1 | WP_074723885.1 |
| Bacteroidota | *Flavobacterium frigoritolerans* | WP_264286204.1 | WP_264287449.1 | WP_264287159.1 | WP_264285323.1 |
| Bacteroidota | *Flavobacterium fructosi* | WP_379858511.1 | WP_379857536.1 | WP_379858922.1 | WP_379857307.1 |
| Bacteroidota | *Flavobacterium galactosidilyticum* | WP_229942134.1 | WP_229942020.1 | WP_229941185.1 | WP_229943974.1 |
| Bacteroidota | *Flavobacterium gelatinilyticum* | WP_281234421.1 | WP_281232719.1 | WP_281232539.1 | WP_281232268.1 |
| Bacteroidota | *Flavobacterium gelidilacus* | WP_051191491.1 | WP_026715688.1 | WP_026716442.1 | WP_026716198.1 |
| Bacteroidota | *Flavobacterium gillisiae* | WP_091086804.1 | WP_091086427.1 | WP_091093031.1 | WP_091090389.1 |
| Bacteroidota | *Flavobacterium gilvum* | WP_035634759.1 | WP_035637836.1 | WP_035638905.1 | WP_035640009.1 |
| Bacteroidota | *Flavobacterium ginsengiterrae* | WP_345144824.1 | WP_345146172.1 | WP_345145973.1 | WP_345145719.1 |
| Bacteroidota | *Flavobacterium ginsenosidimutans* | WP_111286560.1 | WP_111285252.1 | WP_111285576.1 | WP_111289076.1 |
| Bacteroidota | *Flavobacterium glaciei* | WP_114754681.1 | WP_114755205.1 | WP_114755304.1 | WP_114753358.1 |
| Bacteroidota | *Flavobacterium glycines* | WP_066330025.1 | WP_066329059.1 | WP_066329643.1 | WP_066324645.1 |
| Bacteroidota | *Flavobacterium granuli* | WP_310003045.1 | WP_072937916.1 | WP_072941919.1 | WP_310008176.1 |
| Bacteroidota | *Flavobacterium gyeonganense* | WP_278009297.1 | WP_278010294.1 | WP_278010560.1 | WP_278010745.1 |
| Bacteroidota | *Flavobacterium haoranii* | WP_072784910.1 | WP_072783792.1 | WP_072785678.1 | WP_072785004.1 |

| | | | | | |
|---|---|---|---|---|---|
| Bacteroidota | *Flavobacterium hercynium* | WP_089047846.1 | WP_089051503.1 | WP_089050641.1 | WP_089051567.1 |
| Bacteroidota | *Flavobacterium hibernum* | WP_041518017.1 | WP_041518361.1 | WP_041515948.1 | WP_041517094.1 |
| Bacteroidota | *Flavobacterium hibisci* | WP_223681520.1 | WP_223680455.1 | WP_223683385.1 | WP_223681683.1 |
| Bacteroidota | *Flavobacterium humi* | WP_135524624.1 | WP_135524654.1 | WP_135526473.1 | WP_135526235.1 |
| Bacteroidota | *Flavobacterium humidisoli* | WP_248726213.1 | WP_248727515.1 | WP_248727640.1 | WP_248727894.1 |
| Bacteroidota | *Flavobacterium hungaricum* | WP_193844941.1 | WP_194139755.1 | WP_194139922.1 | WP_194140192.1 |
| Bacteroidota | *Flavobacterium hydatis* | WP_035627509.1 | WP_035620475.1 | WP_035627907.1 | WP_035625949.1 |
| Bacteroidota | *Flavobacterium hydrocarbonoxydans* | WP_160374738.1 | WP_160373147.1 | WP_160376079.1 | WP_160375866.1 |
| Bacteroidota | *Flavobacterium hydrophilum* | WP_110348265.1 | WP_110345293.1 | WP_110344961.1 | WP_110344757.1 |
| Bacteroidota | *Flavobacterium ichthyis* | WP_166537492.1 | WP_166536024.1 | WP_166535779.1 | WP_166537713.1 |
| Bacteroidota | *Flavobacterium jejuense* | WP_140960659.1 | WP_140963139.1 | WP_140959233.1 | WP_140960741.1 |
| Bacteroidota | *Flavobacterium johnsoniae* | WP_149207923.1 | WP_071638765.1 | WP_289877969.1 | WP_354017319.1 |
| Bacteroidota | *Flavobacterium kayseriense* | WP_187010675.1 | WP_187011015.1 | WP_187011005.1 | WP_187010795.1 |
| Bacteroidota | *Flavobacterium kingsejongi* | WP_108738093.1 | WP_108737943.1 | WP_108737494.1 | WP_108738362.1 |
| Bacteroidota | *Flavobacterium lacus* | WP_112085686.1 | WP_112085796.1 | WP_112085450.1 | WP_112086584.1 |
| Bacteroidota | *Flavobacterium lacustre* | WP_269685402.1 | WP_269684458.1 | WP_269685141.1 | WP_269684167.1 |
| Bacteroidota | *Flavobacterium laiguense* | WP_116762616.1 | WP_116763267.1 | WP_116764434.1 | WP_116760445.1 |
| Bacteroidota | *Flavobacterium limi* | WP_163395014.1 | WP_163396595.1 | WP_163394177.1 | WP_163394387.1 |
| Bacteroidota | *Flavobacterium limicola* | WP_121364806.1 | WP_121365842.1 | WP_121365603.1 | WP_121365642.1 |
| Bacteroidota | *Flavobacterium limnophilum* | WP_281299277.1 | WP_281299364.1 | WP_281299001.1 | WP_281297214.1 |
| Bacteroidota | *Flavobacterium limnosediminis* | WP_023580149.1 | WP_023579171.1 | WP_023579392.1 | WP_023580327.1 |
| Bacteroidota | *Flavobacterium litorale* | WP_220640035.1 | WP_220639998.1 | WP_220640443.1 | WP_220640681.1 |
| Bacteroidota | *Flavobacterium longum* | WP_415579833.1 | WP_415580331.1 | WP_415580030.1 | WP_415581162.1 |
| Bacteroidota | *Flavobacterium lotistagni* | WP_166336344.1 | WP_166335680.1 | WP_166336123.1 | WP_166334538.1 |
| Bacteroidota | *Flavobacterium luminosum* | WP_250592192.1 | WP_250592150.1 | WP_250591819.1 | WP_250591576.1 |
| Bacteroidota | *Flavobacterium luteum* | WP_151108361.1 | WP_151108338.1 | WP_151106222.1 | WP_151107915.1 |
| Bacteroidota | *Flavobacterium macacae* | WP_125012638.1 | WP_125011741.1 | WP_125013480.1 | WP_125012357.1 |
| Bacteroidota | *Flavobacterium magnesitis* | WP_373392694.1 | WP_373391948.1 | WP_373393194.1 | WP_373390870.1 |

| | | | | | |
|---|---|---|---|---|---|
| Bacteroidota | *Flavobacterium magnum* | WP_108371479.1 | WP_108371837.1 | WP_108373551.1 | WP_108370611.1 |
| Bacteroidota | *Flavobacterium maritimum* | WP_348813958.1 | WP_348812361.1 | WP_348811647.1 | WP_348811765.1 |
| Bacteroidota | *Flavobacterium microcysteis* | WP_140001285.1 | WP_140000093.1 | WP_140001393.1 | WP_140002987.1 |
| Bacteroidota | *Flavobacterium micromati* | WP_073018050.1 | WP_073021752.1 | WP_073021956.1 | WP_073021419.1 |
| Bacteroidota | *Flavobacterium muglaense* | WP_187018573.1 | WP_187020500.1 | WP_187019474.1 | WP_187016867.1 |
| Bacteroidota | *Flavobacterium myungsuense* | WP_379756042.1 | WP_379759358.1 | WP_379758019.1 | WP_379755506.1 |
| Bacteroidota | *Flavobacterium nackdongense* | WP_133275741.1 | WP_133277831.1 | WP_133275103.1 | WP_133275856.1 |
| Bacteroidota | *Flavobacterium nakdongensis* | WP_309531785.1 | WP_309533082.1 | WP_309532800.1 | WP_309531613.1 |
| Bacteroidota | *Flavobacterium nitratireducens* | WP_278035989.1 | WP_278035832.1 | WP_278035364.1 | WP_278035249.1 |
| Bacteroidota | *Flavobacterium noncentrifugens* | WP_091394897.1 | WP_091396656.1 | WP_091398032.1 | WP_091393211.1 |
| Bacteroidota | *Flavobacterium odoriferum* | WP_262319179.1 | WP_262318657.1 | WP_262318477.1 | WP_262317886.1 |
| Bacteroidota | *Flavobacterium okayamense* | WP_221257746.1 | WP_221258181.1 | WP_221258373.1 | WP_221257683.1 |
| Bacteroidota | *Flavobacterium olei* | WP_347051710.1 | WP_347049574.1 | WP_347049694.1 | WP_347049942.1 |
| Bacteroidota | *Flavobacterium oreochromis* | WP_235873409.1 | WP_088397939.1 | WP_088399325.1 | WP_088397908.1 |
| Bacteroidota | *Flavobacterium orientale* | WP_188361824.1 | WP_188361792.1 | WP_188361733.1 | WP_188361861.1 |
| Bacteroidota | *Flavobacterium ovatum* | WP_366187315.1 | WP_366183810.1 | WP_366183988.1 | WP_366184218.1 |
| Bacteroidota | *Flavobacterium pallidum* | WP_108904360.1 | WP_108904208.1 | WP_108904408.1 | WP_108903708.1 |
| Bacteroidota | *Flavobacterium palustre* | WP_188495722.1 | WP_188493166.1 | WP_188493705.1 | WP_188493752.1 |
| Bacteroidota | *Flavobacterium paronense* | WP_290285321.1 | WP_290285180.1 | WP_290285908.1 | WP_290284915.1 |
| Bacteroidota | *Flavobacterium pectinovorum* | WP_302095401.1 | WP_302096861.1 | WP_140511608.1 | WP_302097218.1 |
| Bacteroidota | *Flavobacterium petrolei* | WP_395381288.1 | WP_113666649.1 | WP_113666264.1 | WP_395380779.1 |
| Bacteroidota | *Flavobacterium phragmitis* | WP_091491246.1 | WP_091492985.1 | WP_091491738.1 | WP_091492158.1 |
| Bacteroidota | *Flavobacterium phycosphaerae* | WP_162127828.1 | WP_162127750.1 | WP_162126086.1 | WP_162128342.1 |
| Bacteroidota | *Flavobacterium piscinae* | WP_129464091.1 | WP_321538495.1 | WP_129464344.1 | WP_129464983.1 |
| Bacteroidota | *Flavobacterium pisciphilum* | WP_229990811.1 | WP_229987522.1 | WP_229987734.1 | WP_229990525.1 |
| Bacteroidota | *Flavobacterium piscis* | WP_310282236.1 | WP_310284124.1 | WP_065450509.1 | WP_310281901.1 |
| Bacteroidota | *Flavobacterium piscisymbiosum* | WP_230036937.1 | WP_230039643.1 | WP_230039970.1 | WP_230040703.1 |
| Bacteroidota | *Flavobacterium plantiphilum* | WP_408081925.1 | WP_408078841.1 | WP_408078801.1 | WP_408081465.1 |

| | | | | | |
|---|---|---|---|---|---|
| Bacteroidota | *Flavobacterium pokkalii* | WP_188221639.1 | WP_188220226.1 | WP_055092636.1 | WP_188221459.1 |
| Bacteroidota | *Flavobacterium polysaccharolyticum* | WP_342692751.1 | WP_342692940.1 | WP_342690826.1 | WP_342691523.1 |
| Bacteroidota | *Flavobacterium ponti* | WP_379742522.1 | WP_379740641.1 | WP_379741904.1 | WP_379743091.1 |
| Bacteroidota | *Flavobacterium potami* | WP_223705374.1 | WP_223704457.1 | WP_223704287.1 | WP_223704074.1 |
| Bacteroidota | *Flavobacterium poyangense* | WP_166924366.1 | WP_166923811.1 | WP_166925746.1 | WP_166922132.1 |
| Bacteroidota | *Flavobacterium praedii* | WP_281240167.1 | WP_281239829.1 | WP_281240117.1 | WP_281239177.1 |
| Bacteroidota | *Flavobacterium procerum* | WP_379686047.1 | WP_379682565.1 | WP_379682729.1 | WP_379683027.1 |
| Bacteroidota | *Flavobacterium profundi* | WP_140998711.1 | WP_140996821.1 | WP_140999128.1 | WP_140998869.1 |
| Bacteroidota | *Flavobacterium proteolyticum* | WP_194096385.1 | WP_194095444.1 | WP_194095749.1 | WP_194094760.1 |
| Bacteroidota | *Flavobacterium psychraquaticum* | WP_322550477.1 | WP_322549922.1 | WP_322549451.1 | WP_322550351.1 |
| Bacteroidota | *Flavobacterium psychrophilum* | WP_341966078.1 | WP_011962921.1 | WP_046205375.1 | WP_247440291.1 |
| Bacteroidota | *Flavobacterium psychroterrae* | WP_213304194.1 | WP_213298471.1 | WP_213303007.1 | WP_213305312.1 |
| Bacteroidota | *Flavobacterium psychrotolerans* | WP_116724932.1 | WP_116725783.1 | WP_116723503.1 | WP_116723560.1 |
| Bacteroidota | *Flavobacterium psychrotrophum* | WP_116789094.1 | WP_116788075.1 | WP_116788784.1 | WP_116789352.1 |
| Bacteroidota | *Flavobacterium pygoscelis* | WP_248427670.1 | WP_248427562.1 | WP_248429168.1 | WP_248427783.1 |
| Bacteroidota | *Flavobacterium qiangtangense* | WP_379789644.1 | WP_379790684.1 | WP_379791238.1 | WP_379792899.1 |
| Bacteroidota | *Flavobacterium rakeshii* | WP_157481558.1 | WP_157483142.1 | WP_330122173.1 | WP_330122791.1 |
| Bacteroidota | *Flavobacterium reichenbachii* | WP_035683762.1 | WP_035683155.1 | WP_035682855.1 | WP_035690269.1 |
| Bacteroidota | *Flavobacterium restrictum* | WP_144255659.1 | WP_144255860.1 | WP_144254725.1 | WP_144257666.1 |
| Bacteroidota | *Flavobacterium rhizophilum* | WP_408072782.1 | WP_408075709.1 | WP_408073956.1 | WP_408074961.1 |
| Bacteroidota | *Flavobacterium rhizosphaerae* | WP_408083500.1 | WP_408083731.1 | WP_408083897.1 | WP_408083921.1 |
| Bacteroidota | *Flavobacterium rivuli* | WP_020211280.1 | WP_020214032.1 | WP_020212488.1 | WP_020212097.1 |
| Bacteroidota | *Flavobacterium rivulicola* | WP_171223067.1 | WP_171222509.1 | WP_171221989.1 | WP_171223169.1 |
| Bacteroidota | *Flavobacterium saccharophilum* | WP_072971291.1 | WP_072972051.1 | WP_072972649.1 | WP_072973028.1 |
| Bacteroidota | *Flavobacterium saliperosum* | WP_023575827.1 | WP_023576809.1 | WP_023576527.1 | WP_023577062.1 |
| Bacteroidota | *Flavobacterium sandaracinum* | WP_132065033.1 | WP_132066739.1 | WP_132064968.1 | WP_132064488.1 |
| Bacteroidota | *Flavobacterium sangjuense* | WP_136152239.1 | WP_136152510.1 | WP_136152911.1 | WP_136152372.1 |
| Bacteroidota | *Flavobacterium sasangense* | WP_026726258.1 | WP_026725803.1 | WP_026726094.1 | WP_026724613.1 |

| | | | | | |
|---|---|---|---|---|---|
| Bacteroidota | *Flavobacterium sedimenticola* | WP_283240149.1 | WP_283238790.1 | WP_283238321.1 | WP_283238626.1 |
| Bacteroidota | *Flavobacterium segetis* | WP_072991066.1 | WP_072992850.1 | WP_072993824.1 | WP_072987071.1 |
| Bacteroidota | *Flavobacterium selenitireducens* | WP_191019087.1 | WP_191018474.1 | WP_191018872.1 | WP_191020079.1 |
| Bacteroidota | *Flavobacterium seoulense* | WP_035658105.1 | WP_035660257.1 | WP_035661625.1 | WP_035662614.1 |
| Bacteroidota | *Flavobacterium sharifuzzamanii* | WP_111424150.1 | WP_111426281.1 | WP_111426401.1 | WP_111426645.1 |
| Bacteroidota | *Flavobacterium shii* | WP_264205074.1 | WP_264206984.1 | WP_264204354.1 | WP_264206383.1 |
| Bacteroidota | *Flavobacterium silvaticum* | WP_169525831.1 | WP_169528127.1 | WP_169526441.1 | WP_169528369.1 |
| Bacteroidota | *Flavobacterium silvisoli* | WP_131476404.1 | WP_131476214.1 | WP_131476810.1 | WP_131475440.1 |
| Bacteroidota | *Flavobacterium sinopsychrotolerans* | WP_091172020.1 | WP_091168489.1 | WP_091166664.1 | WP_091166756.1 |
| Bacteroidota | *Flavobacterium soli* | WP_026705388.1 | WP_026705573.1 | WP_026705749.1 | WP_026704342.1 |
| Bacteroidota | *Flavobacterium solisilvae* | WP_169523010.1 | WP_169523750.1 | WP_169524792.1 | WP_169524984.1 |
| Bacteroidota | *Flavobacterium soyae* | WP_232682820.1 | WP_406844002.1 | WP_406844131.1 | WP_232678917.1 |
| Bacteroidota | *Flavobacterium soyangense* | WP_194312891.1 | WP_194311707.1 | WP_194311327.1 | WP_194310813.1 |
| Bacteroidota | *Flavobacterium sp.* | WP_324218812.1 | WP_310560047.1 | WP_298143336.1 | WP_326401577.1 |
| Bacteroidota | *Flavobacterium stagni* | WP_129461543.1 | WP_129461143.1 | WP_129462148.1 | WP_129460802.1 |
| Bacteroidota | *Flavobacterium suaedae* | WP_188619340.1 | WP_188621315.1 | WP_188620323.1 | WP_188620523.1 |
| Bacteroidota | *Flavobacterium subsaxonicum* | WP_026992171.1 | WP_026991950.1 | WP_026990105.1 | WP_026993033.1 |
| Bacteroidota | *Flavobacterium succinicans* | WP_024980019.1 | WP_064714826.1 | WP_064715172.1 | WP_064715246.1 |
| Bacteroidota | *Flavobacterium sufflavum* | WP_128197814.1 | WP_128196693.1 | WP_128194842.1 | WP_128194048.1 |
| Bacteroidota | *Flavobacterium suncheonense* | WP_026979736.1 | WP_026981611.1 | WP_026979946.1 | WP_026981151.1 |
| Bacteroidota | *Flavobacterium supellecticarium* | WP_136402496.1 | WP_136404178.1 | WP_136401869.1 | WP_136403284.1 |
| Bacteroidota | *Flavobacterium swingsii* | WP_091474450.1 | WP_091474524.1 | WP_091474723.1 | WP_091472898.1 |
| Bacteroidota | *Flavobacterium tagetis* | WP_202001004.1 | WP_202003963.1 | WP_202003572.1 | WP_202006294.1 |
| Bacteroidota | *Flavobacterium taihuense* | WP_219316622.1 | WP_219318647.1 | WP_219318958.1 | WP_219319062.1 |
| Bacteroidota | *Flavobacterium tegetincola* | WP_026977841.1 | WP_026978617.1 | WP_026977492.1 | WP_026978195.1 |
| Bacteroidota | *Flavobacterium terrae* | WP_073310989.1 | WP_073310928.1 | WP_073311178.1 | WP_073309335.1 |
| Bacteroidota | *Flavobacterium terrigena* | WP_091315599.1 | WP_091313686.1 | WP_091315883.1 | WP_091311805.1 |
| Bacteroidota | *Flavobacterium terrisoli* | WP_284652054.1 | WP_284652565.1 | WP_284653435.1 | WP_284652285.1 |

| | | | | | |
|---|---|---|---|---|---|
| Bacteroidota | *Flavobacterium tibetense* | WP_113988678.1 | WP_113988761.1 | WP_113989048.1 | WP_113988628.1 |
| Bacteroidota | *Flavobacterium tistrianum* | WP_111366714.1 | WP_111365996.1 | WP_111366193.1 | WP_111364009.1 |
| Bacteroidota | *Flavobacterium turcicum* | WP_166137451.1 | WP_166135359.1 | WP_166135473.1 | WP_166134952.1 |
| Bacteroidota | *Flavobacterium undicola* | WP_161643988.1 | WP_161640825.1 | WP_161643606.1 | WP_161643714.1 |
| Bacteroidota | *Flavobacterium urocaniciphilum* | WP_091470295.1 | WP_091465144.1 | WP_091468925.1 | WP_091468962.1 |
| Bacteroidota | *Flavobacterium urumqiense* | WP_103999328.1 | WP_104000151.1 | WP_104000107.1 | WP_103999846.1 |
| Bacteroidota | *Flavobacterium ustbae* | WP_125719466.1 | WP_125722837.1 | WP_125720253.1 | WP_125723234.1 |
| Bacteroidota | *Flavobacterium xinjiangense* | WP_073207932.1 | WP_073211262.1 | WP_073208242.1 | WP_073207460.1 |
| Bacteroidota | *Flavobacterium xueshanense* | WP_091206530.1 | WP_091206099.1 | WP_091207614.1 | WP_091203754.1 |
| Bacteroidota | *Flavobacterium xylosi* | WP_379853312.1 | WP_379854913.1 | WP_379853196.1 | WP_379854551.1 |
| Bacteroidota | *Flavobacterium yafengii* | WP_282727115.1 | WP_282716422.1 | WP_282715470.1 | WP_282727969.1 |
| Bacteroidota | *Flavobacterium zepuense* | WP_143373431.1 | WP_143372536.1 | WP_143375269.1 | WP_143373232.1 |
| Bacteroidota | *Flavobacterium zhairuonense* | WP_132990185.1 | WP_132988062.1 | WP_132989944.1 | WP_132989696.1 |
| Bacteroidota | *Flavobacterium zhouii* | WP_379851541.1 | WP_379852713.1 | WP_379851381.1 | WP_379849947.1 |
| Bacteroidota | *Flavobacterium zubiriense* | WP_373405786.1 | WP_373406039.1 | WP_373406299.1 | WP_373406703.1 |
| Bacteroidota | *Flectobacillus longus* | WP_283326372.1 | WP_283325519.1 | WP_283325847.1 | WP_283328046.1 |
| Bacteroidota | *Flectobacillus major* | WP_026996775.1 | WP_026997808.1 | WP_026994392.1 | WP_026997855.1 |
| Bacteroidota | *Flectobacillus roseus* | WP_283344929.1 | WP_283345382.1 | WP_283343837.1 | WP_283374206.1 |
| Bacteroidota | *Flectobacillus sp.* | WP_421821822.1 | WP_421820662.1 | WP_421819229.1 | WP_421818880.1 |
| Bacteroidota | *Flexibacter flexilis* | WP_091511107.1 | WP_091508130.1 | WP_091505765.1 | WP_091516603.1 |
| Bacteroidota | *Flexibacter sp. ATCC 35103* | WP_083691640.1 | WP_083692816.1 | WP_083692959.1 | WP_083693241.1 |
| Bacteroidota | *Fluviicola chungangensis* | WP_186279992.1 | WP_144331325.1 | WP_186280074.1 | WP_186280185.1 |
| Bacteroidota | *Fluviicola sp.* | WP_300358642.1 | WP_300663211.1 | WP_343636787.1 | WP_300659961.1 |
| Bacteroidota | *Fluviicola taffensis* | WP_013687018.1 | WP_341901437.1 | WP_013687603.1 | WP_013687422.1 |
| Bacteroidota | *Foetidibacter luteolus* | WP_235920994.1 | WP_153796776.1 | WP_153800762.1 | WP_153795974.1 |
| Bacteroidota | *Formosa agariphila* | WP_038528683.1 | WP_038529567.1 | WP_038528827.1 | WP_038527824.1 |
| Bacteroidota | *Formosa algae* | WP_057779463.1 | WP_057779342.1 | WP_057778720.1 | WP_103191423.1 |
| Bacteroidota | *Formosa maritima* | WP_148454452.1 | WP_148452936.1 | WP_148454901.1 | WP_148455345.1 |

| | | | | | |
|---|---|---|---|---|---|
| Bacteroidota | *Formosa sediminum* | WP_143381647.1 | WP_143381701.1 | WP_143382029.1 | WP_143380981.1 |
| Bacteroidota | *Formosa sp. Hel1_33_131* | WP_069677819.1 | WP_069677461.1 | WP_069675596.1 | WP_069675818.1 |
| Bacteroidota | *Formosa undariae* | WP_382381504.1 | WP_382381391.1 | WP_382381605.1 | WP_382382124.1 |
| Bacteroidota | *Frigoriflavimonas asaccharolytica* | WP_173778168.1 | WP_173780209.1 | WP_173780018.1 | WP_173779288.1 |
| Bacteroidota | *Fulvitalea axinellae* | WP_338393436.1 | WP_338392299.1 | WP_338391720.1 | WP_338391676.1 |
| Bacteroidota | *Fulvivirga imtechensis* | WP_009580067.1 | WP_009583631.1 | WP_009577485.1 | WP_009579199.1 |
| Bacteroidota | *Fulvivirga kasyanovii* | WP_155168941.1 | WP_155174132.1 | WP_155171568.1 | WP_343855452.1 |
| Bacteroidota | *Fulvivirga ligni* | WP_233772449.1 | WP_233771349.1 | WP_233770555.1 | WP_233771846.1 |
| Bacteroidota | *Fulvivirga lutea* | WP_205722999.1 | WP_205721864.1 | WP_205721098.1 | WP_205722411.1 |
| Bacteroidota | *Fulvivirga lutimaris* | WP_155186002.1 | WP_155187492.1 | WP_155185506.1 | WP_155188308.1 |
| Bacteroidota | *Fulvivirga marina* | WP_202859358.1 | WP_202854470.1 | WP_202855140.1 | WP_202854505.1 |
| Bacteroidota | *Fulvivirga maritima* | WP_233779464.1 | WP_233778822.1 | WP_233780993.1 | WP_233780020.1 |
| Bacteroidota | *Fulvivirga sedimenti* | WP_225699088.1 | WP_225698760.1 | WP_225696347.1 | WP_225699844.1 |
| Bacteroidota | *Fulvivirga sediminis* | WP_202241851.1 | WP_202244114.1 | WP_202243775.1 | WP_202244855.1 |
| Bacteroidota | *Fulvivirga sp.* | WP_350112549.1 | WP_350112158.1 | WP_350116307.1 | WP_350111703.1 |
| Bacteroidota | *Fulvivirga ulvae* | WP_233766756.1 | WP_233767761.1 | WP_233762065.1 | WP_233765613.1 |
| Bacteroidota | *Gaetbulibacter aquiaggeris* | WP_395438225.1 | WP_395438398.1 | WP_395438696.1 | WP_395437480.1 |
| Bacteroidota | *Gaetbulibacter saemankumensis* | WP_027137536.1 | WP_027136548.1 | WP_027136618.1 | WP_027136785.1 |
| Bacteroidota | *Gaetbulibacter sp. M240* | WP_394875070.1 | WP_394875138.1 | WP_394875025.1 | WP_394875879.1 |
| Bacteroidota | *Gangjinia marincola* | WP_343765107.1 | WP_343762526.1 | WP_343764592.1 | WP_343764901.1 |
| Bacteroidota | *Gelatiniphilus marinus* | WP_388016973.1 | WP_388014987.1 | WP_388013439.1 | WP_388016878.1 |
| Bacteroidota | *Gelidibacter algens* | WP_066430462.1 | WP_066431358.1 | WP_066437546.1 | WP_111625946.1 |
| Bacteroidota | *Gelidibacter gilvus* | WP_129016401.1 | WP_129016144.1 | WP_129017242.1 | WP_129017912.1 |
| Bacteroidota | *Gelidibacter japonicus* | WP_323026877.1 | WP_163515830.1 | WP_250066750.1 | WP_417368856.1 |
| Bacteroidota | *Gelidibacter maritimus* | WP_182204285.1 | WP_182206455.1 | WP_182204067.1 | WP_182205246.1 |
| Bacteroidota | *Gelidibacter mesophilus* | WP_027125198.1 | WP_027126355.1 | WP_027127133.1 | WP_027125360.1 |
| Bacteroidota | *Gelidibacter pelagius* | WP_208232877.1 | WP_208233111.1 | WP_208233378.1 | WP_208232401.1 |
| Bacteroidota | *Gelidibacter salicanalis* | WP_146893854.1 | WP_199602477.1 | WP_146889517.1 | WP_146888335.1 |

| | | | | | |
|---|---|---|---|---|---|
| Bacteroidota | *Gelidibacter sediminis* | WP_133758292.1 | WP_133758180.1 | WP_133757911.1 | WP_133757493.1 |
| Bacteroidota | *Gelidibacter sp.* | WP_325578379.1 | WP_348037356.1 | WP_325448570.1 | WP_348035391.1 |
| Bacteroidota | *Geojedonia litorea* | WP_387961785.1 | WP_387961949.1 | WP_387961441.1 | WP_387960537.1 |
| Bacteroidota | *Gilvibacter sediminis* | WP_272831957.1 | WP_272832536.1 | WP_272832132.1 | WP_272833033.1 |
| Bacteroidota | *Gilvibacter sp.* | WP_291187662.1 | WP_291185014.1 | WP_374960310.1 | WP_420379054.1 |
| Bacteroidota | *Gilvirhabdus luticola* | WP_316663263.1 | WP_316663118.1 | WP_316662346.1 | WP_316661412.1 |
| Bacteroidota | *Ginsengibacter hankyongi* | WP_150412771.1 | WP_150413724.1 | WP_150415216.1 | WP_150413131.1 |
| Bacteroidota | *Haloflavibacter putidus* | WP_141421426.1 | WP_141420749.1 | WP_141421610.1 | WP_141422424.1 |
| Bacteroidota | *Halpernia frigidisoli* | WP_090082314.1 | WP_090079850.1 | WP_090080935.1 | WP_177205450.1 |
| Bacteroidota | *Halpernia humi* | WP_103913750.1 | WP_103913058.1 | WP_103913318.1 | WP_103913510.1 |
| Bacteroidota | *Halpernia sp.* | WP_417430268.1 | WP_417428735.1 | WP_417428411.1 | WP_417427724.1 |
| Bacteroidota | *Hanstruepera flava* | WP_250433844.1 | WP_250436433.1 | WP_250432405.1 | WP_250434084.1 |
| Bacteroidota | *Hanstruepera marina* | WP_223033503.1 | WP_223034528.1 | WP_223033481.1 | WP_223033900.1 |
| Bacteroidota | *Hanstruepera neustonica* | WP_103052605.1 | WP_103052034.1 | WP_103052338.1 | WP_103051335.1 |
| Bacteroidota | *Hanstruepera ponticola* | WP_191859824.1 | WP_191859655.1 | WP_191860971.1 | WP_104734393.1 |
| Bacteroidota | *Hufsiella arboris* | WP_160845686.1 | WP_160844963.1 | WP_160844516.1 | WP_160845831.1 |
| Bacteroidota | *Hugenholtzia roseola* | WP_027003369.1 | WP_027001197.1 | WP_027003762.1 | WP_051203851.1 |
| Bacteroidota | *Hwangdonia lutea* | WP_316983296.1 | WP_316983175.1 | WP_316983375.1 | WP_316982178.1 |
| Bacteroidota | *Hwangdonia seohaensis* | WP_311938766.1 | WP_311938626.1 | WP_311938917.1 | WP_311939926.1 |
| Bacteroidota | *Hwangdonia sp.* | WP_418604510.1 | WP_418604453.1 | WP_418604389.1 | WP_418603464.1 |
| Bacteroidota | *Hydrobacter penzbergensis* | WP_092726552.1 | WP_092724081.1 | WP_092725408.1 | WP_092722670.1 |
| Bacteroidota | *Hymenobacter actinosclerus* | WP_092773040.1 | WP_092772166.1 | WP_092770383.1 | WP_092773236.1 |
| Bacteroidota | *Hymenobacter aerophilus* | WP_019949270.1 | WP_019947749.1 | WP_019946669.1 | WP_019946883.1 |
| Bacteroidota | *Hymenobacter algoricola* | WP_345113875.1 | WP_345113237.1 | WP_345114690.1 | WP_345109111.1 |
| Bacteroidota | *Hymenobacter amundsenii* | WP_088462812.1 | WP_088464214.1 | WP_088464474.1 | WP_088462720.1 |
| Bacteroidota | *Hymenobacter aquaticus* | WP_135463777.1 | WP_210114293.1 | WP_135461986.1 | WP_135463988.1 |
| Bacteroidota | *Hymenobacter busanensis* | WP_151078495.1 | WP_151079239.1 | WP_338074383.1 | WP_151078604.1 |
| Bacteroidota | *Hymenobacter canadensis* | WP_269559131.1 | WP_269560067.1 | WP_269561050.1 | WP_269558985.1 |

| | | | | | |
|---|---|---|---|---|---|
| Bacteroidota | *Hymenobacter chitinivorans* | WP_100337718.1 | WP_100334769.1 | WP_100335002.1 | WP_100337998.1 |
| Bacteroidota | *Hymenobacter daecheongensis* | WP_073105469.1 | WP_073108984.1 | WP_073111368.1 | WP_073105191.1 |
| Bacteroidota | *Hymenobacter duratus* | WP_211539194.1 | WP_190784662.1 | WP_190785735.1 | WP_190783158.1 |
| Bacteroidota | *Hymenobacter edaphi* | WP_111480507.1 | WP_111478724.1 | WP_111476768.1 | WP_111477643.1 |
| Bacteroidota | *Hymenobacter elongatus* | WP_135498979.1 | WP_135499093.1 | WP_135499484.1 | WP_135498247.1 |
| Bacteroidota | *Hymenobacter endophyticus* | WP_315998093.1 | WP_315999127.1 | WP_315997629.1 | WP_315997288.1 |
| Bacteroidota | *Hymenobacter fastidiosus* | WP_345074911.1 | WP_345074853.1 | WP_345072700.1 | WP_345071857.1 |
| Bacteroidota | *Hymenobacter fodinae* | WP_135430104.1 | WP_135433580.1 | WP_135436110.1 | WP_135430550.1 |
| Bacteroidota | *Hymenobacter gelipurpurascens* | WP_088844785.1 | WP_088843741.1 | WP_088842140.1 | WP_088844610.1 |
| Bacteroidota | *Hymenobacter glacieicola* | WP_188557127.1 | WP_188559509.1 | WP_188558562.1 | WP_188556977.1 |
| Bacteroidota | *Hymenobacter gummosus* | WP_126694282.1 | WP_126696445.1 | WP_126694959.1 | WP_126694911.1 |
| Bacteroidota | *Hymenobacter guriensis* | WP_317191392.1 | WP_196953312.1 | WP_231403234.1 | WP_196956148.1 |
| Bacteroidota | *Hymenobacter jeollabukensis* | WP_138082352.1 | WP_138077952.1 | WP_138074696.1 | WP_138076488.1 |
| Bacteroidota | *Hymenobacter koreensis* | WP_345227100.1 | WP_345220804.1 | WP_345225765.1 | WP_345227550.1 |
| Bacteroidota | *Hymenobacter lapidiphilus* | WP_176907463.1 | WP_176907383.1 | WP_176906325.1 | WP_176910239.1 |
| Bacteroidota | *Hymenobacter lucidus* | WP_226170117.1 | WP_226175553.1 | WP_226177909.1 | WP_226170797.1 |
| Bacteroidota | *Hymenobacter metallicola* | WP_135396638.1 | WP_135394722.1 | WP_135394102.1 | WP_135395606.1 |
| Bacteroidota | *Hymenobacter metallilatus* | WP_125431828.1 | WP_125427055.1 | WP_125432977.1 | WP_125426004.1 |
| Bacteroidota | *Hymenobacter nitidus* | WP_226186498.1 | WP_226189119.1 | WP_226183341.1 | WP_226186023.1 |
| Bacteroidota | *Hymenobacter oligotrophus* | WP_119445448.1 | WP_119444743.1 | WP_119443979.1 | WP_119445303.1 |
| Bacteroidota | *Hymenobacter perfusus* | WP_125438156.1 | WP_125439651.1 | WP_125440281.1 | WP_125435945.1 |
| Bacteroidota | *Hymenobacter persicinus* | WP_129920492.1 | WP_129922767.1 | WP_129923205.1 | WP_129923055.1 |
| Bacteroidota | *Hymenobacter pini* | WP_226270965.1 | WP_226265934.1 | WP_226269985.1 | WP_226271625.1 |
| Bacteroidota | *Hymenobacter piscis* | WP_215593080.1 | WP_215594052.1 | WP_215592429.1 | WP_215591981.1 |
| Bacteroidota | *Hymenobacter psychrophilus* | WP_092740064.1 | WP_092739454.1 | WP_092741564.1 | WP_092739666.1 |
| Bacteroidota | *Hymenobacter psychrotolerans* | WP_073287058.1 | WP_073286580.1 | WP_073287465.1 | WP_073285851.1 |
| Bacteroidota | *Hymenobacter qilianensis* | WP_187733482.1 | WP_187732856.1 | WP_187732044.1 | WP_187733782.1 |
| Bacteroidota | *Hymenobacter rigui* | WP_125424374.1 | WP_125420006.1 | WP_125423685.1 | WP_125417329.1 |

| | | | | | |
|---|---|---|---|---|---|
| Bacteroidota | *Hymenobacter roseosalivarius* | WP_084443675.1 | WP_084444424.1 | WP_084447454.1 | WP_084445205.1 |
| Bacteroidota | *Hymenobacter rubripertinctus* | WP_119655126.1 | WP_119656221.1 | WP_119654072.1 | WP_119654547.1 |
| Bacteroidota | *Hymenobacter sediminicola* | WP_260625764.1 | WP_185887369.1 | WP_185886273.1 | WP_185888447.1 |
| Bacteroidota | *Hymenobacter sediminis* | WP_110976193.1 | WP_202910460.1 | WP_110979241.1 | WP_110976343.1 |
| Bacteroidota | *Hymenobacter sp.* | WP_331147061.1 | WP_331059006.1 | WP_331059402.1 | WP_325135416.1 |
| Bacteroidota | *Hymenobacter sublimis* | WP_247975281.1 | WP_247975982.1 | WP_247976812.1 | WP_247975169.1 |
| Bacteroidota | *Hymenobacter swuensis* | WP_044000558.1 | WP_044001162.1 | WP_044002076.1 | WP_044003628.1 |
| Bacteroidota | *Hymenobacter terrestris* | WP_176898131.1 | WP_176900196.1 | WP_176898993.1 | WP_176901120.1 |
| Bacteroidota | *Hymenobacter tibetensis* | WP_243798354.1 | WP_243800611.1 | WP_243802657.1 | WP_243799028.1 |
| Bacteroidota | *Hymenobacter translucens* | WP_227606985.1 | WP_227609277.1 | WP_227606247.1 | WP_227610336.1 |
| Bacteroidota | *Hymenobacter volaticus* | WP_245120172.1 | WP_245122496.1 | WP_245125362.1 | WP_245120586.1 |
| Bacteroidota | *Hymenobacter wooponensis* | WP_135532540.1 | WP_135529106.1 | WP_135530529.1 | WP_135530138.1 |
| Bacteroidota | *Hymenobacter yonginensis* | WP_270126485.1 | WP_270127571.1 | WP_270128601.1 | WP_270126306.1 |
| Bacteroidota | *Hyunsoonleella aestuarii* | WP_139002422.1 | WP_139002770.1 | WP_139000583.1 | WP_139000362.1 |
| Bacteroidota | *Hyunsoonleella aquatilis* | WP_186562002.1 | WP_186563531.1 | WP_186563450.1 | WP_186559073.1 |
| Bacteroidota | *Hyunsoonleella flava* | WP_130962751.1 | WP_130962624.1 | WP_130962429.1 | WP_130963622.1 |
| Bacteroidota | *Hyunsoonleella jejuensis* | WP_092578310.1 | WP_092578219.1 | WP_092578669.1 | WP_092579037.1 |
| Bacteroidota | *Hyunsoonleella pacifica* | WP_130935561.1 | WP_130935536.1 | WP_130935871.1 | WP_130938023.1 |
| Bacteroidota | *Hyunsoonleella rubra* | WP_380293872.1 | WP_380288079.1 | WP_380289797.1 | WP_380292807.1 |
| Bacteroidota | *Hyunsoonleella sp. 2307UL5-6* | WP_413972126.1 | WP_413972052.1 | WP_413973366.1 | WP_413973261.1 |
| Bacteroidota | *Hyunsoonleella ulvae* | WP_203256791.1 | WP_203256821.1 | WP_203258597.1 | WP_203257890.1 |
| Bacteroidota | *Ichthyenterobacterium magnum* | WP_120201057.1 | WP_120200732.1 | WP_120201318.1 | WP_120199279.1 |
| Bacteroidota | *Ichthyobacterium seriolicida* | WP_096686620.1 | WP_096686479.1 | WP_096686568.1 | WP_096685584.1 |
| Bacteroidota | *Imtechella halotolerans* | WP_008239022.1 | WP_008240408.1 | WP_008238821.1 | WP_008238112.1 |
| Bacteroidota | *Jejudonia soesokkakensis* | WP_380217216.1 | WP_380216994.1 | WP_380217385.1 | WP_380218067.1 |
| Bacteroidota | *Jejuia pallidilutea* | WP_042244445.1 | WP_105474530.1 | WP_105474849.1 | WP_105474438.1 |
| Bacteroidota | *Jejuia sp. DST062* | WP_406972028.1 | WP_406971608.1 | WP_406969494.1 | WP_406968753.1 |
| Bacteroidota | *Jejuia spongiicola* | WP_249973538.1 | WP_249972989.1 | WP_249973577.1 | WP_249972470.1 |

| | | | | | |
|---|---|---|---|---|---|
| Bacteroidota | *Kaistella antarctica* | WP_034716011.1 | WP_034718549.1 | WP_034718534.1 | WP_051803858.1 |
| Bacteroidota | *Kaistella carnis* | WP_125022700.1 | WP_313717127.1 | WP_312818541.1 | WP_125024677.1 |
| Bacteroidota | *Kaistella chaponensis* | WP_076388336.1 | WP_076386389.1 | WP_076386374.1 | WP_076387281.1 |
| Bacteroidota | *Kaistella daneshvariae* | WP_124758469.1 | WP_124757691.1 | WP_124757689.1 | WP_123266721.1 |
| Bacteroidota | *Kaistella faecalis* | WP_218249738.1 | WP_218248450.1 | WP_218248442.1 | WP_218250327.1 |
| Bacteroidota | *Kaistella flava (ex Peng et al. 2021)* | WP_193812873.1 | WP_193811537.1 | WP_193811529.1 | WP_193811028.1 |
| Bacteroidota | *Kaistella gelatinilytica* | WP_196079166.1 | WP_196079956.1 | WP_196079963.1 | WP_196080199.1 |
| Bacteroidota | *Kaistella haifensis* | WP_031501825.1 | WP_031501350.1 | WP_031501345.1 | WP_031504621.1 |
| Bacteroidota | *Kaistella jeonii* | WP_039352692.1 | WP_039351184.1 | WP_039351160.1 | WP_039350477.1 |
| Bacteroidota | *Kaistella montana* | WP_255927104.1 | WP_255929243.1 | WP_255929267.1 | WP_255930280.1 |
| Bacteroidota | *Kaistella palustris* | WP_027376429.1 | WP_027376032.1 | WP_027376024.1 | WP_027377070.1 |
| Bacteroidota | *Kaistella polysaccharea* | WP_226064635.1 | WP_226063347.1 | WP_226063355.1 | WP_226063808.1 |
| Bacteroidota | *Kaistella soli* | WP_217199978.1 | WP_088358885.1 | WP_217201330.1 | WP_217201368.1 |
| Bacteroidota | *Kaistella solincola* | WP_039345848.1 | WP_039342678.1 | WP_039341850.1 | WP_039341001.1 |
| Bacteroidota | *Kaistella sp.* | WP_332032649.1 | WP_373708837.1 | WP_423975190.1 | WP_373707369.1 |
| Bacteroidota | *Kaistella treverensis* | WP_089819399.1 | WP_089819909.1 | WP_089819511.1 | WP_089819931.1 |
| Bacteroidota | *Kaistella yananensis* | WP_265144198.1 | WP_265142890.1 | WP_265142884.1 | WP_265144070.1 |
| Bacteroidota | *Kaistella yonginensis* | WP_290220396.1 | WP_290216955.1 | WP_290216940.1 | WP_290217812.1 |
| Bacteroidota | *Kordia aestuariivivens* | WP_187561876.1 | WP_187563470.1 | WP_187561095.1 | WP_187564525.1 |
| Bacteroidota | *Kordia algicida* | WP_007093217.1 | WP_007093984.1 | WP_040559864.1 | WP_007094736.1 |
| Bacteroidota | *Kordia antarctica* | WP_160128858.1 | WP_160128089.1 | WP_160128402.1 | WP_160131354.1 |
| Bacteroidota | *Kordia jejudonensis* | WP_046757610.1 | WP_046758503.1 | WP_046758012.1 | WP_046757471.1 |
| Bacteroidota | *Kordia periserrulae* | WP_108113611.1 | WP_108114659.1 | WP_108114435.1 | WP_108116523.1 |
| Bacteroidota | *Kordia sp.* | WP_420574344.1 | WP_420573743.1 | WP_420572871.1 | WP_290671858.1 |
| Bacteroidota | *Kordia zhangzhouensis* | WP_082106780.1 | WP_046744617.1 | WP_046744409.1 | WP_046745889.1 |
| Bacteroidota | *Lacibacter cauensis* | WP_144884070.1 | WP_144886451.1 | WP_144884137.1 | WP_144886127.1 |
| Bacteroidota | *Lacibacter luteus* | WP_129132569.1 | WP_129131595.1 | WP_129128817.1 | WP_129132186.1 |
| Bacteroidota | *Lacibacter sediminis* | WP_182801160.1 | WP_182803974.1 | WP_182801129.1 | WP_182801935.1 |

| | | | | | |
|---|---|---|---|---|---|
| Bacteroidota | *Lacibacter sp.* | WP_324232101.1 | WP_324231404.1 | WP_324232070.1 | WP_324230093.1 |
| Bacteroidota | *Lacihabitans lacunae* | WP_379838358.1 | WP_379835146.1 | WP_379834768.1 | WP_379838877.1 |
| Bacteroidota | *Lacihabitans soyangensis* | WP_255038614.1 | WP_255038281.1 | WP_255039128.1 | WP_255037505.1 |
| Bacteroidota | *Lacihabitans sp. CS3-21* | WP_255050028.1 | WP_255050370.1 | WP_255047189.1 | WP_255048821.1 |
| Bacteroidota | *Lacinutrix algicola* | WP_055436825.1 | WP_055437101.1 | WP_055437076.1 | WP_055435440.1 |
| Bacteroidota | *Lacinutrix gracilariae* | WP_379900224.1 | WP_379901067.1 | WP_379899746.1 | WP_379902270.1 |
| Bacteroidota | *Lacinutrix himadriensis* | WP_055443177.1 | WP_055444954.1 | WP_055443481.1 | WP_055445236.1 |
| Bacteroidota | *Lacinutrix iliipiscaria* | WP_183489379.1 | WP_183487450.1 | WP_183488070.1 | WP_183486898.1 |
| Bacteroidota | *Lacinutrix jangbogonensis* | WP_034058149.1 | WP_034058557.1 | WP_034060840.1 | WP_034057457.1 |
| Bacteroidota | *Lacinutrix mariniflava* | WP_055445619.1 | WP_055446055.1 | WP_055448589.1 | WP_055448788.1 |
| Bacteroidota | *Lacinutrix sp.* | WP_290696520.1 | WP_340169257.1 | WP_300207521.1 | WP_290698518.1 |
| Bacteroidota | *Lacinutrix venerupis* | WP_121049029.1 | WP_121048868.1 | WP_121048213.1 | WP_076732624.1 |
| Bacteroidota | *Leptobacterium meishanense* | WP_340074781.1 | WP_340077561.1 | WP_340073967.1 | WP_340074705.1 |
| Bacteroidota | *Limibacter armeniacum* | WP_334246972.1 | WP_334246576.1 | WP_334246496.1 | WP_334246523.1 |
| Bacteroidota | *Limnovirga soli* | WP_171607574.1 | WP_171609007.1 | WP_171609473.1 | WP_171609767.1 |
| Bacteroidota | *Lishizhenia sp.* | WP_318959256.1 | WP_318958285.1 | WP_318958486.1 | WP_318958685.1 |
| Bacteroidota | *Lishizhenia tianjinensis* | WP_090247069.1 | WP_090246723.1 | WP_090247519.1 | WP_090245381.1 |
| Bacteroidota | *Litoribacter populi* | WP_143959586.1 | WP_143962451.1 | WP_143960986.1 | WP_143959597.1 |
| Bacteroidota | *Litoribacter ruber* | WP_213944562.1 | WP_213943340.1 | WP_213945420.1 | WP_213944573.1 |
| Bacteroidota | *Litoribaculum gwangyangense* | WP_345276546.1 | WP_345276104.1 | WP_345276075.1 | WP_345276767.1 |
| Bacteroidota | *Longitalea arenae* | WP_205512728.1 | WP_205508771.1 | WP_205512896.1 | WP_205509770.1 |
| Bacteroidota | *Longitalea luteola* | WP_207513380.1 | WP_207511955.1 | WP_207513435.1 | WP_207513616.1 |
| Bacteroidota | *Luteibaculum oceani* | WP_147014622.1 | WP_147014709.1 | WP_147014190.1 | WP_147013057.1 |
| Bacteroidota | *Luteirhabdus pelagi* | WP_203295889.1 | WP_203295780.1 | WP_203296683.1 | WP_203296609.1 |
| Bacteroidota | *Lutibacter agarilyticus* | WP_089379847.1 | WP_089380012.1 | WP_089380156.1 | WP_089381860.1 |
| Bacteroidota | *Lutibacter citreus* | WP_111708422.1 | WP_111708460.1 | WP_111707821.1 | WP_111708609.1 |
| Bacteroidota | *Lutibacter flavus* | WP_089377158.1 | WP_089377124.1 | WP_089376832.1 | WP_089377379.1 |
| Bacteroidota | *Lutibacter holmesii* | WP_386809318.1 | WP_386809209.1 | WP_386809111.1 | WP_386807184.1 |

| | | | | | |
|---|---|---|---|---|---|
| Bacteroidota | *Lutibacter maritimus* | WP_090223066.1 | WP_090222967.1 | WP_090222428.1 | WP_090221845.1 |
| Bacteroidota | *Lutibacter oceani* | WP_115879669.1 | WP_115879279.1 | WP_115879655.1 | WP_115882099.1 |
| Bacteroidota | *Lutibacter oricola* | WP_090121788.1 | WP_090121462.1 | WP_090120722.1 | WP_090123020.1 |
| Bacteroidota | *Lutibacter profundi* | WP_068211365.1 | WP_068205635.1 | WP_068205690.1 | WP_068206420.1 |
| Bacteroidota | *Lutibacter sp.* | WP_291974348.1 | WP_368051282.1 | WP_305600318.1 | WP_372794956.1 |
| Bacteroidota | *Lutimonas halocynthiae* | WP_290292455.1 | WP_290293069.1 | WP_290293732.1 | WP_290289555.1 |
| Bacteroidota | *Lutimonas saemankumensis* | WP_224927108.1 | WP_224927446.1 | WP_224927476.1 | WP_224932014.1 |
| Bacteroidota | *Lutimonas sp.* | WP_424352615.1 | WP_424350756.1 | WP_424350450.1 | WP_424352018.1 |
| Bacteroidota | *Lutimonas vermicola* | WP_342158652.1 | WP_342158194.1 | WP_342159627.1 | WP_342159791.1 |
| Bacteroidota | *Lutimonas zeaxanthinifaciens* | WP_302045662.1 | WP_302043014.1 | WP_302043042.1 | WP_302043417.1 |
| Bacteroidota | *Mangrovimonas aestuarii* | WP_274475618.1 | WP_274475567.1 | WP_274475634.1 | WP_274475942.1 |
| Bacteroidota | *Mangrovimonas cancribranchiae* | WP_338733612.1 | WP_338733894.1 | WP_338733664.1 | WP_338733268.1 |
| Bacteroidota | *Mangrovimonas futianensis* | WP_232395652.1 | WP_232634821.1 | WP_232395550.1 | WP_232634478.1 |
| Bacteroidota | *Mangrovimonas sp. YM274* | WP_307899412.1 | WP_307899351.1 | WP_307902005.1 | WP_307901681.1 |
| Bacteroidota | *Mangrovimonas spongiae* | WP_125467229.1 | WP_125468490.1 | WP_125467189.1 | WP_125467469.1 |
| Bacteroidota | *Mangrovimonas xylaniphaga* | WP_053977740.1 | WP_053977790.1 | WP_053977261.1 | WP_053977459.1 |
| Bacteroidota | *Mangrovimonas yunxiaonensis* | WP_036122858.1 | WP_036123792.1 | WP_036123026.1 | WP_036122376.1 |
| Bacteroidota | *Mangrovivirga halotolerans* | WP_266056956.1 | WP_266058647.1 | WP_266056794.1 | WP_266057182.1 |
| Bacteroidota | *Mariniflexile aquimaris* | WP_379942699.1 | WP_379942415.1 | WP_379942788.1 | WP_379941225.1 |
| Bacteroidota | *Mariniflexile fucanivorans* | WP_132216225.1 | WP_132219156.1 | WP_132215582.1 | WP_132216338.1 |
| Bacteroidota | *Mariniflexile gromovii* | WP_209654374.1 | WP_209656289.1 | WP_209657102.1 | WP_209654042.1 |
| Bacteroidota | *Mariniflexile jejuense* | WP_379926659.1 | WP_379924078.1 | WP_379926576.1 | WP_379926525.1 |
| Bacteroidota | *Mariniflexile litorale* | WP_308992018.1 | WP_308992851.1 | WP_308992244.1 | WP_308992065.1 |
| Bacteroidota | *Mariniflexile maritimum* | WP_157209100.1 | WP_157206636.1 | WP_157208718.1 | WP_157206179.1 |
| Bacteroidota | *Mariniflexile ostreae* | WP_379861387.1 | WP_379860781.1 | WP_379861361.1 | WP_379860627.1 |
| Bacteroidota | *Mariniflexile soesokkakense* | WP_346242364.1 | WP_346242016.1 | WP_346239891.1 | WP_346242332.1 |
| Bacteroidota | *Mariniflexile sp.* | WP_372754610.1 | WP_372755176.1 | WP_372754476.1 | WP_372754686.1 |
| Bacteroidota | *Marinigracilibium pacificum* | WP_169683144.1 | WP_169684667.1 | WP_169684819.1 | WP_169677532.1 |

| | | | | | |
|---|---|---|---|---|---|
| Bacteroidota | *Marinirhabdus gelatinilytica* | WP_115124603.1 | WP_115124645.1 | WP_115123303.1 | WP_115123992.1 |
| Bacteroidota | *Marinoscillum furvescens* | WP_115866421.1 | WP_115866794.1 | WP_245986162.1 | WP_115868181.1 |
| Bacteroidota | *Marinoscillum sp. MHG1-6* | WP_258102520.1 | WP_258103226.1 | WP_258104476.1 | WP_258102800.1 |
| Bacteroidota | *Marivirga arenosa* | WP_308355675.1 | WP_308356037.1 | WP_302125633.1 | WP_322347713.1 |
| Bacteroidota | *Marivirga atlantica* | WP_201918422.1 | WP_201918138.1 | WP_201924100.1 | WP_201916581.1 |
| Bacteroidota | *Marivirga aurantiaca* | WP_201430705.1 | WP_201432316.1 | WP_201429891.1 | WP_201432555.1 |
| Bacteroidota | *Marivirga harenae* | WP_303270671.1 | WP_303267928.1 | WP_303272113.1 | WP_303270879.1 |
| Bacteroidota | *Marivirga lumbricoides* | WP_188464610.1 | WP_188460934.1 | WP_188462202.1 | WP_188465104.1 |
| Bacteroidota | *Marivirga salinae* | WP_308349634.1 | WP_308351288.1 | WP_308347826.1 | WP_308349786.1 |
| Bacteroidota | *Marivirga sericea* | WP_085519183.1 | WP_085517122.1 | WP_085515726.1 | WP_085518009.1 |
| Bacteroidota | *Marivirga sp.* | WP_296620148.1 | WP_296618831.1 | WP_296622034.1 | WP_325121820.1 |
| Bacteroidota | *Marivirga tractuosa* | WP_375579149.1 | WP_013455427.1 | WP_013452380.1 | WP_375579341.1 |
| Bacteroidota | *Marixanthomonas ophiurae* | WP_117160414.1 | WP_117160044.1 | WP_117160475.1 | WP_117159718.1 |
| Bacteroidota | *Marixanthomonas sp. SCSIO 43207* | WP_223108255.1 | WP_223107920.1 | WP_223107512.1 | WP_223109995.1 |
| Bacteroidota | *Marixanthomonas spongiae* | WP_116695087.1 | WP_116693667.1 | WP_116693396.1 | WP_116694569.1 |
| Bacteroidota | *Marnyiella aurantia* | WP_209815840.1 | WP_181886854.1 | WP_181887822.1 | WP_181886945.1 |
| Bacteroidota | *Mesoflavibacter sp.* | WP_370225504.1 | WP_370225638.1 | WP_370228053.1 | WP_370225799.1 |
| Bacteroidota | *Mesoflavibacter zeaxanthinifaciens* | WP_304142908.1 | WP_304144207.1 | WP_106678698.1 | WP_106679414.1 |
| Bacteroidota | *Mesohalobacter halotolerans* | WP_138931837.1 | WP_138931676.1 | WP_138932211.1 | WP_138932670.1 |
| Bacteroidota | *Mesonia aestuariivivens* | WP_219039632.1 | WP_219039071.1 | WP_219040755.1 | WP_219041146.1 |
| Bacteroidota | *Mesonia algae* | WP_111541660.1 | WP_111540188.1 | WP_111541586.1 | WP_111541566.1 |
| Bacteroidota | *Mesonia aquimarina* | WP_121665614.1 | WP_121665309.1 | WP_121665496.1 | WP_121666282.1 |
| Bacteroidota | *Mesonia hippocampi* | WP_183477565.1 | WP_183477614.1 | WP_183475703.1 | WP_183477477.1 |
| Bacteroidota | *Mesonia maritima* | WP_309727274.1 | WP_309729227.1 | WP_309728181.1 | WP_309728893.1 |
| Bacteroidota | *Mesonia mobilis* | WP_304133107.1 | WP_027883935.1 | WP_304157464.1 | WP_304156199.1 |
| Bacteroidota | *Mesonia ostreae* | WP_311400708.1 | WP_311401302.1 | WP_311400781.1 | WP_311400862.1 |
| Bacteroidota | *Mesonia profundi* | WP_308864155.1 | WP_308863989.1 | WP_308864680.1 | WP_308863498.1 |
| Bacteroidota | *Mesonia sediminis* | WP_379044607.1 | WP_379044484.1 | WP_379046647.1 | WP_379045708.1 |

| | | | | | |
|---|---|---|---|---|---|
| Bacteroidota | *Mesonia sp.* | WP_347420137.1 | WP_347419748.1 | WP_292246739.1 | WP_347418592.1 |
| Bacteroidota | *Microcosmobacter mediterraneus* | WP_311426187.1 | WP_311427926.1 | WP_311426391.1 | WP_311425829.1 |
| Bacteroidota | *Microscilla marina* | WP_002705669.1 | WP_002698220.1 | WP_002693779.1 | WP_002700262.1 |
| Bacteroidota | *Mongoliitalea daihaiensis* | WP_236139018.1 | WP_236139112.1 | WP_236136828.1 | WP_236139309.1 |
| Bacteroidota | *Mongoliitalea lutea* | WP_189579359.1 | WP_189579276.1 | WP_189585048.1 | WP_189579197.1 |
| Bacteroidota | *Mucilaginibacter gracilis* | WP_121200601.1 | WP_121201357.1 | WP_121198186.1 | WP_121197576.1 |
| Bacteroidota | *Mucilaginibacter paludis* | WP_008510050.1 | WP_040627802.1 | WP_008511304.1 | WP_008505046.1 |
| Bacteroidota | *Mucilaginibacter sp.* | WP_348053712.1 | WP_348084747.1 | WP_369592252.1 | WP_348082828.1 |
| Bacteroidota | *Myroides fluvii* | WP_158963029.1 | WP_158960886.1 | WP_158963905.1 | WP_158962616.1 |
| Bacteroidota | *Myroides guanonis* | WP_090677389.1 | WP_090677506.1 | WP_090677655.1 | WP_090680943.1 |
| Bacteroidota | *Myroides injenensis* | WP_010256440.1 | WP_010254937.1 | WP_010256156.1 | WP_010255812.1 |
| Bacteroidota | *Myroides marinus* | WP_038986398.1 | WP_038988375.1 | WP_286425173.1 | WP_038985991.1 |
| Bacteroidota | *Myroides odoratimimus* | WP_286415851.1 | WP_025125853.1 | WP_286386351.1 | WP_286436824.1 |
| Bacteroidota | *Myroides odoratus* | WP_060873289.1 | WP_353101116.1 | WP_353101147.1 | WP_353118317.1 |
| Bacteroidota | *Myroides oncorhynchi* | WP_229946658.1 | WP_229948831.1 | WP_229948027.1 | WP_229946215.1 |
| Bacteroidota | *Myroides pelagicus* | WP_155036017.1 | WP_155035200.1 | WP_155036246.1 | WP_155035270.1 |
| Bacteroidota | *Myroides phaeus* | WP_090406831.1 | WP_090406041.1 | WP_090408418.1 | WP_090409129.1 |
| Bacteroidota | *Myroides sp. DW712* | WP_410877564.1 | WP_410879920.1 | WP_410877788.1 | WP_410881116.1 |
| Bacteroidota | *Nafulsella turpanensis* | WP_017730111.1 | WP_017731237.1 | WP_017729874.1 | WP_017731273.1 |
| Bacteroidota | *Namhaeicola litoreus* | WP_377179375.1 | WP_377178661.1 | WP_377178571.1 | WP_377178062.1 |
| Bacteroidota | *Nemorincola caseinilytica* | WP_345083699.1 | WP_345077030.1 | WP_345084461.1 | WP_345085155.1 |
| Bacteroidota | *Neotamlana laminarinivorans* | WP_226539851.1 | WP_226540228.1 | WP_226540714.1 | WP_226544469.1 |
| Bacteroidota | *Neotamlana nanhaiensis* | WP_044624813.1 | WP_044626297.1 | WP_044624653.1 | WP_044627344.1 |
| Bacteroidota | *Neotamlana sargassicola* | WP_226696502.1 | WP_226695741.1 | WP_226695825.1 | WP_226695554.1 |
| Bacteroidota | *Neotamlana sedimentorum* | WP_044633376.1 | WP_044632100.1 | WP_044632017.1 | WP_044633752.1 |
| Bacteroidota | *Neptunitalea chrysea* | WP_281755024.1 | WP_281752677.1 | WP_281751884.1 | WP_281754931.1 |
| Bacteroidota | *Neptunitalea lumnitzerae* | WP_281764548.1 | WP_281765818.1 | WP_281765697.1 | WP_281766268.1 |
| Bacteroidota | *Niastella caeni* | WP_243751633.1 | WP_243751588.1 | WP_136576828.1 | WP_136578394.1 |

| | | | | | |
|---|---|---|---|---|---|
| Bacteroidota | *Niastella koreensis* | WP_014220614.1 | WP_014223133.1 | WP_014220554.1 | WP_014218826.1 |
| Bacteroidota | *Niastella populi* | WP_081163821.1 | WP_081170820.1 | WP_081163880.1 | WP_081168632.1 |
| Bacteroidota | *Niastella sp.* | WP_407746754.1 | WP_407743925.1 | WP_407744206.1 | WP_407743428.1 |
| Bacteroidota | *Niastella vici* | WP_081147921.1 | WP_081151070.1 | WP_081147991.1 | WP_081155202.1 |
| Bacteroidota | *Niastella yeongjuensis* | WP_081203523.1 | WP_081199676.1 | WP_081203469.1 | WP_081202772.1 |
| Bacteroidota | *Nibribacter koreensis* | WP_345167043.1 | WP_345169332.1 | WP_345162107.1 | WP_345165865.1 |
| Bacteroidota | *Nibribacter ruber* | WP_160692106.1 | WP_160690431.1 | WP_160694538.1 | WP_160692977.1 |
| Bacteroidota | *Nonlabens agnitus* | WP_105981819.1 | WP_105982533.1 | WP_105981802.1 | WP_105981856.1 |
| Bacteroidota | *Nonlabens antarcticus* | WP_194852021.1 | WP_194850349.1 | WP_194852000.1 | WP_194850045.1 |
| Bacteroidota | *Nonlabens arenilitoris* | WP_170061736.1 | WP_105071680.1 | WP_245910827.1 | WP_105072145.1 |
| Bacteroidota | *Nonlabens dokdonensis* | WP_303686156.1 | WP_015363711.1 | WP_015360760.1 | WP_304017239.1 |
| Bacteroidota | *Nonlabens marinus* | WP_041497320.1 | WP_041497035.1 | WP_041494828.1 | WP_041497313.1 |
| Bacteroidota | *Nonlabens ponticola* | WP_241234638.1 | WP_126448044.1 | WP_126448506.1 | WP_126448453.1 |
| Bacteroidota | *Nonlabens sp.* | WP_292901267.1 | WP_213523056.1 | WP_415185567.1 | WP_348116523.1 |
| Bacteroidota | *Nonlabens spongiae* | WP_085765855.1 | WP_085766589.1 | WP_085765672.1 | WP_085766476.1 |
| Bacteroidota | *Nonlabens tegetincola* | WP_105018485.1 | WP_338350316.1 | WP_042278472.1 | WP_105018588.1 |
| Bacteroidota | *Nonlabens ulvanivorans* | WP_348378735.1 | WP_348376625.1 | WP_036585357.1 | WP_397300323.1 |
| Bacteroidota | *Nonlabens xiamenensis* | WP_124979799.1 | WP_124981425.1 | WP_124979846.1 | WP_240642324.1 |
| Bacteroidota | *Nubsella zeaxanthinifaciens* | WP_113639010.1 | WP_113637362.1 | WP_113635575.1 | WP_427531266.1 |
| Bacteroidota | *Oceanihabitans sediminis* | WP_113966091.1 | WP_072349045.1 | WP_072351629.1 | WP_318933800.1 |
| Bacteroidota | *Oceanihabitans sp. 2_MG-2023* | WP_303424503.1 | WP_303425751.1 | WP_303424129.1 | WP_303424880.1 |
| Bacteroidota | *Ochrovirga pacifica* | WP_010136408.1 | WP_010136703.1 | WP_010135397.1 | WP_010134725.1 |
| Bacteroidota | *Olivibacter sp. SDN3* | WP_187340080.1 | WP_187341938.1 | WP_187339661.1 | WP_187340539.1 |
| Bacteroidota | *Olleya aquimaris* | WP_111659532.1 | WP_111659383.1 | WP_111659072.1 | WP_111660549.1 |
| Bacteroidota | *Olleya marilimosa* | WP_339875171.1 | WP_028282215.1 | WP_099569000.1 | WP_191100015.1 |
| Bacteroidota | *Olleya namhaensis* | WP_090838608.1 | WP_281848004.1 | WP_272022623.1 | WP_281847549.1 |
| Bacteroidota | *Olleya sp. UBA1516* | WP_292949009.1 | WP_292945675.1 | WP_292949436.1 | WP_292943930.1 |
| Bacteroidota | *Owenweeksia hongkongensis* | WP_417592092.1 | WP_014201498.1 | WP_014201423.1 | WP_417601339.1 |

| | | | | | |
|---|---|---|---|---|---|
| Bacteroidota | *Paenimyroides aestuarii* | WP_257498959.1 | WP_257498498.1 | WP_257498436.1 | WP_257498655.1 |
| Bacteroidota | *Paenimyroides baculatum* | WP_150010046.1 | WP_150014147.1 | WP_150014239.1 | WP_150011125.1 |
| Bacteroidota | *Paenimyroides ceti* | WP_290363757.1 | WP_290362695.1 | WP_290363368.1 | WP_290363635.1 |
| Bacteroidota | *Paenimyroides marinum* | WP_091095447.1 | WP_091095377.1 | WP_091095373.1 | WP_091095038.1 |
| Bacteroidota | *Paenimyroides tangerinum* | WP_125019654.1 | WP_125018708.1 | WP_125017044.1 | WP_125017160.1 |
| Bacteroidota | *Paenimyroides ummariense* | WP_091519820.1 | WP_091522971.1 | WP_091526027.1 | WP_091521721.1 |
| Bacteroidota | *Paenimyroides viscosum* | WP_124898542.1 | WP_124899808.1 | WP_124898830.1 | WP_124898761.1 |
| Bacteroidota | *Panacibacter ginsenosidivorans* | WP_147187667.1 | WP_147189023.1 | WP_147192662.1 | WP_147188223.1 |
| Bacteroidota | *Panacibacter microcysteis* | WP_196991769.1 | WP_196990551.1 | WP_196991319.1 | WP_196991932.1 |
| Bacteroidota | *Paracrocinitomix mangrovi* | WP_221836281.1 | WP_221834572.1 | WP_221834850.1 | WP_221836018.1 |
| Bacteroidota | *Paradesertivirga mongoliensis* | WP_255903222.1 | WP_255897988.1 | WP_255902703.1 | WP_255902177.1 |
| Bacteroidota | *Parafilimonas terrae* | WP_090656488.1 | WP_090655776.1 | WP_245751513.1 | WP_090658732.1 |
| Bacteroidota | *Paraflavisolibacter caeni* | WP_279295253.1 | WP_279295855.1 | WP_279299632.1 | WP_279297706.1 |
| Bacteroidota | *Paraflavisolibacter sp. H34* | WP_336719945.1 | WP_336722650.1 | WP_336725373.1 | WP_336725333.1 |
| Bacteroidota | *Pararcticibacter amylolyticus* | WP_109416670.1 | WP_109416979.1 | WP_109414587.1 | WP_109416868.1 |
| Bacteroidota | *Parasediminibacterium paludis* | WP_379011984.1 | WP_379013407.1 | WP_379014846.1 | WP_379012157.1 |
| Bacteroidota | *Parasediminibacterium sp. JCM 36343* | WP_410846444.1 | WP_410844105.1 | WP_410844722.1 | WP_410847215.1 |
| Bacteroidota | *Parasegetibacter sp. NRK P23* | WP_251757843.1 | WP_251761400.1 | WP_251758786.1 | WP_251759009.1 |
| Bacteroidota | *Parvicella tangerina* | WP_258540467.1 | WP_258541382.1 | WP_258543270.1 | WP_258541479.1 |
| Bacteroidota | *Patiriisocius hiemis* | WP_311333773.1 | WP_311331756.1 | WP_311331402.1 | WP_311331573.1 |
| Bacteroidota | *Patiriisocius marinistellae* | WP_151894487.1 | WP_151894385.1 | WP_151894039.1 | WP_151894628.1 |
| Bacteroidota | *Patiriisocius marinus* | WP_151675022.1 | WP_151673679.1 | WP_151673105.1 | WP_151673038.1 |
| Bacteroidota | *Patiriisocius sp. Uisw_047* | WP_415371134.1 | WP_415371185.1 | WP_415370858.1 | WP_415370868.1 |
| Bacteroidota | *Paucihalobacter ruber* | WP_140990968.1 | WP_140990425.1 | WP_140990909.1 | WP_140990640.1 |
| Bacteroidota | *Paucihalobacter sp.* | WP_334111456.1 | WP_334112096.1 | WP_334113206.1 | WP_334111572.1 |
| Bacteroidota | *Pedobacter africanus* | WP_310140594.1 | WP_310142609.1 | WP_310141383.1 | WP_084237520.1 |
| Bacteroidota | *Pedobacter agri* | WP_116167521.1 | WP_316805750.1 | WP_316806296.1 | WP_316808301.1 |
| Bacteroidota | *Pedobacter albus* | WP_330106040.1 | WP_330109376.1 | WP_330105924.1 | WP_330107668.1 |

| | | | | | |
|---|---|---|---|---|---|
| Bacteroidota | *Pedobacter alluvionis* | WP_121286791.1 | WP_121287746.1 | WP_121287178.1 | WP_121283637.1 |
| Bacteroidota | *Pedobacter alpinus* | WP_379040248.1 | WP_379048161.1 | WP_379040262.1 | WP_379047338.1 |
| Bacteroidota | *Pedobacter antarcticus* | WP_037439044.1 | WP_219224099.1 | WP_037442773.1 | WP_316744585.1 |
| Bacteroidota | *Pedobacter aquae* | WP_149073860.1 | WP_149074866.1 | WP_149073748.1 | WP_149074685.1 |
| Bacteroidota | *Pedobacter aquatilis* | WP_316736527.1 | WP_316734886.1 | WP_316738179.1 | WP_316756619.1 |
| Bacteroidota | *Pedobacter arcticus* | WP_017258694.1 | WP_017256750.1 | WP_017259123.1 | WP_040539760.1 |
| Bacteroidota | *Pedobacter borealis* | WP_029275995.1 | WP_029280013.1 | WP_029278913.1 | WP_029276083.1 |
| Bacteroidota | *Pedobacter boryungensis* | WP_173270125.1 | WP_173272792.1 | WP_173273727.1 | WP_173273906.1 |
| Bacteroidota | *Pedobacter caeni* | WP_073233220.1 | WP_073232208.1 | WP_073234076.1 | WP_073229707.1 |
| Bacteroidota | *Pedobacter changchengzhani* | WP_133262608.1 | WP_133263516.1 | WP_133262326.1 | WP_133261648.1 |
| Bacteroidota | *Pedobacter chinensis* | WP_115403602.1 | WP_115402615.1 | WP_115404077.1 | WP_115404238.1 |
| Bacteroidota | *Pedobacter chitinilyticus* | WP_113648656.1 | WP_113648051.1 | WP_113645262.1 | WP_113647052.1 |
| Bacteroidota | *Pedobacter cryoconitis* | WP_111632184.1 | WP_184624986.1 | WP_068402913.1 | WP_183885582.1 |
| Bacteroidota | *Pedobacter cryophilus* | WP_136827635.1 | WP_136825249.1 | WP_136826977.1 | WP_317130548.1 |
| Bacteroidota | *Pedobacter cryotolerans* | WP_136877078.1 | WP_136874354.1 | WP_136876996.1 | WP_136878258.1 |
| Bacteroidota | *Pedobacter duraquae* | WP_133553010.1 | WP_133556412.1 | WP_133553364.1 | WP_133555384.1 |
| Bacteroidota | *Pedobacter endophyticus* | WP_196098957.1 | WP_196100950.1 | WP_196099093.1 | WP_196100596.1 |
| Bacteroidota | *Pedobacter fastidiosus* | WP_187071800.1 | WP_187069658.1 | WP_187071857.1 | WP_187070164.1 |
| Bacteroidota | *Pedobacter flavus* | WP_330146138.1 | WP_330144997.1 | WP_330145864.1 | WP_330145931.1 |
| Bacteroidota | *Pedobacter foliorum* | WP_173091633.1 | WP_173090079.1 | WP_173093305.1 | WP_173091865.1 |
| Bacteroidota | *Pedobacter frigidisoli* | WP_131557085.1 | WP_316798605.1 | WP_131561206.1 | WP_316800050.1 |
| Bacteroidota | *Pedobacter frigiditerrae* | WP_131554366.1 | WP_131552253.1 | WP_316768507.1 | WP_131555756.1 |
| Bacteroidota | *Pedobacter frigoris* | WP_316794001.1 | WP_316793455.1 | WP_316793880.1 | WP_136837743.1 |
| Bacteroidota | *Pedobacter gandavensis* | WP_281029962.1 | WP_316746284.1 | WP_316747424.1 | WP_316840536.1 |
| Bacteroidota | *Pedobacter ghigonis* | WP_175634016.1 | WP_175636134.1 | WP_175633854.1 | WP_175636440.1 |
| Bacteroidota | *Pedobacter ginsengisoli* | WP_099440970.1 | WP_285059228.1 | WP_285056991.1 | WP_285054377.1 |
| Bacteroidota | *Pedobacter glucosidilyticus* | WP_026904798.1 | WP_026905129.1 | WP_304064693.1 | WP_026902661.1 |
| Bacteroidota | *Pedobacter hartonius* | WP_090554705.1 | WP_090555335.1 | WP_090555089.1 | WP_090556570.1 |

| | | | | | |
|---|---|---|---|---|---|
| Bacteroidota | *Pedobacter helvus* | WP_138731280.1 | WP_138729967.1 | WP_138730109.1 | WP_138728912.1 |
| Bacteroidota | *Pedobacter heparinus* | WP_316810839.1 | WP_316813681.1 | WP_012781370.1 | WP_015809326.1 |
| Bacteroidota | *Pedobacter hiemivivus* | WP_131611192.1 | WP_136880229.1 | WP_136879996.1 | WP_131607297.1 |
| Bacteroidota | *Pedobacter insulae* | WP_090992148.1 | WP_090996882.1 | WP_090992110.1 | WP_090996648.1 |
| Bacteroidota | *Pedobacter jamesrossensis* | WP_378960642.1 | WP_378962165.1 | WP_378959945.1 | WP_378962518.1 |
| Bacteroidota | *Pedobacter jejuensis* | WP_123205183.1 | WP_123204999.1 | WP_123205102.1 | WP_123206726.1 |
| Bacteroidota | *Pedobacter jeongneungensis* | WP_025143861.1 | WP_025144181.1 | WP_025143672.1 | WP_025145791.1 |
| Bacteroidota | *Pedobacter kyungheensis* | WP_039480112.1 | WP_039479543.1 | WP_039479903.1 | WP_039479944.1 |
| Bacteroidota | *Pedobacter lithocola* | WP_378981640.1 | WP_378987507.1 | WP_378981496.1 | WP_378984310.1 |
| Bacteroidota | *Pedobacter lusitanus* | WP_041885766.1 | WP_041883719.1 | WP_041885191.1 | WP_041887252.1 |
| Bacteroidota | *Pedobacter mendelii* | WP_188412606.1 | WP_188411375.1 | WP_378957843.1 | WP_188415496.1 |
| Bacteroidota | *Pedobacter metabolipauper* | WP_133577838.1 | WP_133577613.1 | WP_133577989.1 | WP_133577171.1 |
| Bacteroidota | *Pedobacter miscanthi* | WP_113947079.1 | WP_316824526.1 | WP_113952201.1 | WP_113950981.1 |
| Bacteroidota | *Pedobacter montanisoli* | WP_243361101.1 | WP_243360292.1 | WP_243363041.1 | WP_243360692.1 |
| Bacteroidota | *Pedobacter mucosus* | WP_238416438.1 | WP_238414581.1 | WP_238414089.1 | WP_238414324.1 |
| Bacteroidota | *Pedobacter namyangjuensis* | WP_113654792.1 | WP_113654293.1 | WP_113653838.1 | WP_113654129.1 |
| Bacteroidota | *Pedobacter nanyangensis* | WP_113663099.1 | WP_113662147.1 | WP_113662675.1 | WP_113661786.1 |
| Bacteroidota | *Pedobacter nototheniae* | WP_316802806.1 | WP_131539222.1 | WP_316805533.1 | WP_316804932.1 |
| Bacteroidota | *Pedobacter nutrimenti* | WP_316833367.1 | WP_110833957.1 | WP_110834448.1 | WP_316836260.1 |
| Bacteroidota | *Pedobacter nyackensis* | WP_084287073.1 | WP_316816358.1 | WP_316815893.1 | WP_316818807.1 |
| Bacteroidota | *Pedobacter paludis* | WP_109930234.1 | WP_109930788.1 | WP_109932520.1 | WP_109931200.1 |
| Bacteroidota | *Pedobacter petrophilus* | WP_154279375.1 | WP_154281502.1 | WP_154282467.1 | WP_154278704.1 |
| Bacteroidota | *Pedobacter planticolens* | WP_182922054.1 | WP_182922361.1 | WP_182922156.1 | WP_182920612.1 |
| Bacteroidota | *Pedobacter polaris* | WP_136841322.1 | WP_136838023.1 | WP_136844270.1 | WP_136841447.1 |
| Bacteroidota | *Pedobacter psychrodurus* | WP_131532023.1 | WP_131533002.1 | WP_316846007.1 | WP_131530597.1 |
| Bacteroidota | *Pedobacter psychrophilus* | WP_068821378.1 | WP_068822602.1 | WP_068823604.1 | WP_068822891.1 |
| Bacteroidota | *Pedobacter psychroterrae* | WP_131596710.1 | WP_131594357.1 | WP_131597463.1 | WP_131597727.1 |
| Bacteroidota | *Pedobacter psychrotolerans* | WP_132535531.1 | WP_132530194.1 | WP_132536433.1 | WP_132529721.1 |

| | | | | | |
|---|---|---|---|---|---|
| Bacteroidota | *Pedobacter punctiformis* | WP_269427941.1 | WP_269428406.1 | WP_269427832.1 | WP_269428613.1 |
| Bacteroidota | *Pedobacter puniceum* | WP_154286206.1 | WP_154288091.1 | WP_154286091.1 | WP_154288349.1 |
| Bacteroidota | *Pedobacter quisquiliarum* | WP_188627291.1 | WP_188626080.1 | WP_188627188.1 | WP_188624811.1 |
| Bacteroidota | *Pedobacter rhizosphaerae* | WP_090880663.1 | WP_090884952.1 | WP_090881040.1 | WP_090883636.1 |
| Bacteroidota | *Pedobacter rhodius* | WP_269414017.1 | WP_269417121.1 | WP_269413907.1 | WP_269416549.1 |
| Bacteroidota | *Pedobacter roseus* | WP_187591070.1 | WP_187592982.1 | WP_187591282.1 | WP_187591487.1 |
| Bacteroidota | *Pedobacter sandarakinus* | WP_265856552.1 | WP_265854285.1 | WP_265855752.1 | WP_265854120.1 |
| Bacteroidota | *Pedobacter schmidteae* | WP_121273509.1 | WP_121269434.1 | WP_121273731.1 | WP_121271022.1 |
| Bacteroidota | *Pedobacter segetis* | WP_200584548.1 | WP_200587972.1 | WP_200585255.1 | WP_200586207.1 |
| Bacteroidota | *Pedobacter soli* | WP_090771268.1 | WP_090772814.1 | WP_090772601.1 | WP_090766432.1 |
| Bacteroidota | *Pedobacter sp.* | WP_353903522.1 | WP_324370905.1 | WP_421942790.1 | WP_424057527.1 |
| Bacteroidota | *Pedobacter steynii* | WP_069382250.1 | WP_069379562.1 | WP_074606995.1 | WP_074611952.1 |
| Bacteroidota | *Pedobacter suwonensis* | WP_090987543.1 | WP_293742548.1 | WP_293741410.1 | WP_090984736.1 |
| Bacteroidota | *Pedobacter terrae* | WP_090499916.1 | WP_335317504.1 | WP_335320483.1 | WP_090498603.1 |
| Bacteroidota | *Pedobacter ureilyticus* | WP_138724127.1 | WP_138722568.1 | WP_138723946.1 | WP_138721851.1 |
| Bacteroidota | *Pedobacter vanadiisoli* | WP_379080426.1 | WP_379081851.1 | WP_379080222.1 | WP_379081366.1 |
| Bacteroidota | *Pedobacter westerhofensis* | WP_142529649.1 | WP_142528417.1 | WP_142529467.1 | WP_142530983.1 |
| Bacteroidota | *Pedobacter xixiisoli* | WP_097131290.1 | WP_097132640.1 | WP_097127343.1 | WP_097132993.1 |
| Bacteroidota | *Pedobacter yonginense* | WP_109927202.1 | WP_109925110.1 | WP_109927344.1 | WP_109926179.1 |
| Bacteroidota | *Pedobacter yulinensis* | WP_107217006.1 | WP_107214288.1 | WP_107216326.1 | WP_107216633.1 |
| Bacteroidota | *Pedobacter zeae* | WP_183759580.1 | WP_183765082.1 | WP_183759401.1 | WP_183768407.1 |
| Bacteroidota | *Pelobium manganitolerans* | WP_120181569.1 | WP_120181251.1 | WP_120181798.1 | WP_120181363.1 |
| Bacteroidota | *Penaeicola halotolerans* | WP_226390012.1 | WP_226390126.1 | WP_226389474.1 | WP_226390042.1 |
| Bacteroidota | *Persicobacter diffluens* | WP_338235790.1 | WP_338237395.1 | WP_338236170.1 | WP_338235858.1 |
| Bacteroidota | *Persicobacter psychrovividus* | WP_332920859.1 | WP_332922396.1 | WP_338397812.1 | WP_338397565.1 |
| Bacteroidota | *Persicobacter sp. CCB-QB2* | WP_053406022.1 | WP_053406831.1 | WP_053404325.1 | WP_053405934.1 |
| Bacteroidota | *Phaeodactylibacter luteus* | WP_147165650.1 | WP_147168864.1 | WP_147166612.1 | WP_147167063.1 |
| Bacteroidota | *Phaeodactylibacter sp.* | WP_350178551.1 | WP_293565027.1 | WP_293566465.1 | WP_350177613.1 |

| | | | | | |
|---|---|---|---|---|---|
| Bacteroidota | *Phaeodactylibacter xiamenensis* | WP_044226137.1 | WP_044220771.1 | WP_425423020.1 | WP_044226419.1 |
| Bacteroidota | *Phnomibacter sp.* | WP_333798807.1 | WP_333801136.1 | WP_333798932.1 | WP_333799269.1 |
| Bacteroidota | *Pinibacter aurantiacus* | WP_217793815.1 | WP_217789871.1 | WP_217794396.1 | WP_217791097.1 |
| Bacteroidota | *Pinibacter soli* | WP_282335857.1 | WP_282334310.1 | WP_282334518.1 | WP_282335433.1 |
| Bacteroidota | *Planktosalinus lacus* | WP_188441290.1 | WP_188441418.1 | WP_188443026.1 | WP_188439440.1 |
| Bacteroidota | *Planobacterium oryzisoli* | WP_194738628.1 | WP_194739118.1 | WP_194739626.1 | WP_194738760.1 |
| Bacteroidota | *Polaribacter aestuariivivens* | WP_138535047.1 | WP_138534867.1 | WP_138535111.1 | WP_138537235.1 |
| Bacteroidota | *Polaribacter aquimarinus* | WP_235869146.1 | WP_109404621.1 | WP_109405612.1 | WP_109404950.1 |
| Bacteroidota | *Polaribacter atrinae* | WP_282073704.1 | WP_341220606.1 | WP_341222179.1 | WP_282073884.1 |
| Bacteroidota | *Polaribacter batillariae* | WP_207971760.1 | WP_207972030.1 | WP_207972378.1 | WP_207971661.1 |
| Bacteroidota | *Polaribacter butkevichii* | WP_105048181.1 | WP_105048550.1 | WP_105048114.1 | WP_105048036.1 |
| Bacteroidota | *Polaribacter cellanae* | WP_208079091.1 | WP_208079180.1 | WP_208079597.1 | WP_208079626.1 |
| Bacteroidota | *Polaribacter dokdonensis* | WP_053974795.1 | WP_053974708.1 | WP_053974658.1 | WP_053975070.1 |
| Bacteroidota | *Polaribacter filamentus* | WP_104808833.1 | WP_104809256.1 | WP_104808805.1 | WP_104808646.1 |
| Bacteroidota | *Polaribacter gangjinensis* | WP_105045698.1 | WP_105045548.1 | WP_105047271.1 | WP_105047223.1 |
| Bacteroidota | *Polaribacter glomeratus* | WP_105021496.1 | WP_105020554.1 | WP_105020177.1 | WP_105020891.1 |
| Bacteroidota | *Polaribacter haliotis* | WP_088353370.1 | WP_088353541.1 | WP_088353291.1 | WP_088352880.1 |
| Bacteroidota | *Polaribacter huanghezhanensis* | WP_301400178.1 | WP_301400243.1 | WP_301400130.1 | WP_301400796.1 |
| Bacteroidota | *Polaribacter irgensii* | WP_004570937.1 | WP_026288371.1 | WP_018942509.1 | WP_004570425.1 |
| Bacteroidota | *Polaribacter litorisediminis* | WP_223447258.1 | WP_223446044.1 | WP_223446128.1 | WP_223439388.1 |
| Bacteroidota | *Polaribacter marinivivus* | WP_377410462.1 | WP_377410213.1 | WP_377410502.1 | WP_377409873.1 |
| Bacteroidota | *Polaribacter marinus* | WP_242177528.1 | WP_242177238.1 | WP_242177468.1 | WP_242176929.1 |
| Bacteroidota | *Polaribacter pacificus* | WP_229664930.1 | WP_188598849.1 | WP_188598986.1 | WP_188599336.1 |
| Bacteroidota | *Polaribacter pectinis* | WP_187483182.1 | WP_187483341.1 | WP_187483380.1 | WP_187482972.1 |
| Bacteroidota | *Polaribacter ponticola* | WP_265724611.1 | WP_265724554.1 | WP_265724156.1 | WP_274270124.1 |
| Bacteroidota | *Polaribacter porphyrae* | WP_105014403.1 | WP_105014864.1 | WP_105014974.1 | WP_105015495.1 |
| Bacteroidota | *Polaribacter reichenbachii* | WP_068360296.1 | WP_068360224.1 | WP_068360040.1 | WP_068364870.1 |
| Bacteroidota | *Polaribacter sejongensis* | WP_261974070.1 | WP_208889400.1 | WP_261973620.1 | WP_373940948.1 |

| | | | | | |
|---|---|---|---|---|---|
| Bacteroidota | *Polaribacter septentrionalilitoris* | WP_159947717.1 | WP_159947352.1 | WP_159947839.1 | WP_159948279.1 |
| Bacteroidota | *Polaribacter sp.* | WP_348024623.1 | WP_296637153.1 | WP_348023849.1 | WP_404486996.1 |
| Bacteroidota | *Polaribacter staleyi* | WP_343329531.1 | WP_343329764.1 | WP_343329481.1 | WP_343329414.1 |
| Bacteroidota | *Polaribacter tangerinus* | WP_088322715.1 | WP_088322902.1 | WP_088322791.1 | WP_088322699.1 |
| Bacteroidota | *Polaribacter vadi* | WP_065319297.1 | WP_065318254.1 | WP_065318000.1 | WP_065320679.1 |
| Bacteroidota | *Polluticaenibacter yanchengensis* | WP_407031991.1 | WP_407031566.1 | WP_407030962.1 | WP_407032955.1 |
| Bacteroidota | *Polluticoccus soli* | WP_276134737.1 | WP_276134619.1 | WP_276134772.1 | WP_276132203.1 |
| Bacteroidota | *Pontibacter actiniarum* | WP_025608587.1 | WP_025607855.1 | WP_025605102.1 | WP_084196269.1 |
| Bacteroidota | *Pontibacter akesuensis* | WP_068837678.1 | WP_068837038.1 | WP_068839774.1 | WP_082815155.1 |
| Bacteroidota | *Pontibacter amylolyticus* | WP_188500845.1 | WP_188502506.1 | WP_188499622.1 | WP_188500582.1 |
| Bacteroidota | *Pontibacter anaerobius* | WP_266052702.1 | WP_266051738.1 | WP_266054022.1 | WP_266054192.1 |
| Bacteroidota | *Pontibacter aquaedesilientis* | WP_191183522.1 | WP_191182936.1 | WP_191182164.1 | WP_224744371.1 |
| Bacteroidota | *Pontibacter arcticus* | WP_112304594.1 | WP_112306243.1 | WP_112305500.1 | WP_239020921.1 |
| Bacteroidota | *Pontibacter aydingkolensis* | WP_219876992.1 | WP_219876295.1 | WP_219878462.1 | WP_246596896.1 |
| Bacteroidota | *Pontibacter beigongshangensis* | WP_187262260.1 | WP_187260967.1 | WP_187260582.1 | WP_187264052.1 |
| Bacteroidota | *Pontibacter burrus* | WP_163911654.1 | WP_163915162.1 | WP_163914727.1 | WP_163910788.1 |
| Bacteroidota | *Pontibacter cellulosilyticus* | WP_187068082.1 | WP_187067611.1 | WP_187065817.1 | WP_230407259.1 |
| Bacteroidota | *Pontibacter chinhatensis* | WP_092100996.1 | WP_092099925.1 | WP_092098427.1 | WP_092101233.1 |
| Bacteroidota | *Pontibacter chitinilyticus* | WP_347157579.1 | WP_347159781.1 | WP_347157039.1 | WP_347157394.1 |
| Bacteroidota | *Pontibacter diazotrophicus* | WP_115566407.1 | WP_115563627.1 | WP_115565944.1 | WP_246000815.1 |
| Bacteroidota | *Pontibacter fetidus* | WP_162344978.1 | WP_162345166.1 | WP_162344390.1 | WP_162346564.1 |
| Bacteroidota | *Pontibacter flavimaris* | WP_073851502.1 | WP_170866119.1 | WP_073851109.1 | WP_073851885.1 |
| Bacteroidota | *Pontibacter harenae* | WP_229967464.1 | WP_229972779.1 | WP_229969948.1 | WP_229968909.1 |
| Bacteroidota | *Pontibacter indicus* | WP_076668258.1 | WP_170871914.1 | WP_076666330.1 | WP_244554682.1 |
| Bacteroidota | *Pontibacter kalidii* | WP_266203549.1 | WP_266202819.1 | WP_266205387.1 | WP_266203411.1 |
| Bacteroidota | *Pontibacter korlensis* | WP_046309691.1 | WP_046308896.1 | WP_046312400.1 | WP_084694729.1 |
| Bacteroidota | *Pontibacter liquoris* | WP_367615719.1 | WP_242916368.1 | WP_242918685.1 | WP_242921285.1 |
| Bacteroidota | *Pontibacter litorisediminis* | WP_276498473.1 | WP_276499133.1 | WP_276495720.1 | WP_276498631.1 |

| | | | | | |
|---|---|---|---|---|---|
| Bacteroidota | *Pontibacter locisalis* | WP_377506203.1 | WP_377507272.1 | WP_377512209.1 | WP_377503573.1 |
| Bacteroidota | *Pontibacter mangrovi* | WP_140620258.1 | WP_140621972.1 | WP_140618665.1 | WP_239022830.1 |
| Bacteroidota | *Pontibacter mucosus* | WP_108210053.1 | WP_108212430.1 | WP_108213444.1 | WP_108210166.1 |
| Bacteroidota | *Pontibacter oryzae* | WP_119432228.1 | WP_119430834.1 | WP_119431269.1 | WP_119432100.1 |
| Bacteroidota | *Pontibacter pamirensis* | WP_162054115.1 | WP_162053720.1 | WP_162052484.1 | WP_237144869.1 |
| Bacteroidota | *Pontibacter populi* | WP_350413596.1 | WP_350410730.1 | WP_350412632.1 | WP_350411634.1 |
| Bacteroidota | *Pontibacter pudoricolor* | WP_162428158.1 | WP_162428816.1 | WP_162426266.1 | WP_162427917.1 |
| Bacteroidota | *Pontibacter qinzhouensis* | WP_147920392.1 | WP_147922611.1 | WP_147923662.1 | WP_255474196.1 |
| Bacteroidota | *Pontibacter ramchanderi* | WP_101444762.1 | WP_101446093.1 | WP_101442876.1 | WP_180336380.1 |
| Bacteroidota | *Pontibacter roseus* | WP_018476290.1 | WP_211214358.1 | WP_018478075.1 | WP_018476588.1 |
| Bacteroidota | *Pontibacter ruber* | WP_250428584.1 | WP_250431823.1 | WP_250429605.1 | WP_250428291.1 |
| Bacteroidota | *Pontibacter rugosus* | WP_377521968.1 | WP_377526565.1 | WP_377529224.1 | WP_377526710.1 |
| Bacteroidota | *Pontibacter saemangeumensis* | WP_345160897.1 | WP_345161860.1 | WP_345157442.1 | WP_345159741.1 |
| Bacteroidota | *Pontibacter sp. H259* | WP_340327220.1 | WP_340325105.1 | WP_340324301.1 | WP_340326631.1 |
| Bacteroidota | *Pontibacter toksunensis* | WP_377482369.1 | WP_377487128.1 | WP_377479545.1 | WP_377485187.1 |
| Bacteroidota | *Pontibacter ummariensis* | WP_089318033.1 | WP_089319534.1 | WP_089317724.1 | WP_089319831.1 |
| Bacteroidota | *Pontibacter virosus* | WP_116543722.1 | WP_207774867.1 | WP_116543526.1 | WP_116541785.1 |
| Bacteroidota | *Pontibacter vulgaris* | WP_242926206.1 | WP_242927759.1 | WP_242928228.1 | WP_242926450.1 |
| Bacteroidota | *Pontimicrobium aquaticum* | WP_136844426.1 | WP_136844924.1 | WP_136842894.1 | WP_136842336.1 |
| Bacteroidota | *Pontimicrobium sp. IMCC45349* | WP_412984746.1 | WP_412984909.1 | WP_412985655.1 | WP_412984315.1 |
| Bacteroidota | *Porifericola rhodea* | WP_302236952.1 | WP_302241969.1 | WP_302238587.1 | WP_302241977.1 |
| Bacteroidota | *Postechiella marina* | WP_344788461.1 | WP_344786197.1 | WP_344786381.1 | WP_344789558.1 |
| Bacteroidota | *Pseudalgibacter alginicilyticus* | WP_054723694.1 | WP_054724859.1 | WP_054724689.1 | WP_054725144.1 |
| Bacteroidota | *Pseudarcicella hirudinis* | WP_092019125.1 | WP_092018740.1 | WP_092016199.1 | WP_092018609.1 |
| Bacteroidota | *Pseudobacter ginsenosidimutans* | WP_130543858.1 | WP_130542873.1 | WP_130543804.1 | WP_130540330.1 |
| Bacteroidota | *Pseudoflavitalea rhizosphaerae* | WP_127132024.1 | WP_127128830.1 | WP_127132125.1 | WP_127125031.1 |
| Bacteroidota | *Pseudofulvibacter geojedonensis* | WP_377714822.1 | WP_377715660.1 | WP_377712568.1 | WP_377713360.1 |
| Bacteroidota | *Pseudopedobacter beijingensis* | WP_379663189.1 | WP_379661639.1 | WP_379663639.1 | WP_379660747.1 |

| | | | | | |
|---|---|---|---|---|---|
| Bacteroidota | *Pseudopedobacter saltans* | WP_013633114.1 | WP_013634471.1 | WP_013633057.1 | WP_013631349.1 |
| Bacteroidota | *Pseudopedobacter sp.* | WP_353133445.1 | WP_353133078.1 | WP_353136078.1 | WP_353135823.1 |
| Bacteroidota | *Pseudotamlana agarivorans* | WP_216068594.1 | WP_067147999.1 | WP_216068537.1 | WP_067145892.1 |
| Bacteroidota | *Pseudotamlana carrageenivorans* | WP_102996449.1 | WP_102996535.1 | WP_102996350.1 | WP_102996863.1 |
| Bacteroidota | *Pseudotamlana haliotis* | WP_150935729.1 | WP_150935676.1 | WP_150937802.1 | WP_150940490.1 |
| Bacteroidota | *Pseudotenacibaculum haliotis* | WP_379666458.1 | WP_379666268.1 | WP_379666551.1 | WP_379664548.1 |
| Bacteroidota | *Pseudotenacibaculum sp. MALMAid0570* | WP_349553689.1 | WP_349553908.1 | WP_349553590.1 | WP_349554108.1 |
| Bacteroidota | *Psychroflexus maritimus* | WP_166400925.1 | WP_166400947.1 | WP_166400890.1 | WP_166399151.1 |
| Bacteroidota | *Psychroflexus salarius* | WP_073193621.1 | WP_073191291.1 | WP_073190977.1 | WP_073193286.1 |
| Bacteroidota | *Psychroflexus sp. ALD_RP9* | WP_207038279.1 | WP_207038686.1 | WP_207038581.1 | WP_207037867.1 |
| Bacteroidota | *Psychroserpens algicola* | WP_204345880.1 | WP_248413321.1 | WP_248411954.1 | WP_204346159.1 |
| Bacteroidota | *Psychroserpens burtonensis* | WP_147231107.1 | WP_028870830.1 | WP_028872699.1 | WP_028872803.1 |
| Bacteroidota | *Psychroserpens damuponensis* | WP_040282237.1 | WP_040279120.1 | WP_040280953.1 | WP_040281111.1 |
| Bacteroidota | *Psychroserpens jangbogonensis* | WP_052172542.1 | WP_033959147.1 | WP_033959080.1 | WP_033956243.1 |
| Bacteroidota | *Psychroserpens luteolus* | WP_230935730.1 | WP_230935135.1 | WP_230935982.1 | WP_230936893.1 |
| Bacteroidota | *Psychroserpens luteus* | WP_194507887.1 | WP_194507318.1 | WP_194507631.1 | WP_194508324.1 |
| Bacteroidota | *Psychroserpens mesophilus* | WP_040250502.1 | WP_040251637.1 | WP_174435320.1 | WP_040250904.1 |
| Bacteroidota | *Psychroserpens ponticola* | WP_249995719.1 | WP_249995923.1 | WP_249995559.1 | WP_249995438.1 |
| Bacteroidota | *Psychroserpens sp.* | WP_425223521.1 | WP_323789418.1 | WP_335728066.1 | WP_415224132.1 |
| Bacteroidota | *Puia dinghuensis* | WP_188929662.1 | WP_188937963.1 | WP_188929879.1 | WP_188927805.1 |
| Bacteroidota | *Puia sp.* | WP_325593104.1 | WP_331581891.1 | WP_325587851.1 | WP_325588515.1 |
| Bacteroidota | *Pustulibacterium marinum* | WP_093023466.1 | WP_093026409.1 | WP_093025214.1 | WP_093023162.1 |
| Bacteroidota | *Putridiphycobacter roseus* | WP_111063168.1 | WP_111062192.1 | WP_111062136.1 | WP_111061356.1 |
| Bacteroidota | *Raineya orbicola* | WP_101358011.1 | WP_101359701.1 | WP_101357744.1 | WP_165778063.1 |
| Bacteroidota | *Rapidithrix thailandica* | WP_346820272.1 | WP_346820230.1 | WP_346819169.1 | WP_346819304.1 |
| Bacteroidota | *Rasiella rasia* | WP_164678329.1 | WP_164678373.1 | WP_164678848.1 | WP_164679013.1 |
| Bacteroidota | *Reichenbachiella agariperforans* | WP_216011801.1 | WP_073124535.1 | WP_073123459.1 | WP_073122811.1 |
| Bacteroidota | *Reichenbachiella agarivorans* | WP_262310620.1 | WP_262309824.1 | WP_262308525.1 | WP_262310351.1 |

| | | | | | |
|---|---|---|---|---|---|
| Bacteroidota | *Reichenbachiella carrageenanivorans* | WP_263052273.1 | WP_263050107.1 | WP_263049587.1 | WP_263051374.1 |
| Bacteroidota | *Reichenbachiella faecimaris* | WP_084372697.1 | WP_084371036.1 | WP_084371183.1 | WP_084371882.1 |
| Bacteroidota | *Reichenbachiella sp.* | WP_318169581.1 | WP_348120269.1 | WP_348463629.1 | WP_422361535.1 |
| Bacteroidota | *Reichenbachiella ulvae* | WP_264138846.1 | WP_264137898.1 | WP_264137298.1 | WP_264138500.1 |
| Bacteroidota | *Reichenbachiella versicolor* | WP_109832983.1 | WP_109829597.1 | WP_109831061.1 | WP_109832016.1 |
| Bacteroidota | *Rhodoflexus caldus* | WP_250632542.1 | WP_250631486.1 | WP_250632666.1 | WP_250629984.1 |
| Bacteroidota | *Robertkochia marina* | WP_136336582.1 | WP_136336830.1 | WP_136335742.1 | WP_136336997.1 |
| Bacteroidota | *Robertkochia sediminum* | WP_203058860.1 | WP_203061522.1 | WP_203060466.1 | WP_203059512.1 |
| Bacteroidota | *Robertkochia sp. 3YJGBD-33* | WP_224484802.1 | WP_224483128.1 | WP_224484353.1 | WP_224484607.1 |
| Bacteroidota | *Roseivirga sp. BDSF3-8* | WP_371908952.1 | WP_371908042.1 | WP_371910304.1 | WP_371908816.1 |
| Bacteroidota | *Rufibacter aurantiacus* | WP_246853757.1 | WP_210487332.1 | WP_210488992.1 | WP_210488416.1 |
| Bacteroidota | *Rufibacter glacialis* | WP_149096627.1 | WP_149097938.1 | WP_149098143.1 | WP_149098844.1 |
| Bacteroidota | *Rufibacter hautae* | WP_317132021.1 | WP_149088950.1 | WP_149091153.1 | WP_149089626.1 |
| Bacteroidota | *Rufibacter immobilis* | WP_377060888.1 | WP_377062404.1 | WP_123134153.1 | WP_123133193.1 |
| Bacteroidota | *Rufibacter latericius* | WP_123128600.1 | WP_123125038.1 | WP_123125940.1 | WP_123126617.1 |
| Bacteroidota | *Rufibacter psychrotolerans* | WP_205501396.1 | WP_205502872.1 | WP_205500681.1 | WP_205500297.1 |
| Bacteroidota | *Rufibacter radiotolerans* | WP_048922194.1 | WP_048921188.1 | WP_048920482.1 | WP_048921794.1 |
| Bacteroidota | *Rufibacter roseolus* | WP_246849904.1 | WP_210463819.1 | WP_210464973.1 | WP_210463158.1 |
| Bacteroidota | *Rufibacter roseus* | WP_066620381.1 | WP_066618735.1 | WP_066621324.1 | WP_066623342.1 |
| Bacteroidota | *Rufibacter sediminis* | WP_186635039.1 | WP_186640101.1 | WP_186636070.1 | WP_186639587.1 |
| Bacteroidota | *Rufibacter sp. LB8* | WP_192822645.1 | WP_192820106.1 | WP_192820940.1 | WP_192822191.1 |
| Bacteroidota | *Rufibacter tibetensis* | WP_062543512.1 | WP_062542449.1 | WP_062545229.1 | WP_062543027.1 |
| Bacteroidota | *Rurimicrobium arvi* | WP_344824269.1 | WP_344826516.1 | WP_344828551.1 | WP_344822986.1 |
| Bacteroidota | *Sabulibacter ruber* | WP_207433634.1 | WP_207435692.1 | WP_207430476.1 | WP_207434099.1 |
| Bacteroidota | *Salibacter sp.* | WP_310682266.1 | WP_310667847.1 | WP_310667026.1 | WP_310667442.1 |
| Bacteroidota | *Salinimicrobium catena* | WP_093112977.1 | WP_093112685.1 | WP_334213357.1 | WP_334214643.1 |
| Bacteroidota | *Salinimicrobium flavum* | WP_380753729.1 | WP_380752956.1 | WP_380747343.1 | WP_380748127.1 |
| Bacteroidota | *Salinimicrobium gaetbulicola* | WP_380736476.1 | WP_380736752.1 | WP_380736910.1 | WP_380737270.1 |

| | | | | | |
|---|---|---|---|---|---|
| Bacteroidota | *Salinimicrobium marinum* | WP_189606297.1 | WP_189603621.1 | WP_189604486.1 | WP_189604708.1 |
| Bacteroidota | *Salinimicrobium oceani* | WP_168139201.1 | WP_168138491.1 | WP_168136558.1 | WP_168136841.1 |
| Bacteroidota | *Salinimicrobium profundisediminis* | WP_266068498.1 | WP_266068280.1 | WP_266070952.1 | WP_266070336.1 |
| Bacteroidota | *Salinimicrobium sediminis* | WP_097056427.1 | WP_097056067.1 | WP_097056276.1 | WP_097057124.1 |
| Bacteroidota | *Salinimicrobium soli* | WP_418336149.1 | WP_418335852.1 | WP_418335645.1 | WP_418334907.1 |
| Bacteroidota | *Salinimicrobium sp. 3283s* | WP_345598467.1 | WP_345598592.1 | WP_345598933.1 | WP_345598169.1 |
| Bacteroidota | *Salinimicrobium terrae* | WP_029034933.1 | WP_029033011.1 | WP_029033552.1 | WP_029034990.1 |
| Bacteroidota | *Salinimicrobium tongyeongense* | WP_265163318.1 | WP_265163551.1 | WP_265163274.1 | WP_265164049.1 |
| Bacteroidota | *Salinimicrobium xinjiangense* | WP_029036764.1 | WP_029036068.1 | WP_029037520.1 | WP_029038605.1 |
| Bacteroidota | *Sandaracinomonas limnophila* | WP_127803033.1 | WP_127804355.1 | WP_127802513.1 | WP_127803171.1 |
| Bacteroidota | *Sanyastnella coralliicola* | WP_306644150.1 | WP_306643031.1 | WP_306639898.1 | WP_306643246.1 |
| Bacteroidota | *Saprospira grandis* | WP_002661067.1 | WP_270100066.1 | WP_015691616.1 | WP_002660161.1 |
| Bacteroidota | *Saprospira sp. CCB-QB6* | WP_272619536.1 | WP_272620540.1 | WP_272620609.1 | WP_272620088.1 |
| Bacteroidota | *Schleiferia thermophila* | WP_037356799.1 | WP_037355887.1 | WP_037358008.1 | WP_160171979.1 |
| Bacteroidota | *Sediminibacter sp. Hel_I_10* | WP_026755642.1 | WP_026755702.1 | WP_026755964.1 | WP_026753280.1 |
| Bacteroidota | *Sediminibacterium ginsengisoli* | WP_078832212.1 | WP_078831188.1 | WP_078832335.1 | WP_245825700.1 |
| Bacteroidota | *Sediminibacterium goheungense* | WP_133475208.1 | WP_133474678.1 | WP_133475808.1 | WP_133475323.1 |
| Bacteroidota | *Sediminibacterium roseum* | WP_161819353.1 | WP_161817057.1 | WP_161818326.1 | WP_161817922.1 |
| Bacteroidota | *Sediminibacterium salmoneum* | WP_026763428.1 | WP_026764238.1 | WP_026763456.1 | WP_026763181.1 |
| Bacteroidota | *Sediminibacterium soli* | WP_161834845.1 | WP_161835386.1 | WP_161834296.1 | WP_161834578.1 |
| Bacteroidota | *Sediminibacterium sp.* | WP_324483938.1 | WP_324486156.1 | WP_305536514.1 | WP_305534927.1 |
| Bacteroidota | *Sediminitomix flava* | WP_109622381.1 | WP_109616372.1 | WP_109618873.1 | WP_245935572.1 |
| Bacteroidota | *Segetibacter sp. 3557_3* | WP_133268512.1 | WP_133268197.1 | WP_133266548.1 | WP_133267570.1 |
| Bacteroidota | *Seonamhaeicola algicola* | WP_147133561.1 | WP_147130549.1 | WP_147133745.1 | WP_147133851.1 |
| Bacteroidota | *Seonamhaeicola aphaedonensis* | WP_116524709.1 | WP_116039943.1 | WP_116523873.1 | WP_116524046.1 |
| Bacteroidota | *Seonamhaeicola marinus* | WP_148544647.1 | WP_148541872.1 | WP_148544579.1 | WP_262713610.1 |
| Bacteroidota | *Seonamhaeicola maritimus* | WP_147766866.1 | WP_147767208.1 | WP_147769343.1 | WP_282136242.1 |
| Bacteroidota | *Seonamhaeicola sediminis* | WP_133356969.1 | WP_133356602.1 | WP_133357173.1 | WP_133354879.1 |

| | | | | | |
|---|---|---|---|---|---|
| Bacteroidota | *Seonamhaeicola sp.* | WP_299549661.1 | WP_299547549.1 | WP_299550404.1 | WP_372938276.1 |
| Bacteroidota | *Shiella aurantiaca* | WP_320004588.1 | WP_320003240.1 | WP_320003921.1 | WP_320004771.1 |
| Bacteroidota | *Siansivirga zeaxanthinifaciens* | WP_044638673.1 | WP_044638880.1 | WP_044638641.1 | WP_044638743.1 |
| Bacteroidota | *Snuella lapsa* | WP_345003992.1 | WP_345003601.1 | WP_345003664.1 | WP_345007529.1 |
| Bacteroidota | *Snuella sedimenti* | WP_199115323.1 | WP_199115617.1 | WP_199116703.1 | WP_199114629.1 |
| Bacteroidota | *Solirubrum puertoriconensis* | WP_059071577.1 | WP_059067657.1 | WP_059071390.1 | WP_059068878.1 |
| Bacteroidota | *Solitalea canadensis* | WP_014680599.1 | WP_014682578.1 | WP_014680712.1 | WP_014681131.1 |
| Bacteroidota | *Solitalea koreensis* | WP_142603940.1 | WP_142603364.1 | WP_142604438.1 | WP_142600541.1 |
| Bacteroidota | *Solitalea lacus* | WP_237846350.1 | WP_237848968.1 | WP_237846591.1 | WP_237847143.1 |
| Bacteroidota | *Solitalea longa* | WP_103789570.1 | WP_103787391.1 | WP_103790159.1 | WP_103788875.1 |
| Bacteroidota | *Soonwooa buanensis* | WP_079665548.1 | WP_079668157.1 | WP_079667385.1 | WP_079666718.1 |
| Bacteroidota | *Soonwooa sp.* | WP_312324244.1 | WP_312322246.1 | WP_300673687.1 | WP_313027157.1 |
| Bacteroidota | *Spongiivirga citrea* | WP_164029590.1 | WP_164029170.1 | WP_164028867.1 | WP_164032428.1 |
| Bacteroidota | *Spongiivirga sp. MCCC 1A20706* | WP_410484143.1 | WP_410484950.1 | WP_410483661.1 | WP_410485712.1 |
| Bacteroidota | *Sporocytophaga myxococcoides* | WP_045468567.1 | WP_045459397.1 | WP_045464077.1 | WP_045459954.1 |
| Bacteroidota | *Sporocytophaga sp.* | WP_293893448.1 | WP_293897300.1 | WP_293894613.1 | WP_293892478.1 |
| Bacteroidota | *Subsaxibacter sp. CAU 1640* | WP_248024861.1 | WP_248025836.1 | WP_248025383.1 | WP_248024187.1 |
| Bacteroidota | *Subsaximicrobium wynnwilliamsii* | WP_147085143.1 | WP_147084534.1 | WP_147085199.1 | WP_317128650.1 |
| Bacteroidota | *Sungkyunkwania multivorans* | WP_386403032.1 | WP_386407340.1 | WP_386405566.1 | WP_386407050.1 |
| Bacteroidota | *Taibaiella chishuiensis* | WP_106521218.1 | WP_106525076.1 | WP_106521535.1 | WP_181358454.1 |
| Bacteroidota | *Taibaiella helva* | WP_118952493.1 | WP_118953545.1 | WP_118949944.1 | WP_157976869.1 |
| Bacteroidota | *Taibaiella koreensis* | WP_118973686.1 | WP_118972826.1 | WP_118974487.1 | WP_162903180.1 |
| Bacteroidota | *Taibaiella lutea* | WP_150032017.1 | WP_150031081.1 | WP_150033014.1 | WP_190277335.1 |
| Bacteroidota | *Taibaiella soli* | WP_243630355.1 | WP_110999133.1 | WP_110996983.1 | WP_110999509.1 |
| Bacteroidota | *Taibaiella sp. KBW10* | WP_124634811.1 | WP_124636061.1 | WP_124635204.1 | WP_124636155.1 |
| Bacteroidota | *Taishania pollutisoli* | WP_163493010.1 | WP_163492192.1 | WP_163490697.1 | WP_216714193.1 |
| Bacteroidota | *Tamlana crocina* | WP_167917346.1 | WP_167917677.1 | WP_167919677.1 | WP_167918689.1 |
| Bacteroidota | *Tamlana flava* | WP_370480120.1 | WP_370479806.1 | WP_370476504.1 | WP_370476851.1 |

| | | | | | |
|---|---|---|---|---|---|
| Bacteroidota | *Tamlana sp. 2201CG12-4* | WP_326284301.1 | WP_326283243.1 | WP_326284009.1 | WP_326282716.1 |
| Bacteroidota | *Tenacibaculum adriaticum* | WP_148869522.1 | WP_148869874.1 | WP_148871029.1 | WP_148871047.1 |
| Bacteroidota | *Tenacibaculum aestuarii* | WP_408036025.1 | WP_408035614.1 | WP_408036213.1 | WP_408036178.1 |
| Bacteroidota | *Tenacibaculum aestuariivivum* | WP_418650629.1 | WP_418650557.1 | WP_418649929.1 | WP_418649959.1 |
| Bacteroidota | *Tenacibaculum agarivorans* | WP_075340842.1 | WP_075342671.1 | WP_075343661.1 | WP_075343920.1 |
| Bacteroidota | *Tenacibaculum aiptasiae* | WP_150901146.1 | WP_150899186.1 | WP_150899780.1 | WP_272151765.1 |
| Bacteroidota | *Tenacibaculum amylolyticum* | WP_408038028.1 | WP_408039680.1 | WP_408039792.1 | WP_408037409.1 |
| Bacteroidota | *Tenacibaculum ascidiaceicola* | WP_415268602.1 | WP_425658519.1 | WP_415268493.1 | WP_425659246.1 |
| Bacteroidota | *Tenacibaculum caenipelagi* | WP_133534370.1 | WP_133536489.1 | WP_133534486.1 | WP_133534460.1 |
| Bacteroidota | *Tenacibaculum crassostreae* | WP_408046608.1 | WP_408046929.1 | WP_408046466.1 | WP_408046492.1 |
| Bacteroidota | *Tenacibaculum dicentrarchi* | WP_214985934.1 | WP_370407873.1 | WP_101902640.1 | WP_101902347.1 |
| Bacteroidota | *Tenacibaculum discolor* | WP_121147477.1 | WP_121146781.1 | WP_124590251.1 | WP_121148738.1 |
| Bacteroidota | *Tenacibaculum finnmarkense* | WP_239780004.1 | WP_232122331.1 | WP_101915365.1 | WP_232128703.1 |
| Bacteroidota | *Tenacibaculum gallaicum* | WP_115900783.1 | WP_115901618.1 | WP_115900961.1 | WP_115900927.1 |
| Bacteroidota | *Tenacibaculum geojense* | WP_386105148.1 | WP_386104704.1 | WP_386104940.1 | WP_386104979.1 |
| Bacteroidota | *Tenacibaculum holothuriorum* | WP_198938355.1 | WP_086029382.1 | WP_086030732.1 | WP_086030767.1 |
| Bacteroidota | *Tenacibaculum insulae* | WP_418644920.1 | WP_418644914.1 | WP_418644674.1 | WP_418644704.1 |
| Bacteroidota | *Tenacibaculum jejuense* | WP_095071561.1 | WP_095072327.1 | WP_095073068.1 | WP_095072980.1 |
| Bacteroidota | *Tenacibaculum larymnensis* | WP_274641245.1 | WP_274639816.1 | WP_274640215.1 | WP_274640185.1 |
| Bacteroidota | *Tenacibaculum litopenaei* | WP_408044355.1 | WP_408044075.1 | WP_408043301.1 | WP_408044502.1 |
| Bacteroidota | *Tenacibaculum lutimaris* | WP_120187015.1 | WP_120186681.1 | WP_120187165.1 | WP_120187140.1 |
| Bacteroidota | *Tenacibaculum maritimum* | WP_024741845.1 | WP_159246260.1 | WP_159320935.1 | WP_406755780.1 |
| Bacteroidota | *Tenacibaculum mesophilum* | WP_394421808.1 | WP_047788878.1 | WP_253679894.1 | WP_047789209.1 |
| Bacteroidota | *Tenacibaculum ovolyticum* | WP_237275937.1 | WP_064966370.1 | WP_271241654.1 | WP_064968091.1 |
| Bacteroidota | *Tenacibaculum pelagium* | WP_182123875.1 | WP_182123523.1 | WP_182123957.1 | WP_182123931.1 |
| Bacteroidota | *Tenacibaculum piscium* | WP_101917121.1 | WP_233900280.1 | WP_233898313.1 | WP_233901351.1 |
| Bacteroidota | *Tenacibaculum platacis* | WP_348725701.1 | WP_348712205.1 | WP_348742779.1 | WP_348742864.1 |
| Bacteroidota | *Tenacibaculum polynesiense* | WP_348718339.1 | WP_348716780.1 | WP_348715293.1 | WP_348718874.1 |

| | | | | | |
|---|---|---|---|---|---|
| Bacteroidota | *Tenacibaculum sediminilitoris* | WP_408024488.1 | WP_408024139.1 | WP_408023732.1 | WP_408025091.1 |
| Bacteroidota | *Tenacibaculum singaporense* | WP_125067213.1 | WP_125345071.1 | WP_125068073.1 | WP_125344444.1 |
| Bacteroidota | *Tenacibaculum skagerrakense* | WP_132794118.1 | WP_132794789.1 | WP_132795009.1 | WP_132795025.1 |
| Bacteroidota | *Tenacibaculum soleae* | WP_271405274.1 | WP_068705066.1 | WP_303558687.1 | WP_303529621.1 |
| Bacteroidota | *Tenacibaculum sp.* | WP_417785267.1 | WP_273694360.1 | WP_417786261.1 | WP_273694783.1 |
| Bacteroidota | *Tenacibaculum tangerinum* | WP_279652504.1 | WP_279652411.1 | WP_279651372.1 | WP_279651401.1 |
| Bacteroidota | *Tenacibaculum todarodis* | WP_072554693.1 | WP_072554883.1 | WP_072555161.1 | WP_072555142.1 |
| Bacteroidota | *Tenacibaculum vairaonense* | WP_348747241.1 | WP_348740343.1 | WP_348746997.1 | WP_348702758.1 |
| Bacteroidota | *Tenacibaculum xiamenense* | WP_408032529.1 | WP_408031608.1 | WP_408030665.1 | WP_408032154.1 |
| Bacteroidota | *Terrimonas pollutisoli* | WP_276503523.1 | WP_276500859.1 | WP_276504086.1 | WP_276502652.1 |
| Bacteroidota | *Terrimonas sp. R1* | WP_385886284.1 | WP_385894225.1 | WP_385890983.1 | WP_385885141.1 |
| Bacteroidota | *Thalassobellus suaedae* | WP_415866307.1 | WP_415866465.1 | WP_415861774.1 | WP_415866469.1 |
| Bacteroidota | *Thermaurantimonas aggregans* | WP_124396926.1 | WP_124396826.1 | WP_124397494.1 | WP_160160588.1 |
| Bacteroidota | *Thermaurantimonas sp.* | WP_409769666.1 | WP_409769462.1 | WP_409771203.1 | WP_409771132.1 |
| Bacteroidota | *Thermonema lapsum* | WP_166921076.1 | WP_166919763.1 | WP_166920515.1 | WP_166918106.1 |
| Bacteroidota | *Thermonema rossianum* | WP_038032522.1 | WP_051632972.1 | WP_038030079.1 | WP_084147099.1 |
| Bacteroidota | *Thermonema sp.* | WP_288006170.1 | WP_288005729.1 | WP_288007082.1 | WP_288004089.1 |
| Bacteroidota | *Ulvibacter antarcticus* | WP_121906617.1 | WP_121906710.1 | WP_121907624.1 | WP_121907465.1 |
| Bacteroidota | *Ulvibacter litoralis* | WP_093143850.1 | WP_093143547.1 | WP_093144987.1 | WP_093145223.1 |
| Bacteroidota | *Ulvibacter sp. MAR_2010_11* | WP_100804002.1 | WP_100803909.1 | WP_100804196.1 | WP_100803434.1 |
| Bacteroidota | *Urechidicola croceus* | WP_070235653.1 | WP_070235832.1 | WP_070235596.1 | WP_070236178.1 |
| Bacteroidota | *Urechidicola vernalis* | WP_311592092.1 | WP_311591836.1 | WP_311592154.1 | WP_311594012.1 |
| Bacteroidota | *Vaginella massiliensis* | WP_395092052.1 | WP_068598256.1 | WP_395093015.1 | WP_395090947.1 |
| Bacteroidota | *Vicingus serpentipes* | WP_147099218.1 | WP_147101768.1 | WP_147097973.1 | WP_147100110.1 |
| Bacteroidota | *Wandonia haliotis* | WP_343786300.1 | WP_343788488.1 | WP_343787892.1 | WP_343785738.1 |
| Bacteroidota | *Wenyingzhuangia aestuarii* | WP_167898478.1 | WP_167898441.1 | WP_167897819.1 | WP_167896793.1 |
| Bacteroidota | *Wenyingzhuangia fucanilytica* | WP_068825163.1 | WP_068826438.1 | WP_068825555.1 | WP_068826364.1 |
| Bacteroidota | *Wenyingzhuangia gilva* | WP_302884102.1 | WP_302883866.1 | WP_302885059.1 | WP_302883913.1 |

| | | | | | |
|---|---|---|---|---|---|
| Bacteroidota | *Wenyingzhuangia heitensis* | WP_167186003.1 | WP_167185859.1 | WP_167187886.1 | WP_167187308.1 |
| Bacteroidota | *Wenyingzhuangia marina* | WP_073120729.1 | WP_073119375.1 | WP_073118897.1 | WP_073119261.1 |
| Bacteroidota | *Wenyingzhuangia sp.* | WP_347182171.1 | WP_347180341.1 | WP_347180434.1 | WP_347180213.1 |
| Bacteroidota | *Winogradskyella alexanderae* | WP_224526353.1 | WP_224530615.1 | WP_224526489.1 | WP_224526926.1 |
| Bacteroidota | *Winogradskyella algicola* | WP_138434410.1 | WP_138433396.1 | WP_138434160.1 | WP_138434612.1 |
| Bacteroidota | *Winogradskyella aquimaris* | WP_320555137.1 | WP_320556630.1 | WP_320556356.1 | WP_320555584.1 |
| Bacteroidota | *Winogradskyella arenosi* | WP_114309688.1 | WP_114309489.1 | WP_114309772.1 | WP_114310057.1 |
| Bacteroidota | *Winogradskyella aurantia* | WP_094967441.1 | WP_094968397.1 | WP_094967377.1 | WP_094968831.1 |
| Bacteroidota | *Winogradskyella aurantiaca* | WP_115463024.1 | WP_115462847.1 | WP_115463094.1 | WP_317047361.1 |
| Bacteroidota | *Winogradskyella bathintestinalis* | WP_290206016.1 | WP_290206804.1 | WP_290206994.1 | WP_290206180.1 |
| Bacteroidota | *Winogradskyella costae* | WP_179336134.1 | WP_179335048.1 | WP_179334623.1 | WP_179335300.1 |
| Bacteroidota | *Winogradskyella echinorum* | WP_186846218.1 | WP_186845557.1 | WP_186845790.1 | WP_186845877.1 |
| Bacteroidota | *Winogradskyella eckloniae* | WP_173281221.1 | WP_173281061.1 | WP_173280570.1 | WP_173281378.1 |
| Bacteroidota | *Winogradskyella endarachnes* | WP_157362260.1 | WP_157361762.1 | WP_157363281.1 | WP_157362488.1 |
| Bacteroidota | *Winogradskyella epiphytica* | WP_110475366.1 | WP_110475170.1 | WP_110476132.1 | WP_110476237.1 |
| Bacteroidota | *Winogradskyella eximia* | WP_115818299.1 | WP_115817543.1 | WP_115818051.1 | WP_115817452.1 |
| Bacteroidota | *Winogradskyella flava* | WP_282043432.1 | WP_185788902.1 | WP_282043289.1 | WP_282043124.1 |
| Bacteroidota | *Winogradskyella forsetii* | WP_179008373.1 | WP_179020256.1 | WP_179006261.1 | WP_179018484.1 |
| Bacteroidota | *Winogradskyella haliclonae* | WP_188374535.1 | WP_188374348.1 | WP_188374596.1 | WP_188375109.1 |
| Bacteroidota | *Winogradskyella helgolandensis* | WP_179319568.1 | WP_178985666.1 | WP_317167555.1 | WP_179319418.1 |
| Bacteroidota | *Winogradskyella immobilis* | WP_227477884.1 | WP_227477848.1 | WP_227476698.1 | WP_227477668.1 |
| Bacteroidota | *Winogradskyella jejuensis* | WP_073087005.1 | WP_073086886.1 | WP_073087204.1 | WP_073085722.1 |
| Bacteroidota | *Winogradskyella litorisediminis* | WP_386127345.1 | WP_386127183.1 | WP_386132737.1 | WP_386132081.1 |
| Bacteroidota | *Winogradskyella litoriviva* | WP_173300366.1 | WP_173300105.1 | WP_173301905.1 | WP_173300484.1 |
| Bacteroidota | *Winogradskyella ludwigii* | WP_179337864.1 | WP_179338973.1 | WP_179339464.1 | WP_179338155.1 |
| Bacteroidota | *Winogradskyella luteola* | WP_218544163.1 | WP_218544848.1 | WP_218544234.1 | WP_218544490.1 |
| Bacteroidota | *Winogradskyella maritima* | WP_386095730.1 | WP_386101441.1 | WP_386095869.1 | WP_386099159.1 |
| Bacteroidota | *Winogradskyella pacifica* | WP_115808434.1 | WP_115808652.1 | WP_115809554.1 | WP_179349795.1 |

| | | | | | |
|---|---|---|---|---|---|
| Bacteroidota | *Winogradskyella pelagia* | WP_208153639.1 | WP_208154871.1 | WP_208153767.1 | WP_208154100.1 |
| Bacteroidota | *Winogradskyella poriferorum* | WP_331810318.1 | WP_331809773.1 | WP_331810245.1 | WP_331809409.1 |
| Bacteroidota | *Winogradskyella psychrotolerans* | WP_020896912.1 | WP_215925849.1 | WP_215936346.1 | WP_020896739.1 |
| Bacteroidota | *Winogradskyella pulchriflava* | WP_386063557.1 | WP_386061894.1 | WP_386061462.1 | WP_386064116.1 |
| Bacteroidota | *Winogradskyella rapida* | WP_386115939.1 | WP_386114648.1 | WP_386113114.1 | WP_386117398.1 |
| Bacteroidota | *Winogradskyella schleiferi* | WP_178988959.1 | WP_178988830.1 | WP_178988226.1 | WP_178987854.1 |
| Bacteroidota | *Winogradskyella sediminis* | WP_115839258.1 | WP_092443319.1 | WP_417876348.1 | WP_115839130.1 |
| Bacteroidota | *Winogradskyella sp.* | WP_296384836.1 | WP_296352858.1 | WP_391593001.1 | WP_370103802.1 |
| Bacteroidota | *Winogradskyella tangerina* | WP_111684169.1 | WP_111682739.1 | WP_111684095.1 | WP_111683804.1 |
| Bacteroidota | *Winogradskyella thalassocola* | WP_092468787.1 | WP_092469884.1 | WP_092467141.1 | WP_092469187.1 |
| Bacteroidota | *Winogradskyella undariae* | WP_173588526.1 | WP_179315410.1 | WP_179317035.1 | WP_173588138.1 |
| Bacteroidota | *Winogradskyella ursingii* | WP_179346562.1 | WP_179344862.1 | WP_179346499.1 | WP_179346073.1 |
| Bacteroidota | *Winogradskyella vidalii* | WP_179352015.1 | WP_179353704.1 | WP_179352429.1 | WP_179351812.1 |
| Bacteroidota | *Winogradskyella vincentii* | WP_224478894.1 | WP_224478105.1 | WP_224479179.1 | WP_224478740.1 |
| Bacteroidota | *Winogradskyella wandonensis* | WP_132705735.1 | WP_132703720.1 | WP_132705881.1 | WP_132702815.1 |
| Bacteroidota | *Winogradskyella wichelsiae* | WP_179376080.1 | WP_179376959.1 | WP_179376871.1 | WP_179376516.1 |
| Bacteroidota | *Wocania arenilitoris* | WP_237240727.1 | WP_237240459.1 | WP_237239480.1 | WP_237240506.1 |
| Bacteroidota | *Wocania ichthyoenteri* | WP_034044808.1 | WP_034044660.1 | WP_034040352.1 | WP_034041167.1 |
| Bacteroidota | *Xanthocytophaga flavus* | WP_314000089.1 | WP_313978959.1 | WP_313986097.1 | WP_313978846.1 |
| Bacteroidota | *Xanthomarina gelatinilytica* | WP_417873413.1 | WP_417857232.1 | WP_370100095.1 | WP_417882250.1 |
| Bacteroidota | *Xanthomarina sp.* | WP_324312250.1 | WP_324313051.1 | WP_286853545.1 | WP_286854418.1 |
| Bacteroidota | *Xanthomarina spongicola* | WP_109683326.1 | WP_109681702.1 | WP_109681507.1 | WP_109682487.1 |
| Bacteroidota | *Yeosuana aromativorans* | WP_188654342.1 | WP_188653250.1 | WP_188654997.1 | WP_188652436.1 |
| Bacteroidota | *Yeosuana marina* | WP_166961327.1 | WP_166965288.1 | WP_166963373.1 | WP_166965638.1 |
| Balneolota | *Aliifodinibius salipaludis* | WP_095607415.1 | WP_095605785.1 | WP_095607636.1 | WP_255233382.1 |
| Balneolota | *Aliifodinibius sp. S!AR15-10* | WP_310686561.1 | WP_310684926.1 | WP_310682093.1 | WP_310685409.1 |
| Balneolota | *Balneola sp. EhC07* | WP_066223311.1 | WP_066222314.1 | WP_066220100.1 | WP_066217088.1 |
| Balneolota | *Cyclonatronum proteinivorum* | WP_164682415.1 | WP_114985548.1 | WP_114983795.1 | WP_164682663.1 |

| | | | | | |
|---|---|---|---|---|---|
| Balneolota | *Cyclonatronum sp.* | WP_291482411.1 | WP_291482712.1 | WP_291482818.1 | WP_291482004.1 |
| Balneolota | *Fodinibius roseus* | WP_073063831.1 | WP_073061524.1 | WP_073063141.1 | WP_073059135.1 |
| Balneolota | *Fodinibius salicampi* | WP_265791860.1 | WP_265790485.1 | WP_265788309.1 | WP_265787781.1 |
| Balneolota | *Fodinibius salinus* | WP_148899828.1 | WP_148898505.1 | WP_246138157.1 | WP_170245550.1 |
| Balneolota | *Fodinibius saliphilus* | WP_138432167.1 | WP_138430825.1 | WP_138430056.1 | WP_138429484.1 |
| Balneolota | *Fodinibius salsisoli* | WP_265767352.1 | WP_265764074.1 | WP_265766003.1 | WP_265765528.1 |
| Balneolota | *Fodinibius sediminis* | WP_142714531.1 | WP_142712524.1 | WP_142714645.1 | WP_185958215.1 |
| Balneolota | *Fodinibius sp.* | WP_372634043.1 | WP_322574622.1 | WP_322575165.1 | WP_322575107.1 |
| Balneolota | *Halalkalibaculum roseum* | WP_165143800.1 | WP_165138370.1 | WP_165140079.1 | WP_165140357.1 |
| Balneolota | *Halalkalibaculum sp. DA3122* | WP_395257904.1 | WP_395259678.1 | WP_395259273.1 | WP_395261429.1 |
| Balneolota | *Natronogracilivirga saccharolytica* | WP_210509762.1 | WP_210513032.1 | WP_210509515.1 | WP_210511334.1 |
| Balneolota | *Rhodohalobacter barkolensis* | WP_101073171.1 | WP_101073066.1 | WP_101071599.1 | WP_101071812.1 |
| Balneolota | *Rhodohalobacter mucosus* | WP_109647948.1 | WP_109647779.1 | WP_109646781.1 | WP_109645422.1 |
| Balneolota | *Rhodohalobacter sp.* | WP_372902497.1 | WP_322570300.1 | WP_372905583.1 | WP_372904606.1 |
| Balneolota | *Rhodohalobacter sulfatireducens* | WP_237852421.1 | WP_237852468.1 | WP_237853641.1 | WP_237855134.1 |
| Calditrichaeota | *Caldithrix abyssi* | WP_006931011.1 | WP_006929106.1 | WP_044281486.1 | WP_006931009.1 |
| Chlamydiota | *Criblamydia sequanensis* | WP_041017343.1 | WP_041016448.1 | WP_053331647.1 | WP_041017369.1 |
| Chlamydiota | *Estrella lausannensis* | WP_098038228.1 | WP_239414312.1 | WP_098037590.1 | WP_098038239.1 |
| Chlamydiota | *Waddlia chondrophila* | WP_013182233.1 | WP_041941457.1 | WP_013181378.1 | WP_013181809.1 |
| Chlorobiota | *Candidatus Chlorobium masyuteum* | WP_166808611.1 | WP_166808429.1 | WP_166808443.1 | WP_166807795.1 |
| Chlorobiota | *Chlorobaculum limnaeum* | WP_069809554.1 | WP_069810356.1 | WP_069809037.1 | WP_069808910.1 |
| Chlorobiota | *Chlorobaculum parvum* | WP_012502214.1 | WP_012502674.1 | WP_012501996.1 | WP_012501656.1 |
| Chlorobiota | *Chlorobaculum sp. MV4-Y* | WP_260633512.1 | WP_260533492.1 | WP_260633235.1 | WP_260533288.1 |
| Chlorobiota | *Chlorobaculum tepidum* | WP_010932718.1 | WP_164926877.1 | WP_010933086.1 | WP_010933222.1 |
| Chlorobiota | *Chlorobaculum thiosulfatiphilum* | WP_139457526.1 | WP_139456871.1 | WP_139457025.1 | WP_139457127.1 |
| Chlorobiota | *Chlorobium ferrooxidans* | WP_006366651.1 | WP_006366427.1 | WP_006367401.1 | WP_006365343.1 |
| Chlorobiota | *Chlorobium limicola* | WP_012466208.1 | WP_041465847.1 | WP_012465860.1 | WP_276672508.1 |
| Chlorobiota | *Chlorobium phaeobacteroides* | WP_011745192.1 | WP_011744678.1 | WP_011744703.1 | WP_011745851.1 |

| | | | | | |
|---|---|---|---|---|---|
| Chlorobiota | *Chlorobium phaeovibrioides* | WP_151419445.1 | WP_126384076.1 | WP_126342229.1 | WP_126384756.1 |
| Chlorobiota | *Chlorobium sp.* | WP_423931470.1 | WP_331561158.1 | WP_292002077.1 | WP_331561788.1 |
| Chlorobiota | *Chloroherpeton thalassium* | WP_012500949.1 | WP_012499088.1 | WP_012500566.1 | WP_012499687.1 |
| Chlorobiota | *Pelodictyon luteolum* | WP_011357747.1 | WP_303680767.1 | WP_011358150.1 | WP_303682490.1 |
| Chlorobiota | *Pelodictyon phaeoclathratiforme* | WP_012508109.1 | WP_041526422.1 | WP_012508493.1 | WP_012509052.1 |
| Chlorobiota | *Prosthecochloris sp.* | WP_294347425.1 | WP_288031379.1 | WP_294346988.1 | WP_287220752.1 |
| Deinococcota | *Thermus sp. PS18* | WP_219760243.1 | WP_267260854.1 | WP_267260744.1 | WP_267261639.1 |
| Gemmatimonadota | *Candidatus Palauibacter irciniicola* | WP_419858893.1 | WP_419859026.1 | WP_419856829.1 | WP_419858894.1 |
| Gemmatimonadota | *Candidatus Palauibacter polyketidifaciens* | WP_310785101.1 | WP_310783687.1 | WP_310785236.1 | WP_310785103.1 |
| Gemmatimonadota | *Candidatus Palauibacter scopulicola* | WP_310778325.1 | WP_310778401.1 | WP_310775338.1 | WP_310778328.1 |
| Gemmatimonadota | *Candidatus Palauibacter soopunensis* | WP_310759046.1 | WP_310758907.1 | WP_310757617.1 | WP_310759045.1 |
| Gemmatimonadota | *Gemmatimonas sp.* | WP_331050747.1 | WP_396219785.1 | WP_409947910.1 | WP_396206035.1 |
| Gemmatimonadota | *Longimicrobium sp.* | WP_329686406.1 | WP_331128124.1 | WP_331876780.1 | WP_422655644.1 |
| Gemmatimonadota | *Longimicrobium terrae* | WP_170036276.1 | WP_170038756.1 | WP_170035611.1 | WP_170036976.1 |
| Gemmatimonadota | *Pseudogemmatithrix spongiicola* | WP_367885176.1 | WP_367887761.1 | WP_367887364.1 | WP_367887824.1 |
| Ignavibacteriota | *Ignavibacterium album* | WP_014560944.1 | WP_014561555.1 | WP_014560830.1 | WP_014560661.1 |
| Ignavibacteriota | *Ignavibacterium sp.* | WP_297843333.1 | WP_337866183.1 | WP_337872606.1 | WP_297842622.1 |
| Myxococcota | *Stigmatella aurantiaca* | WP_002618112.1 | WP_013374927.1 | WP_075007943.1 | WP_075007944.1 |
| Myxococcota | *Vulgatibacter sp.* | WP_373046082.1 | WP_373045413.1 | WP_373045414.1 | WP_373045415.1 |
| Planctomycetota | *Engelhardtia mirabilis* | WP_145069642.1 | WP_145070674.1 | WP_419191964.1 | WP_419191963.1 |
| Planctomycetota | *Saltatorellus ferox* | WP_419191093.1 | WP_145205850.1 | WP_419190757.1 | WP_145205827.1 |
| Rhodothermota | *Longibacter salinarum* | WP_098076119.1 | WP_098074559.1 | WP_098074888.1 | WP_098074867.1 |
| Rhodothermota | *Longimonas halophila* | WP_098061634.1 | WP_098061945.1 | WP_098063067.1 | WP_098060759.1 |
| Rhodothermota | *Longimonas sp.* | WP_353562514.1 | WP_410338332.1 | WP_410338071.1 | WP_353562900.1 |
| Rhodothermota | *Rhodocaloribacter litoris* | WP_166978522.1 | WP_166974234.1 | WP_228350645.1 | WP_166978060.1 |
| Rhodothermota | *Rhodothermus bifroesti* | WP_210376068.1 | WP_210373907.1 | WP_210375124.1 | WP_210375121.1 |
| Rhodothermota | *Rhodothermus marinus* | WP_161540542.1 | WP_012842763.1 | WP_012843182.1 | WP_014067708.1 |

| | | | | | |
|---|---|---|---|---|---|
| Rhodothermota | *Rhodothermus profundi* | WP_072714519.1 | WP_072715349.1 | WP_072715210.1 | WP_072715214.1 |
| Rhodothermota | *Rubrivirga sp.* | WP_420456952.1 | WP_420455377.1 | WP_424521619.1 | WP_420454540.1 |
| Rhodothermota | *Salinibacter altiplanensis* | WP_103021536.1 | WP_103019979.1 | WP_103027427.1 | WP_103029837.1 |
| Rhodothermota | *Salinibacter grassmerensis* | WP_263787278.1 | WP_263786151.1 | WP_263785401.1 | WP_263786104.1 |
| Rhodothermota | *Salinibacter ruber* | WP_251931083.1 | WP_341481556.1 | WP_112903583.1 | WP_259085182.1 |
| Rhodothermota | Salinibacter sp. | WP_375802491.1 | WP_375802762.1 | WP_263834908.1 | WP_375801842.1 |
| Rhodothermota | Salisaeta longa | WP_022835959.1 | WP_022835967.1 | WP_022836238.1 | WP_028567198.1 |

Supplementary Dataset 1B – Unique species containing the complete BSH biosynthetic pathway. Phylum, species, and NCBI accession codes for BshA, BshB, and BshC are listed for each organism.

| **Phylum** | **Organism** | **Accession code BshA** | **Accession code BshB** | **Accession code BshC** |
|---|---|---|---|---|
| Acidobacteriota | *Acidicapsa acidisoli* | WP_263355290.1 | WP_263352547.1 | WP_263353948.1 |
| Acidobacteriota | *Acidicapsa dinghuensis* | WP_263332892.1 | WP_263339261.1 | WP_263342256.1 |
| Acidobacteriota | *Acidicapsa ligni* | WP_263357988.1 | WP_317890557.1 | WP_263356903.1 |
| Acidobacteriota | *Acidipila rosea* | WP_131991325.1 | WP_131994826.1 | WP_131994166.1 |
| Acidobacteriota | *Acidisarcina polymorpha* | WP_114206173.1 | WP_338026767.1 | WP_114205520.1 |
| Acidobacteriota | *Acidobacterium sp. S8* | WP_158748945.1 | WP_158749030.1 | WP_158751191.1 |
| Acidobacteriota | *Alloacidobacterium dinghuense* | WP_186747285.1 | WP_186745795.1 | WP_186744667.1 |
| Acidobacteriota | *Alloacidobacterium sp.* | WP_324830802.1 | WP_324832157.1 | WP_324831241.1 |
| Acidobacteriota | *Bryocella elongata* | WP_103931846.1 | WP_103931120.1 | WP_103933629.1 |
| Acidobacteriota | *Candidatus Korobacter versatilis* | WP_041857031.1 | WP_011524233.1 | WP_011522983.1 |
| Acidobacteriota | *Chloracidobacterium aggregatum* | WP_211423704.1 | WP_211423485.1 | WP_211426094.1 |
| Acidobacteriota | *Chloracidobacterium sp. D* | WP_211432787.1 | WP_211433022.1 | WP_211432012.1 |
| Acidobacteriota | *Chloracidobacterium thermophilum* | WP_058866111.1 | WP_058865916.1 | WP_014098599.1 |
| Acidobacteriota | *Chloracidobacterium validum* | WP_211428433.1 | WP_211430360.1 | WP_211428837.1 |
| Acidobacteriota | *Edaphobacter aggregans* | WP_035355527.1 | WP_125484521.1 | WP_125484223.1 |
| Acidobacteriota | *Edaphobacter bradus* | WP_263366346.1 | WP_263365139.1 | WP_263368437.1 |

| | | | | |
|---|---|---|---|---|
| Acidobacteriota | *Edaphobacter dinghuensis* | WP_188552996.1 | WP_188553926.1 | WP_188554963.1 |
| Acidobacteriota | *Edaphobacter flagellatus* | WP_260705587.1 | WP_260706006.1 | WP_260704451.1 |
| Acidobacteriota | *Edaphobacter modestus* | WP_130418350.1 | WP_130421490.1 | WP_130418550.1 |
| Acidobacteriota | *Edaphobacter paludis* | WP_348268774.1 | WP_348266716.1 | WP_348269426.1 |
| Acidobacteriota | *Edaphobacter sp.* | WP_325386905.1 | WP_299377192.1 | WP_326445757.1 |
| Acidobacteriota | *Granulicella aggregans* | WP_184213714.1 | WP_184217056.1 | WP_184217029.1 |
| Acidobacteriota | *Granulicella arctica* | WP_179492651.1 | WP_179490700.1 | WP_263381590.1 |
| Acidobacteriota | *Granulicella cerasi* | WP_390236537.1 | WP_263370744.1 | WP_263371471.1 |
| Acidobacteriota | *Granulicella mallensis* | WP_014264654.1 | WP_184252288.1 | WP_184255776.1 |
| Acidobacteriota | *Granulicella paludicola* | WP_263378267.1 | WP_263379900.1 | WP_263379336.1 |
| Acidobacteriota | *Granulicella pectinivorans* | WP_089840539.1 | WP_089838949.1 | WP_089840336.1 |
| Acidobacteriota | *Granulicella rosea* | WP_089407803.1 | WP_089408523.1 | WP_089407018.1 |
| Acidobacteriota | *Granulicella sibirica* | WP_128913431.1 | WP_128911222.1 | WP_128913874.1 |
| Acidobacteriota | *Granulicella sp. 5B5* | WP_182276896.1 | WP_182277462.1 | WP_182276653.1 |
| Acidobacteriota | *Granulicella tundricola* | WP_013579839.1 | WP_013580942.1 | WP_013580988.1 |
| Acidobacteriota | *Luteitalea pratensis* | WP_110169869.1 | WP_110169868.1 | WP_157899524.1 |
| Acidobacteriota | *Occallatibacter riparius* | WP_260791635.1 | WP_260792614.1 | WP_260793218.1 |
| Acidobacteriota | *Occallatibacter savannae* | WP_109484365.1 | WP_204101321.1 | WP_109487914.1 |
| Acidobacteriota | *Paludibaculum fermentans* | WP_194446804.1 | WP_194448789.1 | WP_194448933.1 |
| Acidobacteriota | *Paracidobacterium acidisoli* | WP_117298274.1 | WP_117297597.1 | WP_117297892.1 |
| Acidobacteriota | *Pseudacidobacterium ailaaui* | WP_044933726.1 | WP_026443242.1 | WP_026443683.1 |
| Acidobacteriota | *Pyrinomonas methylaliphatogenes* | WP_041975312.1 | WP_041975311.1 | WP_157770925.1 |
| Acidobacteriota | *Pyrinomonas sp.* | WP_352430502.1 | WP_352430500.1 | WP_352432353.1 |
| Acidobacteriota | *Silvibacterium bohemicum* | WP_050061700.1 | WP_231581164.1 | WP_050057645.1 |
| Acidobacteriota | *Silvibacterium dinghuense* | WP_129208410.1 | WP_129208146.1 | WP_129209111.1 |
| Acidobacteriota | *Silvibacterium sp.* | WP_415197828.1 | WP_415197928.1 | WP_415199972.1 |
| Acidobacteriota | *Telmatobacter bradus* | WP_420236317.1 | WP_420239077.1 | WP_420236529.1 |
| Acidobacteriota | *Telmatobacter sp. DSM 110680* | WP_348261856.1 | WP_348261883.1 | WP_348262743.1 |

| | | | | |
|---|---|---|---|---|
| Acidobacteriota | *Terracidiphilus gabretensis* | WP_058189573.1 | WP_231738106.1 | WP_231738037.1 |
| Acidobacteriota | *Terracidiphilus sp.* | WP_424170589.1 | WP_424165284.1 | WP_424166473.1 |
| Acidobacteriota | *Thermoanaerobaculum aquaticum* | WP_038046342.1 | WP_053334713.1 | WP_038046340.1 |
| Acidobacteriota | *Tunturiibacter empetritectus* | WP_183760154.1 | WP_179639254.1 | WP_353069306.1 |
| Acidobacteriota | *Tunturiibacter gelidoferens* | WP_353073468.1 | WP_183980995.1 | WP_183976422.1 |
| Acidobacteriota | *Tunturiibacter lichenicola* | WP_260737915.1 | WP_260734640.1 | WP_260738339.1 |
| Acidobacteriota | *Tunturiibacter psychrotolerans* | WP_429633432.1 | WP_353063468.1 | WP_429629698.1 |
| Bacillota | *Abyssicoccus albus* | WP_123807515.1 | WP_123808477.1 | WP_077140460.1 |
| Bacillota | *Acetohalobium arabaticum* | WP_013277297.1 | WP_013277296.1 | WP_013277295.1 |
| Bacillota | *Acidaminobacter sp.* | WP_291400587.1 | WP_291404499.1 | WP_300687168.1 |
| Bacillota | *Aciduricibacillus chroicocephali* | WP_348029743.1 | WP_348026512.1 | WP_348029467.1 |
| Bacillota | *Alicyclobacillus acidiphilus* | WP_067618599.1 | WP_067618720.1 | WP_067620725.1 |
| Bacillota | *Alicyclobacillus acidocaldarius* | WP_012810972.1 | WP_012811345.1 | WP_012810636.1 |
| Bacillota | *Alicyclobacillus acidoterrestris* | WP_021296764.1 | WP_021297575.1 | WP_021297415.1 |
| Bacillota | *Alicyclobacillus cellulosilyticus* | WP_188882258.1 | WP_188883153.1 | WP_188880771.1 |
| Bacillota | *Alicyclobacillus contaminans* | WP_026974667.1 | WP_026975382.1 | WP_156900185.1 |
| Bacillota | *Alicyclobacillus cycloheptanicus* | WP_274455965.1 | WP_274454683.1 | WP_274456251.1 |
| Bacillota | *Alicyclobacillus dauci* | WP_268042068.1 | WP_268043292.1 | WP_268042825.1 |
| Bacillota | *Alicyclobacillus fastidiosus* | WP_275476750.1 | WP_275473714.1 | WP_268004344.1 |
| Bacillota | *Alicyclobacillus ferrooxydans* | WP_054968087.1 | WP_054968204.1 | WP_054967649.1 |
| Bacillota | *Alicyclobacillus fodiniaquatilis* | WP_377942297.1 | WP_377940631.1 | WP_377945233.1 |
| Bacillota | *Alicyclobacillus fructus* | WP_206830432.1 | WP_206832021.1 | WP_206831247.1 |
| Bacillota | *Alicyclobacillus herbarius* | WP_026962492.1 | WP_026961893.1 | WP_026961655.1 |
| Bacillota | *Alicyclobacillus hesperidum* | WP_281758772.1 | WP_100218590.1 | WP_058094035.1 |
| Bacillota | *Alicyclobacillus kakegawensis* | WP_067935298.1 | WP_067931723.1 | WP_067931593.1 |
| Bacillota | *Alicyclobacillus macrosporangiidus* | WP_303798311.1 | WP_074948571.1 | WP_074950252.1 |
| Bacillota | *Alicyclobacillus mali (ex Roth et al. 2021)* | WP_195867399.1 | WP_067847766.1 | WP_067850001.1 |
| Bacillota | *Alicyclobacillus mengziensis* | WP_206655069.1 | WP_206658284.1 | WP_206658773.1 |

| | | | | |
|---|---|---|---|---|
| Bacillota | *Alicyclobacillus pomorum* | WP_026965004.1 | WP_035466845.1 | WP_051375186.1 |
| Bacillota | *Alicyclobacillus sacchari* | WP_134158628.1 | WP_134159108.1 | WP_134159532.1 |
| Bacillota | *Alicyclobacillus sendaiensis* | WP_062305587.1 | WP_283202282.1 | WP_062307820.1 |
| Bacillota | *Alicyclobacillus shizuokensis* | WP_067929041.1 | WP_067927753.1 | WP_067922161.1 |
| Bacillota | *Alicyclobacillus suci* | WP_146824268.1 | WP_146822294.1 | WP_236018018.1 |
| Bacillota | *Alicyclobacillus tolerans* | WP_236811744.1 | WP_236812631.1 | WP_236813424.1 |
| Bacillota | *Alicyclobacillus vulcanalis* | WP_076347670.1 | WP_076346339.1 | WP_076347978.1 |
| Bacillota | *Aliibacillus thermotolerans* | WP_270897557.1 | WP_270897558.1 | WP_270898399.1 |
| Bacillota | *Aliicoccus persicus* | WP_091473304.1 | WP_091475302.1 | WP_091472699.1 |
| Bacillota | *Alkalibacillus aidingensis* | WP_188206003.1 | WP_188206685.1 | WP_188205716.1 |
| Bacillota | *Alkalibacillus almallahensis* | WP_167262394.1 | WP_167262627.1 | WP_167259867.1 |
| Bacillota | *Alkalibacillus filiformis* | WP_307066048.1 | WP_307069212.1 | WP_307065437.1 |
| Bacillota | *Alkalibacillus flavidus* | WP_354218829.1 | WP_354219555.1 | WP_354218417.1 |
| Bacillota | *Alkalibacillus haloalkaliphilus* | WP_146814819.1 | WP_146817884.1 | WP_317093364.1 |
| Bacillota | *Alkalibacillus salilacus* | WP_306978150.1 | WP_306976254.1 | WP_306975461.1 |
| Bacillota | *Alkalibacillus silvisoli* | WP_343782538.1 | WP_343785100.1 | WP_343782819.1 |
| Bacillota | *Alkalibacillus sp. S2W* | WP_411953315.1 | WP_411953871.1 | WP_411955377.1 |
| Bacillota | *Alkalicella caledoniensis* | WP_213168081.1 | WP_213168082.1 | WP_213168084.1 |
| Bacillota | *Alkalicoccobacillus gibsonii* | WP_369875514.1 | WP_343130382.1 | WP_203087334.1 |
| Bacillota | *Alkalicoccobacillus murimartini* | WP_306979994.1 | WP_306979993.1 | WP_306980876.1 |
| Bacillota | *Alkalicoccobacillus plakortidis* | WP_251603919.1 | WP_251603916.1 | WP_251604935.1 |
| Bacillota | *Alkalicoccobacillus porphyridii* | WP_143848168.1 | WP_143848167.1 | WP_143848764.1 |
| Bacillota | *Alkalicoccus chagannorensis* | WP_026696511.1 | WP_026696512.1 | WP_026698190.1 |
| Bacillota | *Alkalicoccus daliensis* | WP_090840435.1 | WP_090840437.1 | WP_090839767.1 |
| Bacillota | *Alkalicoccus halolimnae* | WP_147804768.1 | WP_147804769.1 | WP_147802416.1 |
| Bacillota | *Alkalicoccus luteus* | WP_168008843.1 | WP_168008845.1 | WP_168004737.1 |
| Bacillota | *Alkalicoccus saliphilus* | WP_107582712.1 | WP_107582714.1 | WP_107583975.1 |
| Bacillota | *Alkalicoccus urumqiensis* | WP_105957872.1 | WP_105957871.1 | WP_105959832.1 |

| | | | | |
|---|---|---|---|---|
| Bacillota | *Alkalihalobacillus alcalophilus* | WP_003322410.1 | WP_003322411.1 | WP_003324267.1 |
| Bacillota | *Alkalihalobacillus algicola* | WP_224880004.1 | WP_224880007.1 | WP_224881858.1 |
| Bacillota | *Alkalihalobacillus deserti* | WP_227936217.1 | WP_227936216.1 | WP_227935718.1 |
| Bacillota | *Alkalihalobacillus hemicentroti* | WP_301550589.1 | WP_301551104.1 | WP_301552758.1 |
| Bacillota | *Alkalihalobacillus macyae* | WP_048309264.1 | WP_305911367.1 | WP_305913697.1 |
| Bacillota | *Alkalihalobacillus pseudalcaliphilus* | WP_047989292.1 | WP_047989293.1 | WP_047987769.1 |
| Bacillota | *Alkalihalobacillus sp. AL-G* | WP_304977622.1 | WP_304977624.1 | WP_304980912.1 |
| Bacillota | *Alkalihalobacillus trypoxylicola* | WP_045484026.1 | WP_061948118.1 | WP_061947378.1 |
| Bacillota | *Alkalihalobacterium alkalinitrilicum* | WP_078428479.1 | WP_078429650.1 | WP_158211770.1 |
| Bacillota | *Alkalihalobacterium bogoriense* | WP_035178327.1 | WP_035178325.1 | WP_026674384.1 |
| Bacillota | *Alkalihalobacterium chitinilyticum* | WP_275116532.1 | WP_275119241.1 | WP_275117687.1 |
| Bacillota | *Alkalihalobacterium elongatum* | WP_216829224.1 | WP_216830119.1 | WP_216830881.1 |
| Bacillota | *Alkalihalobacterium sp. APHAB7* | WP_418199901.1 | WP_418197894.1 | WP_418198652.1 |
| Bacillota | *Alkalihalophilus lindianensis* | WP_317120797.1 | WP_317120796.1 | WP_317122523.1 |
| Bacillota | *Alkalihalophilus marmarensis* | WP_326238215.1 | WP_022627179.1 | WP_022628259.1 |
| Bacillota | *Alkalihalophilus pseudofirmus* | WP_323465924.1 | WP_012958526.1 | WP_012959498.1 |
| Bacillota | *Alkalihalophilus sp. As8PL* | WP_368504480.1 | WP_368504481.1 | WP_368503982.1 |
| Bacillota | *Alteribacillus bidgolensis* | WP_091579714.1 | WP_245917746.1 | WP_091582817.1 |
| Bacillota | *Alteribacillus iranensis* | WP_091656672.1 | WP_091656669.1 | WP_091657528.1 |
| Bacillota | *Alteribacillus persepolensis* | WP_091270857.1 | WP_245705149.1 | WP_091270224.1 |
| Bacillota | *Alteribacillus sp. HJP-4* | WP_375534816.1 | WP_375534815.1 | WP_375533795.1 |
| Bacillota | *Alteribacter aurantiacus* | WP_026689511.1 | WP_026689510.1 | WP_026690006.1 |
| Bacillota | *Alteribacter keqinensis* | WP_122896690.1 | WP_122896689.1 | WP_122897211.1 |
| Bacillota | *Alteribacter lacisalsi* | WP_110518270.1 | WP_110518271.1 | WP_110517475.1 |
| Bacillota | *Alteribacter natronophilus* | WP_138809359.1 | WP_138809360.1 | WP_138808907.1 |
| Bacillota | *Alteribacter populi* | WP_096434897.1 | WP_096434896.1 | WP_157811962.1 |
| Bacillota | *Alteribacter salitolerans* | WP_205165103.1 | WP_205165104.1 | WP_205166118.1 |
| Bacillota | *Ammoniphilus oxalaticus* | WP_120188418.1 | WP_120188417.1 | WP_120189348.1 |

| | | | | |
|---|---|---|---|---|
| Bacillota | *Ammoniphilus resinae* | WP_209811430.1 | WP_209811431.1 | WP_209808725.1 |
| Bacillota | *Ammoniphilus sp. CFH 90114* | WP_129198961.1 | WP_129198958.1 | WP_164984851.1 |
| Bacillota | *Amphibacillus cookii* | WP_204439232.1 | WP_204437696.1 | WP_204435918.1 |
| Bacillota | *Amphibacillus jilinensis* | WP_017472183.1 | WP_017472445.1 | WP_017471791.1 |
| Bacillota | *Amphibacillus marinus* | WP_091493446.1 | WP_091499335.1 | WP_091494518.1 |
| Bacillota | *Amphibacillus sediminis* | WP_067836485.1 | WP_067840260.1 | WP_067838861.1 |
| Bacillota | *Anaerobacillus alkalilacustris* | WP_071309040.1 | WP_071308954.1 | WP_071309864.1 |
| Bacillota | *Anaerobacillus alkaliphilus* | WP_129076319.1 | WP_129076318.1 | WP_129079471.1 |
| Bacillota | *Anaerobacillus arseniciselenatis* | WP_071311749.1 | WP_071311750.1 | WP_071314187.1 |
| Bacillota | *Anaerobacillus isosaccharinicus* | WP_071319369.1 | WP_108721444.1 | WP_182081065.1 |
| Bacillota | *Anaerobacillus sp.* | WP_409852530.1 | WP_409852635.1 | WP_409849617.1 |
| Bacillota | *Anaerobranca californiensis* | WP_072908573.1 | WP_084672538.1 | WP_072908575.1 |
| Bacillota | *Anaerobranca gottschalkii* | WP_091349737.1 | WP_091349735.1 | WP_091349733.1 |
| Bacillota | *Aneurinibacillus aneurinilyticus* | WP_276912594.1 | WP_168974337.1 | WP_021624582.1 |
| Bacillota | *Aneurinibacillus migulanus* | WP_043067898.1 | WP_235355982.1 | WP_043066087.1 |
| Bacillota | *Aneurinibacillus soli* | WP_096466561.1 | WP_096466562.1 | WP_096464848.1 |
| Bacillota | *Aneurinibacillus sp. Ricciae_BoGa-3* | WP_272442497.1 | WP_272442496.1 | WP_272439498.1 |
| Bacillota | *Aneurinibacillus terranovensis* | WP_027417519.1 | WP_035102034.1 | WP_027414484.1 |
| Bacillota | *Aneurinibacillus tyrosinisolvens* | WP_047152598.1 | WP_047152599.1 | WP_052947533.1 |
| Bacillota | *Aneurinibacillus uraniidurans* | WP_272560005.1 | WP_272560006.1 | WP_272562398.1 |
| Bacillota | *Anoxybacillus ayderensis* | WP_085788008.1 | WP_327867808.1 | WP_181520507.1 |
| Bacillota | *Anoxybacillus calidus* | WP_181535810.1 | WP_181535808.1 | WP_181536607.1 |
| Bacillota | *Anoxybacillus flavithermus* | WP_006319484.1 | WP_049754019.1 | WP_003395102.1 |
| Bacillota | *Anoxybacillus mongoliensis* | WP_183243126.1 | WP_246346286.1 | WP_183241713.1 |
| Bacillota | *Anoxybacillus pushchinoensis* | WP_091702736.1 | WP_091702733.1 | WP_091703699.1 |
| Bacillota | *Anoxybacillus sp.* | WP_297990735.1 | WP_297990736.1 | WP_297992505.1 |
| Bacillota | *Anoxybacillus suryakundensis* | WP_032099862.1 | WP_055440630.1 | WP_055441894.1 |
| Bacillota | *Anoxybacillus tengchongensis* | WP_183246662.1 | WP_246348803.1 | WP_183247171.1 |

| | | | | |
|---|---|---|---|---|
| Bacillota | *Anoxybacillus thermarum* | WP_043968235.1 | WP_235341152.1 | WP_043963574.1 |
| Bacillota | *Anoxybacter fermentans* | WP_127015566.1 | WP_205665684.1 | WP_127016546.1 |
| Bacillota | *Anoxybacteroides amylolyticum* | WP_066325763.1 | WP_066325769.1 | WP_066323566.1 |
| Bacillota | *Anoxybacteroides rupiense* | WP_212387093.1 | WP_183186419.1 | WP_183186293.1 |
| Bacillota | *Anoxybacteroides tepidamans* | WP_027408554.1 | WP_183251100.1 | WP_183252222.1 |
| Bacillota | *Anoxybacteroides voinovskiense* | WP_183183362.1 | WP_183183363.1 | WP_183182980.1 |
| Bacillota | *Aquibacillus albus* | WP_204497294.1 | WP_204502172.1 | WP_239584314.1 |
| Bacillota | *Aquibacillus halophilus* | WP_153736072.1 | WP_338079246.1 | WP_153735819.1 |
| Bacillota | *Aquibacillus kalidii* | WP_186577852.1 | WP_186580881.1 | WP_186577561.1 |
| Bacillota | *Aquibacillus koreensis* | WP_259871040.1 | WP_259870913.1 | WP_259872153.1 |
| Bacillota | *Aquibacillus rhizosphaerae* | WP_285931966.1 | WP_285930127.1 | WP_285932499.1 |
| Bacillota | *Aquibacillus saliphilus* | WP_226034707.1 | WP_226036032.1 | WP_226037929.1 |
| Bacillota | *Aquibacillus salsiterrae* | WP_272444767.1 | WP_272445133.1 | WP_272445492.1 |
| Bacillota | *Aquibacillus sediminis* | WP_138417006.1 | WP_138418335.1 | WP_138415026.1 |
| Bacillota | *Aquisalibacillus elongatus* | WP_124220368.1 | WP_124221327.1 | WP_124220187.1 |
| Bacillota | *Aureibacillus halotolerans* | WP_133579378.1 | WP_133579379.1 | WP_133579192.1 |
| Bacillota | *Bacillus aerius* | WP_309172889.1 | WP_343489429.1 | WP_343489326.1 |
| Bacillota | *Bacillus aerolatus* | WP_191991728.1 | WP_152149454.1 | WP_152149950.1 |
| Bacillota | *Bacillus albus* | WP_166703425.1 | WP_130067555.1 | WP_166701986.1 |
| Bacillota | *Bacillus alkalicellulosilyticus* | WP_078553300.1 | WP_078556248.1 | WP_078554865.1 |
| Bacillota | *Bacillus altitudinis* | WP_212047044.1 | WP_377863049.1 | WP_268449497.1 |
| Bacillota | *Bacillus alveayuensis* | WP_044749095.1 | WP_044748078.1 | WP_044893864.1 |
| Bacillota | *Bacillus amyloliquefaciens* | WP_013352659.1 | WP_132105281.1 | WP_416522088.1 |
| Bacillota | *Bacillus andreraoultii* | WP_033828563.1 | WP_033827693.1 | WP_033829314.1 |
| Bacillota | *Bacillus anthracis* | WP_160832241.1 | WP_042511954.1 | WP_420952557.1 |
| Bacillota | *Bacillus aquiflavi* | WP_163239921.1 | WP_163239918.1 | WP_163242269.1 |
| Bacillota | *Bacillus arachidis* | WP_208016830.1 | WP_208016829.1 | WP_286015891.1 |
| Bacillota | *Bacillus atrophaeus* | WP_219946263.1 | WP_268524735.1 | WP_268512546.1 |

| | | | | |
|---|---|---|---|---|
| Bacillota | *Bacillus australimaris* | WP_413597544.1 | WP_413597545.1 | WP_060699163.1 |
| Bacillota | *Bacillus badius* | WP_223661648.1 | WP_063385664.1 | WP_223663117.1 |
| Bacillota | *Bacillus benzoevorans* | WP_184523980.1 | WP_184525618.1 | WP_184521970.1 |
| Bacillota | *Bacillus bingmayongensis* | WP_017152494.1 | WP_017152495.1 | WP_221783100.1 |
| Bacillota | *Bacillus cabrialesii* | WP_103746458.1 | WP_103746457.1 | WP_129505784.1 |
| Bacillota | *Bacillus canaveralius* | WP_101575838.1 | WP_180956474.1 | WP_101576518.1 |
| Bacillota | *Bacillus carboniphilus* | WP_343803993.1 | WP_343804291.1 | WP_343799035.1 |
| Bacillota | *Bacillus cereus* | WP_098540396.1 | WP_336451787.1 | WP_098491221.1 |
| Bacillota | *Bacillus changyiensis* | WP_270798899.1 | WP_271339824.1 | WP_270799347.1 |
| Bacillota | *Bacillus chungangensis* | WP_307229127.1 | WP_370875661.1 | WP_307231862.1 |
| Bacillota | *Bacillus cihuensis* | WP_028391531.1 | WP_028391532.1 | WP_028392954.1 |
| Bacillota | *Bacillus clarus* | WP_042982873.1 | WP_042982872.1 | WP_042984148.1 |
| Bacillota | *Bacillus coahuilensis* | WP_059282924.1 | WP_059351084.1 | WP_059350700.1 |
| Bacillota | *Bacillus cytotoxicus* | WP_251226243.1 | WP_251226244.1 | WP_087095433.1 |
| Bacillota | *Bacillus dakarensis* | WP_077213779.1 | WP_077215132.1 | WP_077211426.1 |
| Bacillota | *Bacillus daqingensis* | WP_377908507.1 | WP_377908508.1 | WP_377908126.1 |
| Bacillota | *Bacillus dicomae* | WP_140969083.1 | WP_140969082.1 | WP_140970586.1 |
| Bacillota | *Bacillus ectoiniformans* | WP_204553449.1 | WP_204553281.1 | WP_204552057.1 |
| Bacillota | *Bacillus enclensis* | WP_058297630.1 | WP_336511350.1 | WP_058298361.1 |
| Bacillota | *Bacillus fonticola* | WP_170007451.1 | WP_170007452.1 | WP_170006931.1 |
| Bacillota | *Bacillus fungorum* | WP_353767081.1 | WP_098759501.1 | WP_353768741.1 |
| Bacillota | *Bacillus gaemokensis* | WP_033676929.1 | WP_033676931.1 | WP_033672194.1 |
| Bacillota | *Bacillus glycinifermentans* | WP_057957431.1 | WP_232517718.1 | WP_048353108.1 |
| Bacillota | *Bacillus haikouensis* | WP_172252472.1 | WP_172252470.1 | WP_172248231.1 |
| Bacillota | *Bacillus halotolerans* | WP_105954959.1 | WP_105954958.1 | WP_101861185.1 |
| Bacillota | *Bacillus haynesii* | WP_268344774.1 | WP_411809870.1 | WP_268307532.1 |
| Bacillota | *Bacillus hominis* | WP_289358627.1 | WP_002117713.1 | WP_289360060.1 |
| Bacillota | *Bacillus horti* | WP_307392687.1 | WP_307392685.1 | WP_307390445.1 |

| | | | | |
|---|---|---|---|---|
| Bacillota | *Bacillus inaquosorum* | WP_327810411.1 | WP_019258742.1 | WP_268279082.1 |
| Bacillota | *Bacillus infantis* | WP_257991664.1 | WP_426878124.1 | WP_224904057.1 |
| Bacillota | *Bacillus kandeliae* | WP_338749638.1 | WP_338749640.1 | WP_338753894.1 |
| Bacillota | *Bacillus kexueae* | WP_243386618.1 | WP_243386617.1 | WP_243386721.1 |
| Bacillota | *Bacillus licheniformis* | WP_145614123.1 | WP_422110169.1 | WP_420951217.1 |
| Bacillota | *Bacillus litorisediminis* | WP_243296802.1 | WP_313897641.1 | WP_243298167.1 |
| Bacillota | *Bacillus lumedeiriae* | WP_404313695.1 | WP_404313696.1 | WP_404314937.1 |
| Bacillota | *Bacillus luti* | WP_156576229.1 | WP_312502853.1 | WP_427229069.1 |
| Bacillota | *Bacillus manliponensis* | WP_034636717.1 | WP_369900347.1 | WP_034635795.1 |
| Bacillota | *Bacillus marasmi* | WP_147533767.1 | WP_147533768.1 | WP_147531919.1 |
| Bacillota | *Bacillus marinisedimentorum* | WP_070121118.1 | WP_070121796.1 | WP_169823916.1 |
| Bacillota | *Bacillus massiliglaciei* | WP_110927635.1 | WP_110927634.1 | WP_110929323.1 |
| Bacillota | *Bacillus massiliigorillae* | WP_042346443.1 | WP_042346440.1 | WP_042349036.1 |
| Bacillota | *Bacillus massilionigeriensis* | WP_075982109.1 | WP_075982110.1 | WP_075981027.1 |
| Bacillota | *Bacillus mediterraneensis* | WP_071459948.1 | WP_071459949.1 | WP_071459349.1 |
| Bacillota | *Bacillus mesophilum* | WP_151573066.1 | WP_151573671.1 | WP_151571977.1 |
| Bacillota | *Bacillus mesophilus* | WP_163178608.1 | WP_163178610.1 | WP_163176861.1 |
| Bacillota | *Bacillus methanolicus* | WP_274854886.1 | WP_004435240.1 | WP_003348869.1 |
| Bacillota | *Bacillus mobilis* | WP_426833822.1 | WP_426833821.1 | WP_427265026.1 |
| Bacillota | *Bacillus mojavensis* | WP_326192655.1 | WP_326334022.1 | WP_326223470.1 |
| Bacillota | *Bacillus mycoides* | WP_240380238.1 | WP_256484637.1 | WP_215551778.1 |
| Bacillota | *Bacillus nakamurai* | WP_229976555.1 | WP_229976556.1 | WP_229977878.1 |
| Bacillota | *Bacillus ndiopicus* | WP_042478014.1 | WP_042470698.1 | WP_042471675.1 |
| Bacillota | *Bacillus niameyensis* | WP_062104823.1 | WP_062104822.1 | WP_062105168.1 |
| Bacillota | *Bacillus nitratireducens* | WP_001237358.1 | WP_000015677.1 | WP_097809294.1 |
| Bacillota | *Bacillus norwichensis* | WP_191811341.1 | WP_191811340.1 | WP_191809666.1 |
| Bacillota | *Bacillus oleivorans* | WP_097157564.1 | WP_097157563.1 | WP_097159486.1 |
| Bacillota | *Bacillus pacificus* | WP_208754974.1 | WP_228920269.1 | WP_173601325.1 |

| | | | | |
|---|---|---|---|---|
| Bacillota | *Bacillus pakistanensis* | WP_205168494.1 | WP_205168495.1 | WP_205173913.1 |
| Bacillota | *Bacillus paralicheniformis* | WP_143260978.1 | WP_020451935.1 | WP_205802241.1 |
| Bacillota | *Bacillus paramycoides* | WP_098472977.1 | WP_328064577.1 | WP_342715544.1 |
| Bacillota | *Bacillus paranthracis* | WP_412838602.1 | WP_229136520.1 | WP_270363186.1 |
| Bacillota | *Bacillus pinisoli* | WP_246942140.1 | WP_246942143.1 | WP_246940210.1 |
| Bacillota | *Bacillus piscicola* | WP_240375564.1 | WP_338021128.1 | WP_240375160.1 |
| Bacillota | *Bacillus proteolyticus* | WP_071743869.1 | WP_340506901.1 | WP_327817159.1 |
| Bacillota | *Bacillus pseudomycoides* | WP_098644666.1 | WP_098165788.1 | WP_243523993.1 |
| Bacillota | *Bacillus pumilus* | WP_226509125.1 | WP_224925693.1 | WP_409481788.1 |
| Bacillota | *Bacillus rubiinfantis* | WP_042355288.1 | WP_042355287.1 | WP_042356345.1 |
| Bacillota | *Bacillus rugosus* | WP_166851399.1 | WP_166851395.1 | WP_326220590.1 |
| Bacillota | *Bacillus safensis* | WP_212043074.1 | WP_224426689.1 | WP_377970795.1 |
| Bacillota | *Bacillus salacetis* | WP_119547185.1 | WP_421379192.1 | WP_421384454.1 |
| Bacillota | *Bacillus salipaludis* | WP_406581955.1 | WP_133335929.1 | WP_406580921.1 |
| Bacillota | *Bacillus salitolerans* | WP_377929730.1 | WP_377929731.1 | WP_377926817.1 |
| Bacillota | *Bacillus seohaeanensis* | WP_377932177.1 | WP_377932179.1 | WP_377937664.1 |
| Bacillota | *Bacillus shivajii* | WP_226517787.1 | WP_226518265.1 | WP_226514851.1 |
| Bacillota | *Bacillus siamensis* | WP_415273134.1 | WP_326352018.1 | WP_269775133.1 |
| Bacillota | *Bacillus sinesaloumensis* | WP_077621011.1 | WP_077618088.1 | WP_077617725.1 |
| Bacillota | *Bacillus solimangrovi* | WP_069716330.1 | WP_083249078.1 | WP_069716086.1 |
| Bacillota | *Bacillus solitudinis* | WP_100407236.1 | WP_100408361.1 | WP_100404574.1 |
| Bacillota | *Bacillus songklensis* | WP_377915469.1 | WP_377915472.1 | WP_377912371.1 |
| Bacillota | *Bacillus sonorensis* | WP_326187081.1 | WP_276561419.1 | WP_077736879.1 |
| Bacillota | *Bacillus sp. 03113* | WP_141433313.1 | WP_141433314.1 | WP_141431292.1 |
| Bacillota | *Bacillus spizizenii* | WP_014114192.1 | WP_019714760.1 | WP_268470174.1 |
| Bacillota | *Bacillus spongiae* | WP_336586651.1 | WP_336586785.1 | WP_336585491.1 |
| Bacillota | *Bacillus stercoris* | WP_071580476.1 | WP_136654146.1 | WP_326274014.1 |
| Bacillota | *Bacillus suaedae* | WP_210596305.1 | WP_210596304.1 | WP_210597829.1 |

| | | | | |
|---|---|---|---|---|
| Bacillota | *Bacillus suaedaesalsae* | WP_204205316.1 | WP_204205317.1 | WP_204203888.1 |
| Bacillota | *Bacillus subtilis* | WP_041336364.1 | WP_336805235.1 | WP_032725648.1 |
| Bacillota | *Bacillus swezeyi* | WP_414555380.1 | WP_239693662.1 | WP_148955705.1 |
| Bacillota | *Bacillus taeanensis* | WP_113805161.1 | WP_113805162.1 | WP_181833047.1 |
| Bacillota | *Bacillus tequilensis* | WP_174227222.1 | WP_167872758.1 | WP_174228065.1 |
| Bacillota | *Bacillus testis* | WP_050615402.1 | WP_050615403.1 | WP_050614681.1 |
| Bacillota | *Bacillus thermotolerans* | WP_175286644.1 | WP_040037628.1 | WP_039231947.1 |
| Bacillota | *Bacillus thuringiensis* | WP_263701529.1 | WP_131934799.1 | WP_197230479.1 |
| Bacillota | *Bacillus tianshenii* | WP_204414591.1 | WP_224840203.1 | WP_224840737.1 |
| Bacillota | *Bacillus timonensis* | WP_136377963.1 | WP_010281490.1 | WP_136378255.1 |
| Bacillota | *Bacillus toyonensis* | WP_098637734.1 | WP_390190602.1 | WP_098837687.1 |
| Bacillota | *Bacillus tropicus* | WP_340031223.1 | WP_337679143.1 | WP_227571775.1 |
| Bacillota | *Bacillus tuaregi* | WP_071395570.1 | WP_071396756.1 | WP_071395437.1 |
| Bacillota | *Bacillus vallismortis* | WP_252685135.1 | WP_394716094.1 | WP_367387382.1 |
| Bacillota | *Bacillus velezensis* | WP_326155360.1 | WP_138092526.1 | WP_041481861.1 |
| Bacillota | *Bacillus weihaiensis* | WP_273125988.1 | WP_273125986.1 | WP_273125371.1 |
| Bacillota | *Bacillus wiedmannii* | WP_098047379.1 | WP_262745050.1 | WP_098707497.1 |
| Bacillota | *Bacillus wudalianchiensis* | WP_065409237.1 | WP_065409236.1 | WP_065411549.1 |
| Bacillota | *Bacillus xiapuensis* | WP_198508479.1 | WP_100332054.1 | WP_100330817.1 |
| Bacillota | *Bacillus yunxiaonensis* | WP_336482823.1 | WP_336482822.1 | WP_336483199.1 |
| Bacillota | *Baia soyae* | WP_131848545.1 | WP_131848544.1 | WP_131847547.1 |
| Bacillota | *Bhargavaea beijingensis* | WP_092095105.1 | WP_092097408.1 | WP_264556923.1 |
| Bacillota | *Bhargavaea cecembensis* | WP_063179153.1 | WP_040227486.1 | WP_197462205.1 |
| Bacillota | *Bhargavaea ginsengi* | WP_251239798.1 | WP_251238947.1 | WP_177168219.1 |
| Bacillota | *Bhargavaea massiliensis* | WP_213423159.1 | WP_213422174.1 | WP_213423353.1 |
| Bacillota | *Bhargavaea ullalensis* | WP_354196044.1 | WP_354197139.1 | WP_354197405.1 |
| Bacillota | Brevibacillus agri | WP_327944617.1 | WP_197187672.1 | WP_005833010.1 |
| Bacillota | Brevibacillus antibioticus | WP_137029308.1 | WP_137029309.1 | WP_137028103.1 |

| | | | | |
|---|---|---|---|---|
| Bacillota | *Brevibacillus borstelensis* | WP_171506301.1 | WP_311067562.1 | WP_327925907.1 |
| Bacillota | *Brevibacillus brevis* | WP_048032664.1 | WP_289677705.1 | WP_310764353.1 |
| Bacillota | *Brevibacillus centrosporus* | WP_400582021.1 | WP_122961388.1 | WP_327930787.1 |
| Bacillota | *Brevibacillus choshinensis* | WP_055743623.1 | WP_055743622.1 | WP_203255281.1 |
| Bacillota | *Brevibacillus composti* | WP_198826360.1 | WP_198826361.1 | WP_198829395.1 |
| Bacillota | *Brevibacillus daliensis* | WP_232696377.1 | WP_232696376.1 | WP_232698373.1 |
| Bacillota | *Brevibacillus dissolubilis* | WP_139489136.1 | WP_139489137.1 | WP_139489967.1 |
| Bacillota | *Brevibacillus fluminis* | WP_122920747.1 | WP_122920746.1 | WP_122916979.1 |
| Bacillota | *Brevibacillus formosus* | WP_197245195.1 | WP_047069015.1 | WP_197244069.1 |
| Bacillota | *Brevibacillus fortis* | WP_327950004.1 | WP_409178656.1 | WP_409174669.1 |
| Bacillota | *Brevibacillus fulvus* | WP_204516473.1 | WP_204516472.1 | WP_204517035.1 |
| Bacillota | *Brevibacillus gelatini* | WP_122904259.1 | WP_122904258.1 | WP_122906096.1 |
| Bacillota | *Brevibacillus ginsengisoli* | WP_426927693.1 | WP_426927692.1 | WP_426928466.1 |
| Bacillota | *Brevibacillus humidisoli* | WP_230046444.1 | WP_230046959.1 | WP_230045536.1 |
| Bacillota | *Brevibacillus invocatus* | WP_122908283.1 | WP_122908779.1 | WP_251243155.1 |
| Bacillota | *Brevibacillus laterosporus* | WP_301802748.1 | WP_096887578.1 | WP_277545643.1 |
| Bacillota | *Brevibacillus marinus* | WP_126427242.1 | WP_126427244.1 | WP_126425988.1 |
| Bacillota | *Brevibacillus massiliensis* | WP_019122419.1 | WP_019122420.1 | WP_019120713.1 |
| Bacillota | *Brevibacillus migulae* | WP_134686654.1 | WP_134686655.1 | WP_134687458.1 |
| Bacillota | *Brevibacillus nitrificans* | WP_328204720.1 | WP_310227962.1 | WP_310236248.1 |
| Bacillota | *Brevibacillus panacihumi* | WP_427329340.1 | WP_122912325.1 | WP_122912984.1 |
| Bacillota | *Brevibacillus parabrevis* | WP_328147717.1 | WP_063227869.1 | WP_122964991.1 |
| Bacillota | *Brevibacillus porteri* | WP_106835973.1 | WP_376898458.1 | WP_106835688.1 |
| Bacillota | *Brevibacillus reuszeri* | WP_376908201.1 | WP_312108564.1 | WP_312114022.1 |
| Bacillota | *Brevibacillus ruminantium* | WP_251870562.1 | WP_251870563.1 | WP_251874646.1 |
| Bacillota | *Brevibacillus sp. B_LB10_24* | WP_408108360.1 | WP_408108361.1 | WP_408106202.1 |
| Bacillota | *Brevibacillus thermoruber* | WP_271139233.1 | WP_029099570.1 | WP_035295256.1 |
| Bacillota | *Caenibacillus caldisaponilyticus* | WP_077616334.1 | WP_077616392.1 | WP_077614480.1 |

| | | | | |
|---|---|---|---|---|
| Bacillota | *Caldalkalibacillus mannanilyticus* | WP_025026795.1 | WP_025026794.1 | WP_025028057.1 |
| Bacillota | *Caldalkalibacillus salinus* | WP_202079974.1 | WP_202079975.1 | WP_202076947.1 |
| Bacillota | *Caldalkalibacillus thermarum* | WP_007505592.1 | WP_042685701.1 | WP_188622502.1 |
| Bacillota | *Caldalkalibacillus uzonensis* | WP_307334880.1 | WP_307334882.1 | WP_307338827.1 |
| Bacillota | *Calderihabitans maritimus* | WP_088553107.1 | WP_088554238.1 | WP_088554237.1 |
| Bacillota | *Caldibacillus debilis* | WP_276674373.1 | WP_333524186.1 | WP_061569350.1 |
| Bacillota | *Caldibacillus thermoamylovorans* | WP_227091017.1 | WP_251250642.1 | WP_251245618.1 |
| Bacillota | *Caldifermentibacillus hisashii* | WP_341288069.1 | WP_404997150.1 | WP_368997560.1 |
| Bacillota | *Caldinitratiruptor microaerophilus* | WP_264843609.1 | WP_264843608.1 | WP_264841576.1 |
| Bacillota | *Calditerricola satsumensis* | WP_054672757.1 | WP_188816842.1 | WP_188816587.1 |
| Bacillota | *Calidifontibacillus erzurumensis* | WP_173731473.1 | WP_173730150.1 | WP_173729957.1 |
| Bacillota | *Calidifontibacillus oryziterrae* | WP_017755555.1 | WP_017755556.1 | WP_017753621.1 |
| Bacillota | *Camelliibacillus cellulosilyticus* | WP_376844808.1 | WP_376844809.1 | WP_376844302.1 |
| Bacillota | *Candidatus Contubernalis alkalaceticus* | WP_241420565.1 | WP_241420566.1 | WP_241420567.1 |
| Bacillota | *Carboxydothermus ferrireducens* | WP_028052299.1 | WP_028052298.1 | WP_028052058.1 |
| Bacillota | *Carboxydothermus hydrogenoformans* | WP_011344056.1 | WP_011344055.1 | WP_011344794.1 |
| Bacillota | *Carboxydothermus islandicus* | WP_075866452.1 | WP_075866451.1 | WP_075865582.1 |
| Bacillota | *Carboxydothermus pertinax* | WP_075859531.1 | WP_075859530.1 | WP_075858767.1 |
| Bacillota | *Caryophanon latum* | WP_066465105.1 | WP_066466106.1 | WP_066463252.1 |
| Bacillota | *Caryophanon tenue* | WP_066548704.1 | WP_066547131.1 | WP_066543228.1 |
| Bacillota | *Cerasibacillus quisquiliarum* | WP_146938177.1 | WP_146934397.1 | WP_146935506.1 |
| Bacillota | *Cerasibacillus sp.* | WP_331643333.1 | WP_331642528.1 | WP_331639209.1 |
| Bacillota | *Cerasibacillus terrae* | WP_147665579.1 | WP_147666711.1 | WP_147665147.1 |
| Bacillota | *Chengkuizengella axinellae* | WP_305991132.1 | WP_305991131.1 | WP_305990434.1 |
| Bacillota | *Chengkuizengella marina* | WP_160647308.1 | WP_160647307.1 | WP_160646215.1 |
| Bacillota | *Chengkuizengella sediminis* | WP_162035760.1 | WP_162035759.1 | WP_162035053.1 |
| Bacillota | *Chengkuizengella sp. SCS-71B* | WP_349248815.1 | WP_349248814.1 | WP_349249855.1 |
| Bacillota | *Chryseomicrobium aureum* | WP_204589171.1 | WP_204589836.1 | WP_392456146.1 |

| | | | | |
|---|---|---|---|---|
| Bacillota | *Chryseomicrobium excrementi* | WP_245858730.1 | WP_100353346.1 | WP_100352793.1 |
| Bacillota | *Chryseomicrobium palamuruense* | WP_378140865.1 | WP_378140089.1 | WP_378140646.1 |
| Bacillota | *Chryseomicrobium sp. FSL W7-1435* | WP_342525698.1 | WP_342527989.1 | WP_342525912.1 |
| Bacillota | *Chungangia koreensis* | WP_378155353.1 | WP_378154905.1 | WP_378151913.1 |
| Bacillota | *Cohnella abietis* | WP_130608606.1 | WP_130608603.1 | WP_157994083.1 |
| Bacillota | *Cohnella boryungensis* | WP_204603271.1 | WP_204603272.1 | WP_204601225.1 |
| Bacillota | *Cohnella caldifontis* | WP_276351556.1 | WP_276351558.1 | WP_276355939.1 |
| Bacillota | *Cohnella candidum* | WP_123040770.1 | WP_123040771.1 | WP_164472625.1 |
| Bacillota | *Cohnella cellulosilytica* | WP_378047127.1 | WP_378047129.1 | WP_378054137.1 |
| Bacillota | *Cohnella endophytica* | WP_120974641.1 | WP_120974642.1 | WP_158602250.1 |
| Bacillota | *Cohnella faecalis* | WP_119147875.1 | WP_119147874.1 | WP_158593973.1 |
| Bacillota | *Cohnella fermenti* | WP_136370498.1 | WP_136370497.1 | WP_168735792.1 |
| Bacillota | *Cohnella ginsengisoli* | WP_277563559.1 | WP_277563560.1 | WP_277567073.1 |
| Bacillota | *Cohnella hashimotonis* | WP_282908436.1 | WP_282908435.1 | WP_282909408.1 |
| Bacillota | *Cohnella herbarum* | WP_169278750.1 | WP_169278749.1 | WP_169279893.1 |
| Bacillota | *Cohnella hongkongensis* | WP_378096723.1 | WP_378096104.1 | WP_378097115.1 |
| Bacillota | *Cohnella kolymensis* | WP_041065952.1 | WP_041065955.1 | WP_161793416.1 |
| Bacillota | *Cohnella lubricantis* | WP_185180865.1 | WP_185180864.1 | WP_185178599.1 |
| Bacillota | *Cohnella luojiensis* | WP_135150944.1 | WP_135150943.1 | WP_167746931.1 |
| Bacillota | *Cohnella lupini* | WP_115993120.1 | WP_115993121.1 | WP_115990748.1 |
| Bacillota | *Cohnella mopanensis* | WP_239614212.1 | WP_239614211.1 | WP_239617380.1 |
| Bacillota | *Cohnella nanjingensis* | WP_185671970.1 | WP_185671971.1 | WP_185671088.1 |
| Bacillota | *Cohnella panacarvi* | WP_027084921.1 | WP_084760938.1 | WP_027088337.1 |
| Bacillota | *Cohnella pontilimi* | WP_136776128.1 | WP_136776129.1 | WP_246042130.1 |
| Bacillota | *Cohnella silvisoli* | WP_232183992.1 | WP_232183994.1 | WP_232187431.1 |
| Bacillota | *Cohnella soli* | WP_378133467.1 | WP_378133469.1 | WP_378132313.1 |
| Bacillota | *Cohnella sp.* | WP_372634402.1 | WP_391571901.1 | WP_391570815.1 |
| Bacillota | *Cohnella suwonensis* | WP_209743106.1 | WP_209743108.1 | WP_378081019.1 |

| | | | | |
|---|---|---|---|---|
| Bacillota | *Cohnella terricola* | WP_144698270.1 | WP_144698268.1 | WP_186438347.1 |
| Bacillota | *Cohnella thailandensis* | WP_185120590.1 | WP_185120591.1 | WP_185121488.1 |
| Bacillota | *Cohnella thermotolerans* | WP_027093617.1 | WP_027093618.1 | WP_027091085.1 |
| Bacillota | *Cohnella xylanilytica* | WP_185134228.1 | WP_212959663.1 | WP_185139369.1 |
| Bacillota | *Cohnella yongneupensis* | WP_378113744.1 | WP_378113743.1 | WP_378114142.1 |
| Bacillota | *Cohnella zeiphila* | WP_185133116.1 | WP_185133115.1 | WP_185127330.1 |
| Bacillota | *Compostibacillus humi* | WP_188392063.1 | WP_188391814.1 | WP_188391400.1 |
| Bacillota | *Corticicoccus populi* | WP_377771250.1 | WP_377773644.1 | WP_377772180.1 |
| Bacillota | *Croceifilum oryzae* | WP_307254358.1 | WP_307254361.1 | WP_307250766.1 |
| Bacillota | *Cytobacillus citreus* | WP_213102480.1 | WP_420828161.1 | WP_213100596.1 |
| Bacillota | *Cytobacillus dafuensis* | WP_057771339.1 | WP_407643411.1 | WP_057769642.1 |
| Bacillota | *Cytobacillus depressus* | WP_151533025.1 | WP_425481971.1 | WP_151534101.1 |
| Bacillota | *Cytobacillus eiseniae* | WP_066395413.1 | WP_066395414.1 | WP_066393352.1 |
| Bacillota | *Cytobacillus firmus* | WP_258758697.1 | WP_404355338.1 | WP_335525087.1 |
| Bacillota | *Cytobacillus gottheilii* | WP_080845475.1 | WP_214478849.1 | WP_426860528.1 |
| Bacillota | *Cytobacillus horneckiae* | WP_066200529.1 | WP_412860526.1 | WP_412859853.1 |
| Bacillota | *Cytobacillus kochii* | WP_281202324.1 | WP_404292258.1 | WP_328199021.1 |
| Bacillota | *Cytobacillus luteolus* | WP_193539513.1 | WP_193539514.1 | WP_193536846.1 |
| Bacillota | *Cytobacillus massiliigabonensis* | WP_102274775.1 | WP_102274774.1 | WP_102271555.1 |
| Bacillota | *Cytobacillus oceanisediminis* | WP_144541601.1 | WP_110064210.1 | WP_144542403.1 |
| Bacillota | *Cytobacillus praedii* | WP_328165307.1 | WP_242691024.1 | WP_232426301.1 |
| Bacillota | *Cytobacillus pseudoceanisediminis* | WP_423375193.1 | WP_258743562.1 | WP_423373599.1 |
| Bacillota | *Cytobacillus purgationiresistens* | WP_307475962.1 | WP_307475964.1 | WP_307480611.1 |
| Bacillota | *Cytobacillus solani* | WP_053476782.1 | WP_056687224.1 | WP_231690049.1 |
| Bacillota | *Cytobacillus sp.* | WP_313798514.1 | WP_313798513.1 | WP_313798771.1 |
| Bacillota | *Cytobacillus spongiae* | WP_233806574.1 | WP_233806576.1 | WP_233811026.1 |
| Bacillota | *Desertibacillus haloalkaliphilus* | WP_217224796.1 | WP_217224795.1 | WP_217224580.1 |
| Bacillota | *Desmospora activa* | WP_107725504.1 | WP_245891082.1 | WP_107726050.1 |

| | | | | |
|---|---|---|---|---|
| Bacillota | *Desmospora profundinema* | WP_309861822.1 | WP_309861824.1 | WP_309862629.1 |
| Bacillota | *Desulforamulus aeronauticus* | WP_072912166.1 | WP_238456746.1 | WP_072913709.1 |
| Bacillota | *Desulforamulus aquiferis* | WP_304544453.1 | WP_304544451.1 | WP_304541554.1 |
| Bacillota | *Desulforamulus ferrireducens* | WP_077714789.1 | WP_077714790.1 | WP_077713567.1 |
| Bacillota | *Desulforamulus hydrothermalis* | WP_008413214.1 | WP_235695578.1 | WP_008412797.1 |
| Bacillota | *Desulforamulus putei* | WP_073238853.1 | WP_073238856.1 | WP_073238516.1 |
| Bacillota | *Desulforamulus reducens* | WP_011878631.1 | WP_011878632.1 | WP_011877534.1 |
| Bacillota | *Desulforamulus ruminis* | WP_013841388.1 | WP_013841387.1 | WP_013842872.1 |
| Bacillota | *Desulfosporosinus acididurans* | WP_047810799.1 | WP_047808470.1 | WP_047810809.1 |
| Bacillota | *Desulfosporosinus acidiphilus* | WP_014828645.1 | WP_041276197.1 | WP_014828658.1 |
| Bacillota | *Desulfosporosinus fructosivorans* | WP_135544966.1 | WP_135544965.1 | WP_243454315.1 |
| Bacillota | *Desulfosporosinus hippei* | WP_092328753.1 | WP_092331157.1 | WP_092328781.1 |
| Bacillota | *Desulfosporosinus lacus* | WP_073028355.1 | WP_073031454.1 | WP_073028368.1 |
| Bacillota | *Desulfosporosinus meridiei* | WP_014904613.1 | WP_014904331.1 | WP_014904626.1 |
| Bacillota | *Desulfosporosinus metallidurans* | WP_075363497.1 | WP_075365157.1 | WP_075363507.1 |
| Bacillota | *Desulfosporosinus nitroreducens* | WP_301997921.1 | WP_252470087.1 | WP_301997935.1 |
| Bacillota | *Desulfosporosinus orientis* | WP_014187046.1 | WP_014186735.1 | WP_014187063.1 |
| Bacillota | *Desulfosporosinus paludis* | WP_407307408.1 | WP_407305579.1 | WP_407307383.1 |
| Bacillota | *Desulfosporosinus sp.* | WP_291352846.1 | WP_291349225.1 | WP_298201967.1 |
| Bacillota | *Desulfosporosinus youngiae* | WP_007786549.1 | WP_007786032.1 | WP_007786564.1 |
| Bacillota | *Desulfotomaculum nigrificans* | WP_013810337.1 | WP_003543278.1 | WP_234702063.1 |
| Bacillota | *Desulfotomaculum sp. 1211_IL3151* | WP_333530773.1 | WP_333530162.1 | WP_333529912.1 |
| Bacillota | *Dethiobacter alkaliphilus* | WP_008516141.1 | WP_008516142.1 | WP_008516143.1 |
| Bacillota | *Domibacillus aminovorans* | WP_063966893.1 | WP_063975283.1 | WP_063975316.1 |
| Bacillota | *Domibacillus antri* | WP_075397513.1 | WP_425429437.1 | WP_075398213.1 |
| Bacillota | *Domibacillus enclensis* | WP_045851444.1 | WP_045852543.1 | WP_045851456.1 |
| Bacillota | *Domibacillus epiphyticus* | WP_076767700.1 | WP_083711496.1 | WP_076762848.1 |
| Bacillota | *Domibacillus indicus* | WP_251270905.1 | WP_197076570.1 | WP_046175139.1 |

| | | | | |
|---|---|---|---|---|
| Bacillota | *Domibacillus iocasae* | WP_069939096.1 | WP_069938662.1 | WP_069939483.1 |
| Bacillota | *Domibacillus mangrovi* | WP_073709983.1 | WP_073710831.1 | WP_073711090.1 |
| Bacillota | *Domibacillus robiginosus* | WP_050180706.1 | WP_050180456.1 | WP_050181536.1 |
| Bacillota | *Domibacillus sp.* | WP_309087748.1 | WP_309089351.1 | WP_309087016.1 |
| Bacillota | *Domibacillus tundrae* | WP_353913610.1 | WP_245618035.1 | WP_046179655.1 |
| Bacillota | *Ectobacillus antri* | WP_124563901.1 | WP_124563900.1 | WP_124563576.1 |
| Bacillota | *Ectobacillus funiculus* | WP_379947984.1 | WP_379947985.1 | WP_410775892.1 |
| Bacillota | *Ectobacillus panaciterrae* | WP_028402725.1 | WP_028402724.1 | WP_028398909.1 |
| Bacillota | *Ectobacillus polymachus* | WP_416827846.1 | WP_416827847.1 | WP_416826947.1 |
| Bacillota | *Ectobacillus ponti* | WP_254758693.1 | WP_254758692.1 | WP_254760171.1 |
| Bacillota | *Ectobacillus sp. sgz5001026* | WP_379970274.1 | WP_379970275.1 | WP_379967020.1 |
| Bacillota | *Edaphobacillus lindanitolerans* | WP_076756388.1 | WP_076756839.1 | WP_076756591.1 |
| Bacillota | *Evansella cellulosilytica* | WP_013488498.1 | WP_013488497.1 | WP_013489193.1 |
| Bacillota | *Evansella clarkii* | WP_088035635.1 | WP_078596894.1 | WP_078595466.1 |
| Bacillota | *Evansella halocellulosilytica* | WP_096188656.1 | WP_096188655.1 | WP_096189230.1 |
| Bacillota | *Evansella sp. LMS18* | WP_255242124.1 | WP_255242123.1 | WP_255243417.1 |
| Bacillota | *Evansella tamaricis* | WP_217069329.1 | WP_217069385.1 | WP_217066593.1 |
| Bacillota | *Evansella vedderi* | WP_307331180.1 | WP_307331183.1 | WP_307322987.1 |
| Bacillota | *Exiguobacterium algae* | WP_214826552.1 | WP_240544489.1 | WP_338082714.1 |
| Bacillota | *Exiguobacterium alkaliphilum* | WP_367660416.1 | WP_034815198.1 | WP_246514194.1 |
| Bacillota | *Exiguobacterium aurantiacum* | WP_070327419.1 | WP_255176439.1 | WP_029335464.1 |
| Bacillota | *Exiguobacterium chiriqhucha* | WP_276693499.1 | WP_276693498.1 | WP_031200163.1 |
| Bacillota | *Exiguobacterium flavidum* | WP_114570975.1 | WP_114572129.1 | WP_114571240.1 |
| Bacillota | *Exiguobacterium indicum* | WP_058713743.1 | WP_411212175.1 | WP_075642557.1 |
| Bacillota | *Exiguobacterium oxidotolerans* | WP_029330621.1 | WP_029332029.1 | WP_029330917.1 |
| Bacillota | *Exiguobacterium qingdaonense* | WP_215141475.1 | WP_215141476.1 | WP_338082249.1 |
| Bacillota | *Falsibacillus albus* | WP_121680777.1 | WP_121680776.1 | WP_121680364.1 |
| Bacillota | *Falsibacillus pallidus* | WP_114745338.1 | WP_114745337.1 | WP_114743777.1 |

| | | | | |
|---|---|---|---|---|
| Bacillota | *Ferdinandcohnia quinoae* | WP_240253376.1 | WP_240253378.1 | WP_240251968.1 |
| Bacillota | *Ferroacidibacillus organovorans* | WP_067564900.1 | WP_067560444.1 | WP_079290518.1 |
| Bacillota | *Fervidibacillus albus* | WP_420842691.1 | WP_275417391.1 | WP_275418420.1 |
| Bacillota | *Fervidibacillus halotolerans* | WP_275421935.1 | WP_275420367.1 | WP_275421882.1 |
| Bacillota | *Ferviditalea candida* | WP_371753413.1 | WP_371753412.1 | WP_371752757.1 |
| Bacillota | *Fictibacillus aquaticus* | WP_205727361.1 | WP_094250508.1 | WP_094250912.1 |
| Bacillota | *Fictibacillus arsenicus* | WP_077364071.1 | WP_077364069.1 | WP_066289143.1 |
| Bacillota | *Fictibacillus barbaricus* | WP_188402068.1 | WP_188402067.1 | WP_310261459.1 |
| Bacillota | *Fictibacillus enclensis* | WP_283889052.1 | WP_289485947.1 | WP_289485337.1 |
| Bacillota | *Fictibacillus fluitans* | WP_301163916.1 | WP_301164878.1 | WP_301164551.1 |
| Bacillota | *Fictibacillus gelatini* | WP_026676973.1 | WP_026676972.1 | WP_026675974.1 |
| Bacillota | *Fictibacillus halophilus* | WP_379834442.1 | WP_226534979.1 | WP_379833681.1 |
| Bacillota | *Fictibacillus iocasae* | WP_379746025.1 | WP_379746023.1 | WP_379749286.1 |
| Bacillota | *Fictibacillus macauensis* | WP_007200809.1 | WP_007200808.1 | WP_007202444.1 |
| Bacillota | *Fictibacillus marinisediminis* | WP_248252980.1 | WP_248252981.1 | WP_248252504.1 |
| Bacillota | *Fictibacillus nanhaiensis* | WP_251320614.1 | WP_328084593.1 | WP_251322428.1 |
| Bacillota | *Fictibacillus norfolkensis* | WP_191753992.1 | WP_191753991.1 | WP_191753699.1 |
| Bacillota | *Fictibacillus phosphorivorans* | WP_251312548.1 | WP_251312549.1 | WP_251310970.1 |
| Bacillota | *Fictibacillus solisalsi* | WP_090234070.1 | WP_090234071.1 | WP_090233142.1 |
| Bacillota | *Fictibacillus sp. FJAT-27399* | WP_062235415.1 | WP_062238660.1 | WP_062231614.1 |
| Bacillota | *Fictibacillus terranigra* | WP_290400662.1 | WP_290400663.1 | WP_290399658.1 |
| Bacillota | *Filibacter tadaridae* | WP_124070649.1 | WP_203229615.1 | WP_124071008.1 |
| Bacillota | *Filobacillus milosensis* | WP_134339040.1 | WP_134338713.1 | WP_134339379.1 |
| Bacillota | *Fodinisporobacter ferrooxydans* | WP_347436385.1 | WP_347436386.1 | WP_347435826.1 |
| Bacillota | *Fontibacillus panacisegetis* | WP_091229626.1 | WP_091229628.1 | WP_091226566.1 |
| Bacillota | *Fontibacillus phaseoli* | WP_114495195.1 | WP_114495194.1 | WP_114494769.1 |
| Bacillota | *Fontibacillus solani* | WP_182535212.1 | WP_182535213.1 | WP_182535122.1 |
| Bacillota | *Fontibacillus sp. BL9* | WP_410768391.1 | WP_410768390.1 | WP_410768037.1 |

| | | | | |
|---|---|---|---|---|
| Bacillota | *Fredinandcohnia onubensis* | WP_099362744.1 | WP_099363396.1 | WP_099363096.1 |
| Bacillota | *Fredinandcohnia salidurans* | WP_388034606.1 | WP_304214230.1 | WP_388035390.1 |
| Bacillota | *Fredinandcohnia sp. QZ13* | WP_309536700.1 | WP_309534390.1 | WP_309534811.1 |
| Bacillota | *Fuchsiella alkaliacetigena* | WP_248662616.1 | WP_248662617.1 | WP_248662618.1 |
| Bacillota | *Geobacillus genomosp. 3* | WP_020960296.1 | WP_041267944.1 | WP_020959237.1 |
| Bacillota | *Geobacillus jurassicus* | WP_066229184.1 | WP_066229281.1 | WP_066233987.1 |
| Bacillota | *Geobacillus kaustophilus* | WP_020278320.1 | WP_044731029.1 | WP_044736192.1 |
| Bacillota | *Geobacillus sp. JS12* | WP_063192935.1 | WP_063192936.1 | WP_063192325.1 |
| Bacillota | *Geobacillus stearothermophilus* | WP_277391634.1 | WP_063210378.1 | WP_237420055.1 |
| Bacillota | *Geobacillus subterraneus* | WP_184318169.1 | WP_168369042.1 | WP_184317307.1 |
| Bacillota | *Geobacillus thermodenitrificans* | WP_327986912.1 | WP_328191681.1 | WP_100660330.1 |
| Bacillota | *Geomicrobium halophilum* | WP_184403564.1 | WP_184404824.1 | WP_184403282.1 |
| Bacillota | *Geomicrobium sediminis* | WP_204698194.1 | WP_204696653.1 | WP_204696779.1 |
| Bacillota | *Gordoniibacillus kamchatkensis* | WP_041046862.1 | WP_041046861.1 | WP_041046255.1 |
| Bacillota | *Gorillibacterium sp. CAU 1737* | WP_347768118.1 | WP_347768119.1 | WP_347772139.1 |
| Bacillota | *Gorillibacterium timonense* | WP_058303859.1 | WP_058303858.1 | WP_082651929.1 |
| Bacillota | *Gottfriedia acidiceleris* | WP_335486042.1 | WP_088010671.1 | WP_088013047.1 |
| Bacillota | *Gottfriedia endophytica* | WP_209402472.1 | WP_209402474.1 | WP_209403689.1 |
| Bacillota | *Gottfriedia luciferensis* | WP_088069167.1 | WP_088069169.1 | WP_088067724.1 |
| Bacillota | *Gottfriedia solisilvae* | WP_087999426.1 | WP_087999425.1 | WP_088000236.1 |
| Bacillota | *Gottfriedia sp. NPDC056225* | WP_172444130.1 | WP_172444023.1 | WP_375949226.1 |
| Bacillota | *Gracilibacillus alcaliphilus* | WP_204667049.1 | WP_204667842.1 | WP_204670532.1 |
| Bacillota | *Gracilibacillus boraciitolerans* | WP_035721112.1 | WP_035722829.1 | WP_084040548.1 |
| Bacillota | *Gracilibacillus caseinilyticus* | WP_244716889.1 | WP_244718528.1 | WP_244717793.1 |
| Bacillota | *Gracilibacillus halophilus* | WP_003466106.1 | WP_003462699.1 | WP_003463201.1 |
| Bacillota | *Gracilibacillus halotolerans* | WP_184244035.1 | WP_184249427.1 | WP_184243471.1 |
| Bacillota | *Gracilibacillus kekensis* | WP_073202086.1 | WP_073201074.1 | WP_073201781.1 |
| Bacillota | *Gracilibacillus lacisalsi* | WP_018933731.1 | WP_018930976.1 | WP_018932777.1 |

| | | | | |
|---|---|---|---|---|
| Bacillota | *Gracilibacillus marinus* | WP_390197772.1 | WP_390196412.1 | WP_390196950.1 |
| Bacillota | *Gracilibacillus massiliensis* | WP_058307601.1 | WP_058306746.1 | WP_058305611.1 |
| Bacillota | *Gracilibacillus orientalis* | WP_091483469.1 | WP_091484803.1 | WP_091483945.1 |
| Bacillota | *Gracilibacillus oryzae* | WP_153401852.1 | WP_153403976.1 | WP_153401574.1 |
| Bacillota | *Gracilibacillus salinarum* | WP_244746134.1 | WP_244743910.1 | WP_244744755.1 |
| Bacillota | *Gracilibacillus saliphilus* | WP_163581994.1 | WP_163579037.1 | WP_163579670.1 |
| Bacillota | *Gracilibacillus salitolerans* | WP_100360378.1 | WP_153791010.1 | WP_153791192.1 |
| Bacillota | *Gracilibacillus sp. S3-1-1* | WP_319957970.1 | WP_319956460.1 | WP_319956720.1 |
| Bacillota | *Gracilibacillus suaedae* | WP_208591048.1 | WP_208587972.1 | WP_208589337.1 |
| Bacillota | *Gracilibacillus thailandensis* | WP_153836484.1 | WP_163578820.1 | WP_153835392.1 |
| Bacillota | *Gracilibacillus ureilyticus* | WP_089738304.1 | WP_089741081.1 | WP_089741886.1 |
| Bacillota | *Gracilibacillus xinjiangensis* | WP_390252955.1 | WP_390254003.1 | WP_390253558.1 |
| Bacillota | *Halalkalibacillus halophilus* | WP_245570102.1 | WP_027965402.1 | WP_035512343.1 |
| Bacillota | *Halalkalibacillus sediminis* | WP_101330576.1 | WP_101331615.1 | WP_101330273.1 |
| Bacillota | *Halalkalibacter akibai* | WP_035663521.1 | WP_035663519.1 | WP_148296803.1 |
| Bacillota | *Halalkalibacter alkaliphilus* | WP_250094869.1 | WP_250094868.1 | WP_250095283.1 |
| Bacillota | *Halalkalibacter alkalisediminis* | WP_273840757.1 | WP_273840758.1 | WP_273839757.1 |
| Bacillota | *Halalkalibacter flavus* | WP_332630148.1 | WP_332630147.1 | WP_332632203.1 |
| Bacillota | *Halalkalibacter hemicellulosilyticus* | WP_035340208.1 | WP_369384506.1 | WP_035340416.1 |
| Bacillota | *Halalkalibacter kiskunsagensis* | WP_335958984.1 | WP_335958985.1 | WP_335960984.1 |
| Bacillota | *Halalkalibacter krulwichiae* | WP_066151466.1 | WP_066151468.1 | WP_066150491.1 |
| Bacillota | *Halalkalibacter lacteus* | WP_332693267.1 | WP_332693270.1 | WP_332690015.1 |
| Bacillota | *Halalkalibacter nanhaiisediminis* | WP_144448998.1 | WP_144448997.1 | WP_158640037.1 |
| Bacillota | *Halalkalibacter oceani* | WP_354571001.1 | WP_251193962.1 | WP_251192048.1 |
| Bacillota | *Halalkalibacter okhensis* | WP_034628459.1 | WP_052144674.1 | WP_034627251.1 |
| Bacillota | *Halalkalibacter urbisdiaboli* | WP_088102184.1 | WP_088102185.1 | WP_088104994.1 |
| Bacillota | *Halalkalibacter wakoensis* | WP_034742903.1 | WP_034742929.1 | WP_052001971.1 |
| Bacillota | *Halalkalibacterium halodurans* | WP_328033658.1 | WP_233444599.1 | WP_010898728.1 |

| | | | | |
|---|---|---|---|---|
| Bacillota | *Halalkalibacterium ligniniphilum* | WP_017728002.1 | WP_017728003.1 | WP_017727198.1 |
| Bacillota | *Halobacillus aidingensis* | WP_089651930.1 | WP_089651334.1 | WP_089652803.1 |
| Bacillota | *Halobacillus alkaliphilus* | WP_089750055.1 | WP_089751867.1 | WP_089751895.1 |
| Bacillota | *Halobacillus amylolyticus* | WP_245033281.1 | WP_245035014.1 | WP_245034112.1 |
| Bacillota | *Halobacillus andaensis* | WP_188375880.1 | WP_188377783.1 | WP_188376172.1 |
| Bacillota | *Halobacillus campisalis* | WP_289214293.1 | WP_289215238.1 | WP_289214028.1 |
| Bacillota | *Halobacillus dabanensis* | WP_075036701.1 | WP_075037406.1 | WP_075036544.1 |
| Bacillota | *Halobacillus faecis* | WP_371860329.1 | WP_146815254.1 | WP_146818693.1 |
| Bacillota | *Halobacillus halophilus* | WP_224892577.1 | WP_224893246.1 | WP_160840210.1 |
| Bacillota | *Halobacillus hunanensis* | WP_079530109.1 | WP_079526548.1 | WP_079529747.1 |
| Bacillota | *Halobacillus ihumii* | WP_163526213.1 | WP_163529223.1 | WP_163530396.1 |
| Bacillota | *Halobacillus karajensis* | WP_035504949.1 | WP_035508604.1 | WP_074733901.1 |
| Bacillota | *Halobacillus kuroshimensis* | WP_206932799.1 | WP_206934211.1 | WP_027954104.1 |
| Bacillota | *Halobacillus litoralis* | WP_226579282.1 | WP_225196267.1 | WP_128522867.1 |
| Bacillota | *Halobacillus locisalis* | WP_181471713.1 | WP_181473500.1 | WP_181471280.1 |
| Bacillota | *Halobacillus mangrovi* | WP_085029742.1 | WP_085030800.1 | WP_085030042.1 |
| Bacillota | *Halobacillus massiliensis* | WP_082234011.1 | WP_082235033.1 | WP_082234268.1 |
| Bacillota | *Halobacillus naozhouensis* | WP_283075109.1 | WP_283077805.1 | WP_283078586.1 |
| Bacillota | *Halobacillus rhizosphaerae* | WP_411787596.1 | WP_411790746.1 | WP_411790457.1 |
| Bacillota | *Halobacillus salinarum* | WP_244708082.1 | WP_244709258.1 | WP_244708500.1 |
| Bacillota | *Halobacillus salinus* | WP_079479978.1 | WP_079479320.1 | WP_135327270.1 |
| Bacillota | *Halobacillus seohaensis* | WP_204706383.1 | WP_204710739.1 | WP_204707910.1 |
| Bacillota | *Halobacillus shinanisalinarum* | WP_244753174.1 | WP_244752164.1 | WP_244752680.1 |
| Bacillota | *Halobacillus sp.* | WP_422165301.1 | WP_422167379.1 | WP_422168518.1 |
| Bacillota | *Halobacillus trueperi* | WP_394217372.1 | WP_394219676.1 | WP_115824858.1 |
| Bacillota | *Halobacillus yeomjeoni* | WP_197316167.1 | WP_197317823.1 | WP_197315676.1 |
| Bacillota | *Hazenella coriacea* | WP_131925631.1 | WP_131925633.1 | WP_131924031.1 |
| Bacillota | *Heliobacterium chlorum* | WP_188038396.1 | WP_188038395.1 | WP_188041012.1 |

| | | | | |
|---|---|---|---|---|
| Bacillota | *Heliobacterium mobile* | WP_170291528.1 | WP_155474654.1 | WP_155477797.1 |
| Bacillota | *Heliomicrobium modesticaldum* | WP_012283098.1 | WP_012283097.1 | WP_012283553.1 |
| Bacillota | *Heliomicrobium undosum* | WP_161255425.1 | WP_161255422.1 | WP_161254746.1 |
| Bacillota | *Heliorestis acidaminivorans* | WP_151618080.1 | WP_170270103.1 | WP_170270043.1 |
| Bacillota | *Heliorestis convoluta* | WP_153725135.1 | WP_153725134.1 | WP_153724431.1 |
| Bacillota | *Heyndrickxia acidicola* | WP_066262737.1 | WP_066262736.1 | WP_066264835.1 |
| Bacillota | *Heyndrickxia acidiproducens* | WP_018660367.1 | WP_018660369.1 | WP_018664087.1 |
| Bacillota | *Heyndrickxia camelliae* | WP_101353257.1 | WP_101353258.1 | WP_101352608.1 |
| Bacillota | *Heyndrickxia coagulans* | WP_308928048.1 | WP_328004612.1 | WP_046721295.1 |
| Bacillota | *Heyndrickxia faecalis* | WP_350346116.1 | WP_350346115.1 | WP_211062182.1 |
| Bacillota | *Heyndrickxia ginsengihumi* | WP_025729133.1 | WP_231473063.1 | WP_025730116.1 |
| Bacillota | *Heyndrickxia oleronia* | WP_280615767.1 | WP_251345137.1 | WP_212942048.1 |
| Bacillota | *Heyndrickxia sp. FSL K6-6286* | WP_342541230.1 | WP_342541229.1 | WP_342539788.1 |
| Bacillota | *Heyndrickxia sporothermodurans* | WP_066228585.1 | WP_066228588.1 | WP_066230041.1 |
| Bacillota | *Heyndrickxia vini* | WP_202780324.1 | WP_202780323.1 | WP_202776866.1 |
| Bacillota | *Hydrogenibacillus schlegelii* | WP_066203583.1 | WP_169816914.1 | WP_082718315.1 |
| Bacillota | *Indiicoccus explosivorum* | WP_088007583.1 | WP_088008730.1 | WP_241535823.1 |
| Bacillota | *Insulibacter thermoxylanivorax* | WP_200965202.1 | WP_200965920.1 | WP_200965739.1 |
| Bacillota | *Jeotgalibacillus alimentarius* | WP_200889064.1 | WP_041122687.1 | WP_041122314.1 |
| Bacillota | *Jeotgalibacillus aurantiacus* | WP_227395293.1 | WP_227395292.1 | WP_227395612.1 |
| Bacillota | *Jeotgalibacillus campisalis* | WP_041057981.1 | WP_041057983.1 | WP_041057289.1 |
| Bacillota | *Jeotgalibacillus haloalkalitolerans* | WP_322420988.1 | WP_322420989.1 | WP_322420580.1 |
| Bacillota | *Jeotgalibacillus malaysiensis* | WP_404403476.1 | WP_404403478.1 | WP_404409007.1 |
| Bacillota | *Jeotgalibacillus marinus* | WP_367777475.1 | WP_367777476.1 | WP_367778874.1 |
| Bacillota | *Jeotgalibacillus proteolyticus* | WP_104056735.1 | WP_104056736.1 | WP_104056138.1 |
| Bacillota | *Jeotgalibacillus salarius* | WP_134380087.1 | WP_134380089.1 | WP_134382059.1 |
| Bacillota | *Jeotgalibacillus soli* | WP_041090089.1 | WP_041090090.1 | WP_041088882.1 |
| Bacillota | *Jeotgalibacillus sp. S-D1* | WP_133375592.1 | WP_133376314.1 | WP_133375963.1 |

| | | | | |
|---|---|---|---|---|
| Bacillota | *Jeotgalibacillus terrae* | WP_204727679.1 | WP_204727680.1 | WP_204728610.1 |
| Bacillota | *Jeotgalicoccus aerolatus* | WP_186090071.1 | WP_092597945.1 | WP_186090400.1 |
| Bacillota | *Jeotgalicoccus coquinae* | WP_184282109.1 | WP_184283941.1 | WP_184281317.1 |
| Bacillota | *Jeotgalicoccus halotolerans* | WP_369890706.1 | WP_115885244.1 | WP_115885716.1 |
| Bacillota | *Jeotgalicoccus meleagridis* | WP_185125737.1 | WP_185125207.1 | WP_185126131.1 |
| Bacillota | *Jeotgalicoccus psychrophilus* | WP_026858060.1 | WP_026858943.1 | WP_026858474.1 |
| Bacillota | *Jeotgalicoccus saudimassiliensis* | WP_035808832.1 | WP_035810548.1 | WP_035807980.1 |
| Bacillota | *Jeotgalicoccus sp. ATCC 8456* | WP_198687648.1 | WP_198687319.1 | WP_198687862.1 |
| Bacillota | *Kroppenstedtia eburnea* | WP_076522904.1 | WP_040387154.1 | WP_076523359.1 |
| Bacillota | *Kroppenstedtia guangzhouensis* | WP_188430415.1 | WP_188430417.1 | WP_188428771.1 |
| Bacillota | *Kroppenstedtia pulmonis* | WP_173221993.1 | WP_173221991.1 | WP_173222804.1 |
| Bacillota | *Kroppenstedtia sanguinis* | WP_380163115.1 | WP_380163114.1 | WP_380164071.1 |
| Bacillota | *Kurthia gibsonii* | WP_325942863.1 | WP_368634274.1 | WP_325921672.1 |
| Bacillota | *Kurthia huakuii* | WP_029498452.1 | WP_029497763.1 | WP_029498146.1 |
| Bacillota | *Kurthia massiliensis* | WP_026021961.1 | WP_010287167.1 | WP_010286585.1 |
| Bacillota | *Kurthia senegalensis* | WP_026022481.1 | WP_010302634.1 | WP_010301772.1 |
| Bacillota | *Kurthia sibirica* | WP_109307152.1 | WP_109307299.1 | WP_109304434.1 |
| Bacillota | *Kurthia sp. FSL E2-0154* | WP_340995847.1 | WP_340998718.1 | WP_340995481.1 |
| Bacillota | *Kurthia zopfii* | WP_109350159.1 | WP_109350915.1 | WP_109349916.1 |
| Bacillota | *Laceyella putida* | WP_379865747.1 | WP_379865748.1 | WP_379866126.1 |
| Bacillota | *Laceyella sacchari* | WP_054096534.1 | WP_259436617.1 | WP_259435664.1 |
| Bacillota | *Laceyella sediminis* | WP_022737412.1 | WP_106342438.1 | WP_106341380.1 |
| Bacillota | *Laceyella tengchongensis* | WP_102992298.1 | WP_333639873.1 | WP_102992870.1 |
| Bacillota | *Lacicoccus alkaliphilus* | WP_072707734.1 | WP_072710414.1 | WP_072708462.1 |
| Bacillota | *Lacicoccus qingdaonensis* | WP_092984866.1 | WP_092985141.1 | WP_092983623.1 |
| Bacillota | *Lederbergia citrea* | WP_213097146.1 | WP_213106924.1 | WP_213107223.1 |
| Bacillota | *Lederbergia citri* | WP_213123659.1 | WP_213123660.1 | WP_213123119.1 |
| Bacillota | *Lederbergia citrisecunda* | WP_213110163.1 | WP_213110164.1 | WP_213109577.1 |

| | | | | |
|---|---|---|---|---|
| Bacillota | *Lederbergia galactosidilytica* | WP_057988245.1 | WP_057988247.1 | WP_057985812.1 |
| Bacillota | *Lederbergia graminis* | WP_382346605.1 | WP_144919802.1 | WP_382347844.1 |
| Bacillota | *Lederbergia lenta* | WP_066137836.1 | WP_251519880.1 | WP_066139938.1 |
| Bacillota | *Lederbergia panacisoli* | WP_257579653.1 | WP_257579654.1 | WP_257580260.1 |
| Bacillota | *Lederbergia ruris* | WP_158322329.1 | WP_212965587.1 | WP_212966198.1 |
| Bacillota | *Lederbergia sp. NSJ-179* | WP_244699947.1 | WP_244699945.1 | WP_244700086.1 |
| Bacillota | *Lederbergia wuyishanensis* | WP_244680233.1 | WP_244680234.1 | WP_244679783.1 |
| Bacillota | *Lentibacillus amyloliquefaciens* | WP_068442481.1 | WP_068447796.1 | WP_068440804.1 |
| Bacillota | *Lentibacillus cibarius* | WP_142791518.1 | WP_142792004.1 | WP_142791314.1 |
| Bacillota | *Lentibacillus halodurans* | WP_090232615.1 | WP_090234410.1 | WP_090238274.1 |
| Bacillota | *Lentibacillus halophilus* | WP_343753122.1 | WP_343754212.1 | WP_343753559.1 |
| Bacillota | *Lentibacillus jeotgali* | WP_010530800.1 | WP_029329174.1 | WP_010530442.1 |
| Bacillota | *Lentibacillus juripiscarius* | WP_382392739.1 | WP_382393513.1 | WP_382392120.1 |
| Bacillota | *Lentibacillus kapialis* | WP_188631484.1 | WP_188634106.1 | WP_188631645.1 |
| Bacillota | *Lentibacillus kimchii* | WP_382361417.1 | WP_382360135.1 | WP_382357189.1 |
| Bacillota | *Lentibacillus persicus* | WP_090087577.1 | WP_090084942.1 | WP_090083743.1 |
| Bacillota | *Lentibacillus salicampi* | WP_135109086.1 | WP_135109587.1 | WP_135108198.1 |
| Bacillota | *Lentibacillus salinarum* | WP_382397247.1 | WP_382399436.1 | WP_382396958.1 |
| Bacillota | *Lentibacillus saliphilus* | WP_217585454.1 | WP_217588552.1 | WP_217585692.1 |
| Bacillota | *Lentibacillus sediminis* | WP_100010811.1 | WP_100010446.1 | WP_100011055.1 |
| Bacillota | *Lentibacillus sp.* | WP_324223546.1 | WP_324224588.1 | WP_324224008.1 |
| Bacillota | *Lihuaxuella thermophila* | WP_244527535.1 | WP_089969006.1 | WP_089966921.1 |
| Bacillota | *Litchfieldia alkalitelluris* | WP_078546123.1 | WP_078546122.1 | WP_078547319.1 |
| Bacillota | *Litchfieldia salsa* | WP_090852835.1 | WP_090852838.1 | WP_090850473.1 |
| Bacillota | *Litoribacterium kuwaitense* | WP_165199021.1 | WP_165199020.1 | WP_165199860.1 |
| Bacillota | *Longirhabdus pacifica* | WP_128894412.1 | WP_128894413.1 | WP_128896012.1 |
| Bacillota | *Lottiidibacillus patelloidae* | WP_094923848.1 | WP_094923846.1 | WP_094921476.1 |
| Bacillota | *Lysinibacillus alkalisoli* | WP_188613934.1 | WP_188613427.1 | WP_188614344.1 |

| | | | | |
|---|---|---|---|---|
| Bacillota | *Lysinibacillus antri* | WP_126657732.1 | WP_126658849.1 | WP_126657660.1 |
| Bacillota | *Lysinibacillus capsici* | WP_320940576.1 | WP_333880215.1 | WP_333880887.1 |
| Bacillota | *Lysinibacillus cavernae* | WP_155590643.1 | WP_155592437.1 | WP_155590130.1 |
| Bacillota | *Lysinibacillus composti* | WP_124764968.1 | WP_124763490.1 | WP_124764103.1 |
| Bacillota | *Lysinibacillus contaminans* | WP_053584004.1 | WP_053585202.1 | WP_053584453.1 |
| Bacillota | *Lysinibacillus endophyticus* | WP_341966957.1 | WP_253423194.1 | WP_253423882.1 |
| Bacillota | *Lysinibacillus fusiformis* | WP_025114698.1 | WP_330493602.1 | WP_143997544.1 |
| Bacillota | *Lysinibacillus louembei* | WP_319836654.1 | WP_319837291.1 | WP_319836882.1 |
| Bacillota | *Lysinibacillus macroides* | WP_053994351.1 | WP_053993183.1 | WP_053993826.1 |
| Bacillota | *Lysinibacillus odysseyi* | WP_036154513.1 | WP_036153162.1 | WP_036156049.1 |
| Bacillota | *Lysinibacillus pakistanensis* | WP_054770305.1 | WP_283870672.1 | WP_283871065.1 |
| Bacillota | *Lysinibacillus parviboronicapiens* | WP_354471761.1 | WP_107947582.1 | WP_107947661.1 |
| Bacillota | *Lysinibacillus piscis* | WP_264986785.1 | WP_264987881.1 | WP_264987283.1 |
| Bacillota | *Lysinibacillus sp. NPDC093210* | WP_401008257.1 | WP_401010871.1 | WP_401009567.1 |
| Bacillota | *Lysinibacillus sphaericus* | WP_197141462.1 | WP_142509402.1 | WP_024362149.1 |
| Bacillota | *Lysinibacillus telephonicus* | WP_126294168.1 | WP_126293219.1 | WP_346235537.1 |
| Bacillota | *Lysinibacillus timonensis* | WP_106784717.1 | WP_106783408.1 | WP_106784036.1 |
| Bacillota | *Lysinibacillus xylanilyticus* | WP_049666747.1 | WP_402983101.1 | WP_400959240.1 |
| Bacillota | *Lysinibacillus yapensis* | WP_118874311.1 | WP_118875069.1 | WP_118874754.1 |
| Bacillota | *Macrococcoides bohemicum* | WP_419743646.1 | WP_199798155.1 | WP_312039520.1 |
| Bacillota | *Macrococcoides canis* | WP_254253336.1 | WP_138072676.1 | WP_086042282.1 |
| Bacillota | *Macrococcoides caseolyticum* | WP_233681270.1 | WP_233681943.1 | WP_157820940.1 |
| Bacillota | *Macrococcoides goetzii* | WP_099578185.1 | WP_205757173.1 | WP_099580771.1 |
| Bacillota | *Macrococcus armenti* | WP_224186387.1 | WP_224187339.1 | WP_224184470.1 |
| Bacillota | *Macrococcus bovicus* | WP_289649503.1 | WP_289650147.1 | WP_289650867.1 |
| Bacillota | *Macrococcus brunensis* | WP_239046908.1 | WP_133431868.1 | WP_239035515.1 |
| Bacillota | *Macrococcus carouselicus* | WP_133417120.1 | WP_133418089.1 | WP_133416794.1 |
| Bacillota | *Macrococcus epidermidis* | WP_254256476.1 | WP_254256905.1 | WP_254257470.1 |

| | | | | |
|---|---|---|---|---|
| Bacillota | *Macrococcus equipercicus* | WP_149458850.1 | WP_149459263.1 | WP_254250417.1 |
| Bacillota | *Macrococcus hajekii* | WP_133429085.1 | WP_133430543.1 | WP_133429380.1 |
| Bacillota | *Macrococcus lamae* | WP_133442833.1 | WP_133443643.1 | WP_133444362.1 |
| Bacillota | *Macrococcus sp. DPC7161* | WP_129064231.1 | WP_129065470.1 | WP_164993313.1 |
| Bacillota | *Mammaliicoccus fleurettii* | WP_115337434.1 | WP_208144256.1 | WP_218702747.1 |
| Bacillota | *Mammaliicoccus lentus* | WP_210618937.1 | WP_210617024.1 | WP_416349754.1 |
| Bacillota | *Mammaliicoccus sciuri* | WP_204187928.1 | WP_206165146.1 | WP_107617817.1 |
| Bacillota | *Mammaliicoccus sp. Dog046* | WP_323702337.1 | WP_323703356.1 | WP_323702670.1 |
| Bacillota | *Mammaliicoccus stepanovicii* | WP_095087978.1 | WP_095089715.1 | WP_095088520.1 |
| Bacillota | *Mammaliicoccus vitulinus* | WP_421040920.1 | WP_103322196.1 | WP_107536473.1 |
| Bacillota | *Mangrovibacillus sp. Mu-81* | WP_334218689.1 | WP_334218688.1 | WP_334217387.1 |
| Bacillota | *Marinicrinis lubricantis* | WP_379893406.1 | WP_379893408.1 | WP_379895403.1 |
| Bacillota | *Marinicrinis sediminis* | WP_379930489.1 | WP_379930488.1 | WP_379929885.1 |
| Bacillota | *Marininema halotolerans* | WP_245838860.1 | WP_091838700.1 | WP_176391884.1 |
| Bacillota | *Marininema mesophilum* | WP_091737720.1 | WP_091737717.1 | WP_091734794.1 |
| Bacillota | *Marinithermofilum abyssi* | WP_188646919.1 | WP_188646918.1 | WP_188648308.1 |
| Bacillota | *Marinococcus halophilus* | WP_079476570.1 | WP_079476571.1 | WP_079476090.1 |
| Bacillota | *Marinococcus halotolerans* | WP_022792248.1 | WP_022792249.1 | WP_022792687.1 |
| Bacillota | *Marinococcus luteus* | WP_091611148.1 | WP_091611152.1 | WP_091610196.1 |
| Bacillota | *Massilibacterium senegalense* | WP_062198225.1 | WP_062198226.1 | WP_062197761.1 |
| Bacillota | *Mechercharimyces sp. CAU 1602* | WP_258839241.1 | WP_258839240.1 | WP_258838868.1 |
| Bacillota | *Melghiribacillus thermohalophilus* | WP_132370988.1 | WP_132372634.1 | WP_132370283.1 |
| Bacillota | *Melghirimyces algeriensis* | WP_142504252.1 | WP_246064816.1 | WP_142504508.1 |
| Bacillota | *Melghirimyces profundicolus* | WP_108023223.1 | WP_108023221.1 | WP_108026146.1 |
| Bacillota | *Melghirimyces thermohalophilus* | WP_091569222.1 | WP_091569219.1 | WP_091565521.1 |
| Bacillota | *Mesobacillus boroniphilus* | WP_213368229.1 | WP_425374501.1 | WP_213366746.1 |
| Bacillota | *Mesobacillus campisalis* | WP_173427554.1 | WP_046524176.1 | WP_046522237.1 |
| Bacillota | *Mesobacillus foraminis* | WP_412919659.1 | WP_412919660.1 | WP_132005738.1 |

| | | | | |
|---|---|---|---|---|
| Bacillota | *Mesobacillus harenae* | WP_174733990.1 | WP_174733991.1 | WP_174733830.1 |
| Bacillota | *Mesobacillus jeotgali* | WP_311071330.1 | WP_079509534.1 | WP_192470257.1 |
| Bacillota | *Mesobacillus maritimus* | WP_221872545.1 | WP_404329097.1 | WP_251425668.1 |
| Bacillota | *Mesobacillus persicus* | WP_090744525.1 | WP_090744528.1 | WP_090747490.1 |
| Bacillota | *Mesobacillus selenatarsenatis* | WP_167831352.1 | WP_084135369.1 | WP_041965511.1 |
| Bacillota | *Mesobacillus sp.* | WP_423991807.1 | WP_423991813.1 | WP_423990406.1 |
| Bacillota | *Mesobacillus subterraneus* | WP_226642288.1 | WP_246017800.1 | WP_125479208.1 |
| Bacillota | *Mesobacillus thioparans* | WP_415817540.1 | WP_415817665.1 | WP_415812484.1 |
| Bacillota | *Mesobacillus zeae* | WP_181213417.1 | WP_119114370.1 | WP_119111472.1 |
| Bacillota | *Metabacillus arenae* | WP_191159270.1 | WP_191159271.1 | WP_191157009.1 |
| Bacillota | *Metabacillus bambusae* | WP_207980778.1 | WP_207980779.1 | WP_207975146.1 |
| Bacillota | *Metabacillus crassostreae* | WP_204954254.1 | WP_204954255.1 | WP_204953265.1 |
| Bacillota | *Metabacillus dongyingensis* | WP_421175340.1 | WP_421175339.1 | WP_421176627.1 |
| Bacillota | *Metabacillus endolithicus* | WP_247344015.1 | WP_247344017.1 | WP_247341532.1 |
| Bacillota | *Metabacillus fastidiosus* | WP_328027515.1 | WP_412758905.1 | WP_326173759.1 |
| Bacillota | *Metabacillus flavus* | WP_211558719.1 | WP_211558721.1 | WP_211557961.1 |
| Bacillota | *Metabacillus halosaccharovorans* | WP_264144479.1 | WP_216774187.1 | WP_216773076.1 |
| Bacillota | *Metabacillus herbersteinensis* | WP_378932858.1 | WP_378933101.1 | WP_378931977.1 |
| Bacillota | *Metabacillus idriensis* | WP_154319159.1 | WP_224773259.1 | WP_309191454.1 |
| Bacillota | *Metabacillus indicus* | WP_411333257.1 | WP_342822342.1 | WP_282033509.1 |
| Bacillota | *Metabacillus iocasae* | WP_205186994.1 | WP_205187002.1 | WP_205183511.1 |
| Bacillota | *Metabacillus kandeliae* | WP_231787999.1 | WP_231788118.1 | WP_231788484.1 |
| Bacillota | *Metabacillus lacus* | WP_154306098.1 | WP_154308074.1 | WP_154305956.1 |
| Bacillota | *Metabacillus litoralis* | WP_226666844.1 | WP_226666846.1 | WP_162987286.1 |
| Bacillota | *Metabacillus malikii* | WP_307335414.1 | WP_307335411.1 | WP_307341933.1 |
| Bacillota | *Metabacillus mangrovi* | WP_155110674.1 | WP_155110675.1 | WP_155111176.1 |
| Bacillota | *Metabacillus niabensis* | WP_226526663.1 | WP_095302204.1 | WP_174879360.1 |
| Bacillota | *Metabacillus rhizosphaerae* | WP_338789327.1 | WP_338789326.1 | WP_338786668.1 |

| | | | | |
|---|---|---|---|---|
| Bacillota | *Metabacillus schmidteae* | WP_102229189.1 | WP_175639695.1 | WP_102229602.1 |
| Bacillota | *Metabacillus sediminilitoris* | WP_136351060.1 | WP_136351061.1 | WP_168733743.1 |
| Bacillota | *Metabacillus sediminis* | WP_338776765.1 | WP_338776767.1 | WP_338781799.1 |
| Bacillota | *Metabacillus sp. cB07* | WP_203287618.1 | WP_242524605.1 | WP_203290116.1 |
| Bacillota | *Metalysinibacillus jejuensis* | WP_108306670.1 | WP_108306092.1 | WP_108305698.1 |
| Bacillota | *Metaplanococcus flavidus* | WP_144836964.1 | WP_144841296.1 | WP_144837543.1 |
| Bacillota | *Metasolibacillus fluoroglycofenilyticus* | WP_107942412.1 | WP_066168100.1 | WP_107942236.1 |
| Bacillota | *Metasolibacillus meyeri* | WP_326123580.1 | WP_326121111.1 | WP_326123935.1 |
| Bacillota | *Metasolibacillus sp.* | WP_293921022.1 | WP_293921394.1 | WP_293922100.1 |
| Bacillota | *Microaerobacter geothermalis* | WP_236405557.1 | WP_407942200.1 | WP_236405793.1 |
| Bacillota | *Natranaerobius thermophilus* | WP_012447734.1 | WP_012447733.1 | WP_012447732.1 |
| Bacillota | *Natranaerobius trueperi* | WP_276207008.1 | WP_089023721.1 | WP_089023720.1 |
| Bacillota | *Natranaerofaba carboxydovora* | WP_241080178.1 | WP_241080177.1 | WP_241080176.1 |
| Bacillota | *Natribacillus halophilus* | WP_090395622.1 | WP_425433695.1 | WP_090397283.1 |
| Bacillota | *Natronobacillus azotifigens* | WP_268778642.1 | WP_268780107.1 | WP_268781301.1 |
| Bacillota | *Neobacillus bataviensis* | WP_223593452.1 | WP_223593454.1 | WP_223595997.1 |
| Bacillota | *Neobacillus citreus* | WP_213146114.1 | WP_213145044.1 | WP_213140845.1 |
| Bacillota | *Neobacillus cucumis* | WP_328052667.1 | WP_309527024.1 | WP_101647346.1 |
| Bacillota | *Neobacillus dielmonensis* | WP_042457792.1 | WP_042457922.1 | WP_042456158.1 |
| Bacillota | *Neobacillus drentensis* | WP_335441998.1 | WP_335441999.1 | WP_335465705.1 |
| Bacillota | *Neobacillus driksii* | WP_374931224.1 | WP_307310470.1 | WP_179597349.1 |
| Bacillota | *Neobacillus endophyticus* | WP_173070301.1 | WP_173070303.1 | WP_173065939.1 |
| Bacillota | *Neobacillus fumarioli* | WP_066369844.1 | WP_066369846.1 | WP_066368341.1 |
| Bacillota | *Neobacillus ginsengisoli* | WP_307404320.1 | WP_307404318.1 | WP_307406315.1 |
| Bacillota | *Neobacillus jeddahensis* | WP_040208386.1 | WP_040208383.1 | WP_040204841.1 |
| Bacillota | *Neobacillus kokaensis* | WP_191271830.1 | WP_191271828.1 | WP_191276915.1 |
| Bacillota | *Neobacillus massiliamazoniensis* | WP_090637063.1 | WP_090637065.1 | WP_090630295.1 |
| Bacillota | *Neobacillus mesonae* | WP_251465883.1 | WP_286232340.1 | WP_066395689.1 |

| | | | | |
|---|---|---|---|---|
| Bacillota | *Neobacillus muris* | WP_251554148.1 | WP_251554150.1 | WP_251549056.1 |
| Bacillota | *Neobacillus niacini* | WP_063251212.1 | WP_335528719.1 | WP_262307682.1 |
| Bacillota | *Neobacillus notoginsengisoli* | WP_118920643.1 | WP_118920642.1 | WP_118922791.1 |
| Bacillota | *Neobacillus novalis* | WP_066089240.1 | WP_066089243.1 | WP_066085070.1 |
| Bacillota | *Neobacillus piezotolerans* | WP_115452360.1 | WP_115452359.1 | WP_115450397.1 |
| Bacillota | *Neobacillus rhizophilus* | WP_213120381.1 | WP_213120380.1 | WP_213116428.1 |
| Bacillota | *Neobacillus rhizosphaerae* | WP_248736036.1 | WP_248736035.1 | WP_248733686.1 |
| Bacillota | *Neobacillus sedimentimangrovi* | WP_163183094.1 | WP_038535832.1 | WP_231314560.1 |
| Bacillota | *Neobacillus soli* | WP_066061022.1 | WP_066061019.1 | WP_066066500.1 |
| Bacillota | *Neobacillus sp.* | WP_312475799.1 | WP_374718208.1 | WP_312469188.1 |
| Bacillota | *Neobacillus terrae* | WP_166022683.1 | WP_166022681.1 | WP_166023729.1 |
| Bacillota | *Neobacillus thermocopriae* | WP_163252363.1 | WP_163252362.1 | WP_163250174.1 |
| Bacillota | *Neobacillus vireti* | WP_024030504.1 | WP_335567265.1 | WP_335497781.1 |
| Bacillota | *Niallia circulans* | WP_047940384.1 | WP_095322317.1 | WP_340461021.1 |
| Bacillota | *Niallia endozanthoxylica* | WP_150439756.1 | WP_150442538.1 | WP_150439624.1 |
| Bacillota | *Niallia nealsonii* | WP_101176769.1 | WP_101176770.1 | WP_101178689.1 |
| Bacillota | *Niallia oryzisoli* | WP_338451815.1 | WP_338453405.1 | WP_394232620.1 |
| Bacillota | *Niallia sp.* | WP_312096806.1 | WP_312096808.1 | WP_312095343.1 |
| Bacillota | *Niallia taxi* | WP_127737632.1 | WP_182104199.1 | WP_182102957.1 |
| Bacillota | *Nosocomiicoccus ampullae* | WP_219522448.1 | WP_068129475.1 | WP_183674126.1 |
| Bacillota | *Nosocomiicoccus massiliensis* | WP_102167308.1 | WP_040928630.1 | WP_052327350.1 |
| Bacillota | *Nosocomiicoccus sp. HMSC09A07* | WP_070456533.1 | WP_070457890.1 | WP_070457575.1 |
| Bacillota | *Novibacillus thermophilus* | WP_077720594.1 | WP_077718877.1 | WP_077719370.1 |
| Bacillota | *Numidum massiliense* | WP_054949644.1 | WP_054949643.1 | WP_054949462.1 |
| Bacillota | *Oceanobacillus alkalisoli* | WP_235760767.1 | WP_235761286.1 | WP_238471747.1 |
| Bacillota | *Oceanobacillus arenosus* | WP_115772703.1 | WP_115774338.1 | WP_115773638.1 |
| Bacillota | *Oceanobacillus bengalensis* | WP_121133105.1 | WP_121128214.1 | WP_121130240.1 |
| Bacillota | *Oceanobacillus chungangensis* | WP_115747896.1 | WP_115751250.1 | WP_115749999.1 |

| | | | | |
|---|---|---|---|---|
| Bacillota | *Oceanobacillus damuensis* | WP_067725496.1 | WP_067727112.1 | WP_067728293.1 |
| Bacillota | *Oceanobacillus halophilus* | WP_121202721.1 | WP_121203191.1 | WP_121202470.1 |
| Bacillota | *Oceanobacillus halotolerans* | WP_163969839.1 | WP_163970678.1 | WP_163970754.1 |
| Bacillota | *Oceanobacillus iheyensis* | WP_011066165.1 | WP_011066878.1 | WP_011065861.1 |
| Bacillota | *Oceanobacillus indicireducens* | WP_188856749.1 | WP_188859558.1 | WP_188855640.1 |
| Bacillota | *Oceanobacillus jeddahense* | WP_040981361.1 | WP_052400892.1 | WP_256706976.1 |
| Bacillota | *Oceanobacillus jordanicus* | WP_238018995.1 | WP_106897347.1 | WP_238020858.1 |
| Bacillota | *Oceanobacillus kapialis* | WP_379564138.1 | WP_404454280.1 | WP_379560979.1 |
| Bacillota | *Oceanobacillus kimchii* | WP_260048388.1 | WP_272031485.1 | WP_317957949.1 |
| Bacillota | *Oceanobacillus limi* | WP_090866871.1 | WP_090868480.1 | WP_090869609.1 |
| Bacillota | *Oceanobacillus locisalsi* | WP_379590821.1 | WP_379593821.1 | WP_379593309.1 |
| Bacillota | *Oceanobacillus longus* | WP_379495450.1 | WP_379496333.1 | WP_379495717.1 |
| Bacillota | *Oceanobacillus luteolus* | WP_251510379.1 | WP_251511152.1 | WP_379596990.1 |
| Bacillota | *Oceanobacillus manasiensis* | WP_042223609.1 | WP_042222495.1 | WP_042223865.1 |
| Bacillota | *Oceanobacillus massiliensis* | WP_010650627.1 | WP_010647567.1 | WP_337019585.1 |
| Bacillota | *Oceanobacillus neutriphilus* | WP_188734211.1 | WP_188732546.1 | WP_188737378.1 |
| Bacillota | *Oceanobacillus oncorhynchi* | WP_343768320.1 | WP_193064296.1 | WP_343769320.1 |
| Bacillota | *Oceanobacillus picturae* | WP_119574346.1 | WP_058950952.1 | WP_379550491.1 |
| Bacillota | *Oceanobacillus piezotolerans* | WP_121521206.1 | WP_121521890.1 | WP_246001176.1 |
| Bacillota | *Oceanobacillus polygoni* | WP_149474075.1 | WP_149473332.1 | WP_149476200.1 |
| Bacillota | *Oceanobacillus profundus* | WP_379601181.1 | WP_251659207.1 | WP_118888966.1 |
| Bacillota | *Oceanobacillus rekensis* | WP_087971512.1 | WP_087972585.1 | WP_087973019.1 |
| Bacillota | *Oceanobacillus salinisoli* | WP_156289216.1 | WP_156288339.1 | WP_156290435.1 |
| Bacillota | *Oceanobacillus saliphilus* | WP_249871848.1 | WP_249871298.1 | WP_249869876.1 |
| Bacillota | *Oceanobacillus senegalensis* | WP_085994422.1 | WP_085991367.1 | WP_085992481.1 |
| Bacillota | *Oceanobacillus sojae* | WP_077603012.1 | WP_077601946.1 | WP_379545238.1 |
| Bacillota | *Oceanobacillus sp. FSL K6-2867* | WP_339227992.1 | WP_339227347.1 | WP_339227598.1 |
| Bacillota | *Oceanobacillus timonensis* | WP_080874393.1 | WP_080873302.1 | WP_080874844.1 |

| | | | | |
|---|---|---|---|---|
| Bacillota | *Oceanobacillus zhaokaii* | WP_114916620.1 | WP_114917281.1 | WP_114916282.1 |
| Bacillota | *Oikeobacillus pervagus* | WP_307255661.1 | WP_307255787.1 | WP_307257227.1 |
| Bacillota | *Ornithinibacillus bavariensis* | WP_212921903.1 | WP_212919871.1 | WP_212919548.1 |
| Bacillota | *Ornithinibacillus californiensis* | WP_047985292.1 | WP_047984527.1 | WP_047984563.1 |
| Bacillota | *Ornithinibacillus caprae* | WP_155666829.1 | WP_155669075.1 | WP_155669501.1 |
| Bacillota | *Ornithinibacillus contaminans* | WP_047980060.1 | WP_047980813.1 | WP_047981359.1 |
| Bacillota | *Ornithinibacillus gellani* | WP_123316431.1 | WP_123316031.1 | WP_123315413.1 |
| Bacillota | *Ornithinibacillus halophilus* | WP_072887307.1 | WP_072892005.1 | WP_072886773.1 |
| Bacillota | *Ornithinibacillus halotolerans* | WP_188382795.1 | WP_188383526.1 | WP_188385364.1 |
| Bacillota | *Ornithinibacillus hominis* | WP_186868106.1 | WP_186870584.1 | WP_186868582.1 |
| Bacillota | *Ornithinibacillus massiliensis* | WP_211741036.1 | WP_211742459.1 | WP_211741530.1 |
| Bacillota | *Ornithinibacillus salinisoli* | WP_377557228.1 | WP_377557623.1 | WP_377555581.1 |
| Bacillota | *Ornithinibacillus scapharcae* | WP_033445626.1 | WP_010098482.1 | WP_010096796.1 |
| Bacillota | *Ornithinibacillus sp. 179-J 7C1 HS* | WP_412975632.1 | WP_412977190.1 | WP_412975348.1 |
| Bacillota | *Ornithinibacillus xuwenensis* | WP_345823651.1 | WP_345826023.1 | WP_345823993.1 |
| Bacillota | *Paenactinomyces guangxiensis* | WP_181751724.1 | WP_309506121.1 | WP_181751500.1 |
| Bacillota | *Paenalkalicoccus suaedae* | WP_176009154.1 | WP_176009153.1 | WP_176009567.1 |
| Bacillota | *Paenibacillus abyssi* | WP_188528309.1 | WP_188528307.1 | WP_188531779.1 |
| Bacillota | *Paenibacillus aceris* | WP_167053849.1 | WP_167053847.1 | WP_167063709.1 |
| Bacillota | *Paenibacillus aceti* | WP_120462256.1 | WP_120462257.1 | WP_120462650.1 |
| Bacillota | *Paenibacillus aestuarii* | WP_270881318.1 | WP_270881317.1 | WP_270885492.1 |
| Bacillota | *Paenibacillus agaridevorans* | WP_108991958.1 | WP_214631096.1 | WP_214629140.1 |
| Bacillota | *Paenibacillus agilis* | WP_144988852.1 | WP_144988849.1 | WP_144991373.1 |
| Bacillota | *Paenibacillus agri* | WP_175373667.1 | WP_175373669.1 | WP_175372874.1 |
| Bacillota | *Paenibacillus agricola* | WP_166145632.1 | WP_166145635.1 | WP_311764869.1 |
| Bacillota | *Paenibacillus alba* | WP_173215900.1 | WP_173215902.1 | WP_173216716.1 |
| Bacillota | *Paenibacillus albicereus* | WP_206110059.1 | WP_168907475.1 | WP_168908071.1 |
| Bacillota | *Paenibacillus albidus* | WP_215176829.1 | WP_215176831.1 | WP_189025022.1 |

| | | | | |
|---|---|---|---|---|
| Bacillota | *Paenibacillus albiflavus* | WP_132416234.1 | WP_132416236.1 | WP_132419223.1 |
| Bacillota | *Paenibacillus albus* | WP_126016142.1 | WP_126016140.1 | WP_126015742.1 |
| Bacillota | *Paenibacillus alginolyticus* | WP_029193046.1 | WP_268615699.1 | WP_029196527.1 |
| Bacillota | *Paenibacillus algorifonticola* | WP_046232076.1 | WP_046232075.1 | WP_046230403.1 |
| Bacillota | *Paenibacillus alkaliterrae* | WP_235235122.1 | WP_235235115.1 | WP_235238666.1 |
| Bacillota | *Paenibacillus alkalitolerans* | WP_199614925.1 | WP_199614927.1 | WP_199618942.1 |
| Bacillota | *Paenibacillus allorhizoplanae* | WP_236284611.1 | WP_236285355.1 | WP_236287531.1 |
| Bacillota | *Paenibacillus allorhizosphaerae* | WP_218100601.1 | WP_218100602.1 | WP_218097063.1 |
| Bacillota | *Paenibacillus alvei* | WP_197266707.1 | WP_420029255.1 | WP_138184889.1 |
| Bacillota | *Paenibacillus amylolyticus* | WP_123066039.1 | WP_347386738.1 | WP_123062536.1 |
| Bacillota | *Paenibacillus andongensis* | WP_261306455.1 | WP_261301090.1 | WP_261305761.1 |
| Bacillota | *Paenibacillus anseongense* | WP_326060907.1 | WP_326062979.1 | WP_326060556.1 |
| Bacillota | *Paenibacillus antarcticus* | WP_068648876.1 | WP_068648878.1 | WP_068651994.1 |
| Bacillota | *Paenibacillus antibioticophila* | WP_212939118.1 | WP_212939119.1 | WP_044479289.1 |
| Bacillota | *Paenibacillus antri* | WP_138193362.1 | WP_138193361.1 | WP_158299477.1 |
| Bacillota | *Paenibacillus apiarius* | WP_087435421.1 | WP_087435420.1 | WP_206096386.1 |
| Bacillota | *Paenibacillus apii* | WP_165094357.1 | WP_165094360.1 | WP_168130059.1 |
| Bacillota | *Paenibacillus apis* | WP_301630174.1 | WP_301627953.1 | WP_301624390.1 |
| Bacillota | *Paenibacillus aquistagni* | WP_085493644.1 | WP_176228866.1 | WP_085492660.1 |
| Bacillota | *Paenibacillus arenilitoris* | WP_190861985.1 | WP_190861983.1 | WP_190862815.1 |
| Bacillota | *Paenibacillus arenosi* | WP_192024005.1 | WP_192024006.1 | WP_192023195.1 |
| Bacillota | *Paenibacillus artemisiicola* | WP_208847075.1 | WP_240545972.1 | WP_208846887.1 |
| Bacillota | *Paenibacillus assamensis* | WP_028593560.1 | WP_028593559.1 | WP_028595832.1 |
| Bacillota | *Paenibacillus athensensis* | WP_134750232.1 | WP_134750231.1 | WP_134757304.1 |
| Bacillota | *Paenibacillus auburnensis* | WP_236331603.1 | WP_236331605.1 | WP_236332662.1 |
| Bacillota | *Paenibacillus aurantiacus* | WP_377500029.1 | WP_377500027.1 | WP_377491245.1 |
| Bacillota | *Paenibacillus aurantius* | WP_315603029.1 | WP_315603030.1 | WP_315607290.1 |
| Bacillota | *Paenibacillus azoreducens* | WP_212979992.1 | WP_212979993.1 | WP_415998835.1 |

| | | | | |
|---|---|---|---|---|
| Bacillota | *Paenibacillus baekrokdamisoli* | WP_125656665.1 | WP_125656668.1 | WP_125657417.1 |
| Bacillota | *Paenibacillus barcinonensis* | WP_110894264.1 | WP_110894265.1 | WP_216517219.1 |
| Bacillota | *Paenibacillus barengoltzii* | WP_127576637.1 | WP_016311834.1 | WP_333556376.1 |
| Bacillota | *Paenibacillus beijingensis* | WP_045672563.1 | WP_045672564.1 | WP_045671783.1 |
| Bacillota | *Paenibacillus borealis* | WP_042215472.1 | WP_042215475.1 | WP_076109039.1 |
| Bacillota | *Paenibacillus bouchesdurhonensis* | WP_110932522.1 | WP_110932521.1 | WP_110932164.1 |
| Bacillota | *Paenibacillus bovis* | WP_060535533.1 | WP_060535534.1 | WP_060535797.1 |
| Bacillota | *Paenibacillus brasilensis* | WP_152381998.1 | WP_152381999.1 | WP_152378823.1 |
| Bacillota | *Paenibacillus brevis* | WP_216481004.1 | WP_216481005.1 | WP_216481043.1 |
| Bacillota | *Paenibacillus camerounensis* | WP_042196114.1 | WP_042204296.1 | WP_042197491.1 |
| Bacillota | *Paenibacillus castaneae* | WP_102710847.1 | WP_102710846.1 | WP_102712600.1 |
| Bacillota | *Paenibacillus catalpae* | WP_091189044.1 | WP_091189043.1 | WP_091180719.1 |
| Bacillota | *Paenibacillus caui* | WP_223066654.1 | WP_223066691.1 | WP_223066358.1 |
| Bacillota | *Paenibacillus cellulosilyticus* | WP_110043359.1 | WP_110043360.1 | WP_110046471.1 |
| Bacillota | *Paenibacillus cellulositrophicus* | WP_152398281.1 | WP_152398282.1 | WP_152398424.1 |
| Bacillota | *Paenibacillus chartarius* | WP_377472238.1 | WP_377472239.1 | WP_377471493.1 |
| Bacillota | *Paenibacillus chibensis* | WP_127607164.1 | WP_328277582.1 | WP_127601684.1 |
| Bacillota | *Paenibacillus chitinolyticus* | WP_405079470.1 | WP_387535989.1 | WP_377646096.1 |
| Bacillota | *Paenibacillus chungangensis* | WP_377562651.1 | WP_377562650.1 | WP_377568441.1 |
| Bacillota | *Paenibacillus cineris* | WP_212984345.1 | WP_212967858.1 | WP_212967761.1 |
| Bacillota | *Paenibacillus contaminans* | WP_113033634.1 | WP_113033636.1 | WP_113032225.1 |
| Bacillota | *Paenibacillus crassostreae* | WP_068657922.1 | WP_068657921.1 | WP_068659503.1 |
| Bacillota | *Paenibacillus cremeus* | WP_144842940.1 | WP_261381216.1 | WP_144847196.1 |
| Bacillota | *Paenibacillus curdlanolyticus* | WP_006039229.1 | WP_006039230.1 | WP_006036336.1 |
| Bacillota | *Paenibacillus cymbidii* | WP_135551466.1 | WP_135551469.1 | WP_135546844.1 |
| Bacillota | *Paenibacillus daejeonensis* | WP_020616645.1 | WP_020616646.1 | WP_020615593.1 |
| Bacillota | *Paenibacillus dakarensis* | WP_054957615.1 | WP_054957616.1 | WP_054956622.1 |
| Bacillota | *Paenibacillus dendritiformis* | WP_111154157.1 | WP_168181507.1 | WP_213469088.1 |

| | | | | |
|---|---|---|---|---|
| Bacillota | *Paenibacillus dendrobii* | WP_202128686.1 | WP_160498316.1 | WP_160498813.1 |
| Bacillota | *Paenibacillus dokdonensis* | WP_136606159.1 | WP_136606160.1 | WP_136606510.1 |
| Bacillota | *Paenibacillus donghaensis* | WP_087914496.1 | WP_194230145.1 | WP_087917553.1 |
| Bacillota | *Paenibacillus durus* | WP_042207282.1 | WP_046723361.1 | WP_046723444.1 |
| Bacillota | *Paenibacillus ehimensis* | WP_127485603.1 | WP_127485605.1 | WP_025853430.1 |
| Bacillota | *Paenibacillus elgii* | WP_010501027.1 | WP_127460064.1 | WP_163858210.1 |
| Bacillota | *Paenibacillus endophyticus* | WP_183561091.1 | WP_183561093.1 | WP_183563669.1 |
| Bacillota | *Paenibacillus endoradicis* | WP_258203218.1 | WP_258203219.1 | WP_258199419.1 |
| Bacillota | *Paenibacillus enshidis* | WP_375353491.1 | WP_375353492.1 | WP_375353779.1 |
| Bacillota | *Paenibacillus etheri* | WP_060622731.1 | WP_060622730.1 | WP_060622030.1 |
| Bacillota | *Paenibacillus eucommiae* | WP_209976595.1 | WP_209976594.1 | WP_209974125.1 |
| Bacillota | *Paenibacillus faecalis* | WP_106767939.1 | WP_106767940.1 | WP_106768318.1 |
| Bacillota | *Paenibacillus faecis* | WP_213504414.1 | WP_213504415.1 | WP_148451589.1 |
| Bacillota | *Paenibacillus farraposensis* | WP_229525501.1 | WP_229525500.1 | WP_229524566.1 |
| Bacillota | *Paenibacillus favisporus* | WP_163879295.1 | WP_326104187.1 | WP_163879530.1 |
| Bacillota | *Paenibacillus ferrarius* | WP_079420736.1 | WP_079420735.1 | WP_079420014.1 |
| Bacillota | *Paenibacillus filicis* | WP_341414414.1 | WP_341414415.1 | WP_341416827.1 |
| Bacillota | *Paenibacillus flagellatus* | WP_110839882.1 | WP_110839883.1 | WP_110840646.1 |
| Bacillota | *Paenibacillus foliorum* | WP_171655782.1 | WP_171655783.1 | WP_312886573.1 |
| Bacillota | *Paenibacillus fonticola* | WP_019640396.1 | WP_019640395.1 | WP_019636244.1 |
| Bacillota | *Paenibacillus forsythiae* | WP_025699477.1 | WP_025699479.1 | WP_025697264.1 |
| Bacillota | *Paenibacillus frigoriresistens* | WP_173184821.1 | WP_173184605.1 | WP_173186448.1 |
| Bacillota | *Paenibacillus gallinarum* | WP_191799275.1 | WP_191799569.1 | WP_191798882.1 |
| Bacillota | *Paenibacillus gansuensis* | WP_377601245.1 | WP_377601243.1 | WP_377602596.1 |
| Bacillota | *Paenibacillus germinis* | WP_171692528.1 | WP_171692346.1 | WP_171688446.1 |
| Bacillota | *Paenibacillus ginsengarvi* | WP_120750693.1 | WP_120750694.1 | WP_120745612.1 |
| Bacillota | *Paenibacillus ginsengihumi* | WP_019534029.1 | WP_019534030.1 | WP_019533053.1 |
| Bacillota | *Paenibacillus glacialis* | WP_068530497.1 | WP_068530494.1 | WP_068537660.1 |

| | | | | |
|---|---|---|---|---|
| Bacillota | *Paenibacillus glucanolyticus* | WP_152825198.1 | WP_076152431.1 | WP_340462073.1 |
| Bacillota | *Paenibacillus glycanilyticus* | WP_251653058.1 | WP_284239737.1 | WP_127498518.1 |
| Bacillota | *Paenibacillus glycinis* | WP_161740498.1 | WP_161740500.1 | WP_161740967.1 |
| Bacillota | *Paenibacillus graminis* | WP_025707470.1 | WP_025707469.1 | WP_025708368.1 |
| Bacillota | *Paenibacillus guangzhouensis* | WP_152393925.1 | WP_152393926.1 | WP_265333443.1 |
| Bacillota | *Paenibacillus gyeongsangnamensis* | WP_269881438.1 | WP_269881437.1 | WP_269879583.1 |
| Bacillota | *Paenibacillus haidiansis* | WP_331844562.1 | WP_331844563.1 | WP_331844893.1 |
| Bacillota | *Paenibacillus hamazuiensis* | WP_248925463.1 | WP_248925462.1 | WP_248926615.1 |
| Bacillota | *Paenibacillus harenae* | WP_307201541.1 | WP_307467111.1 | WP_028611235.1 |
| Bacillota | *Paenibacillus helianthi* | WP_074109058.1 | WP_074107562.1 | WP_074106746.1 |
| Bacillota | *Paenibacillus hemerocallicola* | WP_139604983.1 | WP_139604984.1 | WP_139602082.1 |
| Bacillota | *Paenibacillus herberti* | WP_176444729.1 | WP_089523697.1 | WP_176444650.1 |
| Bacillota | *Paenibacillus hexagrammi* | WP_235118659.1 | WP_235122911.1 | WP_235121698.1 |
| Bacillota | *Paenibacillus hodogayensis* | WP_344908541.1 | WP_344908543.1 | WP_344907027.1 |
| Bacillota | *Paenibacillus humicola* | WP_274651616.1 | WP_274651617.1 | WP_274651867.1 |
| Bacillota | *Paenibacillus hunanensis* | WP_188775443.1 | WP_320544467.1 | WP_320544258.1 |
| Bacillota | *Paenibacillus ihbetae* | WP_077565455.1 | WP_099476792.1 | WP_077565014.1 |
| Bacillota | *Paenibacillus ihuae* | WP_054941331.1 | WP_054941332.1 | WP_054941758.1 |
| Bacillota | *Paenibacillus ihumii* | WP_055107795.1 | WP_055110100.1 | WP_055107462.1 |
| Bacillota | *Paenibacillus illinoisensis* | WP_390582895.1 | WP_337033012.1 | WP_127537776.1 |
| Bacillota | *Paenibacillus jilunlii* | WP_062519683.1 | WP_062519685.1 | WP_062527005.1 |
| Bacillota | *Paenibacillus kobensis* | WP_127530721.1 | WP_127530722.1 | WP_127533050.1 |
| Bacillota | *Paenibacillus koleovorans* | WP_127588578.1 | WP_127588579.1 | WP_127582757.1 |
| Bacillota | *Paenibacillus kribbensis* | WP_179032748.1 | WP_179032749.1 | WP_068502150.1 |
| Bacillota | *Paenibacillus lacisoli* | WP_305022224.1 | WP_305022225.1 | WP_305022581.1 |
| Bacillota | *Paenibacillus larvae* | WP_268570009.1 | WP_023482678.1 | WP_079940202.1 |
| Bacillota | *Paenibacillus lautus* | WP_127594125.1 | WP_216530988.1 | WP_220686041.1 |
| Bacillota | *Paenibacillus lemnae* | WP_169506788.1 | WP_169506789.1 | WP_169503480.1 |

| | | | | |
|---|---|---|---|---|
| Bacillota | *Paenibacillus lentus* | WP_125083238.1 | WP_379392405.1 | WP_379392043.1 |
| Bacillota | *Paenibacillus lignilyticus* | WP_210659859.1 | WP_210659857.1 | WP_210659384.1 |
| Bacillota | *Paenibacillus lupini* | WP_167273182.1 | WP_167273178.1 | WP_167277696.1 |
| Bacillota | *Paenibacillus luteus* | WP_141501475.1 | WP_141501476.1 | WP_141505818.1 |
| Bacillota | *Paenibacillus lutimineralis* | WP_126997891.1 | WP_126997889.1 | WP_126997225.1 |
| Bacillota | *Paenibacillus lutrae* | WP_157335440.1 | WP_157335441.1 | WP_157334343.1 |
| Bacillota | *Paenibacillus lycopersici* | WP_162356891.1 | WP_162356889.1 | WP_162356475.1 |
| Bacillota | *Paenibacillus macerans* | WP_251586340.1 | WP_316560952.1 | WP_036621380.1 |
| Bacillota | *Paenibacillus macquariensis* | WP_068590251.1 | WP_068590249.1 | WP_068587569.1 |
| Bacillota | *Paenibacillus mangrovi* | WP_244721110.1 | WP_244721114.1 | WP_244722015.1 |
| Bacillota | *Paenibacillus marchantiophytorum* | WP_189009728.1 | WP_189014345.1 | WP_189006243.1 |
| Bacillota | *Paenibacillus massiliensis* | WP_028589105.1 | WP_028589104.1 | WP_018886771.1 |
| Bacillota | *Paenibacillus maysiensis* | WP_025685875.1 | WP_025685876.1 | WP_025685742.1 |
| Bacillota | *Paenibacillus medicaginis* | WP_375521638.1 | WP_375521639.1 | WP_375518551.1 |
| Bacillota | *Paenibacillus mellifer* | WP_248550581.1 | WP_248550580.1 | WP_248551931.1 |
| Bacillota | *Paenibacillus mendelii* | WP_204816264.1 | WP_204816265.1 | WP_204816513.1 |
| Bacillota | *Paenibacillus mesophilus* | WP_138878350.1 | WP_138878351.1 | WP_138882220.1 |
| Bacillota | *Paenibacillus mesotrionivorans* | WP_416214544.1 | WP_416214545.1 | WP_416219599.1 |
| Bacillota | *Paenibacillus methanolicus* | WP_148928224.1 | WP_148928225.1 | WP_148928442.1 |
| Bacillota | *Paenibacillus montanisoli* | WP_112882578.1 | WP_112882579.1 | WP_112882883.1 |
| Bacillota | *Paenibacillus montaniterrae* | WP_213513802.1 | WP_213513264.1 | WP_246563468.1 |
| Bacillota | *Paenibacillus monticola* | WP_154118393.1 | WP_154118394.1 | WP_154121344.1 |
| Bacillota | *Paenibacillus motobuensis* | WP_343863633.1 | WP_343863631.1 | WP_343865702.1 |
| Bacillota | *Paenibacillus mucilaginosus* | WP_013916540.1 | WP_014370053.1 | WP_236628849.1 |
| Bacillota | *Paenibacillus nanensis* | WP_119599127.1 | WP_119599126.1 | WP_119598178.1 |
| Bacillota | *Paenibacillus naphthalenovorans* | WP_175472024.1 | WP_074729390.1 | WP_074727048.1 |
| Bacillota | *Paenibacillus nasutitermitis* | WP_188992611.1 | WP_188992612.1 | WP_188992857.1 |
| Bacillota | *Paenibacillus nicotianae* | WP_204823540.1 | WP_204823539.1 | WP_204823265.1 |

| | | | | |
|---|---|---|---|---|
| Bacillota | *Paenibacillus oceani* | WP_190925107.1 | WP_190925109.1 | WP_190926136.1 |
| Bacillota | *Paenibacillus odorifer* | WP_076284576.1 | WP_248548496.1 | WP_111505257.1 |
| Bacillota | *Paenibacillus oenotherae* | WP_219870643.1 | WP_219870644.1 | WP_219870874.1 |
| Bacillota | *Paenibacillus oleatilyticus* | WP_216792181.1 | WP_373949330.1 | WP_373955658.1 |
| Bacillota | *Paenibacillus oralis* | WP_128634089.1 | WP_128634090.1 | WP_128634460.1 |
| Bacillota | *Paenibacillus oryzae* | WP_068683840.1 | WP_068683842.1 | WP_068684506.1 |
| Bacillota | *Paenibacillus oryzisoli* | WP_068670283.1 | WP_068670284.1 | WP_068670715.1 |
| Bacillota | *Paenibacillus ottowii* | WP_064796416.1 | WP_326400406.1 | WP_305164649.1 |
| Bacillota | *Paenibacillus pabuli* | WP_247898189.1 | WP_426251136.1 | WP_062326419.1 |
| Bacillota | *Paenibacillus paeoniae* | WP_116044696.1 | WP_116044698.1 | WP_240644312.1 |
| Bacillota | *Paenibacillus paridis* | WP_139999520.1 | WP_139999522.1 | WP_139997909.1 |
| Bacillota | *Paenibacillus pasadenensis* | WP_251560931.1 | WP_251560930.1 | WP_101809358.1 |
| Bacillota | *Paenibacillus pectinilyticus* | WP_065857228.1 | WP_065857226.1 | WP_244163060.1 |
| Bacillota | *Paenibacillus pedocola* | WP_310829302.1 | WP_310829303.1 | WP_310829725.1 |
| Bacillota | *Paenibacillus peoriae* | WP_010345798.1 | WP_076293050.1 | WP_348623108.1 |
| Bacillota | *Paenibacillus periandrae* | WP_240413846.1 | WP_240413847.1 | WP_240417094.1 |
| Bacillota | *Paenibacillus phocaensis* | WP_068785288.1 | WP_068785287.1 | WP_068784959.1 |
| Bacillota | *Paenibacillus phyllosphaerae* | WP_183598253.1 | WP_183598251.1 | WP_246427471.1 |
| Bacillota | *Paenibacillus physcomitrellae* | WP_094093303.1 | WP_094093304.1 | WP_094093583.1 |
| Bacillota | *Paenibacillus phytohabitans* | WP_171716664.1 | WP_171716665.1 | WP_171721012.1 |
| Bacillota | *Paenibacillus phytorum* | WP_171643358.1 | WP_171643359.1 | WP_171648381.1 |
| Bacillota | *Paenibacillus pinihumi* | WP_028561258.1 | WP_028561259.1 | WP_028561628.1 |
| Bacillota | *Paenibacillus pinisoli* | WP_120113277.1 | WP_424452460.1 | WP_243643891.1 |
| Bacillota | *Paenibacillus pinistramenti* | WP_138492936.1 | WP_138492937.1 | WP_138493228.1 |
| Bacillota | *Paenibacillus piri* | WP_133225913.1 | WP_133225912.1 | WP_342774573.1 |
| Bacillota | *Paenibacillus piscarius* | WP_238652982.1 | WP_238652983.1 | WP_238654633.1 |
| Bacillota | *Paenibacillus plantarum* | WP_171628794.1 | WP_171628795.1 | WP_171635300.1 |
| Bacillota | *Paenibacillus planticolens* | WP_171682055.1 | WP_171682056.1 | WP_246294449.1 |

| | | | | |
|---|---|---|---|---|
| Bacillota | *Paenibacillus plantiphilus* | WP_236344839.1 | WP_371877750.1 | WP_236344089.1 |
| Bacillota | *Paenibacillus polymyxa* | WP_311078506.1 | WP_215076021.1 | WP_075154454.1 |
| Bacillota | *Paenibacillus polysaccharolyticus* | WP_251505656.1 | WP_251505659.1 | WP_253492605.1 |
| Bacillota | *Paenibacillus popilliae* | WP_006285465.1 | WP_142543324.1 | WP_142543523.1 |
| Bacillota | *Paenibacillus prosopidis* | WP_181873345.1 | WP_114378576.1 | WP_114381166.1 |
| Bacillota | *Paenibacillus protaetiae* | WP_129439405.1 | WP_129439407.1 | WP_129444004.1 |
| Bacillota | *Paenibacillus pseudetheri* | WP_234531948.1 | WP_234531942.1 | WP_234533213.1 |
| Bacillota | *Paenibacillus psychroresistens* | WP_155701927.1 | WP_155701928.1 | WP_155700801.1 |
| Bacillota | *Paenibacillus puerhi* | WP_159883059.1 | WP_159883057.1 | WP_235941410.1 |
| Bacillota | *Paenibacillus puldeungensis* | WP_379321196.1 | WP_379321195.1 | WP_379318034.1 |
| Bacillota | *Paenibacillus qinlingensis* | WP_310223539.1 | WP_173110235.1 | WP_310226997.1 |
| Bacillota | *Paenibacillus radicibacter* | WP_258206429.1 | WP_258206430.1 | WP_258206078.1 |
| Bacillota | *Paenibacillus radicis (ex Gao et al. 2016)* | WP_188887732.1 | WP_188887731.1 | WP_229692423.1 |
| Bacillota | *Paenibacillus rhizophilus* | WP_124693619.1 | WP_124693618.1 | WP_124695680.1 |
| Bacillota | *Paenibacillus rhizoplanae* | WP_209994246.1 | WP_209993887.1 | WP_209987956.1 |
| Bacillota | *Paenibacillus rhizosphaerae* | WP_183581975.1 | WP_183581974.1 | WP_183581722.1 |
| Bacillota | *Paenibacillus rhizovicinus* | WP_162638887.1 | WP_162638888.1 | WP_162639117.1 |
| Bacillota | *Paenibacillus rigui* | WP_094013285.1 | WP_094013286.1 | WP_094015209.1 |
| Bacillota | *Paenibacillus riograndensis* | WP_020427657.1 | WP_020427658.1 | WP_020427852.1 |
| Bacillota | *Paenibacillus sabinae* | WP_025335550.1 | WP_025335551.1 | WP_025335875.1 |
| Bacillota | *Paenibacillus sabuli* | WP_190916452.1 | WP_190916454.1 | WP_190915075.1 |
| Bacillota | *Paenibacillus sacheonensis* | WP_161696576.1 | WP_161696574.1 | WP_161703033.1 |
| Bacillota | *Paenibacillus sambharensis* | WP_111149231.1 | WP_111149230.1 | WP_111148711.1 |
| Bacillota | *Paenibacillus sanguinis* | WP_018751371.1 | WP_018751372.1 | WP_018753264.1 |
| Bacillota | *Paenibacillus sedimenti* | WP_188174342.1 | WP_188174343.1 | WP_188172947.1 |
| Bacillota | *Paenibacillus sediminis* | WP_209845359.1 | WP_209845357.1 | WP_209844695.1 |
| Bacillota | *Paenibacillus segetis* | WP_188538934.1 | WP_188538936.1 | WP_188539579.1 |
| Bacillota | *Paenibacillus selenitireducens* | WP_078497211.1 | WP_078497210.1 | WP_078498745.1 |

| | | | | |
|---|---|---|---|---|
| Bacillota | *Paenibacillus senegalensis* | WP_026021375.1 | WP_010273795.1 | WP_010270149.1 |
| Bacillota | *Paenibacillus senegalimassiliensis* | WP_059052566.1 | WP_059052564.1 | WP_059052170.1 |
| Bacillota | *Paenibacillus septentrionalis* | WP_379233282.1 | WP_379232422.1 | WP_379230547.1 |
| Bacillota | *Paenibacillus sepulcri* | WP_210041358.1 | WP_210041361.1 | WP_210037664.1 |
| Bacillota | *Paenibacillus shirakamiensis* | WP_209858989.1 | WP_209858987.1 | WP_209858427.1 |
| Bacillota | *Paenibacillus silagei* | WP_209874571.1 | WP_209869954.1 | WP_209871997.1 |
| Bacillota | *Paenibacillus silvae* | WP_111271384.1 | WP_188591051.1 | WP_111271909.1 |
| Bacillota | *Paenibacillus silvestris* | WP_161408517.1 | WP_161408564.1 | WP_235959055.1 |
| Bacillota | *Paenibacillus silvisoli* | WP_308639168.1 | WP_308639167.1 | WP_308638903.1 |
| Bacillota | *Paenibacillus silviterrae* | WP_284643950.1 | WP_284640174.1 | WP_284638745.1 |
| Bacillota | *Paenibacillus sinopodophylli* | WP_138750981.1 | WP_138750980.1 | WP_138751928.1 |
| Bacillota | *Paenibacillus solanacearum* | WP_218091182.1 | WP_218091183.1 | WP_218090647.1 |
| Bacillota | *Paenibacillus solani* | WP_054401724.1 | WP_054401723.1 | WP_054401438.1 |
| Bacillota | *Paenibacillus solisilvae* | WP_379189360.1 | WP_379189361.1 | WP_379190327.1 |
| Bacillota | *Paenibacillus sonchi* | WP_233183868.1 | WP_039835064.1 | WP_233184314.1 |
| Bacillota | *Paenibacillus sophorae* | WP_036594357.1 | WP_036594356.1 | WP_036596753.1 |
| Bacillota | *Paenibacillus soyae* | WP_257442904.1 | WP_257442906.1 | WP_257444241.1 |
| Bacillota | *Paenibacillus sp.* | WP_325352875.1 | WP_313639716.1 | WP_313642091.1 |
| Bacillota | *Paenibacillus spongiae* | WP_258388478.1 | WP_258388477.1 | WP_258388242.1 |
| Bacillota | *Paenibacillus stellifer* | WP_038697001.1 | WP_038697002.1 | WP_038697658.1 |
| Bacillota | *Paenibacillus swuensis* | WP_068609185.1 | WP_068609188.1 | WP_231891263.1 |
| Bacillota | *Paenibacillus taichungensis* | WP_413372314.1 | WP_376037005.1 | WP_175383163.1 |
| Bacillota | *Paenibacillus taihuensis* | WP_116187053.1 | WP_116187052.1 | WP_116187460.1 |
| Bacillota | *Paenibacillus taiwanensis* | WP_028544285.1 | WP_028544284.1 | WP_245596021.1 |
| Bacillota | *Paenibacillus tarimensis* | WP_235285858.1 | WP_235285857.1 | WP_235285404.1 |
| Bacillota | *Paenibacillus tengchongensis* | WP_151734500.1 | WP_151734499.1 | WP_151735851.1 |
| Bacillota | *Paenibacillus tepidiphilus* | WP_150273581.1 | WP_150273580.1 | WP_150266292.1 |
| Bacillota | *Paenibacillus terrae* | WP_014282065.1 | WP_145158499.1 | WP_014282450.1 |

| | | | | |
|---|---|---|---|---|
| Bacillota | *Paenibacillus terreus* | WP_375523607.1 | WP_375523606.1 | WP_375526893.1 |
| Bacillota | *Paenibacillus terricola* | WP_191203123.1 | WP_191203122.1 | WP_191202242.1 |
| Bacillota | *Paenibacillus terrigena* | WP_018755349.1 | WP_314585441.1 | WP_018756691.1 |
| Bacillota | *Paenibacillus thailandensis* | WP_379275504.1 | WP_379275508.1 | WP_379273623.1 |
| Bacillota | *Paenibacillus thalictri* | WP_131012956.1 | WP_131012957.1 | WP_131014125.1 |
| Bacillota | *Paenibacillus thermoaerophilus* | WP_138787912.1 | WP_138787913.1 | WP_170209381.1 |
| Bacillota | *Paenibacillus thermotolerans* | WP_274364624.1 | WP_274364693.1 | WP_274363310.1 |
| Bacillota | *Paenibacillus thiaminolyticus* | WP_143800797.1 | WP_374018712.1 | WP_087442146.1 |
| Bacillota | *Paenibacillus tianjinensis* | WP_206100817.1 | WP_206100818.1 | WP_206101483.1 |
| Bacillota | *Paenibacillus tianmuensis* | WP_090665888.1 | WP_090665886.1 | WP_090670061.1 |
| Bacillota | *Paenibacillus timonensis* | WP_240268406.1 | WP_240268405.1 | WP_240269054.1 |
| Bacillota | *Paenibacillus tritici* | WP_173129013.1 | WP_173129016.1 | WP_211720737.1 |
| Bacillota | *Paenibacillus tundrae* | WP_307212880.1 | WP_307212877.1 | WP_338540183.1 |
| Bacillota | *Paenibacillus turicensis* | WP_210091285.1 | WP_210091286.1 | WP_210088043.1 |
| Bacillota | *Paenibacillus turpanensis* | WP_166239436.1 | WP_166239434.1 | WP_166243811.1 |
| Bacillota | *Paenibacillus typhae* | WP_090714162.1 | WP_221798010.1 | WP_090712285.1 |
| Bacillota | *Paenibacillus tyrfis* | WP_253409611.1 | WP_088833390.1 | WP_281940216.1 |
| Bacillota | *Paenibacillus uliginis* | WP_208914519.1 | WP_208914520.1 | WP_208915055.1 |
| Bacillota | *Paenibacillus vandeheii* | WP_301246512.1 | WP_301246510.1 | WP_301243803.1 |
| Bacillota | *Paenibacillus vietnamensis* | WP_224722471.1 | WP_224722472.1 | WP_224721890.1 |
| Bacillota | *Paenibacillus vini* | WP_213654996.1 | WP_213654995.1 | WP_213653843.1 |
| Bacillota | *Paenibacillus wenxiniae* | WP_347325917.1 | WP_347325918.1 | WP_347325656.1 |
| Bacillota | *Paenibacillus whitsoniae* | WP_126140510.1 | WP_126140511.1 | WP_126143135.1 |
| Bacillota | *Paenibacillus woosongensis* | WP_155610100.1 | WP_283924922.1 | WP_155610841.1 |
| Bacillota | *Paenibacillus wulumuqiensis* | WP_046215039.1 | WP_046215038.1 | WP_046214747.1 |
| Bacillota | *Paenibacillus wynnii* | WP_036657105.1 | WP_307586656.1 | WP_036657888.1 |
| Bacillota | *Paenibacillus xanthanilyticus* | WP_377722267.1 | WP_377722268.1 | WP_377722477.1 |
| Bacillota | *Paenibacillus xerothermodurans* | WP_089199061.1 | WP_089199062.1 | WP_243633136.1 |

| | | | | |
|---|---|---|---|---|
| Bacillota | *Paenibacillus xylanexedens* | WP_419889253.1 | WP_124116226.1 | WP_145332067.1 |
| Bacillota | *Paenibacillus xylaniclasticus* | WP_127566875.1 | WP_127566877.1 | WP_127567498.1 |
| Bacillota | *Paenibacillus xylanilyticus* | WP_418038329.1 | WP_413035371.1 | WP_175397529.1 |
| Bacillota | *Paenibacillus xylanivorans* | WP_053780711.1 | WP_053780710.1 | WP_053781229.1 |
| Bacillota | *Paenibacillus yanchengensis* | WP_377771597.1 | WP_377772049.1 | WP_377772207.1 |
| Bacillota | *Paenibacillus yonginensis* | WP_068695883.1 | WP_068695885.1 | WP_068696453.1 |
| Bacillota | *Paenibacillus zanthoxyli* | WP_025692677.1 | WP_025692678.1 | WP_025691633.1 |
| Bacillota | *Paenibacillus zeirhizosphaerae* | WP_305754248.1 | WP_305754249.1 | WP_305754549.1 |
| Bacillota | *Paenibacillus zeisoli* | WP_127197751.1 | WP_127197752.1 | WP_127198081.1 |
| Bacillota | *Paenisporosarcina antarctica* | WP_134209959.1 | WP_134210720.1 | WP_134210153.1 |
| Bacillota | *Paenisporosarcina cavernae* | WP_119883146.1 | WP_119882510.1 | WP_119882894.1 |
| Bacillota | *Paenisporosarcina indica* | WP_075617693.1 | WP_075618437.1 | WP_075617467.1 |
| Bacillota | *Paenisporosarcina macmurdoensis* | WP_377732563.1 | WP_377735773.1 | WP_377732245.1 |
| Bacillota | *Paenisporosarcina quisquiliarum* | WP_394191999.1 | WP_269927160.1 | WP_394187284.1 |
| Bacillota | *Paenisporosarcina sp.* | WP_424473098.1 | WP_424472036.1 | WP_424415549.1 |
| Bacillota | *Pallidibacillus pasinlerensis* | WP_161919234.1 | WP_161920773.1 | WP_161919021.1 |
| Bacillota | *Pallidibacillus thermolactis* | WP_173660149.1 | WP_263061301.1 | WP_173662503.1 |
| Bacillota | *Paludifilum halophilum* | WP_094262789.1 | WP_094262790.1 | WP_240511973.1 |
| Bacillota | *Parageobacillus genomosp. 1* | WP_043905239.1 | WP_043905240.1 | WP_043904021.1 |
| Bacillota | *Parageobacillus sp. VR-IP* | WP_175245149.1 | WP_175245148.1 | WP_175243359.1 |
| Bacillota | *Parageobacillus thermantarcticus* | WP_090948791.1 | WP_208601770.1 | WP_090948889.1 |
| Bacillota | *Parageobacillus thermoglucosidasius* | WP_003249633.1 | WP_064550242.1 | WP_064551126.1 |
| Bacillota | *Parageobacillus toebii* | WP_205424629.1 | WP_374718483.1 | WP_374720214.1 |
| Bacillota | *Paraliobacillus quinghaiensis* | WP_117152167.1 | WP_117155941.1 | WP_117153002.1 |
| Bacillota | *Paraliobacillus ryukyuensis* | WP_079709683.1 | WP_113868836.1 | WP_079709353.1 |
| Bacillota | *Paraliobacillus salinarum* | WP_182199625.1 | WP_182201226.1 | WP_182199062.1 |
| Bacillota | *Paraliobacillus sediminis* | WP_117169373.1 | WP_117168819.1 | WP_117169811.1 |
| Bacillota | *Paraliobacillus sp. JSM ZJ581* | WP_374065147.1 | WP_374067258.1 | WP_374067853.1 |

| | | | | |
|---|---|---|---|---|
| Bacillota | *Paranoxybacillus vitaminiphilus* | WP_111643996.1 | WP_111643997.1 | WP_111643621.1 |
| Bacillota | *Paucisalibacillus globulus* | WP_096270445.1 | WP_026909002.1 | WP_096269978.1 |
| Bacillota | *Paucisalibacillus sp. EB02* | WP_042145630.1 | WP_042142703.1 | WP_042145019.1 |
| Bacillota | *Peribacillus acanthi* | WP_108671147.1 | WP_108671146.1 | WP_108669794.1 |
| Bacillota | *Peribacillus alkalitolerans* | WP_163100974.1 | WP_163100975.1 | WP_163101404.1 |
| Bacillota | *Peribacillus asahii* | WP_127761156.1 | WP_252286581.1 | WP_252289885.1 |
| Bacillota | *Peribacillus butanolivorans* | WP_328114255.1 | WP_402878124.1 | WP_053345228.1 |
| Bacillota | *Peribacillus castrilensis* | WP_367409646.1 | WP_367407233.1 | WP_367407051.1 |
| Bacillota | *Peribacillus cavernae* | WP_126864739.1 | WP_126863591.1 | WP_126864368.1 |
| Bacillota | *Peribacillus deserti* | WP_101645019.1 | WP_204542463.1 | WP_204541793.1 |
| Bacillota | *Peribacillus faecalis* | WP_190999950.1 | WP_190999949.1 | WP_190996378.1 |
| Bacillota | *Peribacillus frigoritolerans* | WP_406593472.1 | WP_260358989.1 | WP_289352261.1 |
| Bacillota | *Peribacillus glennii* | WP_117321694.1 | WP_233522557.1 | WP_117322745.1 |
| Bacillota | *Peribacillus huizhouensis* | WP_182503146.1 | WP_182503147.1 | WP_182501307.1 |
| Bacillota | *Peribacillus kribbensis* | WP_026694847.1 | WP_155890147.1 | WP_026692131.1 |
| Bacillota | *Peribacillus loiseleuriae* | WP_049681938.1 | WP_049681939.1 | WP_049680679.1 |
| Bacillota | *Peribacillus muralis* | WP_064463009.1 | WP_377363119.1 | WP_427674549.1 |
| Bacillota | *Peribacillus psychrosaccharolyticus* | WP_040376265.1 | WP_040376266.1 | WP_040374614.1 |
| Bacillota | *Peribacillus saganii* | WP_199478491.1 | WP_117327000.1 | WP_117325393.1 |
| Bacillota | *Peribacillus simplex* | WP_347939832.1 | WP_352920322.1 | WP_144527054.1 |
| Bacillota | *Peribacillus sp. NJ11* | WP_289334827.1 | WP_289334826.1 | WP_289335970.1 |
| Bacillota | *Peribacillus tepidiphilus* | WP_374722489.1 | WP_374722488.1 | WP_153124242.1 |
| Bacillota | *Perspicuibacillus lycopersici* | WP_263072930.1 | WP_263072726.1 | WP_263071344.1 |
| Bacillota | *Phocicoccus pinnipedialis* | WP_186077510.1 | WP_377275301.1 | WP_186078057.1 |
| Bacillota | *Phocicoccus schoeneichii* | WP_186087927.1 | WP_186084444.1 | WP_186086908.1 |
| Bacillota | *Phosphitispora fastidiosa* | WP_231685226.1 | WP_231685227.1 | WP_231685477.1 |
| Bacillota | *Piscibacillus halophilus* | WP_091772403.1 | WP_175614577.1 | WP_091771979.1 |
| Bacillota | *Piscibacillus salipiscarius* | WP_377330544.1 | WP_054751699.1 | WP_377327791.1 |

| | | | | |
|---|---|---|---|---|
| Bacillota | *Planifilum fimeticola* | WP_106345631.1 | WP_106345581.1 | WP_106344368.1 |
| Bacillota | *Planifilum fulgidum* | WP_092035968.1 | WP_092035967.1 | WP_092041673.1 |
| Bacillota | *Planococcus antarcticus* | WP_040852325.1 | WP_006829302.1 | WP_006828840.1 |
| Bacillota | *Planococcus chinensis* | WP_224076780.1 | WP_224076704.1 | WP_204892578.1 |
| Bacillota | *Planococcus dechangensis* | WP_377277442.1 | WP_377278760.1 | WP_377277778.1 |
| Bacillota | *Planococcus donghaensis* | WP_394121751.1 | WP_394121384.1 | WP_065525910.1 |
| Bacillota | *Planococcus glaciei* | WP_200904932.1 | WP_036802959.1 | WP_053166399.1 |
| Bacillota | *Planococcus halocryophilus* | WP_240747830.1 | WP_008496839.1 | WP_008498021.1 |
| Bacillota | *Planococcus halotolerans* | WP_112222673.1 | WP_135815217.1 | WP_233785852.1 |
| Bacillota | *Planococcus kocurii* | WP_058385059.1 | WP_058384407.1 | WP_058384864.1 |
| Bacillota | *Planococcus lenghuensis* | WP_077589509.1 | WP_077590103.1 | WP_077589700.1 |
| Bacillota | *Planococcus liqunii* | WP_300981076.1 | WP_301725568.1 | WP_300981278.1 |
| Bacillota | *Planococcus maitriensis* | WP_205853838.1 | WP_112230374.1 | WP_112231356.1 |
| Bacillota | *Planococcus maritimus* | WP_068465707.1 | WP_068464323.1 | WP_068486073.1 |
| Bacillota | *Planococcus notacanthi* | WP_290214263.1 | WP_290214969.1 | WP_290214207.1 |
| Bacillota | *Planococcus plakortidis* | WP_411361501.1 | WP_405313062.1 | WP_411361557.1 |
| Bacillota | *Planococcus salinarum* | WP_117303781.1 | WP_071151755.1 | WP_117301439.1 |
| Bacillota | *Planococcus salinus* | WP_192892912.1 | WP_123164587.1 | WP_123164382.1 |
| Bacillota | *Planococcus shenhongbingii* | WP_301856043.1 | WP_300990175.1 | WP_301854711.1 |
| Bacillota | *Planococcus shixiaomingii* | WP_301722914.1 | WP_300985885.1 | WP_300985285.1 |
| Bacillota | *Planococcus soli* | WP_142826438.1 | WP_142827080.1 | WP_142826638.1 |
| Bacillota | *Planococcus sp. CAU13* | WP_033541215.1 | WP_033541336.1 | WP_033543526.1 |
| Bacillota | *Planococcus versutus* | WP_065524488.1 | WP_049692979.1 | WP_049693354.1 |
| Bacillota | *Planomicrobium okeanokoites* | WP_117312365.1 | WP_117313702.1 | WP_084244977.1 |
| Bacillota | *Planomicrobium soli* | WP_106531759.1 | WP_106532793.1 | WP_106534878.1 |
| Bacillota | *Planomicrobium sp. Y74* | WP_121632147.1 | WP_121635614.1 | WP_121633166.1 |
| Bacillota | *Planomicrobium stackebrandtii* | WP_308785862.1 | WP_308786454.1 | WP_308786093.1 |
| Bacillota | *Polycladomyces abyssicola* | WP_212772256.1 | WP_212772255.1 | WP_212772617.1 |

| | | | | |
|---|---|---|---|---|
| Bacillota | *Polycladomyces subterraneus* | WP_301237983.1 | WP_301237984.1 | WP_301239880.1 |
| Bacillota | *Polycladomyces zharkentensis* | WP_205494863.1 | WP_302104637.1 | WP_205492524.1 |
| Bacillota | *Polycladospora coralii* | WP_347239741.1 | WP_191138248.1 | WP_191140021.1 |
| Bacillota | *Pontibacillus chungwhensis* | WP_036779359.1 | WP_036782886.1 | WP_036778420.1 |
| Bacillota | *Pontibacillus halophilus* | WP_026799494.1 | WP_026800003.1 | WP_026799673.1 |
| Bacillota | *Pontibacillus litoralis* | WP_036832895.1 | WP_036834667.1 | WP_084600106.1 |
| Bacillota | *Pontibacillus marinus* | WP_027447180.1 | WP_027445858.1 | WP_027448187.1 |
| Bacillota | *Pontibacillus salicampi* | WP_377345715.1 | WP_377351052.1 | WP_377347591.1 |
| Bacillota | *Pontibacillus salipaludis* | WP_188650249.1 | WP_188656141.1 | WP_345238777.1 |
| Bacillota | *Pontibacillus sp. HMF3514* | WP_160101657.1 | WP_160103686.1 | WP_160100870.1 |
| Bacillota | *Pontibacillus yanchengensis* | WP_036818586.1 | WP_160848860.1 | WP_036820012.1 |
| Bacillota | *Pradoshia eiseniae* | WP_104848023.1 | WP_104848022.1 | WP_104849624.1 |
| Bacillota | *Priestia abyssalis* | WP_078409368.1 | WP_078409367.1 | WP_078413473.1 |
| Bacillota | *Priestia aryabhattai* | WP_216142501.1 | WP_226555819.1 | WP_048019823.1 |
| Bacillota | *Priestia endophytica* | WP_268582576.1 | WP_111925402.1 | WP_113749129.1 |
| Bacillota | *Priestia filamentosa* | WP_243513722.1 | WP_120046138.1 | WP_019391719.1 |
| Bacillota | *Priestia flexa* | WP_277716739.1 | WP_281998771.1 | WP_210609843.1 |
| Bacillota | *Priestia koreensis* | WP_367018079.1 | WP_336882656.1 | WP_053403380.1 |
| Bacillota | *Priestia megaterium* | WP_289522846.1 | WP_414184424.1 | WP_097824533.1 |
| Bacillota | *Priestia sp. 179-F W1.4 NHS* | WP_406900744.1 | WP_406901522.1 | WP_406900855.1 |
| Bacillota | *Priestia taiwanensis* | WP_188386729.1 | WP_188386728.1 | WP_188387811.1 |
| Bacillota | *Pseudalkalibacillus berkeleyi* | WP_236333256.1 | WP_236333258.1 | WP_236332239.1 |
| Bacillota | *Pseudalkalibacillus caeni* | WP_138128327.1 | WP_138128328.1 | WP_138129213.1 |
| Bacillota | *Pseudalkalibacillus decolorationis* | WP_257351784.1 | WP_257351783.1 | WP_257350563.1 |
| Bacillota | *Pseudalkalibacillus hwajinpoensis* | WP_136947247.1 | WP_224880743.1 | WP_347550457.1 |
| Bacillota | *Pseudalkalibacillus salsuginis* | WP_236310908.1 | WP_236310910.1 | WP_236313441.1 |
| Bacillota | *Pseudalkalibacillus sedimenti* | WP_273832483.1 | WP_273835075.1 | WP_273834772.1 |
| Bacillota | *Pseudalkalibacillus sp. A8* | WP_408006508.1 | WP_408006509.1 | WP_408010883.1 |

| | | | | |
|---|---|---|---|---|
| Bacillota | *Pseudalkalibacillus spartinae* | WP_273852285.1 | WP_273852284.1 | WP_273850317.1 |
| Bacillota | *Pseudogracilibacillus auburnensis* | WP_110394591.1 | WP_110394227.1 | WP_207520801.1 |
| Bacillota | *Pseudogracilibacillus sp. SO10305* | WP_336265594.1 | WP_336265976.1 | WP_336265821.1 |
| Bacillota | *Pseudoneobacillus rhizosphaerae* | WP_230496521.1 | WP_230496522.1 | WP_230495772.1 |
| Bacillota | *Psychrobacillus antarcticus* | WP_277585418.1 | WP_277586887.1 | WP_277585738.1 |
| Bacillota | *Psychrobacillus insolitus* | WP_111438313.1 | WP_111439258.1 | WP_170122323.1 |
| Bacillota | *Psychrobacillus lasiicapitis* | WP_142537194.1 | WP_142538007.1 | WP_170206445.1 |
| Bacillota | *Psychrobacillus mangrovi* | WP_336495936.1 | WP_336496523.1 | WP_336498714.1 |
| Bacillota | *Psychrobacillus psychrodurans* | WP_269920920.1 | WP_269920784.1 | WP_427138498.1 |
| Bacillota | *Psychrobacillus psychrotolerans* | WP_350473437.1 | WP_422194696.1 | WP_350472967.1 |
| Bacillota | *Psychrobacillus soli* | WP_142607215.1 | WP_142608489.1 | WP_185907990.1 |
| Bacillota | *Psychrobacillus sp.* | WP_313890972.1 | WP_313893284.1 | WP_313891380.1 |
| Bacillota | *Psychrobacillus vulpis* | WP_142642666.1 | WP_142640994.1 | WP_166462473.1 |
| Bacillota | *Pueribacillus sp. YX66* | WP_379668435.1 | WP_379668182.1 | WP_379668012.1 |
| Bacillota | *Pueribacillus theae* | WP_116553981.1 | WP_116554940.1 | WP_116552904.1 |
| Bacillota | *Pullulanibacillus camelliae* | WP_188692193.1 | WP_188692495.1 | WP_188692993.1 |
| Bacillota | *Pullulanibacillus pueri* | WP_188496588.1 | WP_188496589.1 | WP_229745355.1 |
| Bacillota | *Pullulanibacillus sp. KACC 23026* | WP_275280749.1 | WP_275280751.1 | WP_275281084.1 |
| Bacillota | *Radiobacillus deserti* | WP_143894017.1 | WP_143895601.1 | WP_143893315.1 |
| Bacillota | *Radiobacillus kanasensis* | WP_231097091.1 | WP_231094100.1 | WP_231096654.1 |
| Bacillota | *Radiobacillus sp. PE A8.2* | WP_407268678.1 | WP_407270775.1 | WP_407268083.1 |
| Bacillota | *Risungbinella massiliensis* | WP_044640190.1 | WP_044640263.1 | WP_044642465.1 |
| Bacillota | *Robertmurraya andreesenii* | WP_307149767.1 | WP_307149769.1 | WP_307150434.1 |
| Bacillota | *Robertmurraya korlensis* | WP_251514695.1 | WP_251514693.1 | WP_251515900.1 |
| Bacillota | *Robertmurraya kyonggiensis* | WP_136830600.1 | WP_136830601.1 | WP_136829269.1 |
| Bacillota | *Robertmurraya mangrovi* | WP_322447533.1 | WP_322447534.1 | WP_322444982.1 |
| Bacillota | *Robertmurraya massiliosenegalensis* | WP_019156012.1 | WP_019156013.1 | WP_019152893.1 |
| Bacillota | *Robertmurraya sp.* | WP_391559015.1 | WP_391559014.1 | WP_119707447.1 |

| | | | | |
|---|---|---|---|---|
| Bacillota | *Robertmurraya yapensis* | WP_126405731.1 | WP_126409130.1 | WP_126408518.1 |
| Bacillota | *Rossellomorea aquimaris* | WP_064091048.1 | WP_226672924.1 | WP_226674332.1 |
| Bacillota | *Rossellomorea arthrocnemi* | WP_201713300.1 | WP_201713301.1 | WP_230980984.1 |
| Bacillota | *Rossellomorea marisflavi* | WP_290032330.1 | WP_079516756.1 | WP_063191188.1 |
| Bacillota | *Rossellomorea oryzaecorticis* | WP_341983163.1 | WP_341983161.1 | WP_341982479.1 |
| Bacillota | *Rossellomorea sp. NPDC077527* | WP_397549110.1 | WP_397550578.1 | WP_397549808.1 |
| Bacillota | *Rossellomorea vietnamensis* | WP_406686849.1 | WP_426898446.1 | WP_426895544.1 |
| Bacillota | *Rubeoparvulum massiliense* | WP_048601877.1 | WP_315969544.1 | WP_048602591.1 |
| Bacillota | *Rummeliibacillus pycnus* | WP_102692493.1 | WP_397536986.1 | WP_102692757.1 |
| Bacillota | *Rummeliibacillus sp. TYF-LIM-RU47* | WP_150283994.1 | WP_150283419.1 | WP_150283850.1 |
| Bacillota | *Rummeliibacillus stabekisii* | WP_066786104.1 | WP_392995461.1 | WP_251645578.1 |
| Bacillota | *Rummeliibacillus suwonensis* | WP_146549179.1 | WP_146550224.1 | WP_146549454.1 |
| Bacillota | *Saccharibacillus alkalitolerans* | WP_166272347.1 | WP_166272345.1 | WP_166271645.1 |
| Bacillota | *Saccharibacillus deserti* | WP_172250470.1 | WP_172250468.1 | WP_172249784.1 |
| Bacillota | *Saccharibacillus endophyticus* | WP_172239876.1 | WP_172239879.1 | WP_172247082.1 |
| Bacillota | *Saccharibacillus kuerlensis* | WP_018976427.1 | WP_018976426.1 | WP_018975684.1 |
| Bacillota | *Saccharibacillus qingshengii* | WP_172196256.1 | WP_172196255.1 | WP_172194137.1 |
| Bacillota | *Saccharibacillus sacchari* | WP_340227389.1 | WP_037289210.1 | WP_037289904.1 |
| Bacillota | *Saccharibacillus sp. CPCC 101409* | WP_302726611.1 | WP_302726610.1 | WP_302726059.1 |
| Bacillota | *Saccharococcus caldoxylosilyticus* | WP_017435847.1 | WP_017435848.1 | WP_244380647.1 |
| Bacillota | *Saccharococcus sp. Marseille-Q5394* | WP_262173031.1 | WP_262174476.1 | WP_262172861.1 |
| Bacillota | *Saccharococcus thermophilus* | WP_166909718.1 | WP_208404355.1 | WP_166908162.1 |
| Bacillota | *Salibacterium aidingense* | WP_026700929.1 | WP_422787443.1 | WP_026700134.1 |
| Bacillota | *Salibacterium halotolerans* | WP_093335998.1 | WP_093335997.1 | WP_212634930.1 |
| Bacillota | *Salibacterium lacus* | WP_380711740.1 | WP_380711739.1 | WP_380712172.1 |
| Bacillota | *Salibacterium qingdaonense* | WP_090925891.1 | WP_090925890.1 | WP_177195403.1 |
| Bacillota | *Salibacterium salarium* | WP_125555844.1 | WP_348639443.1 | WP_306897393.1 |
| Bacillota | *Salicibibacter cibarius* | WP_200128665.1 | WP_200127415.1 | WP_200123865.1 |

| | | | | |
|---|---|---|---|---|
| Bacillota | *Salicibibacter cibi* | WP_200090038.1 | WP_200088136.1 | WP_200084986.1 |
| Bacillota | *Salicibibacter halophilus* | WP_142086661.1 | WP_142091167.1 | WP_142087858.1 |
| Bacillota | *Salicibibacter kimchii* | WP_114370407.1 | WP_114375494.1 | WP_227002793.1 |
| Bacillota | *Salimicrobium album* | WP_093106267.1 | WP_093107793.1 | WP_176765374.1 |
| Bacillota | *Salimicrobium flavidum* | WP_076556523.1 | WP_200806510.1 | WP_076556754.1 |
| Bacillota | *Salimicrobium halophilum* | WP_093191291.1 | WP_093193794.1 | WP_176757401.1 |
| Bacillota | *Salimicrobium humidisoli* | WP_095822251.1 | WP_095822692.1 | WP_158221305.1 |
| Bacillota | *Salimicrobium jeotgali* | WP_102335850.1 | WP_008588459.1 | WP_180962605.1 |
| Bacillota | *Salimicrobium sp. PL1-032A* | WP_347860490.1 | WP_347861340.1 | WP_347860669.1 |
| Bacillota | *Salinibacillus aidingensis* | WP_425542165.1 | WP_343842300.1 | WP_343840898.1 |
| Bacillota | *Salinibacillus kushneri* | WP_093136767.1 | WP_093131125.1 | WP_093136479.1 |
| Bacillota | *Salinibacillus xinjiangensis* | WP_153728306.1 | WP_323741785.1 | WP_153727443.1 |
| Bacillota | *Salinicoccus albus* | WP_020008327.1 | WP_020007583.1 | WP_020008736.1 |
| Bacillota | *Salinicoccus carnicancri* | WP_017548951.1 | WP_017549953.1 | WP_017548448.1 |
| Bacillota | *Salinicoccus halitifaciens* | WP_230820728.1 | WP_230822019.1 | WP_230820368.1 |
| Bacillota | *Salinicoccus halodurans* | WP_046790275.1 | WP_046791245.1 | WP_046789886.1 |
| Bacillota | *Salinicoccus hispanicus* | WP_160652528.1 | WP_160655972.1 | WP_160651099.1 |
| Bacillota | *Salinicoccus jeotgali* | WP_344702988.1 | WP_344704246.1 | WP_344701231.1 |
| Bacillota | *Salinicoccus kekensis* | WP_097038961.1 | WP_097041364.1 | WP_097038167.1 |
| Bacillota | *Salinicoccus roseus* | WP_271398009.1 | WP_222999666.1 | WP_040105713.1 |
| Bacillota | *Salinicoccus sediminis* | WP_046511680.1 | WP_046513205.1 | WP_046515947.1 |
| Bacillota | *Salinicoccus sesuvii* | WP_380653846.1 | WP_380651283.1 | WP_380655097.1 |
| Bacillota | *Salinicoccus siamensis* | WP_380569492.1 | WP_380570766.1 | WP_380571727.1 |
| Bacillota | *Salinicoccus sp. YB14-2* | WP_052256313.1 | WP_052254894.1 | WP_052256699.1 |
| Bacillota | *Salinithrix halophila* | WP_380705887.1 | WP_380705888.1 | WP_380703580.1 |
| Bacillota | *Salipaludibacillus agaradhaerens* | WP_257832383.1 | WP_078576843.1 | WP_078576125.1 |
| Bacillota | *Salipaludibacillus aurantiacus* | WP_093047103.1 | WP_093047104.1 | WP_093047693.1 |
| Bacillota | *Salipaludibacillus daqingensis* | WP_280768198.1 | WP_280768199.1 | WP_280769090.1 |

| | | | | |
|---|---|---|---|---|
| Bacillota | *Salipaludibacillus keqinensis* | WP_110608171.1 | WP_110608170.1 | WP_110609279.1 |
| Bacillota | *Salipaludibacillus neizhouensis* | WP_110935274.1 | WP_110935273.1 | WP_110936461.1 |
| Bacillota | *Salipaludibacillus sp. CUR1* | WP_230896188.1 | WP_230896189.1 | WP_230898551.1 |
| Bacillota | *Salirhabdus euzebyi* | WP_174495092.1 | WP_174496052.1 | WP_174497492.1 |
| Bacillota | *Salirhabdus salicampi* | WP_254496435.1 | WP_254497455.1 | WP_254496371.1 |
| Bacillota | *Salirhabdus sp. Marseille-P4669* | WP_102027217.1 | WP_102028662.1 | WP_102027290.1 |
| Bacillota | *Salisediminibacterium beveridgei* | WP_069364871.1 | WP_232318275.1 | WP_232318319.1 |
| Bacillota | *Salisediminibacterium halotolerans* | WP_121438815.1 | WP_121438816.1 | WP_121439723.1 |
| Bacillota | *Salisediminibacterium selenitireducens* | WP_013173031.1 | WP_013173032.1 | WP_177304823.1 |
| Bacillota | *Saliterribacillus persicus* | WP_114353814.1 | WP_114353214.1 | WP_114352801.1 |
| Bacillota | *Salsuginibacillus halophilus* | WP_106587852.1 | WP_106587851.1 | WP_106587505.1 |
| Bacillota | *Salsuginibacillus kocurii* | WP_018922278.1 | WP_018922277.1 | WP_018921352.1 |
| Bacillota | *Savagea faecisuis* | WP_381011375.1 | WP_381008702.1 | WP_381010748.1 |
| Bacillota | *Savagea serpentis* | WP_194561555.1 | WP_194562939.1 | WP_194561334.1 |
| Bacillota | *Schinkia azotoformans* | WP_003331124.1 | WP_035194851.1 | WP_328240042.1 |
| Bacillota | *Scopulibacillus cellulosilyticus* | WP_380965033.1 | WP_380965031.1 | WP_380964575.1 |
| Bacillota | *Scopulibacillus daqui* | WP_205003192.1 | WP_205003193.1 | WP_205002233.1 |
| Bacillota | *Scopulibacillus darangshiensis* | WP_132743194.1 | WP_207902904.1 | WP_132745980.1 |
| Bacillota | *Sediminibacillus albus* | WP_093210385.1 | WP_093213481.1 | WP_093210933.1 |
| Bacillota | *Sediminibacillus dalangtanensis* | WP_209369049.1 | WP_209365498.1 | WP_209368594.1 |
| Bacillota | *Sediminibacillus halophilus* | WP_074597154.1 | WP_074599060.1 | WP_074597461.1 |
| Bacillota | *Sediminibacillus massiliensis* | WP_077621753.1 | WP_077624813.1 | WP_077622117.1 |
| Bacillota | *Selenihalanaerobacter shriftii* | WP_078809290.1 | WP_078809291.1 | WP_078809292.1 |
| Bacillota | *Shimazuella alba* | WP_160799579.1 | WP_160799580.1 | WP_160802711.1 |
| Bacillota | *Shimazuella kribbensis* | WP_028775877.1 | WP_037463927.1 | WP_028775472.1 |
| Bacillota | *Shimazuella soli* | WP_240875942.1 | WP_240875941.1 | WP_240873807.1 |
| Bacillota | *Shouchella clausii* | WP_011246913.1 | WP_308899182.1 | WP_094427356.1 |
| Bacillota | *Shouchella hunanensis* | WP_274272691.1 | WP_274272692.1 | WP_274272785.1 |

| | | | | |
|---|---|---|---|---|
| Bacillota | *Shouchella lehensis* | WP_124742136.1 | WP_038480190.1 | WP_051667554.1 |
| Bacillota | *Shouchella lonarensis* | WP_090774967.1 | WP_090774966.1 | WP_090775969.1 |
| Bacillota | *Shouchella miscanthi* | WP_060705393.1 | WP_035398362.1 | WP_144559213.1 |
| Bacillota | *Shouchella patagoniensis* | WP_078391140.1 | WP_078391141.1 | WP_078391589.1 |
| Bacillota | *Shouchella shacheensis* | WP_059105136.1 | WP_245628145.1 | WP_059104709.1 |
| Bacillota | *Shouchella tritolerans* | WP_263706276.1 | WP_212953660.1 | WP_263707492.1 |
| Bacillota | *Shouchella xiaoxiensis* | WP_204467695.1 | WP_204467693.1 | WP_204464544.1 |
| Bacillota | *Siminovitchia acidinfaciens* | WP_126051034.1 | WP_126051035.1 | WP_126049851.1 |
| Bacillota | *Siminovitchia fordii* | WP_144516662.1 | WP_212962979.1 | WP_018705236.1 |
| Bacillota | *Siminovitchia fortis* | WP_144461110.1 | WP_144461111.1 | WP_120073885.1 |
| Bacillota | *Siminovitchia sediminis* | WP_380773864.1 | WP_380773863.1 | WP_380772334.1 |
| Bacillota | *Siminovitchia sp. 179-K 8D1 HS* | WP_413064294.1 | WP_413064293.1 | WP_413062858.1 |
| Bacillota | *Siminovitchia terrae* | WP_120117407.1 | WP_213020524.1 | WP_212948967.1 |
| Bacillota | *Sinobaca qinghaiensis* | WP_120193227.1 | WP_120193228.1 | WP_120192335.1 |
| Bacillota | *Solibacillus cecembensis* | WP_057982258.1 | WP_057986633.1 | WP_427763025.1 |
| Bacillota | *Solibacillus daqui* | WP_274309732.1 | WP_274309051.1 | WP_274309369.1 |
| Bacillota | *Solibacillus isronensis* | WP_079525889.1 | WP_413079058.1 | WP_079525008.1 |
| Bacillota | *Solibacillus kalamii* | WP_087618188.1 | WP_087615704.1 | WP_087615483.1 |
| Bacillota | *Solibacillus palustris* | WP_241368297.1 | WP_241367517.1 | WP_241367882.1 |
| Bacillota | *Solibacillus sp.* | WP_350479019.1 | WP_350478533.1 | WP_350477183.1 |
| Bacillota | *Sporosarcina aquimarina* | WP_317934445.1 | WP_317934225.1 | WP_221836181.1 |
| Bacillota | *Sporosarcina beigongshangi* | WP_212391108.1 | WP_322555715.1 | WP_203246136.1 |
| Bacillota | *Sporosarcina contaminans* | WP_381481929.1 | WP_381480857.1 | WP_381479742.1 |
| Bacillota | *Sporosarcina cyprini* | WP_238083723.1 | WP_238079673.1 | WP_238084261.1 |
| Bacillota | *Sporosarcina gallistercoris* | WP_191688084.1 | WP_191690044.1 | WP_191688376.1 |
| Bacillota | *Sporosarcina globispora* | WP_053434600.1 | WP_053434599.1 | WP_053436572.1 |
| Bacillota | *Sporosarcina highlanderae* | WP_301244972.1 | WP_301241610.1 | WP_301243091.1 |
| Bacillota | *Sporosarcina jeotgali* | WP_323693048.1 | WP_323692251.1 | WP_323692818.1 |

| | | | | |
|---|---|---|---|---|
| Bacillota | *Sporosarcina jiandibaonis* | WP_172369382.1 | WP_172371255.1 | WP_172369776.1 |
| Bacillota | *Sporosarcina koreensis* | WP_060208210.1 | WP_235588630.1 | WP_082713665.1 |
| Bacillota | *Sporosarcina limicola* | WP_192597920.1 | WP_192596793.1 | WP_192597375.1 |
| Bacillota | *Sporosarcina luteola* | WP_147059772.1 | WP_251697812.1 | WP_147054836.1 |
| Bacillota | *Sporosarcina obsidiansis* | WP_153732236.1 | WP_153730310.1 | WP_153731198.1 |
| Bacillota | *Sporosarcina oncorhynchi* | WP_317970799.1 | WP_317970857.1 | WP_317965138.1 |
| Bacillota | *Sporosarcina pasteurii* | WP_115360685.1 | WP_115359879.1 | WP_115360375.1 |
| Bacillota | *Sporosarcina quadrami* | WP_191695924.1 | WP_191695183.1 | WP_191693077.1 |
| Bacillota | *Sporosarcina siberiensis* | WP_381537257.1 | WP_381539562.1 | WP_381538696.1 |
| Bacillota | *Sporosarcina soli* | WP_381431378.1 | WP_381434861.1 | WP_381431890.1 |
| Bacillota | *Sporosarcina sp.* | WP_301107046.1 | WP_301107252.1 | WP_301108213.1 |
| Bacillota | *Sporosarcina thermotolerans* | WP_283731851.1 | WP_381426396.1 | WP_283731650.1 |
| Bacillota | *Sporosarcina trichiuri* | WP_290148243.1 | WP_290148363.1 | WP_290150366.1 |
| Bacillota | *Sporosarcina ureae* | WP_029053023.1 | WP_303969287.1 | WP_029052381.1 |
| Bacillota | *Sporosarcina ureilytica* | WP_075528193.1 | WP_075529057.1 | WP_075528509.1 |
| Bacillota | *Staphylococcus agnetis* | WP_105995156.1 | WP_186444660.1 | WP_037564834.1 |
| Bacillota | *Staphylococcus americanisciuri* | WP_259197863.1 | WP_259199938.1 | WP_259199379.1 |
| Bacillota | *Staphylococcus argenteus* | WP_000690034.1 | WP_088839925.1 | WP_000340487.1 |
| Bacillota | *Staphylococcus arlettae* | WP_313693173.1 | WP_107377050.1 | WP_002510904.1 |
| Bacillota | *Staphylococcus aureus* | WP_061651898.1 | WP_064139785.1 | WP_046400101.1 |
| Bacillota | *Staphylococcus auricularis* | WP_278925602.1 | WP_059107383.1 | WP_059106611.1 |
| Bacillota | *Staphylococcus borealis* | WP_325968792.1 | WP_320795828.1 | WP_053029182.1 |
| Bacillota | *Staphylococcus caeli* | WP_069996062.1 | WP_069994236.1 | WP_069995665.1 |
| Bacillota | *Staphylococcus caledonicus* | WP_423584440.1 | WP_423585406.1 | WP_198611262.1 |
| Bacillota | *Staphylococcus canis* | WP_198618143.1 | WP_198617867.1 | WP_198617317.1 |
| Bacillota | *Staphylococcus capitis* | WP_227938630.1 | WP_367121073.1 | WP_103156962.1 |
| Bacillota | *Staphylococcus caprae* | WP_206416781.1 | WP_206416920.1 | WP_002442407.1 |
| Bacillota | *Staphylococcus carnosus* | WP_103211976.1 | WP_012664257.1 | WP_015900042.1 |

| | | | | |
|---|---|---|---|---|
| Bacillota | *Staphylococcus casei* | WP_069824026.1 | WP_069823448.1 | WP_103267406.1 |
| Bacillota | *Staphylococcus chromogenes* | WP_037573738.1 | WP_105985726.1 | WP_145401049.1 |
| Bacillota | *Staphylococcus coagulans* | WP_182317341.1 | WP_050346061.1 | WP_241415290.1 |
| Bacillota | *Staphylococcus cohnii* | WP_119561916.1 | WP_107504835.1 | WP_103211509.1 |
| Bacillota | *Staphylococcus cornubiensis* | WP_086427769.1 | WP_086428521.1 | WP_086429092.1 |
| Bacillota | *Staphylococcus croceilyticus* | WP_103328611.1 | WP_103329602.1 | WP_103330022.1 |
| Bacillota | *Staphylococcus debuckii* | WP_123144728.1 | WP_123145576.1 | WP_123144472.1 |
| Bacillota | *Staphylococcus delphini* | WP_160215323.1 | WP_212574807.1 | WP_262612528.1 |
| Bacillota | *Staphylococcus devriesei* | WP_119526972.1 | WP_103166080.1 | WP_103167490.1 |
| Bacillota | *Staphylococcus durrellii* | WP_195720778.1 | WP_195721453.1 | WP_195721025.1 |
| Bacillota | *Staphylococcus edaphicus* | WP_099090706.1 | WP_099089595.1 | WP_099089336.1 |
| Bacillota | *Staphylococcus epidermidis* | WP_193390020.1 | WP_139689271.1 | WP_060556068.1 |
| Bacillota | *Staphylococcus equorum* | WP_277580753.1 | WP_197911036.1 | WP_021338687.1 |
| Bacillota | *Staphylococcus felis* | WP_262632082.1 | WP_349421082.1 | WP_103208719.1 |
| Bacillota | *Staphylococcus gallinarum* | WP_119485698.1 | WP_042738051.1 | WP_232154922.1 |
| Bacillota | *Staphylococcus haemolyticus* | WP_070111072.1 | WP_053026387.1 | WP_057504811.1 |
| Bacillota | *Staphylococcus hominis* | WP_172473662.1 | WP_311045257.1 | WP_119651854.1 |
| Bacillota | *Staphylococcus hyicus* | WP_039645720.1 | WP_167694023.1 | WP_039646206.1 |
| Bacillota | *Staphylococcus intermedius* | WP_019167612.1 | WP_019169043.1 | WP_019169118.1 |
| Bacillota | *Staphylococcus kloosii* | WP_195708453.1 | WP_103295076.1 | WP_313544265.1 |
| Bacillota | *Staphylococcus lloydii* | WP_316670929.1 | WP_048792414.1 | WP_195718300.1 |
| Bacillota | *Staphylococcus lugdunensis* | WP_002459091.1 | WP_002459906.1 | WP_037545920.1 |
| Bacillota | *Staphylococcus lutrae* | WP_085238316.1 | WP_085237590.1 | WP_085238060.1 |
| Bacillota | *Staphylococcus marylandisciuri* | WP_262854638.1 | WP_262856279.1 | WP_262855078.1 |
| Bacillota | *Staphylococcus massiliensis* | WP_009383771.1 | WP_009383870.1 | WP_009381584.1 |
| Bacillota | *Staphylococcus microti* | WP_044358858.1 | WP_044359276.1 | WP_044359784.1 |
| Bacillota | *Staphylococcus muscae* | WP_095116950.1 | WP_095115276.1 | WP_095116212.1 |
| Bacillota | *Staphylococcus nepalensis* | WP_349790011.1 | WP_103373391.1 | WP_096810346.1 |

| | | | | |
|---|---|---|---|---|
| Bacillota | *Staphylococcus pasteuri* | WP_326002342.1 | WP_259966835.1 | WP_126568352.1 |
| Bacillota | *Staphylococcus petrasii* | WP_242238694.1 | WP_103298240.1 | WP_103366839.1 |
| Bacillota | *Staphylococcus pettenkoferi* | WP_002472138.1 | WP_268216578.1 | WP_002472853.1 |
| Bacillota | *Staphylococcus pseudintermedius* | WP_390575169.1 | WP_199274268.1 | WP_063278835.1 |
| Bacillota | *Staphylococcus pseudoxylosus* | WP_214269407.1 | WP_214307630.1 | WP_326020120.1 |
| Bacillota | *Staphylococcus ratti* | WP_229291658.1 | WP_229292403.1 | WP_229293642.1 |
| Bacillota | *Staphylococcus rostri* | WP_103358010.1 | WP_103358500.1 | WP_103358994.1 |
| Bacillota | *Staphylococcus saccharolyticus* | WP_349546522.1 | WP_130559112.1 | WP_115313559.1 |
| Bacillota | *Staphylococcus saprophyticus* | WP_129319395.1 | WP_326023640.1 | WP_011303331.1 |
| Bacillota | *Staphylococcus schweitzeri* | WP_047552404.1 | WP_047427805.1 | WP_047549618.1 |
| Bacillota | *Staphylococcus shinii* | WP_101049392.1 | WP_318760341.1 | WP_107548731.1 |
| Bacillota | *Staphylococcus simiae* | WP_002463071.1 | WP_207517890.1 | WP_002465050.1 |
| Bacillota | *Staphylococcus simulans* | WP_107604644.1 | WP_412519330.1 | WP_119551259.1 |
| Bacillota | *Staphylococcus sp.* | WP_287173972.1 | WP_293735409.1 | WP_287236828.1 |
| Bacillota | *Staphylococcus warneri* | WP_270232372.1 | WP_107546604.1 | WP_058709479.1 |
| Bacillota | *Staphylococcus xylosus* | WP_119614750.1 | WP_101104024.1 | WP_042362890.1 |
| Bacillota | *Staphylospora marina* | WP_124727366.1 | WP_124727365.1 | WP_124727719.1 |
| Bacillota | *Sulfoacidibacillus ferrooxidans* | WP_241715794.1 | WP_241715796.1 | WP_241711416.1 |
| Bacillota | *Sulfoacidibacillus thermotolerans* | WP_109431150.1 | WP_109431149.1 | WP_109430457.1 |
| Bacillota | *Sutcliffiella cohnii* | WP_066415539.1 | WP_412766311.1 | WP_328225366.1 |
| Bacillota | *Sutcliffiella deserti* | WP_223702018.1 | WP_223702019.1 | WP_223700326.1 |
| Bacillota | *Sutcliffiella halmapala* | WP_078380210.1 | WP_078380209.1 | WP_169865011.1 |
| Bacillota | *Sutcliffiella horikoshii* | WP_426910852.1 | WP_251429387.1 | WP_237661957.1 |
| Bacillota | *Sutcliffiella rhizosphaerae* | WP_230503349.1 | WP_230503351.1 | WP_230499742.1 |
| Bacillota | *Sutcliffiella sp. NPDC057660* | WP_386638742.1 | WP_386638745.1 | WP_386644139.1 |
| Bacillota | *Symbiobacterium terraclitae* | WP_209467923.1 | WP_209467487.1 | WP_209467439.1 |
| Bacillota | *Symbiobacterium thermophilum* | WP_011195753.1 | WP_273380602.1 | WP_043714106.1 |
| Bacillota | *Tenuibacillus multivorans* | WP_093857175.1 | WP_093857303.1 | WP_093855768.1 |

| | | | | |
|---|---|---|---|---|
| Bacillota | *Terribacillus aidingensis* | WP_366249817.1 | WP_097039613.1 | WP_097040255.1 |
| Bacillota | *Terribacillus halophilus* | WP_093728367.1 | WP_093725557.1 | WP_411230823.1 |
| Bacillota | *Terribacillus saccharophilus* | WP_411213887.1 | WP_411216852.1 | WP_411218624.1 |
| Bacillota | *Terribacillus sp. 179-K 1B1 HS* | WP_413088889.1 | WP_413088162.1 | WP_413091881.1 |
| Bacillota | *Terrihalobacillus insolitus* | WP_272436575.1 | WP_272434696.1 | WP_272435445.1 |
| Bacillota | *Terrilactibacillus laevilacticus* | WP_141603530.1 | WP_141189502.1 | WP_141189187.1 |
| Bacillota | *Terrilactibacillus tamarindi* | WP_155220068.1 | WP_155220070.1 | WP_155218608.1 |
| Bacillota | *Tetzosporium hominis* | WP_243370631.1 | WP_094943548.1 | WP_094941578.1 |
| Bacillota | *Texcoconibacillus texcoconensis* | WP_184664026.1 | WP_184664025.1 | WP_184662655.1 |
| Bacillota | *Thalassobacillus cyri* | WP_093044700.1 | WP_176791319.1 | WP_093046616.1 |
| Bacillota | *Thalassobacillus devorans* | WP_085508675.1 | WP_037986477.1 | WP_085508061.1 |
| Bacillota | *Thalassobacillus hwangdonensis* | WP_386056987.1 | WP_386062940.1 | WP_386056301.1 |
| Bacillota | *Thalassobacillus pellis* | WP_205091761.1 | WP_205092936.1 | WP_205092104.1 |
| Bacillota | *Thalassobacillus sp. FIB228* | WP_394460240.1 | WP_394460934.1 | WP_394459867.1 |
| Bacillota | *Thermaerobacillus caldiproteolyticus* | WP_181555922.1 | WP_194520666.1 | WP_181555025.1 |
| Bacillota | *Thermaerobacter composti* | WP_318751533.1 | WP_318751084.1 | WP_318751445.1 |
| Bacillota | *Thermaerobacter marianensis* | WP_013495781.1 | WP_013495780.1 | WP_013495475.1 |
| Bacillota | *Thermaerobacter sp. FW80* | WP_135225485.1 | WP_135225000.1 | WP_135225372.1 |
| Bacillota | *Thermaerobacter subterraneus* | WP_006903784.1 | WP_006902983.1 | WP_423219092.1 |
| Bacillota | *Thermicanus aegyptius* | WP_028986721.1 | WP_005582017.1 | WP_156920558.1 |
| Bacillota | *Thermincola ferriacetica* | WP_052218853.1 | WP_052218852.1 | WP_052218485.1 |
| Bacillota | *Thermincola potens* | WP_013121178.1 | WP_013121179.1 | WP_013119642.1 |
| Bacillota | *Thermoactinomyces daqus* | WP_033101403.1 | WP_033101447.1 | WP_052154059.1 |
| Bacillota | *Thermoactinomyces mirandus* | WP_181740253.1 | WP_181740251.1 | WP_181737370.1 |
| Bacillota | *Thermoactinomyces sp. DSM 45891* | WP_072329871.1 | WP_072329873.1 | WP_072331764.1 |
| Bacillota | *Thermobacillus xylanilyticus* | WP_213484588.1 | WP_213484587.1 | WP_213484019.1 |
| Bacillota | *Thermoflavimicrobium daqui* | WP_113660335.1 | WP_113660334.1 | WP_113658619.1 |
| Bacillota | *Thermoflavimicrobium dichotomicum* | WP_093230852.1 | WP_093230851.1 | WP_175482318.1 |

| | | | | |
|---|---|---|---|---|
| Bacillota | *Thermolongibacillus altinsuensis* | WP_132947109.1 | WP_132947108.1 | WP_132948044.1 |
| Bacillota | *Tuberibacillus calidus* | WP_027723795.1 | WP_027725603.1 | WP_027726143.1 |
| Bacillota | *Tuberibacillus sp. Marseille-P3662* | WP_085523572.1 | WP_085523628.1 | WP_085524079.1 |
| Bacillota | *Ureibacillus acetophenoni* | WP_097148931.1 | WP_097147578.1 | WP_097147917.1 |
| Bacillota | *Ureibacillus aquaedulcis* | WP_301138178.1 | WP_301136159.1 | WP_301136504.1 |
| Bacillota | *Ureibacillus chungkukjangi* | WP_107934324.1 | WP_107936772.1 | WP_251397087.1 |
| Bacillota | *Ureibacillus galli* | WP_191705773.1 | WP_191706082.1 | WP_191706386.1 |
| Bacillota | *Ureibacillus manganicus* | WP_036183726.1 | WP_036182408.1 | WP_036181806.1 |
| Bacillota | *Ureibacillus massiliensis* | WP_036176731.1 | WP_036171102.1 | WP_036171897.1 |
| Bacillota | *Ureibacillus sinduriensis* | WP_036203690.1 | WP_036200036.1 | WP_036202732.1 |
| Bacillota | *Ureibacillus sp. GCM10028918* | WP_390284955.1 | WP_390286420.1 | WP_390285722.1 |
| Bacillota | *Ureibacillus thermophilus* | WP_208649562.1 | WP_208651565.1 | WP_208651930.1 |
| Bacillota | *Ureibacillus thermosphaericus* | WP_168411771.1 | WP_340436786.1 | WP_168411946.1 |
| Bacillota | *Ureibacillus xyleni* | WP_097074454.1 | WP_097071621.1 | WP_097072042.1 |
| Bacillota | *Virgibacillus alimentarius* | WP_029267903.1 | WP_226370754.1 | WP_226370470.1 |
| Bacillota | *Virgibacillus byunsanensis* | WP_390361856.1 | WP_390362218.1 | WP_390362069.1 |
| Bacillota | *Virgibacillus dokdonensis* | WP_116277069.1 | WP_077706929.1 | WP_077704362.1 |
| Bacillota | *Virgibacillus doumboii* | WP_164668779.1 | WP_164668118.1 | WP_164669103.1 |
| Bacillota | *Virgibacillus halodenitrificans* | WP_121614998.1 | WP_019377733.1 | WP_283875060.1 |
| Bacillota | *Virgibacillus halophilus* | WP_390353554.1 | WP_390357844.1 | WP_390352980.1 |
| Bacillota | *Virgibacillus halotolerans* | WP_205135334.1 | WP_205137470.1 | WP_205135629.1 |
| Bacillota | *Virgibacillus ihumii* | WP_174613585.1 | WP_174614267.1 | WP_174613153.1 |
| Bacillota | *Virgibacillus indicus* | WP_094883727.1 | WP_094885865.1 | WP_094884008.1 |
| Bacillota | *Virgibacillus kekensis* | WP_390293313.1 | WP_390293469.1 | WP_390292948.1 |
| Bacillota | *Virgibacillus litoralis* | WP_209481076.1 | WP_209479546.1 | WP_209482083.1 |
| Bacillota | *Virgibacillus natechei* | WP_209462782.1 | WP_209462604.1 | WP_209461433.1 |
| Bacillota | *Virgibacillus ndiopensis* | WP_099158340.1 | WP_099159900.1 | WP_099157985.1 |
| Bacillota | *Virgibacillus necropolis* | WP_404452830.1 | WP_404454248.1 | WP_089532707.1 |

| | | | | |
|---|---|---|---|---|
| Bacillota | *Virgibacillus oceani* | WP_188453881.1 | WP_188454115.1 | WP_188454492.1 |
| Bacillota | *Virgibacillus pantothenticus* | WP_077303373.1 | WP_077301211.1 | WP_077300414.1 |
| Bacillota | *Virgibacillus phasianinus* | WP_089062613.1 | WP_089061994.1 | WP_089062850.1 |
| Bacillota | *Virgibacillus profundi* | WP_095655681.1 | WP_095657386.1 | WP_095655102.1 |
| Bacillota | *Virgibacillus proomii* | WP_077320600.1 | WP_077320023.1 | WP_077320327.1 |
| Bacillota | *Virgibacillus sediminis* | WP_390305164.1 | WP_390302484.1 | WP_390308289.1 |
| Bacillota | *Virgibacillus senegalensis* | WP_053219421.1 | WP_053218627.1 | WP_053219771.1 |
| Bacillota | *Virgibacillus siamensis* | WP_077326649.1 | WP_077328044.1 | WP_077325994.1 |
| Bacillota | *Virgibacillus soli* | WP_390185828.1 | WP_320379603.1 | WP_320378859.1 |
| Bacillota | *Virgibacillus sp. MSJ-26* | WP_216487690.1 | WP_216489624.1 | WP_216489590.1 |
| Bacillota | *Virgibacillus tibetensis* | WP_327606045.1 | WP_327608938.1 | WP_327606340.1 |
| Bacillota | *Virgibacillus xinjiangensis* | WP_390267290.1 | WP_390273709.1 | WP_390271245.1 |
| Bacillota | *Viridibacillus arvi* | WP_390008744.1 | WP_053417950.1 | WP_053417543.1 |
| Bacillota | *Viridibacillus soli* | WP_200750120.1 | WP_200749327.1 | WP_200748351.1 |
| Bacillota | *Weizmannia acidilactici* | WP_151680746.1 | WP_151697946.1 | WP_151681373.1 |
| Bacillota | *Xylanibacillus composti* | WP_213410179.1 | WP_213410178.1 | WP_213410594.1 |
| Bacillota | *Zhaonella formicivorans* | WP_227766183.1 | WP_227767134.1 | WP_227767136.1 |
| Bacteroidota | *Abyssalbus ytuae* | WP_255842322.1 | WP_255842580.1 | WP_255843861.1 |
| Bacteroidota | *Acidiluteibacter ferrifornacis* | WP_160634009.1 | WP_160633238.1 | WP_160632436.1 |
| Bacteroidota | *Adhaeribacter aerolatus* | WP_146903393.1 | WP_146896245.1 | WP_146900283.1 |
| Bacteroidota | *Adhaeribacter aquaticus* | WP_026462874.1 | WP_026464488.1 | WP_026461962.1 |
| Bacteroidota | *Adhaeribacter arboris* | WP_106925362.1 | WP_106927209.1 | WP_106933576.1 |
| Bacteroidota | *Adhaeribacter pallidiroseus* | WP_115373291.1 | WP_115373435.1 | WP_115375762.1 |
| Bacteroidota | *Adhaeribacter radiodurans* | WP_182414787.1 | WP_182415598.1 | WP_182414468.1 |
| Bacteroidota | *Adhaeribacter rhizoryzae* | WP_150087427.1 | WP_150087644.1 | WP_150086256.1 |
| Bacteroidota | *Adhaeribacter soli* | WP_150904862.1 | WP_150905911.1 | WP_150903363.1 |
| Bacteroidota | *Adhaeribacter swui* | WP_185270322.1 | WP_185274619.1 | WP_185272839.1 |
| Bacteroidota | *Adhaeribacter terreus* | WP_378015753.1 | WP_378016465.1 | WP_378017653.1 |

| | | | | |
|---|---|---|---|---|
| Bacteroidota | *Adhaeribacter terrigena* | WP_200504364.1 | WP_200505621.1 | WP_200506036.1 |
| Bacteroidota | *Aegicerativicinus sediminis* | WP_228851488.1 | WP_228851243.1 | WP_228851022.1 |
| Bacteroidota | *Aequorivita antarctica* | WP_111843834.1 | WP_111845043.1 | WP_111844811.1 |
| Bacteroidota | *Aequorivita aquimaris* | WP_062621430.1 | WP_062619548.1 | WP_062619660.1 |
| Bacteroidota | *Aequorivita aurantiaca* | WP_290253605.1 | WP_290255165.1 | WP_290255086.1 |
| Bacteroidota | *Aequorivita capsosiphonis* | WP_026449899.1 | WP_026451927.1 | WP_026450721.1 |
| Bacteroidota | *Aequorivita ciconiae* | WP_128249060.1 | WP_128249027.1 | WP_128251637.1 |
| Bacteroidota | *Aequorivita echinoideorum* | WP_214113607.1 | WP_214113075.1 | WP_214113159.1 |
| Bacteroidota | *Aequorivita lipolytica* | WP_111815969.1 | WP_111816248.1 | WP_111816891.1 |
| Bacteroidota | *Aequorivita marina* | WP_310991922.1 | WP_310992157.1 | WP_310993816.1 |
| Bacteroidota | *Aequorivita nionensis* | WP_410005228.1 | WP_410004785.1 | WP_410007104.1 |
| Bacteroidota | *Aequorivita sinensis* | WP_271424522.1 | WP_271393576.1 | WP_271393529.1 |
| Bacteroidota | *Aequorivita soesokkakensis* | WP_068760878.1 | WP_068761363.1 | WP_068760990.1 |
| Bacteroidota | *Aequorivita sp. Q41* | WP_347374707.1 | WP_347372786.1 | WP_347374868.1 |
| Bacteroidota | *Aequorivita sublithincola* | WP_014783441.1 | WP_014781031.1 | WP_014780829.1 |
| Bacteroidota | *Aequorivita todarodis* | WP_272854907.1 | WP_272856410.1 | WP_272856446.1 |
| Bacteroidota | *Aequorivita viscosa* | WP_073219450.1 | WP_073221462.1 | WP_073214181.1 |
| Bacteroidota | *Aequorivita vitellina* | WP_237602884.1 | WP_237604082.1 | WP_237604065.1 |
| Bacteroidota | *Aequorivita vladivostokensis* | WP_045080904.1 | WP_045080108.1 | WP_045081657.1 |
| Bacteroidota | *Aequorivita xiaoshiensis* | WP_237607055.1 | WP_237608260.1 | WP_237608850.1 |
| Bacteroidota | *Aestuariibaculum lutulentum* | WP_240572916.1 | WP_240574573.1 | WP_240573340.1 |
| Bacteroidota | *Aestuariibaculum marinum* | WP_188223804.1 | WP_188221776.1 | WP_188222367.1 |
| Bacteroidota | *Aestuariibaculum sediminum* | WP_188229943.1 | WP_188229495.1 | WP_188230458.1 |
| Bacteroidota | *Aestuariibaculum sp. YM273* | WP_307881004.1 | WP_307881203.1 | WP_307881758.1 |
| Bacteroidota | *Aestuariibaculum suncheonense* | WP_188215830.1 | WP_188215657.1 | WP_188216389.1 |
| Bacteroidota | *Aestuariivivens insulae* | WP_242204282.1 | WP_242206092.1 | WP_242205612.1 |
| Bacteroidota | *Aestuariivivens marinum* | WP_242132290.1 | WP_242135394.1 | WP_242135503.1 |
| Bacteroidota | *Aestuariivivens sediminicola* | WP_242120336.1 | WP_242121371.1 | WP_242119724.1 |

| | | | | |
|---|---|---|---|---|
| Bacteroidota | *Aestuariivivens sediminis* | WP_242086932.1 | WP_242158599.1 | WP_242085353.1 |
| Bacteroidota | *Aestuariivivens sp. NBU2969* | WP_223548296.1 | WP_223551508.1 | WP_223552062.1 |
| Bacteroidota | *Agaribacillus aureus* | WP_346757862.1 | WP_346755833.1 | WP_346761836.1 |
| Bacteroidota | *Aggregatimonas sangjinii* | WP_138852003.1 | WP_138852101.1 | WP_138851661.1 |
| Bacteroidota | *Agriterribacter humi* | WP_152268168.1 | WP_152270501.1 | WP_152266726.1 |
| Bacteroidota | *Agriterribacter sp.* | WP_325404972.1 | WP_325403507.1 | WP_324556562.1 |
| Bacteroidota | *Albibacterium bauzanense* | WP_132222972.1 | WP_132221473.1 | WP_132222344.1 |
| Bacteroidota | *Albibacterium sp.* | WP_325357452.1 | WP_325358942.1 | WP_325359824.1 |
| Bacteroidota | *Algibacter agarivorans* | WP_345189876.1 | WP_345193043.1 | WP_345193815.1 |
| Bacteroidota | *Algibacter amylolyticus* | WP_144118081.1 | WP_144117049.1 | WP_144116980.1 |
| Bacteroidota | *Algibacter aquimarinus* | WP_345169810.1 | WP_345170338.1 | WP_345169783.1 |
| Bacteroidota | *Algibacter lectus* | WP_074936123.1 | WP_304407024.1 | WP_282148460.1 |
| Bacteroidota | *Algibacter luteus* | WP_019386403.1 | WP_289194526.1 | WP_019388342.1 |
| Bacteroidota | *Algibacter marinivivus* | WP_109353276.1 | WP_109353074.1 | WP_109353815.1 |
| Bacteroidota | *Algibacter mikhailovii* | WP_189358466.1 | WP_189362427.1 | WP_189362358.1 |
| Bacteroidota | *Algibacter miyuki* | WP_290273454.1 | WP_290271960.1 | WP_290273464.1 |
| Bacteroidota | *Algibacter pacificus* | WP_147678726.1 | WP_147678533.1 | WP_147678795.1 |
| Bacteroidota | *Algibacter pectinivorans* | WP_092853154.1 | WP_092852392.1 | WP_092851984.1 |
| Bacteroidota | *Algibacter sp.* | WP_424201046.1 | WP_348451855.1 | WP_348451771.1 |
| Bacteroidota | *Algivirga pacifica* | WP_345373764.1 | WP_345373213.1 | WP_345371319.1 |
| Bacteroidota | *Algoriella sp.* | WP_332544110.1 | WP_332546255.1 | WP_332543467.1 |
| Bacteroidota | *Algoriphagus aestuariicola* | WP_206570757.1 | WP_206568757.1 | WP_206569828.1 |
| Bacteroidota | *Algoriphagus alkaliphilus* | WP_092730293.1 | WP_092732049.1 | WP_092729109.1 |
| Bacteroidota | *Algoriphagus antarcticus* | WP_086543870.1 | WP_086542156.1 | WP_086541399.1 |
| Bacteroidota | *Algoriphagus aquaeductus* | WP_409682485.1 | WP_406949959.1 | WP_406949864.1 |
| Bacteroidota | *Algoriphagus aquatilis* | WP_377917023.1 | WP_377912258.1 | WP_377917659.1 |
| Bacteroidota | *Algoriphagus aquimarinus* | WP_092894558.1 | WP_146919054.1 | WP_092894759.1 |
| Bacteroidota | *Algoriphagus aquimaris* | WP_133390194.1 | WP_133391627.1 | WP_240686591.1 |

| | | | | |
|---|---|---|---|---|
| Bacteroidota | *Algoriphagus boritolerans* | WP_332911655.1 | WP_103925912.1 | WP_103924010.1 |
| Bacteroidota | *Algoriphagus boseongensis* | WP_133555455.1 | WP_133552919.1 | WP_133555600.1 |
| Bacteroidota | *Algoriphagus chordae* | WP_111316191.1 | WP_111323425.1 | WP_111318264.1 |
| Bacteroidota | *Algoriphagus confluentis* | WP_338223545.1 | WP_338223409.1 | WP_338223464.1 |
| Bacteroidota | *Algoriphagus faecimaris* | WP_087938083.1 | WP_087938226.1 | WP_087940368.1 |
| Bacteroidota | *Algoriphagus formosus* | WP_100628282.1 | WP_100627774.1 | WP_232724787.1 |
| Bacteroidota | *Algoriphagus halophilus* | WP_074226768.1 | WP_074224025.1 | WP_373400315.1 |
| Bacteroidota | *Algoriphagus hitonicola* | WP_092791543.1 | WP_092789748.1 | WP_092791970.1 |
| Bacteroidota | *Algoriphagus iocasae* | WP_184494711.1 | WP_184497394.1 | WP_184495802.1 |
| Bacteroidota | *Algoriphagus jejuensis* | WP_343852720.1 | WP_343853072.1 | WP_343850080.1 |
| Bacteroidota | *Algoriphagus kandeliae* | WP_135075615.1 | WP_135070713.1 | WP_135072807.1 |
| Bacteroidota | *Algoriphagus lacus* | WP_119477098.1 | WP_119477573.1 | WP_119477184.1 |
| Bacteroidota | *Algoriphagus limi* | WP_259415152.1 | WP_259413116.1 | WP_259414277.1 |
| Bacteroidota | *Algoriphagus litoralis* | WP_111670835.1 | WP_111669801.1 | WP_111671671.1 |
| Bacteroidota | *Algoriphagus locisalis* | WP_091692921.1 | WP_091691830.1 | WP_091695981.1 |
| Bacteroidota | *Algoriphagus lutimaris* | WP_205646464.1 | WP_205644250.1 | WP_205645590.1 |
| Bacteroidota | *Algoriphagus machipongonensis* | WP_008203198.1 | WP_008200935.1 | WP_008198563.1 |
| Bacteroidota | *Algoriphagus mannitolivorans* | WP_026951984.1 | WP_026951705.1 | WP_026951897.1 |
| Bacteroidota | *Algoriphagus marinus* | WP_075350005.1 | WP_075352665.1 | WP_075350307.1 |
| Bacteroidota | *Algoriphagus namhaensis* | WP_377903089.1 | WP_377906576.1 | WP_377903203.1 |
| Bacteroidota | *Algoriphagus oliviformis* | WP_206576919.1 | WP_206577607.1 | WP_206579989.1 |
| Bacteroidota | *Algoriphagus ornithinivorans* | WP_091652719.1 | WP_091650703.1 | WP_091651060.1 |
| Bacteroidota | *Algoriphagus pacificus* | WP_206584645.1 | WP_206587470.1 | WP_206585619.1 |
| Bacteroidota | *Algoriphagus ratkowskyi* | WP_086500922.1 | WP_086501427.1 | WP_086501982.1 |
| Bacteroidota | *Algoriphagus resistens* | WP_057936911.1 | WP_057936316.1 | WP_057939717.1 |
| Bacteroidota | *Algoriphagus sanaruensis* | WP_067550178.1 | WP_067550458.1 | WP_067544506.1 |
| Bacteroidota | *Algoriphagus sediminis* | WP_290000812.1 | WP_289999935.1 | WP_290000871.1 |
| Bacteroidota | *Algoriphagus sp.* | WP_348275023.1 | WP_300316368.1 | WP_348277066.1 |

| | | | | |
|---|---|---|---|---|
| Bacteroidota | *Algoriphagus taiwanensis* | WP_338230563.1 | WP_338227733.1 | WP_338229737.1 |
| Bacteroidota | *Algoriphagus terrigena* | WP_026968918.1 | WP_026967199.1 | WP_026967940.1 |
| Bacteroidota | *Algoriphagus vanfongensis* | WP_026956411.1 | WP_026954082.1 | WP_026955295.1 |
| Bacteroidota | *Algoriphagus winogradskyi* | WP_283412018.1 | WP_283413402.1 | WP_283414707.1 |
| Bacteroidota | *Algoriphagus yeomjeoni* | WP_111609732.1 | WP_425639078.1 | WP_425636259.1 |
| Bacteroidota | *Algoriphagus zhangzhouensis* | WP_073572735.1 | WP_073570160.1 | WP_073571012.1 |
| Bacteroidota | *Allomuricauda sp. SCSIO 65647* | WP_235265680.1 | WP_235265793.1 | WP_235266678.1 |
| Bacteroidota | *Allotamlana fucoidanivorans* | WP_139694982.1 | WP_139698433.1 | WP_139696462.1 |
| Bacteroidota | *Altibacter lentus* | WP_034258177.1 | WP_034257797.1 | WP_034259005.1 |
| Bacteroidota | *Altibacter sp.* | WP_290787002.1 | WP_290785786.1 | WP_290785445.1 |
| Bacteroidota | *Anditalea andensis* | WP_035075984.1 | WP_035072741.1 | WP_035075153.1 |
| Bacteroidota | *Anseongella ginsenosidimutans* | WP_132129421.1 | WP_132128381.1 | WP_132129763.1 |
| Bacteroidota | *Antarcticibacterium sp. 1MA-6-2* | WP_235328210.1 | WP_235329999.1 | WP_235330129.1 |
| Bacteroidota | *Apibacter adventoris* | WP_105193923.1 | WP_181042806.1 | WP_105245879.1 |
| Bacteroidota | *Apibacter mensalis* | WP_303849861.1 | WP_303839970.1 | WP_303822039.1 |
| Bacteroidota | *Apibacter muscae* | WP_146292415.1 | WP_146292170.1 | WP_146311829.1 |
| Bacteroidota | *Apibacter sp.* | WP_293917054.1 | WP_293918312.1 | WP_293918330.1 |
| Bacteroidota | *Aquaticitalea lipolytica* | WP_283636607.1 | WP_188604491.1 | WP_188606921.1 |
| Bacteroidota | *Aquiflexum balticum* | WP_084118598.1 | WP_084121789.1 | WP_084118744.1 |
| Bacteroidota | *Aquiflexum gelatinilyticum* | WP_258422916.1 | WP_258424477.1 | WP_259263102.1 |
| Bacteroidota | *Aquiflexum lacus* | WP_194972829.1 | WP_194976628.1 | WP_194975748.1 |
| Bacteroidota | *Aquiflexum sp.* | WP_373522937.1 | WP_373523815.1 | WP_373521777.1 |
| Bacteroidota | *Aquimarina acroporae* | WP_248325971.1 | WP_248327184.1 | WP_248329473.1 |
| Bacteroidota | *Aquimarina addita* | WP_344928121.1 | WP_344926945.1 | WP_344927438.1 |
| Bacteroidota | *Aquimarina agarilytica* | WP_010182235.1 | WP_010178350.1 | WP_010182594.1 |
| Bacteroidota | *Aquimarina agarivorans* | WP_010521832.1 | WP_010522891.1 | WP_010520711.1 |
| Bacteroidota | *Aquimarina aggregata* | WP_281986244.1 | WP_066317910.1 | WP_281990476.1 |
| Bacteroidota | *Aquimarina algicola* | WP_140589169.1 | WP_140595569.1 | WP_140595083.1 |

| | | | | |
|---|---|---|---|---|
| Bacteroidota | *Aquimarina algiphila* | WP_271767707.1 | WP_271766777.1 | WP_282086395.1 |
| Bacteroidota | *Aquimarina amphilecti* | WP_091404647.1 | WP_091403964.1 | WP_091410641.1 |
| Bacteroidota | *Aquimarina aquimarini* | WP_108869272.1 | WP_108868242.1 | WP_108868066.1 |
| Bacteroidota | *Aquimarina atlantica* | WP_034238049.1 | WP_034241498.1 | WP_034246824.1 |
| Bacteroidota | *Aquimarina brevivitae* | WP_130286860.1 | WP_130287960.1 | WP_130286092.1 |
| Bacteroidota | *Aquimarina celericrescens* | WP_378318285.1 | WP_378318615.1 | WP_378321627.1 |
| Bacteroidota | *Aquimarina gracilis* | WP_324180621.1 | WP_324180101.1 | WP_324178525.1 |
| Bacteroidota | *Aquimarina intermedia* | WP_148782085.1 | WP_148782431.1 | WP_148783651.1 |
| Bacteroidota | *Aquimarina litoralis* | WP_219008440.1 | WP_219009007.1 | WP_219010050.1 |
| Bacteroidota | *Aquimarina longa* | WP_062057664.1 | WP_062058925.1 | WP_062059664.1 |
| Bacteroidota | *Aquimarina macrocephali* | WP_024770031.1 | WP_024771131.1 | WP_024771276.1 |
| Bacteroidota | *Aquimarina muelleri* | WP_027411125.1 | WP_027411711.1 | WP_027411300.1 |
| Bacteroidota | *Aquimarina mytili* | WP_201917632.1 | WP_201924142.1 | WP_201922323.1 |
| Bacteroidota | *Aquimarina pacifica* | WP_025742060.1 | WP_025743084.1 | WP_025742517.1 |
| Bacteroidota | *Aquimarina rubra* | WP_378290376.1 | WP_378288455.1 | WP_378295432.1 |
| Bacteroidota | *Aquimarina sediminis* | WP_103068351.1 | WP_103070573.1 | WP_103070728.1 |
| Bacteroidota | *Aquimarina sp. I32.4* | WP_103863893.1 | WP_103864881.1 | WP_103866837.1 |
| Bacteroidota | *Aquimarina spongiae* | WP_073314409.1 | WP_073314355.1 | WP_073320496.1 |
| Bacteroidota | *Aquirufa antheringensis* | WP_130922775.1 | WP_130923145.1 | WP_269012307.1 |
| Bacteroidota | *Aquirufa aurantiipilula* | WP_276344163.1 | WP_223142429.1 | WP_223143335.1 |
| Bacteroidota | *Aquirufa avitistagni* | WP_377983723.1 | WP_377982913.1 | WP_377983456.1 |
| Bacteroidota | *Aquirufa beregesia* | WP_166230096.1 | WP_166228982.1 | WP_166229514.1 |
| Bacteroidota | *Aquirufa ecclesiirivi* | WP_275612994.1 | WP_166375689.1 | WP_275612331.1 |
| Bacteroidota | *Aquirufa echingensis* | WP_377976744.1 | WP_377976516.1 | WP_377974603.1 |
| Bacteroidota | *Aquirufa lenticrescens* | WP_223131088.1 | WP_223130902.1 | WP_223129962.1 |
| Bacteroidota | *Aquirufa nivalisilvae* | WP_269013300.1 | WP_269024951.1 | WP_109324451.1 |
| Bacteroidota | *Aquirufa novilacunae* | WP_406800710.1 | WP_406776800.1 | WP_406800572.1 |
| Bacteroidota | *Aquirufa originis* | WP_377979066.1 | WP_377978324.1 | WP_377978916.1 |

| | | | | |
|---|---|---|---|---|
| Bacteroidota | *Aquirufa regiilacus* | WP_315576891.1 | WP_315575113.1 | WP_315574784.1 |
| Bacteroidota | *Aquirufa rosea* | WP_129026517.1 | WP_129027631.1 | WP_129027468.1 |
| Bacteroidota | *Aquirufa salirivi* | WP_406749496.1 | WP_406751867.1 | WP_406750838.1 |
| Bacteroidota | *Aquirufa sp.* | WP_395767743.1 | WP_395785771.1 | WP_395785934.1 |
| Bacteroidota | *Arachidicoccus ginsenosidivorans* | WP_146779924.1 | WP_146780403.1 | WP_158639367.1 |
| Bacteroidota | *Arachidicoccus rhizosphaerae* | WP_091397094.1 | WP_091395310.1 | WP_091399636.1 |
| Bacteroidota | *Arachidicoccus soli* | WP_119987445.1 | WP_119989760.1 | WP_119984939.1 |
| Bacteroidota | *Arachidicoccus sp. BS20* | WP_066202682.1 | WP_066206247.1 | WP_066208570.1 |
| Bacteroidota | *Arachidicoccus terrestris* | WP_224070549.1 | WP_224071115.1 | WP_224070363.1 |
| Bacteroidota | *Arcicella aquatica* | WP_323246239.1 | WP_323249301.1 | WP_323247214.1 |
| Bacteroidota | *Arcicella aurantiaca* | WP_109743205.1 | WP_109741526.1 | WP_109744417.1 |
| Bacteroidota | *Arcicella gelida* | WP_323328161.1 | WP_323697626.1 | WP_323697301.1 |
| Bacteroidota | *Arcicella rigui* | WP_323297640.1 | WP_323296787.1 | WP_323294779.1 |
| Bacteroidota | *Arcicella rosea* | WP_184128752.1 | WP_184134267.1 | WP_367285748.1 |
| Bacteroidota | *Arcicella sp. LKC2W* | WP_323347125.1 | WP_323345881.1 | WP_323348259.1 |
| Bacteroidota | *Arcticibacter eurypsychrophilus* | WP_069660197.1 | WP_069658220.1 | WP_069658143.1 |
| Bacteroidota | *Arcticibacter pallidicorallinus* | WP_106295235.1 | WP_106293321.1 | WP_106293700.1 |
| Bacteroidota | *Arcticibacter sp.* | WP_407426349.1 | WP_407428255.1 | WP_407430250.1 |
| Bacteroidota | *Arcticibacter svalbardensis* | WP_016197349.1 | WP_016193496.1 | WP_016195818.1 |
| Bacteroidota | *Arcticibacter tournemirensis* | WP_141816717.1 | WP_128769017.1 | WP_141816323.1 |
| Bacteroidota | *Arcticibacterium luteifluviistationis* | WP_111371990.1 | WP_111374148.1 | WP_111373335.1 |
| Bacteroidota | *Arenibacter algicola* | WP_093979083.1 | WP_215950131.1 | WP_142189624.1 |
| Bacteroidota | *Arenibacter amylolyticus* | WP_086478617.1 | WP_086478264.1 | WP_086477215.1 |
| Bacteroidota | *Arenibacter aquaticus* | WP_126161809.1 | WP_126163894.1 | WP_126162343.1 |
| Bacteroidota | *Arenibacter arenosicollis* | WP_187583026.1 | WP_187584347.1 | WP_187584983.1 |
| Bacteroidota | *Arenibacter certesii* | WP_026815150.1 | WP_026812673.1 | WP_026812862.1 |
| Bacteroidota | *Arenibacter echinorum* | WP_111622257.1 | WP_111623164.1 | WP_111621924.1 |
| Bacteroidota | *Arenibacter lacus* | WP_150452474.1 | WP_150451320.1 | WP_150450833.1 |

| | | | | |
|---|---|---|---|---|
| Bacteroidota | *Arenibacter latericius* | WP_026811139.1 | WP_026810798.1 | WP_026809646.1 |
| Bacteroidota | *Arenibacter nanhaiticus* | WP_072764415.1 | WP_072765535.1 | WP_072763073.1 |
| Bacteroidota | *Arenibacter palladensis* | WP_332927960.1 | WP_303585876.1 | WP_072862337.1 |
| Bacteroidota | *Arenibacter sp. GZD-96* | WP_322931903.1 | WP_322930698.1 | WP_322929451.1 |
| Bacteroidota | *Arenibacter troitsensis* | WP_085497318.1 | WP_085500282.1 | WP_085495928.1 |
| Bacteroidota | *Aridibaculum aurantiacum* | WP_207496171.1 | WP_207493540.1 | WP_207493862.1 |
| Bacteroidota | *Arsenicibacter rosenii* | WP_071502120.1 | WP_071502670.1 | WP_071505339.1 |
| Bacteroidota | *Arthrospiribacter ruber* | WP_219291266.1 | WP_219289258.1 | WP_219288141.1 |
| Bacteroidota | *Arundinibacter roseus* | WP_132114301.1 | WP_132116095.1 | WP_132121505.1 |
| Bacteroidota | *Ascidiimonas aurantiaca* | WP_340063238.1 | WP_340065951.1 | WP_340063664.1 |
| Bacteroidota | *Ascidiimonas meishanensis* | WP_340202346.1 | WP_340201815.1 | WP_340203336.1 |
| Bacteroidota | *Asinibacterium sp. OR53* | WP_026768116.1 | WP_026770043.1 | WP_026770422.1 |
| Bacteroidota | *Asprobacillus argus* | WP_349240976.1 | WP_349241222.1 | WP_349242390.1 |
| Bacteroidota | *Aurantibacillus circumpalustris* | WP_317897266.1 | WP_317899147.1 | WP_317898617.1 |
| Bacteroidota | *Aurantibacter aestuarii* | WP_106463421.1 | WP_106463824.1 | WP_106463759.1 |
| Bacteroidota | *Aurantibacter crassamenti* | WP_203252934.1 | WP_203253053.1 | WP_203254514.1 |
| Bacteroidota | *Aurantibacter sp.* | WP_348148950.1 | WP_375239853.1 | WP_375239007.1 |
| Bacteroidota | *Aureibacter tunicatorum* | WP_309937555.1 | WP_309941955.1 | WP_309937614.1 |
| Bacteroidota | *Aureibaculum algae* | WP_138950357.1 | WP_138950222.1 | WP_138948862.1 |
| Bacteroidota | *Aureibaculum flavum* | WP_198839813.1 | WP_198839693.1 | WP_198842913.1 |
| Bacteroidota | *Aureibaculum luteum* | WP_117883329.1 | WP_117883048.1 | WP_117885144.1 |
| Bacteroidota | *Aureibaculum marinum* | WP_123896962.1 | WP_123897081.1 | WP_123896527.1 |
| Bacteroidota | *Aureibaculum sp. 2210JD6-5* | WP_321974792.1 | WP_321976430.1 | WP_321975147.1 |
| Bacteroidota | *Aureicoccus marinus* | WP_105000905.1 | WP_105000782.1 | WP_105000272.1 |
| Bacteroidota | *Aureisphaera galaxeae* | WP_272862417.1 | WP_272863078.1 | WP_272861625.1 |
| Bacteroidota | *Aureispira anguillae* | WP_264788426.1 | WP_264791552.1 | WP_264791511.1 |
| Bacteroidota | *Aureitalea sp. L0-47* | WP_265198881.1 | WP_265199191.1 | WP_265200177.1 |
| Bacteroidota | *Aureivirga marina* | WP_196892692.1 | WP_196894274.1 | WP_196894243.1 |

| | | | | |
|---|---|---|---|---|
| Bacteroidota | *Aureivirga sp. CE67* | WP_196888611.1 | WP_196888714.1 | WP_196887798.1 |
| Bacteroidota | *Autumnicola edwardsiae* | WP_311484892.1 | WP_311485020.1 | WP_311484481.1 |
| Bacteroidota | *Autumnicola lenta* | WP_311495763.1 | WP_311493373.1 | WP_311493415.1 |
| Bacteroidota | *Autumnicola musiva* | WP_311503703.1 | WP_311502472.1 | WP_311502434.1 |
| Bacteroidota | *Autumnicola patrickiae* | WP_311684452.1 | WP_311679862.1 | WP_311679937.1 |
| Bacteroidota | *Autumnicola psychrophila* | WP_311499815.1 | WP_311499532.1 | WP_311500054.1 |
| Bacteroidota | *Autumnicola tepida* | WP_311533564.1 | WP_311534518.1 | WP_311535150.1 |
| Bacteroidota | *Avrilella dinanensis* | WP_277631760.1 | WP_100677930.1 | WP_100678779.1 |
| Bacteroidota | *Belliella alkalica* | WP_241411179.1 | WP_241409255.1 | WP_241411311.1 |
| Bacteroidota | *Belliella aquatica* | WP_188442687.1 | WP_188439546.1 | WP_188439325.1 |
| Bacteroidota | *Belliella baltica* | WP_014772208.1 | WP_014773237.1 | WP_014771954.1 |
| Bacteroidota | *Belliella buryatensis* | WP_089237558.1 | WP_089239844.1 | WP_089240194.1 |
| Bacteroidota | *Belliella calami* | WP_241276453.1 | WP_241275745.1 | WP_241275082.1 |
| Bacteroidota | *Belliella filtrata* | WP_241347112.1 | WP_241346774.1 | WP_241350002.1 |
| Bacteroidota | *Belliella kenyensis* | WP_241296136.1 | WP_241291243.1 | WP_241292937.1 |
| Bacteroidota | *Belliella marina* | WP_376886587.1 | WP_376887078.1 | WP_376886810.1 |
| Bacteroidota | *Belliella pelovolcani* | WP_409629045.1 | WP_076502952.1 | WP_076502588.1 |
| Bacteroidota | *Bergeyella cardium* | WP_120489413.1 | WP_160223991.1 | WP_160224032.1 |
| Bacteroidota | *Bergeyella porcorum* | WP_327985048.1 | WP_327984028.1 | WP_376832879.1 |
| Bacteroidota | *Bergeyella sp. RCAD1439* | WP_327609899.1 | WP_327610773.1 | WP_327610132.1 |
| Bacteroidota | *Bergeyella zoohelcum* | WP_125150706.1 | WP_002686932.1 | WP_002689017.1 |
| Bacteroidota | *Bernardetia litoralis* | WP_014799449.1 | WP_014796014.1 | WP_014797464.1 |
| Bacteroidota | *Bernardetia sp.* | WP_291727868.1 | WP_291727328.1 | WP_291721233.1 |
| Bacteroidota | *Bizionia arctica* | WP_188463855.1 | WP_188463365.1 | WP_188465515.1 |
| Bacteroidota | *Bizionia argentinensis* | WP_008634538.1 | WP_008636439.1 | WP_008637444.1 |
| Bacteroidota | *Bizionia echini* | WP_092207375.1 | WP_339635095.1 | WP_092209708.1 |
| Bacteroidota | *Bizionia hallyeonensis* | WP_376859852.1 | WP_376859305.1 | WP_376860208.1 |
| Bacteroidota | *Bizionia myxarmorum* | WP_148404171.1 | WP_148405154.1 | WP_148403799.1 |

| | | | | |
|---|---|---|---|---|
| Bacteroidota | *Bizionia paragorgiae* | WP_318908187.1 | WP_092135828.1 | WP_417236466.1 |
| Bacteroidota | *Bizionia sediminis* | WP_376891938.1 | WP_376891554.1 | WP_376891096.1 |
| Bacteroidota | *Bizionia sp.* | WP_417195014.1 | WP_417199749.1 | WP_417238594.1 |
| Bacteroidota | *Botryobacter ruber* | WP_114781647.1 | WP_114781543.1 | WP_240676079.1 |
| Bacteroidota | *Brumimicrobium aurantiacum* | WP_116880684.1 | WP_116880515.1 | WP_116881141.1 |
| Bacteroidota | *Brumimicrobium glaciale* | WP_130094022.1 | WP_242494536.1 | WP_165366249.1 |
| Bacteroidota | *Brumimicrobium mesophilum* | WP_107040258.1 | WP_107039225.1 | WP_107039918.1 |
| Bacteroidota | *Brumimicrobium oceani* | WP_109358790.1 | WP_233244208.1 | WP_109360517.1 |
| Bacteroidota | *Brumimicrobium salinarum* | WP_101335045.1 | WP_101334281.1 | WP_101335294.1 |
| Bacteroidota | *Brumimicrobium sp.* | WP_417265319.1 | WP_417266280.1 | WP_417265785.1 |
| Bacteroidota | *Candidatus Arcticimaribacter forsetii* | WP_252158231.1 | WP_252158092.1 | WP_252158673.1 |
| Bacteroidota | *Candidatus Brachybacter algidus* | WP_287401746.1 | WP_287400426.1 | WP_287402622.1 |
| Bacteroidota | *Candidatus Kaistella beijingensis* | WP_224135010.1 | WP_224134386.1 | WP_224134929.1 |
| Bacteroidota | *Candidatus Ornithobacterium hominis* | WP_119058188.1 | WP_119058084.1 | WP_119058210.1 |
| Bacteroidota | *Candidatus Ulvibacter alkanivorans* | WP_114491340.1 | WP_114491714.1 | WP_114491961.1 |
| Bacteroidota | *Capnocytophaga bilenii* | WP_208058769.1 | WP_288898588.1 | WP_208059362.1 |
| Bacteroidota | *Capnocytophaga canimorsus* | WP_095918297.1 | WP_212891728.1 | WP_212901490.1 |
| Bacteroidota | *Capnocytophaga canis* | WP_119653065.1 | WP_172919092.1 | WP_042347325.1 |
| Bacteroidota | *Capnocytophaga catalasegens* | WP_264846145.1 | WP_264846002.1 | WP_264846610.1 |
| Bacteroidota | *Capnocytophaga cynodegmi* | WP_405251651.1 | WP_238253373.1 | WP_018278526.1 |
| Bacteroidota | *Capnocytophaga felis* | WP_155285449.1 | WP_155284653.1 | WP_155284893.1 |
| Bacteroidota | *Capnocytophaga genosp. AHN8471* | WP_203093402.1 | WP_203080770.1 | WP_203081813.1 |
| Bacteroidota | *Capnocytophaga gingivalis* | WP_273084178.1 | WP_424651423.1 | WP_095911379.1 |
| Bacteroidota | *Capnocytophaga granulosa* | WP_424655771.1 | WP_314810622.1 | WP_424656280.1 |
| Bacteroidota | *Capnocytophaga haemolytica* | WP_066429520.1 | WP_066429263.1 | WP_066430160.1 |
| Bacteroidota | *Capnocytophaga leadbetteri* | WP_095912921.1 | WP_304348063.1 | WP_311338546.1 |
| Bacteroidota | *Capnocytophaga ochracea* | WP_264854235.1 | WP_002676140.1 | WP_264852541.1 |
| Bacteroidota | *Capnocytophaga periodontitidis* | WP_198472540.1 | WP_198475329.1 | WP_198472515.1 |

| | | | | |
|---|---|---|---|---|
| Bacteroidota | *Capnocytophaga sp.* | WP_316374321.1 | WP_303076861.1 | WP_308766617.1 |
| Bacteroidota | *Capnocytophaga sputigena* | WP_424653469.1 | WP_314278883.1 | WP_002681527.1 |
| Bacteroidota | *Capnocytophaga stomatis* | WP_095894594.1 | WP_405254939.1 | WP_203969405.1 |
| Bacteroidota | *Catalinimonas alkaloidigena* | WP_089681298.1 | WP_089683750.1 | WP_277486639.1 |
| Bacteroidota | *Catalinimonas niigatensis* | WP_302248781.1 | WP_302253366.1 | WP_302248794.1 |
| Bacteroidota | *Catalinimonas sp. 4WD22* | WP_341296119.1 | WP_341294730.1 | WP_341297279.1 |
| Bacteroidota | *Cecembia calidifontis* | WP_130274964.1 | WP_130276585.1 | WP_130274130.1 |
| Bacteroidota | *Cecembia lonarensis* | WP_009186108.1 | WP_009185929.1 | WP_009186875.1 |
| Bacteroidota | *Cecembia rubra* | WP_106568491.1 | WP_106565721.1 | WP_106568070.1 |
| Bacteroidota | *Cecembia sp.* | WP_291784869.1 | WP_291778684.1 | WP_291784052.1 |
| Bacteroidota | *Cellulophaga baltica* | WP_025614771.1 | WP_282164778.1 | WP_025614484.1 |
| Bacteroidota | *Cellulophaga fucicola* | WP_072301909.1 | WP_072302457.1 | WP_072302783.1 |
| Bacteroidota | *Cellulophaga lytica* | WP_303519488.1 | WP_038506972.1 | WP_075694236.1 |
| Bacteroidota | *Cellulophaga omnivescoria* | WP_077402429.1 | WP_271081074.1 | WP_077398634.1 |
| Bacteroidota | *Cellulophaga sp. HaHaR_3_176* | WP_216785145.1 | WP_216785278.1 | WP_216784904.1 |
| Bacteroidota | *Cellulophaga tyrosinoxydans* | WP_084061103.1 | WP_084060995.1 | WP_084062724.1 |
| Bacteroidota | *Cerina litoralis* | WP_317902400.1 | WP_317902167.1 | WP_317901971.1 |
| Bacteroidota | *Cesiribacter sp. SM1* | WP_224999480.1 | WP_224998922.1 | WP_224999441.1 |
| Bacteroidota | *Changchengzhania lutea* | WP_142785956.1 | WP_142783246.1 | WP_142783659.1 |
| Bacteroidota | *Chishuiella changwenlii* | WP_072931684.1 | WP_333662326.1 | WP_333660450.1 |
| Bacteroidota | *Chishuiella sp.* | WP_313375460.1 | WP_313373552.1 | WP_313579345.1 |
| Bacteroidota | *Chitinophaga agri* | WP_162333742.1 | WP_162331142.1 | WP_238430125.1 |
| Bacteroidota | *Chitinophaga agrisoli* | WP_149836969.1 | WP_149837932.1 | WP_149840340.1 |
| Bacteroidota | *Chitinophaga alhagiae* | WP_119077216.1 | WP_119079661.1 | WP_157986662.1 |
| Bacteroidota | *Chitinophaga arvensicola* | WP_089889289.1 | WP_089896020.1 | WP_177192308.1 |
| Bacteroidota | *Chitinophaga barathri* | WP_120518200.1 | WP_120514463.1 | WP_162946241.1 |
| Bacteroidota | *Chitinophaga caeni* | WP_098192515.1 | WP_098194845.1 | WP_157760992.1 |
| Bacteroidota | *Chitinophaga caseinilytica* | WP_341842633.1 | WP_341839141.1 | WP_341840093.1 |

| | | | | |
|---|---|---|---|---|
| Bacteroidota | *Chitinophaga costaii* | WP_089715512.1 | WP_089710726.1 | WP_089715615.1 |
| Bacteroidota | *Chitinophaga cymbidii* | WP_146858207.1 | WP_146866479.1 | WP_186831128.1 |
| Bacteroidota | *Chitinophaga defluvii* | WP_354661261.1 | WP_354660547.1 | WP_354659270.1 |
| Bacteroidota | *Chitinophaga deserti* | WP_109694755.1 | WP_109699103.1 | WP_157962811.1 |
| Bacteroidota | *Chitinophaga dinghuensis* | WP_111591532.1 | WP_111592684.1 | WP_111595453.1 |
| Bacteroidota | *Chitinophaga eiseniae* | WP_078670110.1 | WP_078670884.1 | WP_078668293.1 |
| Bacteroidota | *Chitinophaga filiformis* | WP_247813009.1 | WP_089836010.1 | WP_089830986.1 |
| Bacteroidota | *Chitinophaga flava* | WP_113616485.1 | WP_113614148.1 | WP_113618438.1 |
| Bacteroidota | *Chitinophaga fulva* | WP_169222926.1 | WP_169225324.1 | WP_169227571.1 |
| Bacteroidota | *Chitinophaga ginsengisoli* | WP_106600515.1 | WP_106605529.1 | WP_106600880.1 |
| Bacteroidota | *Chitinophaga horti* | WP_244842465.1 | WP_264282605.1 | WP_264282541.1 |
| Bacteroidota | *Chitinophaga hostae* | WP_211973759.1 | WP_211974175.1 | WP_211973914.1 |
| Bacteroidota | *Chitinophaga japonensis* | WP_145711536.1 | WP_145715336.1 | WP_145717070.1 |
| Bacteroidota | *Chitinophaga jiangningensis* | WP_073084114.1 | WP_073086198.1 | WP_073087053.1 |
| Bacteroidota | *Chitinophaga lutea* | WP_123845536.1 | WP_123849538.1 | WP_158618180.1 |
| Bacteroidota | *Chitinophaga niabensis* | WP_343304213.1 | WP_343308041.1 | WP_343308270.1 |
| Bacteroidota | *Chitinophaga niastensis* | WP_106526122.1 | WP_106528908.1 | WP_106530351.1 |
| Bacteroidota | *Chitinophaga nivalis* | WP_264735150.1 | WP_264728856.1 | WP_264732223.1 |
| Bacteroidota | *Chitinophaga oryzae* | WP_168807646.1 | WP_168802561.1 | WP_168804602.1 |
| Bacteroidota | *Chitinophaga oryziterrae* | WP_157302517.1 | WP_157302627.1 | WP_157300745.1 |
| Bacteroidota | *Chitinophaga parva* | WP_108684680.1 | WP_108687857.1 | WP_108687548.1 |
| Bacteroidota | *Chitinophaga pinensis* | WP_012788863.1 | WP_146306657.1 | WP_012792326.1 |
| Bacteroidota | *Chitinophaga pollutisoli* | WP_341838311.1 | WP_341836676.1 | WP_341835840.1 |
| Bacteroidota | *Chitinophaga polysaccharea* | WP_145660751.1 | WP_142685119.1 | WP_142688553.1 |
| Bacteroidota | *Chitinophaga qingshengii* | WP_188091016.1 | WP_188086724.1 | WP_188091470.1 |
| Bacteroidota | *Chitinophaga rhizophila* | WP_220251837.1 | WP_220253029.1 | WP_220252641.1 |
| Bacteroidota | *Chitinophaga rhizosphaerae* | WP_126245588.1 | WP_126249185.1 | WP_164714245.1 |
| Bacteroidota | *Chitinophaga rupis* | WP_089920443.1 | WP_089919493.1 | WP_089909602.1 |

| | | | | |
|---|---|---|---|---|
| Bacteroidota | *Chitinophaga sancti* | WP_320580275.1 | WP_320572784.1 | WP_072357357.1 |
| Bacteroidota | *Chitinophaga silvatica* | WP_116975180.1 | WP_116973669.1 | WP_116976693.1 |
| Bacteroidota | *Chitinophaga silvisoli* | WP_116853605.1 | WP_116854710.1 | WP_116854545.1 |
| Bacteroidota | *Chitinophaga skermanii* | WP_111595753.1 | WP_111597493.1 | WP_111596564.1 |
| Bacteroidota | *Chitinophaga solisilvae* | WP_127037710.1 | WP_160712735.1 | WP_160716070.1 |
| Bacteroidota | *Chitinophaga sp.* | WP_326461457.1 | WP_343703673.1 | WP_291942926.1 |
| Bacteroidota | *Chitinophaga terrae (ex Kim and Jung 2007)* | WP_089758629.1 | WP_089759461.1 | WP_089760460.1 |
| Bacteroidota | *Chitinophaga tropicalis* | WP_157304928.1 | WP_157309866.1 | WP_157306277.1 |
| Bacteroidota | *Chitinophaga varians* | WP_188096056.1 | WP_188101673.1 | WP_168873033.1 |
| Bacteroidota | *Chitinophaga vietnamensis* | WP_143310755.1 | WP_143310472.1 | WP_143306111.1 |
| Bacteroidota | *Chondrinema litorale* | WP_284682294.1 | WP_284684089.1 | WP_284684822.1 |
| Bacteroidota | *Christiangramia aestuarii* | WP_156276367.1 | WP_156277420.1 | WP_156273350.1 |
| Bacteroidota | *Christiangramia aquimixticola* | WP_417907117.1 | WP_417907863.1 | WP_417905913.1 |
| Bacteroidota | *Christiangramia crocea* | WP_240099472.1 | WP_240100482.1 | WP_240100745.1 |
| Bacteroidota | *Christiangramia echinicola* | WP_089663554.1 | WP_026934815.1 | WP_089662858.1 |
| Bacteroidota | *Christiangramia flava* | WP_083645604.1 | WP_083645022.1 | WP_083644987.1 |
| Bacteroidota | *Christiangramia forsetii* | WP_011709514.1 | WP_011708676.1 | WP_011710100.1 |
| Bacteroidota | *Christiangramia fulva* | WP_107013167.1 | WP_107012684.1 | WP_107012643.1 |
| Bacteroidota | *Christiangramia gaetbulicola* | WP_108171519.1 | WP_108172085.1 | WP_108172706.1 |
| Bacteroidota | *Christiangramia lutea* | WP_240713313.1 | WP_240714481.1 | WP_240713769.1 |
| Bacteroidota | *Christiangramia marina* | WP_417920970.1 | WP_417921409.1 | WP_417920483.1 |
| Bacteroidota | *Christiangramia oceanisediminis* | WP_241551229.1 | WP_241552613.1 | WP_241552703.1 |
| Bacteroidota | *Christiangramia portivictoriae* | WP_026916233.1 | WP_026914858.1 | WP_026915542.1 |
| Bacteroidota | *Christiangramia sabulilitoris* | WP_143409212.1 | WP_143409686.1 | WP_143411055.1 |
| Bacteroidota | *Christiangramia salexigens* | WP_072552188.1 | WP_072551724.1 | WP_072552596.1 |
| Bacteroidota | *Christiangramia sediminis* | WP_229337945.1 | WP_229337042.1 | WP_229339163.1 |
| Bacteroidota | *Christiangramia sp.* | WP_300437579.1 | WP_347410098.1 | WP_300434533.1 |

| | | | | |
|---|---|---|---|---|
| Bacteroidota | *Chryseobacterium angstadtii* | WP_048504691.1 | WP_048508403.1 | WP_048504839.1 |
| Bacteroidota | *Chryseobacterium aquaeductus* | WP_162087455.1 | WP_162088291.1 | WP_162087563.1 |
| Bacteroidota | *Chryseobacterium aquaticum* | WP_332453612.1 | WP_332454044.1 | WP_332453903.1 |
| Bacteroidota | *Chryseobacterium arachidis* | WP_072953422.1 | WP_072962042.1 | WP_072953694.1 |
| Bacteroidota | *Chryseobacterium arthrosphaerae* | WP_065401126.1 | WP_065399462.1 | WP_278377839.1 |
| Bacteroidota | *Chryseobacterium artocarpi* | WP_065393438.1 | WP_065395308.1 | WP_065395850.1 |
| Bacteroidota | *Chryseobacterium aureum* | WP_126653405.1 | WP_126652494.1 | WP_126653278.1 |
| Bacteroidota | *Chryseobacterium binzhouense* | WP_143884827.1 | WP_312344148.1 | WP_143885655.1 |
| Bacteroidota | *Chryseobacterium camelliae* | WP_100076556.1 | WP_100075739.1 | WP_100076441.1 |
| Bacteroidota | *Chryseobacterium carnipullorum* | WP_073335484.1 | WP_073329474.1 | WP_123877096.1 |
| Bacteroidota | *Chryseobacterium cheonjiense* | WP_169231258.1 | WP_169232082.1 | WP_169231368.1 |
| Bacteroidota | *Chryseobacterium contaminans* | WP_066699585.1 | WP_336963978.1 | WP_336962821.1 |
| Bacteroidota | *Chryseobacterium culicis* | WP_193546784.1 | WP_105681681.1 | WP_089692756.1 |
| Bacteroidota | *Chryseobacterium daecheongense* | WP_123263640.1 | WP_245143878.1 | WP_245142849.1 |
| Bacteroidota | *Chryseobacterium daeguense* | WP_027380310.1 | WP_027381351.1 | WP_027380381.1 |
| Bacteroidota | *Chryseobacterium defluvii* | WP_184192087.1 | WP_121459854.1 | WP_121462258.1 |
| Bacteroidota | *Chryseobacterium echinoideorum* | WP_144283498.1 | WP_144281767.1 | WP_144281870.1 |
| Bacteroidota | *Chryseobacterium edaphi* | WP_263004635.1 | WP_263003815.1 | WP_263003315.1 |
| Bacteroidota | *Chryseobacterium endalhagicum* | WP_202092377.1 | WP_202088381.1 | WP_202092137.1 |
| Bacteroidota | *Chryseobacterium flavum* | WP_115962526.1 | WP_115959980.1 | WP_333596648.1 |
| Bacteroidota | *Chryseobacterium fluminis* | WP_266173542.1 | WP_266172508.1 | WP_266173392.1 |
| Bacteroidota | *Chryseobacterium foetidum* | WP_262150941.1 | WP_262149356.1 | WP_262150698.1 |
| Bacteroidota | *Chryseobacterium formosense* | WP_034678117.1 | WP_034679053.1 | WP_034677951.1 |
| Bacteroidota | *Chryseobacterium formosus* | WP_267266397.1 | WP_267266997.1 | WP_267266551.1 |
| Bacteroidota | *Chryseobacterium gambrini* | WP_421744305.1 | WP_421745152.1 | WP_271160214.1 |
| Bacteroidota | *Chryseobacterium gilvum* | WP_262989998.1 | WP_262991445.1 | WP_262990122.1 |
| Bacteroidota | *Chryseobacterium ginsengisoli* | WP_345204417.1 | WP_345207700.1 | WP_345204007.1 |
| Bacteroidota | *Chryseobacterium ginsenosidimutans* | WP_306617673.1 | WP_344823491.1 | WP_344828136.1 |

| | | | | |
|---|---|---|---|---|
| Bacteroidota | *Chryseobacterium gleum* | WP_002981824.1 | WP_002976287.1 | WP_002982977.1 |
| Bacteroidota | *Chryseobacterium gotjawalense* | WP_282903957.1 | WP_282903963.1 | WP_282903911.1 |
| Bacteroidota | *Chryseobacterium gregarium* | WP_027387127.1 | WP_027387487.1 | WP_027387596.1 |
| Bacteroidota | *Chryseobacterium group* | WP_063971113.1 | WP_063971246.1 | WP_063971373.1 |
| Bacteroidota | *Chryseobacterium gwangjuense* | WP_233112162.1 | WP_233112332.1 | WP_233112287.1 |
| Bacteroidota | *Chryseobacterium hagamense* | WP_146939999.1 | WP_146942032.1 | WP_146939862.1 |
| Bacteroidota | *Chryseobacterium herbae* | WP_259839341.1 | WP_259838793.1 | WP_259839482.1 |
| Bacteroidota | *Chryseobacterium indologenes* | WP_213277952.1 | WP_061084393.1 | WP_213277829.1 |
| Bacteroidota | *Chryseobacterium jejuense* | WP_209919674.1 | WP_209916422.1 | WP_209918947.1 |
| Bacteroidota | *Chryseobacterium koreense* | WP_304343474.1 | WP_304343482.1 | WP_304343634.1 |
| Bacteroidota | *Chryseobacterium kwangjuense* | WP_062650765.1 | WP_062650258.1 | WP_409358204.1 |
| Bacteroidota | *Chryseobacterium lactis* | WP_047098204.1 | WP_185288522.1 | WP_103289094.1 |
| Bacteroidota | *Chryseobacterium lacus* | WP_114304519.1 | WP_114302695.1 | WP_114303026.1 |
| Bacteroidota | *Chryseobacterium lathyri* | WP_307211175.1 | WP_306845405.1 | WP_111956929.1 |
| Bacteroidota | *Chryseobacterium limigenitum* | WP_072407513.1 | WP_072411150.1 | WP_072407276.1 |
| Bacteroidota | *Chryseobacterium luquanense* | WP_267280638.1 | WP_267282236.1 | WP_267280772.1 |
| Bacteroidota | *Chryseobacterium luteum* | WP_034702980.1 | WP_034701920.1 | WP_034702712.1 |
| Bacteroidota | *Chryseobacterium manosquense* | WP_123248881.1 | WP_188322246.1 | WP_188320621.1 |
| Bacteroidota | *Chryseobacterium mucoviscidosis* | WP_087707876.1 | WP_336717035.1 | WP_087706563.1 |
| Bacteroidota | *Chryseobacterium muglaense* | WP_191178277.1 | WP_191179160.1 | WP_191178394.1 |
| Bacteroidota | *Chryseobacterium nakagawai* | WP_123858886.1 | WP_123857931.1 | WP_123858751.1 |
| Bacteroidota | *Chryseobacterium nepalense* | WP_248390388.1 | WP_326988922.1 | WP_326988498.1 |
| Bacteroidota | *Chryseobacterium oleae* | WP_090025259.1 | WP_090025667.1 | WP_090025098.1 |
| Bacteroidota | *Chryseobacterium oranimense* | WP_260540510.1 | WP_260542181.1 | WP_040993467.1 |
| Bacteroidota | *Chryseobacterium oryctis* | WP_264743935.1 | WP_264742216.1 | WP_264744072.1 |
| Bacteroidota | *Chryseobacterium panacisoli* | WP_047382874.1 | WP_149387241.1 | WP_149387698.1 |
| Bacteroidota | *Chryseobacterium paridis* | WP_200241446.1 | WP_200248911.1 | WP_200241814.1 |
| Bacteroidota | *Chryseobacterium pennipullorum* | WP_115929148.1 | WP_115928623.1 | WP_115927319.1 |

| | | | | |
|---|---|---|---|---|
| Bacteroidota | *Chryseobacterium phocaeense* | WP_080777849.1 | WP_080776683.1 | WP_080777716.1 |
| Bacteroidota | *Chryseobacterium phosphatilyticum* | WP_103248064.1 | WP_103247118.1 | WP_109713458.1 |
| Bacteroidota | *Chryseobacterium piperi* | WP_034687838.1 | WP_034681116.1 | WP_034680385.1 |
| Bacteroidota | *Chryseobacterium piscicola* | WP_076449729.1 | WP_076452144.1 | WP_076449478.1 |
| Bacteroidota | *Chryseobacterium polytrichastri* | WP_073295856.1 | WP_073293519.1 | WP_073296113.1 |
| Bacteroidota | *Chryseobacterium populi* | WP_007846994.1 | WP_034669435.1 | WP_007847155.1 |
| Bacteroidota | *Chryseobacterium profundimaris* | WP_283421978.1 | WP_283422706.1 | WP_283421026.1 |
| Bacteroidota | *Chryseobacterium salviniae* | WP_326320873.1 | WP_326319633.1 | WP_326321868.1 |
| Bacteroidota | *Chryseobacterium scophthalmum* | WP_427317969.1 | WP_210150477.1 | WP_427313732.1 |
| Bacteroidota | *Chryseobacterium sediminis* | WP_309876686.1 | WP_184557784.1 | WP_184558456.1 |
| Bacteroidota | *Chryseobacterium shandongense* | WP_123852675.1 | WP_123854981.1 | WP_123854495.1 |
| Bacteroidota | *Chryseobacterium shigense* | WP_184164982.1 | WP_076508834.1 | WP_076507142.1 |
| Bacteroidota | *Chryseobacterium soldanellicola* | WP_089754460.1 | WP_089756315.1 | WP_089754742.1 |
| Bacteroidota | *Chryseobacterium sp.* | WP_312396624.1 | WP_375180805.1 | WP_317132566.1 |
| Bacteroidota | *Chryseobacterium suipulveris* | WP_243548920.1 | WP_243548777.1 | WP_243550068.1 |
| Bacteroidota | *Chryseobacterium taeanense* | WP_374458709.1 | WP_374460750.1 | WP_089856336.1 |
| Bacteroidota | *Chryseobacterium tagetis* | WP_225686926.1 | WP_225690509.1 | WP_225687090.1 |
| Bacteroidota | *Chryseobacterium taichungense* | WP_089999797.1 | WP_312899911.1 | WP_312902982.1 |
| Bacteroidota | *Chryseobacterium taihuense* | WP_089744917.1 | WP_089744036.1 | WP_089741825.1 |
| Bacteroidota | *Chryseobacterium taiwanense* | WP_039364300.1 | WP_039365139.1 | WP_039364551.1 |
| Bacteroidota | *Chryseobacterium takakiae* | WP_072883998.1 | WP_072883461.1 | WP_072883882.1 |
| Bacteroidota | *Chryseobacterium taklimakanense* | WP_095071275.1 | WP_095071556.1 | WP_277111743.1 |
| Bacteroidota | *Chryseobacterium terrae* | WP_408090487.1 | WP_408091166.1 | WP_408090916.1 |
| Bacteroidota | *Chryseobacterium tructae* | WP_290301454.1 | WP_290295476.1 | WP_290301759.1 |
| Bacteroidota | *Chryseobacterium turcicum* | WP_230672020.1 | WP_230667182.1 | WP_230672276.1 |
| Bacteroidota | *Chryseobacterium ureilyticum* | WP_076553735.1 | WP_076551667.1 | WP_076552180.1 |
| Bacteroidota | *Chryseobacterium vietnamense* | WP_047429091.1 | WP_310405478.1 | WP_310404856.1 |
| Bacteroidota | *Chryseobacterium vrystaatense* | WP_034738641.1 | WP_034746935.1 | WP_034738968.1 |

| | | | | |
|---|---|---|---|---|
| Bacteroidota | *Chryseobacterium wanjuense* | WP_089793124.1 | WP_089795693.1 | WP_089793430.1 |
| Bacteroidota | *Chryseobacterium zhengzhouense* | WP_378177090.1 | WP_378181010.1 | WP_378176574.1 |
| Bacteroidota | *Chryseolinea lacunae* | WP_202010734.1 | WP_202013028.1 | WP_202008930.1 |
| Bacteroidota | *Chryseolinea serpens* | WP_073133656.1 | WP_073130757.1 | WP_073134550.1 |
| Bacteroidota | *Chryseolinea soli* | WP_119754613.1 | WP_119757032.1 | WP_119754129.1 |
| Bacteroidota | *Chryseolinea sp. H1M3-3* | WP_276373192.1 | WP_276374865.1 | WP_276368923.1 |
| Bacteroidota | *Chryseosolibacter histidini* | WP_254170009.1 | WP_254162248.1 | WP_254168819.1 |
| Bacteroidota | *Chryseosolibacter indicus* | WP_254155073.1 | WP_254151535.1 | WP_254152595.1 |
| Bacteroidota | *Chryseotalea sanaruensis* | WP_127124308.1 | WP_127123771.1 | WP_127123286.1 |
| Bacteroidota | *Cloacibacterium caeni* | WP_213197414.1 | WP_213196799.1 | WP_213196929.1 |
| Bacteroidota | *Cloacibacterium normanense* | WP_371488702.1 | WP_409623128.1 | WP_069800177.1 |
| Bacteroidota | *Cloacibacterium rupense* | WP_188618261.1 | WP_188616825.1 | WP_229663381.1 |
| Bacteroidota | *Cloacibacterium sp.* | WP_424305291.1 | WP_424304899.1 | WP_414982983.1 |
| Bacteroidota | *Cnuella takakiae* | WP_073048635.1 | WP_073047240.1 | WP_073046215.1 |
| Bacteroidota | *Cochleicola gelatinilyticus* | WP_068592651.1 | WP_068591820.1 | WP_068588824.1 |
| Bacteroidota | *Cognataquiflexum aquatile* | WP_113923999.1 | WP_113922158.1 | WP_113923884.1 |
| Bacteroidota | *Cognataquiflexum nitidum* | WP_241124226.1 | WP_241126391.1 | WP_241124347.1 |
| Bacteroidota | *Cognataquiflexum rubidum* | WP_241139927.1 | WP_241143543.1 | WP_241140045.1 |
| Bacteroidota | *Cognatitamlana onchidii* | WP_136481225.1 | WP_136481392.1 | WP_136480656.1 |
| Bacteroidota | *Compostibacter hankyongensis* | WP_344975939.1 | WP_344977829.1 | WP_344977805.1 |
| Bacteroidota | *Confluentibacter citreus* | WP_100614548.1 | WP_100614611.1 | WP_100615531.1 |
| Bacteroidota | *Confluentibacter flavum* | WP_106660758.1 | WP_106658464.1 | WP_106659928.1 |
| Bacteroidota | *Confluentibacter lentus* | WP_100611433.1 | WP_100611373.1 | WP_100612254.1 |
| Bacteroidota | *Confluentibacter sediminis* | WP_111307457.1 | WP_111307398.1 | WP_111309144.1 |
| Bacteroidota | *Constantimarinum furrinae* | WP_186988759.1 | WP_186989389.1 | WP_186987917.1 |
| Bacteroidota | *Corallibacter sp.* | WP_417290984.1 | WP_418509762.1 | WP_418509659.1 |
| Bacteroidota | *Costertonia aggregata* | WP_179242954.1 | WP_179243042.1 | WP_179242674.1 |
| Bacteroidota | *Croceibacter atlanticus* | WP_348661630.1 | WP_341200636.1 | WP_348661122.1 |

| | | | | |
|---|---|---|---|---|
| Bacteroidota | *Croceimicrobium hydrocarbonivorans* | WP_210758119.1 | WP_210758057.1 | WP_210758765.1 |
| Bacteroidota | *Croceimicrobium sp.* | WP_421753003.1 | WP_421752888.1 | WP_421753676.1 |
| Bacteroidota | *Croceitalea dokdonensis* | WP_054560488.1 | WP_054557427.1 | WP_054559144.1 |
| Bacteroidota | *Croceitalea marina* | WP_377767746.1 | WP_377766699.1 | WP_377765282.1 |
| Bacteroidota | *Croceitalea rosinachiae* | WP_311353170.1 | WP_311350950.1 | WP_311353314.1 |
| Bacteroidota | *Croceitalea sp. MTPC5* | WP_339144497.1 | WP_339140946.1 | WP_339144417.1 |
| Bacteroidota | *Croceivirga lutea* | WP_185783657.1 | WP_185781643.1 | WP_185783995.1 |
| Bacteroidota | *Croceivirga radicis* | WP_010517378.1 | WP_010517241.1 | WP_080319389.1 |
| Bacteroidota | *Croceivirga sp. JEA036* | WP_167987617.1 | WP_167987710.1 | WP_167989412.1 |
| Bacteroidota | *Croceivirga thetidis* | WP_168552228.1 | WP_168552311.1 | WP_168553752.1 |
| Bacteroidota | *Crocinitomix algicola* | WP_070138319.1 | WP_066755789.1 | WP_066757945.1 |
| Bacteroidota | *Crocinitomix catalasitica* | WP_027419381.1 | WP_027421155.1 | WP_051568672.1 |
| Bacteroidota | *Cruoricaptor ignavus* | WP_392437594.1 | WP_073177968.1 | WP_073179178.1 |
| Bacteroidota | *Cryomorpha ignava* | WP_163283374.1 | WP_163285353.1 | WP_163285151.1 |
| Bacteroidota | *Cyclobacterium amurskyense* | WP_048642148.1 | WP_048642687.1 | WP_048644384.1 |
| Bacteroidota | *Cyclobacterium jeungdonense* | WP_163383924.1 | WP_163385926.1 | WP_163385429.1 |
| Bacteroidota | *Cyclobacterium lianum* | WP_073091641.1 | WP_073096623.1 | WP_073090639.1 |
| Bacteroidota | *Cyclobacterium marinum* | WP_149393370.1 | WP_014021153.1 | WP_014019414.1 |
| Bacteroidota | *Cyclobacterium plantarum* | WP_166151612.1 | WP_166149796.1 | WP_166146008.1 |
| Bacteroidota | *Cyclobacterium qasimii* | WP_020888752.1 | WP_020893013.1 | WP_020890698.1 |
| Bacteroidota | *Cyclobacterium roseum* | WP_162419248.1 | WP_162416044.1 | WP_162418251.1 |
| Bacteroidota | *Cyclobacterium salsum* | WP_162341569.1 | WP_162343545.1 | WP_162343434.1 |
| Bacteroidota | *Cyclobacterium sp.* | WP_291369886.1 | WP_410265425.1 | WP_291372784.1 |
| Bacteroidota | *Cyclobacterium xiamenense* | WP_375584005.1 | WP_375583490.1 | WP_092172779.1 |
| Bacteroidota | *Cytophaga aurantiaca* | WP_018341432.1 | WP_018344950.1 | WP_018344260.1 |
| Bacteroidota | *Cytophaga hutchinsonii* | WP_011583422.1 | WP_011585160.1 | WP_011584089.1 |
| Bacteroidota | *Cytophaga sp. FL35* | WP_187456895.1 | WP_187458128.1 | WP_222937458.1 |
| Bacteroidota | *Daejeonella lutea* | WP_079702447.1 | WP_079700741.1 | WP_079703645.1 |

| | | | | |
|---|---|---|---|---|
| Bacteroidota | *Daejeonella oryzae* | WP_026896934.1 | WP_026898669.1 | WP_026896820.1 |
| Bacteroidota | *Daejeonella rubra* | WP_090706014.1 | WP_090700380.1 | WP_090706315.1 |
| Bacteroidota | *Daejeonella sp.* | WP_411274008.1 | WP_340380637.1 | WP_324485892.1 |
| Bacteroidota | *Danxiaibacter flavus* | WP_369329328.1 | WP_369329440.1 | WP_369330972.1 |
| Bacteroidota | *Dawidia cretensis* | WP_254085524.1 | WP_254087815.1 | WP_254082658.1 |
| Bacteroidota | *Dawidia soli* | WP_254090799.1 | WP_254093836.1 | WP_254092065.1 |
| Bacteroidota | *Deminuibacter soli* | WP_116845410.1 | WP_116847565.1 | WP_116846911.1 |
| Bacteroidota | *Desertivirga arenae* | WP_207535454.1 | WP_207532598.1 | WP_207534650.1 |
| Bacteroidota | *Desertivirga brevis* | WP_207424218.1 | WP_207422904.1 | WP_207424873.1 |
| Bacteroidota | *Desertivirga xinjiangensis* | WP_256013323.1 | WP_256009515.1 | WP_256009196.1 |
| Bacteroidota | *Dinghuibacter silviterrae* | WP_133989948.1 | WP_133993427.1 | WP_133993337.1 |
| Bacteroidota | *Dokdonia donghaensis* | WP_375251903.1 | WP_035324608.1 | WP_035327796.1 |
| Bacteroidota | *Dokdonia pacifica* | WP_089371124.1 | WP_089371369.1 | WP_308489842.1 |
| Bacteroidota | *Dokdonia ponticola* | WP_379978956.1 | WP_379978462.1 | WP_379977171.1 |
| Bacteroidota | *Dokdonia sinensis* | WP_121916639.1 | WP_121917143.1 | WP_121915746.1 |
| Bacteroidota | *Dokdonia sp.* | WP_348168451.1 | WP_348171663.1 | WP_348163442.1 |
| Bacteroidota | *Dyadobacter alkalitolerans* | WP_026629948.1 | WP_026629725.1 | WP_026632001.1 |
| Bacteroidota | *Dyadobacter arcticus* | WP_167269929.1 | WP_167274150.1 | WP_167266676.1 |
| Bacteroidota | *Dyadobacter beijingensis* | WP_019940798.1 | WP_019944208.1 | WP_019944815.1 |
| Bacteroidota | *Dyadobacter bucti* | WP_138479005.1 | WP_138481385.1 | WP_138482428.1 |
| Bacteroidota | *Dyadobacter chenwenxiniae* | WP_234607536.1 | WP_234655679.1 | WP_234658388.1 |
| Bacteroidota | *Dyadobacter crusticola* | WP_031526644.1 | WP_031527697.1 | WP_031528518.1 |
| Bacteroidota | *Dyadobacter diqingensis* | WP_254411353.1 | WP_254412135.1 | WP_254561571.1 |
| Bacteroidota | *Dyadobacter endophyticus* | WP_188935609.1 | WP_188927460.1 | WP_188938595.1 |
| Bacteroidota | *Dyadobacter fanqingshengii* | WP_234612655.1 | WP_235138158.1 | WP_234615245.1 |
| Bacteroidota | *Dyadobacter fermentans* | WP_223403261.1 | WP_012779703.1 | WP_050774706.1 |
| Bacteroidota | *Dyadobacter flavalbus* | WP_139010466.1 | WP_139010683.1 | WP_139012196.1 |
| Bacteroidota | *Dyadobacter frigoris* | WP_137343065.1 | WP_137339996.1 | WP_137341069.1 |

| | | | | |
|---|---|---|---|---|
| Bacteroidota | *Dyadobacter helix* | WP_215237805.1 | WP_215238273.1 | WP_406566865.1 |
| Bacteroidota | *Dyadobacter jejuensis* | WP_109674238.1 | WP_109676885.1 | WP_310588680.1 |
| Bacteroidota | *Dyadobacter koreensis* | WP_090334368.1 | WP_090338592.1 | WP_090340372.1 |
| Bacteroidota | *Dyadobacter luteus* | WP_115833335.1 | WP_115829263.1 | WP_115832780.1 |
| Bacteroidota | *Dyadobacter luticola* | WP_138367717.1 | WP_138363610.1 | WP_138368071.1 |
| Bacteroidota | *Dyadobacter pollutisoli* | WP_244822797.1 | WP_244819587.1 | WP_244822558.1 |
| Bacteroidota | *Dyadobacter psychrophilus* | WP_082212856.1 | WP_082214096.1 | WP_082216753.1 |
| Bacteroidota | *Dyadobacter psychrotolerans* | WP_131955926.1 | WP_131957012.1 | WP_131960551.1 |
| Bacteroidota | *Dyadobacter sandarakinus* | WP_204663031.1 | WP_204660248.1 | WP_204658485.1 |
| Bacteroidota | *Dyadobacter sediminis* | WP_138279644.1 | WP_138281443.1 | WP_138283311.1 |
| Bacteroidota | *Dyadobacter soli* | WP_090147482.1 | WP_090146956.1 | WP_090149517.1 |
| Bacteroidota | *Dyadobacter sp.* | WP_348067880.1 | WP_291206586.1 | WP_291200977.1 |
| Bacteroidota | *Dyadobacter subterraneus* | WP_194122670.1 | WP_194120119.1 | WP_194124010.1 |
| Bacteroidota | *Dyadobacter tibetensis* | WP_025762109.1 | WP_025761910.1 | WP_229236027.1 |
| Bacteroidota | *Echinicola arenosa* | WP_192009333.1 | WP_192010197.1 | WP_192010504.1 |
| Bacteroidota | *Echinicola jeungdonensis* | WP_290247152.1 | WP_290249494.1 | WP_290247245.1 |
| Bacteroidota | *Echinicola marina* | WP_226334858.1 | WP_226333278.1 | WP_226335464.1 |
| Bacteroidota | *Echinicola pacifica* | WP_018472072.1 | WP_018473938.1 | WP_018475493.1 |
| Bacteroidota | *Echinicola rosea* | WP_137401295.1 | WP_137403783.1 | WP_137401942.1 |
| Bacteroidota | *Echinicola salinicaeni* | WP_186753604.1 | WP_186757912.1 | WP_186755834.1 |
| Bacteroidota | *Echinicola shivajiensis* | WP_215225948.1 | WP_215226756.1 | WP_215225276.1 |
| Bacteroidota | *Echinicola soli* | WP_141613416.1 | WP_141615741.1 | WP_141614086.1 |
| Bacteroidota | *Echinicola sp. 20G* | WP_200974775.1 | WP_200978684.1 | WP_200975499.1 |
| Bacteroidota | *Echinicola strongylocentroti* | WP_112783852.1 | WP_112782133.1 | WP_112784437.1 |
| Bacteroidota | *Echinicola vietnamensis* | WP_015264618.1 | WP_015267203.1 | WP_015265338.1 |
| Bacteroidota | *Edaphocola aurantiacus* | WP_222166449.1 | WP_222167030.1 | WP_222166341.1 |
| Bacteroidota | *Edaphocola flava* | WP_129021451.1 | WP_129019585.1 | WP_161971320.1 |
| Bacteroidota | *Eisenibacter elegans* | WP_027000537.1 | WP_027000503.1 | WP_027000333.1 |

| | | | | |
|---|---|---|---|---|
| Bacteroidota | *Ekhidna lutea* | WP_089356447.1 | WP_089357853.1 | WP_089355072.1 |
| Bacteroidota | *Ekhidna sp.* | WP_350219655.1 | WP_420575267.1 | WP_424962706.1 |
| Bacteroidota | *Elizabethkingia anophelis* | WP_407478991.1 | WP_367604863.1 | WP_024567208.1 |
| Bacteroidota | *Elizabethkingia argenteiflava* | WP_166519254.1 | WP_166519561.1 | WP_166520285.1 |
| Bacteroidota | *Elizabethkingia meningoseptica* | WP_326516950.1 | WP_249063179.1 | WP_078770067.1 |
| Bacteroidota | *Elizabethkingia sp. YR214* | WP_107809619.1 | WP_107807644.1 | WP_107807205.1 |
| Bacteroidota | *Elizabethkingia ursingii* | WP_249077995.1 | WP_078403340.1 | WP_249078125.1 |
| Bacteroidota | *Empedobacter brevis* | WP_312557397.1 | WP_019976302.1 | WP_353097325.1 |
| Bacteroidota | *Empedobacter falsenii* | WP_219033575.1 | WP_038336617.1 | WP_416403977.1 |
| Bacteroidota | *Empedobacter sedimenti* | WP_282628566.1 | WP_282630778.1 | WP_282629868.1 |
| Bacteroidota | *Empedobacter tilapiae* | WP_314241946.1 | WP_135834524.1 | WP_314244267.1 |
| Bacteroidota | *Emticicia agri* | WP_130024155.1 | WP_130019593.1 | WP_130020679.1 |
| Bacteroidota | *Emticicia aquatica* | WP_238804023.1 | WP_238806179.1 | WP_238807146.1 |
| Bacteroidota | *Emticicia aquatilis* | WP_188768899.1 | WP_188770815.1 | WP_188765468.1 |
| Bacteroidota | *Emticicia oligotrophica* | WP_305951787.1 | WP_015027285.1 | WP_015028296.1 |
| Bacteroidota | *Emticicia soli* | WP_340239816.1 | WP_340237658.1 | WP_340233778.1 |
| Bacteroidota | *Emticicia sp.* | WP_421769690.1 | WP_421772207.1 | WP_421771868.1 |
| Bacteroidota | *Epilithonimonas arachidiradicis* | WP_120213936.1 | WP_120213882.1 | WP_120212215.1 |
| Bacteroidota | *Epilithonimonas bovis* | WP_076781783.1 | WP_076783416.1 | WP_076783565.1 |
| Bacteroidota | *Epilithonimonas caeni* | WP_027383614.1 | WP_027383664.1 | WP_034671664.1 |
| Bacteroidota | *Epilithonimonas hispanica* | WP_116033190.1 | WP_116033058.1 | WP_116033454.1 |
| Bacteroidota | *Epilithonimonas hominis* | WP_089768928.1 | WP_312750933.1 | WP_313306453.1 |
| Bacteroidota | *Epilithonimonas hungarica* | WP_089874829.1 | WP_307311947.1 | WP_307310474.1 |
| Bacteroidota | *Epilithonimonas lactis* | WP_034976505.1 | WP_034976395.1 | WP_311316404.1 |
| Bacteroidota | *Epilithonimonas mollis* | WP_072997678.1 | WP_072996781.1 | WP_072997439.1 |
| Bacteroidota | *Epilithonimonas pallida* | WP_283416258.1 | WP_283416210.1 | WP_283415997.1 |
| Bacteroidota | *Epilithonimonas sp.* | WP_374440986.1 | WP_312768802.1 | WP_374441781.1 |
| Bacteroidota | *Epilithonimonas tenax* | WP_028121144.1 | WP_028121200.1 | WP_316928783.1 |

| | | | | |
|---|---|---|---|---|
| Bacteroidota | *Epilithonimonas vandammei* | WP_124801570.1 | WP_124801530.1 | WP_312194444.1 |
| Bacteroidota | *Epilithonimonas xixisoli* | WP_133945463.1 | WP_133945376.1 | WP_133944863.1 |
| Bacteroidota | *Epilithonimonas zeae* | WP_074236880.1 | WP_074236937.1 | WP_248875071.1 |
| Bacteroidota | *Eudoraea adriatica* | WP_019671452.1 | WP_019671554.1 | WP_026348505.1 |
| Bacteroidota | *Eudoraea chungangensis* | WP_276390869.1 | WP_276391857.1 | WP_276391193.1 |
| Bacteroidota | *Eudoraea sp.* | WP_424290759.1 | WP_424281830.1 | WP_424285184.1 |
| Bacteroidota | *Euzebyella marina* | WP_121849468.1 | WP_121849330.1 | WP_121850639.1 |
| Bacteroidota | *Euzebyella saccharophila* | WP_192462928.1 | WP_192463011.1 | WP_192462489.1 |
| Bacteroidota | *Faecalibacter bovis* | WP_230477862.1 | WP_230476304.1 | WP_230477768.1 |
| Bacteroidota | *Faecalibacter macacae* | WP_121935498.1 | WP_121934504.1 | WP_121933187.1 |
| Bacteroidota | *Faecalibacter rhinopitheci* | WP_194183591.1 | WP_194181660.1 | WP_194183128.1 |
| Bacteroidota | *Faecalibacter sp. LW9* | WP_323674094.1 | WP_322969619.1 | WP_322971732.1 |
| Bacteroidota | *Ferruginibacter albus* | WP_224015377.1 | WP_224019142.1 | WP_224018449.1 |
| Bacteroidota | *Ferruginibacter lapsinanis* | WP_229761199.1 | WP_229759758.1 | WP_229760830.1 |
| Bacteroidota | *Ferruginibacter sp.* | WP_301920859.1 | WP_301932169.1 | WP_301928244.1 |
| Bacteroidota | *Ferruginibacter yonginensis* | WP_379710127.1 | WP_379706315.1 | WP_379706877.1 |
| Bacteroidota | *Fibrella aestuarina* | WP_015329526.1 | WP_015331947.1 | WP_015329586.1 |
| Bacteroidota | *Fibrella aquatilis* | WP_207338346.1 | WP_207337558.1 | WP_207336838.1 |
| Bacteroidota | *Fibrella forsythiae* | WP_207329357.1 | WP_207327138.1 | WP_207329926.1 |
| Bacteroidota | *Fibrella rubiginis* | WP_207364873.1 | WP_207366683.1 | WP_207365617.1 |
| Bacteroidota | *Fibrella sp. GW2-5* | WP_370726049.1 | WP_370722175.1 | WP_370724314.1 |
| Bacteroidota | *Fibrisoma limi* | WP_009283656.1 | WP_009283245.1 | WP_009284943.1 |
| Bacteroidota | *Fibrisoma montanum* | WP_119666868.1 | WP_119667212.1 | WP_119666485.1 |
| Bacteroidota | *Fibrivirga algicola* | WP_085411010.1 | WP_166693286.1 | WP_166692858.1 |
| Bacteroidota | *Filimonas effusa* | WP_129005128.1 | WP_246022425.1 | WP_129001150.1 |
| Bacteroidota | *Filimonas lacunae* | WP_076379672.1 | WP_076378918.1 | WP_231940298.1 |
| Bacteroidota | *Filimonas zeae* | WP_188957660.1 | WP_188951939.1 | WP_188952473.1 |
| Bacteroidota | *Flagellimonas aequoris* | WP_119638374.1 | WP_119641752.1 | WP_119639284.1 |

| | | | | |
|---|---|---|---|---|
| Bacteroidota | *Flagellimonas algicola* | WP_138832362.1 | WP_138832555.1 | WP_138831947.1 |
| Bacteroidota | *Flagellimonas allohymeniacidonis* | WP_130608790.1 | WP_130609031.1 | WP_130607915.1 |
| Bacteroidota | *Flagellimonas alvinocaridis* | WP_136567100.1 | WP_136567470.1 | WP_136566847.1 |
| Bacteroidota | *Flagellimonas amoyensis* | WP_108424753.1 | WP_108424665.1 | WP_108425404.1 |
| Bacteroidota | *Flagellimonas aquimarina* | WP_109664998.1 | WP_109663516.1 | WP_109664479.1 |
| Bacteroidota | *Flagellimonas aurea* | WP_225604307.1 | WP_225604384.1 | WP_207036507.1 |
| Bacteroidota | *Flagellimonas baculiformis* | WP_318345083.1 | WP_318344797.1 | WP_318344658.1 |
| Bacteroidota | *Flagellimonas beolgyonensis* | WP_127019374.1 | WP_127019184.1 | WP_127020011.1 |
| Bacteroidota | *Flagellimonas crocea* | WP_318308088.1 | WP_318308224.1 | WP_318312414.1 |
| Bacteroidota | *Flagellimonas eckloniae* | WP_055393033.1 | WP_055392895.1 | WP_055393334.1 |
| Bacteroidota | *Flagellimonas flava* | WP_073182112.1 | WP_073176911.1 | WP_073181092.1 |
| Bacteroidota | *Flagellimonas halotolerans* | WP_326278964.1 | WP_293162775.1 | WP_326279016.1 |
| Bacteroidota | *Flagellimonas hymeniacidonis* | WP_147744767.1 | WP_147744681.1 | WP_147745025.1 |
| Bacteroidota | *Flagellimonas iocasae* | WP_379829643.1 | WP_379829557.1 | WP_379830012.1 |
| Bacteroidota | *Flagellimonas lutaonensis* | WP_045802637.1 | WP_045802737.1 | WP_045802577.1 |
| Bacteroidota | *Flagellimonas lutimaris* | WP_119609232.1 | WP_119609392.1 | WP_414459438.1 |
| Bacteroidota | *Flagellimonas marinaquae* | WP_338194641.1 | WP_127140727.1 | WP_338195901.1 |
| Bacteroidota | *Flagellimonas maritima* | WP_112379135.1 | WP_112379253.1 | WP_112378974.1 |
| Bacteroidota | *Flagellimonas meishanensis* | WP_222983916.1 | WP_222983994.1 | WP_222983733.1 |
| Bacteroidota | *Flagellimonas meridianipacifica* | WP_106147075.1 | WP_106146871.1 | WP_106147422.1 |
| Bacteroidota | *Flagellimonas myxillae* | WP_249969211.1 | WP_249969116.1 | WP_249969413.1 |
| Bacteroidota | *Flagellimonas nanhaiensis* | WP_116185870.1 | WP_116183645.1 | WP_116185290.1 |
| Bacteroidota | *Flagellimonas oceanensis* | WP_421824386.1 | WP_127138859.1 | WP_421824618.1 |
| Bacteroidota | *Flagellimonas oceani* | WP_166247842.1 | WP_166247919.1 | WP_166247384.1 |
| Bacteroidota | *Flagellimonas ochracea* | WP_166522626.1 | WP_166522542.1 | WP_166524569.1 |
| Bacteroidota | *Flagellimonas okinawensis* | WP_275650265.1 | WP_275650364.1 | WP_275650583.1 |
| Bacteroidota | *Flagellimonas olearia* | WP_129654481.1 | WP_152130461.1 | WP_152129927.1 |
| Bacteroidota | *Flagellimonas onchidii* | WP_136465462.1 | WP_136465354.1 | WP_136468868.1 |

| | | | | |
|---|---|---|---|---|
| Bacteroidota | *Flagellimonas pacifica* | WP_097045101.1 | WP_097045209.1 | WP_097044884.1 |
| Bacteroidota | *Flagellimonas pelagia* | WP_119646572.1 | WP_119646449.1 | WP_119648244.1 |
| Bacteroidota | *Flagellimonas profundi* | WP_207026198.1 | WP_207026359.1 | WP_207025555.1 |
| Bacteroidota | *Flagellimonas ruestringensis* | WP_014033624.1 | WP_014034266.1 | WP_224836465.1 |
| Bacteroidota | *Flagellimonas sediminis* | WP_163633580.1 | WP_163633770.1 | WP_163636359.1 |
| Bacteroidota | *Flagellimonas sp.* | WP_420600871.1 | WP_418499172.1 | WP_420322096.1 |
| Bacteroidota | *Flagellimonas spongiicola* | WP_249658705.1 | WP_249656751.1 | WP_249658493.1 |
| Bacteroidota | *Flagellimonas yonaguniensis* | WP_275616580.1 | WP_275615732.1 | WP_275616724.1 |
| Bacteroidota | *Flagellimonas zhangzhouensis* | WP_090298686.1 | WP_090297115.1 | WP_090298155.1 |
| Bacteroidota | *Flammeovirga kamogawensis* | WP_144073762.1 | WP_144073022.1 | WP_144074686.1 |
| Bacteroidota | *Flammeovirga pacifica* | WP_044227781.1 | WP_044219798.1 | WP_044228724.1 |
| Bacteroidota | *Flammeovirga pectinis* | WP_126616529.1 | WP_126610898.1 | WP_126614491.1 |
| Bacteroidota | *Flammeovirga sp. MY04* | WP_066210782.1 | WP_066212474.1 | WP_066205624.1 |
| Bacteroidota | *Flammeovirga yaeyamensis* | WP_169664813.1 | WP_169662846.1 | WP_169665136.1 |
| Bacteroidota | *Flaviaesturariibacter amylovorans* | WP_345252897.1 | WP_345255691.1 | WP_345253761.1 |
| Bacteroidota | *Flaviaesturariibacter aridisoli* | WP_131850298.1 | WP_131850986.1 | WP_131853005.1 |
| Bacteroidota | *Flaviaesturariibacter flavus* | WP_131449966.1 | WP_243647739.1 | WP_131449650.1 |
| Bacteroidota | *Flavicella marina* | WP_152286514.1 | WP_152286678.1 | WP_226913082.1 |
| Bacteroidota | *Flavicella sediminum* | WP_139959103.1 | WP_139959517.1 | WP_139959645.1 |
| Bacteroidota | *Flavicella sp.* | WP_336891546.1 | WP_300326719.1 | WP_336891758.1 |
| Bacteroidota | *Flavihumibacter cheonanensis* | WP_239602470.1 | WP_239595074.1 | WP_239595392.1 |
| Bacteroidota | *Flavihumibacter fluminis* | WP_234867994.1 | WP_234865708.1 | WP_234865854.1 |
| Bacteroidota | *Flavihumibacter fluvii* | WP_214458631.1 | WP_214457879.1 | WP_214457758.1 |
| Bacteroidota | *Flavihumibacter petaseus* | WP_046367457.1 | WP_046370235.1 | WP_046370374.1 |
| Bacteroidota | *Flavihumibacter profundi* | WP_224117810.1 | WP_224115123.1 | WP_224115252.1 |
| Bacteroidota | *Flavihumibacter rivuli* | WP_214447400.1 | WP_214449400.1 | WP_239804314.1 |
| Bacteroidota | *Flavihumibacter solisilvae* | WP_039138289.1 | WP_039140031.1 | WP_039141623.1 |
| Bacteroidota | *Flavihumibacter sp.* | WP_332732135.1 | WP_332734476.1 | WP_332734753.1 |

| | | | | |
|---|---|---|---|---|
| Bacteroidota | *Flavihumibacter stibioxidans* | WP_187255864.1 | WP_187255311.1 | WP_187255180.1 |
| Bacteroidota | *Flavilitoribacter nigricans* | WP_099155523.1 | WP_099155433.1 | WP_099148935.1 |
| Bacteroidota | *Flavimarina sp. Hel_I_48* | WP_031427791.1 | WP_031425752.1 | WP_031427820.1 |
| Bacteroidota | *Flavipsychrobacter stenotrophus* | WP_105041010.1 | WP_105040190.1 | WP_105037087.1 |
| Bacteroidota | *Flaviramulus aquimarinus* | WP_345272414.1 | WP_345274344.1 | WP_345272394.1 |
| Bacteroidota | *Flaviramulus basaltis* | WP_072401320.1 | WP_171946619.1 | WP_072402259.1 |
| Bacteroidota | *Flaviramulus multivorans* | WP_237230794.1 | WP_237232560.1 | WP_237230925.1 |
| Bacteroidota | *Flaviramulus sp. BrNp1-15* | WP_238673743.1 | WP_238676151.1 | WP_238673896.1 |
| Bacteroidota | *Flavisolibacter ginsengisoli* | WP_072835562.1 | WP_072835367.1 | WP_072835393.1 |
| Bacteroidota | *Flavisolibacter ginsenosidimutans* | WP_146785371.1 | WP_146781711.1 | WP_146781592.1 |
| Bacteroidota | *Flavisolibacter nicotianae* | WP_121354209.1 | WP_121356021.1 | WP_121356571.1 |
| Bacteroidota | *Flavisolibacter tropicus* | WP_066404609.1 | WP_066401941.1 | WP_066402027.1 |
| Bacteroidota | *Flavitalea sp. BT771* | WP_303588332.1 | WP_303587254.1 | WP_303591133.1 |
| Bacteroidota | *Flavivirga abyssicola* | WP_303317827.1 | WP_303315654.1 | WP_303319200.1 |
| Bacteroidota | *Flavivirga algicola* | WP_169673212.1 | WP_169669341.1 | WP_169676338.1 |
| Bacteroidota | *Flavivirga amylovorans* | WP_303282627.1 | WP_303283341.1 | WP_303283293.1 |
| Bacteroidota | *Flavivirga aquatica* | WP_069830633.1 | WP_069831205.1 | WP_069830554.1 |
| Bacteroidota | *Flavivirga aquimarina* | WP_303279803.1 | WP_303277252.1 | WP_303276492.1 |
| Bacteroidota | *Flavivirga eckloniae* | WP_102758028.1 | WP_102754615.1 | WP_102754426.1 |
| Bacteroidota | *Flavivirga jejuensis* | WP_303303193.1 | WP_303301116.1 | WP_303302142.1 |
| Bacteroidota | *Flavivirga rizhaonensis* | WP_135878189.1 | WP_135874426.1 | WP_135874598.1 |
| Bacteroidota | *Flavivirga sp. 57AJ16* | WP_274184694.1 | WP_274186157.1 | WP_274185721.1 |
| Bacteroidota | *Flavivirga spongiicola* | WP_303307549.1 | WP_303308519.1 | WP_303308337.1 |
| Bacteroidota | *Flavobacterium acetivorans* | WP_230062547.1 | WP_230061645.1 | WP_230061457.1 |
| Bacteroidota | *Flavobacterium aciduliphilum* | WP_112111771.1 | WP_112113984.1 | WP_112111999.1 |
| Bacteroidota | *Flavobacterium adhaerens* | WP_348800068.1 | WP_348799652.1 | WP_348800379.1 |
| Bacteroidota | *Flavobacterium aerium* | WP_306350197.1 | WP_306352814.1 | WP_306352934.1 |
| Bacteroidota | *Flavobacterium aestivum* | WP_268846490.1 | WP_268846139.1 | WP_281322463.1 |

| | | | | |
|---|---|---|---|---|
| Bacteroidota | *Flavobacterium agri* | WP_176004808.1 | WP_176004297.1 | WP_176006078.1 |
| Bacteroidota | *Flavobacterium agrisoli* | WP_200104139.1 | WP_200104943.1 | WP_200105035.1 |
| Bacteroidota | *Flavobacterium ajazii* | WP_163410375.1 | WP_163408156.1 | WP_163408377.1 |
| Bacteroidota | *Flavobacterium akiainvivens* | WP_054406461.1 | WP_054409670.1 | WP_054409766.1 |
| Bacteroidota | *Flavobacterium album* | WP_108778230.1 | WP_108778512.1 | WP_108779563.1 |
| Bacteroidota | *Flavobacterium algicola* | WP_239865649.1 | WP_239867780.1 | WP_239866461.1 |
| Bacteroidota | *Flavobacterium alkalisoli* | WP_147583880.1 | WP_147583557.1 | WP_417351809.1 |
| Bacteroidota | *Flavobacterium alvei* | WP_103805007.1 | WP_304197511.1 | WP_304201011.1 |
| Bacteroidota | *Flavobacterium ammonificans* | WP_229324657.1 | WP_229323846.1 | WP_229325007.1 |
| Bacteroidota | *Flavobacterium ammoniigenes* | WP_229316756.1 | WP_229316411.1 | WP_229316318.1 |
| Bacteroidota | *Flavobacterium amnicola* | WP_129435686.1 | WP_129436520.1 | WP_129436066.1 |
| Bacteroidota | *Flavobacterium amniphilum* | WP_250595108.1 | WP_250597894.1 | WP_250598363.1 |
| Bacteroidota | *Flavobacterium anhuiense* | WP_091133020.1 | WP_091133221.1 | WP_129745120.1 |
| Bacteroidota | *Flavobacterium antarcticum* | WP_022827233.1 | WP_022828706.1 | WP_022828531.1 |
| Bacteroidota | *Flavobacterium aquariorum* | WP_111410466.1 | WP_111409112.1 | WP_111409004.1 |
| Bacteroidota | *Flavobacterium aquaticum* | WP_111567236.1 | WP_111565803.1 | WP_111565483.1 |
| Bacteroidota | *Flavobacterium aquatile* | WP_035127441.1 | WP_035126550.1 | WP_035126315.1 |
| Bacteroidota | *Flavobacterium aquicola* | WP_115812379.1 | WP_115809349.1 | WP_115810936.1 |
| Bacteroidota | *Flavobacterium aquidurense* | WP_055096489.1 | WP_055096091.1 | WP_055095569.1 |
| Bacteroidota | *Flavobacterium aquiphilum* | WP_281228934.1 | WP_281226080.1 | WP_281228415.1 |
| Bacteroidota | *Flavobacterium araucananum* | WP_089480192.1 | WP_089481795.1 | WP_089478665.1 |
| Bacteroidota | *Flavobacterium arcticum* | WP_114678538.1 | WP_114678788.1 | WP_114676491.1 |
| Bacteroidota | *Flavobacterium ardleyense* | WP_318640429.1 | WP_379804239.1 | WP_318639709.1 |
| Bacteroidota | *Flavobacterium arsenatis* | WP_310025109.1 | WP_310028268.1 | WP_310023609.1 |
| Bacteroidota | *Flavobacterium artemisiae* | WP_379816507.1 | WP_379817049.1 | WP_379816923.1 |
| Bacteroidota | *Flavobacterium arundinis* | WP_341695324.1 | WP_341696461.1 | WP_341696278.1 |
| Bacteroidota | *Flavobacterium aurantiibacter* | WP_094485081.1 | WP_094486620.1 | WP_094487510.1 |
| Bacteroidota | *Flavobacterium aureirubrum* | WP_342695397.1 | WP_342696707.1 | WP_342696298.1 |

| | | | | |
|---|---|---|---|---|
| Bacteroidota | *Flavobacterium azizsancarii* | WP_271336029.1 | WP_271338417.1 | WP_271334397.1 |
| Bacteroidota | *Flavobacterium azooxidireducens* | WP_248434158.1 | WP_248433393.1 | WP_248433145.1 |
| Bacteroidota | *Flavobacterium beibuense* | WP_035133659.1 | WP_129750265.1 | WP_417366338.1 |
| Bacteroidota | *Flavobacterium bernardetii* | WP_166125315.1 | WP_166126073.1 | WP_166126115.1 |
| Bacteroidota | *Flavobacterium bizetiae* | WP_263759732.1 | WP_173971207.1 | WP_395368299.1 |
| Bacteroidota | *Flavobacterium branchiarum* | WP_290262884.1 | WP_290262510.1 | WP_290266136.1 |
| Bacteroidota | *Flavobacterium branchiicola* | WP_213258301.1 | WP_213258164.1 | WP_213257900.1 |
| Bacteroidota | *Flavobacterium branchiophilum* | WP_014083711.1 | WP_014083730.1 | WP_089081437.1 |
| Bacteroidota | *Flavobacterium buctense* | WP_341432331.1 | WP_187660755.1 | WP_187660569.1 |
| Bacteroidota | *Flavobacterium caeni* | WP_091141017.1 | WP_091140782.1 | WP_091139974.1 |
| Bacteroidota | *Flavobacterium calami* | WP_341690908.1 | WP_341692856.1 | WP_341692452.1 |
| Bacteroidota | *Flavobacterium capsici* | WP_313321563.1 | WP_313323035.1 | WP_313325783.1 |
| Bacteroidota | *Flavobacterium caseinilyticum* | WP_131908690.1 | WP_131910707.1 | WP_131909271.1 |
| Bacteroidota | *Flavobacterium cauense* | WP_023571266.1 | WP_023571697.1 | WP_023570087.1 |
| Bacteroidota | *Flavobacterium celericrescens* | WP_166235729.1 | WP_166236799.1 | WP_166235918.1 |
| Bacteroidota | *Flavobacterium cellulosilyticum* | WP_132006492.1 | WP_132006986.1 | WP_132004434.1 |
| Bacteroidota | *Flavobacterium cerinum* | WP_256550675.1 | WP_128391093.1 | WP_128388847.1 |
| Bacteroidota | *Flavobacterium channae* | WP_231834529.1 | WP_231836887.1 | WP_231836528.1 |
| Bacteroidota | *Flavobacterium cheonanense* | WP_344816363.1 | WP_344816714.1 | WP_344816211.1 |
| Bacteroidota | *Flavobacterium cheongpyeongense* | WP_110307633.1 | WP_110304959.1 | WP_110304752.1 |
| Bacteroidota | *Flavobacterium chilense* | WP_068842693.1 | WP_068842835.1 | WP_068844503.1 |
| Bacteroidota | *Flavobacterium chuncheonense* | WP_379810082.1 | WP_379811473.1 | WP_379811128.1 |
| Bacteroidota | *Flavobacterium chungnamense* | WP_345094576.1 | WP_345095656.1 | WP_345093744.1 |
| Bacteroidota | *Flavobacterium circumlabens* | WP_348080008.1 | WP_132034782.1 | WP_348078518.1 |
| Bacteroidota | *Flavobacterium collinsii* | WP_173968038.1 | WP_263361526.1 | WP_173966104.1 |
| Bacteroidota | *Flavobacterium columnare* | WP_014165519.1 | WP_107317836.1 | WP_324712696.1 |
| Bacteroidota | *Flavobacterium coralii* | WP_223052048.1 | WP_348679228.1 | WP_223053151.1 |
| Bacteroidota | *Flavobacterium crassostreae* | WP_066335673.1 | WP_066334792.1 | WP_066334607.1 |

| | | | | |
|---|---|---|---|---|
| Bacteroidota | *Flavobacterium croceum* | WP_103725045.1 | WP_103726256.1 | WP_103726419.1 |
| Bacteroidota | *Flavobacterium crocinum* | WP_109194175.1 | WP_109194355.1 | WP_109194604.1 |
| Bacteroidota | *Flavobacterium cucumis* | WP_073584520.1 | WP_073582549.1 | WP_073580360.1 |
| Bacteroidota | *Flavobacterium cupreum* | WP_127338736.1 | WP_127338606.1 | WP_127339709.1 |
| Bacteroidota | *Flavobacterium cupriresistens* | WP_230002453.1 | WP_230002663.1 | WP_230002945.1 |
| Bacteroidota | *Flavobacterium cutihirudinis* | WP_115888499.1 | WP_115888668.1 | WP_115888903.1 |
| Bacteroidota | *Flavobacterium cyanobacteriorum* | WP_094416373.1 | WP_094412463.1 | WP_094417039.1 |
| Bacteroidota | *Flavobacterium cyclinae* | WP_231843804.1 | WP_231841638.1 | WP_231841964.1 |
| Bacteroidota | *Flavobacterium daejeonense* | WP_026714153.1 | WP_026715500.1 | WP_026714113.1 |
| Bacteroidota | *Flavobacterium daemonense* | WP_144339228.1 | WP_144337841.1 | WP_144338222.1 |
| Bacteroidota | *Flavobacterium dankookense* | WP_133531808.1 | WP_133531489.1 | WP_133533978.1 |
| Bacteroidota | *Flavobacterium dauae* | WP_129758015.1 | WP_129757852.1 | WP_129757042.1 |
| Bacteroidota | *Flavobacterium defluvii* | WP_073416481.1 | WP_073416641.1 | WP_073417082.1 |
| Bacteroidota | *Flavobacterium degerlachei* | WP_091434817.1 | WP_091429243.1 | WP_091434165.1 |
| Bacteroidota | *Flavobacterium denitrificans* | WP_026730144.1 | WP_026730335.1 | WP_026728034.1 |
| Bacteroidota | *Flavobacterium difficile* | WP_166077145.1 | WP_166078006.1 | WP_166077778.1 |
| Bacteroidota | *Flavobacterium eburneipallidum* | WP_281336077.1 | WP_269225176.1 | WP_281337225.1 |
| Bacteroidota | *Flavobacterium enshiense* | WP_367230005.1 | WP_367230350.1 | WP_367230791.1 |
| Bacteroidota | *Flavobacterium erciyesense* | WP_210788498.1 | WP_210788461.1 | WP_210788629.1 |
| Bacteroidota | *Flavobacterium facile* | WP_329804786.1 | WP_329806052.1 | WP_329806076.1 |
| Bacteroidota | *Flavobacterium faecale* | WP_108741200.1 | WP_108740786.1 | WP_108740713.1 |
| Bacteroidota | *Flavobacterium filum* | WP_276979020.1 | WP_026709932.1 | WP_026711516.1 |
| Bacteroidota | *Flavobacterium flavigenum* | WP_269236635.1 | WP_269236927.1 | WP_269237110.1 |
| Bacteroidota | *Flavobacterium flavipallidum* | WP_341700249.1 | WP_341700978.1 | WP_341701690.1 |
| Bacteroidota | *Flavobacterium flevense* | WP_073245585.1 | WP_073245546.1 | WP_073244749.1 |
| Bacteroidota | *Flavobacterium fluviale* | WP_113679386.1 | WP_113680010.1 | WP_113679743.1 |
| Bacteroidota | *Flavobacterium fluviatile* | WP_163399805.1 | WP_163401290.1 | WP_163400519.1 |
| Bacteroidota | *Flavobacterium fluvii* | WP_073370950.1 | WP_073371333.1 | WP_073370273.1 |

| | | | | |
|---|---|---|---|---|
| Bacteroidota | *Flavobacterium foetidum* | WP_135223692.1 | WP_135220578.1 | WP_135220849.1 |
| Bacteroidota | *Flavobacterium fontis* | WP_073362329.1 | WP_073365076.1 | WP_073362776.1 |
| Bacteroidota | *Flavobacterium fragile* | WP_250581805.1 | WP_250582693.1 | WP_250582588.1 |
| Bacteroidota | *Flavobacterium franklandianum* | WP_143391881.1 | WP_143391434.1 | WP_143391604.1 |
| Bacteroidota | *Flavobacterium frigidarium* | WP_371570782.1 | WP_339654805.1 | WP_026706692.1 |
| Bacteroidota | *Flavobacterium frigidimaris* | WP_074658449.1 | WP_074658704.1 | WP_074659117.1 |
| Bacteroidota | *Flavobacterium frigoris* | WP_074722389.1 | WP_074723038.1 | WP_007138112.1 |
| Bacteroidota | *Flavobacterium frigoritolerans* | WP_264287449.1 | WP_264287159.1 | WP_264285323.1 |
| Bacteroidota | *Flavobacterium fructosi* | WP_379857536.1 | WP_379858922.1 | WP_379857307.1 |
| Bacteroidota | *Flavobacterium fryxellicola* | WP_066078745.1 | WP_066078551.1 | WP_066078063.1 |
| Bacteroidota | *Flavobacterium galactosidilyticum* | WP_229942020.1 | WP_229941185.1 | WP_229943974.1 |
| Bacteroidota | *Flavobacterium gelatinilyticum* | WP_281232719.1 | WP_281232539.1 | WP_281232268.1 |
| Bacteroidota | *Flavobacterium gelidilacus* | WP_026715688.1 | WP_026716442.1 | WP_026716198.1 |
| Bacteroidota | *Flavobacterium gillisiae* | WP_091086427.1 | WP_091093031.1 | WP_091090389.1 |
| Bacteroidota | *Flavobacterium gilvum* | WP_035637836.1 | WP_035638905.1 | WP_035640009.1 |
| Bacteroidota | *Flavobacterium ginsengiterrae* | WP_345146172.1 | WP_345145973.1 | WP_345145719.1 |
| Bacteroidota | *Flavobacterium ginsenosidimutans* | WP_111285252.1 | WP_111285576.1 | WP_111289076.1 |
| Bacteroidota | *Flavobacterium glaciei* | WP_114755205.1 | WP_114755304.1 | WP_114753358.1 |
| Bacteroidota | *Flavobacterium glycines* | WP_066329059.1 | WP_066329643.1 | WP_066324645.1 |
| Bacteroidota | *Flavobacterium granuli* | WP_072937916.1 | WP_310002898.1 | WP_072941334.1 |
| Bacteroidota | *Flavobacterium gyeonganense* | WP_278010294.1 | WP_278010560.1 | WP_278010745.1 |
| Bacteroidota | *Flavobacterium haoranii* | WP_072783792.1 | WP_072785678.1 | WP_072785004.1 |
| Bacteroidota | *Flavobacterium hercynium* | WP_089051503.1 | WP_089050641.1 | WP_089051567.1 |
| Bacteroidota | *Flavobacterium hibernum* | WP_041518361.1 | WP_041515948.1 | WP_041517094.1 |
| Bacteroidota | *Flavobacterium hibisci* | WP_223680455.1 | WP_223683385.1 | WP_223681683.1 |
| Bacteroidota | *Flavobacterium hiemivividum* | WP_132111183.1 | WP_132111004.1 | WP_132110648.1 |
| Bacteroidota | *Flavobacterium humi* | WP_135524654.1 | WP_135526473.1 | WP_135526235.1 |
| Bacteroidota | *Flavobacterium humidisoli* | WP_248727515.1 | WP_248727640.1 | WP_248727894.1 |

| | | | | |
|---|---|---|---|---|
| Bacteroidota | *Flavobacterium hungaricum* | WP_194139755.1 | WP_194139922.1 | WP_194140192.1 |
| Bacteroidota | *Flavobacterium hydatis* | WP_035620475.1 | WP_035627907.1 | WP_035625949.1 |
| Bacteroidota | *Flavobacterium hydrocarbonoxydans* | WP_160373147.1 | WP_160376079.1 | WP_160375866.1 |
| Bacteroidota | *Flavobacterium hydrophilum* | WP_110345293.1 | WP_110344961.1 | WP_110344757.1 |
| Bacteroidota | *Flavobacterium ichthyis* | WP_166536024.1 | WP_166535779.1 | WP_166537713.1 |
| Bacteroidota | *Flavobacterium jejuense* | WP_140963139.1 | WP_140959233.1 | WP_140960741.1 |
| Bacteroidota | *Flavobacterium johnsoniae* | WP_073408372.1 | WP_289877969.1 | WP_289878288.1 |
| Bacteroidota | *Flavobacterium kayseriense* | WP_187011015.1 | WP_187011005.1 | WP_187010795.1 |
| Bacteroidota | *Flavobacterium kingsejongi* | WP_108737943.1 | WP_108737494.1 | WP_108738362.1 |
| Bacteroidota | *Flavobacterium lacisediminis* | WP_264369442.1 | WP_264369666.1 | WP_264368558.1 |
| Bacteroidota | *Flavobacterium lacus* | WP_112085796.1 | WP_112085450.1 | WP_112086584.1 |
| Bacteroidota | *Flavobacterium lacustre* | WP_269684458.1 | WP_269685141.1 | WP_269684167.1 |
| Bacteroidota | *Flavobacterium laiguense* | WP_116763267.1 | WP_116763918.1 | WP_116760445.1 |
| Bacteroidota | *Flavobacterium limi* | WP_163396595.1 | WP_163394177.1 | WP_163394387.1 |
| Bacteroidota | *Flavobacterium limicola* | WP_121365842.1 | WP_121365603.1 | WP_121365642.1 |
| Bacteroidota | *Flavobacterium limnophilum* | WP_281299364.1 | WP_281299001.1 | WP_281297214.1 |
| Bacteroidota | *Flavobacterium limnosediminis* | WP_023579171.1 | WP_023579392.1 | WP_023580327.1 |
| Bacteroidota | *Flavobacterium litorale* | WP_220639998.1 | WP_220640443.1 | WP_220640681.1 |
| Bacteroidota | *Flavobacterium longum* | WP_415580331.1 | WP_415580030.1 | WP_415581162.1 |
| Bacteroidota | *Flavobacterium lotistagni* | WP_166335680.1 | WP_166336123.1 | WP_166334538.1 |
| Bacteroidota | *Flavobacterium luminosum* | WP_250592150.1 | WP_250591819.1 | WP_250591576.1 |
| Bacteroidota | *Flavobacterium luteum* | WP_151108338.1 | WP_151106222.1 | WP_151107915.1 |
| Bacteroidota | *Flavobacterium macacae* | WP_125011741.1 | WP_125013480.1 | WP_125012357.1 |
| Bacteroidota | *Flavobacterium magnesitis* | WP_373391948.1 | WP_373393194.1 | WP_373390870.1 |
| Bacteroidota | *Flavobacterium magnum* | WP_108371837.1 | WP_108373551.1 | WP_108370611.1 |
| Bacteroidota | *Flavobacterium maritimum* | WP_348812361.1 | WP_348811647.1 | WP_348811765.1 |
| Bacteroidota | *Flavobacterium microcysteis* | WP_140000093.1 | WP_140001393.1 | WP_140002987.1 |
| Bacteroidota | *Flavobacterium micromati* | WP_073021752.1 | WP_073021956.1 | WP_073021419.1 |

| | | | | |
|---|---|---|---|---|
| Bacteroidota | *Flavobacterium muglaense* | WP_187020500.1 | WP_187019474.1 | WP_187016867.1 |
| Bacteroidota | *Flavobacterium myungsuense* | WP_379759358.1 | WP_379758019.1 | WP_379755506.1 |
| Bacteroidota | *Flavobacterium nackdongense* | WP_133277831.1 | WP_133275103.1 | WP_133275856.1 |
| Bacteroidota | *Flavobacterium nakdongensis* | WP_309533082.1 | WP_309532800.1 | WP_309531613.1 |
| Bacteroidota | *Flavobacterium nitratireducens* | WP_278035832.1 | WP_278035364.1 | WP_278035249.1 |
| Bacteroidota | *Flavobacterium noncentrifugens* | WP_091396656.1 | WP_091398032.1 | WP_091393211.1 |
| Bacteroidota | *Flavobacterium odoriferum* | WP_262318657.1 | WP_262318477.1 | WP_262317886.1 |
| Bacteroidota | *Flavobacterium okayamense* | WP_221258181.1 | WP_221258373.1 | WP_221257683.1 |
| Bacteroidota | *Flavobacterium olei* | WP_347049574.1 | WP_347049694.1 | WP_347049942.1 |
| Bacteroidota | *Flavobacterium omnivorum* | WP_091258353.1 | WP_091257743.1 | WP_091259028.1 |
| Bacteroidota | *Flavobacterium oreochromis* | WP_088397939.1 | WP_088399325.1 | WP_088397908.1 |
| Bacteroidota | *Flavobacterium orientale* | WP_188361792.1 | WP_188361733.1 | WP_188361861.1 |
| Bacteroidota | *Flavobacterium ovatum* | WP_366183810.1 | WP_366183988.1 | WP_366184218.1 |
| Bacteroidota | *Flavobacterium pallidum* | WP_108904208.1 | WP_108904408.1 | WP_108903708.1 |
| Bacteroidota | *Flavobacterium palustre* | WP_188493166.1 | WP_188493705.1 | WP_188493752.1 |
| Bacteroidota | *Flavobacterium paronense* | WP_290285180.1 | WP_290285908.1 | WP_290284915.1 |
| Bacteroidota | *Flavobacterium pectinovorum* | WP_140511595.1 | WP_243232481.1 | WP_073395410.1 |
| Bacteroidota | *Flavobacterium petrolei* | WP_113666649.1 | WP_395380808.1 | WP_395380779.1 |
| Bacteroidota | *Flavobacterium phragmitis* | WP_091492985.1 | WP_091491738.1 | WP_091492158.1 |
| Bacteroidota | *Flavobacterium phycosphaerae* | WP_162127750.1 | WP_162126086.1 | WP_162128342.1 |
| Bacteroidota | *Flavobacterium piscinae* | WP_129464813.1 | WP_129464344.1 | WP_129464983.1 |
| Bacteroidota | *Flavobacterium pisciphilum* | WP_229987522.1 | WP_229987734.1 | WP_229990525.1 |
| Bacteroidota | *Flavobacterium piscis* | WP_310284124.1 | WP_065450509.1 | WP_310281901.1 |
| Bacteroidota | *Flavobacterium piscisymbiosum* | WP_230039643.1 | WP_230039970.1 | WP_230040703.1 |
| Bacteroidota | *Flavobacterium plantiphilum* | WP_408078841.1 | WP_408078801.1 | WP_408081465.1 |
| Bacteroidota | *Flavobacterium pokkalii* | WP_188220226.1 | WP_055092636.1 | WP_188221459.1 |
| Bacteroidota | *Flavobacterium polysaccharolyticum* | WP_342692940.1 | WP_342690826.1 | WP_342691523.1 |
| Bacteroidota | *Flavobacterium ponti* | WP_379740641.1 | WP_379741904.1 | WP_379743091.1 |

| | | | | |
|---|---|---|---|---|
| Bacteroidota | *Flavobacterium potami* | WP_223704457.1 | WP_223704287.1 | WP_223704074.1 |
| Bacteroidota | *Flavobacterium poyangense* | WP_166923811.1 | WP_166925746.1 | WP_166922132.1 |
| Bacteroidota | *Flavobacterium praedii* | WP_281239829.1 | WP_281240117.1 | WP_281239177.1 |
| Bacteroidota | *Flavobacterium procerum* | WP_379682565.1 | WP_379682729.1 | WP_379683027.1 |
| Bacteroidota | *Flavobacterium profundi* | WP_140996821.1 | WP_140999128.1 | WP_140998869.1 |
| Bacteroidota | *Flavobacterium proteolyticum* | WP_194095444.1 | WP_194095749.1 | WP_194094760.1 |
| Bacteroidota | *Flavobacterium psychraquaticum* | WP_322549922.1 | WP_322549451.1 | WP_322550351.1 |
| Bacteroidota | *Flavobacterium psychrophilum* | WP_011962921.1 | WP_011962816.1 | WP_203095712.1 |
| Bacteroidota | *Flavobacterium psychroterrae* | WP_213298471.1 | WP_213298967.1 | WP_213305312.1 |
| Bacteroidota | *Flavobacterium psychrotolerans* | WP_116725783.1 | WP_116723503.1 | WP_116723560.1 |
| Bacteroidota | *Flavobacterium psychrotrophum* | WP_116788075.1 | WP_116788784.1 | WP_116789352.1 |
| Bacteroidota | *Flavobacterium pygoscelis* | WP_248427562.1 | WP_248429168.1 | WP_248427783.1 |
| Bacteroidota | *Flavobacterium qiangtangense* | WP_379790684.1 | WP_379791238.1 | WP_379792899.1 |
| Bacteroidota | *Flavobacterium rakeshii* | WP_157483142.1 | WP_157481072.1 | WP_157483317.1 |
| Bacteroidota | *Flavobacterium reichenbachii* | WP_035683155.1 | WP_035682855.1 | WP_035690269.1 |
| Bacteroidota | *Flavobacterium restrictum* | WP_144255860.1 | WP_144254725.1 | WP_144257666.1 |
| Bacteroidota | *Flavobacterium rhizophilum* | WP_408075709.1 | WP_408073956.1 | WP_408074961.1 |
| Bacteroidota | *Flavobacterium rhizosphaerae* | WP_408083731.1 | WP_408083897.1 | WP_408083921.1 |
| Bacteroidota | *Flavobacterium rivuli* | WP_020214032.1 | WP_020212488.1 | WP_020212097.1 |
| Bacteroidota | *Flavobacterium rivulicola* | WP_171222509.1 | WP_171221989.1 | WP_171223169.1 |
| Bacteroidota | *Flavobacterium saccharophilum* | WP_072972051.1 | WP_072972649.1 | WP_072973028.1 |
| Bacteroidota | *Flavobacterium saliperosum* | WP_023576809.1 | WP_023576527.1 | WP_023577062.1 |
| Bacteroidota | *Flavobacterium sandaracinum* | WP_132066739.1 | WP_132064968.1 | WP_132064488.1 |
| Bacteroidota | *Flavobacterium sangjuense* | WP_136152510.1 | WP_136152911.1 | WP_136152372.1 |
| Bacteroidota | *Flavobacterium sasangense* | WP_026725803.1 | WP_026726094.1 | WP_026724613.1 |
| Bacteroidota | *Flavobacterium sedimenticola* | WP_283238790.1 | WP_283238321.1 | WP_283238626.1 |
| Bacteroidota | *Flavobacterium sediminis* | WP_109570272.1 | WP_109567860.1 | WP_109569801.1 |
| Bacteroidota | *Flavobacterium segetis* | WP_072992850.1 | WP_072993824.1 | WP_072987071.1 |

| | | | | |
|---|---|---|---|---|
| Bacteroidota | *Flavobacterium selenitireducens* | WP_191018474.1 | WP_191018872.1 | WP_191020079.1 |
| Bacteroidota | *Flavobacterium seoulense* | WP_035660257.1 | WP_035661625.1 | WP_035662614.1 |
| Bacteroidota | *Flavobacterium sharifuzzamanii* | WP_111426281.1 | WP_111426401.1 | WP_111426645.1 |
| Bacteroidota | *Flavobacterium shii* | WP_264206984.1 | WP_264204354.1 | WP_264206383.1 |
| Bacteroidota | *Flavobacterium silvaticum* | WP_169528127.1 | WP_169526441.1 | WP_169528369.1 |
| Bacteroidota | *Flavobacterium silvisoli* | WP_131476214.1 | WP_131476810.1 | WP_131475440.1 |
| Bacteroidota | *Flavobacterium sinopsychrotolerans* | WP_091168489.1 | WP_091166664.1 | WP_091166756.1 |
| Bacteroidota | *Flavobacterium soli* | WP_026705573.1 | WP_026705749.1 | WP_026704342.1 |
| Bacteroidota | *Flavobacterium solisilvae* | WP_169523750.1 | WP_169524792.1 | WP_169524984.1 |
| Bacteroidota | *Flavobacterium soyae* | WP_232682279.1 | WP_232679388.1 | WP_406844274.1 |
| Bacteroidota | *Flavobacterium soyangense* | WP_194311707.1 | WP_194312282.1 | WP_194310813.1 |
| Bacteroidota | *Flavobacterium sp.* | WP_371266015.1 | WP_396184720.1 | WP_324217648.1 |
| Bacteroidota | *Flavobacterium stagni* | WP_129461143.1 | WP_129462148.1 | WP_129460802.1 |
| Bacteroidota | *Flavobacterium suaedae* | WP_188621315.1 | WP_188620323.1 | WP_188620523.1 |
| Bacteroidota | *Flavobacterium subsaxonicum* | WP_026991950.1 | WP_026990105.1 | WP_026993033.1 |
| Bacteroidota | *Flavobacterium succinicans* | WP_064714826.1 | WP_024980033.1 | WP_024980258.1 |
| Bacteroidota | *Flavobacterium sufflavum* | WP_128196693.1 | WP_128194842.1 | WP_128194048.1 |
| Bacteroidota | *Flavobacterium suncheonense* | WP_026981611.1 | WP_026979946.1 | WP_026981151.1 |
| Bacteroidota | *Flavobacterium supellecticarium* | WP_136404178.1 | WP_136401869.1 | WP_136403284.1 |
| Bacteroidota | *Flavobacterium swingsii* | WP_091474524.1 | WP_091474723.1 | WP_091472898.1 |
| Bacteroidota | *Flavobacterium tagetis* | WP_202003963.1 | WP_202003572.1 | WP_202006294.1 |
| Bacteroidota | *Flavobacterium taihuense* | WP_219318647.1 | WP_219318958.1 | WP_219319062.1 |
| Bacteroidota | *Flavobacterium tegetincola* | WP_026978617.1 | WP_026977492.1 | WP_026978195.1 |
| Bacteroidota | *Flavobacterium terrae* | WP_073310928.1 | WP_073311178.1 | WP_073309335.1 |
| Bacteroidota | *Flavobacterium terrigena* | WP_091313686.1 | WP_091315883.1 | WP_091311805.1 |
| Bacteroidota | *Flavobacterium terrisoli* | WP_284652565.1 | WP_284653435.1 | WP_284652285.1 |
| Bacteroidota | *Flavobacterium tibetense* | WP_113988761.1 | WP_113989048.1 | WP_113988628.1 |
| Bacteroidota | *Flavobacterium tistrianum* | WP_111365996.1 | WP_111366193.1 | WP_111364009.1 |

| | | | | |
|---|---|---|---|---|
| Bacteroidota | *Flavobacterium tructae* | WP_374173388.1 | WP_374173498.1 | WP_070907458.1 |
| Bacteroidota | *Flavobacterium turcicum* | WP_166135359.1 | WP_166135473.1 | WP_166134952.1 |
| Bacteroidota | *Flavobacterium undicola* | WP_161640825.1 | WP_161643606.1 | WP_161643714.1 |
| Bacteroidota | *Flavobacterium urocaniciphilum* | WP_091465144.1 | WP_091468925.1 | WP_091468962.1 |
| Bacteroidota | *Flavobacterium urumqiense* | WP_104000151.1 | WP_104000107.1 | WP_103999846.1 |
| Bacteroidota | *Flavobacterium ustbae* | WP_125722837.1 | WP_125720253.1 | WP_125723234.1 |
| Bacteroidota | *Flavobacterium xanthum* | WP_073353088.1 | WP_073353397.1 | WP_073355174.1 |
| Bacteroidota | *Flavobacterium xinjiangense* | WP_073211262.1 | WP_073208242.1 | WP_073207460.1 |
| Bacteroidota | *Flavobacterium xueshanense* | WP_091206099.1 | WP_091207614.1 | WP_091203754.1 |
| Bacteroidota | *Flavobacterium xylosi* | WP_379854913.1 | WP_379853196.1 | WP_379854551.1 |
| Bacteroidota | *Flavobacterium yafengii* | WP_282728797.1 | WP_282715470.1 | WP_282730033.1 |
| Bacteroidota | *Flavobacterium zepuense* | WP_143372536.1 | WP_143375269.1 | WP_143373232.1 |
| Bacteroidota | *Flavobacterium zhairuonense* | WP_132988062.1 | WP_132989944.1 | WP_132989696.1 |
| Bacteroidota | *Flavobacterium zhouii* | WP_379852713.1 | WP_379851381.1 | WP_379849947.1 |
| Bacteroidota | *Flavobacterium zubiriense* | WP_373406039.1 | WP_373406299.1 | WP_373406703.1 |
| Bacteroidota | *Flectobacillus longus* | WP_283371972.1 | WP_283325847.1 | WP_283328046.1 |
| Bacteroidota | *Flectobacillus major* | WP_026997808.1 | WP_026994392.1 | WP_026997855.1 |
| Bacteroidota | *Flectobacillus roseus* | WP_283373285.1 | WP_283373442.1 | WP_283346348.1 |
| Bacteroidota | *Flectobacillus sp.* | WP_421820662.1 | WP_421819229.1 | WP_421818880.1 |
| Bacteroidota | *Flexibacter flexilis* | WP_091508130.1 | WP_091505765.1 | WP_091516603.1 |
| Bacteroidota | *Flexibacter sp. ATCC 35103* | WP_083692816.1 | WP_083692959.1 | WP_083693241.1 |
| Bacteroidota | *Flexithrix dorotheae* | WP_020532198.1 | WP_020528523.1 | WP_020533907.1 |
| Bacteroidota | *Fluviicola chungangensis* | WP_144331325.1 | WP_186280074.1 | WP_186280185.1 |
| Bacteroidota | *Fluviicola sp.* | WP_309280893.1 | WP_300359153.1 | WP_300659961.1 |
| Bacteroidota | *Fluviicola taffensis* | WP_013688266.1 | WP_341903383.1 | WP_013687422.1 |
| Bacteroidota | *Foetidibacter luteolus* | WP_153796776.1 | WP_153800762.1 | WP_153795974.1 |
| Bacteroidota | *Fontibacter flavus* | WP_382385908.1 | WP_382387759.1 | WP_382387480.1 |
| Bacteroidota | *Formosa agariphila* | WP_038529567.1 | WP_038528827.1 | WP_038527824.1 |

| | | | | |
|---|---|---|---|---|
| Bacteroidota | *Formosa algae* | WP_057779342.1 | WP_057778720.1 | WP_057778503.1 |
| Bacteroidota | *Formosa maritima* | WP_148452936.1 | WP_148454901.1 | WP_148455345.1 |
| Bacteroidota | *Formosa sediminum* | WP_143381701.1 | WP_143382029.1 | WP_143380981.1 |
| Bacteroidota | *Formosa sp. PL04* | WP_318498907.1 | WP_318498557.1 | WP_318501113.1 |
| Bacteroidota | *Formosa undariae* | WP_382381391.1 | WP_382381605.1 | WP_382382124.1 |
| Bacteroidota | *Frigoriflavimonas asaccharolytica* | WP_173780209.1 | WP_173780018.1 | WP_173779288.1 |
| Bacteroidota | *Fulvitalea axinellae* | WP_338392299.1 | WP_338391720.1 | WP_338391676.1 |
| Bacteroidota | *Fulvivirga aurantia* | WP_185152832.1 | WP_155210970.1 | WP_155212377.1 |
| Bacteroidota | *Fulvivirga imtechensis* | WP_009583631.1 | WP_009577485.1 | WP_009579199.1 |
| Bacteroidota | *Fulvivirga kasyanovii* | WP_155174132.1 | WP_155171568.1 | WP_343855452.1 |
| Bacteroidota | *Fulvivirga ligni* | WP_233771349.1 | WP_233770555.1 | WP_233771846.1 |
| Bacteroidota | *Fulvivirga lutea* | WP_205721864.1 | WP_205721098.1 | WP_205722411.1 |
| Bacteroidota | *Fulvivirga lutimaris* | WP_155187492.1 | WP_155185506.1 | WP_155188308.1 |
| Bacteroidota | *Fulvivirga marina* | WP_202854470.1 | WP_202855140.1 | WP_202854505.1 |
| Bacteroidota | *Fulvivirga maritima* | WP_233778822.1 | WP_233780993.1 | WP_233780020.1 |
| Bacteroidota | *Fulvivirga sedimenti* | WP_225698760.1 | WP_225696347.1 | WP_225699844.1 |
| Bacteroidota | *Fulvivirga sediminis* | WP_202244114.1 | WP_202243775.1 | WP_202244855.1 |
| Bacteroidota | *Fulvivirga sp.* | WP_350067676.1 | WP_350116307.1 | WP_350098834.1 |
| Bacteroidota | *Fulvivirga ulvae* | WP_233767761.1 | WP_233762065.1 | WP_233765613.1 |
| Bacteroidota | *Gaetbulibacter aestuarii* | WP_344741495.1 | WP_344741373.1 | WP_344742218.1 |
| Bacteroidota | *Gaetbulibacter aquiaggeris* | WP_395438398.1 | WP_395438696.1 | WP_395437480.1 |
| Bacteroidota | *Gaetbulibacter jejuensis* | WP_335976051.1 | WP_343799672.1 | WP_335974384.1 |
| Bacteroidota | *Gaetbulibacter saemankumensis* | WP_027136548.1 | WP_027136618.1 | WP_027136785.1 |
| Bacteroidota | *Gaetbulibacter sp. M235* | WP_394881643.1 | WP_394882940.1 | WP_394882766.1 |
| Bacteroidota | *Galbibacter marinus* | WP_008991857.1 | WP_008991011.1 | WP_008991389.1 |
| Bacteroidota | *Galbibacter mesophilus* | WP_202029560.1 | WP_202027634.1 | WP_202029510.1 |
| Bacteroidota | *Galbibacter orientalis* | WP_335588164.1 | WP_008614805.1 | WP_337900065.1 |
| Bacteroidota | *Galbibacter pacificus* | WP_277898839.1 | WP_277899085.1 | WP_277898211.1 |

| | | | | |
|---|---|---|---|---|
| Bacteroidota | *Galbibacter sp.* | WP_417360466.1 | WP_417364129.1 | WP_324301206.1 |
| Bacteroidota | *Gangjinia marincola* | WP_343762526.1 | WP_343764592.1 | WP_343764901.1 |
| Bacteroidota | *Gelatiniphilus marinus* | WP_388014987.1 | WP_388013439.1 | WP_388016878.1 |
| Bacteroidota | *Gelidibacter algens* | WP_066431358.1 | WP_066437546.1 | WP_111625946.1 |
| Bacteroidota | *Gelidibacter gilvus* | WP_129016144.1 | WP_129017242.1 | WP_129017912.1 |
| Bacteroidota | *Gelidibacter japonicus* | WP_163515830.1 | WP_250066750.1 | WP_323027882.1 |
| Bacteroidota | *Gelidibacter maritimus* | WP_182206455.1 | WP_182204067.1 | WP_182205246.1 |
| Bacteroidota | *Gelidibacter mesophilus* | WP_027126355.1 | WP_027127133.1 | WP_027125360.1 |
| Bacteroidota | *Gelidibacter pelagius* | WP_208233111.1 | WP_208233378.1 | WP_208232401.1 |
| Bacteroidota | *Gelidibacter salicanalis* | WP_146893995.1 | WP_199597625.1 | WP_146888335.1 |
| Bacteroidota | *Gelidibacter sediminis* | WP_133758180.1 | WP_133757911.1 | WP_133757493.1 |
| Bacteroidota | *Gelidibacter sp.* | WP_348037356.1 | WP_348037583.1 | WP_325452820.1 |
| Bacteroidota | *Geojedonia litorea* | WP_387961949.1 | WP_387961441.1 | WP_387960537.1 |
| Bacteroidota | *Gillisia hiemivivida* | WP_146931822.1 | WP_146931177.1 | WP_146933916.1 |
| Bacteroidota | *Gillisia limnaea* | WP_006987563.1 | WP_006990115.1 | WP_006987928.1 |
| Bacteroidota | *Gillisia lutea* | WP_236133762.1 | WP_236133415.1 | WP_236133176.1 |
| Bacteroidota | *Gillisia mitskevichiae* | WP_121345154.1 | WP_121345645.1 | WP_121345880.1 |
| Bacteroidota | *Gillisia sp. Hel_I_86* | WP_144963149.1 | WP_144962968.1 | WP_144958730.1 |
| Bacteroidota | *Gilvibacter sediminis* | WP_272832536.1 | WP_272832132.1 | WP_272833033.1 |
| Bacteroidota | *Gilvibacter sp.* | WP_291185014.1 | WP_420379623.1 | WP_374958602.1 |
| Bacteroidota | *Gilvirhabdus luticola* | WP_316663118.1 | WP_316662346.1 | WP_316661412.1 |
| Bacteroidota | *Ginsengibacter hankyongi* | WP_150413724.1 | WP_150415216.1 | WP_150413131.1 |
| Bacteroidota | *Gramella jeungdoensis* | WP_252113824.1 | WP_252110723.1 | WP_252110617.1 |
| Bacteroidota | *Gramella sp. MAR_2010_147* | WP_089664712.1 | WP_089665144.1 | WP_089664164.1 |
| Bacteroidota | *Gynurincola endophyticus* | WP_126973031.1 | WP_126971669.1 | WP_126971071.1 |
| Bacteroidota | *Haliscomenobacter hydrossis* | WP_013765591.1 | WP_013766642.1 | WP_044234184.1 |
| Bacteroidota | *Haliscomenobacter sp.* | WP_421797522.1 | WP_353485078.1 | WP_353483610.1 |
| Bacteroidota | *Haloflavibacter putidus* | WP_141420749.1 | WP_141421610.1 | WP_141422424.1 |

| | | | | |
|---|---|---|---|---|
| Bacteroidota | *Halomarinibacterium sedimenti* | WP_219052331.1 | WP_219052695.1 | WP_219052270.1 |
| Bacteroidota | *Halpernia frigidisoli* | WP_090079850.1 | WP_090080935.1 | WP_177205450.1 |
| Bacteroidota | *Halpernia humi* | WP_103913058.1 | WP_103913318.1 | WP_103913510.1 |
| Bacteroidota | *Halpernia sp.* | WP_417428735.1 | WP_417428411.1 | WP_417427724.1 |
| Bacteroidota | *Hanamia caeni* | WP_123121589.1 | WP_123121851.1 | WP_123122442.1 |
| Bacteroidota | *Hanstruepera flava* | WP_250436433.1 | WP_250432405.1 | WP_250434084.1 |
| Bacteroidota | *Hanstruepera marina* | WP_223034528.1 | WP_223033481.1 | WP_223033900.1 |
| Bacteroidota | *Hanstruepera neustonica* | WP_103052034.1 | WP_103052338.1 | WP_103051335.1 |
| Bacteroidota | *Hanstruepera ponticola* | WP_104733670.1 | WP_104733978.1 | WP_191859243.1 |
| Bacteroidota | *Haoranjiania flava* | WP_263039019.1 | WP_263036497.1 | WP_263036413.1 |
| Bacteroidota | *Hufsiella arboris* | WP_160844963.1 | WP_160844516.1 | WP_160845831.1 |
| Bacteroidota | *Hufsiella ginkgonis* | WP_160907940.1 | WP_160906799.1 | WP_160908011.1 |
| Bacteroidota | *Hugenholtzia roseola* | WP_027001197.1 | WP_027003762.1 | WP_051203851.1 |
| Bacteroidota | *Hwangdonia lutea* | WP_316983175.1 | WP_316983375.1 | WP_316982178.1 |
| Bacteroidota | *Hwangdonia seohaensis* | WP_311938626.1 | WP_311938917.1 | WP_311939926.1 |
| Bacteroidota | *Hwangdonia sp.* | WP_418604453.1 | WP_418604389.1 | WP_418603464.1 |
| Bacteroidota | *Hydrobacter penzbergensis* | WP_092724081.1 | WP_092725408.1 | WP_092722670.1 |
| Bacteroidota | *Hydrotalea flava* | WP_068247804.1 | WP_068244712.1 | WP_068244803.1 |
| Bacteroidota | *Hydrotalea sandarakina* | WP_111297305.1 | WP_211307670.1 | WP_111293278.1 |
| Bacteroidota | *Hydrotalea sp.* | WP_298412767.1 | WP_298298368.1 | WP_288071285.1 |
| Bacteroidota | *Hymenobacter actinosclerus* | WP_092772166.1 | WP_092770383.1 | WP_092773236.1 |
| Bacteroidota | *Hymenobacter aerilatus* | WP_245091095.1 | WP_245096394.1 | WP_245092938.1 |
| Bacteroidota | *Hymenobacter aerophilus* | WP_019947749.1 | WP_019946669.1 | WP_019946883.1 |
| Bacteroidota | *Hymenobacter algoricola* | WP_345113237.1 | WP_345114690.1 | WP_345109111.1 |
| Bacteroidota | *Hymenobacter amundsenii* | WP_088464214.1 | WP_088464474.1 | WP_088462720.1 |
| Bacteroidota | *Hymenobacter antarcticus* | WP_345121348.1 | WP_345123254.1 | WP_345125416.1 |
| Bacteroidota | *Hymenobacter aquaticus* | WP_210114293.1 | WP_135461986.1 | WP_135463988.1 |
| Bacteroidota | *Hymenobacter aranciens* | WP_305005511.1 | WP_305005013.1 | WP_305006523.1 |

| | | | | |
|---|---|---|---|---|
| Bacteroidota | *Hymenobacter arizonensis* | WP_092673250.1 | WP_092670262.1 | WP_092670509.1 |
| Bacteroidota | *Hymenobacter armeniacus* | WP_190923097.1 | WP_190922562.1 | WP_190924915.1 |
| Bacteroidota | *Hymenobacter artigasi* | WP_168675017.1 | WP_168672938.1 | WP_168672520.1 |
| Bacteroidota | *Hymenobacter baengnokdamensis* | WP_151087692.1 | WP_151088453.1 | WP_151086729.1 |
| Bacteroidota | *Hymenobacter bucti* | WP_382316155.1 | WP_382313791.1 | WP_382315386.1 |
| Bacteroidota | *Hymenobacter busanensis* | WP_151079239.1 | WP_338074383.1 | WP_151078604.1 |
| Bacteroidota | *Hymenobacter caeli* | WP_173808102.1 | WP_317171099.1 | WP_173808997.1 |
| Bacteroidota | *Hymenobacter canadensis* | WP_269560067.1 | WP_269561050.1 | WP_269558985.1 |
| Bacteroidota | *Hymenobacter cavernae* | WP_188815435.1 | WP_188813621.1 | WP_188811646.1 |
| Bacteroidota | *Hymenobacter chitinivorans* | WP_100334769.1 | WP_100336020.1 | WP_100337998.1 |
| Bacteroidota | *Hymenobacter citatus* | WP_187319983.1 | WP_187321137.1 | WP_187319347.1 |
| Bacteroidota | *Hymenobacter coccineus* | WP_070739286.1 | WP_070744121.1 | WP_070747050.1 |
| Bacteroidota | *Hymenobacter crusticola* | WP_086594948.1 | WP_086593780.1 | WP_086592258.1 |
| Bacteroidota | *Hymenobacter cyanobacteriorum* | WP_241937162.1 | WP_241935769.1 | WP_241936268.1 |
| Bacteroidota | *Hymenobacter daecheongensis* | WP_073108984.1 | WP_073111368.1 | WP_073105191.1 |
| Bacteroidota | *Hymenobacter duratus* | WP_190784662.1 | WP_190785735.1 | WP_190783158.1 |
| Bacteroidota | *Hymenobacter edaphi* | WP_111478724.1 | WP_111476768.1 | WP_111477643.1 |
| Bacteroidota | *Hymenobacter elongatus* | WP_135499093.1 | WP_135499484.1 | WP_135498247.1 |
| Bacteroidota | *Hymenobacter endophyticus* | WP_315999127.1 | WP_315997629.1 | WP_315997288.1 |
| Bacteroidota | *Hymenobacter fastidiosus* | WP_345074853.1 | WP_345072700.1 | WP_345071857.1 |
| Bacteroidota | *Hymenobacter fodinae* | WP_135433580.1 | WP_135436110.1 | WP_135430550.1 |
| Bacteroidota | *Hymenobacter frigidus* | WP_188561526.1 | WP_188560312.1 | WP_188561206.1 |
| Bacteroidota | *Hymenobacter gelipurpurascens* | WP_088843741.1 | WP_088842140.1 | WP_088844610.1 |
| Bacteroidota | *Hymenobacter ginkgonis* | WP_157568925.1 | WP_157562471.1 | WP_157563796.1 |
| Bacteroidota | *Hymenobacter glacialis* | WP_070734572.1 | WP_070731728.1 | WP_070735962.1 |
| Bacteroidota | *Hymenobacter glaciei* | WP_345051671.1 | WP_345050062.1 | WP_345056698.1 |
| Bacteroidota | *Hymenobacter glacieicola* | WP_188559509.1 | WP_188558562.1 | WP_188556977.1 |
| Bacteroidota | *Hymenobacter gummosus* | WP_126696445.1 | WP_126695648.1 | WP_126694911.1 |

| | | | | |
|---|---|---|---|---|
| Bacteroidota | *Hymenobacter guriensis* | WP_196953312.1 | WP_196955704.1 | WP_196956148.1 |
| Bacteroidota | *Hymenobacter jeollabukensis* | WP_138077952.1 | WP_138074696.1 | WP_138076488.1 |
| Bacteroidota | *Hymenobacter jeongseonensis* | WP_196281111.1 | WP_196283366.1 | WP_196282795.1 |
| Bacteroidota | *Hymenobacter koreensis* | WP_345220804.1 | WP_345225765.1 | WP_345227550.1 |
| Bacteroidota | *Hymenobacter lapidiphilus* | WP_176907383.1 | WP_176906325.1 | WP_176910239.1 |
| Bacteroidota | *Hymenobacter lucidus* | WP_226175553.1 | WP_226177909.1 | WP_226170797.1 |
| Bacteroidota | *Hymenobacter lutimineralis* | WP_149070077.1 | WP_149069064.1 | WP_149069601.1 |
| Bacteroidota | *Hymenobacter mellowenesis* | WP_305011453.1 | WP_305014459.1 | WP_305010093.1 |
| Bacteroidota | *Hymenobacter metallicola* | WP_135394722.1 | WP_135392312.1 | WP_135395606.1 |
| Bacteroidota | *Hymenobacter metallilatus* | WP_125427055.1 | WP_125432977.1 | WP_125426004.1 |
| Bacteroidota | *Hymenobacter montanus* | WP_191003208.1 | WP_191005164.1 | WP_191003835.1 |
| Bacteroidota | *Hymenobacter monticola* | WP_243508845.1 | WP_243512292.1 | WP_243516219.1 |
| Bacteroidota | *Hymenobacter negativus* | WP_198066871.1 | WP_198068458.1 | WP_198069842.1 |
| Bacteroidota | *Hymenobacter nitidus* | WP_226189119.1 | WP_226183341.1 | WP_226186023.1 |
| Bacteroidota | *Hymenobacter nivis* | WP_140465004.1 | WP_342767440.1 | WP_109657402.1 |
| Bacteroidota | *Hymenobacter norwichensis* | WP_022823951.1 | WP_022823749.1 | WP_022825075.1 |
| Bacteroidota | *Hymenobacter oligotrophus* | WP_119444743.1 | WP_119443979.1 | WP_119445303.1 |
| Bacteroidota | *Hymenobacter perfusus* | WP_125439651.1 | WP_125440281.1 | WP_125435945.1 |
| Bacteroidota | *Hymenobacter persicinus* | WP_129922767.1 | WP_129922804.1 | WP_129923055.1 |
| Bacteroidota | *Hymenobacter pini* | WP_226265934.1 | WP_226268672.1 | WP_226271625.1 |
| Bacteroidota | *Hymenobacter piscis* | WP_215594052.1 | WP_215592429.1 | WP_215591981.1 |
| Bacteroidota | *Hymenobacter polaris* | WP_169528951.1 | WP_169530971.1 | WP_169532678.1 |
| Bacteroidota | *Hymenobacter profundi* | WP_219160458.1 | WP_219159751.1 | WP_219161744.1 |
| Bacteroidota | *Hymenobacter properus* | WP_196287755.1 | WP_196284682.1 | WP_196285685.1 |
| Bacteroidota | *Hymenobacter psoromatis* | WP_223648793.1 | WP_223649629.1 | WP_223653225.1 |
| Bacteroidota | *Hymenobacter psychrophilus* | WP_092739454.1 | WP_092741564.1 | WP_092739666.1 |
| Bacteroidota | *Hymenobacter psychrotolerans* | WP_073286580.1 | WP_073287465.1 | WP_073285851.1 |
| Bacteroidota | *Hymenobacter qilianensis* | WP_187732856.1 | WP_187732044.1 | WP_187733782.1 |

| | | | | |
|---|---|---|---|---|
| Bacteroidota | *Hymenobacter rigui* | WP_125420006.1 | WP_125423685.1 | WP_125417329.1 |
| Bacteroidota | *Hymenobacter roseosalivarius* | WP_084444424.1 | WP_084447454.1 | WP_084445205.1 |
| Bacteroidota | *Hymenobacter rubidus* | WP_201986503.1 | WP_201984882.1 | WP_201984100.1 |
| Bacteroidota | *Hymenobacter rubripertinctus* | WP_119656221.1 | WP_119654072.1 | WP_119654547.1 |
| Bacteroidota | *Hymenobacter ruricola* | WP_196293568.1 | WP_196291767.1 | WP_196293999.1 |
| Bacteroidota | *Hymenobacter saemangeumensis* | WP_345238436.1 | WP_345237286.1 | WP_345237946.1 |
| Bacteroidota | *Hymenobacter sedentarius* | WP_068197752.1 | WP_068195151.1 | WP_068191064.1 |
| Bacteroidota | *Hymenobacter sediminicola* | WP_185887369.1 | WP_185886273.1 | WP_185888447.1 |
| Bacteroidota | *Hymenobacter sediminis* | WP_202910460.1 | WP_110979241.1 | WP_110976343.1 |
| Bacteroidota | *Hymenobacter segetis* | WP_342300887.1 | WP_342301024.1 | WP_342295151.1 |
| Bacteroidota | *Hymenobacter setariae* | WP_144851550.1 | WP_144845586.1 | WP_144847603.1 |
| Bacteroidota | *Hymenobacter siberiensis* | WP_216727714.1 | WP_216688865.1 | WP_216679529.1 |
| Bacteroidota | *Hymenobacter sp.* | WP_325135645.1 | WP_310397190.1 | WP_310393716.1 |
| Bacteroidota | *Hymenobacter sublimis* | WP_247975982.1 | WP_247976812.1 | WP_247975169.1 |
| Bacteroidota | *Hymenobacter swuensis* | WP_044001162.1 | WP_044002076.1 | WP_044003628.1 |
| Bacteroidota | *Hymenobacter telluris* | WP_206982133.1 | WP_206984410.1 | WP_206986445.1 |
| Bacteroidota | *Hymenobacter terrenus* | WP_046243211.1 | WP_046243040.1 | WP_046246906.1 |
| Bacteroidota | *Hymenobacter terrestris* | WP_176900196.1 | WP_176898993.1 | WP_176901120.1 |
| Bacteroidota | *Hymenobacter terricola* | WP_210518078.1 | WP_210515764.1 | WP_210513663.1 |
| Bacteroidota | *Hymenobacter tibetensis* | WP_243800611.1 | WP_243802657.1 | WP_243799028.1 |
| Bacteroidota | *Hymenobacter translucens* | WP_227609277.1 | WP_227606247.1 | WP_227610336.1 |
| Bacteroidota | *Hymenobacter volaticus* | WP_245122496.1 | WP_245125362.1 | WP_245120586.1 |
| Bacteroidota | *Hymenobacter wooponensis* | WP_135529106.1 | WP_135530529.1 | WP_135530138.1 |
| Bacteroidota | *Hymenobacter yonginensis* | WP_270127571.1 | WP_270128601.1 | WP_270126306.1 |
| Bacteroidota | *Hyunsoonleella aestuarii* | WP_139002770.1 | WP_139000583.1 | WP_139000362.1 |
| Bacteroidota | *Hyunsoonleella aquatilis* | WP_186563531.1 | WP_186563450.1 | WP_186559073.1 |
| Bacteroidota | *Hyunsoonleella flava* | WP_130962624.1 | WP_130962429.1 | WP_130963622.1 |
| Bacteroidota | *Hyunsoonleella jejuensis* | WP_092578219.1 | WP_092578669.1 | WP_092579037.1 |

| | | | | |
|---|---|---|---|---|
| Bacteroidota | *Hyunsoonleella pacifica* | WP_130935536.1 | WP_130935871.1 | WP_130938023.1 |
| Bacteroidota | *Hyunsoonleella rubra* | WP_380288079.1 | WP_380289797.1 | WP_380292807.1 |
| Bacteroidota | *Hyunsoonleella sp. 2307UL5-6* | WP_413972052.1 | WP_413973366.1 | WP_413973261.1 |
| Bacteroidota | *Hyunsoonleella ulvae* | WP_203256821.1 | WP_203258597.1 | WP_203257890.1 |
| Bacteroidota | *Ichthyenterobacterium magnum* | WP_120200732.1 | WP_120201318.1 | WP_120199279.1 |
| Bacteroidota | *Ichthyobacterium seriolicida* | WP_096686479.1 | WP_096686568.1 | WP_096685584.1 |
| Bacteroidota | *Ilyomonas limi* | WP_137261080.1 | WP_137261789.1 | WP_137263876.1 |
| Bacteroidota | *Imperialibacter roseus* | WP_317488528.1 | WP_317492116.1 | WP_317489223.1 |
| Bacteroidota | *Imperialibacter sp.* | WP_350078291.1 | WP_350095892.1 | WP_350117640.1 |
| Bacteroidota | *Imtechella halotolerans* | WP_008240408.1 | WP_008238821.1 | WP_008238112.1 |
| Bacteroidota | *Indibacter alkaliphilus* | WP_009032885.1 | WP_009036264.1 | WP_009033840.1 |
| Bacteroidota | *Jejudonia soesokkakensis* | WP_380216994.1 | WP_380217385.1 | WP_380218067.1 |
| Bacteroidota | *Jejuia pallidilutea* | WP_042243901.1 | WP_105474849.1 | WP_105474438.1 |
| Bacteroidota | *Jejuia sp. DST062* | WP_406971608.1 | WP_406969494.1 | WP_406968753.1 |
| Bacteroidota | *Jejuia spongiicola* | WP_249972989.1 | WP_249973577.1 | WP_249972470.1 |
| Bacteroidota | *Jiulongibacter sediminis* | WP_304235437.1 | WP_055145038.1 | WP_304237026.1 |
| Bacteroidota | *Joostella atrarenae* | WP_236959290.1 | WP_236959699.1 | WP_236958707.1 |
| Bacteroidota | *Joostella sp.* | WP_417444730.1 | WP_417444046.1 | WP_417444373.1 |
| Bacteroidota | *Kaistella antarctica* | WP_034718549.1 | WP_034718534.1 | WP_051803858.1 |
| Bacteroidota | *Kaistella carnis* | WP_125025455.1 | WP_125025449.1 | WP_313502080.1 |
| Bacteroidota | *Kaistella chaponensis* | WP_324472738.1 | WP_076386374.1 | WP_076387281.1 |
| Bacteroidota | *Kaistella daneshvariae* | WP_123266406.1 | WP_123266400.1 | WP_123266721.1 |
| Bacteroidota | *Kaistella faecalis* | WP_218248450.1 | WP_218248442.1 | WP_218250327.1 |
| Bacteroidota | *Kaistella flava (ex Peng et al. 2021)* | WP_193811537.1 | WP_193811529.1 | WP_193811028.1 |
| Bacteroidota | *Kaistella gelatinilytica* | WP_196079956.1 | WP_196079963.1 | WP_196080199.1 |
| Bacteroidota | *Kaistella haifensis* | WP_031501350.1 | WP_031501345.1 | WP_031504621.1 |
| Bacteroidota | *Kaistella jeonii* | WP_039351184.1 | WP_039351160.1 | WP_039350477.1 |
| Bacteroidota | *Kaistella montana* | WP_255929243.1 | WP_255929267.1 | WP_255930280.1 |

| | | | | |
|---|---|---|---|---|
| Bacteroidota | *Kaistella palustris* | WP_027376032.1 | WP_027376024.1 | WP_027377070.1 |
| Bacteroidota | *Kaistella polysaccharea* | WP_226063347.1 | WP_226063355.1 | WP_226063808.1 |
| Bacteroidota | *Kaistella soli* | WP_088358885.1 | WP_217201330.1 | WP_217201368.1 |
| Bacteroidota | *Kaistella solincola* | WP_039342678.1 | WP_039341850.1 | WP_039341001.1 |
| Bacteroidota | *Kaistella sp.* | WP_332031973.1 | WP_332031967.1 | WP_423974762.1 |
| Bacteroidota | *Kaistella treverensis* | WP_089819909.1 | WP_089819511.1 | WP_089819931.1 |
| Bacteroidota | *Kaistella yananensis* | WP_265142890.1 | WP_265142884.1 | WP_265144070.1 |
| Bacteroidota | *Kaistella yonginensis* | WP_290216955.1 | WP_290216940.1 | WP_290217812.1 |
| Bacteroidota | *Kordia aestuariivivens* | WP_187563470.1 | WP_187561095.1 | WP_187564525.1 |
| Bacteroidota | *Kordia algicida* | WP_007093984.1 | WP_040559864.1 | WP_007094736.1 |
| Bacteroidota | *Kordia antarctica* | WP_160128089.1 | WP_160128402.1 | WP_160131354.1 |
| Bacteroidota | *Kordia jejudonensis* | WP_046758503.1 | WP_046758012.1 | WP_046757471.1 |
| Bacteroidota | *Kordia periserrulae* | WP_108114659.1 | WP_108114435.1 | WP_108116523.1 |
| Bacteroidota | *Kordia sp.* | WP_405302502.1 | WP_420572871.1 | WP_405302795.1 |
| Bacteroidota | *Kordia zhangzhouensis* | WP_046744617.1 | WP_046744409.1 | WP_046745889.1 |
| Bacteroidota | *Kriegella aquimaris* | WP_089885287.1 | WP_089884735.1 | WP_089894501.1 |
| Bacteroidota | *Lacibacter cauensis* | WP_144886451.1 | WP_144884137.1 | WP_144886127.1 |
| Bacteroidota | *Lacibacter luteus* | WP_129131595.1 | WP_129128817.1 | WP_129132186.1 |
| Bacteroidota | *Lacibacter sediminis* | WP_182803974.1 | WP_182801129.1 | WP_182801935.1 |
| Bacteroidota | *Lacibacter sp.* | WP_324231404.1 | WP_324232070.1 | WP_324230093.1 |
| Bacteroidota | *Lacihabitans lacunae* | WP_379835146.1 | WP_379834768.1 | WP_379838877.1 |
| Bacteroidota | *Lacihabitans soyangensis* | WP_255038281.1 | WP_255039040.1 | WP_255037505.1 |
| Bacteroidota | *Lacihabitans sp. CCS-44* | WP_255080260.1 | WP_255076565.1 | WP_255077870.1 |
| Bacteroidota | *Lacinutrix algicola* | WP_055437101.1 | WP_055437076.1 | WP_055435440.1 |
| Bacteroidota | *Lacinutrix gracilariae* | WP_379901067.1 | WP_379899746.1 | WP_379902270.1 |
| Bacteroidota | *Lacinutrix himadriensis* | WP_055444954.1 | WP_055443481.1 | WP_055445236.1 |
| Bacteroidota | *Lacinutrix iliipiscaria* | WP_183487450.1 | WP_183488070.1 | WP_183486898.1 |
| Bacteroidota | *Lacinutrix jangbogonensis* | WP_034058557.1 | WP_034059251.1 | WP_034057457.1 |

| | | | | |
|---|---|---|---|---|
| Bacteroidota | *Lacinutrix mariniflava* | WP_055446055.1 | WP_055448589.1 | WP_055448788.1 |
| Bacteroidota | *Lacinutrix sp.* | WP_340169257.1 | WP_340169113.1 | WP_290698518.1 |
| Bacteroidota | *Lacinutrix venerupis* | WP_121048868.1 | WP_076733424.1 | WP_121052254.1 |
| Bacteroidota | *Larkinella arboricola* | WP_111628622.1 | WP_111628276.1 | WP_111628983.1 |
| Bacteroidota | *Larkinella bovis* | WP_379847279.1 | WP_379840490.1 | WP_379846488.1 |
| Bacteroidota | *Larkinella humicola* | WP_150876445.1 | WP_150874987.1 | WP_150877174.1 |
| Bacteroidota | *Larkinella insperata* | WP_265992729.1 | WP_265990224.1 | WP_265993129.1 |
| Bacteroidota | *Larkinella knui* | WP_124910643.1 | WP_124904358.1 | WP_124909924.1 |
| Bacteroidota | *Larkinella rosea* | WP_124875391.1 | WP_124870784.1 | WP_124874597.1 |
| Bacteroidota | *Larkinella soli* | WP_128548767.1 | WP_128546727.1 | WP_128545547.1 |
| Bacteroidota | *Larkinella sp.* | WP_421828999.1 | WP_421830674.1 | WP_421827887.1 |
| Bacteroidota | *Larkinella terrae* | WP_154172480.1 | WP_154176098.1 | WP_154174654.1 |
| Bacteroidota | *Leadbetterella byssophila* | WP_013407413.1 | WP_412814525.1 | WP_412814608.1 |
| Bacteroidota | *Leadbetterella sp. DM7* | WP_367915067.1 | WP_367915579.1 | WP_367913745.1 |
| Bacteroidota | *Leeuwenhoekiella aequorea* | WP_128758357.1 | WP_128757498.1 | WP_128758662.1 |
| Bacteroidota | *Leeuwenhoekiella aestuarii* | WP_128760275.1 | WP_128762123.1 | WP_128760454.1 |
| Bacteroidota | *Leeuwenhoekiella marinoflava* | WP_335620825.1 | WP_073099091.1 | WP_335620663.1 |
| Bacteroidota | *Leeuwenhoekiella nanhaiensis* | WP_099646209.1 | WP_099646328.1 | WP_099645389.1 |
| Bacteroidota | *Leeuwenhoekiella palythoae* | WP_225597923.1 | WP_072979912.1 | WP_370174603.1 |
| Bacteroidota | *Leeuwenhoekiella parthenopeia* | WP_228230864.1 | WP_228230944.1 | WP_228228275.1 |
| Bacteroidota | *Leeuwenhoekiella polynyae* | WP_128764228.1 | WP_128763846.1 | WP_128765667.1 |
| Bacteroidota | *Leeuwenhoekiella sp. NPDC079379* | WP_396511161.1 | WP_396512396.1 | WP_396510153.1 |
| Bacteroidota | *Lentiprolixibacter aurantiacus* | WP_266013079.1 | WP_266012786.1 | WP_266013347.1 |
| Bacteroidota | *Leptobacterium flavescens* | WP_163607265.1 | WP_163607288.1 | WP_163605276.1 |
| Bacteroidota | *Leptobacterium meishanense* | WP_340077561.1 | WP_340073967.1 | WP_340074705.1 |
| Bacteroidota | *Lewinella cohaerens* | WP_020536610.1 | WP_020536903.1 | WP_020538753.1 |
| Bacteroidota | *Lewinella sp. LCG006* | WP_367391533.1 | WP_367388024.1 | WP_367390254.1 |
| Bacteroidota | *Limibacter armeniacum* | WP_334246576.1 | WP_334246496.1 | WP_334246523.1 |

| | | | | |
|---|---|---|---|---|
| Bacteroidota | *Limnovirga soli* | WP_171609007.1 | WP_171609473.1 | WP_171609767.1 |
| Bacteroidota | *Lishizhenia sp.* | WP_318958285.1 | WP_318958486.1 | WP_318958685.1 |
| Bacteroidota | *Lishizhenia tianjinensis* | WP_090246723.1 | WP_090247519.1 | WP_090245381.1 |
| Bacteroidota | *Litoribacter populi* | WP_143962451.1 | WP_143960986.1 | WP_143959597.1 |
| Bacteroidota | *Litoribacter ruber* | WP_213943340.1 | WP_213945420.1 | WP_214162654.1 |
| Bacteroidota | *Litoribaculum gwangyangense* | WP_345276104.1 | WP_345276075.1 | WP_345276767.1 |
| Bacteroidota | *Longitalea arenae* | WP_205508771.1 | WP_205512896.1 | WP_205509770.1 |
| Bacteroidota | *Longitalea luteola* | WP_207511955.1 | WP_207513435.1 | WP_207513616.1 |
| Bacteroidota | *Lunatibacter salilacus* | WP_158858628.1 | WP_158857098.1 | WP_158855923.1 |
| Bacteroidota | *Lunatimonas lonarensis* | WP_010853832.1 | WP_035805042.1 | WP_010856401.1 |
| Bacteroidota | *Lunatimonas salinarum* | WP_209329637.1 | WP_209330338.1 | WP_209332122.1 |
| Bacteroidota | *Lunatimonas sp.* | WP_228693011.1 | WP_228691852.1 | WP_228692448.1 |
| Bacteroidota | *Luteibaculum oceani* | WP_147014709.1 | WP_147014190.1 | WP_147013057.1 |
| Bacteroidota | *Luteirhabdus pelagi* | WP_203295780.1 | WP_203296683.1 | WP_203296609.1 |
| Bacteroidota | *Lutibacter agarilyticus* | WP_089380012.1 | WP_089380156.1 | WP_089381860.1 |
| Bacteroidota | *Lutibacter citreus* | WP_111708460.1 | WP_111707821.1 | WP_111708609.1 |
| Bacteroidota | *Lutibacter flavus* | WP_089377124.1 | WP_089376832.1 | WP_089377379.1 |
| Bacteroidota | *Lutibacter holmesii* | WP_386809209.1 | WP_386809111.1 | WP_386807184.1 |
| Bacteroidota | *Lutibacter maritimus* | WP_090222967.1 | WP_090222428.1 | WP_090221845.1 |
| Bacteroidota | *Lutibacter oceani* | WP_115879279.1 | WP_115879655.1 | WP_115882099.1 |
| Bacteroidota | *Lutibacter oricola* | WP_090121462.1 | WP_090120722.1 | WP_090123020.1 |
| Bacteroidota | *Lutibacter profundi* | WP_068205635.1 | WP_068205690.1 | WP_068206420.1 |
| Bacteroidota | *Lutibacter sp.* | WP_408670705.1 | WP_291970443.1 | WP_291975456.1 |
| Bacteroidota | *Lutimonas halocynthiae* | WP_290293069.1 | WP_290293732.1 | WP_290289555.1 |
| Bacteroidota | *Lutimonas saemankumensis* | WP_224927446.1 | WP_224927476.1 | WP_224932014.1 |
| Bacteroidota | *Lutimonas sp.* | WP_424350756.1 | WP_424350450.1 | WP_424352018.1 |
| Bacteroidota | *Lutimonas vermicola* | WP_342158194.1 | WP_342159627.1 | WP_342159791.1 |
| Bacteroidota | *Lutimonas zeaxanthinifaciens* | WP_302043014.1 | WP_302043042.1 | WP_302043417.1 |

| | | | | |
|---|---|---|---|---|
| Bacteroidota | *Mangrovimonas aestuarii* | WP_274475567.1 | WP_274475634.1 | WP_274475942.1 |
| Bacteroidota | *Mangrovimonas cancribranchiae* | WP_338733894.1 | WP_338733664.1 | WP_338733268.1 |
| Bacteroidota | *Mangrovimonas futianensis* | WP_232634821.1 | WP_232395550.1 | WP_232634478.1 |
| Bacteroidota | *Mangrovimonas sp. DI 80* | WP_076663859.1 | WP_076664802.1 | WP_076663450.1 |
| Bacteroidota | *Mangrovimonas spongiae* | WP_125468490.1 | WP_125467189.1 | WP_125467469.1 |
| Bacteroidota | *Mangrovimonas xylaniphaga* | WP_053977790.1 | WP_053977261.1 | WP_053977459.1 |
| Bacteroidota | *Mangrovimonas yunxiaonensis* | WP_036123792.1 | WP_036123026.1 | WP_036122376.1 |
| Bacteroidota | *Mangrovivirga halotolerans* | WP_266058647.1 | WP_266056794.1 | WP_266057182.1 |
| Bacteroidota | *Maribacter aestuarii* | WP_289644541.1 | WP_281543132.1 | WP_289644842.1 |
| Bacteroidota | *Maribacter algarum* | WP_138659431.1 | WP_138658534.1 | WP_138658835.1 |
| Bacteroidota | *Maribacter algicola* | WP_125223587.1 | WP_125223285.1 | WP_125224015.1 |
| Bacteroidota | *Maribacter antarcticus* | WP_027077533.1 | WP_027075876.1 | WP_027075275.1 |
| Bacteroidota | *Maribacter aquimaris* | WP_188244283.1 | WP_188241762.1 | WP_188244462.1 |
| Bacteroidota | *Maribacter aquivivus* | WP_073246261.1 | WP_073243963.1 | WP_282051155.1 |
| Bacteroidota | *Maribacter arcticus* | WP_079512125.1 | WP_335637209.1 | WP_335637955.1 |
| Bacteroidota | *Maribacter arenosus* | WP_188312956.1 | WP_188312492.1 | WP_188315229.1 |
| Bacteroidota | *Maribacter aurantiacus* | WP_138259457.1 | WP_138258082.1 | WP_138259233.1 |
| Bacteroidota | *Maribacter caenipelagi* | WP_133673642.1 | WP_133671069.1 | WP_133673480.1 |
| Bacteroidota | *Maribacter chungangensis* | WP_379935338.1 | WP_379934434.1 | WP_379934574.1 |
| Bacteroidota | *Maribacter cobaltidurans* | WP_094995491.1 | WP_094998851.1 | WP_094997970.1 |
| Bacteroidota | *Maribacter confluentis* | WP_304437470.1 | WP_304437268.1 | WP_304435015.1 |
| Bacteroidota | *Maribacter dokdonensis* | WP_210709020.1 | WP_348427586.1 | WP_321828019.1 |
| Bacteroidota | *Maribacter forsetii* | WP_036153520.1 | WP_036154123.1 | WP_036155838.1 |
| Bacteroidota | *Maribacter halichondriae* | WP_273566554.1 | WP_273566662.1 | WP_337251723.1 |
| Bacteroidota | *Maribacter huludaoensis* | WP_276473091.1 | WP_276473340.1 | WP_276472642.1 |
| Bacteroidota | *Maribacter hydrothermalis* | WP_068482505.1 | WP_068484963.1 | WP_068484075.1 |
| Bacteroidota | *Maribacter litoralis* | WP_116772091.1 | WP_116768890.1 | WP_159301745.1 |
| Bacteroidota | *Maribacter luteus* | WP_282056634.1 | WP_282054775.1 | WP_154368496.1 |

| | | | | |
|---|---|---|---|---|
| Bacteroidota | *Maribacter orientalis* | WP_091627189.1 | WP_091625859.1 | WP_091626195.1 |
| Bacteroidota | *Maribacter polysaccharolyticus* | WP_274826574.1 | WP_274823406.1 | WP_274824402.1 |
| Bacteroidota | *Maribacter polysiphoniae* | WP_109652914.1 | WP_109654071.1 | WP_273276891.1 |
| Bacteroidota | *Maribacter sp.* | WP_291960873.1 | WP_291964572.1 | WP_348369336.1 |
| Bacteroidota | *Maribacter spongiicola* | WP_133687150.1 | WP_133687444.1 | WP_133688997.1 |
| Bacteroidota | *Maribacter stanieri* | WP_282113329.1 | WP_339725903.1 | WP_282113675.1 |
| Bacteroidota | *Maribacter thermophilus* | WP_047245800.1 | WP_047245436.1 | WP_047244756.1 |
| Bacteroidota | *Maribacter ulvicola* | WP_076548560.1 | WP_076549865.1 | WP_076549380.1 |
| Bacteroidota | *Maribacter vaceletii* | WP_121069140.1 | WP_121065987.1 | WP_121069339.1 |
| Bacteroidota | *Maribacter zhoushanensis* | WP_276494026.1 | WP_276493126.1 | WP_276494673.1 |
| Bacteroidota | *Mariniflexile aquimaris* | WP_379942415.1 | WP_379942788.1 | WP_379941225.1 |
| Bacteroidota | *Mariniflexile fucanivorans* | WP_132219156.1 | WP_132215582.1 | WP_132216338.1 |
| Bacteroidota | *Mariniflexile gromovii* | WP_209656289.1 | WP_209657102.1 | WP_209654042.1 |
| Bacteroidota | *Mariniflexile jejuense* | WP_379924078.1 | WP_379926576.1 | WP_379926525.1 |
| Bacteroidota | *Mariniflexile litorale* | WP_308992851.1 | WP_308992244.1 | WP_308992065.1 |
| Bacteroidota | *Mariniflexile maritimum* | WP_157206636.1 | WP_157208718.1 | WP_157206179.1 |
| Bacteroidota | *Mariniflexile ostreae* | WP_379860781.1 | WP_379861361.1 | WP_379860627.1 |
| Bacteroidota | *Mariniflexile soesokkakense* | WP_346242016.1 | WP_346239891.1 | WP_346242332.1 |
| Bacteroidota | *Mariniflexile sp.* | WP_372755176.1 | WP_372754476.1 | WP_372754686.1 |
| Bacteroidota | *Marinigracilibium pacificum* | WP_169684667.1 | WP_169684819.1 | WP_169677532.1 |
| Bacteroidota | *Marinilongibacter aquaticus* | WP_261451105.1 | WP_261449973.1 | WP_261447952.1 |
| Bacteroidota | *Mariniradius saccharolyticus* | WP_008627210.1 | WP_008628996.1 | WP_040480123.1 |
| Bacteroidota | *Mariniradius sediminis* | WP_234862381.1 | WP_234862495.1 | WP_234861300.1 |
| Bacteroidota | *Marinirhabdus gelatinilytica* | WP_115124645.1 | WP_115123303.1 | WP_115123992.1 |
| Bacteroidota | *Marinoscillum furvescens* | WP_115866794.1 | WP_245986162.1 | WP_115868181.1 |
| Bacteroidota | *Marinoscillum pacificum* | WP_258099391.1 | WP_258099852.1 | WP_258100254.1 |
| Bacteroidota | *Marinoscillum sp.* | WP_421873655.1 | WP_421878018.1 | WP_421892226.1 |
| Bacteroidota | *Marivirga arenosa* | WP_308356037.1 | WP_308356793.1 | WP_308357795.1 |

| | | | | |
|---|---|---|---|---|
| Bacteroidota | *Marivirga atlantica* | WP_201918138.1 | WP_201924100.1 | WP_201916581.1 |
| Bacteroidota | *Marivirga aurantiaca* | WP_201432316.1 | WP_201429891.1 | WP_201432555.1 |
| Bacteroidota | *Marivirga harenae* | WP_303267928.1 | WP_303272113.1 | WP_303270879.1 |
| Bacteroidota | *Marivirga lumbricoides* | WP_188460934.1 | WP_188462202.1 | WP_188465104.1 |
| Bacteroidota | *Marivirga salinae* | WP_308351288.1 | WP_308347826.1 | WP_308349786.1 |
| Bacteroidota | *Marivirga sericea* | WP_085517122.1 | WP_085515726.1 | WP_085518009.1 |
| Bacteroidota | *Marivirga sp.* | WP_296618831.1 | WP_296622034.1 | WP_325121820.1 |
| Bacteroidota | *Marivirga tractuosa* | WP_375580939.1 | WP_375577921.1 | WP_013454056.1 |
| Bacteroidota | *Marixanthomonas ophiurae* | WP_117160044.1 | WP_117160475.1 | WP_117159718.1 |
| Bacteroidota | *Marixanthomonas sp. SCSIO 43207* | WP_223107920.1 | WP_223107512.1 | WP_223109995.1 |
| Bacteroidota | *Marixanthomonas spongiae* | WP_116693667.1 | WP_116693396.1 | WP_116694569.1 |
| Bacteroidota | *Marixanthotalea marina* | WP_318543194.1 | WP_318543261.1 | WP_318542961.1 |
| Bacteroidota | *Marnyiella aurantia* | WP_181886854.1 | WP_209815644.1 | WP_209814436.1 |
| Bacteroidota | *Membranihabitans marinus* | WP_236970699.1 | WP_236969875.1 | WP_236969502.1 |
| Bacteroidota | *Membranihabitans maritimus* | WP_236974510.1 | WP_236979183.1 | WP_236979239.1 |
| Bacteroidota | *Mesoflavibacter sp.* | WP_370225638.1 | WP_370228053.1 | WP_370225799.1 |
| Bacteroidota | *Mesoflavibacter zeaxanthinifaciens* | WP_417557548.1 | WP_027879723.1 | WP_027880087.1 |
| Bacteroidota | *Mesohalobacter halotolerans* | WP_138931676.1 | WP_138932211.1 | WP_138932670.1 |
| Bacteroidota | *Mesonia aestuariivivens* | WP_219039071.1 | WP_219040755.1 | WP_219041146.1 |
| Bacteroidota | *Mesonia algae* | WP_111540188.1 | WP_111541586.1 | WP_111541566.1 |
| Bacteroidota | *Mesonia aquimarina* | WP_121665309.1 | WP_121665496.1 | WP_121666282.1 |
| Bacteroidota | *Mesonia hippocampi* | WP_183477614.1 | WP_183475703.1 | WP_183477477.1 |
| Bacteroidota | *Mesonia maritima* | WP_309729227.1 | WP_309728181.1 | WP_309728893.1 |
| Bacteroidota | *Mesonia mobilis* | WP_304137891.1 | WP_304038444.1 | WP_304136972.1 |
| Bacteroidota | *Mesonia ostreae* | WP_311401302.1 | WP_311400781.1 | WP_311400862.1 |
| Bacteroidota | *Mesonia phycicola* | WP_073148113.1 | WP_073149804.1 | WP_073149941.1 |
| Bacteroidota | *Mesonia profundi* | WP_308863989.1 | WP_308864680.1 | WP_308863498.1 |
| Bacteroidota | *Mesonia sediminis* | WP_379044484.1 | WP_379046647.1 | WP_379045708.1 |

| | | | | |
|---|---|---|---|---|
| Bacteroidota | *Mesonia sp.* | WP_292248828.1 | WP_292246739.1 | WP_347418592.1 |
| Bacteroidota | *Microcosmobacter mediterraneus* | WP_311427926.1 | WP_311426391.1 | WP_311425829.1 |
| Bacteroidota | *Microscilla marina* | WP_002698220.1 | WP_002693779.1 | WP_002700262.1 |
| Bacteroidota | *Moheibacter lacus* | WP_182044234.1 | WP_182043139.1 | WP_182042476.1 |
| Bacteroidota | *Moheibacter sediminis* | WP_084017991.1 | WP_084017945.1 | WP_159447435.1 |
| Bacteroidota | *Moheibacter sp.* | WP_424368457.1 | WP_424367864.1 | WP_424368122.1 |
| Bacteroidota | *Moheibacter stercoris* | WP_354510703.1 | WP_354506512.1 | WP_354505341.1 |
| Bacteroidota | *Mongoliibacter ruber* | WP_106133080.1 | WP_106132015.1 | WP_106134676.1 |
| Bacteroidota | *Mongoliibacter sp.* | WP_293012583.1 | WP_293012793.1 | WP_293011822.1 |
| Bacteroidota | *Mongoliitalea daihaiensis* | WP_236139112.1 | WP_236136828.1 | WP_236139309.1 |
| Bacteroidota | *Mongoliitalea lutea* | WP_189579276.1 | WP_189585048.1 | WP_189579197.1 |
| Bacteroidota | *Mucilaginibacter agri* | WP_166584801.1 | WP_166584559.1 | WP_166587319.1 |
| Bacteroidota | *Mucilaginibacter angelicae* | WP_377026524.1 | WP_377023939.1 | WP_377021327.1 |
| Bacteroidota | *Mucilaginibacter antarcticus* | WP_377126946.1 | WP_377129763.1 | WP_377128107.1 |
| Bacteroidota | *Mucilaginibacter aquaedulcis* | WP_290327273.1 | WP_290321937.1 | WP_290325587.1 |
| Bacteroidota | *Mucilaginibacter aquariorum* | WP_256540140.1 | WP_256539051.1 | WP_256538611.1 |
| Bacteroidota | *Mucilaginibacter aquatilis* | WP_157540593.1 | WP_317163585.1 | WP_157539959.1 |
| Bacteroidota | *Mucilaginibacter arboris* | WP_157566211.1 | WP_157564779.1 | WP_157567059.1 |
| Bacteroidota | *Mucilaginibacter auburnensis* | WP_100340163.1 | WP_100340579.1 | WP_100342736.1 |
| Bacteroidota | *Mucilaginibacter aurantiaciroseus* | WP_252759152.1 | WP_252758200.1 | WP_252755998.1 |
| Bacteroidota | *Mucilaginibacter boryungensis* | WP_194107392.1 | WP_194105065.1 | WP_194106909.1 |
| Bacteroidota | *Mucilaginibacter calamicampi* | WP_377100119.1 | WP_377096542.1 | WP_377101964.1 |
| Bacteroidota | *Mucilaginibacter celer* | WP_119406908.1 | WP_119410587.1 | WP_119409068.1 |
| Bacteroidota | *Mucilaginibacter conchicola* | WP_117392726.1 | WP_117393950.1 | WP_117394106.1 |
| Bacteroidota | *Mucilaginibacter corticis* | WP_144246735.1 | WP_144249890.1 | WP_144247614.1 |
| Bacteroidota | *Mucilaginibacter daejeonensis* | WP_228170624.1 | WP_228173292.1 | WP_228171161.1 |
| Bacteroidota | *Mucilaginibacter defluvii* | WP_345328736.1 | WP_345331670.1 | WP_345334012.1 |
| Bacteroidota | *Mucilaginibacter dorajii* | WP_259095654.1 | WP_259088923.1 | WP_259092943.1 |

| | | | | |
|---|---|---|---|---|
| Bacteroidota | *Mucilaginibacter endophyticus* | WP_114939907.1 | WP_114936456.1 | WP_114939283.1 |
| Bacteroidota | *Mucilaginibacter flavidus* | WP_252792180.1 | WP_252787898.1 | WP_252790012.1 |
| Bacteroidota | *Mucilaginibacter flavus* | WP_290306153.1 | WP_290305038.1 | WP_290304195.1 |
| Bacteroidota | *Mucilaginibacter frigoritolerans* | WP_144913145.1 | WP_144911351.1 | WP_144914774.1 |
| Bacteroidota | *Mucilaginibacter galii* | WP_188413953.1 | WP_188415640.1 | WP_377169483.1 |
| Bacteroidota | *Mucilaginibacter gilvus* | WP_128535439.1 | WP_128534534.1 | WP_128536213.1 |
| Bacteroidota | *Mucilaginibacter ginkgonis* | WP_157525657.1 | WP_157526112.1 | WP_157525403.1 |
| Bacteroidota | *Mucilaginibacter ginsenosidivorans* | WP_147030842.1 | WP_147033817.1 | WP_147031469.1 |
| Bacteroidota | *Mucilaginibacter ginsenosidivorax* | WP_147054731.1 | WP_147052744.1 | WP_147055663.1 |
| Bacteroidota | *Mucilaginibacter glaciei* | WP_191163871.1 | WP_191164270.1 | WP_191162443.1 |
| Bacteroidota | *Mucilaginibacter gossypiicola* | WP_091215239.1 | WP_091222143.1 | WP_317040865.1 |
| Bacteroidota | *Mucilaginibacter gotjawali* | WP_096349449.1 | WP_096350522.1 | WP_183476404.1 |
| Bacteroidota | *Mucilaginibacter gracilis* | WP_121201357.1 | WP_121198186.1 | WP_121197576.1 |
| Bacteroidota | *Mucilaginibacter gynuensis* | WP_345210895.1 | WP_345213621.1 | WP_345214337.1 |
| Bacteroidota | *Mucilaginibacter hurinus* | WP_114005415.1 | WP_114003385.1 | WP_114005266.1 |
| Bacteroidota | *Mucilaginibacter inviolabilis* | WP_166092465.1 | WP_166089827.1 | WP_166091221.1 |
| Bacteroidota | *Mucilaginibacter jinjuensis* | WP_273630002.1 | WP_273632286.1 | WP_273629463.1 |
| Bacteroidota | *Mucilaginibacter kameinonensis* | WP_121808705.1 | WP_121809185.1 | WP_121808023.1 |
| Bacteroidota | *Mucilaginibacter lacusdianchii* | WP_158825378.1 | WP_158824798.1 | WP_158827222.1 |
| Bacteroidota | *Mucilaginibacter lappiensis* | WP_076371559.1 | WP_076374166.1 | WP_317617672.1 |
| Bacteroidota | *Mucilaginibacter limnophilus* | WP_127707196.1 | WP_127706749.1 | WP_127703902.1 |
| Bacteroidota | *Mucilaginibacter litoreus* | WP_377116600.1 | WP_377111737.1 | WP_377111387.1 |
| Bacteroidota | *Mucilaginibacter lutimaris* | WP_377143437.1 | WP_377145568.1 | WP_377142764.1 |
| Bacteroidota | *Mucilaginibacter mali* | WP_173416735.1 | WP_173414054.1 | WP_173415910.1 |
| Bacteroidota | *Mucilaginibacter mallensis* | WP_091378892.1 | WP_091370817.1 | WP_091376794.1 |
| Bacteroidota | *Mucilaginibacter myungsuensis* | WP_194110220.1 | WP_194112504.1 | WP_194112018.1 |
| Bacteroidota | *Mucilaginibacter oryzae* | WP_109609816.1 | WP_109608412.1 | WP_109609481.1 |
| Bacteroidota | *Mucilaginibacter pallidiroseus* | WP_146380790.1 | WP_146382585.1 | WP_146382325.1 |

| | | | | |
|---|---|---|---|---|
| Bacteroidota | *Mucilaginibacter paludis* | WP_040627802.1 | WP_008511304.1 | WP_008505046.1 |
| Bacteroidota | *Mucilaginibacter panaciglaebae* | WP_345105444.1 | WP_345106671.1 | WP_345104577.1 |
| Bacteroidota | *Mucilaginibacter pankratovii* | WP_191190127.1 | WP_191191809.1 | WP_191191606.1 |
| Bacteroidota | *Mucilaginibacter pedocola* | WP_078349397.1 | WP_078348817.1 | WP_078348021.1 |
| Bacteroidota | *Mucilaginibacter phenanthrenivorans* | WP_258138196.1 | WP_258137176.1 | WP_258138502.1 |
| Bacteroidota | *Mucilaginibacter phyllosphaerae* | WP_134337205.1 | WP_134334969.1 | WP_134335263.1 |
| Bacteroidota | *Mucilaginibacter pineti* | WP_091149399.1 | WP_091145974.1 | WP_091149996.1 |
| Bacteroidota | *Mucilaginibacter pocheonensis* | WP_310095523.1 | WP_310091551.1 | WP_310103339.1 |
| Bacteroidota | *Mucilaginibacter polytrichastri* | WP_074490993.1 | WP_216351114.1 | WP_074490320.1 |
| Bacteroidota | *Mucilaginibacter psychrotolerans* | WP_133233713.1 | WP_133228008.1 | WP_133230210.1 |
| Bacteroidota | *Mucilaginibacter rigui* | WP_191177093.1 | WP_191173972.1 | WP_191173641.1 |
| Bacteroidota | *Mucilaginibacter rivuli* | WP_219489402.1 | WP_219489650.1 | WP_219489433.1 |
| Bacteroidota | *Mucilaginibacter robiniae* | WP_169606372.1 | WP_169605812.1 | WP_169605632.1 |
| Bacteroidota | *Mucilaginibacter roseus* | WP_232179103.1 | WP_232178139.1 | WP_232177772.1 |
| Bacteroidota | *Mucilaginibacter rubeus* | WP_112574782.1 | WP_112567222.1 | WP_112575728.1 |
| Bacteroidota | *Mucilaginibacter sabulilitoris* | WP_321563156.1 | WP_321562246.1 | WP_321562877.1 |
| Bacteroidota | *Mucilaginibacter segetis* | WP_200063558.1 | WP_200063914.1 | WP_200066997.1 |
| Bacteroidota | *Mucilaginibacter sp.* | WP_299359762.1 | WP_343114049.1 | WP_369591216.1 |
| Bacteroidota | *Mucilaginibacter straminoryzae* | WP_245131972.1 | WP_245129948.1 | WP_245129198.1 |
| Bacteroidota | *Mucilaginibacter terrae* | WP_311951396.1 | WP_377161838.1 | WP_311949776.1 |
| Bacteroidota | *Mucilaginibacter terrenus* | WP_117384536.1 | WP_117382517.1 | WP_117382866.1 |
| Bacteroidota | *Mucilaginibacter terrigena* | WP_129878240.1 | WP_129875404.1 | WP_129876957.1 |
| Bacteroidota | *Mucilaginibacter ximonensis* | WP_377184449.1 | WP_377187730.1 | WP_377186185.1 |
| Bacteroidota | *Mucilaginibacter xinganensis* | WP_094572437.1 | WP_094569776.1 | WP_317043860.1 |
| Bacteroidota | *Mucilaginibacter yixingensis* | WP_107827472.1 | WP_107826445.1 | WP_107826477.1 |
| Bacteroidota | *Muricauda lutisoli* | WP_207070618.1 | WP_207070520.1 | WP_207072086.1 |
| Bacteroidota | *Muriicola jejuensis* | WP_163694011.1 | WP_163693910.1 | WP_163691096.1 |
| Bacteroidota | *Muriicola marianensis* | WP_188369167.1 | WP_188369072.1 | WP_188369306.1 |

| | | | | |
|---|---|---|---|---|
| Bacteroidota | *Muriicola soli* | WP_129603256.1 | WP_129603418.1 | WP_129602993.1 |
| Bacteroidota | *Muriicola sp.* | WP_424370009.1 | WP_372944865.1 | WP_372944702.1 |
| Bacteroidota | *Myroides fluvii* | WP_158960886.1 | WP_158963905.1 | WP_158962616.1 |
| Bacteroidota | *Myroides guanonis* | WP_090677506.1 | WP_090677655.1 | WP_090680943.1 |
| Bacteroidota | *Myroides indicus* | WP_133711415.1 | WP_133711710.1 | WP_133712705.1 |
| Bacteroidota | *Myroides injenensis* | WP_010254937.1 | WP_267739598.1 | WP_010255812.1 |
| Bacteroidota | *Myroides marinus* | WP_286405314.1 | WP_038986771.1 | WP_286405152.1 |
| Bacteroidota | *Myroides odoratimimus* | WP_025125853.1 | WP_286386351.1 | WP_286420923.1 |
| Bacteroidota | *Myroides odoratus* | WP_060873191.1 | WP_413511832.1 | WP_002987277.1 |
| Bacteroidota | *Myroides oncorhynchi* | WP_229948831.1 | WP_229948027.1 | WP_229946215.1 |
| Bacteroidota | *Myroides pelagicus* | WP_155035200.1 | WP_155036246.1 | WP_155035270.1 |
| Bacteroidota | *Myroides phaeus* | WP_090406041.1 | WP_090408418.1 | WP_090409129.1 |
| Bacteroidota | *Myroides sp. N17-2* | WP_121965180.1 | WP_121966521.1 | WP_121967414.1 |
| Bacteroidota | *Nafulsella turpanensis* | WP_017731237.1 | WP_017729874.1 | WP_017731273.1 |
| Bacteroidota | *Namhaeicola litoreus* | WP_377178661.1 | WP_377178571.1 | WP_377178062.1 |
| Bacteroidota | *Negadavirga shengliensis* | WP_377060634.1 | WP_377064796.1 | WP_377067651.1 |
| Bacteroidota | *Nemorincola caseinilytica* | WP_345077030.1 | WP_345084461.1 | WP_345085155.1 |
| Bacteroidota | *Neolewinella agarilytica* | WP_090169585.1 | WP_090165148.1 | WP_090169212.1 |
| Bacteroidota | *Neolewinella antarctica* | WP_168039119.1 | WP_168035921.1 | WP_168035849.1 |
| Bacteroidota | *Neolewinella aquimaris* | WP_183495849.1 | WP_183493761.1 | WP_183494136.1 |
| Bacteroidota | *Neolewinella aurantiaca* | WP_147932863.1 | WP_147928689.1 | WP_147932599.1 |
| Bacteroidota | *Neolewinella lacunae* | WP_187464858.1 | WP_187468026.1 | WP_187465779.1 |
| Bacteroidota | *Neolewinella litorea* | WP_136459915.1 | WP_136455929.1 | WP_136456325.1 |
| Bacteroidota | *Neolewinella marina* | WP_099107199.1 | WP_099106183.1 | WP_099104521.1 |
| Bacteroidota | *Neolewinella maritima* | WP_238748977.1 | WP_238752014.1 | WP_238750331.1 |
| Bacteroidota | *Neolewinella persica* | WP_020567182.1 | WP_020567298.1 | WP_020568668.1 |
| Bacteroidota | *Neolewinella sp.* | WP_420458282.1 | WP_420459922.1 | WP_420460317.1 |
| Bacteroidota | *Neolewinella xylanilytica* | WP_104418305.1 | WP_104421724.1 | WP_104420911.1 |

| | | | | |
|---|---|---|---|---|
| Bacteroidota | *Neotamlana laminarinivorans* | WP_226540228.1 | WP_226540714.1 | WP_226544469.1 |
| Bacteroidota | *Neotamlana nanhaiensis* | WP_044626297.1 | WP_044624653.1 | WP_044627344.1 |
| Bacteroidota | *Neotamlana sargassicola* | WP_226695741.1 | WP_226695825.1 | WP_226695554.1 |
| Bacteroidota | *Neotamlana sedimentorum* | WP_044632100.1 | WP_044632017.1 | WP_044633752.1 |
| Bacteroidota | *Neptunitalea chrysea* | WP_281752677.1 | WP_281751884.1 | WP_281754931.1 |
| Bacteroidota | *Neptunitalea lumnitzerae* | WP_281765818.1 | WP_281765697.1 | WP_281766268.1 |
| Bacteroidota | *Niabella agricola* | WP_235554510.1 | WP_235553365.1 | WP_235553023.1 |
| Bacteroidota | *Niabella aurantiaca* | WP_018627122.1 | WP_018630035.1 | WP_018626360.1 |
| Bacteroidota | *Niabella beijingensis* | WP_223708802.1 | WP_223707379.1 | WP_223713486.1 |
| Bacteroidota | *Niabella drilacis* | WP_090389250.1 | WP_090388761.1 | WP_090389126.1 |
| Bacteroidota | *Niabella ginsenosidivorans* | WP_067761319.1 | WP_067752730.1 | WP_067753521.1 |
| Bacteroidota | *Niabella pedocola* | WP_231007196.1 | WP_231004163.1 | WP_231003342.1 |
| Bacteroidota | *Niabella soli* | WP_008587685.1 | WP_008584562.1 | WP_008582146.1 |
| Bacteroidota | *Niabella sp.* | WP_300602989.1 | WP_300602285.1 | WP_300600842.1 |
| Bacteroidota | *Niabella yanshanensis* | WP_114791872.1 | WP_114789308.1 | WP_114789230.1 |
| Bacteroidota | *Niastella caeni* | WP_243751588.1 | WP_136576828.1 | WP_136578394.1 |
| Bacteroidota | *Niastella koreensis* | WP_014223133.1 | WP_014220554.1 | WP_014218826.1 |
| Bacteroidota | *Niastella populi* | WP_081170820.1 | WP_081163880.1 | WP_081168632.1 |
| Bacteroidota | *Niastella sp.* | WP_407743925.1 | WP_407744206.1 | WP_407743428.1 |
| Bacteroidota | *Niastella vici* | WP_081151070.1 | WP_081147991.1 | WP_081155202.1 |
| Bacteroidota | *Niastella yeongjuensis* | WP_081199676.1 | WP_081203469.1 | WP_081202772.1 |
| Bacteroidota | *Nibrella saemangeumensis* | WP_345240722.1 | WP_345243353.1 | WP_345248971.1 |
| Bacteroidota | *Nibrella viscosa* | WP_345269585.1 | WP_345263130.1 | WP_345268198.1 |
| Bacteroidota | *Nibribacter koreensis* | WP_345169332.1 | WP_345162107.1 | WP_345165865.1 |
| Bacteroidota | *Nibribacter ruber* | WP_160690431.1 | WP_160694538.1 | WP_160692977.1 |
| Bacteroidota | *Nitritalea halalkaliphila* | WP_009055316.1 | WP_009057114.1 | WP_009057364.1 |
| Bacteroidota | *Nonlabens agnitus* | WP_105982533.1 | WP_105981802.1 | WP_105981856.1 |
| Bacteroidota | *Nonlabens antarcticus* | WP_194850349.1 | WP_194852000.1 | WP_194850045.1 |

| | | | | |
|---|---|---|---|---|
| Bacteroidota | *Nonlabens arenilitoris* | WP_105071680.1 | WP_245910827.1 | WP_105072145.1 |
| Bacteroidota | *Nonlabens dokdonensis* | WP_015363711.1 | WP_303685940.1 | WP_304017239.1 |
| Bacteroidota | *Nonlabens marinus* | WP_041497035.1 | WP_041494828.1 | WP_041497313.1 |
| Bacteroidota | *Nonlabens ponticola* | WP_126448044.1 | WP_126448506.1 | WP_126448453.1 |
| Bacteroidota | *Nonlabens sp.* | WP_415181852.1 | WP_347196734.1 | WP_347197301.1 |
| Bacteroidota | *Nonlabens spongiae* | WP_085766589.1 | WP_085765672.1 | WP_085766476.1 |
| Bacteroidota | *Nonlabens tegetincola* | WP_085774733.1 | WP_042278472.1 | WP_042278067.1 |
| Bacteroidota | *Nonlabens ulvanivorans* | WP_348021067.1 | WP_317291836.1 | WP_036584679.1 |
| Bacteroidota | *Nonlabens xiamenensis* | WP_124981425.1 | WP_124979846.1 | WP_240642324.1 |
| Bacteroidota | *Nubsella zeaxanthinifaciens* | WP_113637362.1 | WP_427529302.1 | WP_113637772.1 |
| Bacteroidota | *Oceanihabitans sediminis* | WP_113966153.1 | WP_072351629.1 | WP_113966430.1 |
| Bacteroidota | *Oceanihabitans sp. 2_MG-2023* | WP_303425751.1 | WP_303424129.1 | WP_303424880.1 |
| Bacteroidota | *Ochrovirga pacifica* | WP_010136703.1 | WP_010135397.1 | WP_010134725.1 |
| Bacteroidota | *Ohtaekwangia koreensis* | WP_079687037.1 | WP_079688647.1 | WP_079688007.1 |
| Bacteroidota | *Ohtaekwangia kribbensis* | WP_377580141.1 | WP_377585692.1 | WP_377579531.1 |
| Bacteroidota | *Ohtaekwangia sp.* | WP_331979302.1 | WP_331959849.1 | WP_331964706.1 |
| Bacteroidota | *Olivibacter domesticus* | WP_093328473.1 | WP_093321673.1 | WP_317040863.1 |
| Bacteroidota | *Olivibacter ginsenosidimutans* | WP_345230593.1 | WP_345234589.1 | WP_345232325.1 |
| Bacteroidota | *Olivibacter sitiensis* | WP_028297179.1 | WP_028295449.1 | WP_028296240.1 |
| Bacteroidota | *Olivibacter sp. XZL3* | WP_134089489.1 | WP_134089330.1 | WP_134088390.1 |
| Bacteroidota | *Olleya aquimaris* | WP_111659383.1 | WP_111659072.1 | WP_111660549.1 |
| Bacteroidota | *Olleya marilimosa* | WP_028282215.1 | WP_028282949.1 | WP_191101322.1 |
| Bacteroidota | *Olleya namhaensis* | WP_281848004.1 | WP_272022623.1 | WP_272023392.1 |
| Bacteroidota | *Olleya sp. YS* | WP_279301442.1 | WP_279301861.1 | WP_279301014.1 |
| Bacteroidota | *Ornithobacterium rhinotracheale* | WP_153828604.1 | WP_247117320.1 | WP_413371110.1 |
| Bacteroidota | *Owenweeksia hongkongensis* | WP_417609137.1 | WP_417591921.1 | WP_417589851.1 |
| Bacteroidota | *Paenimyroides aestuarii* | WP_257498498.1 | WP_257498436.1 | WP_257498655.1 |
| Bacteroidota | *Paenimyroides baculatum* | WP_150014147.1 | WP_150014239.1 | WP_150011125.1 |

| | | | | |
|---|---|---|---|---|
| Bacteroidota | *Paenimyroides ceti* | WP_290362695.1 | WP_290363368.1 | WP_290363635.1 |
| Bacteroidota | *Paenimyroides marinum* | WP_091095377.1 | WP_091095373.1 | WP_091095038.1 |
| Bacteroidota | *Paenimyroides tangerinum* | WP_125018708.1 | WP_125017044.1 | WP_125017160.1 |
| Bacteroidota | *Paenimyroides ummariense* | WP_091522971.1 | WP_091526027.1 | WP_091521721.1 |
| Bacteroidota | *Paenimyroides viscosum* | WP_124899808.1 | WP_124898830.1 | WP_124898761.1 |
| Bacteroidota | *Panacibacter ginsenosidivorans* | WP_147189023.1 | WP_147192662.1 | WP_147188223.1 |
| Bacteroidota | *Panacibacter microcysteis* | WP_196990551.1 | WP_196991319.1 | WP_196991932.1 |
| Bacteroidota | *Parachryseolinea silvisoli* | WP_267298128.1 | WP_267296791.1 | WP_267293654.1 |
| Bacteroidota | *Paracnuella aquatica* | WP_114236417.1 | WP_241558418.1 | WP_114236187.1 |
| Bacteroidota | *Paracrocinitomix mangrovi* | WP_221834572.1 | WP_221834850.1 | WP_221836018.1 |
| Bacteroidota | *Paradesertivirga mongoliensis* | WP_255897988.1 | WP_255902703.1 | WP_255902177.1 |
| Bacteroidota | *Parafilimonas sp.* | WP_415161251.1 | WP_415157779.1 | WP_415157992.1 |
| Bacteroidota | *Parafilimonas terrae* | WP_090655776.1 | WP_245751513.1 | WP_090658732.1 |
| Bacteroidota | *Paraflavisolibacter caeni* | WP_279295855.1 | WP_279299632.1 | WP_279297706.1 |
| Bacteroidota | *Paraflavisolibacter sp. H34* | WP_336722650.1 | WP_336725373.1 | WP_336725333.1 |
| Bacteroidota | *Paraflavitalea devenefica* | WP_167287404.1 | WP_167290942.1 | WP_167291895.1 |
| Bacteroidota | *Paraflavitalea pollutisoli* | WP_276480049.1 | WP_276486104.1 | WP_276485702.1 |
| Bacteroidota | *Paraflavitalea soli* | WP_119051981.1 | WP_119050955.1 | WP_119050606.1 |
| Bacteroidota | *Paraflavitalea sp. CAU 1676* | WP_275834834.1 | WP_275832317.1 | WP_275834035.1 |
| Bacteroidota | *Parapedobacter composti* | WP_090970143.1 | WP_090972664.1 | WP_090971754.1 |
| Bacteroidota | *Parapedobacter defluvii* | WP_353197832.1 | WP_188753847.1 | WP_353195674.1 |
| Bacteroidota | *Parapedobacter deserti* | WP_379023536.1 | WP_379021282.1 | WP_379020811.1 |
| Bacteroidota | *Parapedobacter indicus* | WP_090627862.1 | WP_090624180.1 | WP_090626682.1 |
| Bacteroidota | *Parapedobacter koreensis* | WP_090608055.1 | WP_090608876.1 | WP_090609168.1 |
| Bacteroidota | *Parapedobacter lycopersici* | WP_353181428.1 | WP_353181923.1 | WP_257658627.1 |
| Bacteroidota | *Parapedobacter pyrenivorans* | WP_188504305.1 | WP_353127354.1 | WP_188507784.1 |
| Bacteroidota | *Parapedobacter soli* | WP_262246792.1 | WP_262248233.1 | WP_262245092.1 |
| Bacteroidota | *Parapedobacter sp.* | WP_325635624.1 | WP_325636015.1 | WP_325637971.1 |

| | | | | |
|---|---|---|---|---|
| Bacteroidota | *Parapedobacter tibetensis* | WP_257666263.1 | WP_257667877.1 | WP_257667435.1 |
| Bacteroidota | *Pararcticibacter amylolyticus* | WP_109416979.1 | WP_109414587.1 | WP_109416868.1 |
| Bacteroidota | *Pararhodonellum marinum* | WP_194774176.1 | WP_194775591.1 | WP_194774932.1 |
| Bacteroidota | *Parasediminibacterium paludis* | WP_379013407.1 | WP_379014846.1 | WP_379012157.1 |
| Bacteroidota | *Parasediminibacterium sp. JCM 36343* | WP_410844105.1 | WP_410844722.1 | WP_410847215.1 |
| Bacteroidota | *Parasegetibacter sp. NRK P23* | WP_251761400.1 | WP_251758786.1 | WP_251759009.1 |
| Bacteroidota | *Pareuzebyella sediminis* | WP_149277078.1 | WP_149275542.1 | WP_149277551.1 |
| Bacteroidota | *Parvicella tangerina* | WP_258541382.1 | WP_258543270.1 | WP_258541479.1 |
| Bacteroidota | *Patiriisocius hiemis* | WP_311331756.1 | WP_311331402.1 | WP_311331573.1 |
| Bacteroidota | *Patiriisocius marinistellae* | WP_151894385.1 | WP_151894039.1 | WP_151894628.1 |
| Bacteroidota | *Patiriisocius marinus* | WP_151673679.1 | WP_151673105.1 | WP_151673038.1 |
| Bacteroidota | *Patiriisocius sp. Uisw_047* | WP_415371185.1 | WP_415370858.1 | WP_415370868.1 |
| Bacteroidota | *Paucihalobacter ruber* | WP_140990425.1 | WP_140990909.1 | WP_140990640.1 |
| Bacteroidota | *Paucihalobacter sp.* | WP_334112096.1 | WP_334113206.1 | WP_334111572.1 |
| Bacteroidota | *Pedobacter africanus* | WP_084237760.1 | WP_084240834.1 | WP_310143232.1 |
| Bacteroidota | *Pedobacter agri* | WP_316805750.1 | WP_029204146.1 | WP_307292130.1 |
| Bacteroidota | *Pedobacter albus* | WP_330109376.1 | WP_330105924.1 | WP_330107668.1 |
| Bacteroidota | *Pedobacter alluvionis* | WP_121287746.1 | WP_121287178.1 | WP_121283637.1 |
| Bacteroidota | *Pedobacter alpinus* | WP_379048161.1 | WP_379040262.1 | WP_379047338.1 |
| Bacteroidota | *Pedobacter antarcticus* | WP_219224099.1 | WP_219226882.1 | WP_074591307.1 |
| Bacteroidota | *Pedobacter aquae* | WP_149074866.1 | WP_149073748.1 | WP_149074685.1 |
| Bacteroidota | *Pedobacter aquatilis* | WP_290241275.1 | WP_316830188.1 | WP_316734565.1 |
| Bacteroidota | *Pedobacter arcticus* | WP_017256750.1 | WP_017259123.1 | WP_040539760.1 |
| Bacteroidota | *Pedobacter borealis* | WP_029280013.1 | WP_029278913.1 | WP_029276083.1 |
| Bacteroidota | *Pedobacter boryungensis* | WP_173272792.1 | WP_173273727.1 | WP_173273906.1 |
| Bacteroidota | *Pedobacter caeni* | WP_073232208.1 | WP_073234076.1 | WP_073229707.1 |
| Bacteroidota | *Pedobacter changchengzhani* | WP_133263516.1 | WP_133262326.1 | WP_133261648.1 |
| Bacteroidota | *Pedobacter chinensis* | WP_115402615.1 | WP_115404077.1 | WP_115404238.1 |

| | | | | |
|---|---|---|---|---|
| Bacteroidota | *Pedobacter chitinilyticus* | WP_113648051.1 | WP_113645262.1 | WP_113647052.1 |
| Bacteroidota | *Pedobacter cryoconitis* | WP_183881087.1 | WP_068402913.1 | WP_068406289.1 |
| Bacteroidota | *Pedobacter cryophilus* | WP_136825249.1 | WP_136826977.1 | WP_317130548.1 |
| Bacteroidota | *Pedobacter cryotolerans* | WP_136874354.1 | WP_136876996.1 | WP_136878258.1 |
| Bacteroidota | *Pedobacter deserti* | WP_256001905.1 | WP_256007584.1 | WP_256007120.1 |
| Bacteroidota | *Pedobacter duraquae* | WP_133556412.1 | WP_133553364.1 | WP_133555384.1 |
| Bacteroidota | *Pedobacter endophyticus* | WP_196100950.1 | WP_196099093.1 | WP_196100596.1 |
| Bacteroidota | *Pedobacter faecalis* | WP_285010636.1 | WP_285011385.1 | WP_285010782.1 |
| Bacteroidota | *Pedobacter fastidiosus* | WP_187069658.1 | WP_187071857.1 | WP_187070164.1 |
| Bacteroidota | *Pedobacter flavus* | WP_330144997.1 | WP_330145864.1 | WP_330145931.1 |
| Bacteroidota | *Pedobacter foliorum* | WP_173090079.1 | WP_173093305.1 | WP_173091865.1 |
| Bacteroidota | *Pedobacter frigidisoli* | WP_131560922.1 | WP_131561206.1 | WP_316800050.1 |
| Bacteroidota | *Pedobacter frigiditerrae* | WP_316768949.1 | WP_131554479.1 | WP_316772800.1 |
| Bacteroidota | *Pedobacter frigoris* | WP_136835606.1 | WP_136837513.1 | WP_316791726.1 |
| Bacteroidota | *Pedobacter gandavensis* | WP_316842888.1 | WP_182960205.1 | WP_316840536.1 |
| Bacteroidota | *Pedobacter ghigonis* | WP_175636134.1 | WP_175633854.1 | WP_175636440.1 |
| Bacteroidota | *Pedobacter ginsengisoli* | WP_099440654.1 | WP_285056991.1 | WP_285054377.1 |
| Bacteroidota | *Pedobacter glucosidilyticus* | WP_026905129.1 | WP_304064693.1 | WP_026902661.1 |
| Bacteroidota | *Pedobacter hartonius* | WP_090555335.1 | WP_090555089.1 | WP_090556570.1 |
| Bacteroidota | *Pedobacter helvus* | WP_138729967.1 | WP_138730109.1 | WP_138728912.1 |
| Bacteroidota | *Pedobacter heparinus* | WP_012780673.1 | WP_012781370.1 | WP_316812515.1 |
| Bacteroidota | *Pedobacter hiemivivus* | WP_131610061.1 | WP_136881953.1 | WP_131607297.1 |
| Bacteroidota | *Pedobacter indicus* | WP_118193771.1 | WP_118196262.1 | WP_118193992.1 |
| Bacteroidota | *Pedobacter insulae* | WP_090996882.1 | WP_090992110.1 | WP_090996648.1 |
| Bacteroidota | *Pedobacter jamesrossensis* | WP_378962165.1 | WP_378958657.1 | WP_378962518.1 |
| Bacteroidota | *Pedobacter jejuensis* | WP_123204999.1 | WP_123205102.1 | WP_123206726.1 |
| Bacteroidota | *Pedobacter jeongneungensis* | WP_344853497.1 | WP_344850380.1 | WP_344852960.1 |
| Bacteroidota | *Pedobacter kyungheensis* | WP_039479543.1 | WP_039479903.1 | WP_039479944.1 |

| | | | | |
|---|---|---|---|---|
| Bacteroidota | *Pedobacter lithocola* | WP_378987507.1 | WP_378981496.1 | WP_378984310.1 |
| Bacteroidota | *Pedobacter lusitanus* | WP_041883719.1 | WP_041885191.1 | WP_041887252.1 |
| Bacteroidota | *Pedobacter mendelii* | WP_188411375.1 | WP_188412809.1 | WP_188415496.1 |
| Bacteroidota | *Pedobacter metabolipauper* | WP_133577613.1 | WP_133577989.1 | WP_133577171.1 |
| Bacteroidota | *Pedobacter miscanthi* | WP_316824526.1 | WP_113952201.1 | WP_316827092.1 |
| Bacteroidota | *Pedobacter montanisoli* | WP_243360292.1 | WP_243363041.1 | WP_243360692.1 |
| Bacteroidota | *Pedobacter mucosus* | WP_238414581.1 | WP_238414089.1 | WP_238414324.1 |
| Bacteroidota | *Pedobacter namyangjuensis* | WP_113654293.1 | WP_113653838.1 | WP_113654129.1 |
| Bacteroidota | *Pedobacter nanyangensis* | WP_113662147.1 | WP_113662675.1 | WP_113661786.1 |
| Bacteroidota | *Pedobacter nototheniae* | WP_316801765.1 | WP_131538947.1 | WP_131535627.1 |
| Bacteroidota | *Pedobacter nutrimenti* | WP_110833957.1 | WP_110834448.1 | WP_110831824.1 |
| Bacteroidota | *Pedobacter nyackensis* | WP_316816358.1 | WP_084286656.1 | WP_084289426.1 |
| Bacteroidota | *Pedobacter paludis* | WP_109930788.1 | WP_109932520.1 | WP_109931200.1 |
| Bacteroidota | *Pedobacter panaciterrae* | WP_172661574.1 | WP_172659420.1 | WP_337717474.1 |
| Bacteroidota | *Pedobacter petrophilus* | WP_154281502.1 | WP_154282467.1 | WP_154278704.1 |
| Bacteroidota | *Pedobacter planticolens* | WP_182922361.1 | WP_182922156.1 | WP_182920612.1 |
| Bacteroidota | *Pedobacter polaris* | WP_136838023.1 | WP_136844270.1 | WP_136841447.1 |
| Bacteroidota | *Pedobacter psychrodurus* | WP_131533002.1 | WP_131534490.1 | WP_131530597.1 |
| Bacteroidota | *Pedobacter psychrophilus* | WP_068822602.1 | WP_068823604.1 | WP_068822891.1 |
| Bacteroidota | *Pedobacter psychroterrae* | WP_131594357.1 | WP_131597463.1 | WP_131597727.1 |
| Bacteroidota | *Pedobacter psychrotolerans* | WP_132530194.1 | WP_132536433.1 | WP_132529721.1 |
| Bacteroidota | *Pedobacter punctiformis* | WP_269428406.1 | WP_269427832.1 | WP_269428613.1 |
| Bacteroidota | *Pedobacter puniceum* | WP_154288091.1 | WP_154286091.1 | WP_154288349.1 |
| Bacteroidota | *Pedobacter quisquiliarum* | WP_188626080.1 | WP_188627188.1 | WP_188624811.1 |
| Bacteroidota | *Pedobacter rhizosphaerae* | WP_090884952.1 | WP_090881040.1 | WP_090883636.1 |
| Bacteroidota | *Pedobacter rhodius* | WP_269417121.1 | WP_269413907.1 | WP_269416549.1 |
| Bacteroidota | *Pedobacter roseus* | WP_187592982.1 | WP_187591282.1 | WP_187591487.1 |
| Bacteroidota | *Pedobacter sandarakinus* | WP_265854285.1 | WP_265855752.1 | WP_265854120.1 |

| | | | | |
|---|---|---|---|---|
| Bacteroidota | *Pedobacter schmidteae* | WP_121269434.1 | WP_121273731.1 | WP_121271022.1 |
| Bacteroidota | *Pedobacter segetis* | WP_200587972.1 | WP_200585255.1 | WP_200586207.1 |
| Bacteroidota | *Pedobacter soli* | WP_090772814.1 | WP_090772601.1 | WP_090766432.1 |
| Bacteroidota | *Pedobacter sp.* | WP_421938346.1 | WP_343522411.1 | WP_353903901.1 |
| Bacteroidota | *Pedobacter steynii* | WP_069379562.1 | WP_074606995.1 | WP_069378909.1 |
| Bacteroidota | *Pedobacter suwonensis* | WP_145857173.1 | WP_293741410.1 | WP_145860125.1 |
| Bacteroidota | *Pedobacter terrae* | WP_335317504.1 | WP_335320483.1 | WP_335318463.1 |
| Bacteroidota | *Pedobacter ureilyticus* | WP_138722568.1 | WP_138723946.1 | WP_138721851.1 |
| Bacteroidota | *Pedobacter vanadiisoli* | WP_379081851.1 | WP_379080222.1 | WP_379081366.1 |
| Bacteroidota | *Pedobacter westerhofensis* | WP_142528417.1 | WP_142529467.1 | WP_142530983.1 |
| Bacteroidota | *Pedobacter xixiisoli* | WP_097132640.1 | WP_097127343.1 | WP_097132993.1 |
| Bacteroidota | *Pedobacter yonginense* | WP_109925110.1 | WP_109927344.1 | WP_109926179.1 |
| Bacteroidota | *Pedobacter yulinensis* | WP_107214288.1 | WP_107216326.1 | WP_107216633.1 |
| Bacteroidota | *Pedobacter zeae* | WP_183765082.1 | WP_183759401.1 | WP_183768407.1 |
| Bacteroidota | *Pelagihabitans pacificus* | WP_152575596.1 | WP_152574733.1 | WP_152575827.1 |
| Bacteroidota | *Pelobium manganitolerans* | WP_120181251.1 | WP_120181798.1 | WP_120181363.1 |
| Bacteroidota | *Penaeicola halotolerans* | WP_226390126.1 | WP_226389474.1 | WP_226390042.1 |
| Bacteroidota | *Persicitalea jodogahamensis* | WP_189563363.1 | WP_189566008.1 | WP_189562809.1 |
| Bacteroidota | *Persicitalea sp.* | WP_373514763.1 | WP_373514327.1 | WP_373511538.1 |
| Bacteroidota | *Persicobacter diffluens* | WP_338237395.1 | WP_338236170.1 | WP_338235858.1 |
| Bacteroidota | *Persicobacter psychrovividus* | WP_332922396.1 | WP_338397812.1 | WP_338397565.1 |
| Bacteroidota | *Persicobacter sp. CCB-QB2* | WP_053406831.1 | WP_053404325.1 | WP_053405934.1 |
| Bacteroidota | *Phaeocystidibacter luteus* | WP_151668120.1 | WP_151668392.1 | WP_170266297.1 |
| Bacteroidota | *Phaeocystidibacter marisrubri* | WP_151691796.1 | WP_151692856.1 | WP_151692644.1 |
| Bacteroidota | *Phaeodactylibacter luteus* | WP_147168864.1 | WP_147166612.1 | WP_147167063.1 |
| Bacteroidota | *Phaeodactylibacter sp.* | WP_350176227.1 | WP_350177840.1 | WP_293564349.1 |
| Bacteroidota | *Phaeodactylibacter xiamenensis* | WP_044220771.1 | WP_345894640.1 | WP_421948296.1 |
| Bacteroidota | Phnomibacter ginsenosidimutans | WP_157479401.1 | WP_157478826.1 | WP_157478517.1 |

| | | | | |
|---|---|---|---|---|
| Bacteroidota | *Phnomibacter sp.* | WP_333801136.1 | WP_333798932.1 | WP_333799269.1 |
| Bacteroidota | *Pinibacter aurantiacus* | WP_217789871.1 | WP_217794396.1 | WP_217791097.1 |
| Bacteroidota | *Pinibacter soli* | WP_282334310.1 | WP_282334518.1 | WP_282335433.1 |
| Bacteroidota | *Planktosalinus lacus* | WP_188441418.1 | WP_188443026.1 | WP_188439440.1 |
| Bacteroidota | *Planobacterium oryzisoli* | WP_194739118.1 | WP_194739626.1 | WP_194738760.1 |
| Bacteroidota | *Pleomorphovibrio marinus* | WP_114751276.1 | WP_114750181.1 | WP_114747671.1 |
| Bacteroidota | *Polaribacter aestuariivivens* | WP_138534867.1 | WP_138535111.1 | WP_138537235.1 |
| Bacteroidota | *Polaribacter aquimarinus* | WP_109404621.1 | WP_109405612.1 | WP_109404950.1 |
| Bacteroidota | *Polaribacter atrinae* | WP_068452971.1 | WP_068451824.1 | WP_341222082.1 |
| Bacteroidota | *Polaribacter batillariae* | WP_207972030.1 | WP_207972378.1 | WP_207971661.1 |
| Bacteroidota | *Polaribacter butkevichii* | WP_105048550.1 | WP_105048114.1 | WP_105048036.1 |
| Bacteroidota | *Polaribacter cellanae* | WP_208079180.1 | WP_208079597.1 | WP_208079626.1 |
| Bacteroidota | *Polaribacter dokdonensis* | WP_053974708.1 | WP_053974658.1 | WP_053975070.1 |
| Bacteroidota | *Polaribacter filamentus* | WP_104809256.1 | WP_104808805.1 | WP_104808646.1 |
| Bacteroidota | *Polaribacter gangjinensis* | WP_105045548.1 | WP_105047271.1 | WP_105047223.1 |
| Bacteroidota | *Polaribacter glomeratus* | WP_105020554.1 | WP_105020177.1 | WP_105020891.1 |
| Bacteroidota | *Polaribacter haliotis* | WP_088353541.1 | WP_088353291.1 | WP_088352880.1 |
| Bacteroidota | *Polaribacter huanghezhanensis* | WP_301400243.1 | WP_301400130.1 | WP_301400796.1 |
| Bacteroidota | *Polaribacter irgensii* | WP_026288371.1 | WP_018942509.1 | WP_004570425.1 |
| Bacteroidota | *Polaribacter litorisediminis* | WP_223446044.1 | WP_223446128.1 | WP_223439388.1 |
| Bacteroidota | *Polaribacter marinivivus* | WP_377410213.1 | WP_377410502.1 | WP_377409873.1 |
| Bacteroidota | *Polaribacter marinus* | WP_242177238.1 | WP_242177468.1 | WP_242176929.1 |
| Bacteroidota | *Polaribacter pacificus* | WP_188598849.1 | WP_188598986.1 | WP_188599336.1 |
| Bacteroidota | *Polaribacter pectinis* | WP_187483341.1 | WP_187483380.1 | WP_187482972.1 |
| Bacteroidota | *Polaribacter ponticola* | WP_265724554.1 | WP_265724573.1 | WP_274270124.1 |
| Bacteroidota | *Polaribacter porphyrae* | WP_105014864.1 | WP_105014974.1 | WP_105015495.1 |
| Bacteroidota | *Polaribacter reichenbachii* | WP_068360224.1 | WP_068360040.1 | WP_068364870.1 |
| Bacteroidota | *Polaribacter sejongensis* | WP_208889400.1 | WP_208888940.1 | WP_208888764.1 |

| | | | | |
|---|---|---|---|---|
| Bacteroidota | *Polaribacter septentrionalilitoris* | WP_159947352.1 | WP_159947839.1 | WP_159948279.1 |
| Bacteroidota | *Polaribacter sp.* | WP_422162747.1 | WP_422162769.1 | WP_298948141.1 |
| Bacteroidota | *Polaribacter staleyi* | WP_343329764.1 | WP_343329481.1 | WP_343329414.1 |
| Bacteroidota | *Polaribacter tangerinus* | WP_088322902.1 | WP_088322791.1 | WP_088322699.1 |
| Bacteroidota | *Polaribacter vadi* | WP_065318254.1 | WP_339883837.1 | WP_339884802.1 |
| Bacteroidota | *Pollutibacter soli* | WP_336516241.1 | WP_336514272.1 | WP_336514248.1 |
| Bacteroidota | *Polluticaenibacter yanchengensis* | WP_407031566.1 | WP_407030962.1 | WP_407032955.1 |
| Bacteroidota | *Polluticoccus soli* | WP_276134619.1 | WP_276134772.1 | WP_276132203.1 |
| Bacteroidota | *Pontibacter actiniarum* | WP_025607855.1 | WP_025605102.1 | WP_025608381.1 |
| Bacteroidota | *Pontibacter akesuensis* | WP_068837038.1 | WP_068839774.1 | WP_082815155.1 |
| Bacteroidota | *Pontibacter amylolyticus* | WP_188502506.1 | WP_188499622.1 | WP_188500582.1 |
| Bacteroidota | *Pontibacter anaerobius* | WP_266051738.1 | WP_266054022.1 | WP_266054192.1 |
| Bacteroidota | *Pontibacter aquaedesilientis* | WP_191182936.1 | WP_191182164.1 | WP_224744371.1 |
| Bacteroidota | *Pontibacter arcticus* | WP_112306243.1 | WP_112305500.1 | WP_239020921.1 |
| Bacteroidota | *Pontibacter aydingkolensis* | WP_219876295.1 | WP_219878462.1 | WP_246596896.1 |
| Bacteroidota | *Pontibacter beigongshangensis* | WP_187260967.1 | WP_187260582.1 | WP_187264052.1 |
| Bacteroidota | *Pontibacter burrus* | WP_163915162.1 | WP_163914727.1 | WP_163910788.1 |
| Bacteroidota | *Pontibacter cellulosilyticus* | WP_187067611.1 | WP_187065817.1 | WP_230407259.1 |
| Bacteroidota | *Pontibacter chinhatensis* | WP_092099925.1 | WP_092098427.1 | WP_092101233.1 |
| Bacteroidota | *Pontibacter chitinilyticus* | WP_347159781.1 | WP_347157039.1 | WP_347157394.1 |
| Bacteroidota | *Pontibacter diazotrophicus* | WP_115563627.1 | WP_115565944.1 | WP_246000815.1 |
| Bacteroidota | *Pontibacter fetidus* | WP_162345166.1 | WP_162344390.1 | WP_162346564.1 |
| Bacteroidota | *Pontibacter flavimaris* | WP_170866119.1 | WP_073851109.1 | WP_073851885.1 |
| Bacteroidota | *Pontibacter harenae* | WP_229972779.1 | WP_229969948.1 | WP_229968909.1 |
| Bacteroidota | *Pontibacter indicus* | WP_170871914.1 | WP_076666330.1 | WP_244554682.1 |
| Bacteroidota | *Pontibacter kalidii* | WP_266202819.1 | WP_266205387.1 | WP_266203411.1 |
| Bacteroidota | *Pontibacter korlensis* | WP_046308896.1 | WP_046312400.1 | WP_084694729.1 |
| Bacteroidota | *Pontibacter liquoris* | WP_242916368.1 | WP_242918685.1 | WP_242921285.1 |

| | | | | |
|---|---|---|---|---|
| Bacteroidota | *Pontibacter litorisediminis* | WP_276499133.1 | WP_276495720.1 | WP_276498631.1 |
| Bacteroidota | *Pontibacter locisalis* | WP_377507272.1 | WP_377512209.1 | WP_377503573.1 |
| Bacteroidota | *Pontibacter mangrovi* | WP_140621972.1 | WP_140618665.1 | WP_239022830.1 |
| Bacteroidota | *Pontibacter mucosus* | WP_108212430.1 | WP_108213444.1 | WP_108210166.1 |
| Bacteroidota | *Pontibacter oryzae* | WP_119430834.1 | WP_119431269.1 | WP_119432100.1 |
| Bacteroidota | *Pontibacter pamirensis* | WP_162053720.1 | WP_162052484.1 | WP_237144869.1 |
| Bacteroidota | *Pontibacter populi* | WP_350410730.1 | WP_350412632.1 | WP_350411634.1 |
| Bacteroidota | *Pontibacter pudoricolor* | WP_162428816.1 | WP_162426266.1 | WP_162427917.1 |
| Bacteroidota | *Pontibacter qinzhouensis* | WP_147922611.1 | WP_147923662.1 | WP_255474196.1 |
| Bacteroidota | *Pontibacter ramchanderi* | WP_101446093.1 | WP_101442876.1 | WP_180336380.1 |
| Bacteroidota | *Pontibacter roseus* | WP_211214358.1 | WP_018478075.1 | WP_018476588.1 |
| Bacteroidota | *Pontibacter ruber* | WP_250431823.1 | WP_250429605.1 | WP_250428291.1 |
| Bacteroidota | *Pontibacter rugosus* | WP_377526565.1 | WP_377529224.1 | WP_377526710.1 |
| Bacteroidota | *Pontibacter russatus* | WP_161890271.1 | WP_161887828.1 | WP_237586605.1 |
| Bacteroidota | *Pontibacter saemangeumensis* | WP_345161860.1 | WP_345157442.1 | WP_345159741.1 |
| Bacteroidota | *Pontibacter silvestris* | WP_229962237.1 | WP_229958046.1 | WP_229960458.1 |
| Bacteroidota | *Pontibacter sp. HSC-36F09* | WP_253402509.1 | WP_253405866.1 | WP_253401128.1 |
| Bacteroidota | *Pontibacter toksunensis* | WP_377487128.1 | WP_377479545.1 | WP_377485187.1 |
| Bacteroidota | *Pontibacter ummariensis* | WP_089319534.1 | WP_089317724.1 | WP_089319831.1 |
| Bacteroidota | *Pontibacter virosus* | WP_207774867.1 | WP_116543526.1 | WP_116541785.1 |
| Bacteroidota | *Pontibacter vulgaris* | WP_242927759.1 | WP_242928228.1 | WP_242926450.1 |
| Bacteroidota | *Pontimicrobium aquaticum* | WP_136844924.1 | WP_136842894.1 | WP_136842336.1 |
| Bacteroidota | *Pontimicrobium sp. IMCC45349* | WP_412984909.1 | WP_412985655.1 | WP_412984315.1 |
| Bacteroidota | *Porifericola rhodea* | WP_302241969.1 | WP_302238587.1 | WP_302241977.1 |
| Bacteroidota | *Poritiphilus flavus* | WP_161435387.1 | WP_161436029.1 | WP_161435034.1 |
| Bacteroidota | *Portibacter lacus* | WP_235295426.1 | WP_235292611.1 | WP_235293306.1 |
| Bacteroidota | *Portibacter marinus* | WP_235298559.1 | WP_235296124.1 | WP_235298989.1 |
| Bacteroidota | *Postechiella marina* | WP_344786197.1 | WP_344786381.1 | WP_344789558.1 |

| | | | | |
|---|---|---|---|---|
| Bacteroidota | *Pricia antarctica* | WP_091870366.1 | WP_091870156.1 | WP_091868748.1 |
| Bacteroidota | *Pricia mediterranea* | WP_314016435.1 | WP_314016377.1 | WP_314012617.1 |
| Bacteroidota | *Pricia sp.* | WP_373518980.1 | WP_373518823.1 | WP_373517750.1 |
| Bacteroidota | *Profundicola chukchiensis* | WP_304420887.1 | WP_304420718.1 | WP_304416377.1 |
| Bacteroidota | *Pseudalgibacter alginicilyticus* | WP_054724859.1 | WP_054724689.1 | WP_054725144.1 |
| Bacteroidota | *Pseudarcicella hirudinis* | WP_092018740.1 | WP_092016199.1 | WP_092018609.1 |
| Bacteroidota | *Pseudobacter ginsenosidimutans* | WP_130542873.1 | WP_130543804.1 | WP_130540330.1 |
| Bacteroidota | *Pseudochryseolinea flava* | WP_112748661.1 | WP_112749846.1 | WP_112745532.1 |
| Bacteroidota | *Pseudocnuella soli* | WP_132050710.1 | WP_240040623.1 | WP_132055477.1 |
| Bacteroidota | *Pseudoflavitalea rhizosphaerae* | WP_127128830.1 | WP_127132125.1 | WP_127125031.1 |
| Bacteroidota | *Pseudoflavitalea sp. G-6-1-2* | WP_169423241.1 | WP_169427293.1 | WP_169427852.1 |
| Bacteroidota | *Pseudofulvibacter geojedonensis* | WP_377715660.1 | WP_377712568.1 | WP_377713360.1 |
| Bacteroidota | *Pseudopedobacter beijingensis* | WP_379661639.1 | WP_379663639.1 | WP_379660747.1 |
| Bacteroidota | *Pseudopedobacter saltans* | WP_013634471.1 | WP_013633057.1 | WP_013631349.1 |
| Bacteroidota | *Pseudopedobacter sp.* | WP_353133078.1 | WP_353136078.1 | WP_353135823.1 |
| Bacteroidota | *Pseudotamlana agarivorans* | WP_067147999.1 | WP_216068537.1 | WP_216068325.1 |
| Bacteroidota | *Pseudotamlana carrageenivorans* | WP_102996535.1 | WP_102996350.1 | WP_102996863.1 |
| Bacteroidota | *Pseudotamlana haliotis* | WP_150935676.1 | WP_150937802.1 | WP_150940490.1 |
| Bacteroidota | *Pseudotenacibaculum haliotis* | WP_379666268.1 | WP_379666551.1 | WP_379664548.1 |
| Bacteroidota | *Pseudotenacibaculum sp. MALMAid0570* | WP_349553908.1 | WP_349553590.1 | WP_349554108.1 |
| Bacteroidota | *Pseudozobellia sp. WGM2* | WP_209399679.1 | WP_209403114.1 | WP_209405363.1 |
| Bacteroidota | *Pseudozobellia thermophila* | WP_072994189.1 | WP_072994881.1 | WP_072995245.1 |
| Bacteroidota | *Psychroflexus aestuariivivens* | WP_127844959.1 | WP_127845023.1 | WP_127846452.1 |
| Bacteroidota | *Psychroflexus aurantiacus* | WP_164005076.1 | WP_164004848.1 | WP_164004585.1 |
| Bacteroidota | *Psychroflexus curvus* | WP_224486199.1 | WP_224509615.1 | WP_224487413.1 |
| Bacteroidota | *Psychroflexus gondwanensis* | WP_003438426.1 | WP_003442182.1 | WP_003440994.1 |
| Bacteroidota | *Psychroflexus halocasei* | WP_093239311.1 | WP_093241794.1 | WP_093244459.1 |
| Bacteroidota | *Psychroflexus lacisalsi* | WP_224454393.1 | WP_224454992.1 | WP_224454775.1 |

| | | | | |
|---|---|---|---|---|
| Bacteroidota | *Psychroflexus longus* | WP_224461640.1 | WP_224461157.1 | WP_224461978.1 |
| Bacteroidota | *Psychroflexus maritimus* | WP_166400947.1 | WP_166400890.1 | WP_166399151.1 |
| Bacteroidota | *Psychroflexus montanilacus* | WP_224462274.1 | WP_224465188.1 | WP_224464209.1 |
| Bacteroidota | *Psychroflexus planctonicus* | WP_188457486.1 | WP_188458122.1 | WP_188459313.1 |
| Bacteroidota | *Psychroflexus salarius* | WP_073191291.1 | WP_073190977.1 | WP_073193286.1 |
| Bacteroidota | *Psychroflexus salinarum* | WP_379657626.1 | WP_379658489.1 | WP_379658701.1 |
| Bacteroidota | *Psychroflexus salis* | WP_188405766.1 | WP_188405842.1 | WP_188406275.1 |
| Bacteroidota | *Psychroflexus sediminis* | WP_093369230.1 | WP_093367813.1 | WP_093367458.1 |
| Bacteroidota | *Psychroflexus sp. MES1-P1E* | WP_102089321.1 | WP_102087993.1 | WP_102089097.1 |
| Bacteroidota | *Psychroflexus torquis* | WP_015025785.1 | WP_015025542.1 | WP_015024936.1 |
| Bacteroidota | *Psychroflexus tropicus* | WP_019038808.1 | WP_019039259.1 | WP_019038943.1 |
| Bacteroidota | *Psychroserpens algicola* | WP_204345061.1 | WP_204345741.1 | WP_204346159.1 |
| Bacteroidota | *Psychroserpens burtonensis* | WP_028870830.1 | WP_028872699.1 | WP_028872803.1 |
| Bacteroidota | *Psychroserpens damuponensis* | WP_040279120.1 | WP_040280953.1 | WP_040281111.1 |
| Bacteroidota | *Psychroserpens jangbogonensis* | WP_033959147.1 | WP_033960345.1 | WP_033956243.1 |
| Bacteroidota | *Psychroserpens luteolus* | WP_230935135.1 | WP_230935982.1 | WP_230936893.1 |
| Bacteroidota | *Psychroserpens luteus* | WP_194507318.1 | WP_194507631.1 | WP_194508324.1 |
| Bacteroidota | *Psychroserpens mesophilus* | WP_040251637.1 | WP_174435320.1 | WP_040250904.1 |
| Bacteroidota | *Psychroserpens ponticola* | WP_249995923.1 | WP_249995559.1 | WP_249995438.1 |
| Bacteroidota | *Psychroserpens sp.* | WP_425223194.1 | WP_405335216.1 | WP_424516448.1 |
| Bacteroidota | *Puia dinghuensis* | WP_188937963.1 | WP_188929879.1 | WP_188927805.1 |
| Bacteroidota | *Puia sp.* | WP_331576494.1 | WP_331578006.1 | WP_331576466.1 |
| Bacteroidota | *Pukyongia salina* | WP_105217152.1 | WP_105216883.1 | WP_105214167.1 |
| Bacteroidota | *Pustulibacterium marinum* | WP_093026409.1 | WP_093025214.1 | WP_093023162.1 |
| Bacteroidota | *Putridiphycobacter roseus* | WP_111062192.1 | WP_111062136.1 | WP_111061356.1 |
| Bacteroidota | *Raineya orbicola* | WP_101359701.1 | WP_101357744.1 | WP_165778063.1 |
| Bacteroidota | *Rapidithrix thailandica* | WP_346820230.1 | WP_346819169.1 | WP_346819304.1 |
| Bacteroidota | *Rasiella rasia* | WP_164678373.1 | WP_164678848.1 | WP_164679013.1 |

| | | | | |
|---|---|---|---|---|
| Bacteroidota | *Ravibacter arvi* | WP_345028311.1 | WP_345026905.1 | WP_345026675.1 |
| Bacteroidota | *Reichenbachiella agariperforans* | WP_073124535.1 | WP_073123459.1 | WP_216011389.1 |
| Bacteroidota | *Reichenbachiella agarivorans* | WP_262309824.1 | WP_262308525.1 | WP_262310351.1 |
| Bacteroidota | *Reichenbachiella carrageenanivorans* | WP_263050107.1 | WP_263049587.1 | WP_263051374.1 |
| Bacteroidota | *Reichenbachiella faecimaris* | WP_084371036.1 | WP_084371183.1 | WP_084371882.1 |
| Bacteroidota | *Reichenbachiella sp.* | WP_348179303.1 | WP_420580935.1 | WP_348183144.1 |
| Bacteroidota | *Reichenbachiella ulvae* | WP_264137898.1 | WP_264137298.1 | WP_264138500.1 |
| Bacteroidota | *Reichenbachiella versicolor* | WP_109829597.1 | WP_109831061.1 | WP_109832016.1 |
| Bacteroidota | *Rhabdobacter roseus* | WP_184174142.1 | WP_184172287.1 | WP_184174737.1 |
| Bacteroidota | *Rhinopithecimicrobium faecis* | WP_353546519.1 | WP_353546293.1 | WP_353547657.1 |
| Bacteroidota | *Rhizosphaericola mali* | WP_131328136.1 | WP_131330415.1 | WP_131330649.1 |
| Bacteroidota | *Rhodocytophaga aerolata* | WP_302039324.1 | WP_302037742.1 | WP_302038683.1 |
| Bacteroidota | *Rhodocytophaga rosea* | WP_162443911.1 | WP_162448146.1 | WP_232064797.1 |
| Bacteroidota | *Rhodoflexus caldus* | WP_250631486.1 | WP_250632666.1 | WP_250629984.1 |
| Bacteroidota | *Riemerella anatipestifer* | WP_340305057.1 | WP_004917237.1 | WP_310492218.1 |
| Bacteroidota | *Riemerella columbina* | WP_302505457.1 | WP_302505408.1 | WP_018675101.1 |
| Bacteroidota | *Riemerella columbipharyngis* | WP_092736975.1 | WP_092737568.1 | WP_092736482.1 |
| Bacteroidota | *Robertkochia marina* | WP_224490187.1 | WP_224491073.1 | WP_136336997.1 |
| Bacteroidota | *Robertkochia sediminum* | WP_203061522.1 | WP_203060466.1 | WP_203059512.1 |
| Bacteroidota | *Robertkochia solimangrovi* | WP_143955679.1 | WP_143955932.1 | WP_143956131.1 |
| Bacteroidota | *Robertkochia sp. 3YJGBD-33* | WP_224483128.1 | WP_224484353.1 | WP_224484607.1 |
| Bacteroidota | *Robiginitalea aestuariiviva* | WP_277707881.1 | WP_277707969.1 | WP_277707719.1 |
| Bacteroidota | *Robiginitalea aurantiaca* | WP_289725935.1 | WP_289725543.1 | WP_289725990.1 |
| Bacteroidota | *Robiginitalea marina* | WP_252739687.1 | WP_252739769.1 | WP_252740229.1 |
| Bacteroidota | *Robiginitalea myxolifaciens* | WP_092982559.1 | WP_092982473.1 | WP_092982694.1 |
| Bacteroidota | *Robiginitalea sediminis* | WP_088340947.1 | WP_088341021.1 | WP_088340817.1 |
| Bacteroidota | *Robiginitalea sp.* | WP_424498482.1 | WP_424497095.1 | WP_424498923.1 |
| Bacteroidota | *Roseivirga echinicomitans* | WP_068416355.1 | WP_068413391.1 | WP_068414968.1 |

| | | | | |
|---|---|---|---|---|
| Bacteroidota | *Roseivirga ehrenbergii* | WP_062588049.1 | WP_062591750.1 | WP_062594148.1 |
| Bacteroidota | *Roseivirga misakiensis* | WP_069833988.1 | WP_069835979.1 | WP_069833712.1 |
| Bacteroidota | *Roseivirga pacifica* | WP_252937943.1 | WP_422354073.1 | WP_268123344.1 |
| Bacteroidota | *Roseivirga seohaensis* | WP_421977005.1 | WP_421976220.1 | WP_421976594.1 |
| Bacteroidota | *Roseivirga sp.* | WP_421986854.1 | WP_273274618.1 | WP_323756790.1 |
| Bacteroidota | *Rudanella lutea* | WP_019988432.1 | WP_027302744.1 | WP_019989311.1 |
| Bacteroidota | *Rudanella paleaurantiibacter* | WP_152125379.1 | WP_152125921.1 | WP_152123479.1 |
| Bacteroidota | *Rufibacter aurantiacus* | WP_210487332.1 | WP_210488992.1 | WP_210488416.1 |
| Bacteroidota | *Rufibacter glacialis* | WP_149097938.1 | WP_149098143.1 | WP_149098844.1 |
| Bacteroidota | *Rufibacter hautae* | WP_149088950.1 | WP_149091153.1 | WP_149089626.1 |
| Bacteroidota | *Rufibacter immobilis* | WP_123132599.1 | WP_377059760.1 | WP_123133193.1 |
| Bacteroidota | *Rufibacter latericius* | WP_123125038.1 | WP_123125940.1 | WP_123126617.1 |
| Bacteroidota | *Rufibacter psychrotolerans* | WP_205502872.1 | WP_205500681.1 | WP_205500297.1 |
| Bacteroidota | *Rufibacter radiotolerans* | WP_048921188.1 | WP_048920482.1 | WP_048921794.1 |
| Bacteroidota | *Rufibacter roseolus* | WP_210463819.1 | WP_210464973.1 | WP_210463158.1 |
| Bacteroidota | *Rufibacter roseus* | WP_066618735.1 | WP_066621324.1 | WP_066623342.1 |
| Bacteroidota | *Rufibacter sediminis* | WP_186640101.1 | WP_186636070.1 | WP_186639587.1 |
| Bacteroidota | *Rufibacter sp. XAAS-G3-1* | WP_181305783.1 | WP_181306455.1 | WP_181305147.1 |
| Bacteroidota | *Rufibacter tibetensis* | WP_062542449.1 | WP_062545229.1 | WP_062543027.1 |
| Bacteroidota | *Runella aurantiaca* | WP_114461994.1 | WP_114463800.1 | WP_114460029.1 |
| Bacteroidota | *Runella defluvii* | WP_183970994.1 | WP_183975005.1 | WP_183975628.1 |
| Bacteroidota | *Runella limosa* | WP_051397804.1 | WP_028522048.1 | WP_028525003.1 |
| Bacteroidota | *Runella slithyformis* | WP_013928840.1 | WP_013925979.1 | WP_013926801.1 |
| Bacteroidota | *Runella sp.* | WP_298349780.1 | WP_298356035.1 | WP_298353356.1 |
| Bacteroidota | *Runella zeae* | WP_028666985.1 | WP_273215829.1 | WP_028664036.1 |
| Bacteroidota | *Rurimicrobium arvi* | WP_344826516.1 | WP_344828551.1 | WP_344822986.1 |
| Bacteroidota | *Sabulibacter ruber* | WP_207435692.1 | WP_207430476.1 | WP_207434099.1 |
| Bacteroidota | *Salegentibacter chungangensis* | WP_380745092.1 | WP_380744225.1 | WP_380744138.1 |

| | | | | |
|---|---|---|---|---|
| Bacteroidota | *Salegentibacter echinorum* | WP_072880367.1 | WP_072876739.1 | WP_072877766.1 |
| Bacteroidota | *Salegentibacter holothuriorum* | WP_079719038.1 | WP_079719586.1 | WP_079721408.1 |
| Bacteroidota | *Salegentibacter lacus* | WP_224470366.1 | WP_224469924.1 | WP_224471196.1 |
| Bacteroidota | *Salegentibacter maritimus* | WP_198637701.1 | WP_198638642.1 | WP_198639097.1 |
| Bacteroidota | *Salegentibacter mishustinae* | WP_057483160.1 | WP_057481422.1 | WP_282019038.1 |
| Bacteroidota | *Salegentibacter salarius* | WP_070054325.1 | WP_070053585.1 | WP_070054088.1 |
| Bacteroidota | *Salegentibacter salegens* | WP_079734098.1 | WP_079734565.1 | WP_079734433.1 |
| Bacteroidota | *Salegentibacter salinarum* | WP_079712555.1 | WP_079713981.1 | WP_079712005.1 |
| Bacteroidota | *Salegentibacter sediminis* | WP_081211134.1 | WP_081212117.1 | WP_081212067.1 |
| Bacteroidota | *Salegentibacter sp. Hel_I_6* | WP_037322480.1 | WP_037315191.1 | WP_037320299.1 |
| Bacteroidota | *Salegentibacter tibetensis* | WP_224473551.1 | WP_224474186.1 | WP_224474207.1 |
| Bacteroidota | *Salibacter halophilus* | WP_151167347.1 | WP_151166061.1 | WP_151165890.1 |
| Bacteroidota | *Salibacter sp.* | WP_310667847.1 | WP_310667026.1 | WP_310683725.1 |
| Bacteroidota | *Salinimicrobium catena* | WP_093112685.1 | WP_093112502.1 | WP_093113704.1 |
| Bacteroidota | *Salinimicrobium flavum* | WP_380752956.1 | WP_380747343.1 | WP_380748127.1 |
| Bacteroidota | *Salinimicrobium gaetbulicola* | WP_380736752.1 | WP_380736910.1 | WP_380737270.1 |
| Bacteroidota | *Salinimicrobium marinum* | WP_189603621.1 | WP_189604486.1 | WP_189604708.1 |
| Bacteroidota | *Salinimicrobium oceani* | WP_168138491.1 | WP_168136558.1 | WP_168136841.1 |
| Bacteroidota | *Salinimicrobium profundisediminis* | WP_266068280.1 | WP_266070952.1 | WP_266070336.1 |
| Bacteroidota | *Salinimicrobium sediminis* | WP_097056067.1 | WP_097056276.1 | WP_097057124.1 |
| Bacteroidota | *Salinimicrobium soli* | WP_418335852.1 | WP_418335645.1 | WP_418334907.1 |
| Bacteroidota | *Salinimicrobium sp. HB62* | WP_324719156.1 | WP_324719385.1 | WP_324721883.1 |
| Bacteroidota | *Salinimicrobium terrae* | WP_029033011.1 | WP_029033552.1 | WP_029034990.1 |
| Bacteroidota | *Salinimicrobium tongyeongense* | WP_265163551.1 | WP_265163274.1 | WP_265164049.1 |
| Bacteroidota | *Salinimicrobium xinjiangense* | WP_029036068.1 | WP_029037520.1 | WP_029038605.1 |
| Bacteroidota | *Salmonirosea aquatica* | WP_152762569.1 | WP_152764206.1 | WP_152759519.1 |
| Bacteroidota | *Sandaracinomonas limnophila* | WP_127804355.1 | WP_127802513.1 | WP_127803171.1 |
| Bacteroidota | *Sanyastnella coralliicola* | WP_306643031.1 | WP_306639898.1 | WP_306643246.1 |

| | | | | |
|---|---|---|---|---|
| Bacteroidota | *Saonia flava* | WP_167960989.1 | WP_167960713.1 | WP_167961412.1 |
| Bacteroidota | *Saprospira grandis* | WP_270100066.1 | WP_015691616.1 | WP_014373498.1 |
| Bacteroidota | *Saprospira sp. CCB-QB6* | WP_272620540.1 | WP_272620609.1 | WP_272620088.1 |
| Bacteroidota | *Schleiferia thermophila* | WP_037355887.1 | WP_037358008.1 | WP_160171979.1 |
| Bacteroidota | *Sediminibacter sp. Hel_I_10* | WP_026755702.1 | WP_026755964.1 | WP_026753280.1 |
| Bacteroidota | *Sediminibacterium ginsengisoli* | WP_078831188.1 | WP_078832335.1 | WP_245825700.1 |
| Bacteroidota | *Sediminibacterium goheungense* | WP_133474678.1 | WP_133475808.1 | WP_133475323.1 |
| Bacteroidota | *Sediminibacterium roseum* | WP_161817057.1 | WP_161818326.1 | WP_161817922.1 |
| Bacteroidota | *Sediminibacterium salmoneum* | WP_026764238.1 | WP_026763456.1 | WP_026763181.1 |
| Bacteroidota | *Sediminibacterium soli* | WP_161835386.1 | WP_161834296.1 | WP_161834578.1 |
| Bacteroidota | *Sediminibacterium sp.* | WP_322194853.1 | WP_322196213.1 | WP_295204650.1 |
| Bacteroidota | *Sediminicola arcticus* | WP_354615426.1 | WP_354615298.1 | WP_354615560.1 |
| Bacteroidota | *Sediminicola luteus* | WP_097443678.1 | WP_097442876.1 | WP_097440349.1 |
| Bacteroidota | *Sediminicola sp. 1XM1-17* | WP_337300318.1 | WP_337300228.1 | WP_337299993.1 |
| Bacteroidota | *Sediminitomix flava* | WP_109616372.1 | WP_109618873.1 | WP_245935572.1 |
| Bacteroidota | *Segetibacter aerophilus* | WP_147203107.1 | WP_147204351.1 | WP_147202618.1 |
| Bacteroidota | *Segetibacter koreensis* | WP_018617501.1 | WP_018610874.1 | WP_018612798.1 |
| Bacteroidota | *Segetibacter sp.* | WP_297821751.1 | WP_297821841.1 | WP_297822188.1 |
| Bacteroidota | *Seonamhaeicola algicola* | WP_147130549.1 | WP_147133745.1 | WP_147133851.1 |
| Bacteroidota | *Seonamhaeicola aphaedonensis* | WP_116039943.1 | WP_116523873.1 | WP_116524046.1 |
| Bacteroidota | *Seonamhaeicola marinus* | WP_148541872.1 | WP_148544579.1 | WP_262713610.1 |
| Bacteroidota | *Seonamhaeicola maritimus* | WP_282135950.1 | WP_147769343.1 | WP_147769247.1 |
| Bacteroidota | *Seonamhaeicola sediminis* | WP_133356602.1 | WP_133357173.1 | WP_133354879.1 |
| Bacteroidota | *Seonamhaeicola sp.* | WP_299547549.1 | WP_299550404.1 | WP_299551035.1 |
| Bacteroidota | *Shiella aurantiaca* | WP_320003240.1 | WP_320003921.1 | WP_320004771.1 |
| Bacteroidota | *Shivajiella indica* | WP_380799824.1 | WP_380800894.1 | WP_380799678.1 |
| Bacteroidota | *Siansivirga zeaxanthinifaciens* | WP_044638880.1 | WP_044638641.1 | WP_044638743.1 |
| Bacteroidota | *Sinomicrobium kalidii* | WP_233030326.1 | WP_233028330.1 | WP_233028368.1 |

| | | | | |
|---|---|---|---|---|
| Bacteroidota | *Sinomicrobium oceani* | WP_072317176.1 | WP_072315514.1 | WP_326492528.1 |
| Bacteroidota | *Sinomicrobium pectinilyticum* | WP_123214022.1 | WP_123218411.1 | WP_123214924.1 |
| Bacteroidota | *Sinomicrobium sp. N-1-3-6* | WP_112372356.1 | WP_112370791.1 | WP_112373290.1 |
| Bacteroidota | *Sinomicrobium weinanense* | WP_187964674.1 | WP_187966821.1 | WP_187964446.1 |
| Bacteroidota | *Siphonobacter aquaeclarae* | WP_093207878.1 | WP_093203666.1 | WP_093200151.1 |
| Bacteroidota | *Snuella lapsa* | WP_345003601.1 | WP_345003664.1 | WP_345007529.1 |
| Bacteroidota | *Snuella sedimenti* | WP_199115617.1 | WP_199116703.1 | WP_199114629.1 |
| Bacteroidota | *Solirubrum puertoriconensis* | WP_059067657.1 | WP_059071390.1 | WP_059068878.1 |
| Bacteroidota | *Solitalea canadensis* | WP_014682578.1 | WP_014680712.1 | WP_014681131.1 |
| Bacteroidota | *Solitalea koreensis* | WP_142603364.1 | WP_142604438.1 | WP_142600541.1 |
| Bacteroidota | *Solitalea lacus* | WP_237848968.1 | WP_237846591.1 | WP_237847143.1 |
| Bacteroidota | *Solitalea longa* | WP_103787391.1 | WP_103790159.1 | WP_103788875.1 |
| Bacteroidota | *Soonwooa buanensis* | WP_079668157.1 | WP_079667385.1 | WP_079666718.1 |
| Bacteroidota | *Soonwooa sp.* | WP_300672259.1 | WP_300673687.1 | WP_300674339.1 |
| Bacteroidota | *Sphingobacterium alimentarium* | WP_132776151.1 | WP_132777594.1 | WP_132777741.1 |
| Bacteroidota | *Sphingobacterium alkalisoli* | WP_136819553.1 | WP_136821884.1 | WP_136818617.1 |
| Bacteroidota | *Sphingobacterium allocomposti* | WP_148907204.1 | WP_148908684.1 | WP_148909427.1 |
| Bacteroidota | *Sphingobacterium arenae* | WP_190310305.1 | WP_190308564.1 | WP_190307389.1 |
| Bacteroidota | *Sphingobacterium bambusae* | WP_320185828.1 | WP_320184487.1 | WP_320186568.1 |
| Bacteroidota | *Sphingobacterium bovisgrunnientis* | WP_160068841.1 | WP_160069544.1 | WP_160070284.1 |
| Bacteroidota | *Sphingobacterium bovistauri* | WP_225551901.1 | WP_225550927.1 | WP_225554940.1 |
| Bacteroidota | *Sphingobacterium cavernae* | WP_149913121.1 | WP_149915679.1 | WP_149913630.1 |
| Bacteroidota | *Sphingobacterium chungjuense* | WP_166332274.1 | WP_166333473.1 | WP_166332900.1 |
| Bacteroidota | *Sphingobacterium chuzhouense* | WP_190315297.1 | WP_190314194.1 | WP_190312758.1 |
| Bacteroidota | *Sphingobacterium composti* | WP_159634226.1 | WP_159636159.1 | WP_159634454.1 |
| Bacteroidota | *Sphingobacterium corticibacter* | WP_116776166.1 | WP_116774497.1 | WP_116775555.1 |
| Bacteroidota | *Sphingobacterium corticibacterium* | WP_130143238.1 | WP_130139939.1 | WP_130142176.1 |
| Bacteroidota | *Sphingobacterium corticis* | WP_380867056.1 | WP_380869886.1 | WP_380868428.1 |

| | | | | |
|---|---|---|---|---|
| Bacteroidota | *Sphingobacterium daejeonense* | WP_260027722.1 | WP_260033050.1 | WP_380894417.1 |
| Bacteroidota | *Sphingobacterium deserti* | WP_037495749.1 | WP_037503513.1 | WP_037497633.1 |
| Bacteroidota | *Sphingobacterium detergens* | WP_120261583.1 | WP_120259190.1 | WP_418359127.1 |
| Bacteroidota | *Sphingobacterium endophyticum* | WP_156306153.1 | WP_156309128.1 | WP_156307566.1 |
| Bacteroidota | *Sphingobacterium faecale* | WP_202101697.1 | WP_202102283.1 | WP_202103280.1 |
| Bacteroidota | *Sphingobacterium faecium* | WP_108160791.1 | WP_346604585.1 | WP_108159374.1 |
| Bacteroidota | *Sphingobacterium gobiense* | WP_105727300.1 | WP_105723300.1 | WP_105725618.1 |
| Bacteroidota | *Sphingobacterium griseoflavum* | WP_189625151.1 | WP_189627229.1 | WP_189625520.1 |
| Bacteroidota | *Sphingobacterium haloxyli* | WP_105717127.1 | WP_105716264.1 | WP_105715364.1 |
| Bacteroidota | *Sphingobacterium hotanense* | WP_149527035.1 | WP_260044612.1 | WP_260039951.1 |
| Bacteroidota | *Sphingobacterium humi* | WP_160370061.1 | WP_160367591.1 | WP_160369867.1 |
| Bacteroidota | *Sphingobacterium hungaricum* | WP_196935180.1 | WP_196936872.1 | WP_196935741.1 |
| Bacteroidota | *Sphingobacterium kyonggiense* | WP_344673062.1 | WP_344674680.1 | WP_344675295.1 |
| Bacteroidota | *Sphingobacterium lactis* | WP_103904721.1 | WP_103906279.1 | WP_103906628.1 |
| Bacteroidota | *Sphingobacterium lumbrici* | WP_140937507.1 | WP_140938382.1 | WP_140937032.1 |
| Bacteroidota | *Sphingobacterium micropteri* | WP_190995033.1 | WP_190993283.1 | WP_190994339.1 |
| Bacteroidota | *Sphingobacterium multivorum* | WP_312744889.1 | WP_313529588.1 | WP_201666921.1 |
| Bacteroidota | *Sphingobacterium nematocida* | WP_079641174.1 | WP_079641716.1 | WP_079644723.1 |
| Bacteroidota | *Sphingobacterium olei* | WP_136900903.1 | WP_136899696.1 | WP_136900632.1 |
| Bacteroidota | *Sphingobacterium oryzagri* | WP_274266794.1 | WP_274269451.1 | WP_274267454.1 |
| Bacteroidota | *Sphingobacterium paludis* | WP_133639213.1 | WP_133640419.1 | WP_133639947.1 |
| Bacteroidota | *Sphingobacterium paucimobilis* | WP_021068833.1 | WP_021070158.1 | WP_021071774.1 |
| Bacteroidota | *Sphingobacterium pedocola* | WP_196937863.1 | WP_196938004.1 | WP_196939367.1 |
| Bacteroidota | *Sphingobacterium phlebotomi* | WP_148919322.1 | WP_148918764.1 | WP_148918117.1 |
| Bacteroidota | *Sphingobacterium prati* | WP_172393777.1 | WP_172394118.1 | WP_172392574.1 |
| Bacteroidota | *Sphingobacterium psychroaquaticum* | WP_085472502.1 | WP_085471717.1 | WP_134430177.1 |
| Bacteroidota | *Sphingobacterium puteale* | WP_121126256.1 | WP_121123514.1 | WP_121125361.1 |
| Bacteroidota | *Sphingobacterium rhinopitheci* | WP_241790133.1 | WP_241788344.1 | WP_241790031.1 |

| | | | | |
|---|---|---|---|---|
| Bacteroidota | *Sphingobacterium shayense* | WP_172389207.1 | WP_172389514.1 | WP_172389689.1 |
| Bacteroidota | *Sphingobacterium siyangense* | WP_424545452.1 | WP_346068928.1 | WP_205400216.1 |
| Bacteroidota | *Sphingobacterium sp.* | WP_312337529.1 | WP_343566882.1 | WP_313261371.1 |
| Bacteroidota | *Sphingobacterium spiritivorum* | WP_002992708.1 | WP_115171584.1 | WP_201681064.1 |
| Bacteroidota | *Sphingobacterium suaedae* | WP_380903687.1 | WP_380901132.1 | WP_380900344.1 |
| Bacteroidota | *Sphingobacterium tabacisoli* | WP_210355260.1 | WP_210353897.1 | WP_210355779.1 |
| Bacteroidota | *Sphingobacterium thalpophilum* | WP_424543296.1 | WP_282636042.1 | WP_370481172.1 |
| Bacteroidota | *Sphingobacterium thermophilum* | WP_345067253.1 | WP_345068116.1 | WP_345069121.1 |
| Bacteroidota | *Sphingobacterium wenxiniae* | WP_093365916.1 | WP_093363732.1 | WP_093364962.1 |
| Bacteroidota | *Sphingobacterium yanglingense* | WP_133585347.1 | WP_133584895.1 | WP_133586886.1 |
| Bacteroidota | *Sphingobacterium zhuxiongii* | WP_153509740.1 | WP_153511565.1 | WP_153509167.1 |
| Bacteroidota | *Spirosoma aerolatum* | WP_080058510.1 | WP_080057299.1 | WP_080053908.1 |
| Bacteroidota | *Spirosoma agri* | WP_164036195.1 | WP_164036465.1 | WP_164041052.1 |
| Bacteroidota | *Spirosoma arboris* | WP_157586299.1 | WP_157589715.1 | WP_157587946.1 |
| Bacteroidota | *Spirosoma aureum* | WP_167215690.1 | WP_167214910.1 | WP_167204164.1 |
| Bacteroidota | *Spirosoma endbachense* | WP_162391124.1 | WP_162391301.1 | WP_162388914.1 |
| Bacteroidota | *Spirosoma endophyticum* | WP_093823250.1 | WP_093828459.1 | WP_093825618.1 |
| Bacteroidota | *Spirosoma flavum* | WP_381497588.1 | WP_381506423.1 | WP_381508764.1 |
| Bacteroidota | *Spirosoma fluviale* | WP_097128538.1 | WP_097124341.1 | WP_097130254.1 |
| Bacteroidota | *Spirosoma foliorum* | WP_182457958.1 | WP_182461395.1 | WP_182459763.1 |
| Bacteroidota | *Spirosoma lacussanchae* | WP_138499346.1 | WP_138500323.1 | WP_138507235.1 |
| Bacteroidota | *Spirosoma linguale* | WP_012928013.1 | WP_012925276.1 | WP_012929824.1 |
| Bacteroidota | *Spirosoma luteum* | WP_018621355.1 | WP_018619126.1 | WP_018617925.1 |
| Bacteroidota | *Spirosoma montaniterrae* | WP_077131233.1 | WP_077131579.1 | WP_077132411.1 |
| Bacteroidota | *Spirosoma oryzae* | WP_106140358.1 | WP_106137905.1 | WP_106136314.1 |
| Bacteroidota | *Spirosoma panaciterrae* | WP_020600438.1 | WP_020595618.1 | WP_020598523.1 |
| Bacteroidota | *Spirosoma pollinicola* | WP_100987614.1 | WP_100986176.1 | WP_100988906.1 |
| Bacteroidota | *Spirosoma profusum* | WP_190888856.1 | WP_190888710.1 | WP_190889448.1 |

| | | | | |
|---|---|---|---|---|
| Bacteroidota | *Spirosoma radiotolerans* | WP_046576037.1 | WP_046376221.1 | WP_046578030.1 |
| Bacteroidota | *Spirosoma rigui* | WP_080238489.1 | WP_080241085.1 | WP_080238045.1 |
| Bacteroidota | *Spirosoma soli* | WP_381520133.1 | WP_381518935.1 | WP_381524593.1 |
| Bacteroidota | *Spirosoma sp.* | WP_420149467.1 | WP_420150375.1 | WP_420151292.1 |
| Bacteroidota | *Spirosoma spitsbergense* | WP_020601793.1 | WP_020602422.1 | WP_020607185.1 |
| Bacteroidota | *Spirosoma taeanense* | WP_171740860.1 | WP_171739650.1 | WP_171741738.1 |
| Bacteroidota | *Spirosoma terrae* | WP_163946965.1 | WP_163944895.1 | WP_163941605.1 |
| Bacteroidota | *Spirosoma utsteinense* | WP_186739525.1 | WP_186739119.1 | WP_186737346.1 |
| Bacteroidota | *Spirosoma validum* | WP_191036995.1 | WP_191039184.1 | WP_191040336.1 |
| Bacteroidota | *Splendidivirga corallicola* | WP_346754110.1 | WP_346754927.1 | WP_346750622.1 |
| Bacteroidota | *Spongiimicrobium salis* | WP_394749418.1 | WP_394749267.1 | WP_394749767.1 |
| Bacteroidota | *Spongiimicrobium sp. 3-5* | WP_411032168.1 | WP_411030488.1 | WP_411031717.1 |
| Bacteroidota | *Spongiivirga citrea* | WP_164029170.1 | WP_164028867.1 | WP_164032428.1 |
| Bacteroidota | *Spongiivirga sp. MCCC 1A20706* | WP_410484950.1 | WP_410483661.1 | WP_410485712.1 |
| Bacteroidota | *Sporocytophaga myxococcoides* | WP_045459397.1 | WP_045464077.1 | WP_045459954.1 |
| Bacteroidota | *Sporocytophaga sp.* | WP_293897300.1 | WP_293894613.1 | WP_293892478.1 |
| Bacteroidota | *Subsaxibacter sp. CAU 1640* | WP_248025836.1 | WP_248025383.1 | WP_248024187.1 |
| Bacteroidota | *Subsaximicrobium wynnwilliamsii* | WP_147084534.1 | WP_147085199.1 | WP_317128650.1 |
| Bacteroidota | *Sungkyunkwania multivorans* | WP_386407340.1 | WP_386405566.1 | WP_386407050.1 |
| Bacteroidota | *Taibaiella chishuiensis* | WP_106525076.1 | WP_106521535.1 | WP_181358454.1 |
| Bacteroidota | *Taibaiella helva* | WP_118953545.1 | WP_118949944.1 | WP_157976869.1 |
| Bacteroidota | *Taibaiella koreensis* | WP_118972826.1 | WP_118974487.1 | WP_162903180.1 |
| Bacteroidota | *Taibaiella lutea* | WP_150031081.1 | WP_150033014.1 | WP_190277335.1 |
| Bacteroidota | *Taibaiella soli* | WP_110999133.1 | WP_110996983.1 | WP_110999509.1 |
| Bacteroidota | *Taibaiella sp. KBW10* | WP_124636061.1 | WP_124635204.1 | WP_124636155.1 |
| Bacteroidota | *Taishania pollutisoli* | WP_163492192.1 | WP_163490697.1 | WP_216714193.1 |
| Bacteroidota | *Tamlana crocina* | WP_167917677.1 | WP_167919677.1 | WP_167918689.1 |
| Bacteroidota | *Tamlana flava* | WP_370479806.1 | WP_370476504.1 | WP_370476851.1 |

| | | | | |
|---|---|---|---|---|
| Bacteroidota | *Tamlana sp. I1* | WP_194767652.1 | WP_194767741.1 | WP_194768077.1 |
| Bacteroidota | *Telluribacter humicola* | WP_207505055.1 | WP_207507151.1 | WP_310586155.1 |
| Bacteroidota | *Telluribacter sp.* | WP_327487111.1 | WP_327487727.1 | WP_327484708.1 |
| Bacteroidota | *Tellurirhabdus bombi* | WP_234735125.1 | WP_234734379.1 | WP_234736306.1 |
| Bacteroidota | *Tellurirhabdus rosea* | WP_266368610.1 | WP_266365853.1 | WP_266369101.1 |
| Bacteroidota | *Tenacibaculum adriaticum* | WP_148869874.1 | WP_148871029.1 | WP_148871047.1 |
| Bacteroidota | *Tenacibaculum aestuarii* | WP_408035614.1 | WP_408036213.1 | WP_408036178.1 |
| Bacteroidota | *Tenacibaculum aestuariivivum* | WP_418650557.1 | WP_418649929.1 | WP_418649959.1 |
| Bacteroidota | *Tenacibaculum agarivorans* | WP_075342671.1 | WP_075343661.1 | WP_075343920.1 |
| Bacteroidota | *Tenacibaculum aiptasiae* | WP_420552946.1 | WP_420551224.1 | WP_272151765.1 |
| Bacteroidota | *Tenacibaculum amylolyticum* | WP_408039680.1 | WP_408039792.1 | WP_408037409.1 |
| Bacteroidota | *Tenacibaculum ascidiaceicola* | WP_425658519.1 | WP_415268493.1 | WP_415268506.1 |
| Bacteroidota | *Tenacibaculum caenipelagi* | WP_133536489.1 | WP_133534486.1 | WP_133534460.1 |
| Bacteroidota | *Tenacibaculum crassostreae* | WP_408046929.1 | WP_408046466.1 | WP_408046492.1 |
| Bacteroidota | *Tenacibaculum dicentrarchi* | WP_101902041.1 | WP_101902640.1 | WP_370407980.1 |
| Bacteroidota | *Tenacibaculum discolor* | WP_124588550.1 | WP_124590251.1 | WP_099213840.1 |
| Bacteroidota | *Tenacibaculum finnmarkense* | WP_101915721.1 | WP_232125399.1 | WP_193707775.1 |
| Bacteroidota | *Tenacibaculum gallaicum* | WP_115901618.1 | WP_115900961.1 | WP_115900927.1 |
| Bacteroidota | *Tenacibaculum geojense* | WP_386104704.1 | WP_386104940.1 | WP_386104979.1 |
| Bacteroidota | *Tenacibaculum holothuriorum* | WP_086029382.1 | WP_086030732.1 | WP_086030767.1 |
| Bacteroidota | *Tenacibaculum insulae* | WP_418644914.1 | WP_418644674.1 | WP_418644704.1 |
| Bacteroidota | *Tenacibaculum jejuense* | WP_095072327.1 | WP_095073068.1 | WP_095072980.1 |
| Bacteroidota | *Tenacibaculum larymnensis* | WP_274639816.1 | WP_274640215.1 | WP_274640185.1 |
| Bacteroidota | *Tenacibaculum litopenaei* | WP_408044075.1 | WP_408043301.1 | WP_408044502.1 |
| Bacteroidota | *Tenacibaculum lutimaris* | WP_120186681.1 | WP_120187165.1 | WP_120187140.1 |
| Bacteroidota | *Tenacibaculum maritimum* | WP_406741777.1 | WP_232641412.1 | WP_159245217.1 |
| Bacteroidota | *Tenacibaculum mesophilum* | WP_281979306.1 | WP_253679894.1 | WP_281980437.1 |
| Bacteroidota | *Tenacibaculum ovolyticum* | WP_422091942.1 | WP_422092133.1 | WP_237276067.1 |

| | | | | |
|---|---|---|---|---|
| Bacteroidota | *Tenacibaculum pelagium* | WP_182123523.1 | WP_182123957.1 | WP_182123931.1 |
| Bacteroidota | *Tenacibaculum piscium* | WP_101917182.1 | WP_101916480.1 | WP_101916845.1 |
| Bacteroidota | *Tenacibaculum platacis* | WP_348725202.1 | WP_348742779.1 | WP_348724435.1 |
| Bacteroidota | *Tenacibaculum polynesiense* | WP_348716780.1 | WP_348715293.1 | WP_348718874.1 |
| Bacteroidota | *Tenacibaculum sediminilitoris* | WP_408024139.1 | WP_408023732.1 | WP_408025091.1 |
| Bacteroidota | *Tenacibaculum singaporense* | WP_125067654.1 | WP_125344468.1 | WP_125068044.1 |
| Bacteroidota | *Tenacibaculum skagerrakense* | WP_132794789.1 | WP_132795009.1 | WP_132795025.1 |
| Bacteroidota | *Tenacibaculum soleae* | WP_068705066.1 | WP_418654330.1 | WP_271407244.1 |
| Bacteroidota | *Tenacibaculum sp.* | WP_273694360.1 | WP_417799530.1 | WP_417786282.1 |
| Bacteroidota | *Tenacibaculum tangerinum* | WP_279652411.1 | WP_279651372.1 | WP_279651401.1 |
| Bacteroidota | *Tenacibaculum todarodis* | WP_072554883.1 | WP_072555161.1 | WP_072555142.1 |
| Bacteroidota | *Tenacibaculum vairaonense* | WP_348707420.1 | WP_348702102.1 | WP_348746797.1 |
| Bacteroidota | *Tenacibaculum xiamenense* | WP_408031608.1 | WP_408030665.1 | WP_408032154.1 |
| Bacteroidota | *Terrimonas ferruginea* | WP_028787882.1 | WP_303892481.1 | WP_303891090.1 |
| Bacteroidota | *Terrimonas ginsenosidimutans* | WP_237871313.1 | WP_237876689.1 | WP_237877120.1 |
| Bacteroidota | *Terrimonas pollutisoli* | WP_276500859.1 | WP_276504086.1 | WP_276502652.1 |
| Bacteroidota | *Terrimonas rubra* | WP_386097367.1 | WP_386096749.1 | WP_386099849.1 |
| Bacteroidota | *Terrimonas sp.* | WP_116875316.1 | WP_116872616.1 | WP_116872878.1 |
| Bacteroidota | *Thalassobellus suaedae* | WP_415866465.1 | WP_415861774.1 | WP_415862084.1 |
| Bacteroidota | *Thermaurantimonas aggregans* | WP_124396826.1 | WP_124397494.1 | WP_160160588.1 |
| Bacteroidota | *Thermaurantimonas sp.* | WP_409769462.1 | WP_409771203.1 | WP_409771132.1 |
| Bacteroidota | *Thermoflavifilum aggregans* | WP_100313719.1 | WP_100314561.1 | WP_100314460.1 |
| Bacteroidota | *Thermoflavifilum sp.* | WP_297042760.1 | WP_297044056.1 | WP_297043879.1 |
| Bacteroidota | *Thermoflavifilum thermophilum* | WP_092457765.1 | WP_092461028.1 | WP_092460516.1 |
| Bacteroidota | *Thermoflexibacter ruber* | WP_091545755.1 | WP_091545253.1 | WP_091542748.1 |
| Bacteroidota | *Thermonema lapsum* | WP_166919763.1 | WP_166920515.1 | WP_166918106.1 |
| Bacteroidota | *Thermonema rossianum* | WP_051632972.1 | WP_038030079.1 | WP_084147099.1 |
| Bacteroidota | *Thermonema sp.* | WP_288005729.1 | WP_288007082.1 | WP_288004089.1 |

| | | | | |
|---|---|---|---|---|
| Bacteroidota | *Tunicatimonas pelagia* | WP_302205012.1 | WP_302207186.1 | WP_302204998.1 |
| Bacteroidota | *Ulvibacter antarcticus* | WP_121906710.1 | WP_121907624.1 | WP_121907465.1 |
| Bacteroidota | *Ulvibacter litoralis* | WP_093143547.1 | WP_093144987.1 | WP_093145223.1 |
| Bacteroidota | *Ulvibacter sp. MAR_2010_11* | WP_100803909.1 | WP_100804196.1 | WP_100803434.1 |
| Bacteroidota | *Ulvibacterium marinum* | WP_282162937.1 | WP_120711029.1 | WP_120713983.1 |
| Bacteroidota | *Ulvibacterium sp.* | WP_425236708.1 | WP_422080450.1 | WP_422084048.1 |
| Bacteroidota | *Urechidicola croceus* | WP_070235832.1 | WP_070235596.1 | WP_070236178.1 |
| Bacteroidota | *Urechidicola vernalis* | WP_311591836.1 | WP_311592154.1 | WP_311594012.1 |
| Bacteroidota | *Vaginella massiliensis* | WP_068598256.1 | WP_068595181.1 | WP_395090947.1 |
| Bacteroidota | *Vicingus serpentipes* | WP_147101768.1 | WP_147097973.1 | WP_147100110.1 |
| Bacteroidota | *Wandonia haliotis* | WP_343788488.1 | WP_343787892.1 | WP_343785738.1 |
| Bacteroidota | *Wenyingzhuangia aestuarii* | WP_167898441.1 | WP_167897819.1 | WP_167896793.1 |
| Bacteroidota | *Wenyingzhuangia fucanilytica* | WP_068826438.1 | WP_068825555.1 | WP_068826364.1 |
| Bacteroidota | *Wenyingzhuangia gilva* | WP_302883866.1 | WP_302885059.1 | WP_302883913.1 |
| Bacteroidota | *Wenyingzhuangia heitensis* | WP_167185859.1 | WP_167187886.1 | WP_167187308.1 |
| Bacteroidota | *Wenyingzhuangia marina* | WP_073119375.1 | WP_073118897.1 | WP_073119261.1 |
| Bacteroidota | *Wenyingzhuangia sp.* | WP_347182783.1 | WP_347180434.1 | WP_347181761.1 |
| Bacteroidota | *Winogradskyella alexanderae* | WP_224530615.1 | WP_224526489.1 | WP_224526926.1 |
| Bacteroidota | *Winogradskyella algicola* | WP_138433396.1 | WP_138434160.1 | WP_138434612.1 |
| Bacteroidota | *Winogradskyella aquimaris* | WP_320556630.1 | WP_320556356.1 | WP_320555584.1 |
| Bacteroidota | *Winogradskyella arenosi* | WP_114309489.1 | WP_114309772.1 | WP_114310057.1 |
| Bacteroidota | *Winogradskyella aurantia* | WP_094968397.1 | WP_094967377.1 | WP_094968831.1 |
| Bacteroidota | *Winogradskyella aurantiaca* | WP_115462847.1 | WP_115463094.1 | WP_317047361.1 |
| Bacteroidota | *Winogradskyella bathintestinalis* | WP_290206804.1 | WP_290206994.1 | WP_290206180.1 |
| Bacteroidota | *Winogradskyella costae* | WP_179335048.1 | WP_179334623.1 | WP_179335300.1 |
| Bacteroidota | *Winogradskyella echinorum* | WP_186845557.1 | WP_186845790.1 | WP_186845877.1 |
| Bacteroidota | *Winogradskyella eckloniae* | WP_173281061.1 | WP_173280570.1 | WP_173281378.1 |
| Bacteroidota | *Winogradskyella endarachnes* | WP_157361762.1 | WP_157363281.1 | WP_157362488.1 |

| | | | | |
|---|---|---|---|---|
| Bacteroidota | *Winogradskyella epiphytica* | WP_110475170.1 | WP_110476132.1 | WP_110476237.1 |
| Bacteroidota | *Winogradskyella eximia* | WP_282031934.1 | WP_282032511.1 | WP_115817452.1 |
| Bacteroidota | *Winogradskyella flava* | WP_282042702.1 | WP_185789937.1 | WP_185789761.1 |
| Bacteroidota | *Winogradskyella forsetii* | WP_179008254.1 | WP_179006261.1 | WP_179005818.1 |
| Bacteroidota | *Winogradskyella haliclonae* | WP_188374348.1 | WP_188374596.1 | WP_188375109.1 |
| Bacteroidota | *Winogradskyella helgolandensis* | WP_178985666.1 | WP_218646038.1 | WP_179319418.1 |
| Bacteroidota | *Winogradskyella immobilis* | WP_227477848.1 | WP_227476698.1 | WP_227477668.1 |
| Bacteroidota | *Winogradskyella jejuensis* | WP_073086886.1 | WP_073087204.1 | WP_073085722.1 |
| Bacteroidota | *Winogradskyella litorisediminis* | WP_386127183.1 | WP_386132737.1 | WP_386132081.1 |
| Bacteroidota | *Winogradskyella litoriviva* | WP_173300105.1 | WP_173301905.1 | WP_173300484.1 |
| Bacteroidota | *Winogradskyella ludwigii* | WP_179338973.1 | WP_179339464.1 | WP_179338155.1 |
| Bacteroidota | *Winogradskyella luteola* | WP_218544848.1 | WP_218544234.1 | WP_218544490.1 |
| Bacteroidota | *Winogradskyella maritima* | WP_386101441.1 | WP_386095869.1 | WP_386099159.1 |
| Bacteroidota | *Winogradskyella pacifica* | WP_179347853.1 | WP_179349892.1 | WP_179349795.1 |
| Bacteroidota | *Winogradskyella pelagia* | WP_208154871.1 | WP_208153767.1 | WP_208154100.1 |
| Bacteroidota | *Winogradskyella poriferorum* | WP_331809773.1 | WP_331810245.1 | WP_331809409.1 |
| Bacteroidota | *Winogradskyella psychrotolerans* | WP_215925849.1 | WP_215925180.1 | WP_215936225.1 |
| Bacteroidota | *Winogradskyella pulchriflava* | WP_386061894.1 | WP_386061462.1 | WP_386064116.1 |
| Bacteroidota | *Winogradskyella rapida* | WP_386114648.1 | WP_386113114.1 | WP_386117398.1 |
| Bacteroidota | *Winogradskyella schleiferi* | WP_178988830.1 | WP_178988226.1 | WP_178987854.1 |
| Bacteroidota | *Winogradskyella sediminis* | WP_092443319.1 | WP_115839024.1 | WP_417875597.1 |
| Bacteroidota | *Winogradskyella sp.* | WP_418638804.1 | WP_417871518.1 | WP_370000453.1 |
| Bacteroidota | *Winogradskyella tangerina* | WP_111682739.1 | WP_111684095.1 | WP_111683804.1 |
| Bacteroidota | *Winogradskyella thalassocola* | WP_092469884.1 | WP_092467141.1 | WP_092469187.1 |
| Bacteroidota | *Winogradskyella undariae* | WP_179315410.1 | WP_179317035.1 | WP_173588138.1 |
| Bacteroidota | *Winogradskyella ursingii* | WP_179344862.1 | WP_179346499.1 | WP_179346073.1 |
| Bacteroidota | *Winogradskyella vidalii* | WP_179353704.1 | WP_179352429.1 | WP_179351812.1 |
| Bacteroidota | *Winogradskyella vincentii* | WP_224478105.1 | WP_224479179.1 | WP_224478740.1 |

| | | | | |
|---|---|---|---|---|
| Bacteroidota | *Winogradskyella wandonensis* | WP_132703720.1 | WP_132705881.1 | WP_132702815.1 |
| Bacteroidota | *Winogradskyella wichelsiae* | WP_179376959.1 | WP_179376871.1 | WP_179376516.1 |
| Bacteroidota | *Wocania arenilitoris* | WP_237240459.1 | WP_237239480.1 | WP_237240506.1 |
| Bacteroidota | *Wocania ichthyoenteri* | WP_034044660.1 | WP_034040352.1 | WP_034041167.1 |
| Bacteroidota | *Xanthocytophaga agilis* | WP_314516216.1 | WP_314508697.1 | WP_314512272.1 |
| Bacteroidota | *Xanthocytophaga flavus* | WP_313990804.1 | WP_314031117.1 | WP_313990763.1 |
| Bacteroidota | *Xanthomarina gelatinilytica* | WP_417866332.1 | WP_417858572.1 | WP_417855412.1 |
| Bacteroidota | *Xanthomarina sp.* | WP_324313051.1 | WP_324313525.1 | WP_324313434.1 |
| Bacteroidota | *Xanthomarina spongicola* | WP_109681702.1 | WP_109681507.1 | WP_109682487.1 |
| Bacteroidota | *Yeosuana aromativorans* | WP_188653250.1 | WP_188654997.1 | WP_188652436.1 |
| Bacteroidota | *Yeosuana marina* | WP_339917917.1 | WP_166963373.1 | WP_339917623.1 |
| Bacteroidota | *Zeaxanthinibacter enoshimensis* | WP_373074604.1 | WP_373074286.1 | WP_373072286.1 |
| Bacteroidota | *Zeaxanthinibacter sp. PT1* | WP_272595066.1 | WP_272593691.1 | WP_272594309.1 |
| Bacteroidota | *Zhouia amylolytica* | WP_038262980.1 | WP_038262849.1 | WP_255083157.1 |
| Bacteroidota | *Zhouia sp. PK063* | WP_399473499.1 | WP_399471047.1 | WP_399471254.1 |
| Bacteroidota | *Zhouia spongiae* | WP_242936813.1 | WP_242936238.1 | WP_242937265.1 |
| Bacteroidota | *Zobellia alginiliquefaciens* | WP_276166266.1 | WP_276166540.1 | WP_276169207.1 |
| Bacteroidota | *Zobellia amurskyensis* | WP_038233676.1 | WP_155598589.1 | WP_155600564.1 |
| Bacteroidota | *Zobellia barbeyronii* | WP_214613186.1 | WP_214610970.1 | WP_214611862.1 |
| Bacteroidota | *Zobellia nedashkovskayae* | WP_194534562.1 | WP_194535239.1 | WP_194534115.1 |
| Bacteroidota | *Zobellia roscoffensis* | WP_194526843.1 | WP_194530383.1 | WP_194526398.1 |
| Bacteroidota | *Zobellia russellii* | WP_400073611.1 | WP_400073453.1 | WP_400073937.1 |
| Bacteroidota | *Zobellia sp. B3R18* | WP_215914274.1 | WP_215912230.1 | WP_215912435.1 |
| Bacteroidota | *Zobellia uliginosa* | WP_215938390.1 | WP_215939514.1 | WP_303399556.1 |
| Bacteroidota | *Zunongwangia atlantica* | WP_084840657.1 | WP_084841855.1 | WP_084841803.1 |
| Bacteroidota | *Zunongwangia endophytica* | WP_290231854.1 | WP_290233323.1 | WP_290233428.1 |
| Bacteroidota | *Zunongwangia mangrovi* | WP_092540965.1 | WP_092542863.1 | WP_092542653.1 |
| Bacteroidota | *Zunongwangia profunda* | WP_013072155.1 | WP_041578784.1 | WP_228251206.1 |

| | | | | |
|---|---|---|---|---|
| Bacteroidota | *Zunongwangia sp.* | WP_417887496.1 | WP_347406885.1 | WP_417885621.1 |
| Balneolota | *Aliifodinibius salipaludis* | WP_095605785.1 | WP_095607636.1 | WP_255233382.1 |
| Balneolota | *Aliifodinibius sp. S!AR15-10* | WP_310684926.1 | WP_310682093.1 | WP_310685409.1 |
| Balneolota | *Balneola sp. EhC07* | WP_066222314.1 | WP_066220100.1 | WP_066217088.1 |
| Balneolota | *Balneola vulgaris* | WP_018127418.1 | WP_018126226.1 | WP_018126604.1 |
| Balneolota | *Cyclonatronum proteinivorum* | WP_114985548.1 | WP_114983795.1 | WP_164682663.1 |
| Balneolota | *Cyclonatronum sp.* | WP_291482712.1 | WP_291482818.1 | WP_291482004.1 |
| Balneolota | *Fodinibius halophilus* | WP_165266187.1 | WP_165267505.1 | WP_165265239.1 |
| Balneolota | *Fodinibius roseus* | WP_073061524.1 | WP_073063141.1 | WP_073059135.1 |
| Balneolota | *Fodinibius salicampi* | WP_265790485.1 | WP_265788309.1 | WP_265787781.1 |
| Balneolota | *Fodinibius salinus* | WP_148898505.1 | WP_246138157.1 | WP_170245550.1 |
| Balneolota | *Fodinibius saliphilus* | WP_138430825.1 | WP_138430056.1 | WP_138429484.1 |
| Balneolota | *Fodinibius salsisoli* | WP_265764074.1 | WP_265766003.1 | WP_265765528.1 |
| Balneolota | *Fodinibius sediminis* | WP_142712524.1 | WP_142714645.1 | WP_185958215.1 |
| Balneolota | *Fodinibius sp.* | WP_322574622.1 | WP_322575165.1 | WP_372638409.1 |
| Balneolota | *Gracilimonas amylolytica* | WP_103666029.1 | WP_103663667.1 | WP_103663801.1 |
| Balneolota | *Gracilimonas halophila* | WP_390297046.1 | WP_390300200.1 | WP_390300637.1 |
| Balneolota | *Gracilimonas mengyeensis* | WP_142455253.1 | WP_142455020.1 | WP_142452685.1 |
| Balneolota | *Gracilimonas sediminicola* | WP_255134448.1 | WP_255135815.1 | WP_255135726.1 |
| Balneolota | *Gracilimonas sp.* | WP_395079183.1 | WP_395069629.1 | WP_395080504.1 |
| Balneolota | *Gracilimonas tropica* | WP_020404354.1 | WP_020402163.1 | WP_020402845.1 |
| Balneolota | *Halalkalibaculum roseum* | WP_165138370.1 | WP_165140079.1 | WP_165140357.1 |
| Balneolota | *Halalkalibaculum sp. DA3122* | WP_395259678.1 | WP_395259273.1 | WP_395261429.1 |
| Balneolota | *Natronogracilivirga saccharolytica* | WP_210513032.1 | WP_210509515.1 | WP_210511334.1 |
| Balneolota | *Rhodohalobacter barkolensis* | WP_101073066.1 | WP_101071599.1 | WP_101071812.1 |
| Balneolota | *Rhodohalobacter halophilus* | WP_069131108.1 | WP_069130964.1 | WP_069131403.1 |
| Balneolota | *Rhodohalobacter mucosus* | WP_109647779.1 | WP_109646781.1 | WP_109645422.1 |
| Balneolota | *Rhodohalobacter sp.* | WP_372902930.1 | WP_322571692.1 | WP_322571913.1 |

| | | | | |
|---|---|---|---|---|
| Balneolota | *Rhodohalobacter sulfatireducens* | WP_237852468.1 | WP_237853641.1 | WP_237855134.1 |
| Calditrichaeota | *Caldithrix abyssi* | WP_006929106.1 | WP_044281486.1 | WP_006931009.1 |
| Chlamydiota | *Criblamydia sequanensis* | WP_041016448.1 | WP_053331647.1 | WP_041017369.1 |
| Chlamydiota | *Estrella lausannensis* | WP_239414312.1 | WP_098037590.1 | WP_098038239.1 |
| Chlamydiota | *Waddlia chondrophila* | WP_041941457.1 | WP_013181378.1 | WP_013181809.1 |
| Cyanobacteriota | *Leptolyngbya sp. 7M* | WP_211624753.1 | WP_211627088.1 | WP_211627697.1 |
| Deinococcota | *Allomeiothermus silvanus* | WP_276958151.1 | WP_013159191.1 | WP_276956435.1 |
| Deinococcota | *Calidithermus chliarophilus* | WP_027893954.1 | WP_027892264.1 | WP_027893633.1 |
| Deinococcota | *Calidithermus roseus* | WP_119275938.1 | WP_119278785.1 | WP_119278386.1 |
| Deinococcota | *Calidithermus terrae* | WP_119313774.1 | WP_119316488.1 | WP_119314295.1 |
| Deinococcota | *Calidithermus timidus* | WP_018465601.1 | WP_018466689.1 | WP_018467417.1 |
| Deinococcota | *Deinobacterium chartae* | WP_183983655.1 | WP_183987798.1 | WP_343058308.1 |
| Deinococcota | *Deinococcus actinosclerus* | WP_231724576.1 | WP_062159268.1 | WP_062157899.1 |
| Deinococcota | *Deinococcus aerius* | WP_103129980.1 | WP_103130965.1 | WP_103128680.1 |
| Deinococcota | *Deinococcus aerolatus* | WP_188969226.1 | WP_188974266.1 | WP_188969069.1 |
| Deinococcota | *Deinococcus aerophilus* | WP_188902948.1 | WP_188905080.1 | WP_188901788.1 |
| Deinococcota | *Deinococcus aestuarii* | WP_216317724.1 | WP_216323892.1 | WP_216328636.1 |
| Deinococcota | *Deinococcus aetherius* | WP_264774375.1 | WP_264775877.1 | WP_264774420.1 |
| Deinococcota | *Deinococcus alpinitundrae* | WP_237724992.1 | WP_161882051.1 | WP_161882035.1 |
| Deinococcota | *Deinococcus altitudinis* | WP_407570891.1 | WP_407569033.1 | WP_407570468.1 |
| Deinococcota | *Deinococcus aluminii* | WP_345451717.1 | WP_345458031.1 | WP_345451657.1 |
| Deinococcota | *Deinococcus antarcticus* | WP_380077259.1 | WP_380079366.1 | WP_380076048.1 |
| Deinococcota | *Deinococcus apachensis* | WP_019585681.1 | WP_026332545.1 | WP_019585857.1 |
| Deinococcota | *Deinococcus aquaedulcis* | WP_221088799.1 | WP_328774622.1 | WP_221089802.1 |
| Deinococcota | *Deinococcus aquatilis* | WP_019008767.1 | WP_019010936.1 | WP_019009930.1 |
| Deinococcota | *Deinococcus aquiradiocola* | WP_188960394.1 | WP_188964532.1 | WP_188960829.1 |
| Deinococcota | *Deinococcus arcticus* | WP_199188265.1 | WP_158263797.1 | WP_107136680.1 |
| Deinococcota | *Deinococcus arenae* | WP_229781077.1 | WP_162621388.1 | WP_110829477.1 |

| | | | | |
|---|---|---|---|---|
| Deinococcota | *Deinococcus arenicola* | WP_317639329.1 | WP_317641462.1 | WP_317640175.1 |
| Deinococcota | *Deinococcus budaensis* | WP_184024512.1 | WP_246363429.1 | WP_184024438.1 |
| Deinococcota | *Deinococcus caeni* | WP_345444445.1 | WP_345441182.1 | WP_345441909.1 |
| Deinococcota | *Deinococcus carri* | WP_345461590.1 | WP_345466657.1 | WP_345461676.1 |
| Deinococcota | *Deinococcus cavernae* | WP_119761767.1 | WP_119764245.1 | WP_119762871.1 |
| Deinococcota | *Deinococcus cellulosilyticus* | WP_146882446.1 | WP_146886834.1 | WP_246130609.1 |
| Deinococcota | *Deinococcus daejeonensis* | WP_189053633.1 | WP_189055309.1 | WP_189053065.1 |
| Deinococcota | *Deinococcus depolymerans* | WP_343760772.1 | WP_343758711.1 | WP_343755778.1 |
| Deinococcota | *Deinococcus deserti* | WP_012693197.1 | WP_162485366.1 | WP_012693436.1 |
| Deinococcota | *Deinococcus detaillensis* | WP_143721285.1 | WP_143719968.1 | WP_143719870.1 |
| Deinococcota | *Deinococcus ficus* | WP_027462627.1 | WP_022801228.1 | WP_027461688.1 |
| Deinococcota | *Deinococcus fonticola* | WP_135227646.1 | WP_135229883.1 | WP_135230392.1 |
| Deinococcota | *Deinococcus frigens* | WP_029476876.1 | WP_029478215.1 | WP_029477453.1 |
| Deinococcota | *Deinococcus gobiensis* | WP_014685004.1 | WP_043800282.1 | WP_014684828.1 |
| Deinococcota | *Deinococcus grandis* | WP_058976710.1 | WP_058974927.1 | WP_058976407.1 |
| Deinococcota | *Deinococcus hohokamensis* | WP_380059839.1 | WP_380062069.1 | WP_380060145.1 |
| Deinococcota | *Deinococcus hopiensis* | WP_084049044.1 | WP_084050723.1 | WP_084048923.1 |
| Deinococcota | *Deinococcus humi* | WP_184129699.1 | WP_184126902.1 | WP_184129920.1 |
| Deinococcota | *Deinococcus irradiatisoli* | WP_245896078.1 | WP_109827479.1 | WP_109827008.1 |
| Deinococcota | *Deinococcus koreensis* | WP_103309064.1 | WP_103313291.1 | WP_103312787.1 |
| Deinococcota | *Deinococcus kurensis* | WP_162393704.1 | WP_235910409.1 | WP_155297836.1 |
| Deinococcota | *Deinococcus malanensis* | WP_189005511.1 | WP_229780756.1 | WP_189004304.1 |
| Deinococcota | *Deinococcus maricopensis* | WP_013557152.1 | WP_013555732.1 | WP_013557142.1 |
| Deinococcota | *Deinococcus marmoris* | WP_075831195.1 | WP_075831896.1 | WP_075835214.1 |
| Deinococcota | *Deinococcus metalli* | WP_184109496.1 | WP_184108687.1 | WP_221275190.1 |
| Deinococcota | *Deinococcus metallilatus* | WP_129119165.1 | WP_129117670.1 | WP_129119133.1 |
| Deinococcota | *Deinococcus misasensis* | WP_034338289.1 | WP_034339836.1 | WP_034337961.1 |
| Deinococcota | *Deinococcus multiflagellatus* | WP_224603931.1 | WP_380056235.1 | WP_224605968.1 |

| | | | | |
|---|---|---|---|---|
| Deinococcota | *Deinococcus navajonensis* | WP_380036978.1 | WP_380041639.1 | WP_380037409.1 |
| Deinococcota | *Deinococcus oregonensis* | WP_380008879.1 | WP_380013361.1 | WP_380009835.1 |
| Deinococcota | *Deinococcus peraridilitoris* | WP_041230988.1 | WP_015234773.1 | WP_015237366.1 |
| Deinococcota | *Deinococcus petrolearius* | WP_380047487.1 | WP_380048497.1 | WP_380047843.1 |
| Deinococcota | *Deinococcus phoenicis* | WP_034358017.1 | WP_034354440.1 | WP_034358065.1 |
| Deinococcota | *Deinococcus piscis* | WP_189643464.1 | WP_189641973.1 | WP_189643702.1 |
| Deinococcota | *Deinococcus planocerae* | WP_102125389.1 | WP_102126075.1 | WP_102125342.1 |
| Deinococcota | *Deinococcus psychrotolerans* | WP_124872848.1 | WP_124867313.1 | WP_124867048.1 |
| Deinococcota | *Deinococcus puniceus* | WP_157451107.1 | WP_064013629.1 | WP_064014463.1 |
| Deinococcota | *Deinococcus radiodurans* | WP_051618804.1 | WP_010888991.1 | WP_027479890.1 |
| Deinococcota | *Deinococcus radiomollis* | WP_407541532.1 | WP_407540184.1 | WP_407538780.1 |
| Deinococcota | *Deinococcus radiophilus* | WP_126351677.1 | WP_126352099.1 | WP_229253139.1 |
| Deinococcota | *Deinococcus radiopugnans* | WP_221265535.1 | WP_039686665.1 | WP_039683546.1 |
| Deinococcota | *Deinococcus radiotolerans* | WP_189067161.1 | WP_189068310.1 | WP_189067021.1 |
| Deinococcota | *Deinococcus reticulitermitis* | WP_092265124.1 | WP_177183183.1 | WP_092262682.1 |
| Deinococcota | *Deinococcus roseus* | WP_229684599.1 | WP_188999069.1 | WP_229684699.1 |
| Deinococcota | *Deinococcus rubellus* | WP_260561884.1 | WP_260559421.1 | WP_260559226.1 |
| Deinococcota | *Deinococcus ruber* | WP_189087755.1 | WP_189092038.1 | WP_189092761.1 |
| Deinococcota | *Deinococcus saxicola* | WP_415784007.1 | WP_415785443.1 | WP_415785112.1 |
| Deinococcota | *Deinococcus soli (ex Cha et al. 2016)* | WP_046844773.1 | WP_309855596.1 | WP_309849145.1 |
| Deinococcota | *Deinococcus sonorensis* | WP_350242827.1 | WP_350244490.1 | WP_350245292.1 |
| Deinococcota | *Deinococcus sp.* | WP_303100937.1 | WP_291426961.1 | WP_291425733.1 |
| Deinococcota | *Deinococcus taeanensis* | WP_225477265.1 | WP_225475795.1 | WP_225477183.1 |
| Deinococcota | *Deinococcus taklimakanensis* | WP_386843827.1 | WP_386843423.1 | WP_386842091.1 |
| Deinococcota | *Deinococcus terrestris* | WP_152868895.1 | WP_152871367.1 | WP_322618497.1 |
| Deinococcota | *Deinococcus wulumuqiensis* | WP_114671372.1 | WP_025568165.1 | WP_278911727.1 |
| Deinococcota | *Deinococcus xinjiangensis* | WP_353540710.1 | WP_353541224.1 | WP_353543416.1 |
| Deinococcota | *Deinococcus yavapaiensis* | WP_110886036.1 | WP_110886876.1 | WP_110887518.1 |

| | | | | |
|---|---|---|---|---|
| Deinococcota | *Deinococcus yunweiensis* | WP_412029121.1 | WP_412029680.1 | WP_412029553.1 |
| Deinococcota | *Marinithermus hydrothermalis* | WP_013703448.1 | WP_013704541.1 | WP_013702922.1 |
| Deinococcota | *Meiothermus granaticius* | WP_119357307.1 | WP_240631290.1 | WP_119358206.1 |
| Deinococcota | *Meiothermus sp.* | WP_337868470.1 | WP_295397524.1 | WP_314135743.1 |
| Deinococcota | *Oceanithermus desulfurans* | WP_147148152.1 | WP_147145011.1 | WP_183677849.1 |
| Deinococcota | *Oceanithermus profundus* | WP_013457204.1 | WP_013458147.1 | WP_013456947.1 |
| Deinococcota | *Oceanithermus sp.* | WP_293171335.1 | WP_287372649.1 | WP_287408025.1 |
| Deinococcota | *Thermus albus* | WP_243027396.1 | WP_243027258.1 | WP_243029035.1 |
| Deinococcota | *Thermus altitudinis* | WP_243029601.1 | WP_243029378.1 | WP_243031917.1 |
| Deinococcota | *Thermus amyloliquefaciens* | WP_038058845.1 | WP_038056350.1 | WP_038058292.1 |
| Deinococcota | *Thermus antranikianii* | WP_028493307.1 | WP_028493165.1 | WP_028493524.1 |
| Deinococcota | *Thermus aquaticus* | WP_053768318.1 | WP_053767951.1 | WP_003043996.1 |
| Deinococcota | *Thermus arciformis* | WP_093007318.1 | WP_093005597.1 | WP_093004707.1 |
| Deinococcota | *Thermus brockianus* | WP_071676276.1 | WP_071677682.1 | WP_071677286.1 |
| Deinococcota | *Thermus caldifontis* | WP_205387548.1 | WP_114312579.1 | WP_114312959.1 |
| Deinococcota | *Thermus caliditerrae* | WP_038046263.1 | WP_234553299.1 | WP_234554877.1 |
| Deinococcota | *Thermus composti* | WP_188847462.1 | WP_188845202.1 | WP_188847932.1 |
| Deinococcota | *Thermus filiformis* | WP_038061065.1 | WP_038060589.1 | WP_038067687.1 |
| Deinococcota | *Thermus igniterrae* | WP_018112247.1 | WP_018112198.1 | WP_018111336.1 |
| Deinococcota | *Thermus islandicus* | WP_022798861.1 | WP_022797977.1 | WP_022797816.1 |
| Deinococcota | *Thermus oshimai* | WP_018461480.1 | WP_016329451.1 | WP_016328891.1 |
| Deinococcota | *Thermus parvatiensis* | WP_060384412.1 | WP_060384356.1 | WP_008633192.1 |
| Deinococcota | *Thermus scotoductus* | WP_172955583.1 | WP_038068734.1 | WP_038070430.1 |
| Deinococcota | *Thermus sediminis* | WP_117237037.1 | WP_117236739.1 | WP_117238527.1 |
| Deinococcota | *Thermus sp.* | WP_026328956.1 | WP_370533807.1 | WP_347241700.1 |
| Deinococcota | *Thermus tengchongensis* | WP_234557131.1 | WP_234557249.1 | WP_135343133.1 |
| Deinococcota | *Thermus thermamylovorans* | WP_130839858.1 | WP_130839531.1 | WP_130841377.1 |
| Deinococcota | *Thermus thermophilus* | WP_165738579.1 | WP_203970898.1 | WP_223903362.1 |

| | | | | |
|---|---|---|---|---|
| Deinococcota | *Truepera radiovictrix* | WP_013178337.1 | WP_013177075.1 | WP_013177872.1 |
| Gemmatimonadota | *Candidatus Palauibacter irciniicola* | WP_419859026.1 | WP_419856829.1 | WP_419858894.1 |
| Gemmatimonadota | *Candidatus Palauibacter polyketidifaciens* | WP_310783687.1 | WP_310785236.1 | WP_310785103.1 |
| Gemmatimonadota | *Candidatus Palauibacter scopulicola* | WP_310777503.1 | WP_310775338.1 | WP_310778328.1 |
| Gemmatimonadota | *Candidatus Palauibacter soopunensis* | WP_310758907.1 | WP_310757617.1 | WP_310759045.1 |
| Gemmatimonadota | *Gemmatimonas aurantiaca* | WP_337170776.1 | WP_012682624.1 | WP_012683085.1 |
| Gemmatimonadota | *Gemmatimonas groenlandica* | WP_171224594.1 | WP_171225058.1 | WP_171224533.1 |
| Gemmatimonadota | *Gemmatimonas phototrophica* | WP_026850599.1 | WP_026850308.1 | WP_158514790.1 |
| Gemmatimonadota | *Gemmatimonas sp.* | WP_396213143.1 | WP_396214343.1 | WP_309669755.1 |
| Gemmatimonadota | *Gemmatirosa kalamazoonensis* | WP_025412160.1 | WP_025411901.1 | WP_025412232.1 |
| Gemmatimonadota | *Longimicrobium sp.* | WP_331875729.1 | WP_331053369.1 | WP_331074245.1 |
| Gemmatimonadota | *Longimicrobium terrae* | WP_170038756.1 | WP_170035611.1 | WP_170036976.1 |
| Gemmatimonadota | *Pseudogemmatithrix spongiicola* | WP_367887761.1 | WP_367887364.1 | WP_367887824.1 |
| Gemmatimonadota | *Roseisolibacter agri* | WP_284351024.1 | WP_284350742.1 | WP_284351645.1 |
| Ignavibacteriota | *Ignavibacterium album* | WP_304142912.1 | WP_014560830.1 | WP_304130095.1 |
| Ignavibacteriota | *Ignavibacterium sp.* | WP_297836878.1 | WP_337867012.1 | WP_337872978.1 |
| Myxococcota | *Archangium lipolyticum* | WP_257455266.1 | WP_257455265.1 | WP_257455264.1 |
| Myxococcota | *Archangium minus* | WP_395823165.1 | WP_395823167.1 | WP_395823169.1 |
| Myxococcota | *Archangium primigenium* | WP_204487858.1 | WP_204487860.1 | WP_204487862.1 |
| Myxococcota | *Archangium sp.* | WP_331119032.1 | WP_331102339.1 | WP_331102341.1 |
| Myxococcota | *Archangium violaceum* | WP_204220669.1 | WP_395843542.1 | WP_239015428.1 |
| Myxococcota | *Corallococcus aberystwythensis* | WP_120553254.1 | WP_120553253.1 | WP_208721103.1 |
| Myxococcota | *Corallococcus coralloides* | WP_128795524.1 | WP_014394710.1 | WP_014394711.1 |
| Myxococcota | *Corallococcus exercitus* | WP_375759901.1 | WP_375759900.1 | WP_375759899.1 |
| Myxococcota | *Corallococcus exiguus* | WP_171810442.1 | WP_206879771.1 | WP_206879769.1 |
| Myxococcota | *Corallococcus interemptor* | WP_427290402.1 | WP_120546908.1 | WP_120546907.1 |
| Myxococcota | *Corallococcus macrosporus* | WP_207050954.1 | WP_013938150.1 | WP_193364466.1 |

| | | | | |
|---|---|---|---|---|
| Myxococcota | *Corallococcus sicarius* | WP_120623473.1 | WP_120623474.1 | WP_208718773.1 |
| Myxococcota | *Corallococcus soli* | WP_193347660.1 | WP_193347661.1 | WP_193347662.1 |
| Myxococcota | *Corallococcus sp. BB11-1* | WP_267556363.1 | WP_267556362.1 | WP_267556361.1 |
| Myxococcota | *Hyalangium gracile* | WP_224246637.1 | WP_224246631.1 | WP_224246630.1 |
| Myxococcota | *Hyalangium minutum* | WP_044181231.1 | WP_044181229.1 | WP_044181226.1 |
| Myxococcota | *Hyalangium rubrum* | WP_321546729.1 | WP_321546728.1 | WP_321546727.1 |
| Myxococcota | *Hyalangium sp.* | WP_324988804.1 | WP_324988805.1 | WP_422653662.1 |
| Myxococcota | *Hyalangium versicolor* | WP_224364566.1 | WP_224364567.1 | WP_224364568.1 |
| Myxococcota | *Melittangium boletus* | WP_095980233.1 | WP_095980232.1 | WP_095983038.1 |
| Myxococcota | *Myxococcus dinghuensis* | WP_253972760.1 | WP_253972761.1 | WP_253972790.1 |
| Myxococcota | *Myxococcus eversor* | WP_163870315.1 | WP_163870316.1 | WP_163870317.1 |
| Myxococcota | *Myxococcus fulvus* | WP_408889278.1 | WP_074951182.1 | WP_408889280.1 |
| Myxococcota | *Myxococcus guangdongensis* | WP_254042490.1 | WP_254042489.1 | WP_254042488.1 |
| Myxococcota | *Myxococcus hansupus* | WP_002635932.1 | WP_002635933.1 | WP_002635934.1 |
| Myxococcota | *Myxococcus landrumus* | WP_206714871.1 | WP_206714872.1 | WP_241757966.1 |
| Myxococcota | *Myxococcus qinghaiensis* | WP_253996877.1 | WP_253996878.1 | WP_253996879.1 |
| Myxococcota | *Myxococcus sp. AM010* | WP_176417252.1 | WP_176417253.1 | WP_176417254.1 |
| Myxococcota | *Myxococcus stipitatus* | WP_338863836.1 | WP_338863837.1 | WP_338863838.1 |
| Myxococcota | *Myxococcus vastator* | WP_163777287.1 | WP_163777286.1 | WP_176412746.1 |
| Myxococcota | *Myxococcus xanthus* | WP_140795178.1 | WP_140857059.1 | WP_174259366.1 |
| Myxococcota | *Pyxidicoccus caerfyrddinensis* | WP_164000549.1 | WP_164000547.1 | WP_164000545.1 |
| Myxococcota | *Pyxidicoccus fallax* | WP_169350108.1 | WP_169350107.1 | WP_169350106.1 |
| Myxococcota | *Pyxidicoccus parkwaysis* | WP_206724901.1 | WP_206724902.1 | WP_371877666.1 |
| Myxococcota | *Pyxidicoccus sp. MSG2* | WP_267859784.1 | WP_267859785.1 | WP_267859786.1 |
| Myxococcota | *Pyxidicoccus trucidator* | WP_164018915.1 | WP_164018914.1 | WP_164018913.1 |
| Myxococcota | *Pyxidicoccus xibeiensis* | WP_253983710.1 | WP_253983709.1 | WP_253983708.1 |
| Myxococcota | *Stigmatella ashevillena* | WP_272144062.1 | WP_272144064.1 | WP_272144065.1 |
| Myxococcota | *Stigmatella aurantiaca* | WP_013374927.1 | WP_002613833.1 | WP_013374928.1 |

| | | | | |
|---|---|---|---|---|
| Myxococcota | *Stigmatella erecta* | WP_093519928.1 | WP_093519930.1 | WP_093519932.1 |
| Myxococcota | *Stigmatella hybrida* | WP_225411804.1 | WP_225411805.1 | WP_225411806.1 |
| Myxococcota | *Vulgatibacter sp.* | WP_373045413.1 | WP_373045414.1 | WP_373045415.1 |
| Planctomycetota | *Engelhardtia mirabilis* | WP_145070674.1 | WP_419191964.1 | WP_419191963.1 |
| Planctomycetota | *Saltatorellus ferox* | WP_145205850.1 | WP_419190757.1 | WP_145205827.1 |
| Rhodothermota | *Longibacter salinarum* | WP_098074559.1 | WP_098074888.1 | WP_098074867.1 |
| Rhodothermota | *Longimonas halophila* | WP_098061945.1 | WP_098063067.1 | WP_098060759.1 |
| Rhodothermota | *Longimonas sp.* | WP_353562849.1 | WP_353564194.1 | WP_353562900.1 |
| Rhodothermota | *Rhodocaloribacter litoris* | WP_166974234.1 | WP_228350645.1 | WP_166978060.1 |
| Rhodothermota | *Rhodothermus bifroesti* | WP_210373907.1 | WP_210375124.1 | WP_210375121.1 |
| Rhodothermota | *Rhodothermus marinus* | WP_029933186.1 | WP_161540516.1 | WP_012843179.1 |
| Rhodothermota | *Rhodothermus profundi* | WP_072715349.1 | WP_072715210.1 | WP_072715214.1 |
| Rhodothermota | *Rubricoccus marinus* | WP_094547598.1 | WP_094548922.1 | WP_094545390.1 |
| Rhodothermota | *Rubrivirga marina* | WP_095512051.1 | WP_095512152.1 | WP_095511188.1 |
| Rhodothermota | *Rubrivirga sp.* | WP_424519443.1 | WP_420456587.1 | WP_424520788.1 |
| Rhodothermota | *Salinibacter altiplanensis* | WP_103029746.1 | WP_103018943.1 | WP_103020602.1 |
| Rhodothermota | *Salinibacter grassmerensis* | WP_263786151.1 | WP_263785401.1 | WP_263786104.1 |
| Rhodothermota | *Salinibacter ruber* | WP_259170794.1 | WP_259083461.1 | WP_251923059.1 |
| Rhodothermota | *Salinibacter sp.* | WP_375800574.1 | WP_332316875.1 | WP_263834564.1 |
| Rhodothermota | *Salisaeta longa* | WP_022835967.1 | WP_022836238.1 | WP_028567198.1 |
